# A new lineage of *Ranavirus micropterus1* infects ornamental wrasses from the Great Barrier Reef and causes severe disease in captivity

**DOI:** 10.64898/2026.02.05.704086

**Authors:** Vincenzo A. Costa, Rina Li, David Furner, Richard J. Whittington, Lee K. Campbell, Annabelle Olsson, Francisca Samsing

## Abstract

The Great Barrier Reef (GBR) supports remarkable vertebrate diversity but is among the most fragile ecosystems globally, and diseases affecting reef fishes remain poorly understood. We employed traditional veterinary diagnostic techniques and metatranscriptomics to investigate a disease outbreak with 100% mortality in captive wrasses (*Macropharyngodon choati*) from the GBR. Histopathology revealed multifocal necrosis of renal tubules, and cytopathic effect was observed in bluegill fry cell lines. A novel ranavirus isolate, Macropharyngodon choati ranavirus (McRV), was identified at exceptionally high transcript abundance across different host tissues, alongside strong expression of host genes linked to cellular stress and immune function. Phylogenetic analysis placed McRV in a distinct clade of wrasse ranaviruses, clustering with a virus previously identified on the GBR, suggesting that wild wrasses serve as natural hosts. These data emphasise the need for enhanced pathogen monitoring in marine wildlife increasingly threatened by anthropogenic change.

## Introduction

Despite occupying only a small fraction of the world’s surface area, tropical coral reefs are recognised as cradles for biodiversity, supporting one-third of all currently described marine fishes^1^. The Great Barrier Reef (GBR), the largest coral reef ecosystem in the world, contributes $6.4 billion to the Australian economy, supporting over 64,000 jobs^2,3^. The GBR provides a variety of ecosystem services including coastal protection, fisheries, tourism, and recreation. Moreover, as one of the most speciose groups of vertebrates, with evolutionary histories spanning over 60 million years, fishes of the GBR provide cultural, educational, and scientific benefits^4^.

Wrasses (Labridae) are among the most evolutionarily successful and ecologically diverse vertebrate groups on tropical coral reefs, with over 600 species arranged into 82 genera^5^. Wrasses exhibit remarkable variation in colour, body shape, and feeding ecology, alongside complex mating systems^5^. Owing to their extraordinary diversity in colouration, wrasses are a popular group in the global aquarium industry^6^. They also play key ecological roles through cleaning symbioses supporting fish health and recruitment and through invertebrate predation that shapes benthic communities^7,8^. Despite their economic and ecological importance, diseases affecting wrasses, and reef fishes more broadly, have received far less attention than those impacting corals, even though wrasses similarly face increasing pressures from anthropogenic stressors^9^.

Few diseases have been reported in fishes on the GBR. They include *Streptococcus agalactiae* infections in wild groupers (*Epinephelus lanceolatus*)^10^, serranid pigment abnormality syndrome, a disease of unknown aetiology affecting wire netting cod (*Epinephelus quoyanus*) on the southern GBR^11^, and viral erythrocytic necrosis, caused by erythrocytic necrosis virus (ENV; *Iridoviridae*), in a juvenile triggerfish (*Rhinecanthus aculeatus*) from Lizard Island^12^. Moreover, recent explorations of the reef fish virome – using metatranscriptomic sequencing – have identified a variety of RNA and DNA viruses in apparently healthy fishes, including iridoviruses at Orpheus Island and Lizard Island, suggesting that they are widespread on the GBR^13,14^.

The *Iridoviridae* are a family of double-stranded DNA viruses that infect a wide range of ectothermic vertebrates and invertebrates^15^. The family is divided into two subfamilies: the *Alphairidovirinae*, which infect bony fishes, amphibians, and reptiles; and the *Betairidovirinae*, which also include viruses infecting fishes (e.g. ENV) but are primarily composed of those associated with invertebrate hosts, including insects, crustaceans, and molluscs^15^. The *Alphairidovirinae* incorporates important pathogens of bony fish, including those listed by the World Organisation for Animal Health (WOAH): infectious spleen and kidney necrosis virus, red sea bream iridovirus and turbot reddish body iridovirus (genus *Megalocytivirus*), and epizootic haematopoietic necrosis virus (genus *Ranavirus*). Other ranaviruses of concern include viruses belonging to the *Ranavirus micropterus1* species, including largemouth bass virus (LMBV), which was first identified in North America as the causative agent of fish kills in wild largemouth bass (*Micropterus salmoides*) and has since emerged, along with related isolates, as a pathogen affecting a variety of farmed fish species in Asia^16,17^.

Here, we investigated a disease affecting ornamental red leopard wrasses (*M. choati*) from the GBR showing 100% mortality upon onset of clinical signs. Histopathological analysis identified kidney lesions that were suggestive of an infectious process. To investigate further, we performed metatranscriptomic analysis on a variety of tissue samples which identified a ranavirus with notably high transcript abundance. We isolated the virus in cell culture and validated its replication through PCR and whole genome sequencing.

## Results

Tissue samples of dead wrasses (n=11) were received by our laboratory for disease investigation from a commercial wholesale marine ornamental fish facility. All animals were reported to have exhibited abnormal swimming behaviour and impaired respiration nine days after transfer to captivity, with 100% mortality occurring within 1-2 days of onset of clinical signs. No other species at the facility exhibited clinical signs.

### Histopathology

Relevant lesions were identified in the kidney where there was acute multifocal tubulointerstitial necrosis. This mostly spared the densely cellular haematopoietic tissue (Figure 1A). Some glomeruli were involved within some of the necrotic areas. There were pyknotic epithelial cells in early-affected renal tubules, and some tubules contained casts of cell debris.

**Figure 1.**
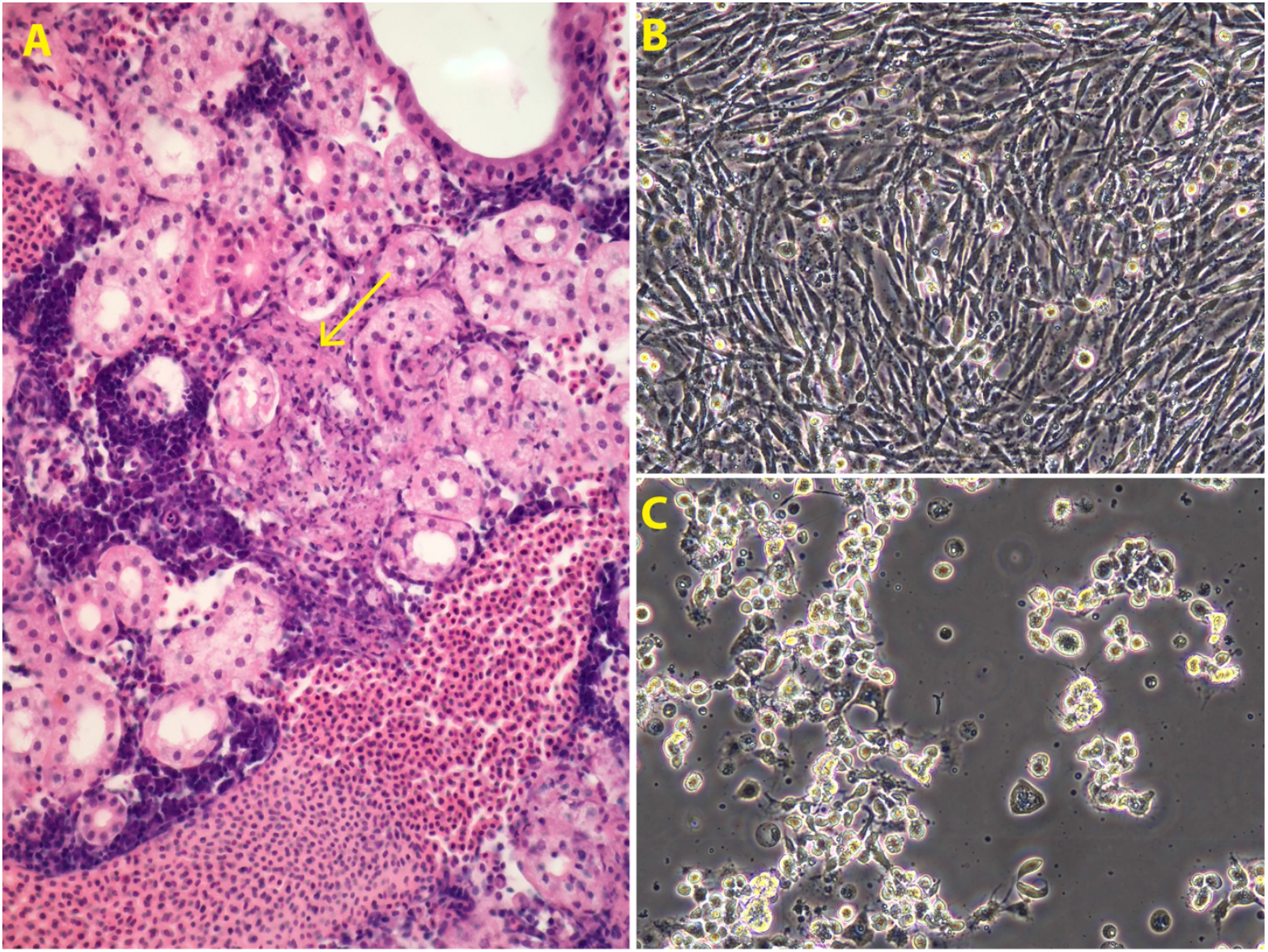
Isolation of McRV in BF-2 cells and histological lesions. **(A)** Uninfected control BF-2 monolayer at 6 dpi in the first passage. (**B)** BF-2 cells inoculated with wrasse spleen homogenate showing cytopathic effect at 6dpi in the primary passage. Both images taken at 20x magnification. (**C)** Kidney, with acute tubulointerstitial necrosis (arrow). There is multifocal to locally extensive necrosis, largely sparing the densely cellular basophilic haematopoietic tissue, but rather concentrated on renal tubules and intertubular (interstitial) connective tissues.

### Identification of a ranavirus in wrasse metatranscriptomes

Shotgun metatranscriptomic sequencing was performed on spleen, liver, and brain samples, resulting in a total of 979 million reads (median 55 million reads per library) (Figure 2A). Sequencing reads were assembled into a total of 2.9 million contigs (median 149,246 per library) with an average N50 of 1452. Metatranscriptomic analysis identified a ranavirus, tentatively named Macropharyngodon choati ranavirus (McRV) (Figure 2B; Table S1). Importantly, McRV was detected in all sample types, and no other vertebrate-associated viruses were identified. McRV reads dominated spleen samples (median 2.72 million reads) and made up a very large proportion of non-host reads in samples W1.1 (71%), W4.1 (85%), and W5.1 (81%). McRV exhibited significantly higher abundance than all other top non-host genera, including opportunistic bacteria and parasites (Wilcoxon signed-rank test, all p < 0.05). For instance, median abundance of McRV was 29-fold higher than the entire *Vibrio* genus (p = 0.01) and over 1000-fold higher than the myxozoan parasite *Enteromyxum leei* (p < 0.001) (Figure 2B). Because of its exceptionally high abundance and presence in all samples relative to other potential infectious agents, we focused our analysis on McRV infection, as described in the sections below.

**Figure 2.**
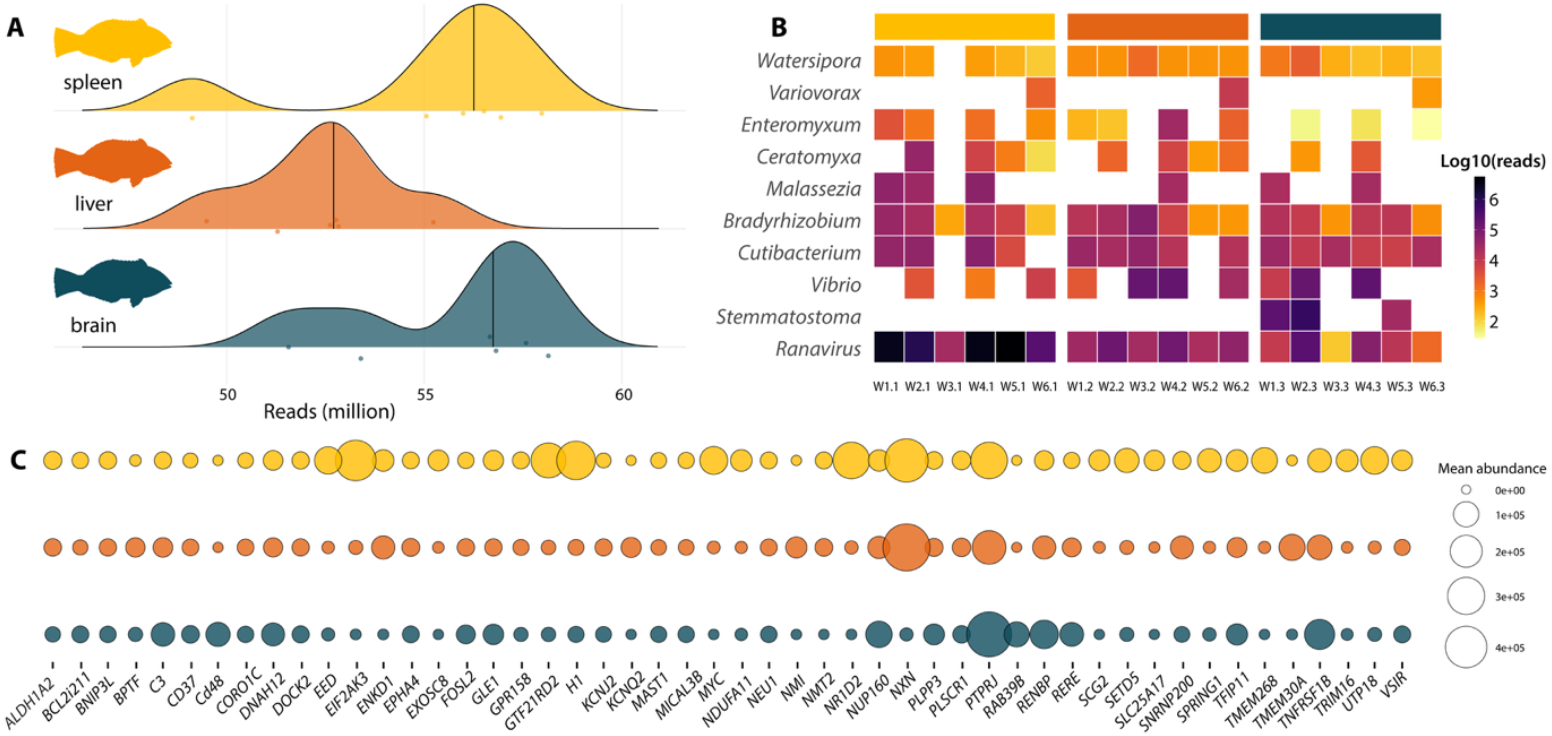
Composition of wrasse metatranscriptomes. **(A)** Number of reads per library. (**B)** Normalised abundance of top 10 genera following the removal of host reads. (**C)** Mean transcript abundance of the top 50 expressed genes identified in samples.

### High expression of genes linked to cellular stress and immune function

To complement pathogen discovery, we examined host transcript abundance. Principal coordinate analysis (PCoA) based on Bray–Curtis dissimilarity revealed differences in transcriptional profiles among tissue types (PERMANOVA, p = 0.001, R^2^ = 0.418). Pairwise comparisons showed that brain tissue was significantly different from both liver (p = 0.002, R^2^ = 0.478) and spleen (p = 0.003, R^2^ = 0.325). While liver and spleen were also significantly different from each other (p = 0.006, R^2^ = 0.262), they clustered more closely together (Figure S1). Among all samples, the 50 most highly expressed genes were dominated by transcripts associated with cellular stress and transcriptional regulation including *NXN, EIF2AK3, MYC, H1, GTF2IRD2*, and *EED* (Table S2). Several immune genes were also among the top 50 expressed. These fell into four broad categories reflecting Gene Ontology (GO) terms (Biological Process): (i) innate immunity and antiviral defence, including toll-like receptor signalling, macrophage activation, and interferon responses (*NMI, PLSCR1, FOSL2*); (ii) complement activation (*C3)* ; (iii) adaptive immune activation, with regulation of T- and B-cell proliferation and differentiation (*CD48, CD37, VSIR, DOCK2, FOSL2*); and (iv) cytokine-mediated and inflammatory signalling, including interleukin and tumour necrosis factor (TNF) signalling (*TRIM16, TNFRSF1B, PTPRJ, NMI, PLSCR1*) (Figure 2C).

### Replication of McRV in bluegill fry cell lines

We inoculated bluegill fry (BF-2) fish cell lines using splenic homogenates, as these exhibited the highest transcript abundance of McRV in our metatranscriptomic analysis. Cytopathic effect (CPE) was observed at 5 days post-inoculation (dpi) in a culture well inoculated with diluted, clarified homogenate. Supernatant from this culture was used for a single passage once onto fresh BF-2 monolayers at the same dilution, where similar CPE developed by 6 dpi (Figure 1C). No CPE was observed in control wells (Figure 1B).

### PCR confirmation of McRV

We performed PCR using *Ranavirus micropterus1* primers targeting the major capsid protein (380 bp)^13^ for validation. This was performed on tissue samples, tissue homogenates used for cell culture, and tissue culture supernatant. Importantly, all tissue samples and homogenates were positive for McRV. Among tissue culture supernatant samples, 63% were positive, with the strongest bands detected in wells exhibiting CPE (Figure 1C). All negative controls tested negative.

### Genomic, transcriptomic, and phylogenetic analysis of McRV

Total DNA was sequenced from the tissue culture supernatant of McRV-infected BF-2 cells (Figure 2B). From this, we recovered a complete virus genome of 98,903 nt, assembled into a single contig with a mean coverage of 5036x (total 3.4 million reads), which accounted for 68% of the total sequence reads (Figure 3A). Consistent with other ranaviruses, the McRV genome is circularly permuted and terminally redundant, as indicated by direct terminal repeats (DTRs) and the presence of the RNaseIII gene across the genome termini (Figure 3A). The virus genome encodes 130 predicted open reading frames (ORFs) and has a G+C content of 52.1%.

**Figure 3.**
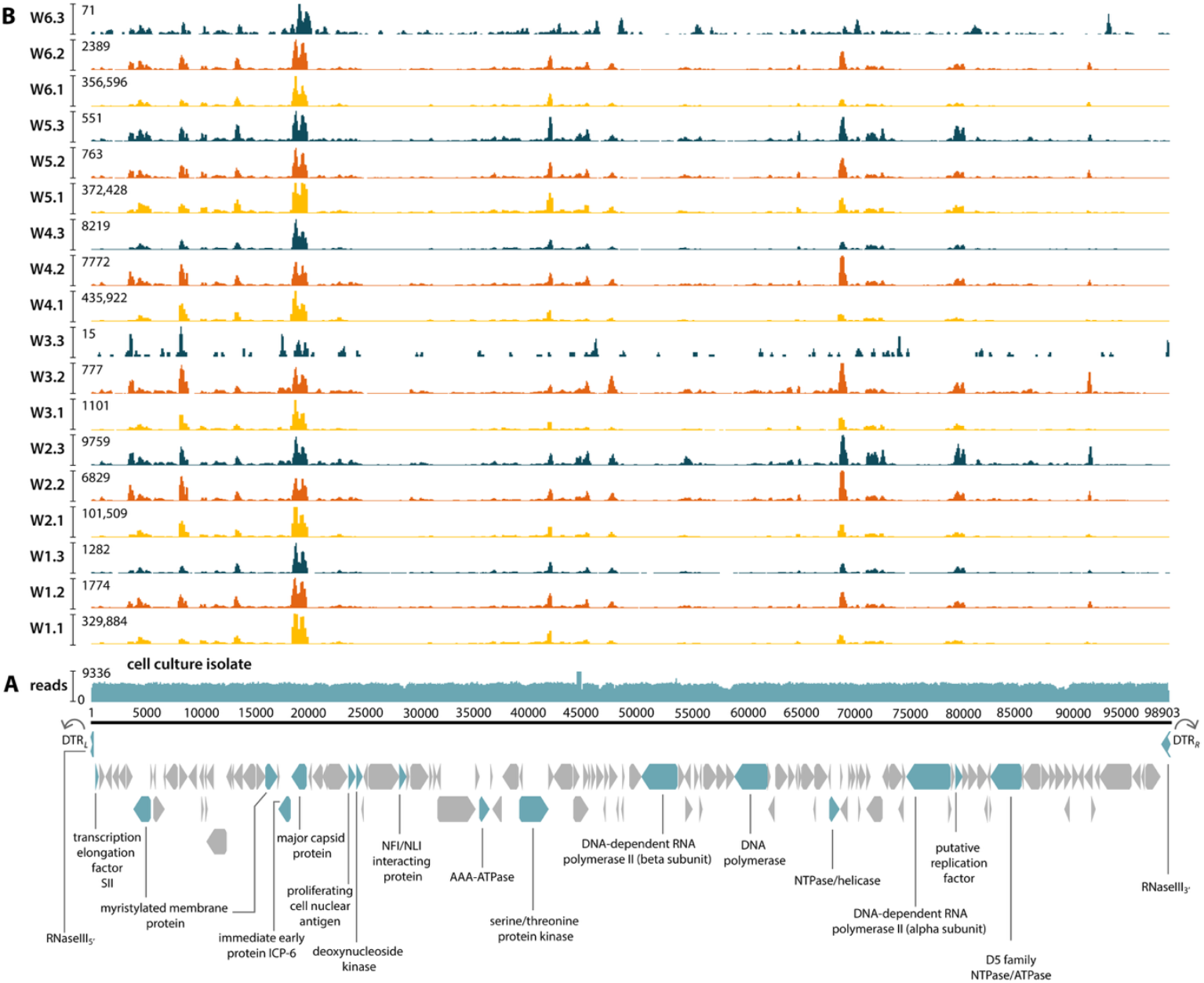
McRV genome structure and coverage. **(A)** McRV genome recovered from cell culture. The circularly permuted and terminally redundant genome is displayed in its linear arbitrary order from 5’ to 3’ as indicated by DTRs and the RNaseIII gene at the genome termini. Genome coverage is depicted as the number of mapped reads. Core genes are highlighted and labelled below. **(B)** McRV coverage across wrasse metatranscriptomes. Read mapping illustrates viral gene expression at the time of sampling. Coverage colour-coded by sample type, consistent with Figure 2.

The complete coding region shared ∼98% nucleotide identity with members of the *Ranavirus micropterus1* (e.g. LMBV) previously reported from China and ∼97% identity with those from North America. The cell culture isolate was almost identical (∼99.7%) to McRV genes detected in wrasse metatranscriptomes. In these samples, coverage was consistently highest for the major capsid protein. Other genes showing similarly uniform coverage included the myristylated membrane protein, the p31K protein, ORF54 and ORF84 (both of unknown function), and ORF24, which encodes a predicted immunoglobulin-like domain (Figure 3B).

Phylogenetic analysis of the virus major capsid protein revealed two clades broadly corresponding to geographic region: clade I, which incorporates viruses exclusively found in Asia and clade II, incorporating those primarily from North America. McRV fell within clade I and was identical to Halichoeres melanurus ranavirus (NCBI/GenBank: ON595973), which was identified in April 2021 from Orpheus Island on the GBR (Figure 4B).

**Figure 4.**
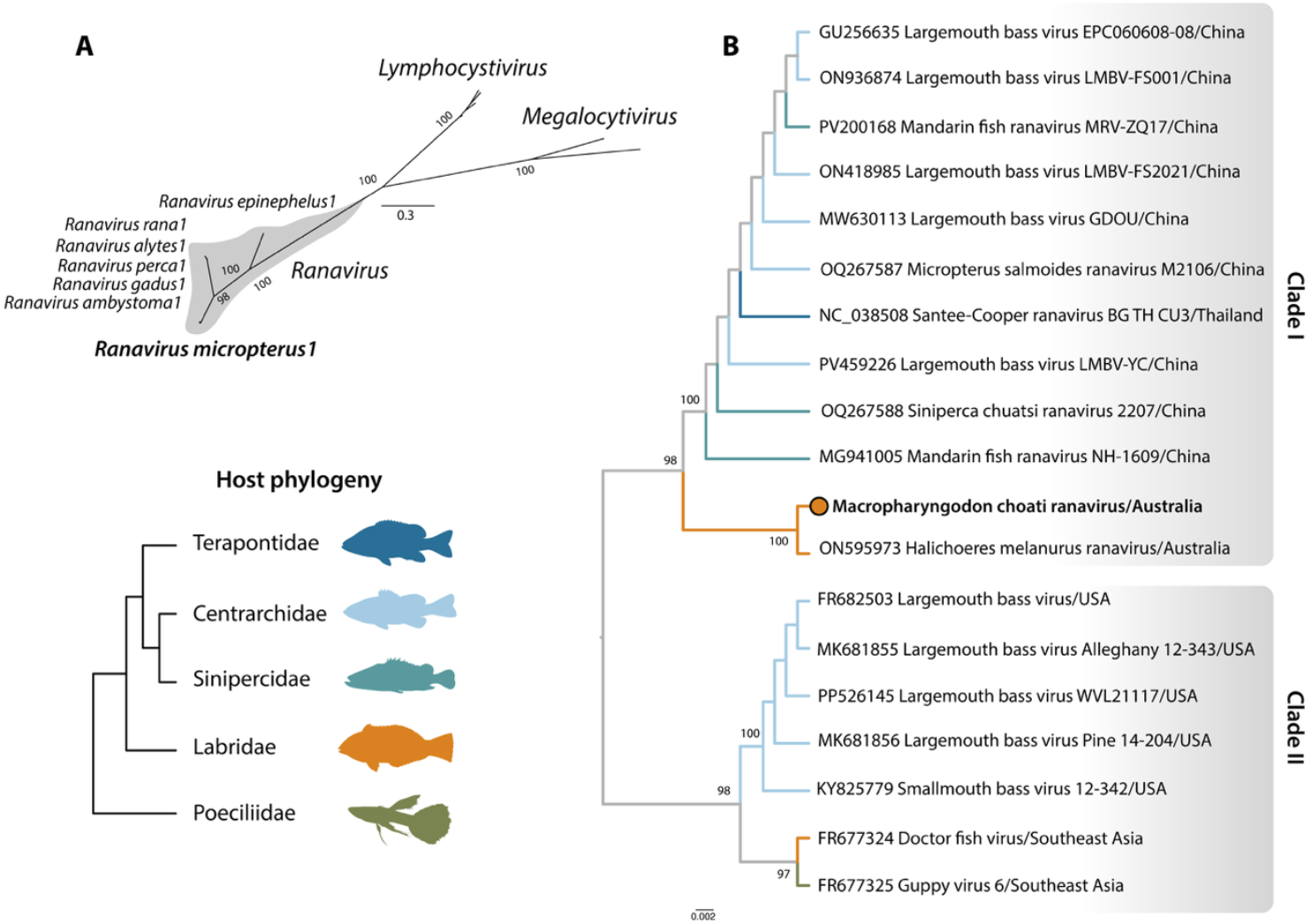
Phylogenetic relationships. **(A)** Unrooted phylogeny of the *Alphairidovirinae* (*Iridoviridae*) based on nucleotide sequences of the major capsid protein. The *Ranavirus* genus is shaded to highlight its placement within the subfamily. Scale bar indicates nucleotide substitutions per site. **(B)** Phylogeny of *Ranavirus micropterus1* using nucleotide sequences of the major capsid protein. Branches are colour-coded by host family, with McRV indicated by a coloured circle. The tree is midpoint-rooted for clarity only. Scale bar indicates nucleotide substitutions per site.

## Discussion

We used a combination of traditional veterinary diagnostic techniques and metatranscriptomics to investigate a disease of unknown aetiology in captive *M. choati* from the GBR, Australia^18^. This led to the identification of McRV, a new member of the virus species *Ranavirus micropterus1*. Several lines of evidence suggest that McRV was the likely causative agent of the disease outbreak observed in this study. McRV was the most dominant infectious agent in our metatranscriptomic analysis, with significantly higher transcript abundance than opportunistic pathogens and parasites including *Vibrio spp., Enteromyxum leei*, and *Stemmatostoma* which were detected at low abundance and only sporadically across tissue samples. Transcript abundance of McRV was extremely high in splenic samples (∼2 million reads), a primary lymphoid organ in fish and a common site of ranavirus infection^19^. McRV was isolated in fish cell lines, indicating the presence of an active virus, and PCR confirmed its presence in all brain, liver, and spleen samples as well as in tissue homogenates and tissue culture supernatant samples. Moreover, the lesion observed in the wrasse kidney, being an acute tubulointerstitial necrosis was not typical of a primary tubular nephrosis (e.g. that associated with a renal toxin), and furthermore it is likely that the tubules were co-involved or affected by extension from events in the surrounding interstitial tissue. This is supported by experimental studies of LMBV, which illustrate necrosis of renal tubules as well virus detection in kidney tissue^20^. Despite this strong association, the clinical signs observed here were non-specific, and only a subset of organs was necessarily examined histologically, such that additional data are needed to fully establish causation.

While a genetically identical virus to McRV has been previously reported in Australia – Halichoeres melanurus ranavirus, which was detected in apparently healthy tail-spot wrasses at Orpheus Island on the GBR^13^ – this study provides the first evidence of *Ranavirus micropterus1* disease and the first complete virus genome in Australia. The identification of both viruses approximately four years apart from different reef locations on the GBR, suggests that *Ranavirus micropterus1* may circulate in wild wrasses on the GBR and that these species may be natural hosts. For instance, despite sequencing the metatranscriptomes for virus discovery in over 60 diverse species (representing 16 families) from the same fish community in a spatially restricted ecosystem, the virus was only identified in *H. melanurus*^13^. Similarly, while there were other reef fish species at the holding facility in question, clinical signs and disease were observed exclusively in *M. choati*. As such, the disease outbreak described here may reflect progression from subclinical infection on the reef to overt disease following transfer to captivity. For instance, transport and stocking density are well documented drivers of disease emergence, known to elevate physiological stress, suppress immune function, and increase susceptibility to infectious disease in captive fish relative to their wild counterparts^21^. As a case in point, common carp (*Cyprinus carpio*) exposed to transport at high stocking densities exhibited increased cortisol and glucose levels, together with reductions in lysozyme activity and circulating white blood cell counts^22^. Similarly, a variety of salmonid species exposed to handling and transport stress^23^ all exhibited increases in plasma cortisol and glucose that persisted for hours to days, indicating a universal physiological stress response with the potential to compromise immune function.

Both McRV and Halichoeres melanurus ranavirus fell within a clade in the virus phylogeny comprised predominantly of ranaviruses isolated from farmed largemouth bass in China. Largemouth bass, a fish native to North America, was introduced to China during the 1980s for aquaculture. China now dominates largemouth bass production, accounting for over 99% of the world’s production^17^. In line with this translocation history, *Ranavirus micropterus1* emerged in multiple fish farms in Guangdong Province during 2006-2008, suggesting that it was introduced from North America. Since its detection in China, the virus has since been identified in other parts of Asia in a variety of species including mandarin fish (*Siniperca chuatsi*), barcoo grunter (*Scortum barcoo*), koi (*C. carpio*), damselfish (*Pomacentrus similis*), and snakehead (*Channa maculata*)^24,25,26,27,28^.

A separate translocation route of *Ranavirus micropterus1* is evident via the ornamental fish trade. Viruses such as doctor fish virus and guppy virus 6 were isolated in the USA from ornamental fishes (*Labroides dimidiatus* and *Poecilia reticulata*) imported from Southeast Asia approximately 17 years prior to the emergence of *Ranavirus micropterus1* in China^17,29^. These isolates form a sister group to traditional LMBV isolates within the American clade (clade II), suggesting that *Ranavirus micropterus1* may have originated in Asia, translocated to North America through the ornamental fish trade, and subsequently reintroduced to Asia via largemouth bass aquaculture. The phylogenetic divergence between clades I and II (∼97% nucleotide similarity) is consistent with these host translocation events. Notably, the clustering of Australian wrasse isolates within the Asian clade suggests that this virus may be widely distributed among wild marine fishes throughout the Asia–Pacific region. While the precise pathways of domestic-wild transmission are unknown, it is notable that *Ranavirus micropterus1* can survive in water for over 7 days during which it can be horizontally transmitted among a variety of host species^17^. For example, doctor fish virus was shown to successfully infect rainbow trout (*Oncorhynchus mykiss*) and channel catfish (*Ictalurus punctatus*), two distantly related freshwater species separated by millions of years of evolution, under experimental conditions^29^.

We leveraged our metatranscriptomic approach to examine host RNA – which often make up a very large portion of the data in metatranscriptomic libraries – alongside pathogen discovery. This analysis strongly supported the activity of McRV through the high expression of immune loci, including those linked to antiviral defence (*NMI, PLSCR1, FOSL2*). While this analysis was necessarily exploratory due to the absence of healthy controls, many of these immune-related genes ranked among the 50 most highly expressed transcripts across samples (Figure 2C; Table S2). For example, *TNFRSF1B*, a key mediator of TNF signalling, was among the top five expressed genes, consistent with observations from experimental LMBV infections, where *TNFRSF1B* was also strongly upregulated^30^. Also of note was the extremely high expression of nucleoredoxin (*NXN*), consistent with high viral abundance in spleen samples. Nucleoredoxin is involved in regulating redox balance and was recently shown to inhibit cellular apoptosis in red-lip mullet (*Planiliza haematocheilus*) challenged with viral haemorrhagic septicaemia virus, suggesting a potential role in teleost antiviral immunity^31^. Collectively, these data are strongly suggestive of an active host immune response at the time of sampling.

Taken together, this study reports the first detection of *Ranavirus micropterus1* in diseased captive wrasses in Australia. Its association with wild tropical wrasses strongly suggests that these species are natural hosts, highlighting the need for surveillance to better understand the risk of virus emergence and translocation in domestic fish populations. This is of particular importance given the broad host range of related virus isolates as well as the severe impact of *Ranavirus micropterus1* in farmed fishes in Asia. Overall, we demonstrate the power of combining traditional veterinary diagnostic techniques with metatranscriptomics for investigating unexplained mortalities and show that integrating host transcriptomic data can support evidence of active infection.

## Methods

### Animal Ethics Statement

Animal ethics was approved by The University of Sydney Animal Ethics Committee (AEC) under project number 2025/AE000024.

### Sample collection

A total of 30 leopard wrasses (*M. choati*) were collected from the GBR, Australia, in March 2025 and brought into a multispecies holding facility. Fish showed clinical signs after nine days post transfer, including abnormal swimming behaviour and impaired respiration, with 100% mortality occurring within 1–2 days. Importantly, no other fish species at the facility exhibited these clinical signs. Spleen, liver, and brain tissues were dissected from ten affected individuals and preserved in RNAlater. One individual was placed in fixed formalin. Samples were transferred to the University of Sydney (Camden) and those preserved in RNAlater were immediately stored at −80 °C.

### Histopathology

Whole fish was fixed in 10% neutral buffered formalin for a minimum of 48 h, demineralised in 12.5% EDTA (pH 7.0), and processed for routine histopathology. Tissues were dehydrated through graded ethanol (70%, 95%, and 100%), cleared in xylene, and infiltrated with paraffin wax (TPC-Trio, Medite, Germany). The specimen was embedded in paraffin (Tissue-Tek TEC, Sakura, Japan), sectioned at 4 µm using a rotary microtome (RM2235, Leica, Germany), mounted on glass slides, and stained with haematoxylin and eosin (H&E) using an automated staining system (Tissue-Tek Prisma, Sakura, Japan). Slides were examined by light microscopy with an Olympus BX41 microscope.

### RNA extraction, library preparation, and metatranscriptomic sequencing

Tissue samples (spleen, liver, and brain tissue) from six individuals were processed individually in separate extractions. For homogenisation, each sample was placed in a bashing tube containing 0.1 and 2.0 mm lysis beads (Cat# S9014-50, Zymo Research) and 800 µL of TRIzol (Zymo Research), then vortexed at maximum speed for 10 minutes (Vortex-Genie 2). The homogenate was centrifuged at 12,000 × g for 5 minutes to remove tissue debris. Total RNA was extracted from the clarified supernatant using the Direct-zol RNA Miniprep Plus Kit (Zymo Research, CA, USA) following the manufacturer’s instructions. Library preparation for each sample (n=18) was prepared using the TruSeq Total RNA Library Preparation Protocol (Illumina). Ribo-Zero Plus Kit (Illumina) was employed for ribosomal RNA depletion and paired-end sequencing (150 bp) was conducted on the NovaSeq 6000 platform (Illumina). Library construction and metatranscriptomic sequencing were performed by Macrogen (Korea).

### Virus discovery and transcript abundance

Raw RNA reads were quality trimmed using Trimmomatic (v.0.38)^32^ with the parameters SLIDINGWINDOW:4:5, LEADING:5, TRAILING:5, and MINLEN:25, and assembled into contigs using MEGAHIT (v.1.2.9)^33^, with default parameter settings. Assembled contigs were compared against the protein version of the Reference Viral Database (RVDB-prot) (v.30.0)^34^, NCBI non-redundant protein (nr) and nucleotide (nt) databases (May 2025) using DIAMOND (BLASTX) (v.2.0.9)^35^ and BLASTn with an E-value cut-off of 1 x 10^-5^. Contigs with top matches to the kingdom “Viruses” (NCBI taxid: 10239) were predicted into open reading frames (ORFs) using Geneious Prime (v.2025.2.2)^36^. These contigs were used a query to perform a second search against the NCBI nr/nt databases for validation using Geneious Prime. Viral contig abundances were calculated using RNA-Seq by Expectation Maximization (RSEM) (v.1.3.0)^37^.

### Profiling of bacteria and microbial eukaryotes

To screen for potential bacterial and eukaryotic pathogens, we used Kraken2 (v2.1.3)^38^ with a custom database comprising all nucleotide sequences available on NCBI (with the removal of environmental or artificial sequences). Trimmed reads were classified by applying a minimum confidence threshold of 0.2 and requiring at least one minimum hit group. Bracken (v3.0)^39^ was used to estimate taxon abundances from Kraken2 output files at the species level and taxonomic profiles were initially inspected using Krona^40^ then subsequently analysed in R (v4.3.0) (see below).

### Analysis of potential pathogens

We used measures of transcript abundance, prevalence across tissues, and taxonomy to prioritise candidate pathogens for further investigation. For each detected viral, bacterial, and eukaryotic taxon, we compared relative transcript abundance among samples and tissue types, focusing on agents that were (i) consistently detected across individuals, (ii) present at high abundance relative to other non-host taxa, and (iii) known to infect vertebrate hosts. Taxa detected at low abundance or inconsistently across samples were deprioritised. Statistical comparisons of transcript abundance were performed in R (v4.3.0) and plots were made with the ggplot2 and viridis packages. This analysis identified McRV as the primary candidate pathogen, which was further examined using phylogenetics, host transcriptomics, histopathology, cell culture, and whole-genome sequencing, as described in the sections below.

### Phylogenetic analysis

We constructed phylogenies of the viral subfamily *Alphairidovirinae* and viral species *Ranavirus micropterus1* using nucleotide sequences of the major capsid protein. McRV was aligned with all related complete sequences available on NCBI/GenBank using MAFFT (v7.450) with the E-INS-i algorithm^41^. Alignments were trimmed using TrimAl (v1.2)^42^ with a gap threshold of 0.9 and variable conserve value. The best-fit model of nucleotide substitution was estimated using ModelFinder Plus (-m MFP)^43^ in IQ-TREE (v1.6.12)^44^. Phylogenetic trees were estimated using a maximum-likelihood approach with 1000 bootstrap replicates. Trees were annotated using FigTree (v1.4.4) and Adobe Illustrator.

### Host transcriptomic analysis

Host transcriptomes were assembled de novo using the nf-core/denovotranscript workflow^45^. Briefly, raw reads were trimmed using fastp^46^, and rRNA was removed with SortMeRNA^47^. Transcriptome assembly was performed using a combination of Trinity^48^ and rnaSPAdes^49^, and redundancy reduced using EvidentialGene^50^. Assembly quality was assessed with BUSCO^51^, rnaQUAST^52^, and TransRate^53^. Transcript abundance was quantified with Salmon^54^. Transcriptome annotation was performed using Trinotate (v4.0.2)^55^, executed as a standalone workflow via a Singularity (v3.11.3) container. The annotation integrated DIAMOND/BLAST-based homology searches against UniProtKB/Swiss-Prot (October 2025), Gene Ontology (GO) assignment using the GO slim subset (October 2025), and protein domain annotation using HMMER against the Pfam-A database (July 2025).

Downstream analyses were conducted in R (v4.3.0). Bray–Curtis dissimilarities were calculated from normalised transcript abundance data and visualised using PCoA. Differences in transcriptional profiles among tissue types were tested using PERMANOVA, as implemented in the vegan package^56^. Figures were generated using ggplot2.

### Propagation of BF-2 cell lines

BF-2 cells were maintained at 25 °C in T25 and T75 cm2 flasks using Leibovitz’s L15 medium supplemented (L15) (Gibco, ThermoFisher) with 10% foetal bovine serum (FBS) (Gibco, ThermoFisher). Cells were passaged weekly by removing medium from confluent flasks, rinsing the monolayer with 1X Dulbecco’s phosphate-buffered saline (PBS; Gibco, ThermoFisher), and adding TrypLE Express Enzyme (Gibco, ThermoFisher). After detachment, fresh medium (10% FBS/L15) was added, and cells were split at a 1:3 or 1:4 ratio. Whole spleen tissue from *M. choati* stored in RNAlater at −80 °C was used for virus isolation. Each sample was placed in a bashing tube containing 0.1 mm and 2.0 mm lysis beads and homogenised in 1 mL of homogenising medium (L15 + 2% Anti-anti; Gibco, ThermoFisher) using a TissueLyser (30 Hz, 2 × 40 s cycles; QIAGEN). Homogenates were incubated at 4 °C for 2 hours and centrifuged at 3,100 × g for 10 minutes at 4 °C to collect clarified tissue homogenate supernatants, which were stored at 4 °C until use.

### McRV isolation in BF-2 cells

All challenges were performed under Physical Containment Level 2 (PC2) conditions, in accordance with the Work Health & Safety Regulations at the University of Sydney, Sydney Microscopy & Microanalysis (2022 edition). Virus isolation was performed following the procedures described in the WOAH Aquatic Manual for Infection with Ranavirus species^57^.

BF-2 cells were seeded in 6-well plates at ∼1 × 10^5^ cells per well using growth medium (10% FBS/L15) and incubated at 25 °C for 2 days until 90% confluent. Before inoculation, the growth medium was replaced with maintenance medium (L15 + 2% FBS 1X Anti-anti, Gibco). In one plate (Plate A), two replicate wells were inoculated with 10 µL each of neat (clarified tissue homogenate supernatant) while two wells in a second plate (Plate B) were inoculated with 200 µL each of a 1:10 dilution of clarified wrasse tissue homogenate prepared in homogenising medium. This dilution corresponded to 100 µL of sample per mL of culture medium and represents a 1:100 dilution of a 0.1 mg mL^⁻1^ tissue homogenate. Additional control wells in each plate were inoculated with equivalent volumes of sterile culture medium (negative control) and fish tissue homogenate (spleen from clinically healthy barramundi stored in RNAlater and prepared using the same protocol) to control for potential cytotoxic effects associated with RNAlater.

Plates were incubated at 25 °C and monitored daily for CPE and changes in cell morphology for up to 6 days post-inoculation (dpi). At 2 dpi, culture media was replaced with fresh maintenance medium to maintain nutrient availability. A first passage (P1) was performed by transferring supernatant from a P0 well inoculated with diluted wrasse tissue homogenate to fresh BF-2 monolayers at the same dilution. P1 cultures were incubated at 25 °C and monitored daily for CPE. Supernatants from P0 and P1 cultures were collected and stored at −80 °C for downstream molecular analyses.

### PCR validation

Conventional PCR was performed to confirm the detection of McRV in wrasse tissue samples, wrasse tissue clarified homogenates used for virus isolation and tissue culture supernatants from the virus isolation experiment. DNA was extracted from tissue samples using the QIAamp DNA Micro Kit (QIAGEN) following the manufacturer’s instructions. PCR amplification was carried out using the *H. melanurus* ranavirus (HmRav) primer set (forward: TCAGATGGACCAAAAATCTC; reverse: AGTGTAGTTGGAACCCACAG) (IDT, IA, USA) with an annealing temperature of 50 °C, as previously described^13^. PCR products were analysed by gel electrophoresis using HyperLadder− 100 bp Plus (BIOLINE; Cat. No. BIO-33073; Batch No. H4P5-112I) as the DNA size standard.

### Viral genome sequencing

Total DNA was extracted from tissue culture supernatant using the QIAamp DNA Micro Kit (QIAGEN) following the manufacturer’s instructions. DNA libraries were constructed using the Illumina DNA Prep kit and paired-end sequencing (150 bp) was conducted on the MiSeq i100 Plus (Illumina). Library construction and sequencing were performed by the Ramaciotti Centre for Genomics (University of New South Wales, Australia). Raw DNA reads were quality trimmed using Trimmomatic (v.0.38)^32^ and assembled into contigs using MEGAHIT (v.1.2.9)^33^, with default parameter settings. This revealed a contig of 98,903 nt, which we annotated with the ‘‘Live Annotate and Predict’’ tool implemented in Geneious using related sequences from NCBI/GenBank, with a similarity threshold of 90%. To validate the assembly and estimate coverage, trimmed paired-end reads were mapped to the annotated McRV genome using BBMap (v39.13)^58^.

## Acknowledgements

This work was funded by the Richards Vet Path Bequest by the Sydney School of Veterinary Science (SSVS), The University of Sydney, Australia. The authors acknowledge the staff of the SSVS and thank Anna Waldron and Natalie Schiller for their assistance in the laboratory. We also acknowledge the staff of Veterinary Pathology Diagnostic Services (VPDS) for their support with histopathology, in particular Elaine Chew. We thank Cairns Marine for their assistance with sampling and Dr Luisa Monteiro de Miranda for her contribution to histological slide review.

## Data availability

Raw sequence reads have been deposited in the Sequence Read Archive (NCBI/SRA) under BioProject PRJNA1403177 and the complete viral genome sequence has been deposited in GenBank under accession number PX883867.

## Supplementary data

**Figure S1**. Principal coordinate analysis of host transcriptomes by tissue sample.

**Table S1**. Viral contigs, their abundance, and closest matches on NCBI/GenBank (BLAST).

**Table S2**. Top 1000 expressed host genes, their abundance, and functions.

**Table S3**. Read depth and contigs among sample types.

