## Supplementary Information for "A new lineage of *Ranavirus micropterus1* infects ornamental wrasses from the Great Barrier Reef and causes severe disease in captivity"

**Figure S1. Principal coordinate analysis of host transcriptomes by tissue sample.**

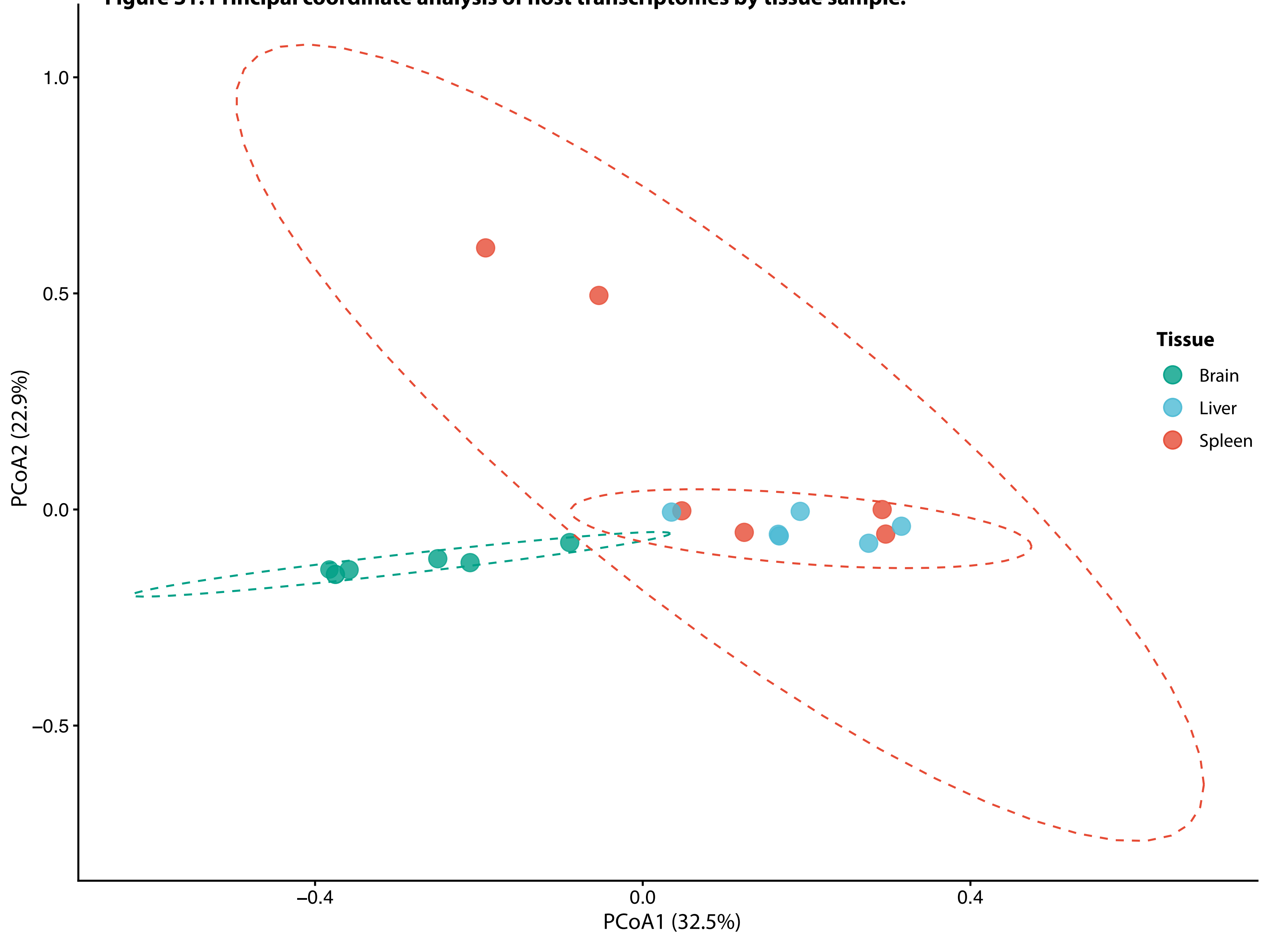

**Table S1. Viral contigs, their abundance, and closest matches on NCBI/GenBank (BLAST).**

| Contig | Library | Length | Top Hit | Percentage | BitScore | E.value | Abundance (reads) |
|---|---|---|---|---|---|---|---|
| k141_104076_flag1_multi4009.0000_len7540 | W1-1 | 7540 | QJE49076.1 putative myristylated membrane protein [Largemouth bass virus] | 99.8 | 503 | 0 | 156397 |
| k141_11782_flag0_multi1729.6004_len3499 | W1-1 | 3499 | WXI69454.1 hypothetical protein [Largemouth bass virus] | 99.3 | 563 | 0 | 22738 |
| k141_123371_flag1_multi1217.6985_len5215 | W1-1 | 5215 | UUY86269.1 hypothetical protein [Largemouth bass virus] | 99.1 | 949 | 0 | 22857 |
| k141_14017_flag0_multi2385.0592_len5308 | W1-1 | 5308 | WEI28960.1 putative D5 family NTPase/ATPase [Largemouth bass virus] | 99.7 | 955 | 0 | 44446 |
| k141_24902_flag0_multi7767.7579_len6672 | W1-1 | 6672 | UUY86232.1 hypothetical protein [Largemouth bass virus] | 99.4 | 492 | 0 | 136039.9 |
| k141_3154_flag1_multi1309.0000_len10308 | W1-1 | 10308 | QJE49122.1 putative DNA-dependent RNA polymerase b subunit [Largemouth bass virus] | 99.6 | 1094 | 0 | 36559 |
| k141_350_flag0_multi2739.6577_len11441 | W1-1 | 11441 | QJE49137.1 hypothetical protein LMBV_074 [Largemouth bass virus] | 99.1 | 1162 | 0 | 108950 |
| k141_56498_flag1_multi3565.8636_len15921 | W1-1 | 15921 | QJE49219.1 hypothetical protein LMBV_070 [Largemouth bass virus] | 99.8 | 879 | 0 | 145873.3 |
| k141_66726_flag0_multi4450.6891_len3776 | W1-1 | 3776 | QJE49145.1 putative proliferating cell nuclear antigen [Largemouth bass virus] | 98.7 | 594 | 0 | 60026 |
| k141_77380_flag0_multi4707.0000_len3152 | W1-1 | 3152 | UUY86258.1 hypothetical protein [Largemouth bass virus] | 99.6 | 566 | 0 | 66684 |
| k141_30232_flag0_multi1159.3025_len3625 | W1-1 | 3625 | AYV88170.1 putative DNA polymerase [Mandarin fish ranavirus] | 99.5 | 1004 | 0 | 17020 |
| k141_70260_flag0_multi1586.0000_len7683 | W1-1 | 7683 | AYV88134.2 putative tyrosine kinase [Mandarin fish ranavirus] | 99.6 | 957 | 0 | 35334 |
| k141_131099_flag1_multi3574.6533_len11182 | W1-1 | 11182 | XRL22828.1 DNA-directed RNA polymerase subunit beta [Siniperca chuatsi ranavirus] | 98.7 | 1369 | 0 | 119903.48 |
| k141_11817_flag1_multi65.5505_len3783 | W1-2 | 3783 | UUY86258.1 hypothetical protein [Largemouth bass virus] | 99.6 | 559 | 0 | 1348 |
| k141_118671_flag1_multi18.4460_len5155 | W1-2 | 5155 | WXI69454.1 hypothetical protein [Largemouth bass virus] | 99.3 | 563 | 0 | 342 |
| k141_126933_flag1_multi148.0000_len2271 | W1-2 | 2271 | QJE49132.1 hypothetical protein LMBV_069 [Largemouth bass virus] | 99.5 | 597 | 0 | 1190 |
| k141_14299_flag1_multi38.9297_len3670 | W1-2 | 3670 | QJE49145.1 putative proliferating cell nuclear antigen [Largemouth bass virus] | 98.7 | 594 | 0 | 502 |
| k141_19382_flag1_multi152.0000_len1759 | W1-2 | 1759 | QJE49076.1 putative myristylated membrane protein [Largemouth bass virus] | 99.8 | 503 | 0 | 961 |
| k141_24832_flag1_multi23.0000_len3956 | W1-2 | 3956 | QJE49137.1 hypothetical protein LMBV_074 [Largemouth bass virus] | 99.1 | 1158 | 0 | 334 |
| k141_33718_flag1_multi67.0000_len3844 | W1-2 | 3844 | UUY86232.1 hypothetical protein [Largemouth bass virus] | 99.4 | 492 | 0 | 912 |
| k141_81627_flag1_multi33.0000_len1752 | W1-2 | 1752 | UUY86267.1 ribonucleotide reductase alpha subunit [Largemouth bass virus] | 100 | 562 | 0 | 197 |
| k141_98543_flag1_multi15.0000_len3769 | W1-2 | 3769 | UUY86269.1 hypothetical protein [Largemouth bass virus] | 99.1 | 949 | 0 | 197 |
| k141_14764_flag1_multi21.0000_len3088 | W1-2 | 3088 | AYV88170.1 putative DNA polymerase [Mandarin fish ranavirus] | 99.5 | 1004 | 0 | 217 |
| k141_49324_flag1_multi20.0000_len3405 | W1-2 | 3405 | AYV88134.2 putative tyrosine kinase [Mandarin fish ranavirus] | 99.6 | 957 | 0 | 243 |
| k141_107887_flag0_multi25.6789_len3290 | W1-2 | 3290 | XRL22821.1 hypothetical protein [Siniperca chuatsi ranavirus] | 99.7 | 952 | 0 | 286 |
| k141_7433_flag1_multi52.9715_len5611 | W1-2 | 5611 | XRL22828.1 DNA-directed RNA polymerase subunit beta [Siniperca chuatsi ranavirus] | 98.6 | 1311 | 0 | 1057 |
| k141_130924_flag1_multi27.0000_len1712 | W1-3 | 1712 | UUY86261.1 hypothetical protein [Largemouth bass virus] | 99.8 | 503 | 0 | 154 |
| k141_213587_flag1_multi23.0000_len1897 | W1-3 | 1897 | QJE49145.1 putative proliferating cell nuclear antigen [Largemouth bass virus] | 98.7 | 594 | 0 | 168 |
| k141_240836_flag1_multi10.0000_len2352 | W1-3 | 2352 | WXI69454.1 hypothetical protein [Largemouth bass virus] | 99.3 | 561 | 0 | 78 |
| k141_250055_flag1_multi8.0000_len2156 | W1-3 | 2156 | WXI69547.1 DNA-dependent RNA polymerase II alpha subunit [Largemouth bass virus] | 99 | 718 | 0 | 57 |
| k141_72834_flag1_multi11.9295_len1843 | W1-3 | 1843 | UUY86267.1 ribonucleotide reductase alpha subunit [Largemouth bass virus] | 99.8 | 547 | 0 | 70 |
| k141_7620_flag0_multi19.0000_len2224 | W1-3 | 2224 | UUY86258.1 hypothetical protein [Largemouth bass virus] | 99.6 | 566 | 0 | 151 |

| | | | | | | | |
|---|---|---|---|---|---|---|---|
| k141_78202_flag1_multi62.0000_len2151 | W1-3 | 2151 | UUY86232.1 hypothetical protein [Largemouth bass virus] | 99.4 | 492 | 0 | 459 |
| k141_25368_flag1_multi416.0000_len1486 | W1-3 | 1486 | UVF58785.1 MAG: major capsid protein [Halichoeres melanurus ranavirus] | 98.3 | 459 | 0 | 2210 |
| k141_94625_flag1_multi66.0000_len1634 | W1-3 | 1634 | UVF58790.1 MAG: myristylated membrane protein, partial [Halichoeres melanurus ranavirus] | 100 | 496 | 0 | 395 |
| k141_1107_flag1_multi10.0000_len1982 | W1-3 | 1982 | AYV88134.2 putative tyrosine kinase [Mandarin fish ranavirus] | 99.4 | 642 | 0 | 72 |
| k141_149749_flag1_multi13.0000_len2499 | W1-3 | 2499 | AYV88199.1 hypothetical protein [Mandarin fish ranavirus] | 99.5 | 746 | 0 | 126 |
| k141_164954_flag1_multi52.0000_len2484 | W1-3 | 2484 | WHA35678.1 hypothetical protein SCRaV_86R [Siniperca chuatsi ranavirus] | 95.8 | 660 | 0 | 493 |
| k141_108285_flag1_multi597.5245_len5340 | W2-1 | 5340 | WXI69454.1 hypothetical protein [Largemouth bass virus] | 99.3 | 556 | 0 | 10782 |
| k141_26739_flag1_multi926.6753_len4872 | W2-1 | 4872 | QJE49122.1 putative DNA-dependent RNA polymerase b subunit [Largemouth bass virus] | 99.6 | 1094 | 0 | 15422 |
| k141_335_flag0_multi1664.4011_len3701 | W2-1 | 3701 | QJE49145.1 putative proliferating cell nuclear antigen [Largemouth bass virus] | 98.7 | 594 | 0 | 22407 |
| k141_35379_flag1_multi2963.4315_len4275 | W2-1 | 4275 | UUY86232.1 hypothetical protein [Largemouth bass virus] | 99.4 | 492 | 0 | 43093 |
| k141_57335_flag1_multi3436.5611_len5297 | W2-1 | 5297 | QJE49219.1 hypothetical protein LMBV_070 [Largemouth bass virus] | 99.8 | 879 | 0 | 119745.15 |
| k141_85599_flag1_multi571.0000_len7666 | W2-1 | 7666 | QJE49137.1 hypothetical protein LMBV_074 [Largemouth bass virus] | 99.1 | 1162 | 0 | 15584 |
| k141_94752_flag1_multi442.7351_len7846 | W2-1 | 7846 | UUY86269.1 hypothetical protein [Largemouth bass virus] | 99.1 | 949 | 0 | 11857 |
| k141_9957_flag1_multi870.0000_len7268 | W2-1 | 7268 | UUY86258.1 hypothetical protein [Largemouth bass virus] | 99.6 | 566 | 0 | 20795.45 |
| k141_50607_flag1_multi13727.0000_len4186 | W2-1 | 4186 | UVF58785.1 MAG: major capsid protein [Halichoeres melanurus ranavirus] | 99.8 | 462 | 0 | 218588.45 |
| k141_108732_flag1_multi513.0632_len12644 | W2-1 | 12644 | AYV88170.1 putative DNA polymerase [Mandarin fish ranavirus] | 99.5 | 1004 | 0 | 20583.24 |
| k141_82684_flag1_multi479.0000_len8192 | W2-1 | 8192 | AYV88134.2 putative tyrosine kinase [Mandarin fish ranavirus] | 99.6 | 957 | 0 | 13572 |
| k141_110649_flag0_multi1198.8755_len3184 | W2-1 | 3184 | XRL22821.1 hypothetical protein [Siniperca chuatsi ranavirus] | 99.7 | 933 | 0 | 11131 |
| k141_50351_flag1_multi1577.9690_len7565 | W2-1 | 7565 | XRL22828.1 DNA-directed RNA polymerase subunit beta [Siniperca chuatsi ranavirus] | 98.7 | 1369 | 0 | 41964.37 |
| k141_108957_flag1_multi161.8953_len3236 | W2-2 | 3236 | WEI28960.1 putative D5 family NTPase/ATPase [Largemouth bass virus] | 99.7 | 955 | 0 | 1893 |
| k141_14040_flag0_multi447.0000_len9258 | W2-2 | 9258 | QJE49076.1 putative myristylated membrane protein [Largemouth bass virus] | 99.8 | 503 | 0 | 12373 |
| k141_1415_flag1_multi277.6054_len3813 | W2-2 | 3813 | UUY86258.1 hypothetical protein [Largemouth bass virus] | 99.6 | 566 | 0 | 5014.41 |
| k141_38373_flag1_multi107.0000_len6712 | W2-2 | 6712 | QJE49122.1 putative DNA-dependent RNA polymerase b subunit [Largemouth bass virus] | 99.6 | 1094 | 0 | 2507 |
| k141_65096_flag1_multi284.6073_len4847 | W2-2 | 4847 | QJE49219.1 hypothetical protein LMBV_070 [Largemouth bass virus] | 99.8 | 879 | 0 | 4896 |
| k141_71280_flag0_multi193.7902_len3669 | W2-2 | 3669 | QJE49145.1 putative proliferating cell nuclear antigen [Largemouth bass virus] | 98.7 | 594 | 0 | 3902 |
| k141_88771_flag1_multi425.0000_len9603 | W2-2 | 9603 | UUY86232.1 hypothetical protein [Largemouth bass virus] | 99.4 | 492 | 0 | 14483 |
| k141_97130_flag1_multi111.0000_len12670 | W2-2 | 12670 | QJE49137.1 hypothetical protein LMBV_074 [Largemouth bass virus] | 99.1 | 1162 | 0 | 4869 |
| k141_137359_flag1_multi175.0991_len6398 | W2-2 | 6398 | AYV88170.1 putative DNA polymerase [Mandarin fish ranavirus] | 99.5 | 1004 | 0 | 4310 |
| k141_7301_flag1_multi101.7366_len15158 | W2-2 | 15158 | AYV88134.2 putative tyrosine kinase [Mandarin fish ranavirus] | 99.6 | 957 | 0 | 5111 |
| k141_32707_flag1_multi244.0000_len5956 | W2-2 | 5956 | XRL22828.1 DNA-directed RNA polymerase subunit beta [Siniperca chuatsi ranavirus] | 98.7 | 1369 | 0 | 5175 |
| k141_127614_flag1_multi1038.1650_len3760 | W2-3 | 3760 | UUY86232.1 hypothetical protein [Largemouth bass virus] | 99.4 | 492 | 0 | 12938 |
| k141_132806_flag1_multi177.6741_len6659 | W2-3 | 6659 | QJE49137.1 hypothetical protein LMBV_074 [Largemouth bass virus] | 99.1 | 1162 | 0 | 3873.86 |
| k141_144911_flag0_multi268.0201_len6158 | W2-3 | 6158 | QJE49122.1 putative DNA-dependent RNA polymerase b subunit [Largemouth bass virus] | 99.6 | 1094 | 0 | 5524 |
| k141_25177_flag1_multi261.0000_len4927 | W2-3 | 4927 | WEI28960.1 putative D5 family NTPase/ATPase [Largemouth bass virus] | 99.7 | 955 | 0 | 4209 |

| | | | | | | | |
|---|---|---|---|---|---|---|---|
| k141_51782_flag1_multi267.0000_len5316 | W2-3 | 5316 | WXI69454.1 hypothetical protein [Largemouth bass virus] | 99.3 | 563 | 0 | 4611 |
| k141_73830_flag1_multi296.0000_len6590 | W2-3 | 6590 | QJE49145.1 putative proliferating cell nuclear antigen [Largemouth bass virus] | 98.7 | 594 | 0 | 6777 |
| k141_77026_flag1_multi396.0000_len4992 | W2-3 | 4992 | QJE49219.1 hypothetical protein LMBV_070 [Largemouth bass virus] | 99.8 | 879 | 0 | 6319 |
| k141_88335_flag1_multi509.6523_len3517 | W2-3 | 3517 | UUY86258.1 hypothetical protein [Largemouth bass virus] | 99.6 | 566 | 0 | 4305.52 |
| k141_96175_flag1_multi55.0000_len6081 | W2-3 | 6081 | UUY86269.1 hypothetical protein [Largemouth bass virus] | 99.1 | 949 | 0 | 1147.19 |
| k141_169500_flag0_multi784.0000_len4173 | W2-3 | 4173 | UVF58785.1 MAG: major capsid protein [Halichoeres melanurus ranavirus] | 100 | 463 | 0 | 19404 |
| k141_143344_flag0_multi325.7912_len6344 | W2-3 | 6344 | AYV88170.1 putative DNA polymerase [Mandarin fish ranavirus] | 99.5 | 1004 | 0 | 6555.45 |
| k141_25074_flag1_multi163.4244_len6371 | W2-3 | 6371 | AYV88134.2 putative tyrosine kinase [Mandarin fish ranavirus] | 99.6 | 957 | 0 | 2273 |
| k141_11980_flag0_multi528.6306_len5062 | W2-3 | 5062 | XRL22828.1 DNA-directed RNA polymerase subunit beta [Siniperca chuatsi ranavirus] | 98.5 | 1138 | 0 | 12228 |
| k141_11371_flag1_multi19.0000_len1717 | W3-1 | 1717 | UUY86258.1 hypothetical protein [Largemouth bass virus] | 99.6 | 566 | 0 | 120 |
| k141_19214_flag1_multi7.0000_len1715 | W3-1 | 1715 | UUY86267.1 ribonucleotide reductase alpha subunit [Largemouth bass virus] | 100 | 561 | 0 | 41 |
| k141_22867_flag1_multi71.0000_len2052 | W3-1 | 2052 | UUY86232.1 hypothetical protein [Largemouth bass virus] | 99.4 | 492 | 0 | 470 |
| k141_32023_flag1_multi7.0000_len2105 | W3-1 | 2105 | QJE49086.1 hypothetical protein LMBV_023 [Largemouth bass virus] | 99.4 | 701 | 0 | 42 |
| k141_3496_flag1_multi24.0000_len1712 | W3-1 | 1712 | QJE49145.1 putative proliferating cell nuclear antigen [Largemouth bass virus] | 98.6 | 560 | 0 | 133 |
| k141_3752_flag1_multi52.0000_len2304 | W3-1 | 2304 | QJE49132.1 hypothetical protein LMBV_069 [Largemouth bass virus] | 99.7 | 606 | 0 | 392 |
| k141_979_flag1_multi61.8659_len5042 | W3-1 | 5042 | UVF58790.1 MAG: myristylated membrane protein, partial [Halichoeres melanurus ranavirus] | 100 | 494 | 0 | 1084.76 |
| k141_22052_flag1_multi9.0000_len2072 | W3-1 | 2072 | AYV88199.1 hypothetical protein [Mandarin fish ranavirus] | 98.4 | 548 | 0 | 56 |
| k141_4092_flag1_multi8.0000_len2986 | W3-1 | 2986 | XRL22821.1 hypothetical protein [Siniperca chuatsi ranavirus] | 99.6 | 919 | 0 | 76 |
| k141_122255_flag1_multi32.0000_len3843 | W3-2 | 3843 | UUY86232.1 hypothetical protein [Largemouth bass virus] | 99.4 | 492 | 0 | 420 |
| k141_47706_flag1_multi24.0000_len3823 | W3-2 | 3823 | WXI69454.1 hypothetical protein [Largemouth bass virus] | 99.3 | 556 | 0 | 302 |
| k141_47902_flag1_multi35.0000_len1827 | W3-2 | 1827 | QJE49145.1 putative proliferating cell nuclear antigen [Largemouth bass virus] | 98.7 | 594 | 0 | 223 |
| k141_64244_flag1_multi42.0000_len2204 | W3-2 | 2204 | QJE49132.1 hypothetical protein LMBV_069 [Largemouth bass virus] | 99.7 | 581 | 0 | 331 |
| k141_65531_flag1_multi33.0000_len1715 | W3-2 | 1715 | UUY86258.1 hypothetical protein [Largemouth bass virus] | 99.6 | 563 | 0 | 195 |
| k141_74124_flag1_multi4.0000_len1638 | W3-2 | 1638 | UUY86269.1 hypothetical protein [Largemouth bass virus] | 99.2 | 525 | 0 | 22 |
| k141_79031_flag1_multi9.0000_len2553 | W3-2 | 2553 | QJE49122.1 putative DNA-dependent RNA polymerase b subunit [Largemouth bass virus] | 99.5 | 846 | 0 | 79 |
| k141_80253_flag1_multi26.8755_len2439 | W3-2 | 2439 | UUY86267.1 ribonucleotide reductase alpha subunit [Largemouth bass virus] | 100 | 562 | 0 | 215 |
| k141_9285_flag1_multi54.0000_len4722 | W3-2 | 4722 | QJE49076.1 putative myristylated membrane protein [Largemouth bass virus] | 99.8 | 503 | 0 | 875 |
| k141_114091_flag1_multi124.0000_len3305 | W3-2 | 3305 | UVF58785.1 MAG: major capsid protein [Halichoeres melanurus ranavirus] | 100 | 463 | 0 | 1534 |
| k141_101487_flag1_multi30.0000_len4884 | W3-2 | 4884 | AYV88134.2 putative tyrosine kinase [Mandarin fish ranavirus] | 99.6 | 957 | 0 | 535 |
| k141_122507_flag0_multi28.8746_len5052 | W3-2 | 5052 | AYV88170.1 putative DNA polymerase [Mandarin fish ranavirus] | 99.5 | 1004 | 0 | 421 |
| k141_95189_flag1_multi13.0000_len2500 | W3-2 | 2500 | AYV88199.1 hypothetical protein [Mandarin fish ranavirus] | 99.5 | 777 | 0 | 117 |
| k141_109742_flag1_multi28.0000_len7125 | W3-2 | 7125 | XRL22828.1 DNA-directed RNA polymerase subunit beta [Siniperca chuatsi ranavirus] | 98.6 | 1261 | 0 | 715 |
| k141_89037_flag1_multi20.8536_len3284 | W3-2 | 3284 | XRL22821.1 hypothetical protein [Siniperca chuatsi ranavirus] | 99.7 | 953 | 0 | 228 |
| k141_136_flag0_multi9804.0000_len6482 | W4-1 | 6482 | QJE49219.1 hypothetical protein LMBV_070 [Largemouth bass virus] | 99.8 | 879 | 0 | 261945.95 |
| k141_13896_flag1_multi1558.9071_len4491 | W4-1 | 4491 | UUY86269.1 hypothetical protein [Largemouth bass virus] | 99.1 | 949 | 0 | 24781 |
| k141_58595_flag1_multi1483.0000_len2910 | W4-1 | 2910 | UUY86267.1 ribonucleotide reductase alpha subunit [Largemouth bass virus] | 100 | 562 | 0 | 16310.08 |

| | | | | | | | |
|---|---|---|---|---|---|---|---|
| k141_70699_flag1_multi9037.1091_len9998 | W4-1 | 9998 | QJE49076.1 putative myristylated membrane protein [Largemouth bass virus] | 99.8 | 503 | 0 | 433969.74 |
| k141_71500_flag1_multi4763.2646_len3418 | W4-1 | 3418 | UUY86258.1 hypothetical protein [Largemouth bass virus] | 99.6 | 566 | 0 | 60669.35 |
| k141_88838_flag1_multi1052.9493_len7818 | W4-1 | 7818 | WXI69454.1 hypothetical protein [Largemouth bass virus] | 99.3 | 535 | 0 | 26432 |
| k141_94885_flag1_multi1935.0016_len9106 | W4-1 | 9106 | QJE49122.1 putative DNA-dependent RNA polymerase b subunit [Largemouth bass virus] | 99.6 | 1094 | 0 | 66440.79 |
| k141_96684_flag1_multi7454.3768_len5813 | W4-1 | 5813 | QJE49145.1 putative proliferating cell nuclear antigen [Largemouth bass virus] | 98.7 | 594 | 0 | 123701.87 |
| k141_101816_flag1_multi1894.0000_len12206 | W4-1 | 12206 | AYV88134.2 putative tyrosine kinase [Mandarin fish ranavirus] | 99.6 | 957 | 0 | 81964 |
| k141_40315_flag0_multi1856.0000_len7957 | W4-1 | 7957 | AYV88170.1 putative DNA polymerase [Mandarin fish ranavirus] | 99.5 | 994 | 0 | 43990 |
| k141_67569_flag0_multi1303.0000_len2724 | W4-1 | 2724 | AYV88199.1 hypothetical protein [Mandarin fish ranavirus] | 99.4 | 466 | 0 | 22449.04 |
| k141_31830_flag1_multi7502.0000_len10097 | W4-1 | 10097 | XRL22828.1 DNA-directed RNA polymerase subunit beta [Siniperca chuatsi ranavirus] | 98.7 | 1369 | 0 | 267531 |
| k141_99002_flag1_multi3285.0000_len4973 | W4-1 | 4973 | XRL22821.1 hypothetical protein [Siniperca chuatsi ranavirus] | 99.7 | 952 | 0 | 57863 |
| k141_114409_flag1_multi94.5383_len4824 | W4-2 | 4824 | QJE49137.1 hypothetical protein LMBV_074 [Largemouth bass virus] | 99.1 | 1162 | 0 | 1604 |
| k141_116824_flag1_multi201.9411_len2075 | W4-2 | 2075 | UUY86267.1 ribonucleotide reductase alpha subunit [Largemouth bass virus] | 100 | 557 | 0 | 1402 |
| k141_34734_flag1_multi91.5859_len5195 | W4-2 | 5195 | WXI69454.1 hypothetical protein [Largemouth bass virus] | 99.3 | 556 | 0 | 1613 |
| k141_512_flag1_multi332.0000_len4007 | W4-2 | 4007 | UUY86232.1 hypothetical protein [Largemouth bass virus] | 99.4 | 492 | 0 | 4515 |
| k141_79837_flag1_multi286.3146_len5983 | W4-2 | 5983 | QJE49219.1 hypothetical protein LMBV_070 [Largemouth bass virus] | 99.8 | 879 | 0 | 6104 |
| k141_91778_flag0_multi773.0000_len5473 | W4-2 | 5473 | QJE49145.1 putative proliferating cell nuclear antigen [Largemouth bass virus] | 98.7 | 594 | 0 | 15692 |
| k141_94638_flag0_multi304.0843_len9363 | W4-2 | 9363 | QJE49076.1 putative myristylated membrane protein [Largemouth bass virus] | 99.8 | 503 | 0 | 8574 |
| k141_95585_flag1_multi309.9457_len3880 | W4-2 | 3880 | UUY86258.1 hypothetical protein [Largemouth bass virus] | 99.6 | 566 | 0 | 4457 |
| k141_97921_flag1_multi47.0000_len3776 | W4-2 | 3776 | UUY86269.1 hypothetical protein [Largemouth bass virus] | 99 | 947 | 0 | 601 |
| k141_101067_flag0_multi114.9077_len2525 | W4-2 | 2525 | AYV88179.2 putative DNA dependent RNA polymerase II second largest subunit [Mandarin fish ranavirus] | 99.7 | 792 | 0 | 479 |
| k141_19186_flag1_multi158.9555_len5621 | W4-2 | 5621 | AYV88170.1 putative DNA polymerase [Mandarin fish ranavirus] | 99.5 | 1004 | 0 | 2997 |
| k141_97174_flag1_multi64.0000_len6146 | W4-2 | 6146 | AYV88134.2 putative tyrosine kinase [Mandarin fish ranavirus] | 98.7 | 954 | 0 | 1392 |
| k141_11277_flag0_multi193.6347_len3267 | W4-2 | 3267 | XRL22821.1 hypothetical protein [Siniperca chuatsi ranavirus] | 99.7 | 951 | 0 | 2131 |
| k141_85139_flag0_multi232.2412_len5970 | W4-2 | 5970 | XRL22828.1 DNA-directed RNA polymerase subunit beta [Siniperca chuatsi ranavirus] | 98.7 | 1369 | 0 | 4856 |
| k141_123303_flag1_multi161.0000_len1826 | W4-3 | 1826 | UUY86258.1 hypothetical protein [Largemouth bass virus] | 99.6 | 566 | 0 | 1057 |
| k141_136013_flag1_multi179.0000_len9582 | W4-3 | 9582 | QJE49219.1 hypothetical protein LMBV_070 [Largemouth bass virus] | 99.8 | 879 | 0 | 6044 |
| k141_187617_flag1_multi67.1190_len3973 | W4-3 | 3973 | QJE49137.1 hypothetical protein LMBV_074 [Largemouth bass virus] | 99 | 1153 | 0 | 974 |
| k141_206346_flag1_multi311.0000_len3621 | W4-3 | 3621 | UUY86232.1 hypothetical protein [Largemouth bass virus] | 99.4 | 492 | 0 | 3957 |
| k141_212115_flag0_multi73.2792_len2369 | W4-3 | 2369 | UUY86267.1 ribonucleotide reductase alpha subunit [Largemouth bass virus] | 100 | 562 | 0 | 575 |
| k141_220412_flag0_multi103.4052_len3951 | W4-3 | 3951 | QJE49145.1 putative proliferating cell nuclear antigen [Largemouth bass virus] | 98.6 | 583 | 0 | 1431 |
| k141_66255_flag1_multi66.0000_len4687 | W4-3 | 4687 | QJE49122.1 putative DNA-dependent RNA polymerase b subunit [Largemouth bass virus] | 99.6 | 1090 | 0 | 1070 |
| k141_81724_flag1_multi44.0000_len3681 | W4-3 | 3681 | UUY86269.1 hypothetical protein [Largemouth bass virus] | 99.1 | 948 | 0 | 567 |
| k141_90633_flag1_multi52.0000_len5041 | W4-3 | 5041 | WXI69454.1 hypothetical protein [Largemouth bass virus] | 99.3 | 563 | 0 | 890 |
| k141_183438_flag0_multi1328.0000_len5783 | W4-3 | 5783 | AVJ54673.1 major capsid protein [Smallmouth bass virus] | 98.5 | 462 | 0 | 17888 |

| | | | | | | | |
|---|---|---|---|---|---|---|---|
| k141_10150_flag1_multi450.0000_len1700 | W4-3 | 1700 | UVF58790.1 MAG: myristylated membrane protein, partial [Halichoeres melanurus ranavirus] | 99.8 | 483 | 0 | 2725 |
| k141_122155_flag1_multi42.0000_len3032 | W4-3 | 3032 | AYV88170.1 putative DNA polymerase [Mandarin fish ranavirus] | 99.5 | 1003 | 0 | 405 |
| k141_51354_flag1_multi70.0000_len3395 | W4-3 | 3395 | AYV88134.2 putative tyrosine kinase [Mandarin fish ranavirus] | 99.6 | 954 | 0 | 839 |
| k141_8642_flag1_multi23.0000_len1393 | W4-3 | 1393 | AYV88130.2 putative ATPase-dependent protease [Mandarin fish ranavirus] | 99.1 | 463 | 0 | 113 |
| k141_149432_flag1_multi52.0000_len4448 | W4-3 | 4448 | XRL22828.1 DNA-directed RNA polymerase subunit beta [Siniperca chuatsi ranavirus] | 98.5 | 1303 | 0 | 716 |
| k141_65910_flag1_multi66.0000_len3105 | W4-3 | 3105 | XRL22821.1 hypothetical protein [Siniperca chuatsi ranavirus] | 99.7 | 953 | 0 | 704 |
| k141_151962_flag1_multi2476.0000_len15670 | W5-1 | 15670 | QJE49122.1 putative DNA-dependent RNA polymerase b subunit [Largemouth bass virus] | 99.6 | 1094 | 0 | 106891 |
| k141_16814_flag1_multi7296.9623_len31929 | W5-1 | 31929 | QJE49137.1 hypothetical protein LMBV_074 [Largemouth bass virus] | 99.1 | 1162 | 0 | 909850.07 |
| k141_72672_flag0_multi3360.0000_len9773 | W5-1 | 9773 | WEI28960.1 putative D5 family NTPase/ATPase [Largemouth bass virus] | 99.7 | 955 | 0 | 142045.34 |
| k141_95291_flag1_multi13356.0000_len10588 | W5-1 | 10588 | QJE49076.1 putative myristylated membrane protein [Largemouth bass virus] | 99.8 | 503 | 0 | 512209.23 |
| k141_115668_flag0_multi13643.0000_len4942 | W5-1 | 4942 | UVF58785.1 MAG: major capsid protein [Halichoeres melanurus ranavirus] | 99.6 | 461 | 0 | 948394.8 |
| k141_136027_flag0_multi4045.1620_len12243 | W5-1 | 12243 | AYV88134.2 putative tyrosine kinase [Mandarin fish ranavirus] | 99.6 | 957 | 0 | 144945.87 |
| k141_150520_flag1_multi5953.1972_len22349 | W5-1 | 22349 | AYV88170.1 putative DNA polymerase [Mandarin fish ranavirus] | 99.5 | 1004 | 0 | 354285.82 |
| k141_6970_flag0_multi8017.0000_len13241 | W5-1 | 13241 | XRL22828.1 DNA-directed RNA polymerase subunit beta [Siniperca chuatsi ranavirus] | 98.7 | 1369 | 0 | 527250.93 |
| k141_10305_flag1_multi9.0000_len1469 | W5-2 | 1469 | QJE49122.1 putative DNA-dependent RNA polymerase b subunit [Largemouth bass virus] | 99.8 | 489 | 0 | 45 |
| k141_108124_flag0_multi15.0000_len2780 | W5-2 | 2780 | WXI69454.1 hypothetical protein [Largemouth bass virus] | 99.3 | 563 | 0 | 146 |
| k141_12015_flag1_multi48.0000_len2001 | W5-2 | 2001 | UUY86232.1 hypothetical protein [Largemouth bass virus] | 99.4 | 492 | 0 | 296 |
| k141_132087_flag1_multi7.0000_len2466 | W5-2 | 2466 | UUY86269.1 hypothetical protein [Largemouth bass virus] | 98.9 | 821 | 0 | 63 |
| k141_150044_flag1_multi7.0000_len2068 | W5-2 | 2068 | WXI69547.1 DNA-dependent RNA polymerase II alpha subunit [Largemouth bass virus] | 99 | 689 | 0 | 47 |
| k141_158116_flag1_multi34.8608_len4781 | W5-2 | 4781 | QJE49219.1 hypothetical protein LMBV_070 [Largemouth bass virus] | 99.8 | 879 | 0 | 492 |
| k141_162810_flag1_multi9.0000_len1986 | W5-2 | 1986 | QJE49137.1 hypothetical protein LMBV_074 [Largemouth bass virus] | 99.6 | 470 | 0 | 58 |
| k141_20688_flag1_multi13.0000_len1769 | W5-2 | 1769 | UUY86267.1 ribonucleotide reductase alpha subunit [Largemouth bass virus] | 100 | 562 | 0 | 79 |
| k141_48388_flag1_multi6.0000_len2011 | W5-2 | 2011 | QJE49137.1 hypothetical protein LMBV_074 [Largemouth bass virus] | 98.7 | 670 | 0 | 37 |
| k141_54394_flag1_multi11.0000_len3143 | W5-2 | 3143 | WEI28960.1 putative D5 family NTPase/ATPase [Largemouth bass virus] | 99.7 | 955 | 0 | 109 |
| k141_61317_flag1_multi28.0000_len1670 | W5-2 | 1670 | UUY86258.1 hypothetical protein [Largemouth bass virus] | 99.6 | 556 | 0 | 153 |
| k141_92401_flag1_multi8.0000_len1405 | W5-2 | 1405 | QJE49086.1 hypothetical protein LMBV_023 [Largemouth bass virus] | 99.4 | 468 | 0 | 40 |
| k141_15294_flag1_multi237.4203_len1616 | W5-2 | 1616 | UVF58785.1 MAG: major capsid protein [Halichoeres melanurus ranavirus] | 99.8 | 455 | 0 | 1481 |
| k141_59161_flag1_multi68.0000_len2196 | W5-2 | 2196 | UVF58790.1 MAG: myristylated membrane protein, partial [Halichoeres melanurus ranavirus] | 99.4 | 494 | 0 | 511 |
| k141_62061_flag1_multi10.0000_len2511 | W5-2 | 2511 | AYV88170.1 putative DNA polymerase [Mandarin fish ranavirus] | 99.5 | 837 | 0 | 80 |
| k141_136644_flag1_multi11.9617_len1865 | W5-3 | 1865 | UUY86267.1 ribonucleotide reductase alpha subunit [Largemouth bass virus] | 100 | 562 | 0 | 64 |
| k141_196909_flag1_multi137.0000_len3174 | W5-3 | 3174 | UUY86232.1 hypothetical protein [Largemouth bass virus] | 99.4 | 492 | 0 | 525.04 |
| k141_216948_flag1_multi24.8417_len2637 | W5-3 | 2637 | UUY86258.1 hypothetical protein [Largemouth bass virus] | 99.6 | 566 | 0 | 353 |
| k141_220256_flag1_multi20.9726_len1672 | W5-3 | 1672 | WXI69454.1 hypothetical protein [Largemouth bass virus] | 99.2 | 517 | 0 | 111 |

| | | | | | | | |
|---|---|---|---|---|---|---|---|
| k141_246267_flag1_multi15.0000_len1645 | W5-3 | 1645 | QJE49132.1 hypothetical protein LMBV_069 [Largemouth bass virus] | 99.5 | 547 | 0 | 77 |
| k141_257081_flag1_multi34.0000_len1774 | W5-3 | 1774 | UUY86261.1 hypothetical protein [Largemouth bass virus] | 99.8 | 503 | 0 | 222 |
| k141_7685_flag0_multi16.8382_len2317 | W5-3 | 2317 | QJE49145.1 putative proliferating cell nuclear antigen [Largemouth bass virus] | 98.7 | 594 | 0 | 114 |
| k141_267055_flag1_multi8.0000_len1355 | W5-3 | 1355 | UVF58793.1 MAG: DNA polymerase [Halichoeres melanurus ranavirus] | 99.8 | 451 | 0 | 32 |
| k141_171238_flag1_multi7.0000_len1401 | W5-3 | 1401 | AYV88199.1 hypothetical protein [Mandarin fish ranavirus] | 98.7 | 464 | 0 | 31 |
| k141_24543_flag1_multi6.0000_len1809 | W5-3 | 1809 | WAK75092.1 putative D5 family NTPase/ATPase [Mandarin fish ranavirus] | 99.5 | 584 | 0 | 39 |
| k141_246234_flag1_multi6.0000_len1528 | W5-3 | 1528 | AYV88134.2 putative tyrosine kinase [Mandarin fish ranavirus] | 99.6 | 503 | 0 | 27 |
| k141_19344_flag0_multi429.1133_len5155 | W6-1 | 5155 | UUY86258.1 hypothetical protein [Largemouth bass virus] | 99.6 | 566 | 0 | 7840.26 |
| k141_20527_flag1_multi107.6398_len1965 | W6-1 | 1965 | UUY86267.1 ribonucleotide reductase alpha subunit [Largemouth bass virus] | 100 | 555 | 0 | 696 |
| k141_22548_flag1_multi176.0000_len4596 | W6-1 | 4596 | QJE49137.1 hypothetical protein LMBV_074 [Largemouth bass virus] | 99.1 | 1162 | 0 | 2760 |
| k141_25837_flag1_multi1932.1295_len7925 | W6-1 | 7925 | QJE49145.1 putative proliferating cell nuclear antigen [Largemouth bass virus] | 98.7 | 594 | 0 | 49374 |
| k141_2784_flag1_multi106.0000_len6083 | W6-1 | 6083 | WXI69454.1 hypothetical protein [Largemouth bass virus] | 99.3 | 551 | 0 | 2001 |
| k141_66286_flag0_multi491.0000_len9199 | W6-1 | 9199 | QJE49219.1 hypothetical protein LMBV_070 [Largemouth bass virus] | 99.8 | 879 | 0 | 16678 |
| k141_80119_flag0_multi138.0000_len6844 | W6-1 | 6844 | QJE49122.1 putative DNA-dependent RNA polymerase b subunit [Largemouth bass virus] | 99.6 | 1094 | 0 | 3268 |
| k141_84444_flag1_multi95.4697_len3837 | W6-1 | 3837 | UUY86269.1 hypothetical protein [Largemouth bass virus] | 98.9 | 947 | 0 | 1231 |
| k141_87520_flag1_multi484.9576_len3752 | W6-1 | 3752 | UUY86232.1 hypothetical protein [Largemouth bass virus] | 99.4 | 492 | 0 | 6321 |
| k141_29519_flag1_multi100.0523_len3719 | W6-1 | 3719 | AYV88134.2 putative tyrosine kinase [Mandarin fish ranavirus] | 99.6 | 957 | 0 | 6752.69 |
| k141_30779_flag0_multi175.0000_len3472 | W6-1 | 3472 | AYV88170.1 putative DNA polymerase [Mandarin fish ranavirus] | 99.5 | 1004 | 0 | 1327 |
| k141_5301_flag1_multi40.0000_len2732 | W6-1 | 2732 | AYV88130.2 putative ATPase-dependent protease [Mandarin fish ranavirus] | 99.1 | 453 | 0 | 330 |
| k141_22578_flag0_multi147.7496_len2489 | W6-1 | 2489 | XRL22821.1 hypothetical protein [Siniperca chuatsi ranavirus] | 100 | 704 | 0 | 1285 |
| k141_23266_flag1_multi354.4227_len5797 | W6-1 | 5797 | XRL22828.1 DNA-directed RNA polymerase subunit beta [Siniperca chuatsi ranavirus] | 98.7 | 1369 | 0 | 7134 |
| k141_130447_flag1_multi48.6197_len2208 | W6-2 | 2208 | QJE49145.1 putative proliferating cell nuclear antigen [Largemouth bass virus] | 98.6 | 560 | 0 | 396 |
| k141_131598_flag1_multi30.9528_len3322 | W6-2 | 3322 | WEI28960.1 putative D5 family NTPase/ATPase [Largemouth bass virus] | 99.5 | 955 | 0 | 359 |
| k141_154690_flag1_multi35.0000_len4785 | W6-2 | 4785 | QJE49137.1 hypothetical protein LMBV_074 [Largemouth bass virus] | 99.1 | 1162 | 0 | 432 |
| k141_16749_flag1_multi15.0000_len3615 | W6-2 | 3615 | UUY86269.1 hypothetical protein [Largemouth bass virus] | 99.1 | 949 | 0 | 193 |
| k141_19388_flag1_multi48.0000_len1753 | W6-2 | 1753 | UUY86267.1 ribonucleotide reductase alpha subunit [Largemouth bass virus] | 100 | 562 | 0 | 288 |
| k141_21673_flag1_multi141.0000_len3505 | W6-2 | 3505 | QJE49076.1 putative myristylated membrane protein [Largemouth bass virus] | 99.8 | 503 | 0 | 1765 |
| k141_37376_flag1_multi92.0000_len3641 | W6-2 | 3641 | UUY86232.1 hypothetical protein [Largemouth bass virus] | 99.4 | 492 | 0 | 1149 |
| k141_5715_flag1_multi34.0000_len4706 | W6-2 | 4706 | QJE49122.1 putative DNA-dependent RNA polymerase b subunit [Largemouth bass virus] | 99.6 | 1093 | 0 | 554 |
| k141_73980_flag0_multi1659.0000_len3644 | W6-2 | 3644 | WXI69454.1 hypothetical protein [Largemouth bass virus] | 99.3 | 563 | 0 | 529 |
| k141_90611_flag0_multi67.4518_len2830 | W6-2 | 2830 | UUY86258.1 hypothetical protein [Largemouth bass virus] | 99.6 | 566 | 0 | 657 |
| k141_99825_flag1_multi131.0000_len4738 | W6-2 | 4738 | QJE49219.1 hypothetical protein LMBV_070 [Largemouth bass virus] | 99.8 | 879 | 0 | 1996 |
| k141_154175_flag1_multi14.9977_len6257 | W6-2 | 6257 | AYV88134.2 putative tyrosine kinase [Mandarin fish ranavirus] | 99.6 | 957 | 0 | 331 |
| k141_44950_flag0_multi22.8314_len3143 | W6-2 | 3143 | AYV88170.1 putative DNA polymerase [Mandarin fish ranavirus] | 99.5 | 999 | 0 | 237 |
| k141_767_flag1_multi13.0000_len1469 | W6-2 | 1469 | WAK75099.1 putative DNA dependent RNA polymerase II largest subunit [Mandarin fish ranavirus] | 100 | 448 | 0 | 67 |

| | | | | | | | |
|---|---|---|---|---|---|---|---|
| k141_89951_flag1_multi78.0000_len4367 | W6-2 | 4367 | XRL22828.1 DNA-directed RNA polymerase subunit beta [Siniperca chuatsi ranavirus] | 98.2 | 926 | 0 | 1209 |
| k141_53133_flag1_multi30.0000_len1411 | W6-3 | 1411 | UVF58785.1 MAG: major capsid protein [Halichoeres melanurus ranavirus] | 100 | 462 | 0 | 145 |
| k141_88354_flag1_multi7.0000_len1491 | W6-3 | 1491 | UVF58790.1 MAG: myristylated membrane protein, partial [Halichoeres melanurus ranavirus] | 100 | 492 | 0 | 38 |
| k141_228895_flag0_multi131.9718_len2343 | W5-3 | 2343 | YP_009506759.1 major capsid protein [Santee-Cooper ranavirus] | 96.5 | 455 | 1.05E-305 | 1152.07 |
| k141_8776_flag1_multi14.8167_len2585 | W3-2 | 2585 | UUY86261.1 hypothetical protein [Largemouth bass virus] | 99.8 | 503 | 1.68E-305 | 131 |
| k141_153794_flag1_multi3.7727_len1254 | W6-3 | 1254 | WEI29006.1 putative 2-cysteine adaptor domain protein [Largemouth bass virus] | 99.8 | 417 | 6.42E-305 | 17 |
| k141_138302_flag1_multi5.0000_len1279 | W5-2 | 1279 | WHA35533.1 putative DNA dependent RNA polymerase A subunit [Micropterus salmoides ranavirus] | 100 | 423 | 8.96E-305 | 21 |
| k141_75056_flag0_multi3077.2812_len2573 | W4-1 | 2573 | QJE49137.1 hypothetical protein LMBV_074 [Largemouth bass virus] | 98.7 | 458 | 8.37E-304 | 18491 |
| k141_87621_flag1_multi58.0000_len2238 | W6-2 | 2238 | WEI29033.1 hypothetical protein [Largemouth bass virus] | 98 | 608 | 3.90E-303 | 525 |
| k141_149008_flag1_multi159.0000_len2228 | W4-3 | 2228 | WEI29033.1 hypothetical protein [Largemouth bass virus] | 98 | 608 | 4.95E-303 | 1476 |
| k141_31245_flag0_multi472.0641_len11236 | W2-2 | 11236 | UVF58785.1 MAG: major capsid protein [Halichoeres melanurus ranavirus] | 100 | 463 | 1.83E-302 | 19462 |
| k141_70752_flag1_multi1921.2657_len2418 | W2-1 | 2418 | WEI29033.1 hypothetical protein [Largemouth bass virus] | 98 | 608 | 5.00E-302 | 20174 |
| k141_15453_flag1_multi21.0000_len2614 | W3-1 | 2614 | UUY86261.1 hypothetical protein [Largemouth bass virus] | 98.4 | 498 | 1.02E-300 | 173 |
| k141_7870_flag1_multi24.0000_len2179 | W3-1 | 2179 | WEI29033.1 hypothetical protein [Largemouth bass virus] | 98 | 604 | 2.98E-300 | 194 |
| k141_22204_flag1_multi53.9771_len2457 | W1-2 | 2457 | UUY86261.1 hypothetical protein [Largemouth bass virus] | 99.8 | 487 | 2.87E-299 | 435 |
| k141_229213_flag1_multi13.0000_len1643 | W5-3 | 1643 | UVF58788.1 MAG: helicase, partial [Halichoeres melanurus ranavirus] | 99.8 | 403 | 6.82E-296 | 70 |
| k141_70101_flag1_multi46.0000_len1236 | W1-2 | 1236 | QJE49149.1 putative immediate early protein ICP-46 [Largemouth bass virus] | 100 | 382 | 7.74E-296 | 193 |
| k141_134802_flag1_multi67.0000_len1244 | W6-2 | 1244 | QJE49149.1 putative immediate early protein ICP-46 [Largemouth bass virus] | 100 | 382 | 8.74E-296 | 272 |
| k141_151400_flag1_multi17.9052_len1755 | W5-2 | 1755 | UVF58788.1 MAG: helicase, partial [Halichoeres melanurus ranavirus] | 99.8 | 403 | 2.85E-295 | 99 |
| k141_51796_flag1_multi8.0000_len1275 | W1-3 | 1275 | UUY86269.1 hypothetical protein [Largemouth bass virus] | 99.1 | 424 | 4.15E-295 | 34 |
| k141_2247_flag1_multi165.8378_len1362 | W4-2 | 1362 | QJE49149.1 putative immediate early protein ICP-46 [Largemouth bass virus] | 100 | 382 | 5.93E-295 | 732 |
| k141_6757_flag1_multi27.0000_len1204 | W3-1 | 1204 | QJE49149.1 putative immediate early protein ICP-46 [Largemouth bass virus] | 99.7 | 382 | 1.73E-294 | 106 |
| k141_71610_flag1_multi17.9846_len1312 | W5-2 | 1312 | QJE49149.1 putative immediate early protein ICP-46 [Largemouth bass virus] | 100 | 380 | 4.26E-293 | 72 |
| k141_69739_flag1_multi84.0000_len4471 | W1-2 | 4471 | WEI29033.1 hypothetical protein [Largemouth bass virus] | 98 | 608 | 2.70E-292 | 1450 |
| k141_203133_flag1_multi18.0000_len1246 | W5-3 | 1246 | QJE49149.1 putative immediate early protein ICP-46 [Largemouth bass virus] | 100 | 378 | 1.69E-291 | 65 |
| k141_94858_flag0_multi126.8455_len2561 | W2-3 | 2561 | UVF58788.1 MAG: helicase, partial [Halichoeres melanurus ranavirus] | 99.8 | 403 | 5.97E-291 | 1021 |
| k141_112871_flag1_multi10.0000_len1658 | W5-2 | 1658 | QJE49110.1 hypothetical protein LMBV_047 [Largemouth bass virus] | 99.5 | 405 | 5.47E-290 | 59 |
| k141_37166_flag1_multi148.0000_len1250 | W4-2 | 1250 | AYV88173.2 putative ribonucleotide reductase beta subunit [Mandarin fish ranavirus] | 100 | 387 | 5.84E-289 | 622 |
| k141_194666_flag1_multi27.0000_len1254 | W4-3 | 1254 | AYV88173.2 putative ribonucleotide reductase beta subunit [Mandarin fish ranavirus] | 100 | 387 | 6.08E-289 | 121 |
| k141_67495_flag1_multi21.0000_len1285 | W1-2 | 1285 | AYV88173.2 putative ribonucleotide reductase beta subunit [Mandarin fish ranavirus] | 100 | 387 | 9.45E-289 | 96 |
| k141_144846_flag1_multi156.0000_len1294 | W2-3 | 1294 | AYV88173.2 putative ribonucleotide reductase beta subunit [Mandarin fish ranavirus] | 100 | 387 | 1.07E-288 | 642 |
| k141_64406_flag1_multi9.0000_len1307 | W5-3 | 1307 | AYV88173.2 putative ribonucleotide reductase beta subunit [Mandarin fish ranavirus] | 100 | 387 | 1.78E-288 | 44 |
| k141_212268_flag1_multi11.0000_len2321 | W5-3 | 2321 | QJE49110.1 hypothetical protein LMBV_047 [Largemouth bass virus] | 99.5 | 405 | 2.16E-286 | 87 |
| k141_52448_flag0_multi784.0000_len6091 | W2-3 | 6091 | WEI29033.1 hypothetical protein [Largemouth bass virus] | 98 | 608 | 2.67E-286 | 8383 |
| k141_86222_flag1_multi13.0000_len1218 | W5-2 | 1218 | AYV88173.2 putative ribonucleotide reductase beta subunit [Mandarin fish ranavirus] | 100 | 384 | 3.07E-286 | 54 |
| k141_128460_flag1_multi29.0000_len2453 | W6-2 | 2453 | QJE49110.1 hypothetical protein LMBV_047 [Largemouth bass virus] | 99.5 | 405 | 1.05E-285 | 258 |
| k141_103485_flag1_multi281.0000_len6336 | W4-2 | 6336 | WEI29033.1 hypothetical protein [Largemouth bass virus] | 98 | 608 | 2.40E-285 | 6612 |

| | | | | | | | |
|---|---|---|---|---|---|---|---|
| k141_31460_flag1_multi9.0000_len3848 | W3-1 | 3848 | UVF58788.1 MAG: helicase, partial [Halichoeres melanurus ranavirus] | 99.8 | 403 | 8.75E-285 | 107 |
| k141_88400_flag1_multi196.4808_len2614 | W6-1 | 2614 | QJE49110.1 hypothetical protein LMBV_047 [Largemouth bass virus] | 99.5 | 405 | 1.43E-284 | 1812 |
| k141_10177_flag1_multi3019.0000_len1951 | W4-1 | 1951 | QJE49149.1 putative immediate early protein ICP-46 [Largemouth bass virus] | 100 | 374 | 1.75E-284 | 20970 |
| k141_70439_flag1_multi44.0000_len2051 | W6-2 | 2051 | AYV88173.2 putative ribonucleotide reductase beta subunit [Mandarin fish ranavirus] | 100 | 387 | 1.79E-284 | 310 |
| k141_140188_flag1_multi9.0000_len1205 | W1-3 | 1205 | UVF58793.1 MAG: DNA polymerase [Halichoeres melanurus ranavirus] | 99.8 | 401 | 3.68E-284 | 33 |
| k141_31296_flag1_multi6.0000_len1179 | W5-2 | 1179 | UUY86211.1 hypothetical protein [Largemouth bass virus] | 99 | 392 | 5.28E-284 | 25 |
| k141_106283_flag1_multi17.0000_len1123 | W1-3 | 1123 | QJE49149.1 putative immediate early protein ICP-46 [Largemouth bass virus] | 100 | 365 | 7.48E-284 | 62 |
| k141_45892_flag1_multi23.5601_len2180 | W3-2 | 2180 | AYV88173.2 putative ribonucleotide reductase beta subunit [Mandarin fish ranavirus] | 100 | 387 | 8.75E-284 | 161 |
| k141_26717_flag1_multi10.0000_len1532 | W3-1 | 1532 | QJE49110.1 hypothetical protein LMBV_047 [Largemouth bass virus] | 99.2 | 398 | 3.09E-283 | 53 |
| k141_99163_flag0_multi85.1957_len2466 | W6-1 | 2466 | AYV88173.2 putative ribonucleotide reductase beta subunit [Mandarin fish ranavirus] | 100 | 387 | 1.55E-281 | 604 |
| k141_71351_flag1_multi26.0000_len3471 | W1-2 | 3471 | QJE49110.1 hypothetical protein LMBV_047 [Largemouth bass virus] | 99.5 | 405 | 7.27E-281 | 325 |
| k141_1143_flag0_multi758.3783_len2639 | W1-1 | 2639 | AYV88173.2 putative ribonucleotide reductase beta subunit [Mandarin fish ranavirus] | 100 | 387 | 1.20E-280 | 6385 |
| k141_140063_flag1_multi59.0000_len2210 | W1-3 | 2210 | UUY86194.1 hypothetical protein [Largemouth bass virus] | 100 | 400 | 1.52E-279 | 501 |
| k141_95404_flag1_multi7.0000_len2414 | W1-2 | 2414 | UUY86211.1 hypothetical protein [Largemouth bass virus] | 99 | 395 | 1.79E-279 | 60 |
| k141_188960_flag1_multi10.0000_len1116 | W1-3 | 1116 | XRL22821.1 hypothetical protein [Siniperca chuatsi ranavirus] | 99.2 | 372 | 2.75E-279 | 38 |
| k141_91115_flag1_multi34.9156_len9001 | W3-2 | 9001 | WEI29033.1 hypothetical protein [Largemouth bass virus] | 98 | 608 | 4.81E-279 | 1036 |
| k141_22886_flag0_multi41.8327_len2621 | W3-1 | 2621 | UUY86194.1 hypothetical protein [Largemouth bass virus] | 99.8 | 401 | 1.41E-277 | 350 |
| k141_103735_flag1_multi87.0000_len4425 | W3-2 | 4425 | QJE49110.1 hypothetical protein LMBV_047 [Largemouth bass virus] | 99.5 | 405 | 1.01E-276 | 1364 |
| k141_162545_flag1_multi21.0000_len1767 | W5-2 | 1767 | QJE49113.1 hypothetical protein LMBV_050 [Largemouth bass virus] | 99.7 | 371 | 1.74E-275 | 115 |
| k141_3646_flag1_multi214.4098_len5580 | W4-3 | 5580 | QJE49149.1 putative immediate early protein ICP-46 [Largemouth bass virus] | 100 | 382 | 2.07E-275 | 4362 |
| k141_214585_flag1_multi64.0000_len4768 | W4-3 | 4768 | QJE49110.1 hypothetical protein LMBV_047 [Largemouth bass virus] | 99.5 | 405 | 2.42E-275 | 771 |
| k141_101172_flag1_multi149.0000_len1306 | W6-1 | 1306 | AYV88121.1 hypothetical protein [Mandarin fish ranavirus] | 98.7 | 374 | 2.80E-275 | 657 |
| k141_119466_flag1_multi29.0000_len1207 | W5-2 | 1207 | QJE49145.1 putative proliferating cell nuclear antigen [Largemouth bass virus] | 100 | 387 | 3.37E-275 | 118 |
| k141_6081_flag1_multi17.0000_len1763 | W5-3 | 1763 | QJE49113.1 hypothetical protein LMBV_050 [Largemouth bass virus] | 99.7 | 371 | 3.38E-275 | 90 |
| k141_122171_flag1_multi245.3165_len2258 | W2-2 | 2258 | AYV88173.2 putative ribonucleotide reductase beta subunit [Mandarin fish ranavirus] | 99.7 | 379 | 5.15E-275 | 1237 |
| k141_113085_flag1_multi25.5077_len1309 | W6-2 | 1309 | AYV88121.1 hypothetical protein [Mandarin fish ranavirus] | 98.7 | 374 | 1.18E-274 | 106 |
| k141_133060_flag1_multi5.0000_len1217 | W5-3 | 1217 | QJE49102.1 putative DNA dependent RNA polymerase a subunit [Largemouth bass virus] | 98.3 | 405 | 5.53E-273 | 19 |
| k141_52929_flag1_multi51.0000_len2435 | W6-2 | 2435 | QJE49113.1 hypothetical protein LMBV_050 [Largemouth bass virus] | 99.7 | 371 | 8.85E-272 | 422 |
| k141_109800_flag1_multi444.0000_len5408 | W2-1 | 5408 | QJE49110.1 hypothetical protein LMBV_047 [Largemouth bass virus] | 99.5 | 405 | 9.16E-272 | 8543 |
| k141_126922_flag1_multi56.0000_len2454 | W4-3 | 2454 | QJE49113.1 hypothetical protein LMBV_050 [Largemouth bass virus] | 99.7 | 371 | 1.09E-271 | 485 |
| k141_59622_flag1_multi171.0000_len5790 | W2-3 | 5790 | QJE49110.1 hypothetical protein LMBV_047 [Largemouth bass virus] | 99.5 | 405 | 1.25E-271 | 3447 |
| k141_118046_flag1_multi515.1995_len5896 | W4-2 | 5896 | QJE49110.1 hypothetical protein LMBV_047 [Largemouth bass virus] | 99.5 | 405 | 2.74E-271 | 10763 |
| k141_72709_flag0_multi9814.7139_len5965 | W4-1 | 5965 | QJE49110.1 hypothetical protein LMBV_047 [Largemouth bass virus] | 99.5 | 405 | 4.72E-271 | 160115 |
| k141_148402_flag1_multi30.0000_len2593 | W1-2 | 2593 | QJE49113.1 hypothetical protein LMBV_050 [Largemouth bass virus] | 99.7 | 371 | 5.48E-271 | 280 |
| k141_85491_flag1_multi145.0000_len4316 | W6-1 | 4316 | QJE49149.1 putative immediate early protein ICP-46 [Largemouth bass virus] | 100 | 368 | 3.46E-270 | 1936 |
| k141_221601_flag1_multi12.0000_len2294 | W5-3 | 2294 | AYV88121.1 hypothetical protein [Mandarin fish ranavirus] | 98.7 | 374 | 7.10E-270 | 83 |
| k141_70364_flag1_multi38.0000_len2651 | W1-2 | 2651 | QJE49122.1 putative DNA-dependent RNA polymerase b subunit [Largemouth bass virus] | 100 | 379 | 1.55E-269 | 361 |
| k141_99425_flag1_multi131.4283_len3349 | W6-2 | 3349 | UUY86194.1 hypothetical protein [Largemouth bass virus] | 100 | 394 | 2.32E-269 | 1305 |

| | | | | | | | |
|---|---|---|---|---|---|---|---|
| k141_191652_flag1_multi10.0000_len1328 | W5-3 | 1328 | XRL22821.1 hypothetical protein [Siniperca chuatsi ranavirus] | 100 | 366 | 1.61E-268 | 39 |
| k141_99763_flag1_multi24.0000_len2262 | W1-2 | 2262 | AYV88121.1 hypothetical protein [Mandarin fish ranavirus] | 98.7 | 372 | 2.24E-268 | 196 |
| k141_75207_flag0_multi172.1696_len3184 | W6-1 | 3184 | QJE49113.1 hypothetical protein LMBV_050 [Largemouth bass virus] | 99.7 | 371 | 4.22E-268 | 2263.37 |
| k141_242733_flag1_multi5.0000_len1107 | W6-3 | 1107 | WEI28972.1 CTD-phosphotransferase [Largemouth bass virus] | 99.2 | 368 | 1.13E-267 | 13 |
| k141_21420_flag1_multi144.0000_len2159 | W4-2 | 2159 | UUY86210.1 putative RNAseIII [Largemouth bass virus] | 99.7 | 385 | 1.15E-267 | 1031 |
| k141_52312_flag1_multi14.0000_len1529 | W1-3 | 1529 | XRL22828.1 DNA-directed RNA polymerase subunit beta [Siniperca chuatsi ranavirus] | 99.2 | 385 | 7.50E-267 | 69 |
| k141_83252_flag1_multi1833.0000_len9377 | W1-1 | 9377 | QJE49149.1 putative immediate early protein ICP-46 [Largemouth bass virus] | 100 | 382 | 2.98E-266 | 49513.48 |
| k141_58798_flag1_multi32.7929_len5970 | W5-3 | 5970 | UUY86194.1 hypothetical protein [Largemouth bass virus] | 100 | 404 | 3.64E-266 | 691 |
| k141_57058_flag0_multi12.7320_len1126 | W1-3 | 1126 | XRL22821.1 hypothetical protein [Siniperca chuatsi ranavirus] | 100 | 374 | 6.71E-266 | 32 |
| k141_150395_flag1_multi31.9936_len6057 | W5-2 | 6057 | UUY86194.1 hypothetical protein [Largemouth bass virus] | 100 | 404 | 6.88E-266 | 679 |
| k141_110311_flag1_multi48.0000_len1105 | W4-3 | 1105 | AYV88121.1 hypothetical protein [Mandarin fish ranavirus] | 98.9 | 361 | 1.26E-265 | 190 |
| k141_105077_flag0_multi681.0766_len3341 | W4-2 | 3341 | AYV88121.1 hypothetical protein [Mandarin fish ranavirus] | 98.7 | 374 | 9.27E-265 | 7376 |
| k141_24471_flag1_multi224.7477_len2757 | W2-3 | 2757 | UUY86210.1 putative RNAseIII [Largemouth bass virus] | 99.7 | 385 | 1.25E-264 | 2033 |
| k141_78623_flag0_multi3166.1765_len2878 | W1-1 | 2878 | UUY86210.1 putative RNAseIII [Largemouth bass virus] | 99.7 | 385 | 3.49E-264 | 21543 |
| k141_119494_flag1_multi68.0000_len1066 | W4-3 | 1066 | UVF58786.1 MAG: 3-beta-hydroxy-delta-5-C27 steroid oxidoreductase, partial [Halichoeres melanurus ranavirus] | 99.7 | 353 | 6.70E-263 | 248 |
| k141_49418_flag1_multi1928.9751_len7203 | W2-1 | 7203 | UUY86194.1 hypothetical protein [Largemouth bass virus] | 100 | 404 | 1.23E-262 | 50402 |
| k141_829_flag1_multi118.5173_len1357 | W6-1 | 1357 | AYV88122.1 putative 3-beta-hydroxy-delta-5-C27 steroid oxidoreductase-like protein [Mandarin fish ranavirus] | 100 | 354 | 1.62E-262 | 689 |
| k141_85950_flag1_multi35.0000_len1456 | W6-2 | 1456 | AYV88122.1 putative 3-beta-hydroxy-delta-5-C27 steroid oxidoreductase-like protein [Mandarin fish ranavirus] | 100 | 354 | 2.06E-262 | 167 |
| k141_223599_flag1_multi15.0000_len1708 | W1-3 | 1708 | QJE49113.1 hypothetical protein LMBV_050 [Largemouth bass virus] | 99.7 | 353 | 4.73E-261 | 93 |
| k141_109392_flag1_multi18.0000_len1230 | W5-2 | 1230 | AYV88122.1 putative 3-beta-hydroxy-delta-5-C27 steroid oxidoreductase-like protein [Mandarin fish ranavirus] | 100 | 349 | 1.34E-259 | 68 |
| k141_89383_flag1_multi18.0000_len1050 | W1-3 | 1050 | AYV88122.1 putative 3-beta-hydroxy-delta-5-C27 steroid oxidoreductase-like protein [Mandarin fish ranavirus] | 99.7 | 349 | 1.40E-259 | 58 |
| k141_34166_flag1_multi26.0000_len1953 | W1-2 | 1953 | UUY86210.1 putative RNAseIII [Largemouth bass virus] | 99.5 | 373 | 2.44E-259 | 178 |
| k141_22634_flag1_multi7.0000_len1076 | W3-1 | 1076 | UUY86223.1 hypothetical protein [Largemouth bass virus] | 99.4 | 358 | 2.46E-259 | 31 |
| k141_89704_flag0_multi4129.2364_len4641 | W1-1 | 4641 | AYV88121.1 hypothetical protein [Mandarin fish ranavirus] | 98.7 | 374 | 2.72E-259 | 83528 |
| k141_159369_flag1_multi55.0000_len1949 | W4-3 | 1949 | UUY86210.1 putative RNAseIII [Largemouth bass virus] | 99.7 | 372 | 4.74E-259 | 369 |
| k141_82687_flag1_multi128.2945_len4932 | W6-1 | 4932 | UUY86210.1 putative RNAseIII [Largemouth bass virus] | 99.7 | 385 | 2.46E-255 | 2540.65 |
| k141_91552_flag1_multi4.0000_len1491 | W3-2 | 1491 | UUY86269.1 hypothetical protein [Largemouth bass virus] | 98.8 | 409 | 6.12E-255 | 24 |
| k141_27183_flag1_multi6.0000_len1069 | W3-1 | 1069 | UVF58793.1 MAG: DNA polymerase [Halichoeres melanurus ranavirus] | 100 | 350 | 6.71E-254 | 19 |
| k141_35943_flag0_multi82.8514_len9469 | W2-1 | 9469 | AYV88173.2 putative ribonucleotide reductase beta subunit [Mandarin fish ranavirus] | 100 | 379 | 1.24E-252 | 6172.86 |
| k141_10750_flag1_multi11.0000_len1023 | W3-1 | 1023 | AYV88122.1 putative 3-beta-hydroxy-delta-5-C27 steroid oxidoreductase-like protein [Mandarin fish ranavirus] | 100 | 340 | 2.87E-252 | 33 |
| k141_35002_flag0_multi153.8863_len13594 | W6-1 | 13594 | QYU76034.1 putative myristylated membrane protein, partial [Koi ranavirus] | 100 | 400 | 4.19E-252 | 7264 |
| k141_257811_flag1_multi3.0000_len1043 | W6-3 | 1043 | AYV88121.1 hypothetical protein [Mandarin fish ranavirus] | 98.8 | 342 | 1.34E-250 | 12 |
| k141_12802_flag1_multi43.0000_len1971 | W4-2 | 1971 | QJE49123.1 putative DNA repair protein RAD2 [Largemouth bass virus] | 99.4 | 353 | 6.35E-249 | 211 |

| | | | | | | | |
|---|---|---|---|---|---|---|---|
| k141_31135_flag0_multi41.0000_len1827 | W3-1 | 1827 | QJE49113.1 hypothetical protein LMBV_050 [Largemouth bass virus] | 99.1 | 341 | 6.31E-248 | 71 |
| k141_258307_flag1_multi14.0000_len1491 | W5-3 | 1491 | QJE49137.1 hypothetical protein LMBV_074 [Largemouth bass virus] | 99.4 | 354 | 7.65E-248 | 61 |
| k141_99651_flag0_multi908.7785_len1202 | W6-2 | 1202 | UVF58785.1 MAG: major capsid protein [Halichoeres melanurus ranavirus] | 100 | 350 | 2.92E-246 | 3387 |
| k141_106459_flag0_multi124.8789_len2594 | W4-2 | 2594 | QJE49122.1 putative DNA-dependent RNA polymerase b subunit [Largemouth bass virus] | 100 | 349 | 9.06E-246 | 1587 |
| k141_66854_flag1_multi9.0000_len1005 | W5-2 | 1005 | QJE49109.1 hypothetical protein LMBV_046 [Largemouth bass virus] | 100 | 324 | 3.05E-245 | 25 |
| k141_184146_flag1_multi33.0000_len1059 | W4-3 | 1059 | QJE49109.1 hypothetical protein LMBV_046 [Largemouth bass virus] | 100 | 324 | 3.26E-245 | 122 |
| k141_53546_flag1_multi44.0000_len1024 | W6-2 | 1024 | QJE49109.1 hypothetical protein LMBV_046 [Largemouth bass virus] | 100 | 324 | 3.90E-245 | 168 |
| k141_166544_flag1_multi59.0000_len3192 | W2-3 | 3192 | QJE49123.1 putative DNA repair protein RAD2 [Largemouth bass virus] | 99.4 | 352 | 3.51E-242 | 601 |
| k141_56684_flag0_multi5668.0000_len1740 | W2-1 | 1740 | QJE49109.1 hypothetical protein LMBV_046 [Largemouth bass virus] | 100 | 324 | 2.10E-241 | 44358 |
| k141_122318_flag0_multi6207.6862_len1846 | W1-1 | 1846 | QJE49109.1 hypothetical protein LMBV_046 [Largemouth bass virus] | 100 | 324 | 1.54E-240 | 101503 |
| k141_168238_flag0_multi583.2094_len3542 | W2-3 | 3542 | QYU76034.1 putative myristylated membrane protein, partial [Koi ranavirus] | 100 | 357 | 3.47E-240 | 8445 |
| k141_154539_flag1_multi84.0000_len1785 | W5-3 | 1785 | QJE49109.1 hypothetical protein LMBV_046 [Largemouth bass virus] | 100 | 318 | 4.44E-235 | 512 |
| k141_112307_flag1_multi436.1117_len6451 | W1-1 | 6451 | QJE49123.1 putative DNA repair protein RAD2 [Largemouth bass virus] | 99.2 | 354 | 6.16E-232 | 8880 |
| k141_102627_flag1_multi274.4999_len5620 | W2-1 | 5620 | AYV88122.1 putative 3-beta-hydroxy-delta-5-C27 steroid oxidoreductase-like protein [Mandarin fish ranavirus] | 100 | 336 | 1.40E-230 | 5970 |
| k141_19190_flag1_multi4.0000_len976 | W3-1 | 976 | QJE49102.1 putative DNA dependent RNA polymerase a subunit [Largemouth bass virus] | 99.4 | 325 | 6.40E-228 | 15 |
| k141_137607_flag1_multi7.0000_len995 | W5-2 | 995 | AYV88134.2 putative tyrosine kinase [Mandarin fish ranavirus] | 100 | 312 | 5.55E-227 | 23 |
| k141_141923_flag1_multi7.0000_len1407 | W5-3 | 1407 | QJE49086.1 hypothetical protein LMBV_023 [Largemouth bass virus] | 99.7 | 313 | 3.86E-226 | 26 |
| k141_78591_flag1_multi11.0000_len1428 | W1-3 | 1428 | QJE49110.1 hypothetical protein LMBV_047 [Largemouth bass virus] | 99.4 | 320 | 1.29E-222 | 55 |
| k141_23331_flag1_multi6.0000_len1020 | W3-1 | 1020 | AYV88179.2 putative DNA dependent RNA polymerase II second largest subunit [Mandarin fish ranavirus] | 99.4 | 317 | 3.31E-222 | 20 |
| k141_33607_flag0_multi302.2347_len5957 | W2-1 | 5957 | QJE49123.1 putative DNA repair protein RAD2 [Largemouth bass virus] | 99.4 | 336 | 4.48E-220 | 2233 |
| k141_121424_flag1_multi9.6115_len1114 | W6-2 | 1114 | QJE49123.1 putative DNA repair protein RAD2 [Largemouth bass virus] | 97.5 | 314 | 5.34E-220 | 39 |
| k141_109491_flag0_multi4415.5043_len1180 | W2-1 | 1180 | AYV88121.1 hypothetical protein [Mandarin fish ranavirus] | 99 | 303 | 8.31E-220 | 2087 |
| k141_89885_flag1_multi13.0000_len899 | W1-3 | 899 | QJE49109.1 hypothetical protein LMBV_046 [Largemouth bass virus] | 100 | 294 | 8.66E-220 | 41 |
| k141_20370_flag1_multi5.0000_len930 | W5-3 | 930 | QJE49122.1 putative DNA-dependent RNA polymerase b subunit [Largemouth bass virus] | 100 | 309 | 1.00E-219 | 14 |
| k141_85822_flag1_multi7.0000_len951 | W1-2 | 951 | AYV88179.2 putative DNA dependent RNA polymerase II second largest subunit [Mandarin fish ranavirus] | 99.7 | 312 | 1.33E-219 | 18 |
| k141_41740_flag1_multi23.0000_len1866 | W1-3 | 1866 | WEI29033.1 hypothetical protein [Largemouth bass virus] | 97.6 | 495 | 2.29E-216 | 169 |
| k141_100681_flag1_multi17.0000_len1071 | W1-3 | 1071 | UVF58788.1 MAG: helicase, partial [Halichoeres melanurus ranavirus] | 100 | 305 | 4.50E-215 | 69 |
| k141_21218_flag0_multi148.0000_len1025 | W6-1 | 1025 | WAK75092.1 putative D5 family NTPase/ATPase [Mandarin fish ranavirus] | 98.3 | 297 | 4.87E-212 | 291 |
| k141_124415_flag1_multi41.0000_len1865 | W1-2 | 1865 | UUY86193.1 hypothetical protein [Largemouth bass virus] | 100 | 290 | 1.69E-211 | 253 |
| k141_73519_flag1_multi3.0000_len971 | W5-3 | 971 | AYV88179.2 putative DNA dependent RNA polymerase II second largest subunit [Mandarin fish ranavirus] | 100 | 299 | 2.38E-208 | 10 |
| k141_89944_flag1_multi211.0000_len1999 | W4-2 | 1999 | WHA35577.1 putative ABC-ATPase [Micropterus salmoides ranavirus] | 97.7 | 300 | 3.24E-207 | 1471 |
| k141_4641_flag1_multi17.0000_len1805 | W3-1 | 1805 | UUY86193.1 hypothetical protein [Largemouth bass virus] | 100 | 281 | 5.67E-205 | 92 |
| k141_54207_flag1_multi7.0000_len852 | W1-2 | 852 | QJE49123.1 putative DNA repair protein RAD2 [Largemouth bass virus] | 99.6 | 283 | 4.25E-203 | 19 |

| | | | | | | | |
|---|---|---|---|---|---|---|---|
| k141_66094_flag1_multi74.0000_len1732 | W4-3 | 1732 | QJE49097.1 hypothetical protein LMBV_034 [Largemouth bass virus] | 100 | 297 | 1.95E-201 | 457 |
| k141_73874_flag1_multi34.0000_len1718 | W6-2 | 1718 | QJE49097.1 hypothetical protein LMBV_034 [Largemouth bass virus] | 100 | 297 | 2.31E-201 | 200 |
| k141_107520_flag1_multi116.0000_len1731 | W2-2 | 1731 | QJE49097.1 hypothetical protein LMBV_034 [Largemouth bass virus] | 100 | 297 | 2.67E-201 | 692 |
| k141_148463_flag1_multi43.0000_len1971 | W1-2 | 1971 | UUY86263.1 hypothetical protein [Largemouth bass virus] | 97.6 | 292 | 4.79E-201 | 296 |
| k141_22343_flag1_multi25.0000_len1968 | W1-3 | 1968 | UUY86263.1 hypothetical protein [Largemouth bass virus] | 97.6 | 292 | 4.79E-201 | 187 |
| k141_93444_flag1_multi3915.0000_len4338 | W4-1 | 4338 | UUY86193.1 hypothetical protein [Largemouth bass virus] | 100 | 290 | 6.17E-201 | 58875.14 |
| k141_212791_flag1_multi15.0000_len1545 | W1-3 | 1545 | UUY86192.1 putative myristylated membrane protein [Largemouth bass virus] | 99.6 | 269 | 9.77E-201 | 84 |
| k141_152258_flag1_multi38.1159_len1728 | W6-2 | 1728 | UUY86192.1 putative myristylated membrane protein [Largemouth bass virus] | 99.6 | 270 | 1.07E-200 | 243 |
| k141_29360_flag1_multi15.0000_len1923 | W3-1 | 1923 | UUY86263.1 hypothetical protein [Largemouth bass virus] | 97.6 | 291 | 1.63E-200 | 95 |
| k141_29722_flag1_multi210.0507_len1898 | W6-1 | 1898 | QJE49097.1 hypothetical protein LMBV_034 [Largemouth bass virus] | 100 | 297 | 1.95E-200 | 1400.17 |
| k141_111157_flag1_multi96.8236_len1955 | W4-2 | 1955 | QJE49097.1 hypothetical protein LMBV_034 [Largemouth bass virus] | 100 | 297 | 3.80E-200 | 661 |
| k141_150271_flag1_multi54.9155_len2200 | W6-2 | 2200 | UUY86263.1 hypothetical protein [Largemouth bass virus] | 97.6 | 292 | 6.65E-200 | 438 |
| k141_28312_flag1_multi10.0000_len1608 | W3-1 | 1608 | QJE49097.1 hypothetical protein LMBV_034 [Largemouth bass virus] | 100 | 295 | 8.39E-200 | 48 |
| k141_39200_flag1_multi5.0000_len1254 | W5-2 | 1254 | AYV88131.1 putative eIF-2 alpha-like protein [Mandarin fish ranavirus] | 99.6 | 270 | 1.95E-199 | 16 |
| k141_15001_flag1_multi16.0000_len1667 | W5-3 | 1667 | QJE49097.1 hypothetical protein LMBV_034 [Largemouth bass virus] | 100 | 294 | 2.75E-198 | 88 |
| k141_27532_flag0_multi69.0000_len1484 | W5-3 | 1484 | QYU76034.1 putative myristylated membrane protein, partial [Koi ranavirus] | 100 | 273 | 1.08E-196 | 331 |
| k141_11746_flag1_multi16.0000_len1738 | W1-2 | 1738 | QJE49097.1 hypothetical protein LMBV_034 [Largemouth bass virus] | 100 | 291 | 1.47E-196 | 106 |
| k141_250457_flag1_multi20.5879_len4446 | W5-3 | 4446 | WHA35577.1 putative ABC-ATPase [Micropterus salmoides ranavirus] | 97.7 | 299 | 1.80E-196 | 295 |
| k141_77103_flag0_multi662.3691_len2747 | W2-1 | 2747 | QJE49097.1 hypothetical protein LMBV_034 [Largemouth bass virus] | 100 | 297 | 2.14E-196 | 6638 |
| k141_108012_flag1_multi7.0000_len1181 | W5-3 | 1181 | QJE49141.1 putative NTPase [Largemouth bass virus] | 98.9 | 279 | 8.47E-195 | 23 |
| k141_101067_flag1_multi29.0000_len1951 | W5-2 | 1951 | UUY86263.1 hypothetical protein [Largemouth bass virus] | 97.5 | 283 | 1.30E-194 | 198 |
| k141_122121_flag1_multi139.0000_len1944 | W4-3 | 1944 | UUY86263.1 hypothetical protein [Largemouth bass virus] | 97.5 | 282 | 4.89E-194 | 982 |
| k141_135793_flag0_multi310.8017_len2294 | W2-3 | 2294 | UUY86263.1 hypothetical protein [Largemouth bass virus] | 97.2 | 283 | 9.74E-192 | 2466 |
| k141_32698_flag1_multi18.0000_len1930 | W3-2 | 1930 | AYV88194.1 hypothetical protein [Mandarin fish ranavirus] | 99.6 | 281 | 9.99E-192 | 127 |
| k141_116161_flag1_multi296.4162_len2217 | W6-1 | 2217 | UUY86263.1 hypothetical protein [Largemouth bass virus] | 97.2 | 281 | 3.35E-191 | 2271 |
| k141_158938_flag0_multi495.7268_len983 | W2-3 | 983 | WHA35533.1 putative DNA dependent RNA polymerase A subunit [Micropterus salmoides ranavirus] | 100 | 277 | 9.34E-190 | 94 |
| k141_184467_flag1_multi4.0000_len1197 | W6-3 | 1197 | WAK75070.1 putative neurofilament triplet H1-like protein [Mandarin fish ranavirus] | 100 | 398 | 1.43E-185 | 14 |
| k141_258402_flag1_multi16.0000_len1160 | W5-3 | 1160 | XRL22828.1 DNA-directed RNA polymerase subunit beta [Siniperca chuatsi ranavirus] | 98.9 | 281 | 1.89E-185 | 57 |
| k141_223620_flag1_multi6.0000_len820 | W5-3 | 820 | QJE49123.1 putative DNA repair protein RAD2 [Largemouth bass virus] | 99.6 | 253 | 2.73E-180 | 14 |
| k141_27788_flag1_multi15.0000_len1296 | W5-2 | 1296 | QJE49098.1 hypothetical protein LMBV_035 [Largemouth bass virus] | 99.2 | 250 | 3.12E-179 | 67 |
| k141_162408_flag1_multi5.0000_len750 | W5-2 | 750 | QJE49123.1 putative DNA repair protein RAD2 [Largemouth bass virus] | 99.6 | 249 | 1.70E-177 | 7 |
| k141_22533_flag1_multi19.0000_len1287 | W3-1 | 1287 | AYV88176.1 putative tumor necrosis factor receptor [Mandarin fish ranavirus] | 100 | 237 | 4.21E-177 | 80 |
| k141_3833_flag1_multi4.0000_len751 | W3-1 | 751 | XPZ21295.1 putative RNAseIII [Mandarin fish ranavirus] | 100 | 249 | 3.26E-176 | 9 |
| k141_63892_flag1_multi26.0000_len1460 | W5-2 | 1460 | AYV88176.1 putative tumor necrosis factor receptor [Mandarin fish ranavirus] | 100 | 237 | 5.09E-176 | 161 |
| k141_8377_flag1_multi28.0000_len1441 | W1-3 | 1441 | AYV88176.1 putative tumor necrosis factor receptor [Mandarin fish ranavirus] | 100 | 237 | 8.24E-176 | 142 |
| k141_113380_flag1_multi39.9125_len1478 | W3-2 | 1478 | AYV88176.1 putative tumor necrosis factor receptor [Mandarin fish ranavirus] | 100 | 237 | 8.98E-176 | 205 |
| k141_12772_flag1_multi30.4192_len2004 | W6-2 | 2004 | UUY86210.1 putative RNAseIII [Largemouth bass virus] | 100 | 258 | 1.40E-174 | 204 |
| k141_213726_flag1_multi29.0000_len1377 | W5-3 | 1377 | AYV88176.1 putative tumor necrosis factor receptor [Mandarin fish ranavirus] | 100 | 234 | 1.73E-174 | 132 |

| | | | | | | | |
|---|---|---|---|---|---|---|---|
| k141_12973_flag0_multi9.8718_len1685 | W3-1 | 1685 | QJE49141.1 putative NTPase [Largemouth bass virus] | 98.8 | 253 | 2.03E-172 | 58 |
| k141_81191_flag0_multi16.0000_len1208 | W5-3 | 1208 | WAK75136.1 hypothetical protein [Mandarin fish ranavirus] | 100 | 243 | 2.89E-172 | 60 |
| k141_90160_flag1_multi9.0000_len1170 | W1-3 | 1170 | WAK75136.1 hypothetical protein [Mandarin fish ranavirus] | 100 | 243 | 7.15E-172 | 30 |
| k141_32708_flag1_multi43.0000_len1290 | W6-2 | 1290 | WAK75136.1 hypothetical protein [Mandarin fish ranavirus] | 100 | 243 | 7.97E-172 | 186 |
| k141_157714_flag1_multi139.0000_len896 | W5-2 | 896 | QJE49073.1 putative p31K protein [Largemouth bass virus] | 99.2 | 262 | 8.24E-171 | 387 |
| k141_35782_flag1_multi10.7099_len1675 | W3-1 | 1675 | WAK75136.1 hypothetical protein [Mandarin fish ranavirus] | 100 | 243 | 8.16E-170 | 65 |
| k141_140860_flag1_multi7.0000_len742 | W5-2 | 742 | AYV88179.2 putative DNA dependent RNA polymerase II second largest subunit [Mandarin fish ranavirus] | 99.2 | 247 | 3.09E-169 | 16 |
| k141_8215_flag1_multi208.0368_len1038 | W3-2 | 1038 | XRB52768.1 P31K protein, partial [Largemouth bass virus] | 98 | 246 | 4.49E-169 | 801 |
| k141_213102_flag1_multi98.0000_len881 | W5-3 | 881 | QJE49073.1 putative p31K protein [Largemouth bass virus] | 99.2 | 242 | 6.46E-169 | 269 |
| k141_253_flag1_multi11.0000_len1204 | W1-3 | 1204 | QJE49098.1 hypothetical protein LMBV_035 [Largemouth bass virus] | 98.7 | 235 | 2.51E-168 | 42 |
| k141_88962_flag0_multi69.0000_len1056 | W5-3 | 1056 | UUY86204.1 hypothetical protein [Largemouth bass virus] | 100 | 274 | 2.94E-167 | 167 |
| k141_154919_flag1_multi18.3374_len2278 | W5-2 | 2278 | WAK75136.1 hypothetical protein [Mandarin fish ranavirus] | 100 | 243 | 6.37E-167 | 150 |
| k141_127817_flag1_multi321.0000_len988 | W6-2 | 988 | XRB52768.1 P31K protein, partial [Largemouth bass virus] | 96.3 | 246 | 9.00E-167 | 1159 |
| k141_75134_flag1_multi51.0000_len869 | W4-3 | 869 | UUY86216.1 hypothetical protein [Largemouth bass virus] | 99.6 | 248 | 9.07E-167 | 158 |
| k141_8920_flag0_multi142.8769_len1701 | W1-2 | 1701 | QJE49073.1 putative p31K protein [Largemouth bass virus] | 99.2 | 262 | 1.25E-166 | 879.17 |
| k141_56578_flag1_multi28.6446_len1016 | W5-2 | 1016 | UUY86216.1 hypothetical protein [Largemouth bass virus] | 99.6 | 248 | 6.20E-166 | 105 |
| k141_127394_flag1_multi622.5455_len977 | W4-3 | 977 | AYV88120.1 putative p31K protein [Mandarin fish ranavirus] | 97.9 | 241 | 6.99E-165 | 2402 |
| k141_107697_flag1_multi102.6397_len1007 | W6-2 | 1007 | UUY86216.1 hypothetical protein [Largemouth bass virus] | 99.2 | 248 | 1.84E-164 | 330 |
| k141_15534_flag1_multi39.0000_len1009 | W3-1 | 1009 | UUY86216.1 hypothetical protein [Largemouth bass virus] | 99.2 | 248 | 1.91E-164 | 41 |
| k141_53718_flag1_multi2.0000_len719 | W6-3 | 719 | XPZ21261.1 hypothetical protein MRVORF002 [Mandarin fish ranavirus] | 99.6 | 239 | 2.77E-164 | 3 |
| k141_129963_flag0_multi1505.4252_len1411 | W1-1 | 1411 | UUY86216.1 hypothetical protein [Largemouth bass virus] | 99.6 | 248 | 2.94E-164 | 12239 |
| k141_32404_flag1_multi15.0000_len1296 | W4-2 | 1296 | ADB77862.1 DNA methyltransferase [Largemouth bass ulcerative syndrome virus] | 99.5 | 220 | 3.17E-164 | 63 |
| k141_104114_flag0_multi322.1731_len1198 | W6-1 | 1198 | UUY86216.1 hypothetical protein [Largemouth bass virus] | 99.6 | 247 | 1.01E-163 | 1759 |
| k141_35669_flag1_multi6.0000_len726 | W3-1 | 726 | AYV88179.2 putative DNA dependent RNA polymerase II second largest subunit [Mandarin fish ranavirus] | 100 | 241 | 8.83E-163 | 12 |
| k141_71093_flag1_multi588.2269_len1697 | W2-3 | 1697 | UUY86216.1 hypothetical protein [Largemouth bass virus] | 99.6 | 248 | 1.75E-162 | 3781.33 |
| k141_43277_flag1_multi6.0000_len800 | W3-2 | 800 | QJE49083.1 hypothetical protein LMBV_020 [Largemouth bass virus] | 99.2 | 244 | 1.80E-162 | 15 |
| k141_13407_flag1_multi39.0000_len1361 | W1-2 | 1361 | UUY86216.1 hypothetical protein [Largemouth bass virus] | 99.2 | 246 | 4.44E-162 | 220 |
| k141_34592_flag1_multi428.9965_len2407 | W3-1 | 2407 | XRB52768.1 P31K protein, partial [Largemouth bass virus] | 98.4 | 245 | 5.77E-162 | 587 |
| k141_91010_flag1_multi98.0000_len835 | W1-3 | 835 | QJE49073.1 putative p31K protein [Largemouth bass virus] | 99.6 | 250 | 2.02E-161 | 227 |
| k141_103027_flag1_multi10.0000_len659 | W5-2 | 659 | UUY86238.1 hypothetical protein [Largemouth bass virus] | 99.5 | 219 | 4.15E-161 | 22 |
| k141_97128_flag1_multi16258.1688_len3305 | W4-1 | 3305 | UUY86197.1 hypothetical protein [Largemouth bass virus] | 100 | 331 | 4.18E-161 | 181883.17 |
| k141_107474_flag1_multi1004.3072_len1902 | W2-1 | 1902 | UUY86216.1 hypothetical protein [Largemouth bass virus] | 99.6 | 248 | 5.09E-161 | 6841 |
| k141_45526_flag1_multi85.0000_len1952 | W4-3 | 1952 | ADB77862.1 DNA methyltransferase [Largemouth bass ulcerative syndrome virus] | 99.5 | 220 | 6.22E-161 | 564 |
| k141_70128_flag1_multi21.2250_len6302 | W3-2 | 6302 | QJE49123.1 putative DNA repair protein RAD2 [Largemouth bass virus] | 92.7 | 273 | 1.61E-160 | 301 |
| k141_12543_flag1_multi12.0000_len1371 | W3-1 | 1371 | AYV88210.2 putative proliferating cell nuclear antigen [Mandarin fish ranavirus] | 100 | 240 | 5.34E-160 | 53 |
| k141_168285_flag1_multi16.0000_len1347 | W5-3 | 1347 | AYV88210.2 putative proliferating cell nuclear antigen [Mandarin fish ranavirus] | 100 | 241 | 8.03E-160 | 68 |
| k141_15188_flag1_multi20.0000_len1359 | W1-3 | 1359 | AYV88210.2 putative proliferating cell nuclear antigen [Mandarin fish ranavirus] | 100 | 240 | 9.29E-160 | 98 |
| k141_115771_flag1_multi66.0000_len1361 | W4-3 | 1361 | AYV88210.2 putative proliferating cell nuclear antigen [Mandarin fish ranavirus] | 100 | 240 | 9.64E-160 | 319 |

| | | | | | | | |
|---|---|---|---|---|---|---|---|
| k141_79051_flag1_multi24.0000_len1360 | W5-2 | 1360 | AYV88210.2 putative proliferating cell nuclear antigen [Mandarin fish ranavirus] | 100 | 240 | 9.64E-160 | 117 |
| k141_70625_flag1_multi980.0000_len1517 | W2-1 | 1517 | AYV88210.2 putative proliferating cell nuclear antigen [Mandarin fish ranavirus] | 100 | 242 | 2.17E-159 | 5265 |
| k141_95025_flag1_multi473.4332_len1600 | W2-3 | 1600 | AYV88210.2 putative proliferating cell nuclear antigen [Mandarin fish ranavirus] | 100 | 242 | 5.79E-159 | 2535 |
| k141_7594_flag1_multi6.0000_len1229 | W3-1 | 1229 | WHA35533.1 putative DNA dependent RNA polymerase A subunit [Micropterus salmoides ranavirus] | 93.5 | 260 | 7.33E-159 | 17 |
| k141_166887_flag1_multi3.0000_len706 | W5-3 | 706 | WHA35533.1 putative DNA dependent RNA polymerase A subunit [Micropterus salmoides ranavirus] | 100 | 234 | 7.71E-158 | 7 |
| k141_85479_flag1_multi30.0000_len1400 | W3-2 | 1400 | AYV88210.2 putative proliferating cell nuclear antigen [Mandarin fish ranavirus] | 100 | 239 | 1.03E-157 | 141 |
| k141_80469_flag1_multi243.0000_len2992 | W6-1 | 2992 | ADB77862.1 DNA methyltransferase [Largemouth bass ulcerative syndrome virus] | 99.5 | 220 | 1.63E-156 | 2032 |
| k141_4095_flag1_multi10.0000_len705 | W1-3 | 705 | QJE49083.1 hypothetical protein LMBV_020 [Largemouth bass virus] | 99.6 | 234 | 3.40E-156 | 22 |
| k141_4696_flag1_multi6078.0000_len5033 | W1-1 | 5033 | UUY86197.1 hypothetical protein [Largemouth bass virus] | 100 | 331 | 3.55E-156 | 93127.92 |
| k141_64825_flag1_multi233.2356_len1703 | W2-2 | 1703 | AYV88210.2 putative proliferating cell nuclear antigen [Mandarin fish ranavirus] | 100 | 237 | 4.03E-155 | 1548 |
| k141_29827_flag1_multi1573.1261_len2402 | W1-1 | 2402 | AYV88210.2 putative proliferating cell nuclear antigen [Mandarin fish ranavirus] | 99.6 | 242 | 6.15E-155 | 14045.63 |
| k141_50685_flag1_multi98.0000_len1209 | W1-3 | 1209 | WXI69548.1 hypothetical protein [Largemouth bass virus] | 99.6 | 224 | 1.47E-154 | 459 |
| k141_199569_flag1_multi510.6626_len1217 | W4-3 | 1217 | WXI69548.1 hypothetical protein [Largemouth bass virus] | 99.6 | 224 | 1.59E-154 | 2358 |
| k141_40951_flag1_multi369.1714_len3151 | W4-2 | 3151 | UUY86216.1 hypothetical protein [Largemouth bass virus] | 99.6 | 246 | 2.72E-154 | 1354 |
| k141_254628_flag1_multi9.0000_len1028 | W6-3 | 1028 | WXI69548.1 hypothetical protein [Largemouth bass virus] | 99.5 | 222 | 4.93E-154 | 31 |
| k141_37707_flag1_multi7825.0000_len4325 | W2-1 | 4325 | QJE49073.1 putative p31K protein [Largemouth bass virus] | 99.2 | 259 | 5.47E-154 | 120377 |
| k141_133264_flag1_multi14.0000_len629 | W1-3 | 629 | AYV88168.1 hypothetical protein [Mandarin fish ranavirus] | 99 | 207 | 1.04E-153 | 31 |
| k141_19935_flag1_multi81.1913_len1343 | W3-1 | 1343 | WXI69548.1 hypothetical protein [Largemouth bass virus] | 99.6 | 223 | 4.24E-153 | 359 |
| k141_89183_flag0_multi12.6292_len853 | W5-2 | 853 | AYV88168.1 hypothetical protein [Mandarin fish ranavirus] | 99 | 207 | 2.90E-152 | 28 |
| k141_32475_flag1_multi50.1649_len2142 | W5-2 | 2142 | WXI69548.1 hypothetical protein [Largemouth bass virus] | 99.6 | 226 | 2.92E-152 | 370 |
| k141_63690_flag1_multi373.9203_len1183 | W6-2 | 1183 | AYV88159.1 hypothetical protein [Mandarin fish ranavirus] | 100 | 214 | 5.49E-152 | 1643 |
| k141_104197_flag1_multi10.0000_len1328 | W5-3 | 1328 | AYV88142.2 hypothetical protein [Mandarin fish ranavirus] | 100 | 223 | 6.04E-151 | 39 |
| k141_208011_flag1_multi70.0000_len1858 | W1-3 | 1858 | AYV88159.1 hypothetical protein [Mandarin fish ranavirus] | 100 | 216 | 3.08E-150 | 467 |
| k141_143040_flag1_multi43.0000_len1446 | W1-2 | 1446 | AYV88210.2 putative proliferating cell nuclear antigen [Mandarin fish ranavirus] | 99.6 | 229 | 4.83E-150 | 217 |
| k141_23750_flag1_multi181.0000_len1968 | W1-2 | 1968 | AYV88159.1 hypothetical protein [Mandarin fish ranavirus] | 100 | 216 | 6.92E-150 | 1318 |
| k141_40919_flag1_multi398.0000_len1957 | W4-3 | 1957 | AYV88159.1 hypothetical protein [Mandarin fish ranavirus] | 100 | 216 | 8.90E-150 | 2813 |
| k141_122964_flag1_multi9.9857_len772 | W6-3 | 772 | AYV88159.1 hypothetical protein [Mandarin fish ranavirus] | 100 | 208 | 2.19E-149 | 28 |
| k141_97747_flag0_multi289.7093_len1135 | W6-1 | 1135 | WEI29033.1 hypothetical protein [Largemouth bass virus] | 100 | 304 | 2.70E-149 | 859 |
| k141_44525_flag1_multi20.0000_len627 | W5-3 | 627 | AYV88168.1 hypothetical protein [Mandarin fish ranavirus] | 99 | 202 | 2.76E-149 | 42 |
| k141_3861_flag1_multi11.0000_len616 | W3-1 | 616 | AYV88168.1 hypothetical protein [Mandarin fish ranavirus] | 99 | 200 | 1.59E-147 | 20 |
| k141_102487_flag0_multi80.2700_len2552 | W5-2 | 2552 | AYV88159.1 hypothetical protein [Mandarin fish ranavirus] | 100 | 216 | 3.26E-147 | 696 |
| k141_144146_flag1_multi8.0000_len949 | W6-3 | 949 | AYV88176.1 putative tumor necrosis factor receptor [Mandarin fish ranavirus] | 100 | 192 | 7.79E-147 | 23 |
| k141_31734_flag1_multi7.9872_len688 | W3-1 | 688 | UUY86269.1 hypothetical protein [Largemouth bass virus] | 98.7 | 229 | 1.40E-145 | 15 |
| k141_114355_flag1_multi107.1101_len1295 | W5-3 | 1295 | QJE49100.1 hypothetical protein LMBV_037 [Largemouth bass virus] | 99.5 | 212 | 1.74E-145 | 475 |
| k141_107414_flag0_multi50.0000_len1230 | W3-2 | 1230 | AYV88168.1 hypothetical protein [Mandarin fish ranavirus] | 99 | 201 | 7.33E-145 | 198 |
| k141_57966_flag1_multi6.0000_len628 | W6-3 | 628 | QJE49149.1 putative immediate early protein ICP-46 [Largemouth bass virus] | 100 | 200 | 8.52E-145 | 11 |
| k141_50210_flag1_multi18.0000_len703 | W1-3 | 703 | AYV88121.1 hypothetical protein [Mandarin fish ranavirus] | 98.6 | 208 | 5.30E-144 | 47 |
| k141_176343_flag1_multi4.0000_len653 | W5-3 | 653 | UVF58793.1 MAG: DNA polymerase [Halichoeres melanurus ranavirus] | 100 | 217 | 6.63E-144 | 6 |

| | | | | | | | |
|---|---|---|---|---|---|---|---|
| k141_23025_flag1_multi5.0000_len668 | W1-3 | 668 | QJE49137.1 hypothetical protein LMBV_074 [Largemouth bass virus] | 98.2 | 222 | 1.81E-143 | 13 |
| k141_5299_flag1_multi33.0000_len615 | W6-2 | 615 | QJE49080.1 hypothetical protein LMBV_017 [Largemouth bass virus] | 98.5 | 196 | 1.07E-142 | 75 |
| k141_123234_flag1_multi10.0000_len607 | W5-2 | 607 | QJE49080.1 hypothetical protein LMBV_017 [Largemouth bass virus] | 98.5 | 196 | 1.40E-142 | 21 |
| k141_230486_flag1_multi6.0000_len592 | W5-3 | 592 | QJE49080.1 hypothetical protein LMBV_017 [Largemouth bass virus] | 98.5 | 195 | 2.30E-142 | 11 |
| k141_219393_flag1_multi12.0000_len589 | W1-3 | 589 | UUY86238.1 hypothetical protein [Largemouth bass virus] | 100 | 196 | 2.40E-142 | 24 |
| k141_120901_flag1_multi19.0000_len610 | W1-2 | 610 | QJE49080.1 hypothetical protein LMBV_017 [Largemouth bass virus] | 98.5 | 195 | 2.94E-142 | 37 |
| k141_78985_flag1_multi61.0000_len702 | W4-2 | 702 | QJE49080.1 hypothetical protein LMBV_017 [Largemouth bass virus] | 98.5 | 196 | 4.90E-142 | 142 |
| k141_15583_flag1_multi6.0000_len624 | W3-1 | 624 | QJE49080.1 hypothetical protein LMBV_017 [Largemouth bass virus] | 98.5 | 196 | 5.11E-142 | 13 |
| k141_21554_flag1_multi37.0000_len611 | W3-2 | 611 | QJE49080.1 hypothetical protein LMBV_017 [Largemouth bass virus] | 98.5 | 195 | 5.92E-142 | 82 |
| k141_193202_flag1_multi26.0000_len645 | W4-3 | 645 | QJE49080.1 hypothetical protein LMBV_017 [Largemouth bass virus] | 98.5 | 196 | 6.79E-142 | 60 |
| k141_174785_flag1_multi771.0000_len4210 | W2-3 | 4210 | AYV88159.1 hypothetical protein [Mandarin fish ranavirus] | 100 | 216 | 9.51E-142 | 12132 |
| k141_215866_flag1_multi12.0000_len586 | W1-3 | 586 | QJE49080.1 hypothetical protein LMBV_017 [Largemouth bass virus] | 98.4 | 193 | 2.47E-141 | 25 |
| k141_102353_flag1_multi7.0000_len582 | W3-2 | 582 | UUY86238.1 hypothetical protein [Largemouth bass virus] | 100 | 193 | 4.06E-140 | 12 |
| k141_34231_flag1_multi7.0000_len588 | W6-3 | 588 | QJE49080.1 hypothetical protein LMBV_017 [Largemouth bass virus] | 98.4 | 191 | 1.18E-139 | 13 |
| k141_225593_flag1_multi10.0000_len623 | W5-3 | 623 | UUY86210.1 putative RNAseIII [Largemouth bass virus] | 100 | 200 | 3.83E-139 | 22 |
| k141_21315_flag1_multi6.8669_len1178 | W3-1 | 1178 | AYV88134.2 putative tyrosine kinase [Mandarin fish ranavirus] | 100 | 201 | 1.99E-138 | 22 |
| k141_33597_flag1_multi15.0000_len621 | W1-3 | 621 | UUY86210.1 putative RNAseIII [Largemouth bass virus] | 99.5 | 200 | 4.27E-138 | 27 |
| k141_60077_flag1_multi32.0000_len3887 | W6-2 | 3887 | AYV88210.2 putative proliferating cell nuclear antigen [Mandarin fish ranavirus] | 99.6 | 224 | 8.95E-138 | 388 |
| k141_112616_flag1_multi453.8178_len849 | W2-1 | 849 | QJE49080.1 hypothetical protein LMBV_017 [Largemouth bass virus] | 97.4 | 189 | 2.53E-134 | 1809 |
| k141_237792_flag1_multi6.0000_len803 | W6-3 | 803 | UUY86261.1 hypothetical protein [Largemouth bass virus] | 100 | 212 | 1.12E-133 | 13 |
| k141_146178_flag1_multi18.0000_len1418 | W5-2 | 1418 | UUY86210.1 putative RNAseIII [Largemouth bass virus] | 99.5 | 200 | 1.36E-133 | 87 |
| k141_98541_flag1_multi6.0000_len647 | W6-3 | 647 | WAK75136.1 hypothetical protein [Mandarin fish ranavirus] | 100 | 189 | 7.22E-133 | 11 |
| k141_44182_flag1_multi5.0000_len1168 | W5-3 | 1168 | QJE49083.1 hypothetical protein LMBV_020 [Largemouth bass virus] | 99.5 | 205 | 1.13E-132 | 21 |
| k141_186552_flag1_multi4.0000_len603 | W1-3 | 603 | AYV88179.2 putative DNA dependent RNA polymerase II second largest subunit [Mandarin fish ranavirus] | 99 | 200 | 2.32E-132 | 7 |
| k141_94270_flag1_multi6.0000_len910 | W1-2 | 910 | WHA35507.1 hypothetical protein MSRaV_19R [Micropterus salmoides ranavirus] | 98.9 | 187 | 1.77E-131 | 18 |
| k141_23824_flag1_multi50.0000_len629 | W1-3 | 629 | AYV88125.2 hypothetical protein [Mandarin fish ranavirus] | 97.9 | 190 | 2.98E-131 | 106 |
| k141_131990_flag1_multi6.0000_len1104 | W1-3 | 1104 | QJE49140.1 putative ribonucleotide reductase alpha subunit [Largemouth bass virus] | 99.5 | 194 | 4.06E-131 | 18 |
| k141_129900_flag1_multi118.0000_len788 | W1-2 | 788 | AYV88125.2 hypothetical protein [Mandarin fish ranavirus] | 97.4 | 191 | 4.24E-131 | 280 |
| k141_15468_flag1_multi40.0000_len857 | W3-1 | 857 | AYV88125.2 hypothetical protein [Mandarin fish ranavirus] | 97.4 | 191 | 2.09E-130 | 100 |
| k141_224937_flag1_multi12.0000_len850 | W6-3 | 850 | WXI69518.1 hypothetical protein [Largemouth bass virus] | 91.8 | 267 | 6.80E-130 | 39 |
| k141_30454_flag1_multi99.6214_len1245 | W3-2 | 1245 | WEI28998.1 hypothetical protein [Largemouth bass virus] | 93 | 284 | 3.65E-129 | 537 |
| k141_30825_flag1_multi54.0000_len1271 | W1-3 | 1271 | WEI28998.1 hypothetical protein [Largemouth bass virus] | 93 | 284 | 4.85E-129 | 289 |
| k141_76549_flag1_multi9.0000_len646 | W6-3 | 646 | XRL22790.1 hypothetical protein [Siniperca chuatsi ranavirus] | 99 | 192 | 5.46E-129 | 19 |
| k141_25742_flag1_multi3.0000_len619 | W6-3 | 619 | UUY86236.1 hypothetical protein [Largemouth bass virus] | 100 | 181 | 8.18E-129 | 4 |
| k141_61635_flag1_multi30.0000_len1379 | W5-3 | 1379 | WEI28998.1 hypothetical protein [Largemouth bass virus] | 93 | 284 | 2.42E-128 | 161 |
| k141_62480_flag1_multi84.0000_len1393 | W1-2 | 1393 | WEI28998.1 hypothetical protein [Largemouth bass virus] | 93 | 284 | 4.07E-128 | 467 |
| k141_105001_flag0_multi345.1931_len1467 | W4-2 | 1467 | WEI28998.1 hypothetical protein [Largemouth bass virus] | 93 | 284 | 4.59E-128 | 2018 |
| k141_154408_flag1_multi59.3321_len1478 | W6-2 | 1478 | WEI28998.1 hypothetical protein [Largemouth bass virus] | 93 | 284 | 5.25E-128 | 667 |
| k141_150661_flag1_multi4.0000_len548 | W5-3 | 548 | AYV88131.1 putative eIF-2 alpha-like protein [Mandarin fish ranavirus] | 99.5 | 182 | 3.68E-127 | 5 |

| | | | | | | | |
|---|---|---|---|---|---|---|---|
| k141_19597_flag1_multi2.0000_len545 | W6-3 | 545 | UUY86263.1 hypothetical protein [Largemouth bass virus] | 100 | 181 | 3.91E-127 | 4 |
| k141_84233_flag1_multi456.0000_len1783 | W6-1 | 1783 | WEI28998.1 hypothetical protein [Largemouth bass virus] | 93 | 284 | 1.40E-126 | 3488 |
| k141_99491_flag0_multi13.0000_len803 | W1-3 | 803 | XRL22821.1 hypothetical protein [Siniperca chuatsi ranavirus] | 100 | 186 | 1.42E-126 | 45 |
| k141_232727_flag1_multi7.0000_len547 | W4-3 | 547 | AYV88131.1 putative eIF-2 alpha-like protein [Mandarin fish ranavirus] | 100 | 181 | 1.43E-126 | 12 |
| k141_116885_flag1_multi30.0000_len588 | W1-3 | 588 | UUY86216.1 hypothetical protein [Largemouth bass virus] | 99.5 | 191 | 4.07E-125 | 65 |
| k141_32673_flag1_multi5.0000_len610 | W3-1 | 610 | QJE49137.1 hypothetical protein LMBV_074 [Largemouth bass virus] | 98.5 | 202 | 1.06E-124 | 9 |
| k141_257294_flag1_multi3.0000_len572 | W6-3 | 572 | UUY86194.1 hypothetical protein [Largemouth bass virus] | 100 | 190 | 1.40E-124 | 4 |
| k141_56393_flag1_multi3.0000_len577 | W6-3 | 577 | UUY86258.1 hypothetical protein [Largemouth bass virus] | 100 | 189 | 1.85E-124 | 4 |
| k141_1780_flag1_multi760.0696_len658 | W3-1 | 658 | AYV88121.1 hypothetical protein [Mandarin fish ranavirus] | 98.4 | 183 | 3.36E-124 | 7 |
| k141_243336_flag1_multi18.0000_len542 | W4-3 | 542 | QJE49123.1 putative DNA repair protein RAD2 [Largemouth bass virus] | 100 | 180 | 3.58E-123 | 34 |
| k141_125706_flag1_multi13.0000_len589 | W5-2 | 589 | AYV88121.1 hypothetical protein [Mandarin fish ranavirus] | 95.2 | 188 | 4.22E-123 | 22 |
| k141_159_flag1_multi2974.6154_len3066 | W4-1 | 3066 | UUY86198.1 hypothetical protein [Largemouth bass virus] | 100 | 213 | 2.41E-122 | 34542 |
| k141_113272_flag1_multi14.0000_len562 | W1-2 | 562 | AYV88179.2 putative DNA dependent RNA polymerase II second largest subunit [Mandarin fish ranavirus] | 98.9 | 186 | 1.47E-121 | 35 |
| k141_43035_flag1_multi5.0000_len546 | W5-3 | 546 | AYV88179.2 putative DNA dependent RNA polymerase II second largest subunit [Mandarin fish ranavirus] | 100 | 181 | 1.71E-120 | 9 |
| k141_86911_flag1_multi28.0698_len1616 | W5-2 | 1616 | WEI28998.1 hypothetical protein [Largemouth bass virus] | 92.8 | 276 | 4.27E-120 | 189 |
| k141_50511_flag1_multi4.0000_len544 | W6-3 | 544 | UUY86267.1 ribonucleotide reductase alpha subunit [Largemouth bass virus] | 100 | 180 | 7.14E-120 | 5 |
| k141_189512_flag1_multi7.0000_len668 | W1-3 | 668 | AYV88133.1 putative DNA-directed RNA polymerase II subunit [Mandarin fish ranavirus] | 100 | 159 | 1.47E-118 | 19 |
| k141_79222_flag1_multi6.0000_len724 | W5-2 | 724 | AYV88133.1 putative DNA-directed RNA polymerase II subunit [Mandarin fish ranavirus] | 100 | 159 | 3.11E-118 | 10 |
| k141_162819_flag0_multi1.0000_len558 | W5-1 | 558 | AIG51690.1 major capsid protein, partial [Koi ranavirus] | 94.6 | 186 | 3.49E-117 | 79.17 |
| k141_134434_flag1_multi5.0000_len525 | W1-3 | 525 | QJE49123.1 putative DNA repair protein RAD2 [Largemouth bass virus] | 100 | 174 | 1.36E-116 | 7 |
| k141_31913_flag1_multi4.0000_len594 | W3-1 | 594 | QJE49083.1 hypothetical protein LMBV_020 [Largemouth bass virus] | 99.4 | 178 | 1.96E-115 | 7 |
| k141_230771_flag1_multi4.0000_len520 | W6-3 | 520 | QJE49145.1 putative proliferating cell nuclear antigen [Largemouth bass virus] | 100 | 173 | 3.06E-115 | 7 |
| k141_13295_flag1_multi7.0000_len501 | W3-1 | 501 | UUY86238.1 hypothetical protein [Largemouth bass virus] | 98.2 | 167 | 3.36E-115 | 9 |
| k141_46866_flag1_multi4.0000_len485 | W3-3 | 485 | QJE49149.1 putative immediate early protein ICP-46 [Largemouth bass virus] | 100 | 161 | 6.22E-115 | 5 |
| k141_53869_flag1_multi5.0000_len584 | W6-3 | 584 | AYV88168.1 hypothetical protein [Mandarin fish ranavirus] | 98.1 | 159 | 7.05E-114 | 10 |
| k141_147870_flag1_multi4.0000_len548 | W5-3 | 548 | QJE49137.1 hypothetical protein LMBV_074 [Largemouth bass virus] | 98.4 | 182 | 2.05E-113 | 8 |
| k141_81444_flag1_multi19.0000_len558 | W6-2 | 558 | AYV88136.2 hypothetical protein [Mandarin fish ranavirus] | 98.7 | 159 | 6.20E-113 | 35 |
| k141_188315_flag1_multi19.0000_len513 | W4-3 | 513 | AYV88136.2 hypothetical protein [Mandarin fish ranavirus] | 98.7 | 159 | 6.54E-113 | 34 |
| k141_9206_flag1_multi23.0301_len1006 | W3-1 | 1006 | WEI28998.1 hypothetical protein [Largemouth bass virus] | 92.3 | 261 | 3.28E-112 | 94 |
| k141_9694_flag1_multi4.0000_len486 | W3-1 | 486 | UVF58787.1 MAG: DNA-dependent RNA polymerase largest subunit, partial [Halichoeres melanurus ranavirus] | 100 | 161 | 3.40E-112 | 5 |
| k141_37889_flag1_multi56.5406_len707 | W4-2 | 707 | AYV88136.2 hypothetical protein [Mandarin fish ranavirus] | 98.7 | 159 | 4.34E-112 | 140.69 |
| k141_146417_flag1_multi8.0000_len520 | W1-2 | 520 | AYV88179.2 putative DNA dependent RNA polymerase II second largest subunit [Mandarin fish ranavirus] | 99.4 | 172 | 1.37E-111 | 12 |
| k141_6351_flag1_multi13.0000_len479 | W1-2 | 479 | AYV88136.2 hypothetical protein [Mandarin fish ranavirus] | 98.7 | 157 | 1.40E-111 | 20 |
| k141_67506_flag1_multi29.4862_len795 | W6-1 | 795 | AYV88136.2 hypothetical protein [Mandarin fish ranavirus] | 98.7 | 159 | 2.68E-111 | 75 |

| | | | | | | | |
|---|---|---|---|---|---|---|---|
| k141_31106_flag1_multi18.0000_len537 | W3-2 | 537 | AYV88136.2 hypothetical protein [Mandarin fish ranavirus] | 98.7 | 151 | 6.38E-111 | 31 |
| k141_36093_flag0_multi14513.5106_len1316 | W1-1 | 1316 | UVF58785.1 MAG: major capsid protein [Halichoeres melanurus ranavirus] | 100 | 166 | 8.23E-111 | 204167 |
| k141_217865_flag1_multi6.8142_len464 | W5-3 | 464 | QYU76035.1 DNA polymerase, partial [Koi ranavirus] | 100 | 154 | 1.11E-110 | 9 |
| k141_26331_flag1_multi36.0000_len967 | W3-1 | 967 | UUY86256.1 putative orf58-like protein [Largemouth bass virus] | 99.1 | 225 | 1.55E-110 | 161 |
| k141_121205_flag1_multi6.0000_len996 | W5-3 | 996 | AYV88136.2 hypothetical protein [Mandarin fish ranavirus] | 98.7 | 159 | 2.37E-110 | 19 |
| k141_100791_flag1_multi5.0000_len496 | W1-3 | 496 | QJE49122.1 putative DNA-dependent RNA polymerase b subunit [Largemouth bass virus] | 100 | 164 | 5.54E-110 | 7 |
| k141_151609_flag1_multi9.0000_len1924 | W5-2 | 1924 | UUY86218.1 hypothetical protein [Largemouth bass virus] | 98.2 | 171 | 1.34E-109 | 46 |
| k141_32021_flag1_multi12.0000_len892 | W1-2 | 892 | XRB52829.1 P8.141C-like protein, partial [Largemouth bass virus] | 98.8 | 160 | 1.59E-109 | 33 |
| k141_66652_flag0_multi358.0000_len2814 | W2-3 | 2814 | UUY86204.1 hypothetical protein [Largemouth bass virus] | 100 | 193 | 9.61E-109 | 1413 |
| k141_14272_flag1_multi4.0000_len481 | W3-1 | 481 | UUY86211.1 hypothetical protein [Largemouth bass virus] | 98.7 | 159 | 3.09E-108 | 6 |
| k141_79644_flag0_multi1662.1574_len2263 | W1-1 | 2263 | UUY86218.1 hypothetical protein [Largemouth bass virus] | 98.8 | 170 | 5.41E-108 | 8476 |
| k141_4619_flag0_multi372.9712_len905 | W6-2 | 905 | UVF58785.1 MAG: major capsid protein [Halichoeres melanurus ranavirus] | 100 | 159 | 1.20E-107 | 872 |
| k141_27804_flag1_multi61.1749_len1010 | W5-3 | 1010 | UUY86256.1 putative orf58-like protein [Largemouth bass virus] | 92.1 | 242 | 1.43E-107 | 260 |
| k141_116449_flag1_multi6.0000_len853 | W5-2 | 853 | QJE49140.1 putative ribonucleotide reductase alpha subunit [Largemouth bass virus] | 100 | 160 | 1.56E-107 | 15 |
| k141_115774_flag1_multi7.0000_len652 | W1-3 | 652 | UUY86221.1 hypothetical protein [Largemouth bass virus] | 98.9 | 177 | 2.90E-107 | 15 |
| k141_61173_flag1_multi90.0000_len1678 | W6-2 | 1678 | WXI69513.1 hypothetical protein [Largemouth bass virus] | 97.9 | 187 | 3.12E-107 | 588 |
| k141_91943_flag1_multi4.0000_len463 | W3-2 | 463 | AYV88130.2 putative ATPase-dependent protease [Mandarin fish ranavirus] | 98.7 | 153 | 7.85E-107 | 6 |
| k141_23771_flag1_multi7.0000_len468 | W5-3 | 468 | AYV88130.2 putative ATPase-dependent protease [Mandarin fish ranavirus] | 98.7 | 153 | 8.53E-107 | 9 |
| k141_64226_flag1_multi190.2107_len1793 | W1-1 | 1793 | AYV88136.2 hypothetical protein [Mandarin fish ranavirus] | 98.7 | 159 | 9.44E-107 | 3238.19 |
| k141_148965_flag0_multi44.2305_len1399 | W1-3 | 1399 | UUY86256.1 putative orf58-like protein [Largemouth bass virus] | 97.8 | 223 | 1.06E-106 | 231 |
| k141_11586_flag1_multi2.0000_len479 | W3-1 | 479 | AYV88173.2 putative ribonucleotide reductase beta subunit [Mandarin fish ranavirus] | 100 | 159 | 2.12E-106 | 3 |
| k141_2498_flag0_multi60.0922_len1595 | W3-2 | 1595 | UUY86256.1 putative orf58-like protein [Largemouth bass virus] | 98.2 | 225 | 3.04E-106 | 411 |
| k141_22544_flag0_multi248.3778_len1131 | W4-3 | 1131 | UUY86256.1 putative orf58-like protein [Largemouth bass virus] | 100 | 160 | 4.81E-106 | 1835.85 |
| k141_114775_flag1_multi36.6247_len1244 | W5-2 | 1244 | UUY86256.1 putative orf58-like protein [Largemouth bass virus] | 100 | 160 | 1.78E-105 | 222 |
| k141_4674_flag1_multi3.0000_len459 | W3-1 | 459 | AYV88121.1 hypothetical protein [Mandarin fish ranavirus] | 98.7 | 151 | 1.79E-105 | 3 |
| k141_202261_flag1_multi3.0000_len503 | W3-3 | 503 | UVF58785.1 MAG: major capsid protein [Halichoeres melanurus ranavirus] | 100 | 154 | 3.11E-105 | 6 |
| k141_13925_flag1_multi5.0000_len463 | W3-1 | 463 | AYV88131.1 putative eIF-2 alpha-like protein [Mandarin fish ranavirus] | 100 | 153 | 3.83E-105 | 6 |
| k141_125040_flag1_multi15.0000_len511 | W6-3 | 511 | UUY86216.1 hypothetical protein [Largemouth bass virus] | 99.4 | 165 | 7.31E-105 | 25 |
| k141_20534_flag1_multi9.0000_len1076 | W3-1 | 1076 | UUY86221.1 hypothetical protein [Largemouth bass virus] | 98.9 | 177 | 8.17E-105 | 29 |
| k141_9376_flag1_multi13.2474_len521 | W3-1 | 521 | AYV88126.1 putative immediate early protein ICP-18 [Mandarin fish ranavirus] | 99.3 | 147 | 1.19E-104 | 17 |
| k141_175103_flag1_multi400.2270_len1709 | W2-3 | 1709 | UUY86256.1 putative orf58-like protein [Largemouth bass virus] | 92.1 | 242 | 3.05E-104 | 5642.75 |
| k141_31432_flag1_multi6243.2500_len1741 | W1-1 | 1741 | UUY86256.1 putative orf58-like protein [Largemouth bass virus] | 92.1 | 242 | 4.09E-104 | 62061.35 |
| k141_127675_flag1_multi7.0000_len593 | W3-2 | 593 | UUY86218.1 hypothetical protein [Largemouth bass virus] | 98.7 | 155 | 7.66E-104 | 11 |
| k141_59753_flag1_multi20495.0000_len1350 | W4-1 | 1350 | UVF58785.1 MAG: major capsid protein [Halichoeres melanurus ranavirus] | 100 | 156 | 1.38E-103 | 171223.32 |
| k141_29370_flag1_multi31.0000_len798 | W3-2 | 798 | AYV88126.1 putative immediate early protein ICP-18 [Mandarin fish ranavirus] | 99.3 | 147 | 3.17E-103 | 81 |
| k141_246427_flag1_multi7.0000_len561 | W1-3 | 561 | UUY86269.1 hypothetical protein [Largemouth bass virus] | 99.4 | 164 | 4.50E-103 | 12 |
| k141_268124_flag1_multi17.9250_len861 | W5-3 | 861 | AYV88126.1 putative immediate early protein ICP-18 [Mandarin fish ranavirus] | 99.3 | 147 | 9.51E-103 | 41 |
| k141_157199_flag1_multi5.0000_len509 | W5-2 | 509 | UVF58793.1 MAG: DNA polymerase [Halichoeres melanurus ranavirus] | 100 | 162 | 1.02E-102 | 8 |
| k141_7178_flag0_multi80.6557_len1233 | W5-3 | 1233 | WEI29006.1 putative 2-cysteine adaptor domain protein [Largemouth bass virus] | 100 | 152 | 1.66E-102 | 345 |

| | | | | | | | |
|---|---|---|---|---|---|---|---|
| k141_21216_flag1_multi9.0000_len1908 | W3-2 | 1908 | UUY86221.1 hypothetical protein [Largemouth bass virus] | 98.9 | 177 | 2.92E-102 | 53 |
| k141_44197_flag1_multi7.0000_len879 | W5-3 | 879 | UUY86218.1 hypothetical protein [Largemouth bass virus] | 98.7 | 155 | 2.92E-102 | 16 |
| k141_225011_flag1_multi13.0000_len446 | W6-3 | 446 | AYV88120.1 putative p31K protein [Mandarin fish ranavirus] | 98.6 | 148 | 1.47E-101 | 19 |
| k141_135881_flag1_multi10759.0000_len2218 | W5-1 | 2218 | UUY86256.1 putative orf58-like protein [Largemouth bass virus] | 100 | 160 | 2.39E-101 | 103510.49 |
| k141_2743_flag1_multi5.0000_len480 | W3-1 | 480 | QJE49137.1 hypothetical protein LMBV_074 [Largemouth bass virus] | 98.1 | 159 | 4.28E-101 | 6 |
| k141_118642_flag1_multi4.0000_len432 | W6-2 | 432 | XRB52806.1 hypothetical protein LMBV_49 [Largemouth bass virus] | 99.3 | 141 | 4.28E-101 | 5 |
| k141_7287_flag1_multi16.0000_len1219 | W3-1 | 1219 | UUY86218.1 hypothetical protein [Largemouth bass virus] | 98.7 | 155 | 5.65E-101 | 52 |
| k141_88665_flag1_multi16.0000_len792 | W1-3 | 792 | AYV88126.1 putative immediate early protein ICP-18 [Mandarin fish ranavirus] | 99.3 | 145 | 8.01E-101 | 40 |
| k141_40633_flag0_multi3.3469_len1000 | W5-1 | 1000 | UUY86192.1 putative myristylated membrane protein [Largemouth bass virus] | 92.5 | 159 | 3.69E-100 | 8 |
| k141_125032_flag1_multi17.6904_len1349 | W6-2 | 1349 | UUY86218.1 hypothetical protein [Largemouth bass virus] | 98.7 | 155 | 6.60E-100 | 72 |
| k141_233065_flag1_multi368.0000_len3611 | W4-3 | 3611 | XRB52829.1 P8.141C-like protein, partial [Largemouth bass virus] | 98.8 | 160 | 7.34E-100 | 4187 |
| k141_17999_flag1_multi5.0000_len501 | W6-1 | 501 | UUY86233.1 hypothetical protein [Largemouth bass virus] | 95.9 | 148 | 4.61E-99 | 5 |
| k141_11670_flag1_multi2.0000_len535 | W3-1 | 535 | UVF58793.1 MAG: DNA polymerase [Halichoeres melanurus ranavirus] | 100 | 155 | 9.02E-99 | 4 |
| k141_7534_flag1_multi7.9699_len673 | W3-2 | 673 | QJE49085.1 hypothetical protein LMBV_022 [Largemouth bass virus] | 97.8 | 139 | 9.71E-99 | 11 |
| k141_33388_flag1_multi2.0000_len425 | W3-3 | 425 | AYV88159.1 hypothetical protein [Mandarin fish ranavirus] | 100 | 141 | 3.07E-98 | 2 |
| k141_18940_flag1_multi10.0000_len709 | W6-3 | 709 | UUY86256.1 putative orf58-like protein [Largemouth bass virus] | 100 | 147 | 3.88E-98 | 22 |
| k141_80021_flag1_multi2793.0000_len3928 | W4-1 | 3928 | UUY86256.1 putative orf58-like protein [Largemouth bass virus] | 100 | 160 | 5.90E-97 | 37904 |
| k141_24529_flag1_multi5.0000_len573 | W1-2 | 573 | QJE49128.1 hypothetical protein LMBV_065 [Largemouth bass virus] | 98.5 | 133 | 1.00E-96 | 10 |
| k141_52856_flag1_multi8.0000_len685 | W4-3 | 685 | QJE49128.1 hypothetical protein LMBV_065 [Largemouth bass virus] | 98.5 | 133 | 4.24E-96 | 20 |
| k141_205111_flag1_multi2.0000_len454 | W3-3 | 454 | AYV88211.1 hypothetical protein [Mandarin fish ranavirus] | 100 | 150 | 6.82E-95 | 3 |
| k141_109364_flag0_multi359.6471_len549 | W2-1 | 549 | AYV88126.1 putative immediate early protein ICP-18 [Mandarin fish ranavirus] | 99.3 | 136 | 6.88E-95 | 1806 |
| k141_110039_flag1_multi3.0000_len414 | W6-3 | 414 | QJE49097.1 hypothetical protein LMBV_034 [Largemouth bass virus] | 100 | 137 | 7.58E-95 | 2 |
| k141_7711_flag1_multi6.0000_len406 | W3-1 | 406 | QJE49109.1 hypothetical protein LMBV_046 [Largemouth bass virus] | 100 | 135 | 1.18E-94 | 7 |
| k141_80951_flag1_multi3.0000_len446 | W6-3 | 446 | WHA35533.1 putative DNA dependent RNA polymerase A subunit [Micropterus salmoides ranavirus] | 100 | 148 | 2.68E-94 | 4 |
| k141_11533_flag1_multi129.4239_len785 | W2-1 | 785 | AYV88185.1 hypothetical protein [Mandarin fish ranavirus] | 97 | 132 | 2.84E-93 | 356 |
| k141_47257_flag1_multi2.0000_len386 | W5-3 | 386 | QJE49128.1 hypothetical protein LMBV_065 [Largemouth bass virus] | 98.4 | 128 | 5.91E-93 | 2 |
| k141_11025_flag0_multi1142.8511_len732 | W2-1 | 732 | QJE49125.1 hypothetical protein LMBV_062 [Largemouth bass virus] | 98.1 | 214 | 6.89E-93 | 8211 |
| k141_43345_flag1_multi10.0000_len381 | W6-3 | 381 | QJE49149.1 putative immediate early protein ICP-46 [Largemouth bass virus] | 100 | 126 | 4.72E-92 | 11 |
| k141_225160_flag1_multi26.8158_len874 | W5-3 | 874 | QJE49119.1 hypothetical protein LMBV_056 [Largemouth bass virus] | 100 | 140 | 1.17E-91 | 78 |
| k141_179303_flag1_multi15.0000_len766 | W1-3 | 766 | QJE49119.1 hypothetical protein LMBV_056 [Largemouth bass virus] | 100 | 139 | 1.22E-91 | 38 |
| k141_14737_flag1_multi22.0000_len788 | W3-1 | 788 | QJE49119.1 hypothetical protein LMBV_056 [Largemouth bass virus] | 100 | 139 | 1.64E-91 | 48 |
| k141_104345_flag1_multi3.0000_len427 | W5-2 | 427 | AYV88179.2 putative DNA dependent RNA polymerase II second largest subunit [Mandarin fish ranavirus] | 100 | 142 | 1.89E-91 | 2 |
| k141_134652_flag3_multi10546.0000_len536 | W1-1 | 536 | AYV88191.1 hypothetical protein [Mandarin fish ranavirus] | 99.2 | 130 | 2.41E-91 | 52361.96 |
| k141_43035_flag1_multi1076.0280_len999 | W2-1 | 999 | QJE49119.1 hypothetical protein LMBV_056 [Largemouth bass virus] | 99.3 | 141 | 3.53E-91 | 3736 |
| k141_21166_flag1_multi26.0000_len763 | W5-2 | 763 | QJE49119.1 hypothetical protein LMBV_056 [Largemouth bass virus] | 100 | 138 | 4.78E-91 | 65 |
| k141_37090_flag1_multi7.0000_len429 | W1-3 | 429 | AYV88179.2 putative DNA dependent RNA polymerase II second largest subunit [Mandarin fish ranavirus] | 100 | 142 | 2.64E-90 | 8 |
| k141_144087_flag1_multi62.0000_len968 | W4-3 | 968 | QJE49119.1 hypothetical protein LMBV_056 [Largemouth bass virus] | 100 | 138 | 4.08E-90 | 220 |

| | | | | | | | |
|---|---|---|---|---|---|---|---|
| k141_119432_flag1_multi163.0000_len1405 | W2-3 | 1405 | QJE49119.1 hypothetical protein LMBV_056 [Largemouth bass virus] | 100 | 141 | 7.36E-90 | 727 |
| k141_19261_flag1_multi3.0000_len423 | W3-1 | 423 | AYV88179.2 putative DNA dependent RNA polymerase II second largest subunit [Mandarin fish ranavirus] | 100 | 140 | 1.26E-89 | 3 |
| k141_21163_flag0_multi441.9775_len585 | W2-2 | 585 | QJE49073.1 putative p31K protein [Largemouth bass virus] | 100 | 138 | 2.46E-89 | 1309 |
| k141_188002_flag1_multi3.0000_len482 | W5-3 | 482 | UUY86269.1 hypothetical protein [Largemouth bass virus] | 99.3 | 147 | 3.12E-89 | 4 |
| k141_188904_flag1_multi3.8041_len432 | W5-3 | 432 | XPZ21295.1 putative RNAseIII [Mandarin fish ranavirus] | 99.3 | 143 | 4.98E-89 | 5 |
| k141_90995_flag1_multi13.4669_len700 | W6-1 | 700 | AYV88208.1 hypothetical protein [Mandarin fish ranavirus] | 100 | 129 | 1.84E-88 | 30 |
| k141_80618_flag1_multi234.9130_len1348 | W6-1 | 1348 | QJE49119.1 hypothetical protein LMBV_056 [Largemouth bass virus] | 100 | 139 | 2.58E-88 | 6294.41 |
| k141_256006_flag1_multi3.0000_len425 | W6-3 | 425 | AYV88211.1 hypothetical protein [Mandarin fish ranavirus] | 100 | 141 | 4.91E-88 | 4 |
| k141_61389_flag1_multi12.0000_len2326 | W6-1 | 2326 | AYV88185.1 hypothetical protein [Mandarin fish ranavirus] | 97.7 | 132 | 9.93E-88 | 94 |
| k141_76455_flag0_multi410.9509_len548 | W6-1 | 548 | AYV88120.1 putative p31K protein [Mandarin fish ranavirus] | 98.5 | 137 | 1.99E-87 | 1059 |
| k141_240804_flag1_multi4.0000_len421 | W6-3 | 421 | UUY86258.1 hypothetical protein [Largemouth bass virus] | 99.3 | 140 | 3.70E-87 | 4 |
| k141_104768_flag0_multi585.0000_len966 | W6-1 | 966 | QJE49215.1 putative orf58-like protein [Largemouth bass virus] | 100 | 132 | 7.26E-87 | 2014 |
| k141_112996_flag1_multi9.0000_len420 | W1-3 | 420 | UVF58793.1 MAG: DNA polymerase [Halichoeres melanurus ranavirus] | 100 | 140 | 1.24E-86 | 11 |
| k141_273023_flag1_multi4.0000_len619 | W5-3 | 619 | AYV88204.1 putative NIF/NLI interacting factor [Mandarin fish ranavirus] | 99.3 | 134 | 1.29E-86 | 7 |
| k141_2470_flag1_multi25.0000_len779 | W3-1 | 779 | XRL22828.1 DNA-directed RNA polymerase subunit beta [Siniperca chuatsi ranavirus] | 98.7 | 152 | 1.62E-86 | 62 |
| k141_224120_flag1_multi5.0000_len433 | W5-3 | 433 | WHA35533.1 putative DNA dependent RNA polymerase A subunit [Micropterus salmoides ranavirus] | 100 | 144 | 3.12E-86 | 5 |
| k141_42247_flag1_multi5.0000_len433 | W5-3 | 433 | UUY86269.1 hypothetical protein [Largemouth bass virus] | 98.6 | 144 | 1.32E-85 | 6 |
| k141_211606_flag1_multi4.0000_len583 | W6-3 | 583 | XRB52842.1 transcription factor TFIIB cyclin-like domain-containing protein, partial [Largemouth bass virus] | 100 | 125 | 2.23E-85 | 6 |
| k141_87762_flag1_multi239.3994_len3008 | W1-1 | 3008 | QJE49119.1 hypothetical protein LMBV_056 [Largemouth bass virus] | 100 | 141 | 2.34E-84 | 5543 |
| k141_98614_flag1_multi4.0000_len353 | W1-3 | 353 | ADB77862.1 DNA methyltransferase [Largemouth bass ulcerative syndrome virus] | 100 | 117 | 3.83E-84 | 3 |
| k141_229916_flag1_multi4.0000_len449 | W6-3 | 449 | AYV88155.1 hypothetical protein [Mandarin fish ranavirus] | 97.7 | 129 | 1.39E-83 | 5 |
| k141_24074_flag1_multi4.0000_len463 | W5-2 | 463 | AYV88179.2 putative DNA dependent RNA polymerase II second largest subunit [Mandarin fish ranavirus] | 100 | 135 | 1.54E-83 | 4 |
| k141_75357_flag1_multi4.0000_len536 | W5-2 | 536 | QJE49128.1 hypothetical protein LMBV_065 [Largemouth bass virus] | 98.3 | 118 | 2.20E-83 | 8 |
| k141_7478_flag1_multi5.0000_len376 | W5-2 | 376 | AYV88136.2 hypothetical protein [Mandarin fish ranavirus] | 98.4 | 123 | 3.25E-83 | 5 |
| k141_142161_flag1_multi6.0000_len369 | W1-3 | 369 | AYV88173.2 putative ribonucleotide reductase beta subunit [Mandarin fish ranavirus] | 100 | 123 | 5.84E-83 | 9 |
| k141_25586_flag0_multi90.2459_len1231 | W6-1 | 1231 | UUY86204.1 hypothetical protein [Largemouth bass virus] | 100 | 149 | 3.05E-82 | 827 |
| k141_34464_flag1_multi4.0000_len410 | W3-1 | 410 | ABA41591.1 DNA-dependent DNA polymerase, partial [Largemouth bass virus] | 98.5 | 136 | 1.28E-81 | 4 |
| k141_120855_flag1_multi1.0000_len663 | W5-1 | 663 | UUY86258.1 hypothetical protein [Largemouth bass virus] | 91.9 | 149 | 1.06E-80 | 1 |
| k141_111147_flag1_multi5.0000_len352 | W4-2 | 352 | AYV88208.1 hypothetical protein [Mandarin fish ranavirus] | 100 | 117 | 2.14E-80 | 3 |
| k141_39172_flag1_multi3.0000_len375 | W6-3 | 375 | XPZ21261.1 hypothetical protein MRVORF002 [Mandarin fish ranavirus] | 99.2 | 125 | 1.59E-79 | 3 |
| k141_145314_flag1_multi1.0000_len366 | W5-1 | 366 | UVF58785.1 MAG: major capsid protein [Halichoeres melanurus ranavirus] | 95.9 | 122 | 3.48E-79 | 0 |
| k141_34524_flag1_multi20.0000_len1146 | W3-1 | 1146 | UUY86234.1 hypothetical protein [Largemouth bass virus] | 100 | 123 | 9.12E-79 | 78 |
| k141_45500_flag1_multi1.0000_len629 | W2-1 | 629 | UUY86256.1 putative orf58-like protein [Largemouth bass virus] | 84.4 | 147 | 1.02E-78 | 2340.16 |
| k141_153608_flag1_multi53.0000_len1171 | W6-2 | 1171 | UUY86234.1 hypothetical protein [Largemouth bass virus] | 100 | 123 | 1.21E-78 | 224 |
| k141_245910_flag1_multi16.0000_len790 | W1-3 | 790 | WAK75073.1 hypothetical protein [Mandarin fish ranavirus] | 100 | 119 | 1.33E-78 | 33 |

| | | | | | | | |
|---|---|---|---|---|---|---|---|
| k141_84366_flag1_multi17.1461_len1236 | W5-3 | 1236 | UUY86234.1 hypothetical protein [Largemouth bass virus] | 100 | 123 | 2.33E-78 | 76 |
| k141_32018_flag1_multi1.0000_len428 | W5-1 | 428 | QYU76034.1 putative myristylated membrane protein, partial [Koi ranavirus] | 95.8 | 142 | 3.65E-78 | 0 |
| k141_48751_flag1_multi10.0000_len348 | W6-3 | 348 | QJE49109.1 hypothetical protein LMBV_046 [Largemouth bass virus] | 100 | 115 | 2.05E-77 | 12 |
| k141_7920_flag0_multi7384.8387_len1468 | W2-1 | 1468 | UUY86234.1 hypothetical protein [Largemouth bass virus] | 100 | 123 | 2.21E-77 | 7696 |
| k141_267941_flag1_multi4.0000_len387 | W5-3 | 387 | UUY86269.1 hypothetical protein [Largemouth bass virus] | 99.2 | 129 | 2.21E-77 | 4 |
| k141_102429_flag1_multi33.8577_len1097 | W5-2 | 1097 | WAK75073.1 hypothetical protein [Mandarin fish ranavirus] | 100 | 119 | 3.12E-77 | 132 |
| k141_91875_flag1_multi153.1910_len1073 | W6-2 | 1073 | UUY86251.1 hypothetical protein [Largemouth bass virus] | 99.2 | 120 | 1.10E-76 | 718 |
| k141_73340_flag1_multi32.0000_len1335 | W1-2 | 1335 | WAK75073.1 hypothetical protein [Mandarin fish ranavirus] | 100 | 119 | 3.51E-76 | 207 |
| k141_100_flag1_multi2.0000_len353 | W6-3 | 353 | UUY86238.1 hypothetical protein [Largemouth bass virus] | 99.1 | 117 | 6.11E-76 | 0 |
| k141_78704_flag0_multi25.0028_len1194 | W5-3 | 1194 | WAK75073.1 hypothetical protein [Mandarin fish ranavirus] | 99.2 | 118 | 9.75E-76 | 92 |
| k141_128726_flag1_multi1.0000_len340 | W1-1 | 340 | WEI29006.1 putative 2-cysteine adaptor domain protein [Largemouth bass virus] | 99.1 | 112 | 1.61E-75 | 0 |
| k141_160409_flag1_multi3.0000_len422 | W5-3 | 422 | ADB77862.1 DNA methyltransferase [Largemouth bass ulcerative syndrome virus] | 100 | 105 | 3.28E-75 | 2 |
| k141_75522_flag1_multi1.0000_len359 | W4-1 | 359 | AYV88120.1 putative p31K protein [Mandarin fish ranavirus] | 95.8 | 119 | 3.68E-75 | 0 |
| k141_165032_flag1_multi3.0000_len362 | W1-3 | 362 | AYV88179.2 putative DNA dependent RNA polymerase II second largest subunit [Mandarin fish ranavirus] | 100 | 120 | 4.04E-75 | 4 |
| k141_15852_flag1_multi364.0703_len1692 | W6-1 | 1692 | WAK75073.1 hypothetical protein [Mandarin fish ranavirus] | 100 | 119 | 8.64E-75 | 1609 |
| k141_9408_flag0_multi20.0000_len515 | W3-2 | 515 | UUY86257.1 hypothetical protein [Largemouth bass virus] | 92.7 | 123 | 1.22E-74 | 50 |
| k141_33957_flag1_multi28.0000_len1709 | W3-1 | 1709 | WAK75073.1 hypothetical protein [Mandarin fish ranavirus] | 99.2 | 119 | 3.95E-74 | 148 |
| k141_48689_flag0_multi322.0000_len498 | W4-3 | 498 | UUY86257.1 hypothetical protein [Largemouth bass virus] | 92.6 | 122 | 5.56E-74 | 146.45 |
| k141_86554_flag1_multi6.0000_len446 | W3-2 | 446 | AYV88183.1 hypothetical protein [Mandarin fish ranavirus] | 99.1 | 113 | 7.64E-74 | 7 |
| k141_87806_flag0_multi302.2527_len513 | W2-1 | 513 | AYV88121.1 hypothetical protein [Mandarin fish ranavirus] | 98.3 | 117 | 1.09E-73 | 682 |
| k141_55135_flag1_multi9437.0000_len1687 | W4-1 | 1687 | UVF58785.1 MAG: major capsid protein [Halichoeres melanurus ranavirus] | 100 | 127 | 1.45E-73 | 88052.16 |
| k141_16508_flag1_multi3.0000_len353 | W3-2 | 353 | AYV88179.2 putative DNA dependent RNA polymerase II second largest subunit [Mandarin fish ranavirus] | 100 | 117 | 1.70E-73 | 2 |
| k141_41165_flag1_multi3.0000_len359 | W5-3 | 359 | WHA35533.1 putative DNA dependent RNA polymerase A subunit [Micropterus salmoides ranavirus] | 100 | 119 | 3.86E-73 | 3 |
| k141_121765_flag1_multi30.5923_len872 | W5-2 | 872 | UUY86257.1 hypothetical protein [Largemouth bass virus] | 92.7 | 123 | 9.74E-73 | 218 |
| k141_18640_flag1_multi2.0000_len331 | W6-3 | 331 | UUY86267.1 ribonucleotide reductase alpha subunit [Largemouth bass virus] | 100 | 109 | 2.22E-72 | 1 |
| k141_32584_flag1_multi88.0000_len2644 | W4-2 | 2644 | WAK75073.1 hypothetical protein [Mandarin fish ranavirus] | 100 | 119 | 3.57E-72 | 739 |
| k141_30403_flag0_multi15.0000_len446 | W5-2 | 446 | UUY86221.1 hypothetical protein [Largemouth bass virus] | 100 | 122 | 6.99E-72 | 9 |
| k141_214089_flag1_multi7.0000_len316 | W1-3 | 316 | ACP19247.1 DNA polymerase, partial [Doctor fish virus] | 100 | 104 | 7.80E-72 | 8 |
| k141_123470_flag1_multi31.3813_len1038 | W1-2 | 1038 | QJE49119.1 hypothetical protein LMBV_056 [Largemouth bass virus] | 100 | 114 | 9.76E-72 | 113 |
| k141_131987_flag1_multi1.0000_len533 | W1-1 | 533 | AIG51690.1 major capsid protein, partial [Koi ranavirus] | 93.1 | 131 | 2.62E-71 | 3 |
| k141_101207_flag1_multi1.0000_len373 | W5-1 | 373 | AYV88120.1 putative p31K protein [Mandarin fish ranavirus] | 94.3 | 123 | 4.02E-71 | 0 |
| k141_19304_flag1_multi5.0000_len382 | W3-1 | 382 | QJE49137.1 hypothetical protein LMBV_074 [Largemouth bass virus] | 99.2 | 127 | 1.58E-70 | 5 |
| k141_139620_flag1_multi42.6660_len1147 | W6-2 | 1147 | WAK75073.1 hypothetical protein [Mandarin fish ranavirus] | 100 | 110 | 3.32E-70 | 155 |
| k141_30533_flag1_multi5.4376_len630 | W1-3 | 630 | UUY86251.1 hypothetical protein [Largemouth bass virus] | 100 | 106 | 4.00E-70 | 10 |
| k141_34768_flag1_multi24.0000_len369 | W5-3 | 369 | UUY86216.1 hypothetical protein [Largemouth bass virus] | 99.1 | 111 | 5.72E-70 | 57 |
| k141_99928_flag1_multi8.0000_len333 | W3-3 | 333 | XRB52768.1 P31K protein, partial [Largemouth bass virus] | 98.2 | 110 | 7.39E-70 | 9 |
| k141_9753_flag1_multi4.0000_len446 | W6-3 | 446 | UUY86269.1 hypothetical protein [Largemouth bass virus] | 100 | 108 | 1.23E-69 | 4 |

| | | | | | | | |
|---|---|---|---|---|---|---|---|
| k141_18034_flag1_multi6.0000_len341 | W3-1 | 341 | UVF58793.1 MAG: DNA polymerase [Halichoeres melanurus ranavirus] | 100 | 113 | 3.13E-69 | 5 |
| k141_128539_flag1_multi49.0000_len350 | W1-2 | 350 | QJE49138.1 hypothetical protein LMBV_075 [Largemouth bass virus] | 98.1 | 106 | 6.93E-69 | 60 |
| k141_70112_flag1_multi3.0000_len361 | W3-2 | 361 | UUY86207.1 hypothetical protein [Largemouth bass virus] | 96.3 | 107 | 1.02E-68 | 2 |
| k141_133485_flag1_multi5.0000_len591 | W6-2 | 591 | XPZ21295.1 putative RNAseIII [Mandarin fish ranavirus] | 99.2 | 120 | 1.84E-68 | 8 |
| k141_201648_flag1_multi3.0000_len328 | W6-3 | 328 | QJE49095.1 hypothetical protein LMBV_032 [Largemouth bass virus] | 100 | 109 | 3.87E-68 | 1 |
| k141_144322_flag1_multi1.0000_len506 | W5-1 | 506 | AYV88159.1 hypothetical protein [Mandarin fish ranavirus] | 87.2 | 125 | 8.59E-68 | 2331.15 |
| k141_56646_flag1_multi13.0000_len619 | W5-2 | 619 | UUY86251.1 hypothetical protein [Largemouth bass virus] | 100 | 103 | 2.77E-67 | 22 |
| k141_51626_flag1_multi6.0000_len349 | W3-2 | 349 | XPZ21295.1 putative RNAseIII [Mandarin fish ranavirus] | 98.3 | 116 | 5.46E-67 | 6 |
| k141_108857_flag1_multi13.0000_len796 | W5-3 | 796 | QJE49138.1 hypothetical protein LMBV_075 [Largemouth bass virus] | 98.1 | 106 | 6.79E-67 | 40 |
| k141_104129_flag1_multi12.9310_len663 | W1-3 | 663 | QJE49138.1 hypothetical protein LMBV_075 [Largemouth bass virus] | 98.1 | 105 | 1.12E-66 | 33 |
| k141_23560_flag1_multi10.0000_len338 | W3-1 | 338 | QJE49138.1 hypothetical protein LMBV_075 [Largemouth bass virus] | 98.1 | 104 | 1.14E-66 | 10 |
| k141_36962_flag0_multi1.0000_len494 | W5-1 | 494 | AIG51690.1 major capsid protein, partial [Koi ranavirus] | 94.1 | 118 | 1.48E-66 | 29.74 |
| k141_74_flag1_multi1.0000_len322 | W5-1 | 322 | QIZ30887.1 major capsid protein, partial [Largemouth bass virus] | 92.5 | 107 | 1.72E-66 | 0 |
| k141_150034_flag0_multi1.0000_len433 | W5-1 | 433 | AIG51690.1 major capsid protein, partial [Koi ranavirus] | 96.4 | 111 | 1.90E-66 | 7.72 |
| k141_106064_flag1_multi5.0000_len346 | W5-2 | 346 | QJE49097.1 hypothetical protein LMBV_034 [Largemouth bass virus] | 99.1 | 114 | 2.01E-66 | 3 |
| k141_82359_flag1_multi1.0000_len312 | W5-1 | 312 | UVF58785.1 MAG: major capsid protein [Halichoeres melanurus ranavirus] | 97.1 | 104 | 2.62E-66 | 0 |
| k141_166270_flag1_multi7.0000_len324 | W1-3 | 324 | AYV88134.2 putative tyrosine kinase [Mandarin fish ranavirus] | 100 | 107 | 5.85E-66 | 10 |
| k141_17386_flag0_multi2.8553_len445 | W5-3 | 445 | UUY86198.1 hypothetical protein [Largemouth bass virus] | 100 | 103 | 6.50E-66 | 4 |
| k141_168731_flag1_multi1.0000_len360 | W5-1 | 360 | QJE49137.1 hypothetical protein LMBV_074 [Largemouth bass virus] | 95 | 119 | 1.37E-65 | 0 |
| k141_7626_flag1_multi3.0000_len328 | W6-3 | 328 | UUY86258.1 hypothetical protein [Largemouth bass virus] | 100 | 108 | 1.62E-65 | 3 |
| k141_42104_flag1_multi9.1679_len671 | W5-2 | 671 | QJE49138.1 hypothetical protein LMBV_075 [Largemouth bass virus] | 99 | 104 | 2.05E-65 | 26 |
| k141_150090_flag1_multi1.0000_len328 | W5-1 | 328 | XRL22790.1 hypothetical protein [Siniperca chuatsi ranavirus] | 96.3 | 108 | 2.83E-65 | 1 |
| k141_181039_flag1_multi5.0000_len340 | W1-3 | 340 | WEI28968.1 putative DNA dependent RNA polymerase a subunit [Largemouth bass virus] | 100 | 112 | 3.12E-65 | 5 |
| k141_20214_flag1_multi1.0000_len349 | W5-1 | 349 | QYU76034.1 putative myristylated membrane protein, partial [Koi ranavirus] | 91.3 | 115 | 4.24E-65 | 0 |
| k141_40012_flag1_multi1.0000_len310 | W1-1 | 310 | UVF58785.1 MAG: major capsid protein [Halichoeres melanurus ranavirus] | 98.1 | 103 | 5.57E-65 | 0 |
| k141_105146_flag1_multi1.0000_len357 | W5-1 | 357 | UUY86256.1 putative orf58-like protein [Largemouth bass virus] | 93.8 | 112 | 7.53E-65 | 130.93 |
| k141_57538_flag1_multi5.0000_len321 | W5-3 | 321 | QJE49137.1 hypothetical protein LMBV_074 [Largemouth bass virus] | 100 | 106 | 1.36E-64 | 1 |
| k141_174145_flag1_multi3.0000_len311 | W3-3 | 311 | AYV88149.2 hypothetical protein [Mandarin fish ranavirus] | 98 | 100 | 3.10E-64 | 2 |
| k141_10703_flag0_multi26.6576_len544 | W1-1 | 544 | AYV88191.1 hypothetical protein [Mandarin fish ranavirus] | 93.4 | 106 | 3.48E-64 | 16366.04 |
| k141_31211_flag1_multi3.0000_len393 | W3-1 | 393 | QJE49085.1 hypothetical protein LMBV_022 [Largemouth bass virus] | 98.9 | 91 | 9.12E-64 | 3 |
| k141_199634_flag1_multi3.0000_len316 | W1-3 | 316 | UVF58793.1 MAG: DNA polymerase [Halichoeres melanurus ranavirus] | 100 | 105 | 9.56E-64 | 2 |
| k141_5600_flag1_multi7.0000_len302 | W3-1 | 302 | QJE49123.1 putative DNA repair protein RAD2 [Largemouth bass virus] | 100 | 100 | 1.59E-63 | 7 |
| k141_155744_flag1_multi1.0000_len451 | W5-1 | 451 | AYV88191.1 hypothetical protein [Mandarin fish ranavirus] | 86.4 | 118 | 1.73E-63 | 1 |
| k141_38143_flag1_multi3.0000_len348 | W1-2 | 348 | AYV88208.1 hypothetical protein [Mandarin fish ranavirus] | 100 | 93 | 2.46E-63 | 2 |
| k141_95083_flag1_multi1.0000_len332 | W4-1 | 332 | AYV88120.1 putative p31K protein [Mandarin fish ranavirus] | 91.8 | 110 | 7.11E-63 | 1 |
| k141_44813_flag0_multi1.0000_len528 | W4-1 | 528 | UZV46047.1 major capsid protein, partial [Largemouth bass virus] | 89.6 | 115 | 8.24E-63 | 29079 |
| k141_2544_flag1_multi5.0000_len305 | W3-3 | 305 | AIG51690.1 major capsid protein, partial [Koi ranavirus] | 100 | 101 | 1.08E-62 | 4 |
| k141_18935_flag1_multi1.0000_len337 | W5-1 | 337 | UUY86261.1 hypothetical protein [Largemouth bass virus] | 98.2 | 112 | 4.91E-62 | 0 |
| k141_38492_flag0_multi1.0000_len282 | W5-1 | 282 | AYV88159.1 hypothetical protein [Mandarin fish ranavirus] | 98.9 | 93 | 1.65E-61 | 899.06 |

| | | | | | | | |
|---|---|---|---|---|---|---|---|
| k141_82181_flag1_multi1.0000_len324 | W5-1 | 324 | UUY86241.1 hypothetical protein [Largemouth bass virus] | 96.5 | 86 | 2.11E-61 | 1 |
| k141_68259_flag1_multi1.0000_len371 | W5-1 | 371 | AIG51690.1 major capsid protein, partial [Koi ranavirus] | 94.5 | 110 | 2.13E-61 | 1 |
| k141_120880_flag1_multi1.0000_len327 | W5-1 | 327 | QJE49215.1 putative orf58-like protein [Largemouth bass virus] | 96.9 | 96 | 2.86E-61 | 0 |
| k141_26109_flag1_multi4.0000_len309 | W3-1 | 309 | WHA35533.1 putative DNA dependent RNA polymerase A subunit [Micropterus salmoides ranavirus] | 100 | 102 | 3.22E-61 | 3 |
| k141_201663_flag1_multi16.0000_len591 | W5-3 | 591 | UUY86251.1 hypothetical protein [Largemouth bass virus] | 100 | 94 | 3.37E-61 | 24 |
| k141_122015_flag1_multi1.0000_len295 | W5-1 | 295 | UVF58785.1 MAG: major capsid protein [Halichoeres melanurus ranavirus] | 96.9 | 98 | 1.37E-60 | 0 |
| k141_70526_flag1_multi1.0000_len281 | W1-1 | 281 | AYV88159.1 hypothetical protein [Mandarin fish ranavirus] | 100 | 93 | 2.72E-60 | 0 |
| k141_43473_flag0_multi1.0000_len282 | W5-1 | 282 | AYV88159.1 hypothetical protein [Mandarin fish ranavirus] | 98.9 | 93 | 5.48E-60 | 149.14 |
| k141_50251_flag0_multi1.0000_len312 | W5-1 | 312 | AIG51690.1 major capsid protein, partial [Koi ranavirus] | 95.1 | 103 | 8.46E-60 | 121.23 |
| k141_41688_flag1_multi1.0000_len341 | W5-1 | 341 | UUY86258.1 hypothetical protein [Largemouth bass virus] | 94.7 | 113 | 1.10E-59 | 3 |
| k141_134553_flag1_multi1.0000_len286 | W5-1 | 286 | UVF58785.1 MAG: major capsid protein [Halichoeres melanurus ranavirus] | 97.9 | 94 | 1.33E-59 | 0 |
| k141_45372_flag0_multi1.0000_len329 | W1-1 | 329 | XRL22790.1 hypothetical protein [Siniperca chuatsi ranavirus] | 91.7 | 108 | 1.65E-59 | 3995.08 |
| k141_74891_flag1_multi1.0000_len342 | W5-1 | 342 | QJE49153.1 hypothetical protein LMBV_004 [Largemouth bass virus] | 96.1 | 103 | 2.03E-59 | 0 |
| k141_201037_flag1_multi2.0000_len305 | W6-3 | 305 | AYV88134.2 putative tyrosine kinase [Mandarin fish ranavirus] | 100 | 101 | 4.45E-59 | 1 |
| k141_126897_flag0_multi1.7538_len535 | W5-2 | 535 | XPZ21295.1 putative RNAseIII [Mandarin fish ranavirus] | 82.4 | 136 | 7.48E-59 | 1 |
| k141_48031_flag1_multi1.0000_len586 | W4-1 | 586 | XRL22790.1 hypothetical protein [Siniperca chuatsi ranavirus] | 93.4 | 106 | 8.25E-59 | 0 |
| k141_86763_flag1_multi30.2031_len1165 | W4-1 | 1165 | AYV88120.1 putative p31K protein [Mandarin fish ranavirus] | 94.5 | 110 | 1.55E-58 | 133.26 |
| k141_61328_flag1_multi5.5287_len385 | W1-3 | 385 | WAK75094.1 hypothetical protein [Mandarin fish ranavirus] | 100 | 82 | 1.64E-58 | 5 |
| k141_42162_flag1_multi49.6296_len465 | W5-2 | 465 | WAK75112.1 putative LITAF PIG7 possible membrane associated motif in LPS-induced tumor necrosis factor alpha factor [Mandarin fish ranavirus] | 100 | 85 | 2.69E-58 | 69 |
| k141_47692_flag0_multi332.0000_len297 | W6-1 | 297 | UUY86197.1 hypothetical protein [Largemouth bass virus] | 100 | 99 | 2.82E-58 | 120 |
| k141_199098_flag1_multi3.0000_len356 | W4-3 | 356 | QJE49085.1 hypothetical protein LMBV_022 [Largemouth bass virus] | 97.6 | 84 | 3.20E-58 | 3 |
| k141_20649_flag1_multi57.0000_len386 | W3-1 | 386 | WAK75112.1 putative LITAF PIG7 possible membrane associated motif in LPS-induced tumor necrosis factor alpha factor [Mandarin fish ranavirus] | 98.8 | 85 | 3.94E-58 | 72 |
| k141_103636_flag1_multi2.0000_len505 | W5-2 | 505 | AYV88134.2 putative tyrosine kinase [Mandarin fish ranavirus] | 100 | 94 | 4.79E-58 | 2 |
| k141_100332_flag1_multi1.0000_len281 | W5-1 | 281 | QJE49066.1 hypothetical protein LMBV_003 [Largemouth bass virus] | 100 | 93 | 5.33E-58 | 0 |
| k141_26627_flag1_multi1.0000_len293 | W4-1 | 293 | AYV88159.1 hypothetical protein [Mandarin fish ranavirus] | 96.9 | 97 | 8.67E-58 | 0 |
| k141_141614_flag1_multi1.0000_len360 | W5-1 | 360 | QJE49132.1 hypothetical protein LMBV_069 [Largemouth bass virus] | 90.8 | 119 | 1.06E-57 | 1 |
| k141_81646_flag1_multi3.0593_len731 | W4-1 | 731 | XRL22821.1 hypothetical protein [Siniperca chuatsi ranavirus] | 87.9 | 116 | 1.53E-57 | 6 |
| k141_39687_flag1_multi1.0000_len309 | W4-1 | 309 | AIG51690.1 major capsid protein, partial [Koi ranavirus] | 96.1 | 102 | 2.06E-57 | 0 |
| k141_109815_flag1_multi1.0000_len326 | W2-1 | 326 | UUY86227.1 hypothetical protein [Largemouth bass virus] | 82.4 | 108 | 5.62E-57 | 0 |
| k141_50065_flag0_multi1.0000_len282 | W5-1 | 282 | UVF58785.1 MAG: major capsid protein [Halichoeres melanurus ranavirus] | 98.9 | 93 | 5.87E-57 | 958.11 |
| k141_89647_flag1_multi224.5107_len795 | W4-2 | 795 | WAK75112.1 putative LITAF PIG7 possible membrane associated motif in LPS-induced tumor necrosis factor alpha factor [Mandarin fish ranavirus] | 100 | 85 | 6.96E-57 | 543 |
| k141_65082_flag1_multi4.0000_len311 | W6-3 | 311 | UUY86269.1 hypothetical protein [Largemouth bass virus] | 99 | 103 | 7.72E-57 | 4 |
| k141_139920_flag1_multi1.0000_len574 | W5-1 | 574 | XRB52768.1 P31K protein, partial [Largemouth bass virus] | 92.1 | 101 | 1.15E-56 | 2 |
| k141_75887_flag1_multi3.0000_len345 | W1-1 | 345 | UVF58785.1 MAG: major capsid protein [Halichoeres melanurus ranavirus] | 82.5 | 114 | 3.91E-56 | 0 |
| k141_118680_flag1_multi1.0000_len455 | W1-1 | 455 | AIG51690.1 major capsid protein, partial [Koi ranavirus] | 75.4 | 142 | 1.17E-55 | 2 |
| k141_108844_flag0_multi1.0000_len278 | W1-1 | 278 | AIG51690.1 major capsid protein, partial [Koi ranavirus] | 98.9 | 92 | 1.21E-55 | 24726.77 |

| | | | | | | | |
|---|---|---|---|---|---|---|---|
| k141_11082_flag0_multi1.0000_len304 | W1-1 | 304 | AIG51690.1 major capsid protein, partial [Koi ranavirus] | 96 | 101 | 1.76E-55 | 5992.98 |
| k141_62093_flag1_multi1.0000_len281 | W5-1 | 281 | AIG51690.1 major capsid protein, partial [Koi ranavirus] | 98.9 | 93 | 2.53E-55 | 2.98 |
| k141_21637_flag1_multi4.8115_len629 | W3-1 | 629 | UUY86257.1 hypothetical protein [Largemouth bass virus] | 91.8 | 97 | 3.50E-55 | 7 |
| k141_81600_flag0_multi1.0000_len289 | W4-1 | 289 | AIG51690.1 major capsid protein, partial [Koi ranavirus] | 97.9 | 96 | 4.04E-55 | 6407.23 |
| k141_85502_flag1_multi1.0000_len402 | W5-1 | 402 | QJE49215.1 putative orf58-like protein [Largemouth bass virus] | 92.5 | 93 | 6.59E-55 | 2.1 |
| k141_131786_flag1_multi2.0000_len649 | W2-2 | 649 | UUY86227.1 hypothetical protein [Largemouth bass virus] | 83.5 | 109 | 9.86E-55 | 4 |
| k141_108962_flag1_multi4.2099_len970 | W1-1 | 970 | WEI28972.1 CTD-phosphotransferase [Largemouth bass virus] | 76.6 | 124 | 1.11E-54 | 11.1 |
| k141_13338_flag1_multi3.5259_len681 | W5-1 | 681 | XRL22821.1 hypothetical protein [Siniperca chuatsi ranavirus] | 94.9 | 99 | 1.26E-54 | 9.09 |
| k141_120741_flag1_multi4.9222_len1272 | W6-3 | 1272 | AYV88191.1 hypothetical protein [Mandarin fish ranavirus] | 98.9 | 89 | 1.71E-54 | 30 |
| k141_78661_flag1_multi1.0000_len361 | W4-1 | 361 | AYV88176.1 putative tumor necrosis factor receptor [Mandarin fish ranavirus] | 98.8 | 80 | 2.18E-54 | 0 |
| k141_9129_flag1_multi1.0000_len318 | W1-1 | 318 | QYU76034.1 putative myristylated membrane protein, partial [Koi ranavirus] | 100 | 89 | 4.60E-54 | 0 |
| k141_243691_flag1_multi2.0000_len317 | W6-3 | 317 | UUY86234.1 hypothetical protein [Largemouth bass virus] | 100 | 86 | 4.71E-54 | 2 |
| k141_87414_flag1_multi1.0000_len418 | W5-1 | 418 | XRL22790.1 hypothetical protein [Siniperca chuatsi ranavirus] | 92.3 | 104 | 5.27E-54 | 0 |
| k141_249205_flag1_multi4.0000_len302 | W6-3 | 302 | AYV88211.1 hypothetical protein [Mandarin fish ranavirus] | 100 | 90 | 4.10E-53 | 4 |
| k141_42308_flag1_multi2.6056_len785 | W5-1 | 785 | AYV88159.1 hypothetical protein [Mandarin fish ranavirus] | 96.6 | 88 | 2.09E-52 | 32.02 |
| k141_45566_flag0_multi1.0000_len458 | W2-1 | 458 | UUY86263.1 hypothetical protein [Largemouth bass virus] | 97.8 | 90 | 2.30E-52 | 0 |
| k141_74968_flag1_multi1.0000_len281 | W1-1 | 281 | AIG51690.1 major capsid protein, partial [Koi ranavirus] | 98.9 | 93 | 2.52E-52 | 0 |
| k141_124057_flag1_multi1.0000_len281 | W5-1 | 281 | XRL22790.1 hypothetical protein [Siniperca chuatsi ranavirus] | 97.8 | 93 | 3.20E-52 | 0 |
| k141_59372_flag1_multi28.0000_len319 | W4-3 | 319 | WXI69503.1 hypothetical protein [Largemouth bass virus] | 95.7 | 92 | 4.77E-52 | 30 |
| k141_94414_flag0_multi1.0000_len282 | W1-1 | 282 | AIG51690.1 major capsid protein, partial [Koi ranavirus] | 98.9 | 94 | 5.23E-52 | 1538.56 |
| k141_19751_flag1_multi1.0000_len304 | W5-1 | 304 | XRL22790.1 hypothetical protein [Siniperca chuatsi ranavirus] | 94.7 | 94 | 6.06E-52 | 0 |
| k141_141310_flag1_multi1.0000_len320 | W5-1 | 320 | QJE49066.1 hypothetical protein LMBV_003 [Largemouth bass virus] | 87.7 | 106 | 6.19E-52 | 2 |
| k141_73662_flag1_multi1.0000_len604 | W4-1 | 604 | WAK75073.1 hypothetical protein [Mandarin fish ranavirus] | 91.5 | 94 | 6.42E-52 | 27 |
| k141_98280_flag1_multi34.9753_len627 | W5-1 | 627 | UUY86227.1 hypothetical protein [Largemouth bass virus] | 94.1 | 85 | 8.01E-52 | 320 |
| k141_159970_flag1_multi1.0000_len281 | W5-1 | 281 | AYV88149.2 hypothetical protein [Mandarin fish ranavirus] | 97.6 | 84 | 8.85E-52 | 0.82 |
| k141_82689_flag1_multi1.0000_len477 | W2-1 | 477 | AYV88133.1 putative DNA-directed RNA polymerase II subunit [Mandarin fish ranavirus] | 82.7 | 98 | 1.39E-51 | 2 |
| k141_96382_flag1_multi4.0000_len317 | W1-3 | 317 | WEI28969.1 hypothetical protein [Largemouth bass virus] | 100 | 83 | 2.33E-51 | 3 |
| k141_104300_flag1_multi1.0000_len281 | W4-1 | 281 | QJE49132.1 hypothetical protein LMBV_069 [Largemouth bass virus] | 98.9 | 93 | 2.44E-51 | 0 |
| k141_19968_flag1_multi2.6058_len826 | W5-1 | 826 | QJE49149.1 putative immediate early protein ICP-46 [Largemouth bass virus] | 95.5 | 88 | 3.98E-51 | 7 |
| k141_121029_flag0_multi1.0000_len311 | W5-1 | 311 | XRB52831.1 hypothetical protein LMBV_74 [Largemouth bass virus] | 96.1 | 102 | 8.40E-51 | 239.86 |
| k141_51064_flag1_multi1.0000_len590 | W2-1 | 590 | WAK75073.1 hypothetical protein [Mandarin fish ranavirus] | 95.5 | 88 | 9.12E-51 | 6 |
| k141_135603_flag1_multi1.0000_len373 | W5-1 | 373 | UUY86235.1 hypothetical protein [Largemouth bass virus] | 93.3 | 89 | 1.24E-50 | 0 |
| k141_63726_flag1_multi1.0000_len380 | W5-1 | 380 | WHU98721.1 MCP, partial [Hybrid snakehead ranavirus] | 95.5 | 88 | 2.84E-50 | 0 |
| k141_132639_flag1_multi1.0000_len281 | W5-1 | 281 | QJE49105.1 hypothetical protein LMBV_042 [Largemouth bass virus] | 98.8 | 86 | 3.48E-50 | 0 |
| k141_56714_flag1_multi1.0000_len322 | W1-1 | 322 | AYV88120.1 putative p31K protein [Mandarin fish ranavirus] | 80.4 | 107 | 4.68E-50 | 0 |
| k141_162859_flag1_multi1.0000_len373 | W5-1 | 373 | QJE49066.1 hypothetical protein LMBV_003 [Largemouth bass virus] | 79 | 124 | 5.60E-50 | 0 |
| k141_3762_flag0_multi1.0000_len364 | W4-1 | 364 | AIG51690.1 major capsid protein, partial [Koi ranavirus] | 94.4 | 89 | 9.81E-50 | 11616.05 |
| k141_23684_flag1_multi4.0000_len715 | W5-1 | 715 | AYV88211.1 hypothetical protein [Mandarin fish ranavirus] | 92.9 | 98 | 1.06E-49 | 11 |
| k141_57398_flag1_multi1.0000_len324 | W5-1 | 324 | QJE49077.1 hypothetical protein LMBV_014 [Largemouth bass virus] | 93.6 | 78 | 1.95E-49 | 0 |

| | | | | | | | |
|---|---|---|---|---|---|---|---|
| k141_102859_flag1_multi1.0000_len281 | W4-1 | 281 | QJE49132.1 hypothetical protein LMBV_069 [Largemouth bass virus] | 97.8 | 93 | 2.88E-49 | 0 |
| k141_146377_flag1_multi5.8708_len1031 | W5-1 | 1031 | AYV88191.1 hypothetical protein [Mandarin fish ranavirus] | 93.3 | 89 | 5.41E-49 | 27 |
| k141_10492_flag1_multi126.7056_len389 | W6-1 | 389 | AYV88191.1 hypothetical protein [Mandarin fish ranavirus] | 100 | 78 | 8.88E-49 | 161 |
| k141_26280_flag0_multi385.0289_len971 | W6-1 | 971 | XRB52836.1 SAP domain-containing protein [Largemouth bass virus] | 70.9 | 213 | 1.35E-48 | 1519 |
| k141_84172_flag1_multi1.0000_len318 | W1-1 | 318 | QJE49101.1 hypothetical protein LMBV_038 [Largemouth bass virus] | 91.4 | 93 | 2.22E-48 | 0 |
| k141_44802_flag0_multi3.1024_len688 | W4-1 | 688 | AIG51690.1 major capsid protein, partial [Koi ranavirus] | 97.6 | 83 | 2.71E-48 | 3438.12 |
| k141_32218_flag1_multi3.0000_len337 | W3-2 | 337 | AYV88198.1 hypothetical protein [Mandarin fish ranavirus] | 98.9 | 87 | 6.60E-48 | 2 |
| k141_120202_flag1_multi5.0000_len310 | W3-3 | 310 | QJE49215.1 putative orf58-like protein [Largemouth bass virus] | 100 | 72 | 4.70E-47 | 3 |
| k141_9048_flag1_multi1.7041_len577 | W3-1 | 577 | AYV88167.1 hypothetical protein [Mandarin fish ranavirus] | 98.6 | 73 | 9.91E-47 | 56 |
| k141_77612_flag0_multi3.7700_len428 | W6-2 | 428 | AYV88167.1 hypothetical protein [Mandarin fish ranavirus] | 100 | 71 | 1.84E-46 | 5 |
| k141_130366_flag1_multi1.7172_len629 | W5-1 | 629 | XRB52797.1 Bcl-2 Bcl-2 homology region 1-3 domain-containing protein [Largemouth bass virus] | 89.1 | 92 | 3.65E-46 | 3 |
| k141_117052_flag0_multi2058.8519_len357 | W1-1 | 357 | AIG51690.1 major capsid protein, partial [Koi ranavirus] | 100 | 76 | 1.32E-45 | 1391.23 |
| k141_54552_flag0_multi1.0000_len371 | W1-1 | 371 | QJE49080.1 hypothetical protein LMBV_017 [Largemouth bass virus] | 84 | 94 | 1.33E-45 | 1 |
| k141_27638_flag1_multi4.5422_len792 | W4-1 | 792 | XRL22790.1 hypothetical protein [Siniperca chuatsi ranavirus] | 95.5 | 88 | 1.36E-45 | 14 |
| k141_25458_flag1_multi73.2836_len3096 | W4-2 | 3096 | AYV88125.2 hypothetical protein [Mandarin fish ranavirus] | 94 | 84 | 1.63E-45 | 337 |
| k141_61600_flag1_multi17.5367_len985 | W1-1 | 985 | AIG51690.1 major capsid protein, partial [Koi ranavirus] | 97.6 | 85 | 1.78E-45 | 11048.04 |
| k141_153895_flag1_multi89.0000_len425 | W6-2 | 425 | WAK75112.1 putative LITAF PIG7 possible membrane associated motif in LPS-induced tumor necrosis factor alpha factor [Mandarin fish ranavirus] | 98.6 | 70 | 2.92E-45 | 83 |
| k141_94323_flag0_multi1.0000_len386 | W4-1 | 386 | AIG51690.1 major capsid protein, partial [Koi ranavirus] | 98.8 | 82 | 5.47E-45 | 3 |
| k141_99952_flag1_multi5.0000_len394 | W1-3 | 394 | QJE49132.1 hypothetical protein LMBV_069 [Largemouth bass virus] | 100 | 130 | 6.62E-45 | 6 |
| k141_73746_flag1_multi1.0000_len636 | W5-1 | 636 | UUY86258.1 hypothetical protein [Largemouth bass virus] | 90.3 | 93 | 1.23E-44 | 2 |
| k141_25181_flag1_multi8.5014_len2315 | W5-1 | 2315 | AYV88191.1 hypothetical protein [Mandarin fish ranavirus] | 97.5 | 80 | 2.12E-44 | 71.22 |
| k141_24428_flag1_multi1.0000_len528 | W1-1 | 528 | AIG51690.1 major capsid protein, partial [Koi ranavirus] | 91 | 89 | 3.24E-44 | 65982.9 |
| k141_101459_flag1_multi20.6667_len357 | W4-3 | 357 | AYV88167.1 hypothetical protein [Mandarin fish ranavirus] | 100 | 66 | 1.19E-43 | 30 |
| k141_91087_flag1_multi6.0000_len608 | W1-2 | 608 | AYV88179.2 putative DNA dependent RNA polymerase II second largest subunit [Mandarin fish ranavirus] | 100 | 81 | 1.22E-43 | 12 |
| k141_99444_flag1_multi1.0000_len306 | W5-1 | 306 | QJE49066.1 hypothetical protein LMBV_003 [Largemouth bass virus] | 98.7 | 79 | 1.59E-43 | 1 |
| k141_92601_flag0_multi1612.9560_len1686 | W1-1 | 1686 | UVF58785.1 MAG: major capsid protein [Halichoeres melanurus ranavirus] | 100 | 86 | 4.60E-43 | 30212.01 |
| k141_116726_flag1_multi1.0000_len379 | W5-1 | 379 | QJE49132.1 hypothetical protein LMBV_069 [Largemouth bass virus] | 81.1 | 111 | 6.32E-43 | 0 |
| k141_19373_flag1_multi1.0000_len599 | W1-1 | 599 | QJE49101.1 hypothetical protein LMBV_038 [Largemouth bass virus] | 94.3 | 87 | 9.61E-43 | 1 |
| k141_87091_flag1_multi1.0000_len406 | W4-1 | 406 | QYU76034.1 putative myristylated membrane protein, partial [Koi ranavirus] | 91 | 89 | 1.37E-42 | 13 |
| k141_38297_flag1_multi1.0000_len353 | W4-2 | 353 | QJE49077.1 hypothetical protein LMBV_014 [Largemouth bass virus] | 98.5 | 67 | 1.38E-42 | 36 |
| k141_49952_flag1_multi9.9514_len4399 | W1-1 | 4399 | AYV88120.1 putative p31K protein [Mandarin fish ranavirus] | 96.4 | 83 | 5.43E-42 | 174 |
| k141_116467_flag0_multi7.5291_len468 | W5-3 | 468 | WAK75112.1 putative LITAF PIG7 possible membrane associated motif in LPS-induced tumor necrosis factor alpha factor [Mandarin fish ranavirus] | 98.5 | 66 | 5.46E-42 | 28 |
| k141_67968_flag0_multi1.0000_len787 | W5-1 | 787 | QJE49137.1 hypothetical protein LMBV_074 [Largemouth bass virus] | 96.4 | 84 | 3.85E-41 | 20 |
| k141_54883_flag0_multi590.5915_len939 | W4-1 | 939 | UVF58785.1 MAG: major capsid protein [Halichoeres melanurus ranavirus] | 96.4 | 84 | 1.24E-40 | 3192 |
| k141_13741_flag1_multi1.0000_len411 | W5-1 | 411 | XRB52842.1 transcription factor TFIIB cyclin-like domain-containing protein, partial [Largemouth bass virus] | 94.2 | 69 | 2.67E-40 | 0 |

| | | | | | | | |
|---|---|---|---|---|---|---|---|
| k141_104554_flag0_multi1.0000_len603 | W5-1 | 603 | AYV88120.1 putative p31K protein [Mandarin fish ranavirus] | 85.9 | 85 | 4.33E-40 | 11654.04 |
| k141_13975_flag1_multi1.0000_len358 | W5-1 | 358 | WEI28972.1 CTD-phosphotransferase [Largemouth bass virus] | 94.8 | 77 | 4.61E-40 | 0 |
| k141_30727_flag1_multi4.0000_len398 | W6-3 | 398 | WAK75097.1 hypothetical protein [Mandarin fish ranavirus] | 100 | 68 | 8.47E-39 | 3 |
| k141_31522_flag1_multi1.0000_len519 | W2-1 | 519 | AYV88159.1 hypothetical protein [Mandarin fish ranavirus] | 82.4 | 85 | 1.22E-38 | 5 |
| k141_156459_flag0_multi1.0000_len352 | W5-1 | 352 | AYV88191.1 hypothetical protein [Mandarin fish ranavirus] | 97 | 67 | 1.65E-38 | 31.23 |
| k141_97207_flag1_multi7.0000_len885 | W5-1 | 885 | WEI28998.1 hypothetical protein [Largemouth bass virus] | 92.3 | 78 | 2.01E-38 | 19 |
| k141_58582_flag1_multi1.0000_len465 | W5-1 | 465 | AFD96401.1 major capsid protein, partial [Largemouth bass virus] | 88.5 | 78 | 3.13E-38 | 81.8 |
| k141_162810_flag0_multi3.7326_len399 | W5-1 | 399 | AFD96401.1 major capsid protein, partial [Largemouth bass virus] | 94.3 | 70 | 1.16E-37 | 7.33 |
| k141_53982_flag0_multi59.9830_len729 | W1-1 | 729 | AIG51690.1 major capsid protein, partial [Koi ranavirus] | 100 | 70 | 2.57E-37 | 6774.18 |
| k141_126628_flag1_multi3.8399_len572 | W1-1 | 572 | WEI28972.1 CTD-phosphotransferase [Largemouth bass virus] | 98.6 | 71 | 3.50E-37 | 4 |
| k141_97923_flag1_multi5.6947_len1189 | W2-1 | 1189 | WEI28972.1 CTD-phosphotransferase [Largemouth bass virus] | 84.3 | 89 | 4.55E-37 | 25 |
| k141_6833_flag1_multi13.0000_len377 | W3-1 | 377 | WAK75121.1 hypothetical protein [Mandarin fish ranavirus] | 96.8 | 63 | 5.79E-37 | 12 |
| k141_8180_flag1_multi33.0998_len1423 | W5-1 | 1423 | QYU76034.1 putative myristylated membrane protein, partial [Koi ranavirus] | 98.6 | 73 | 5.85E-37 | 352.02 |
| k141_167168_flag0_multi1.0000_len430 | W5-1 | 430 | QIZ30887.1 major capsid protein, partial [Largemouth bass virus] | 93.9 | 66 | 2.45E-36 | 9398.67 |
| k141_16189_flag1_multi7.0000_len350 | W6-2 | 350 | QJE49203.1 hypothetical protein LMBV_054 [Largemouth bass virus] | 96.3 | 82 | 2.59E-36 | 7 |
| k141_33623_flag1_multi1.0000_len318 | W5-1 | 318 | AYV88159.1 hypothetical protein [Mandarin fish ranavirus] | 93.9 | 66 | 3.97E-36 | 0 |
| k141_101063_flag1_multi1.9922_len655 | W5-1 | 655 | QJE49137.1 hypothetical protein LMBV_074 [Largemouth bass virus] | 98.6 | 71 | 5.15E-36 | 4 |
| k141_160288_flag1_multi1.0000_len462 | W5-1 | 462 | AYV88120.1 putative p31K protein [Mandarin fish ranavirus] | 97 | 66 | 9.95E-36 | 23 |
| k141_75471_flag1_multi7.8802_len1485 | W4-1 | 1485 | AYV88121.1 hypothetical protein [Mandarin fish ranavirus] | 94.6 | 74 | 1.00E-35 | 37 |
| k141_128897_flag1_multi2.0000_len669 | W5-1 | 669 | QJE49077.1 hypothetical protein LMBV_014 [Largemouth bass virus] | 95.2 | 63 | 1.16E-35 | 7 |
| k141_105112_flag0_multi395.0079_len394 | W4-1 | 394 | QJE49203.1 hypothetical protein LMBV_054 [Largemouth bass virus] | 98.7 | 78 | 2.63E-35 | 402 |
| k141_85762_flag1_multi1.0000_len344 | W4-1 | 344 | AIG51690.1 major capsid protein, partial [Koi ranavirus] | 84.9 | 86 | 3.81E-35 | 0 |
| k141_3370_flag0_multi7.8845_len1474 | W5-1 | 1474 | AYP19467.1 hypothetical protein [Frog virus 3] | 70.4 | 108 | 4.45E-35 | 23 |
| k141_124085_flag1_multi1.0000_len336 | W5-1 | 336 | QJE49215.1 putative orf58-like protein [Largemouth bass virus] | 82.1 | 78 | 6.74E-35 | 2 |
| k141_150904_flag1_multi23.0000_len354 | W1-3 | 354 | WAK75072.1 hypothetical protein [Mandarin fish ranavirus] | 100 | 62 | 8.32E-35 | 26 |
| k141_53238_flag0_multi23.2126_len602 | W4-1 | 602 | WEI28972.1 CTD-phosphotransferase [Largemouth bass virus] | 97.2 | 71 | 9.87E-35 | 33 |
| k141_43468_flag0_multi1.0000_len330 | W5-1 | 330 | UUY86235.1 hypothetical protein [Largemouth bass virus] | 96.7 | 61 | 1.06E-34 | 1 |
| k141_81031_flag1_multi14.0000_len889 | W4-2 | 889 | WHA35556.1 putative LPXTG-anchored collagen-like adhesin Scl2/SclB [Micropterus salmoides ranavirus] | 100 | 208 | 1.15E-34 | 37 |
| k141_159434_flag0_multi1.0000_len426 | W5-1 | 426 | AIG51690.1 major capsid protein, partial [Koi ranavirus] | 85.7 | 77 | 2.18E-34 | 16.01 |
| k141_73703_flag1_multi18.0000_len2083 | W5-1 | 2083 | UUY86263.1 hypothetical protein [Largemouth bass virus] | 95.8 | 72 | 3.43E-34 | 131 |
| k141_38104_flag1_multi1.0000_len329 | W4-1 | 329 | UUY86267.1 ribonucleotide reductase alpha subunit [Largemouth bass virus] | 80 | 80 | 5.30E-34 | 0 |
| k141_106679_flag0_multi1.0000_len360 | W2-1 | 360 | WEI28972.1 CTD-phosphotransferase [Largemouth bass virus] | 95.3 | 64 | 1.52E-33 | 35.7 |
| k141_176071_flag0_multi2.3686_len670 | W2-3 | 670 | UUY86235.1 hypothetical protein [Largemouth bass virus] | 95.5 | 66 | 2.89E-33 | 6 |
| k141_164191_flag0_multi51.0000_len4401 | W5-1 | 4401 | AYV88211.1 hypothetical protein [Mandarin fish ranavirus] | 93.9 | 82 | 2.95E-33 | 750 |
| k141_56464_flag1_multi12.0000_len721 | W4-3 | 721 | WHA35556.1 putative LPXTG-anchored collagen-like adhesin Scl2/SclB [Micropterus salmoides ranavirus] | 100 | 205 | 3.80E-33 | 30 |
| k141_74344_flag1_multi19.0000_len2132 | W1-1 | 2132 | AYV88159.1 hypothetical protein [Mandarin fish ranavirus] | 91.8 | 73 | 4.02E-33 | 141 |
| k141_95987_flag1_multi14.0000_len1458 | W2-2 | 1458 | WHA35556.1 putative LPXTG-anchored collagen-like adhesin Scl2/SclB [Micropterus salmoides ranavirus] | 100 | 208 | 6.08E-33 | 67 |

| | | | | | | | |
|---|---|---|---|---|---|---|---|
| k141_145631_flag1_multi1.0000_len368 | W5-1 | 368 | AYV88191.1 hypothetical protein [Mandarin fish ranavirus] | 96.4 | 55 | 6.09E-33 | 1 |
| k141_5749_flag1_multi1.0000_len436 | W5-1 | 436 | AYV88191.1 hypothetical protein [Mandarin fish ranavirus] | 90.8 | 65 | 6.85E-33 | 2 |
| k141_41202_flag0_multi1.0000_len434 | W5-1 | 434 | AIG51690.1 major capsid protein, partial [Koi ranavirus] | 92.6 | 68 | 6.93E-33 | 0 |
| k141_41639_flag1_multi1.0000_len319 | W1-1 | 319 | AYV88191.1 hypothetical protein [Mandarin fish ranavirus] | 96.6 | 59 | 9.02E-33 | 3466.7 |
| k141_64096_flag1_multi33.0000_len1597 | W6-1 | 1597 | WHA35556.1 putative LPXTG-anchored collagen-like adhesin Scl2/SclB [Micropterus salmoides ranavirus] | 100 | 208 | 1.15E-32 | 118 |
| k141_141198_flag1_multi1.0000_len302 | W5-1 | 302 | QJE49067.1 hypothetical protein LMBV_004 [Largemouth bass virus] | 81.8 | 77 | 1.22E-32 | 0 |
| k141_91204_flag0_multi1.0000_len321 | W1-1 | 321 | UUY86266.1 hypothetical protein [Largemouth bass virus] | 96.7 | 61 | 1.52E-32 | 3.03 |
| k141_156332_flag1_multi12.6856_len1019 | W5-1 | 1019 | AYV88191.1 hypothetical protein [Mandarin fish ranavirus] | 95.2 | 62 | 4.62E-32 | 59.09 |
| k141_112637_flag1_multi3.8150_len514 | W2-1 | 514 | UUY86235.1 hypothetical protein [Largemouth bass virus] | 64.1 | 103 | 4.69E-32 | 6 |
| k141_34164_flag1_multi1.0000_len391 | W5-1 | 391 | XRL22790.1 hypothetical protein [Siniperca chuatsi ranavirus] | 93.9 | 66 | 7.45E-32 | 180.41 |
| k141_91965_flag0_multi1.0000_len238 | W2-1 | 238 | WXI69513.1 hypothetical protein [Largemouth bass virus] | 86.1 | 79 | 8.09E-32 | 72 |
| k141_60285_flag1_multi1.0000_len281 | W5-1 | 281 | XRB52842.1 transcription factor TFIIB cyclin-like domain-containing protein, partial [Largemouth bass virus] | 98.4 | 63 | 1.04E-31 | 0 |
| k141_118169_flag1_multi1.0000_len571 | W5-1 | 571 | AIG51690.1 major capsid protein, partial [Koi ranavirus] | 94.2 | 69 | 1.29E-31 | 2 |
| k141_105568_flag1_multi1.6689_len443 | W5-1 | 443 | XRL22821.1 hypothetical protein [Siniperca chuatsi ranavirus] | 96.8 | 62 | 3.33E-31 | 2.57 |
| k141_46899_flag1_multi114.0000_len791 | W6-2 | 791 | WAK75072.1 hypothetical protein [Mandarin fish ranavirus] | 100 | 60 | 4.34E-31 | 142 |
| k141_109619_flag1_multi14.2732_len1682 | W5-1 | 1682 | UVF58785.1 MAG: major capsid protein [Halichoeres melanurus ranavirus] | 97.2 | 71 | 6.43E-31 | 86 |
| k141_152057_flag1_multi1.0000_len446 | W5-3 | 446 | CBW45581.1 hypothetical protein, partial [Doctor fish virus] | 87 | 69 | 1.03E-30 | 13 |
| k141_73402_flag1_multi1.0000_len345 | W5-1 | 345 | AIG51690.1 major capsid protein, partial [Koi ranavirus] | 94.1 | 68 | 1.92E-30 | 0 |
| k141_22725_flag1_multi3.8165_len517 | W3-1 | 517 | UUY86269.1 hypothetical protein [Largemouth bass virus] | 98.4 | 63 | 2.08E-30 | 5 |
| k141_105093_flag1_multi5.3500_len861 | W6-1 | 861 | AIG51690.1 major capsid protein, partial [Koi ranavirus] | 98.5 | 65 | 4.54E-30 | 3 |
| k141_117063_flag0_multi1.0000_len383 | W5-1 | 383 | WAK75136.1 hypothetical protein [Mandarin fish ranavirus] | 100 | 57 | 1.09E-29 | 776.74 |
| k141_52924_flag1_multi4.9574_len704 | W1-1 | 704 | AYV88120.1 putative p31K protein [Mandarin fish ranavirus] | 96.8 | 63 | 1.18E-29 | 11 |
| k141_861_flag1_multi1.0000_len302 | W4-1 | 302 | UUY86195.1 hypothetical protein [Largemouth bass virus] | 94.9 | 59 | 1.21E-29 | 0 |
| k141_44142_flag1_multi1.0000_len320 | W4-1 | 320 | AIG51690.1 major capsid protein, partial [Koi ranavirus] | 94 | 67 | 1.48E-29 | 0 |
| k141_72438_flag1_multi1.0000_len476 | W4-1 | 476 | AYV88191.1 hypothetical protein [Mandarin fish ranavirus] | 77.6 | 76 | 1.69E-29 | 181 |
| k141_77731_flag1_multi5.8750_len557 | W4-1 | 557 | AYV88211.1 hypothetical protein [Mandarin fish ranavirus] | 96.9 | 65 | 1.74E-29 | 17 |
| k141_58569_flag1_multi1.9084_len818 | W5-1 | 818 | AFD96401.1 major capsid protein, partial [Largemouth bass virus] | 96.6 | 59 | 2.10E-29 | 4.83 |
| k141_129193_flag1_multi2.8957_len582 | W5-1 | 582 | WAK75094.1 hypothetical protein [Mandarin fish ranavirus] | 98 | 50 | 2.36E-29 | 1 |
| k141_113044_flag1_multi1.0000_len353 | W5-1 | 353 | AYV88120.1 putative p31K protein [Mandarin fish ranavirus] | 85.9 | 64 | 3.96E-29 | 0 |
| k141_106742_flag1_multi1.0000_len295 | W6-1 | 295 | UUY86241.1 hypothetical protein [Largemouth bass virus] | 100 | 48 | 6.07E-29 | 0 |
| k141_38160_flag1_multi3.0000_len1022 | W5-1 | 1022 | UUY86193.1 hypothetical protein [Largemouth bass virus] | 89.2 | 65 | 6.34E-29 | 9 |
| k141_57293_flag1_multi1.0000_len414 | W4-1 | 414 | QJE49149.1 putative immediate early protein ICP-46 [Largemouth bass virus] | 100 | 54 | 7.58E-29 | 135 |
| k141_82995_flag1_multi1.0000_len280 | W5-1 | 280 | AYV88191.1 hypothetical protein [Mandarin fish ranavirus] | 83.1 | 59 | 9.99E-29 | 0 |
| k141_66706_flag0_multi1.0000_len381 | W5-1 | 381 | UUY86261.1 hypothetical protein [Largemouth bass virus] | 98.4 | 63 | 2.04E-28 | 3 |
| k141_53661_flag1_multi133.1899_len2010 | W2-1 | 2010 | WHA35556.1 putative LPXTG-anchored collagen-like adhesin Scl2/SclB [Micropterus salmoides ranavirus] | 100 | 202 | 2.48E-28 | 648 |
| k141_162761_flag1_multi1.0000_len329 | W5-1 | 329 | WAK75068.1 hypothetical protein [Mandarin fish ranavirus] | 94.4 | 54 | 3.45E-28 | 2814.39 |
| k141_87350_flag1_multi6.6878_len583 | W4-1 | 583 | AYV88120.1 putative p31K protein [Mandarin fish ranavirus] | 66.7 | 96 | 5.78E-28 | 43 |

| | | | | | | | |
|---|---|---|---|---|---|---|---|
| k141_8888_flag1_multi1.0000_len281 | W5-1 | 281 | AYV88191.1 hypothetical protein [Mandarin fish ranavirus] | 96.2 | 53 | 8.12E-28 | 0 |
| k141_43805_flag1_multi1.0000_len481 | W5-1 | 481 | UUY86235.1 hypothetical protein [Largemouth bass virus] | 84.4 | 64 | 1.01E-27 | 11.48 |
| k141_23283_flag1_multi3.0135_len584 | W2-1 | 584 | WEI29006.1 putative 2-cysteine adaptor domain protein [Largemouth bass virus] | 98 | 50 | 1.51E-27 | 8 |
| k141_127062_flag0_multi7.2396_len454 | W1-1 | 454 | AIG51690.1 major capsid protein, partial [Koi ranavirus] | 98.2 | 57 | 1.56E-27 | 0 |
| k141_72180_flag1_multi1.0000_len399 | W4-1 | 399 | UUY86192.1 putative myristylated membrane protein [Largemouth bass virus] | 87.7 | 57 | 2.05E-27 | 9 |
| k141_11300_flag1_multi1.0000_len520 | W4-1 | 520 | AYV88194.1 hypothetical protein [Mandarin fish ranavirus] | 85.5 | 69 | 2.40E-27 | 3 |
| k141_59471_flag1_multi1.0000_len377 | W1-1 | 377 | QYU76034.1 putative myristylated membrane protein, partial [Koi ranavirus] | 79.1 | 67 | 2.57E-27 | 0 |
| k141_33544_flag1_multi6.0000_len353 | W3-1 | 353 | WAK75077.1 hypothetical protein [Mandarin fish ranavirus] | 100 | 52 | 2.90E-27 | 3 |
| k141_108486_flag1_multi3.7745_len447 | W5-1 | 447 | UUY86241.1 hypothetical protein [Largemouth bass virus] | 97.9 | 47 | 3.12E-27 | 2 |
| k141_101007_flag0_multi1.0000_len480 | W5-1 | 480 | UUY86241.1 hypothetical protein [Largemouth bass virus] | 97.9 | 47 | 3.34E-27 | 7 |
| k141_116248_flag1_multi1.0000_len504 | W1-1 | 504 | UVF58787.1 MAG: DNA-dependent RNA polymerase largest subunit, partial [Halichoeres melanurus ranavirus] | 94.5 | 55 | 3.78E-27 | 1 |
| k141_72042_flag1_multi1.0000_len385 | W4-1 | 385 | UUY86267.1 ribonucleotide reductase alpha subunit [Largemouth bass virus] | 85.7 | 63 | 5.76E-27 | 1 |
| k141_66748_flag0_multi1.0000_len348 | W5-1 | 348 | QJE49215.1 putative orf58-like protein [Largemouth bass virus] | 87.1 | 62 | 6.11E-27 | 3 |
| k141_22_flag0_multi7.7600_len941 | W5-1 | 941 | UUY86269.1 hypothetical protein [Largemouth bass virus] | 92.4 | 66 | 7.72E-27 | 27.98 |
| k141_104027_flag0_multi475.0000_len1602 | W1-1 | 1602 | AIG51690.1 major capsid protein, partial [Koi ranavirus] | 100 | 58 | 8.98E-27 | 1212.04 |
| k141_168715_flag1_multi1.0000_len588 | W5-1 | 588 | UUY86241.1 hypothetical protein [Largemouth bass virus] | 61.8 | 89 | 1.07E-26 | 2 |
| k141_78715_flag1_multi1.7372_len415 | W4-1 | 415 | UUY86229.1 putative DNA dependent RNA polymerase a subunit [Largemouth bass virus] | 95.2 | 63 | 1.26E-26 | 1 |
| k141_121153_flag0_multi13643.0000_len2220 | W5-1 | 2220 | CBW45581.1 hypothetical protein, partial [Doctor fish virus] | 87 | 69 | 1.26E-26 | 1345 |
| k141_2249_flag1_multi16.8258_len2254 | W5-1 | 2254 | UUY86227.1 hypothetical protein [Largemouth bass virus] | 94.9 | 59 | 1.43E-26 | 131 |
| k141_67942_flag1_multi2.9064_len739 | W1-1 | 739 | UUY86241.1 hypothetical protein [Largemouth bass virus] | 79.4 | 63 | 1.62E-26 | 12 |
| k141_179472_flag1_multi9.0000_len911 | W1-3 | 911 | QJE49090.1 hypothetical protein LMBV_027 [Largemouth bass virus] | 96.7 | 60 | 1.91E-26 | 27 |
| k141_6913_flag0_multi1.0000_len279 | W5-1 | 279 | WAK75112.1 putative LITAF PIG7 possible membrane associated motif in LPS-induced tumor necrosis factor alpha factor [Mandarin fish ranavirus] | 100 | 46 | 2.03E-26 | 0 |
| k141_86911_flag1_multi1.0000_len373 | W6-1 | 373 | AIG51690.1 major capsid protein, partial [Koi ranavirus] | 74.7 | 79 | 2.41E-26 | 15 |
| k141_128489_flag1_multi36.1614_len3723 | W5-1 | 3723 | WAK75073.1 hypothetical protein [Mandarin fish ranavirus] | 94.7 | 57 | 2.50E-26 | 554 |
| k141_1389_flag1_multi1.0000_len319 | W1-1 | 319 | AIG51690.1 major capsid protein, partial [Koi ranavirus] | 86.6 | 67 | 3.32E-26 | 1146 |
| k141_137807_flag1_multi1.0000_len310 | W5-1 | 310 | AFD96401.1 major capsid protein, partial [Largemouth bass virus] | 85 | 60 | 3.38E-26 | 6 |
| k141_97577_flag1_multi3.7738_len446 | W4-1 | 446 | WHU98721.1 MCP, partial [Hybrid snakehead ranavirus] | 100 | 54 | 3.71E-26 | 948 |
| k141_85098_flag1_multi1.0000_len311 | W5-1 | 311 | AAC79876.1 viral core protein, partial [Labroides dimidatus ranavirus] | 95.9 | 49 | 4.20E-26 | 0 |
| k141_117340_flag1_multi3.0000_len313 | W6-3 | 313 | WAK75103.1 hypothetical protein [Mandarin fish ranavirus] | 100 | 52 | 4.50E-26 | 3 |
| k141_33936_flag1_multi12.5453_len748 | W1-1 | 748 | AYV88191.1 hypothetical protein [Mandarin fish ranavirus] | 97.9 | 47 | 4.68E-26 | 22 |
| k141_76735_flag0_multi1.0000_len270 | W2-1 | 270 | AAC79876.1 viral core protein, partial [Labroides dimidatus ranavirus] | 100 | 46 | 4.91E-26 | 0 |
| k141_73262_flag1_multi1.0000_len280 | W2-2 | 280 | WAK75112.1 putative LITAF PIG7 possible membrane associated motif in LPS-induced tumor necrosis factor alpha factor [Mandarin fish ranavirus] | 97.9 | 47 | 5.80E-26 | 0 |
| k141_65614_flag0_multi99.3625_len301 | W2-1 | 301 | UUY86250.1 hypothetical protein [Largemouth bass virus] | 100 | 57 | 7.75E-26 | 2 |
| k141_65032_flag0_multi71.5825_len335 | W5-1 | 335 | AAC79876.1 viral core protein, partial [Labroides dimidatus ranavirus] | 92.2 | 51 | 8.07E-26 | 0 |
| k141_91258_flag1_multi1.0000_len522 | W1-1 | 522 | WAK75072.1 hypothetical protein [Mandarin fish ranavirus] | 100 | 55 | 9.53E-26 | 210 |
| k141_39458_flag1_multi1.0000_len272 | W1-1 | 272 | AYV88191.1 hypothetical protein [Mandarin fish ranavirus] | 100 | 46 | 9.57E-26 | 0 |

| | | | | | | | |
|---|---|---|---|---|---|---|---|
| k141_44430_flag1_multi1.8674_len488 | W5-1 | 488 | WEI29006.1 putative 2-cysteine adaptor domain protein [Largemouth bass virus] | 97.9 | 47 | 9.86E-26 | 4 |
| k141_82305_flag1_multi1.0000_len324 | W5-1 | 324 | WAK75112.1 putative LITAF PIG7 possible membrane associated motif in LPS-induced tumor necrosis factor alpha factor [Mandarin fish ranavirus] | 100 | 46 | 9.95E-26 | 0 |
| k141_124236_flag1_multi1.9428_len543 | W2-2 | 543 | UUY86241.1 hypothetical protein [Largemouth bass virus] | 97.8 | 46 | 1.05E-25 | 4 |
| k141_153587_flag1_multi1.0000_len281 | W5-1 | 281 | AYV88191.1 hypothetical protein [Mandarin fish ranavirus] | 100 | 46 | 1.08E-25 | 0 |
| k141_115864_flag1_multi2.7263_len521 | W5-1 | 521 | UUY86241.1 hypothetical protein [Largemouth bass virus] | 95.7 | 47 | 1.15E-25 | 5 |
| k141_104940_flag1_multi1.0000_len320 | W5-1 | 320 | QJE49112.1 hypothetical protein LMBV_049 [Largemouth bass virus] | 97.9 | 47 | 1.17E-25 | 0 |
| k141_29471_flag0_multi1.0000_len278 | W5-1 | 278 | WAK75107.1 hypothetical protein [Mandarin fish ranavirus] | 100 | 45 | 1.41E-25 | 0 |
| k141_37838_flag1_multi1.0000_len278 | W5-1 | 278 | WAK75112.1 putative LITAF PIG7 possible membrane associated motif in LPS-induced tumor necrosis factor alpha factor [Mandarin fish ranavirus] | 100 | 46 | 1.60E-25 | 0 |
| k141_122486_flag1_multi1.0000_len279 | W1-1 | 279 | QJE49077.1 hypothetical protein LMBV_014 [Largemouth bass virus] | 91.8 | 49 | 1.79E-25 | 0 |
| k141_9480_flag0_multi1.0000_len292 | W5-1 | 292 | AIG51690.1 major capsid protein, partial [Koi ranavirus] | 92.7 | 55 | 2.42E-25 | 2.07 |
| k141_64324_flag1_multi3.7893_len554 | W4-1 | 554 | WAK75112.1 putative LITAF PIG7 possible membrane associated motif in LPS-induced tumor necrosis factor alpha factor [Mandarin fish ranavirus] | 98 | 49 | 2.45E-25 | 10 |
| k141_96141_flag1_multi4.3359_len531 | W1-1 | 531 | UUY86241.1 hypothetical protein [Largemouth bass virus] | 97.8 | 46 | 2.52E-25 | 9 |
| k141_46707_flag1_multi6.6280_len512 | W4-1 | 512 | AFD96401.1 major capsid protein, partial [Largemouth bass virus] | 98 | 50 | 2.58E-25 | 6 |
| k141_26148_flag1_multi1.0000_len278 | W4-1 | 278 | CBW45583.1 ICP 46 homolog, partial [Doctor fish virus] | 100 | 46 | 2.61E-25 | 0 |
| k141_34813_flag1_multi2.8827_len533 | W5-1 | 533 | AAC79876.1 viral core protein, partial [Labroides dimidatus ranavirus] | 100 | 46 | 2.77E-25 | 4 |
| k141_167045_flag1_multi2.8663_len470 | W5-1 | 470 | UUY86199.1 hypothetical protein [Largemouth bass virus] | 100 | 46 | 2.98E-25 | 6 |
| k141_49335_flag1_multi3.8139_len1092 | W5-1 | 1092 | UUY86241.1 hypothetical protein [Largemouth bass virus] | 92.5 | 53 | 3.06E-25 | 15 |
| k141_178855_flag0_multi58.0000_len1301 | W2-3 | 1301 | WHA35556.1 putative LPXTG-anchored collagen-like adhesin Scl2/SclB [Micropterus salmoides ranavirus] | 99 | 199 | 3.14E-25 | 78 |
| k141_6879_flag0_multi15.4642_len783 | W1-1 | 783 | QJE49077.1 hypothetical protein LMBV_014 [Largemouth bass virus] | 96 | 50 | 3.22E-25 | 22 |
| k141_153370_flag1_multi2.9322_len539 | W5-1 | 539 | WEI29006.1 putative 2-cysteine adaptor domain protein [Largemouth bass virus] | 97.9 | 47 | 3.27E-25 | 4 |
| k141_109588_flag1_multi28.9688_len783 | W5-1 | 783 | WAK75112.1 putative LITAF PIG7 possible membrane associated motif in LPS-induced tumor necrosis factor alpha factor [Mandarin fish ranavirus] | 100 | 48 | 3.69E-25 | 40 |
| k141_66330_flag1_multi1.0000_len280 | W5-1 | 280 | WAK75107.1 hypothetical protein [Mandarin fish ranavirus] | 100 | 45 | 4.03E-25 | 1 |
| k141_14274_flag0_multi1.0000_len280 | W5-1 | 280 | AYV88191.1 hypothetical protein [Mandarin fish ranavirus] | 100 | 46 | 4.35E-25 | 0 |
| k141_142487_flag1_multi4.7923_len584 | W5-1 | 584 | UUY86241.1 hypothetical protein [Largemouth bass virus] | 97.8 | 46 | 4.43E-25 | 4 |
| k141_120826_flag1_multi3.9244_len657 | W1-1 | 657 | AAC79876.1 viral core protein, partial [Labroides dimidatus ranavirus] | 90.4 | 52 | 4.71E-25 | 21 |
| k141_91212_flag1_multi4.7333_len486 | W5-1 | 486 | UUY86199.1 hypothetical protein [Largemouth bass virus] | 100 | 46 | 4.97E-25 | 4 |
| k141_129534_flag1_multi1.0000_len588 | W1-1 | 588 | WAK75112.1 putative LITAF PIG7 possible membrane associated motif in LPS-induced tumor necrosis factor alpha factor [Mandarin fish ranavirus] | 100 | 47 | 5.00E-25 | 0 |
| k141_122367_flag1_multi2.6803_len654 | W1-1 | 654 | AYV88191.1 hypothetical protein [Mandarin fish ranavirus] | 88.7 | 53 | 6.22E-25 | 7 |
| k141_87611_flag1_multi1.0000_len419 | W5-1 | 419 | WAK75112.1 putative LITAF PIG7 possible membrane associated motif in LPS-induced tumor necrosis factor alpha factor [Mandarin fish ranavirus] | 97.9 | 47 | 6.40E-25 | 1 |
| k141_14833_flag1_multi1.0000_len464 | W1-1 | 464 | AYV88191.1 hypothetical protein [Mandarin fish ranavirus] | 97.9 | 47 | 7.02E-25 | 17.09 |
| k141_1040_flag1_multi5.1812_len279 | W1-1 | 279 | WAK75107.1 hypothetical protein [Mandarin fish ranavirus] | 100 | 45 | 8.13E-25 | 0 |
| k141_27467_flag1_multi4.8185_len648 | W1-1 | 648 | AAC79876.1 viral core protein, partial [Labroides dimidatus ranavirus] | 100 | 46 | 8.64E-25 | 8 |
| k141_111406_flag1_multi7.7755_len858 | W2-1 | 858 | AYV88191.1 hypothetical protein [Mandarin fish ranavirus] | 100 | 46 | 9.22E-25 | 13 |

| | | | | | | | |
|---|---|---|---|---|---|---|---|
| k141_76753_flag1_multi2.8909_len581 | W5-1 | 581 | UUY86199.1 hypothetical protein [Largemouth bass virus] | 100 | 46 | 9.62E-25 | 3 |
| k141_67427_flag1_multi1.5756_len617 | W5-1 | 617 | QJE49097.1 hypothetical protein LMBV_034 [Largemouth bass virus] | 89.8 | 59 | 1.07E-24 | 4 |
| k141_36770_flag0_multi62.6134_len1957 | W5-1 | 1957 | QJE49137.1 hypothetical protein LMBV_074 [Largemouth bass virus] | 94.9 | 59 | 1.31E-24 | 309 |
| k141_86746_flag0_multi30.5790_len2065 | W4-1 | 2065 | AAF64582.1 capsid protein, partial [Guppyfish iridovirus] | 93.3 | 60 | 1.41E-24 | 158 |
| k141_40140_flag1_multi1.9450_len559 | W1-1 | 559 | AAC79876.1 viral core protein, partial [Labroides dimidatus ranavirus] | 95.8 | 48 | 1.43E-24 | 3 |
| k141_39552_flag0_multi1.0000_len281 | W1-1 | 281 | QIZ30887.1 major capsid protein, partial [Largemouth bass virus] | 100 | 46 | 1.47E-24 | 0 |
| k141_159064_flag1_multi2.2204_len631 | W5-1 | 631 | QJE49078.1 putative DNA methyltransferase [Largemouth bass virus] | 100 | 46 | 1.84E-24 | 4 |
| k141_131166_flag1_multi2.6945_len691 | W5-1 | 691 | UUY86266.1 hypothetical protein [Largemouth bass virus] | 88.1 | 59 | 2.01E-24 | 14.4 |
| k141_151418_flag1_multi1.0000_len269 | W5-1 | 269 | WAK75094.1 hypothetical protein [Mandarin fish ranavirus] | 100 | 41 | 2.01E-24 | 0 |
| k141_69568_flag1_multi4.8067_len617 | W5-1 | 617 | CBW45583.1 ICP 46 homolog, partial [Doctor fish virus] | 97.9 | 48 | 2.12E-24 | 4 |
| k141_32209_flag1_multi2.8514_len437 | W5-1 | 437 | AYV88120.1 putative p31K protein [Mandarin fish ranavirus] | 70.5 | 78 | 2.25E-24 | 4 |
| k141_108613_flag1_multi39.2249_len590 | W5-1 | 590 | AYV88191.1 hypothetical protein [Mandarin fish ranavirus] | 100 | 46 | 2.67E-24 | 82 |
| k141_131353_flag0_multi1248.4265_len685 | W1-1 | 685 | UUY86227.1 hypothetical protein [Largemouth bass virus] | 87.8 | 49 | 2.79E-24 | 4 |
| k141_125453_flag1_multi5.6858_len507 | W5-1 | 507 | XRB52842.1 transcription factor TFIIB cyclin-like domain-containing protein, partial [Largemouth bass virus] | 100 | 46 | 2.94E-24 | 2 |
| k141_5070_flag1_multi4.7588_len1368 | W5-1 | 1368 | UUY86197.1 hypothetical protein [Largemouth bass virus] | 95.3 | 64 | 3.84E-24 | 32 |
| k141_74904_flag1_multi1.0000_len499 | W1-1 | 499 | WAK75073.1 hypothetical protein [Mandarin fish ranavirus] | 100 | 46 | 4.16E-24 | 4 |
| k141_40092_flag1_multi4.5556_len357 | W5-1 | 357 | AYV88191.1 hypothetical protein [Mandarin fish ranavirus] | 100 | 46 | 4.69E-24 | 0 |
| k141_113299_flag0_multi1.0000_len279 | W5-1 | 279 | WAK75107.1 hypothetical protein [Mandarin fish ranavirus] | 100 | 44 | 4.89E-24 | 0 |
| k141_68654_flag0_multi12.4751_len644 | W4-2 | 644 | AYV88172.2 hypothetical protein [Mandarin fish ranavirus] | 92.2 | 167 | 4.91E-24 | 13 |
| k141_73067_flag1_multi1.5298_len560 | W5-1 | 560 | AYV88176.1 putative tumor necrosis factor receptor [Mandarin fish ranavirus] | 100 | 46 | 5.17E-24 | 3 |
| k141_41922_flag1_multi1.0000_len279 | W1-1 | 279 | QJE49146.1 putative thiol oxidoreductase [Largemouth bass virus] | 87 | 54 | 5.21E-24 | 0 |
| k141_72440_flag1_multi1.0000_len308 | W5-1 | 308 | UUY86235.1 hypothetical protein [Largemouth bass virus] | 100 | 46 | 5.42E-24 | 0 |
| k141_34557_flag1_multi1.0000_len405 | W5-1 | 405 | AYV88191.1 hypothetical protein [Mandarin fish ranavirus] | 100 | 46 | 5.68E-24 | 2 |
| k141_3091_flag1_multi4.8168_len1495 | W4-1 | 1495 | XPZ21295.1 putative RNAseIII [Mandarin fish ranavirus] | 69.3 | 88 | 5.70E-24 | 21 |
| k141_121564_flag1_multi15.6748_len922 | W2-3 | 922 | UUY86241.1 hypothetical protein [Largemouth bass virus] | 100 | 46 | 5.75E-24 | 49 |
| k141_131264_flag1_multi1.0000_len269 | W5-1 | 269 | UUY86241.1 hypothetical protein [Largemouth bass virus] | 100 | 39 | 5.75E-24 | 0 |
| k141_12672_flag1_multi7.8169_len982 | W5-1 | 982 | AYV88191.1 hypothetical protein [Mandarin fish ranavirus] | 100 | 46 | 6.01E-24 | 19 |
| k141_94879_flag0_multi17.5057_len841 | W1-1 | 841 | XRB52785.1 hypothetical protein LMBV_28 [Largemouth bass virus] | 100 | 46 | 6.34E-24 | 21 |
| k141_143126_flag1_multi3.8985_len791 | W5-1 | 791 | UUY86199.1 hypothetical protein [Largemouth bass virus] | 100 | 46 | 6.34E-24 | 9 |
| k141_58772_flag1_multi8.8692_len569 | W1-1 | 569 | UUY86199.1 hypothetical protein [Largemouth bass virus] | 100 | 46 | 6.82E-24 | 14 |
| k141_29715_flag1_multi7.0421_len1163 | W6-1 | 1163 | WAK75094.1 hypothetical protein [Mandarin fish ranavirus] | 100 | 46 | 7.22E-24 | 20 |
| k141_53177_flag1_multi2.4590_len446 | W5-1 | 446 | UUY86235.1 hypothetical protein [Largemouth bass virus] | 83.1 | 59 | 7.24E-24 | 2 |
| k141_197906_flag1_multi7.9472_len3192 | W4-3 | 3192 | AYV88172.2 hypothetical protein [Mandarin fish ranavirus] | 92.9 | 170 | 8.09E-24 | 85 |
| k141_164336_flag0_multi1.0000_len334 | W5-1 | 334 | AFD96401.1 major capsid protein, partial [Largemouth bass virus] | 100 | 46 | 8.13E-24 | 1 |
| k141_102173_flag1_multi1.9010_len434 | W5-1 | 434 | UUY86235.1 hypothetical protein [Largemouth bass virus] | 100 | 47 | 8.89E-24 | 2 |
| k141_42653_flag1_multi1.0000_len277 | W5-1 | 277 | UUY86192.1 putative myristylated membrane protein [Largemouth bass virus] | 100 | 46 | 9.89E-24 | 0 |
| k141_135165_flag1_multi1.0000_len363 | W1-1 | 363 | UUY86235.1 hypothetical protein [Largemouth bass virus] | 78.5 | 65 | 1.06E-23 | 0 |
| k141_133299_flag0_multi1.6756_len440 | W1-1 | 440 | WAK75094.1 hypothetical protein [Mandarin fish ranavirus] | 80 | 55 | 1.14E-23 | 1 |
| k141_35662_flag1_multi15.5312_len877 | W2-1 | 877 | UUY86199.1 hypothetical protein [Largemouth bass virus] | 100 | 46 | 1.22E-23 | 50 |

| | | | | | | | |
|---|---|---|---|---|---|---|---|
| k141_16131_flag1_multi2.8701_len495 | W4-1 | 495 | AYV88120.1 putative p31K protein [Mandarin fish ranavirus] | 100 | 48 | 1.23E-23 | 15 |
| k141_154686_flag0_multi19.3444_len1764 | W5-1 | 1764 | AFD96401.1 major capsid protein, partial [Largemouth bass virus] | 100 | 51 | 1.29E-23 | 97.96 |
| k141_52324_flag1_multi1.0000_len280 | W5-1 | 280 | UUY86263.1 hypothetical protein [Largemouth bass virus] | 100 | 46 | 1.30E-23 | 0 |
| k141_184_flag1_multi2.6613_len389 | W5-1 | 389 | AYV88191.1 hypothetical protein [Mandarin fish ranavirus] | 100 | 44 | 1.35E-23 | 1 |
| k141_82495_flag0_multi1.0000_len277 | W5-1 | 277 | QYU76034.1 putative myristylated membrane protein, partial [Koi ranavirus] | 89.5 | 57 | 1.38E-23 | 1 |
| k141_18245_flag1_multi5.0000_len517 | W6-1 | 517 | AYV88120.1 putative p31K protein [Mandarin fish ranavirus] | 89.3 | 56 | 1.49E-23 | 9 |
| k141_81563_flag0_multi1.0000_len282 | W1-1 | 282 | AIG51690.1 major capsid protein, partial [Koi ranavirus] | 95.9 | 49 | 1.56E-23 | 2 |
| k141_98926_flag1_multi6.5577_len453 | W4-1 | 453 | AYV88155.1 hypothetical protein [Mandarin fish ranavirus] | 100 | 46 | 1.58E-23 | 0 |
| k141_96640_flag1_multi1.0000_len280 | W4-1 | 280 | WEI29006.1 putative 2-cysteine adaptor domain protein [Largemouth bass virus] | 78.3 | 69 | 1.75E-23 | 0 |
| k141_91948_flag1_multi2.9247_len566 | W6-1 | 566 | WEI29006.1 putative 2-cysteine adaptor domain protein [Largemouth bass virus] | 100 | 44 | 1.79E-23 | 3 |
| k141_50773_flag1_multi2.8969_len626 | W1-1 | 626 | WEI28972.1 CTD-phosphotransferase [Largemouth bass virus] | 100 | 46 | 1.81E-23 | 8 |
| k141_5029_flag1_multi3.7125_len381 | W5-1 | 381 | QJE49215.1 putative orf58-like protein [Largemouth bass virus] | 100 | 44 | 1.95E-23 | 4 |
| k141_94017_flag0_multi1.0000_len276 | W2-1 | 276 | UUY86235.1 hypothetical protein [Largemouth bass virus] | 97.9 | 47 | 1.98E-23 | 0 |
| k141_35845_flag1_multi1.0000_len506 | W3-2 | 506 | UUY86235.1 hypothetical protein [Largemouth bass virus] | 100 | 46 | 1.98E-23 | 0 |
| k141_15533_flag1_multi1.0000_len372 | W1-1 | 372 | WAK75106.1 hypothetical protein [Mandarin fish ranavirus] | 75.7 | 70 | 2.11E-23 | 0 |
| k141_36558_flag1_multi1.0000_len316 | W5-1 | 316 | AYV88191.1 hypothetical protein [Mandarin fish ranavirus] | 97.7 | 43 | 2.17E-23 | 0 |
| k141_82698_flag0_multi7.7828_len1591 | W5-1 | 1591 | WAK75112.1 putative LITAF PIG7 possible membrane associated motif in LPS-induced tumor necrosis factor alpha factor [Mandarin fish ranavirus] | 95.8 | 48 | 2.35E-23 | 53 |
| k141_9380_flag1_multi1.0000_len432 | W5-1 | 432 | QJE49128.1 hypothetical protein LMBV_065 [Largemouth bass virus] | 100 | 46 | 2.50E-23 | 0 |
| k141_152259_flag1_multi1.9716_len1021 | W5-1 | 1021 | WEI29006.1 putative 2-cysteine adaptor domain protein [Largemouth bass virus] | 100 | 46 | 2.56E-23 | 18 |
| k141_75588_flag0_multi15.5977_len1993 | W5-1 | 1993 | UUY86241.1 hypothetical protein [Largemouth bass virus] | 100 | 46 | 2.70E-23 | 127.71 |
| k141_107154_flag0_multi1.0000_len208 | W1-1 | 208 | UUY86197.1 hypothetical protein [Largemouth bass virus] | 91.9 | 74 | 2.71E-23 | 3 |
| k141_12140_flag1_multi9.8087_len1223 | W1-1 | 1223 | QJE49149.1 putative immediate early protein ICP-46 [Largemouth bass virus] | 100 | 46 | 2.74E-23 | 36 |
| k141_51991_flag0_multi60.3407_len913 | W1-1 | 913 | AYV88191.1 hypothetical protein [Mandarin fish ranavirus] | 76.6 | 64 | 2.86E-23 | 76 |
| k141_163272_flag0_multi1.0000_len413 | W5-1 | 413 | QJE49077.1 hypothetical protein LMBV_014 [Largemouth bass virus] | 100 | 43 | 3.03E-23 | 97 |
| k141_50367_flag1_multi21.4896_len717 | W5-1 | 717 | WEI29006.1 putative 2-cysteine adaptor domain protein [Largemouth bass virus] | 94 | 50 | 3.22E-23 | 10 |
| k141_126924_flag0_multi2.8945_len615 | W1-1 | 615 | AFD96401.1 major capsid protein, partial [Largemouth bass virus] | 100 | 47 | 3.24E-23 | 7 |
| k141_29759_flag1_multi22.0000_len1645 | W5-1 | 1645 | UUY86241.1 hypothetical protein [Largemouth bass virus] | 95.9 | 49 | 3.24E-23 | 121 |
| k141_123083_flag1_multi3.4458_len870 | W5-1 | 870 | QJE49065.1 hypothetical protein LMBV_002 [Largemouth bass virus] | 100 | 47 | 3.31E-23 | 6 |
| k141_135314_flag1_multi2.4572_len679 | W5-1 | 679 | QJE49080.1 hypothetical protein LMBV_017 [Largemouth bass virus] | 58.2 | 91 | 3.46E-23 | 6 |
| k141_99572_flag1_multi1.0000_len370 | W4-1 | 370 | QJE49215.1 putative orf58-like protein [Largemouth bass virus] | 100 | 46 | 3.52E-23 | 8 |
| k141_159951_flag0_multi1.0000_len264 | W5-1 | 264 | AYV88134.2 putative tyrosine kinase [Mandarin fish ranavirus] | 86.8 | 53 | 3.52E-23 | 0 |
| k141_96993_flag1_multi3.4721_len535 | W5-1 | 535 | AYV88155.1 hypothetical protein [Mandarin fish ranavirus] | 76.6 | 64 | 3.79E-23 | 6 |
| k141_8800_flag1_multi2.6078_len373 | W1-1 | 373 | AFD96401.1 major capsid protein, partial [Largemouth bass virus] | 93.9 | 49 | 3.85E-23 | 11 |
| k141_14688_flag1_multi1.0000_len518 | W1-1 | 518 | UUY86241.1 hypothetical protein [Largemouth bass virus] | 97.7 | 43 | 3.89E-23 | 7 |
| k141_26442_flag1_multi1.0000_len428 | W5-1 | 428 | AYV88211.1 hypothetical protein [Mandarin fish ranavirus] | 94.5 | 55 | 3.92E-23 | 1 |
| k141_35247_flag1_multi1.0000_len390 | W4-1 | 390 | AYV88120.1 putative p31K protein [Mandarin fish ranavirus] | 100 | 47 | 4.01E-23 | 354.92 |
| k141_168940_flag0_multi1.0000_len279 | W5-1 | 279 | QJE49215.1 putative orf58-like protein [Largemouth bass virus] | 97.9 | 48 | 4.44E-23 | 0 |
| k141_68888_flag1_multi1.6978_len280 | W5-1 | 280 | QJE49096.1 putative D5 family NTPase/ATPase [Largemouth bass virus] | 86 | 57 | 4.49E-23 | 0 |
| k141_59301_flag0_multi18.0705_len297 | W1-1 | 297 | QYU76034.1 putative myristylated membrane protein, partial [Koi ranavirus] | 100 | 46 | 4.64E-23 | 0 |

| | | | | | | | |
|---|---|---|---|---|---|---|---|
| k141_74124_flag0_multi1.0000_len347 | W4-1 | 347 | UUY86235.1 hypothetical protein [Largemouth bass virus] | 92.3 | 52 | 4.94E-23 | 21 |
| k141_43565_flag0_multi5.4843_len810 | W5-1 | 810 | AYV88191.1 hypothetical protein [Mandarin fish ranavirus] | 100 | 44 | 5.16E-23 | 16 |
| k141_12341_flag0_multi1.0000_len278 | W4-1 | 278 | AYV88120.1 putative p31K protein [Mandarin fish ranavirus] | 95.7 | 47 | 5.32E-23 | 4 |
| k141_43967_flag0_multi1.0000_len336 | W5-1 | 336 | WAK75068.1 hypothetical protein [Mandarin fish ranavirus] | 97.8 | 46 | 5.58E-23 | 0 |
| k141_87618_flag0_multi1.0000_len774 | W5-1 | 774 | WAK75112.1 putative LITAF PIG7 possible membrane associated motif in LPS-induced tumor necrosis factor alpha factor [Mandarin fish ranavirus] | 97.9 | 47 | 5.82E-23 | 17 |
| k141_54522_flag1_multi1.0000_len280 | W5-1 | 280 | UUY86235.1 hypothetical protein [Largemouth bass virus] | 95.9 | 49 | 6.02E-23 | 0 |
| k141_132256_flag1_multi2.8827_len499 | W1-1 | 499 | QJE49149.1 putative immediate early protein ICP-46 [Largemouth bass virus] | 100 | 46 | 6.36E-23 | 5 |
| k141_24308_flag0_multi1.0000_len305 | W4-1 | 305 | UUY86199.1 hypothetical protein [Largemouth bass virus] | 93.2 | 44 | 6.41E-23 | 0 |
| k141_66722_flag1_multi10.5330_len899 | W5-1 | 899 | QJE49077.1 hypothetical protein LMBV_014 [Largemouth bass virus] | 100 | 46 | 6.43E-23 | 26 |
| k141_36581_flag1_multi3.8889_len762 | W1-1 | 762 | AYV88191.1 hypothetical protein [Mandarin fish ranavirus] | 97.8 | 46 | 6.96E-23 | 9 |
| k141_159139_flag1_multi1.0000_len269 | W5-1 | 269 | UUY86235.1 hypothetical protein [Largemouth bass virus] | 100 | 46 | 7.27E-23 | 0 |
| k141_133371_flag1_multi1.0000_len280 | W5-1 | 280 | WAK75136.1 hypothetical protein [Mandarin fish ranavirus] | 100 | 46 | 7.64E-23 | 0 |
| k141_47002_flag1_multi6.6167_len501 | W4-1 | 501 | AYV88134.2 putative tyrosine kinase [Mandarin fish ranavirus] | 100 | 47 | 7.96E-23 | 7 |
| k141_1855_flag1_multi1.0000_len380 | W1-1 | 380 | AAF64582.1 capsid protein, partial [Guppyfish iridovirus] | 90.6 | 53 | 8.06E-23 | 1 |
| k141_101369_flag0_multi30.9148_len493 | W5-1 | 493 | QJE49119.1 hypothetical protein LMBV_056 [Largemouth bass virus] | 100 | 46 | 8.14E-23 | 11 |
| k141_91018_flag1_multi3.6174_len1223 | W1-1 | 1223 | QJE49114.1 hypothetical protein LMBV_051 [Largemouth bass virus] | 100 | 46 | 8.50E-23 | 15 |
| k141_66567_flag1_multi4.8375_len707 | W1-1 | 707 | AYV88191.1 hypothetical protein [Mandarin fish ranavirus] | 95.9 | 49 | 8.76E-23 | 11 |
| k141_17675_flag1_multi1.0000_len280 | W2-1 | 280 | AYV88126.1 putative immediate early protein ICP-18 [Mandarin fish ranavirus] | 100 | 46 | 8.80E-23 | 0 |
| k141_104896_flag0_multi1.0000_len290 | W4-1 | 290 | AIG51690.1 major capsid protein, partial [Koi ranavirus] | 97.9 | 47 | 9.21E-23 | 60 |
| k141_158123_flag0_multi2.9674_len1369 | W5-1 | 1369 | UUY86199.1 hypothetical protein [Largemouth bass virus] | 88.7 | 53 | 1.02E-22 | 14 |
| k141_18336_flag0_multi1.0000_len405 | W5-1 | 405 | AYV88183.1 hypothetical protein [Mandarin fish ranavirus] | 100 | 46 | 1.04E-22 | 0 |
| k141_154454_flag1_multi1.0000_len280 | W5-1 | 280 | QJE49066.1 hypothetical protein LMBV_003 [Largemouth bass virus] | 100 | 46 | 1.08E-22 | 0 |
| k141_50478_flag0_multi1.0000_len374 | W1-1 | 374 | AYV88134.2 putative tyrosine kinase [Mandarin fish ranavirus] | 100 | 46 | 1.10E-22 | 13.33 |
| k141_108852_flag0_multi1.0000_len376 | W1-1 | 376 | AFD96401.1 major capsid protein, partial [Largemouth bass virus] | 79.7 | 64 | 1.13E-22 | 1 |
| k141_75449_flag0_multi1.0000_len278 | W1-1 | 278 | UUY86235.1 hypothetical protein [Largemouth bass virus] | 100 | 46 | 1.15E-22 | 1 |
| k141_139321_flag0_multi1.8659_len305 | W5-1 | 305 | AYV88159.1 hypothetical protein [Mandarin fish ranavirus] | 100 | 46 | 1.16E-22 | 0 |
| k141_108045_flag1_multi1.0000_len292 | W1-1 | 292 | QYU76034.1 putative myristylated membrane protein, partial [Koi ranavirus] | 100 | 48 | 1.19E-22 | 0 |
| k141_65065_flag1_multi1.0000_len504 | W5-1 | 504 | AYV88176.1 putative tumor necrosis factor receptor [Mandarin fish ranavirus] | 100 | 43 | 1.20E-22 | 2 |
| k141_95380_flag1_multi1.0000_len280 | W6-1 | 280 | QJE49104.1 hypothetical protein LMBV_041 [Largemouth bass virus] | 100 | 46 | 1.26E-22 | 0 |
| k141_4717_flag1_multi1.0000_len387 | W5-1 | 387 | AYV88134.2 putative tyrosine kinase [Mandarin fish ranavirus] | 94.1 | 51 | 1.33E-22 | 0 |
| k141_90620_flag1_multi1.0000_len387 | W6-2 | 387 | UUY86257.1 hypothetical protein [Largemouth bass virus] | 87.5 | 56 | 1.33E-22 | 24 |
| k141_16270_flag1_multi9.0000_len801 | W4-1 | 801 | WAK75112.1 putative LITAF PIG7 possible membrane associated motif in LPS-induced tumor necrosis factor alpha factor [Mandarin fish ranavirus] | 100 | 44 | 1.39E-22 | 139 |
| k141_80610_flag1_multi2.8950_len579 | W5-1 | 579 | AYV88191.1 hypothetical protein [Mandarin fish ranavirus] | 97.9 | 47 | 1.48E-22 | 2 |
| k141_2348_flag1_multi2.4727_len856 | W5-1 | 856 | QJE49215.1 putative orf58-like protein [Largemouth bass virus] | 100 | 46 | 1.60E-22 | 6 |
| k141_96195_flag1_multi1.0000_len708 | W5-1 | 708 | UUY86251.1 hypothetical protein [Largemouth bass virus] | 100 | 46 | 1.62E-22 | 7 |
| k141_96648_flag1_multi4.0864_len662 | W4-1 | 662 | AYV88134.2 putative tyrosine kinase [Mandarin fish ranavirus] | 83.1 | 65 | 1.70E-22 | 9 |
| k141_127361_flag1_multi2.8982_len593 | W5-1 | 593 | AYV88191.1 hypothetical protein [Mandarin fish ranavirus] | 87 | 54 | 1.72E-22 | 9 |
| k141_160390_flag1_multi1.8978_len366 | W5-1 | 366 | UUY86235.1 hypothetical protein [Largemouth bass virus] | 100 | 46 | 1.73E-22 | 1 |

| | | | | | | | |
|---|---|---|---|---|---|---|---|
| k141_79483_flag1_multi2.8783_len519 | W6-1 | 519 | QYU76034.1 putative myristylated membrane protein, partial [Koi ranavirus] | 100 | 46 | 1.76E-22 | 4 |
| k141_125302_flag1_multi2.8655_len483 | W5-1 | 483 | AIG51690.1 major capsid protein, partial [Koi ranavirus] | 95.9 | 49 | 1.77E-22 | 3 |
| k141_142502_flag0_multi1.0000_len330 | W5-1 | 330 | AFD96401.1 major capsid protein, partial [Largemouth bass virus] | 71.6 | 67 | 1.78E-22 | 0 |
| k141_23666_flag1_multi15.7169_len579 | W4-1 | 579 | AFD96401.1 major capsid protein, partial [Largemouth bass virus] | 97.9 | 48 | 1.79E-22 | 11 |
| k141_25661_flag0_multi1.0000_len374 | W5-1 | 374 | AIG51690.1 major capsid protein, partial [Koi ranavirus] | 100 | 47 | 1.90E-22 | 0 |
| k141_45677_flag1_multi1.0000_len311 | W5-1 | 311 | AYV88173.2 putative ribonucleotide reductase beta subunit [Mandarin fish ranavirus] | 85.5 | 55 | 1.94E-22 | 0 |
| k141_22109_flag0_multi9.4314_len345 | W4-1 | 345 | AYV88159.1 hypothetical protein [Mandarin fish ranavirus] | 100 | 47 | 1.96E-22 | 3 |
| k141_168257_flag1_multi1.0000_len276 | W5-1 | 276 | WEI28972.1 CTD-phosphotransferase [Largemouth bass virus] | 100 | 47 | 1.98E-22 | 0.95 |
| k141_91079_flag1_multi14.7007_len1217 | W5-1 | 1217 | AYV88191.1 hypothetical protein [Mandarin fish ranavirus] | 97.9 | 47 | 2.12E-22 | 62 |
| k141_65644_flag1_multi3.8493_len599 | W4-1 | 599 | AFD96401.1 major capsid protein, partial [Largemouth bass virus] | 100 | 46 | 2.20E-22 | 6 |
| k141_101527_flag0_multi1.0000_len279 | W4-1 | 279 | AIG51690.1 major capsid protein, partial [Koi ranavirus] | 100 | 47 | 2.21E-22 | 1 |
| k141_101527_flag0_multi1.0000_len279 | W4-1 | 279 | AIG51690.1 major capsid protein, partial [Koi ranavirus] | 100 | 47 | 2.21E-22 | 0 |
| k141_161467_flag1_multi1.0000_len279 | W5-1 | 279 | AYV88191.1 hypothetical protein [Mandarin fish ranavirus] | 97.8 | 46 | 2.22E-22 | 0 |
| k141_100350_flag0_multi1.0000_len279 | W5-1 | 279 | UUY86235.1 hypothetical protein [Largemouth bass virus] | 100 | 46 | 2.30E-22 | 0 |
| k141_20890_flag0_multi1.0000_len279 | W5-1 | 279 | UUY86235.1 hypothetical protein [Largemouth bass virus] | 100 | 46 | 2.30E-22 | 0 |
| k141_146336_flag1_multi2.9103_len1033 | W5-1 | 1033 | AYV88191.1 hypothetical protein [Mandarin fish ranavirus] | 97.8 | 46 | 2.32E-22 | 11 |
| k141_93020_flag0_multi1.9499_len580 | W5-1 | 580 | UUY86199.1 hypothetical protein [Largemouth bass virus] | 60.5 | 86 | 2.38E-22 | 3 |
| k141_116367_flag1_multi4.7996_len650 | W5-1 | 650 | QIZ30887.1 major capsid protein, partial [Largemouth bass virus] | 100 | 46 | 2.45E-22 | 7 |
| k141_138126_flag0_multi1.0000_len682 | W5-1 | 682 | XPZ21295.1 putative RNAseIII [Mandarin fish ranavirus] | 76.6 | 64 | 2.48E-22 | 2 |
| k141_14590_flag0_multi1.0000_len401 | W1-1 | 401 | UUY86235.1 hypothetical protein [Largemouth bass virus] | 97.9 | 47 | 2.65E-22 | 0 |
| k141_19812_flag0_multi33.5131_len1903 | W5-1 | 1903 | AYV88191.1 hypothetical protein [Mandarin fish ranavirus] | 74.2 | 66 | 2.68E-22 | 211 |
| k141_35856_flag1_multi1.0000_len662 | W5-1 | 662 | QYU76034.1 putative myristylated membrane protein, partial [Koi ranavirus] | 100 | 46 | 2.85E-22 | 10 |
| k141_109485_flag1_multi10.7486_len1223 | W2-2 | 1223 | WAK75112.1 putative LITAF PIG7 possible membrane associated motif in LPS-induced tumor necrosis factor alpha factor [Mandarin fish ranavirus] | 71.6 | 67 | 2.90E-22 | 45 |
| k141_108930_flag0_multi172.0268_len290 | W5-1 | 290 | QYU76034.1 putative myristylated membrane protein, partial [Koi ranavirus] | 100 | 46 | 3.03E-22 | 1 |
| k141_3174_flag1_multi6.7915_len803 | W2-1 | 803 | UUY86235.1 hypothetical protein [Largemouth bass virus] | 100 | 48 | 3.11E-22 | 16 |
| k141_83135_flag1_multi2.9257_len760 | W1-1 | 760 | QIZ30887.1 major capsid protein, partial [Largemouth bass virus] | 100 | 46 | 3.18E-22 | 9 |
| k141_105610_flag1_multi5.8477_len896 | W2-3 | 896 | QJE49125.1 hypothetical protein LMBV_062 [Largemouth bass virus] | 98.1 | 54 | 3.27E-22 | 18 |
| k141_21169_flag1_multi5.6361_len457 | W2-1 | 457 | AYV88191.1 hypothetical protein [Mandarin fish ranavirus] | 97.8 | 46 | 3.37E-22 | 4 |
| k141_125300_flag1_multi3.4303_len729 | W5-1 | 729 | AYV88183.1 hypothetical protein [Mandarin fish ranavirus] | 98 | 49 | 3.38E-22 | 5 |
| k141_10698_flag1_multi1.8515_len1097 | W4-1 | 1097 | UUY86195.1 hypothetical protein [Largemouth bass virus] | 92.3 | 52 | 3.46E-22 | 16 |
| k141_94551_flag0_multi1.0000_len340 | W5-1 | 340 | UUY86235.1 hypothetical protein [Largemouth bass virus] | 100 | 46 | 3.48E-22 | 2 |
| k141_9915_flag1_multi1.0000_len531 | W5-1 | 531 | WEI29006.1 putative 2-cysteine adaptor domain protein [Largemouth bass virus] | 95.7 | 47 | 3.61E-22 | 19 |
| k141_28189_flag0_multi1.0000_len279 | W5-1 | 279 | WEI28972.1 CTD-phosphotransferase [Largemouth bass virus] | 92.2 | 51 | 3.75E-22 | 0 |
| k141_104014_flag1_multi2.9184_len705 | W6-1 | 705 | UUY86235.1 hypothetical protein [Largemouth bass virus] | 66.2 | 77 | 3.78E-22 | 4 |
| k141_106483_flag0_multi3.6786_len561 | W5-1 | 561 | UUY86235.1 hypothetical protein [Largemouth bass virus] | 97.9 | 47 | 3.81E-22 | 1 |
| k141_54376_flag1_multi1.9404_len527 | W4-1 | 527 | QYU76035.1 DNA polymerase, partial [Koi ranavirus] | 100 | 46 | 3.87E-22 | 2 |
| k141_36748_flag1_multi1.0000_len275 | W4-1 | 275 | AYV88120.1 putative p31K protein [Mandarin fish ranavirus] | 93.9 | 49 | 3.96E-22 | 139 |
| k141_3551_flag1_multi3.3323_len797 | W4-1 | 797 | UUY86199.1 hypothetical protein [Largemouth bass virus] | 100 | 44 | 3.99E-22 | 18 |
| k141_29367_flag1_multi1.6429_len589 | W6-1 | 589 | AFD96401.1 major capsid protein, partial [Largemouth bass virus] | 87 | 54 | 4.00E-22 | 3 |

| | | | | | | | |
|---|---|---|---|---|---|---|---|
| k141_95609_flag1_multi1.0000_len279 | W6-1 | 279 | AYV88120.1 putative p31K protein [Mandarin fish ranavirus] | 90.2 | 51 | 4.11E-22 | 0 |
| k141_55465_flag1_multi15.9300_len484 | W6-1 | 484 | QJE49119.1 hypothetical protein LMBV_056 [Largemouth bass virus] | 65.9 | 82 | 4.17E-22 | 26 |
| k141_3808_flag1_multi1.0000_len338 | W5-1 | 338 | WEI29006.1 putative 2-cysteine adaptor domain protein [Largemouth bass virus] | 95.7 | 47 | 4.44E-22 | 3 |
| k141_94187_flag1_multi3.1073_len495 | W4-1 | 495 | WEI29006.1 putative 2-cysteine adaptor domain protein [Largemouth bass virus] | 100 | 46 | 4.49E-22 | 2 |
| k141_72495_flag1_multi1.9510_len610 | W5-1 | 610 | QYU76034.1 putative myristylated membrane protein, partial [Koi ranavirus] | 97.9 | 47 | 4.58E-22 | 6 |
| k141_143602_flag1_multi2.8516_len451 | W5-1 | 451 | AIG51690.1 major capsid protein, partial [Koi ranavirus] | 95.9 | 49 | 4.74E-22 | 1 |
| k141_38550_flag1_multi1.0000_len280 | W4-1 | 280 | QYU76034.1 putative myristylated membrane protein, partial [Koi ranavirus] | 100 | 46 | 4.97E-22 | 4.51 |
| k141_97022_flag0_multi1.0000_len278 | W5-1 | 278 | QYU76034.1 putative myristylated membrane protein, partial [Koi ranavirus] | 100 | 46 | 4.97E-22 | 9 |
| k141_34161_flag0_multi1.0000_len432 | W5-1 | 432 | AYV88120.1 putative p31K protein [Mandarin fish ranavirus] | 100 | 47 | 5.03E-22 | 15.01 |
| k141_99760_flag1_multi1.0000_len276 | W4-1 | 276 | QJE49104.1 hypothetical protein LMBV_041 [Largemouth bass virus] | 97.9 | 47 | 5.10E-22 | 3 |
| k141_9506_flag1_multi6.0848_len471 | W1-1 | 471 | UUY86241.1 hypothetical protein [Largemouth bass virus] | 100 | 40 | 5.15E-22 | 17 |
| k141_49681_flag0_multi3.7850_len448 | W5-1 | 448 | AFD96401.1 major capsid protein, partial [Largemouth bass virus] | 100 | 45 | 5.17E-22 | 3 |
| k141_10120_flag1_multi6.5701_len462 | W2-1 | 462 | AIG51690.1 major capsid protein, partial [Koi ranavirus] | 100 | 48 | 5.42E-22 | 10 |
| k141_3368_flag1_multi1.0000_len353 | W5-1 | 353 | AIG51690.1 major capsid protein, partial [Koi ranavirus] | 95.8 | 48 | 5.51E-22 | 0 |
| k141_61072_flag1_multi1.9644_len590 | W5-1 | 590 | WEI28974.1 hypothetical protein [Largemouth bass virus] | 93.8 | 48 | 5.72E-22 | 3 |
| k141_49065_flag1_multi1.0000_len444 | W6-1 | 444 | AYV88120.1 putative p31K protein [Mandarin fish ranavirus] | 97.9 | 47 | 5.76E-22 | 0 |
| k141_122591_flag1_multi1.0000_len757 | W5-1 | 757 | QJE49215.1 putative orf58-like protein [Largemouth bass virus] | 97.9 | 47 | 5.78E-22 | 11 |
| k141_19474_flag0_multi1.9027_len398 | W5-1 | 398 | AFD96401.1 major capsid protein, partial [Largemouth bass virus] | 97.8 | 46 | 5.78E-22 | 2 |
| k141_239378_flag1_multi6.8475_len967 | W4-3 | 967 | UUY86199.1 hypothetical protein [Largemouth bass virus] | 100 | 45 | 6.33E-22 | 23 |
| k141_92333_flag1_multi174.2957_len1629 | W5-1 | 1629 | WAK75079.1 hypothetical protein [Mandarin fish ranavirus] | 92 | 50 | 6.59E-22 | 700.68 |
| k141_138522_flag1_multi8.8630_len1484 | W5-1 | 1484 | AYV88191.1 hypothetical protein [Mandarin fish ranavirus] | 97.9 | 48 | 6.94E-22 | 42 |
| k141_57199_flag1_multi1.4310_len373 | W5-1 | 373 | QJE49066.1 hypothetical protein LMBV_003 [Largemouth bass virus] | 85.2 | 54 | 6.98E-22 | 1 |
| k141_94390_flag0_multi9.4000_len1146 | W5-1 | 1146 | QJE49077.1 hypothetical protein LMBV_014 [Largemouth bass virus] | 90.6 | 53 | 7.41E-22 | 33 |
| k141_161788_flag0_multi3.4981_len408 | W5-1 | 408 | AYV88191.1 hypothetical protein [Mandarin fish ranavirus] | 100 | 46 | 7.87E-22 | 4 |
| k141_92757_flag0_multi16.8085_len329 | W5-1 | 329 | AYV88120.1 putative p31K protein [Mandarin fish ranavirus] | 100 | 46 | 7.88E-22 | 0 |
| k141_48182_flag1_multi2.8844_len539 | W4-1 | 539 | QYU76034.1 putative myristylated membrane protein, partial [Koi ranavirus] | 100 | 46 | 8.03E-22 | 5 |
| k141_88255_flag1_multi5.6788_len499 | W4-1 | 499 | AIG51690.1 major capsid protein, partial [Koi ranavirus] | 100 | 46 | 8.06E-22 | 7 |
| k141_79804_flag1_multi4.9240_len944 | W4-1 | 944 | QIZ30887.1 major capsid protein, partial [Largemouth bass virus] | 100 | 46 | 8.32E-22 | 18 |
| k141_101922_flag1_multi3.7957_len988 | W5-1 | 988 | QYU76036.1 helicase-like protein, partial [Koi ranavirus] | 98.2 | 55 | 8.93E-22 | 17 |
| k141_5985_flag1_multi1.0000_len422 | W5-1 | 422 | WEI29006.1 putative 2-cysteine adaptor domain protein [Largemouth bass virus] | 100 | 46 | 9.17E-22 | 3 |
| k141_107147_flag0_multi89.3862_len677 | W1-1 | 677 | XRB52842.1 transcription factor TFIIB cyclin-like domain-containing protein, partial [Largemouth bass virus] | 97.9 | 47 | 9.33E-22 | 213 |
| k141_6148_flag0_multi14.6412_len2106 | W1-1 | 2106 | AIG51690.1 major capsid protein, partial [Koi ranavirus] | 88.9 | 63 | 9.40E-22 | 1374.02 |
| k141_145088_flag0_multi1.0000_len280 | W5-1 | 280 | UUY86235.1 hypothetical protein [Largemouth bass virus] | 100 | 46 | 9.46E-22 | 0 |
| k141_151461_flag1_multi1.0000_len279 | W5-1 | 279 | QYU76034.1 putative myristylated membrane protein, partial [Koi ranavirus] | 100 | 46 | 9.51E-22 | 0 |
| k141_67381_flag1_multi2.7469_len623 | W1-1 | 623 | QJE49146.1 putative thiol oxidoreductase [Largemouth bass virus] | 100 | 46 | 9.78E-22 | 9 |
| k141_5505_flag0_multi1.0000_len582 | W1-1 | 582 | AIG51690.1 major capsid protein, partial [Koi ranavirus] | 97.9 | 48 | 1.00E-21 | 32 |
| k141_23153_flag1_multi1.0000_len655 | W2-3 | 655 | AYV88169.1 hypothetical protein [Mandarin fish ranavirus] | 97.9 | 48 | 1.01E-21 | 8 |
| k141_46425_flag1_multi6.3658_len562 | W5-1 | 562 | AYV88120.1 putative p31K protein [Mandarin fish ranavirus] | 88.7 | 53 | 1.02E-21 | 15 |
| k141_115107_flag1_multi9.4732_len589 | W5-1 | 589 | AIG51690.1 major capsid protein, partial [Koi ranavirus] | 82 | 61 | 1.03E-21 | 27 |

| | | | | | | | |
|---|---|---|---|---|---|---|---|
| k141_137877_flag1_multi10.2075_len406 | W5-1 | 406 | WHU98721.1 MCP, partial [Hybrid snakehead ranavirus] | 100 | 46 | 1.06E-21 | 1 |
| k141_77283_flag1_multi1.0000_len269 | W5-1 | 269 | AXB27397.1 major capsid protein, partial [Ranavirus sp.] | 95.7 | 46 | 1.09E-21 | 0 |
| k141_82820_flag0_multi1.0000_len279 | W5-1 | 279 | QJE49153.1 hypothetical protein LMBV_004 [Largemouth bass virus] | 97.9 | 47 | 1.13E-21 | 0 |
| k141_108358_flag1_multi2.8757_len511 | W5-1 | 511 | QJE49097.1 hypothetical protein LMBV_034 [Largemouth bass virus] | 93.9 | 49 | 1.16E-21 | 6 |
| k141_89629_flag1_multi6.8416_len482 | W1-1 | 482 | UUY86192.1 putative myristylated membrane protein [Largemouth bass virus] | 100 | 42 | 1.24E-21 | 7 |
| k141_68650_flag1_multi4.8680_len838 | W5-1 | 838 | QYU76034.1 putative myristylated membrane protein, partial [Koi ranavirus] | 100 | 46 | 1.27E-21 | 16 |
| k141_112452_flag1_multi8.8875_len1563 | W2-1 | 1563 | QYU76035.1 DNA polymerase, partial [Koi ranavirus] | 98 | 50 | 1.31E-21 | 55 |
| k141_14419_flag1_multi9.4570_len2476 | W5-1 | 2476 | QJE49077.1 hypothetical protein LMBV_014 [Largemouth bass virus] | 97.8 | 46 | 1.37E-21 | 74 |
| k141_95499_flag1_multi1.0000_len280 | W1-1 | 280 | XRB52768.1 P31K protein, partial [Largemouth bass virus] | 97.8 | 46 | 1.45E-21 | 0 |
| k141_33355_flag1_multi1.0000_len434 | W4-1 | 434 | AYV88191.1 hypothetical protein [Mandarin fish ranavirus] | 97.8 | 46 | 1.46E-21 | 15.1 |
| k141_15503_flag0_multi1.0000_len363 | W5-1 | 363 | WEI29006.1 putative 2-cysteine adaptor domain protein [Largemouth bass virus] | 100 | 45 | 1.53E-21 | 1 |
| k141_76344_flag1_multi27.6336_len755 | W5-1 | 755 | AYV88120.1 putative p31K protein [Mandarin fish ranavirus] | 87.3 | 55 | 1.53E-21 | 121 |
| k141_53812_flag1_multi2.7074_len640 | W2-1 | 640 | XRL22821.1 hypothetical protein [Siniperca chuatsi ranavirus] | 89.3 | 56 | 1.56E-21 | 3 |
| k141_66974_flag1_multi7.9047_len1831 | W5-1 | 1831 | XRB52842.1 transcription factor TFIIB cyclin-like domain-containing protein, partial [Largemouth bass virus] | 97.9 | 47 | 1.57E-21 | 41 |
| k141_102732_flag1_multi1.0000_len278 | W4-1 | 278 | AYV88120.1 putative p31K protein [Mandarin fish ranavirus] | 93.9 | 49 | 1.61E-21 | 11 |
| k141_23617_flag1_multi2.5211_len734 | W5-1 | 734 | QJE49137.1 hypothetical protein LMBV_074 [Largemouth bass virus] | 95.8 | 48 | 1.62E-21 | 5 |
| k141_100770_flag1_multi2.0000_len572 | W1-1 | 572 | UUY86192.1 putative myristylated membrane protein [Largemouth bass virus] | 95.7 | 47 | 1.65E-21 | 3 |
| k141_45220_flag0_multi2.7606_len517 | W5-1 | 517 | WXI69525.1 caspase recruitment domain protein [Largemouth bass virus] | 95.8 | 48 | 1.67E-21 | 5 |
| k141_129068_flag1_multi1.0000_len352 | W5-1 | 352 | QJE49104.1 hypothetical protein LMBV_041 [Largemouth bass virus] | 97.9 | 47 | 1.85E-21 | 5 |
| k141_109798_flag1_multi3.4627_len932 | W5-1 | 932 | QYU76034.1 putative myristylated membrane protein, partial [Koi ranavirus] | 100 | 46 | 1.87E-21 | 10 |
| k141_82460_flag1_multi3.7621_len431 | W4-1 | 431 | WEI29006.1 putative 2-cysteine adaptor domain protein [Largemouth bass virus] | 100 | 46 | 1.90E-21 | 7 |
| k141_66672_flag1_multi3.0841_len652 | W5-1 | 652 | UUY86221.1 hypothetical protein [Largemouth bass virus] | 65.1 | 83 | 2.00E-21 | 5 |
| k141_36147_flag0_multi41.8846_len297 | W5-1 | 297 | QJE49153.1 hypothetical protein LMBV_004 [Largemouth bass virus] | 92.2 | 51 | 2.01E-21 | 41 |
| k141_18887_flag1_multi4.7942_len588 | W4-1 | 588 | WEI28972.1 CTD-phosphotransferase [Largemouth bass virus] | 100 | 47 | 2.05E-21 | 8 |
| k141_39131_flag0_multi1.0000_len451 | W2-1 | 451 | AFD96401.1 major capsid protein, partial [Largemouth bass virus] | 88.2 | 51 | 2.13E-21 | 2 |
| k141_19588_flag1_multi1.0000_len513 | W5-1 | 513 | UUY86258.1 hypothetical protein [Largemouth bass virus] | 88.9 | 54 | 2.20E-21 | 1 |
| k141_149209_flag1_multi1.0000_len307 | W2-3 | 307 | AIG51690.1 major capsid protein, partial [Koi ranavirus] | 100 | 46 | 2.27E-21 | 1 |
| k141_86101_flag1_multi7.7905_len1678 | W5-1 | 1678 | WAK75112.1 putative LITAF PIG7 possible membrane associated motif in LPS-induced tumor necrosis factor alpha factor [Mandarin fish ranavirus] | 100 | 46 | 2.28E-21 | 43 |
| k141_8349_flag1_multi1.0000_len323 | W1-1 | 323 | WEI29006.1 putative 2-cysteine adaptor domain protein [Largemouth bass virus] | 100 | 46 | 2.36E-21 | 0 |
| k141_6958_flag1_multi7.6873_len1014 | W4-1 | 1014 | WAK75114.1 hypothetical protein [Mandarin fish ranavirus] | 100 | 46 | 2.40E-21 | 23 |
| k141_42120_flag0_multi352.3462_len817 | W2-2 | 817 | AYV88120.1 putative p31K protein [Mandarin fish ranavirus] | 85.5 | 55 | 2.46E-21 | 1578 |
| k141_154855_flag0_multi1.0000_len325 | W5-1 | 325 | WEI28972.1 CTD-phosphotransferase [Largemouth bass virus] | 100 | 46 | 2.49E-21 | 2 |
| k141_57224_flag0_multi1.0000_len303 | W4-1 | 303 | WEI29006.1 putative 2-cysteine adaptor domain protein [Largemouth bass virus] | 100 | 46 | 2.55E-21 | 0 |
| k141_55157_flag1_multi1.0000_len329 | W5-1 | 329 | UVF58793.1 MAG: DNA polymerase [Halichoeres melanurus ranavirus] | 97.9 | 47 | 2.58E-21 | 1 |
| k141_33337_flag1_multi1.0000_len269 | W1-1 | 269 | QJE49104.1 hypothetical protein LMBV_041 [Largemouth bass virus] | 97.9 | 47 | 2.60E-21 | 0 |
| k141_110518_flag1_multi2.8921_len660 | W4-2 | 660 | UUY86235.1 hypothetical protein [Largemouth bass virus] | 100 | 46 | 2.71E-21 | 5 |
| k141_147399_flag1_multi6.0571_len631 | W5-1 | 631 | UUY86223.1 hypothetical protein [Largemouth bass virus] | 76.6 | 64 | 2.74E-21 | 8 |
| k141_28339_flag1_multi18.6487_len2242 | W5-1 | 2242 | UUY86199.1 hypothetical protein [Largemouth bass virus] | 95.9 | 49 | 2.74E-21 | 148 |

| | | | | | | | |
|---|---|---|---|---|---|---|---|
| k141_161137_flag1_multi1.0000_len379 | W5-1 | 379 | XPZ21295.1 putative RNAseIII [Mandarin fish ranavirus] | 100 | 44 | 2.74E-21 | 1 |
| k141_133033_flag0_multi1.0000_len270 | W5-1 | 270 | AIG51690.1 major capsid protein, partial [Koi ranavirus] | 74.6 | 71 | 2.76E-21 | 0 |
| k141_67565_flag1_multi20.3669_len975 | W4-1 | 975 | AYV88120.1 putative p31K protein [Mandarin fish ranavirus] | 98 | 49 | 2.80E-21 | 81 |
| k141_45940_flag0_multi44.2096_len618 | W1-1 | 618 | WAK75139.1 putative myristylated membrane protein [Mandarin fish ranavirus] | 80.8 | 125 | 2.89E-21 | 457 |
| k141_123398_flag0_multi16.8640_len560 | W5-1 | 560 | UUY86218.1 hypothetical protein [Largemouth bass virus] | 85.2 | 54 | 2.94E-21 | 69 |
| k141_43553_flag0_multi1.0000_len332 | W4-1 | 332 | AIG51690.1 major capsid protein, partial [Koi ranavirus] | 95.9 | 49 | 3.08E-21 | 0 |
| k141_42395_flag1_multi1.0000_len379 | W5-1 | 379 | UUY86235.1 hypothetical protein [Largemouth bass virus] | 77.6 | 67 | 3.11E-21 | 0 |
| k141_44649_flag0_multi1.0000_len279 | W5-1 | 279 | AIG51690.1 major capsid protein, partial [Koi ranavirus] | 100 | 47 | 3.11E-21 | 1 |
| k141_43602_flag1_multi28.7494_len568 | W4-1 | 568 | AIG51690.1 major capsid protein, partial [Koi ranavirus] | 100 | 46 | 3.19E-21 | 60 |
| k141_20708_flag1_multi1.9368_len505 | W5-1 | 505 | WHU98721.1 MCP, partial [Hybrid snakehead ranavirus] | 100 | 46 | 3.25E-21 | 80 |
| k141_27209_flag1_multi1.0000_len277 | W5-1 | 277 | WEI29006.1 putative 2-cysteine adaptor domain protein [Largemouth bass virus] | 100 | 45 | 3.33E-21 | 3 |
| k141_21224_flag1_multi1.0000_len279 | W6-1 | 279 | XRL22821.1 hypothetical protein [Siniperca chuatsi ranavirus] | 92.2 | 51 | 3.36E-21 | 0 |
| k141_8097_flag1_multi7.5695_len515 | W4-1 | 515 | AYV88120.1 putative p31K protein [Mandarin fish ranavirus] | 92.3 | 52 | 3.37E-21 | 9 |
| k141_74101_flag1_multi7.4958_len1458 | W5-1 | 1458 | QJE49081.1 hypothetical protein LMBV_018 [Largemouth bass virus] | 96 | 50 | 3.49E-21 | 35 |
| k141_154346_flag0_multi1.0000_len316 | W5-1 | 316 | AIG51690.1 major capsid protein, partial [Koi ranavirus] | 100 | 46 | 3.54E-21 | 1 |
| k141_94539_flag1_multi7.6454_len595 | W4-1 | 595 | QYU76034.1 putative myristylated membrane protein, partial [Koi ranavirus] | 95.8 | 48 | 3.67E-21 | 8 |
| k141_26105_flag1_multi7.7925_len917 | W4-1 | 917 | WEI29006.1 putative 2-cysteine adaptor domain protein [Largemouth bass virus] | 90.7 | 54 | 3.68E-21 | 18 |
| k141_31172_flag0_multi1.0000_len441 | W5-1 | 441 | AYV88176.1 putative tumor necrosis factor receptor [Mandarin fish ranavirus] | 100 | 41 | 3.73E-21 | 5 |
| k141_163799_flag1_multi2.4991_len684 | W5-1 | 684 | AFD96401.1 major capsid protein, partial [Largemouth bass virus] | 100 | 45 | 3.79E-21 | 16 |
| k141_77124_flag1_multi1.0000_len269 | W5-1 | 269 | AYV88120.1 putative p31K protein [Mandarin fish ranavirus] | 100 | 43 | 3.97E-21 | 0 |
| k141_77671_flag1_multi3.8421_len578 | W4-1 | 578 | WHA35533.1 putative DNA dependent RNA polymerase A subunit [Micropterus salmoides ranavirus] | 81.7 | 60 | 4.09E-21 | 7 |
| k141_114607_flag1_multi5.5023_len362 | W5-1 | 362 | QJE49104.1 hypothetical protein LMBV_041 [Largemouth bass virus] | 100 | 46 | 4.12E-21 | 3 |
| k141_131553_flag1_multi5.7935_len698 | W5-1 | 698 | UUY86195.1 hypothetical protein [Largemouth bass virus] | 97.8 | 46 | 4.30E-21 | 17 |
| k141_155402_flag0_multi1.0000_len281 | W5-1 | 281 | QJE49215.1 putative orf58-like protein [Largemouth bass virus] | 100 | 40 | 4.30E-21 | 0 |
| k141_80853_flag1_multi16.6089_len1082 | W4-1 | 1082 | XRB52842.1 transcription factor TFIIB cyclin-like domain-containing protein, partial [Largemouth bass virus] | 100 | 46 | 4.34E-21 | 66 |
| k141_214261_flag1_multi13.3944_len1353 | W5-3 | 1353 | WAK75072.1 hypothetical protein [Mandarin fish ranavirus] | 100 | 49 | 4.43E-21 | 35 |
| k141_62374_flag1_multi3.0116_len571 | W5-1 | 571 | AIG51690.1 major capsid protein, partial [Koi ranavirus] | 100 | 46 | 4.57E-21 | 4 |
| k141_86674_flag1_multi3.3052_len508 | W5-1 | 508 | AYV88191.1 hypothetical protein [Mandarin fish ranavirus] | 88.5 | 52 | 4.63E-21 | 5 |
| k141_104006_flag0_multi1.3692_len401 | W1-1 | 401 | AYV88179.2 putative DNA dependent RNA polymerase II second largest subunit [Mandarin fish ranavirus] | 100 | 46 | 4.71E-21 | 0 |
| k141_55127_flag1_multi3.5462_len1179 | W4-1 | 1179 | AYV88191.1 hypothetical protein [Mandarin fish ranavirus] | 97.8 | 46 | 4.76E-21 | 12 |
| k141_95510_flag0_multi5.1036_len498 | W1-1 | 498 | XRB52768.1 P31K protein, partial [Largemouth bass virus] | 91.8 | 49 | 4.79E-21 | 6 |
| k141_23010_flag1_multi1.0000_len351 | W2-1 | 351 | XRL22790.1 hypothetical protein [Siniperca chuatsi ranavirus] | 100 | 46 | 4.81E-21 | 0 |
| k141_26476_flag0_multi1.0000_len282 | W4-1 | 282 | AXB27397.1 major capsid protein, partial [Ranavirus sp.] | 95.7 | 46 | 5.11E-21 | 0 |
| k141_93117_flag1_multi5.6788_len499 | W4-1 | 499 | QJE49153.1 hypothetical protein LMBV_004 [Largemouth bass virus] | 100 | 46 | 5.16E-21 | 12 |
| k141_29543_flag1_multi19.8303_len2698 | W5-1 | 2698 | QJE49128.1 hypothetical protein LMBV_065 [Largemouth bass virus] | 97.8 | 46 | 5.31E-21 | 147 |
| k141_92989_flag0_multi1.0000_len306 | W5-1 | 306 | XRL22790.1 hypothetical protein [Siniperca chuatsi ranavirus] | 100 | 46 | 5.43E-21 | 1 |
| k141_56200_flag1_multi3.8825_len728 | W5-1 | 728 | UUY86238.1 hypothetical protein [Largemouth bass virus] | 97.9 | 48 | 5.46E-21 | 12 |

| | | | | | | | |
|---|---|---|---|---|---|---|---|
| k141_69996_flag1_multi16.3458_len702 | W1-1 | 702 | AYV88179.2 putative DNA dependent RNA polymerase II second largest subunit [Mandarin fish ranavirus] | 94 | 50 | 5.60E-21 | 34 |
| k141_96149_flag0_multi6.0000_len278 | W5-1 | 278 | UUY86258.1 hypothetical protein [Largemouth bass virus] | 100 | 46 | 5.74E-21 | 0 |
| k141_119322_flag1_multi1.0000_len387 | W5-1 | 387 | QJE49112.1 hypothetical protein LMBV_049 [Largemouth bass virus] | 83 | 53 | 5.76E-21 | 1 |
| k141_28559_flag1_multi1.0000_len402 | W5-1 | 402 | UUY86258.1 hypothetical protein [Largemouth bass virus] | 63.9 | 83 | 5.78E-21 | 0 |
| k141_32572_flag1_multi1.0000_len383 | W5-1 | 383 | AYV88120.1 putative p31K protein [Mandarin fish ranavirus] | 100 | 46 | 5.91E-21 | 1 |
| k141_109281_flag1_multi2.8386_len426 | W5-1 | 426 | UUY86227.1 hypothetical protein [Largemouth bass virus] | 97.7 | 44 | 6.18E-21 | 7 |
| k141_157482_flag1_multi173.9635_len4001 | W5-1 | 4001 | AYV88133.1 putative DNA-directed RNA polymerase II subunit [Mandarin fish ranavirus] | 80.3 | 61 | 6.22E-21 | 466 |
| k141_148639_flag1_multi8.6614_len1157 | W5-1 | 1157 | WEI28974.1 hypothetical protein [Largemouth bass virus] | 97.9 | 48 | 6.31E-21 | 30 |
| k141_147440_flag1_multi1.0000_len685 | W5-1 | 685 | QYU76034.1 putative myristylated membrane protein, partial [Koi ranavirus] | 52.6 | 116 | 6.40E-21 | 7 |
| k141_117852_flag1_multi3.8000_len486 | W5-1 | 486 | QJE49101.1 hypothetical protein LMBV_038 [Largemouth bass virus] | 98 | 49 | 6.63E-21 | 6 |
| k141_139037_flag1_multi2.7594_len889 | W5-1 | 889 | XPZ21295.1 putative RNAseIII [Mandarin fish ranavirus] | 100 | 46 | 6.75E-21 | 7.01 |
| k141_4361_flag1_multi1.6483_len559 | W5-1 | 559 | QYU76034.1 putative myristylated membrane protein, partial [Koi ranavirus] | 100 | 46 | 6.92E-21 | 0 |
| k141_4653_flag0_multi6.2380_len952 | W5-1 | 952 | WAK75107.1 hypothetical protein [Mandarin fish ranavirus] | 93.6 | 47 | 6.97E-21 | 80 |
| k141_43930_flag1_multi5.9103_len999 | W5-1 | 999 | WAK75073.1 hypothetical protein [Mandarin fish ranavirus] | 97.9 | 47 | 7.20E-21 | 18 |
| k141_56288_flag0_multi49.2251_len674 | W5-1 | 674 | UUY86263.1 hypothetical protein [Largemouth bass virus] | 100 | 46 | 7.34E-21 | 92 |
| k141_128876_flag1_multi1.9406_len528 | W5-1 | 528 | AYV88120.1 putative p31K protein [Mandarin fish ranavirus] | 93.9 | 49 | 7.50E-21 | 44 |
| k141_111755_flag1_multi12.8269_len1735 | W5-1 | 1735 | WAK75112.1 putative LITAF PIG7 possible membrane associated motif in LPS-induced tumor necrosis factor alpha factor [Mandarin fish ranavirus] | 100 | 46 | 8.55E-21 | 131 |
| k141_73943_flag1_multi4.7306_len334 | W4-1 | 334 | AIG51690.1 major capsid protein, partial [Koi ranavirus] | 81.8 | 55 | 8.58E-21 | 4 |
| k141_119057_flag0_multi1.0000_len333 | W5-1 | 333 | QJE49153.1 hypothetical protein LMBV_004 [Largemouth bass virus] | 83 | 53 | 8.61E-21 | 2 |
| k141_82572_flag1_multi1.0000_len590 | W5-1 | 590 | WAK75072.1 hypothetical protein [Mandarin fish ranavirus] | 97.9 | 47 | 8.65E-21 | 4 |
| k141_31599_flag1_multi19.6827_len412 | W6-1 | 412 | QJE49125.1 hypothetical protein LMBV_062 [Largemouth bass virus] | 100 | 46 | 8.70E-21 | 1 |
| k141_57745_flag1_multi3.9086_len896 | W5-1 | 896 | AYV88155.1 hypothetical protein [Mandarin fish ranavirus] | 100 | 46 | 8.74E-21 | 8 |
| k141_59037_flag0_multi1.0000_len523 | W5-1 | 523 | WEI28972.1 CTD-phosphotransferase [Largemouth bass virus] | 100 | 46 | 9.23E-21 | 4 |
| k141_99436_flag1_multi3.8357_len561 | W6-2 | 561 | QJE49153.1 hypothetical protein LMBV_004 [Largemouth bass virus] | 87.3 | 55 | 9.49E-21 | 4 |
| k141_41669_flag1_multi6.5197_len928 | W5-1 | 928 | AYV88191.1 hypothetical protein [Mandarin fish ranavirus] | 100 | 45 | 9.58E-21 | 14 |
| k141_96281_flag0_multi1.0000_len377 | W1-1 | 377 | AIG51690.1 major capsid protein, partial [Koi ranavirus] | 100 | 46 | 1.03E-20 | 1 |
| k141_33055_flag1_multi117.7338_len1005 | W2-1 | 1005 | AYV88191.1 hypothetical protein [Mandarin fish ranavirus] | 85.2 | 54 | 1.06E-20 | 696 |
| k141_90309_flag1_multi12.9055_len829 | W1-1 | 829 | WAK75079.1 hypothetical protein [Mandarin fish ranavirus] | 89.8 | 49 | 1.08E-20 | 28 |
| k141_53927_flag1_multi6.8400_len985 | W5-1 | 985 | WEI29006.1 putative 2-cysteine adaptor domain protein [Largemouth bass virus] | 75 | 68 | 1.08E-20 | 22 |
| k141_46035_flag1_multi1.0000_len376 | W5-1 | 376 | QJE49073.1 putative p31K protein [Largemouth bass virus] | 94.3 | 53 | 1.11E-20 | 3 |
| k141_151002_flag0_multi5.0688_len388 | W5-1 | 388 | AIG51690.1 major capsid protein, partial [Koi ranavirus] | 100 | 47 | 1.19E-20 | 5 |
| k141_99971_flag1_multi3.9418_len1327 | W5-1 | 1327 | QIZ30887.1 major capsid protein, partial [Largemouth bass virus] | 100 | 46 | 1.26E-20 | 24 |
| k141_2596_flag1_multi2.9168_len694 | W5-1 | 694 | XPZ21295.1 putative RNAseIII [Mandarin fish ranavirus] | 97.9 | 48 | 1.35E-20 | 17 |
| k141_156269_flag1_multi1.0000_len775 | W5-1 | 775 | QJE49125.1 hypothetical protein LMBV_062 [Largemouth bass virus] | 100 | 46 | 1.39E-20 | 15 |
| k141_132570_flag1_multi3.9605_len748 | W5-1 | 748 | WEI29006.1 putative 2-cysteine adaptor domain protein [Largemouth bass virus] | 68.4 | 76 | 1.41E-20 | 5 |
| k141_142490_flag1_multi3.5877_len745 | W5-1 | 745 | XRL22821.1 hypothetical protein [Siniperca chuatsi ranavirus] | 88.7 | 53 | 1.44E-20 | 13 |
| k141_30513_flag1_multi2.8547_len733 | W5-1 | 733 | WAK75072.1 hypothetical protein [Mandarin fish ranavirus] | 88.7 | 53 | 1.45E-20 | 5 |

| | | | | | | | |
|---|---|---|---|---|---|---|---|
| k141_58995_flag0_multi1.0000_len279 | W1-1 | 279 | XRL22790.1 hypothetical protein [Siniperca chuatsi ranavirus] | 95.8 | 48 | 1.50E-20 | 0 |
| k141_99155_flag1_multi1.0000_len418 | W1-1 | 418 | UUY86197.1 hypothetical protein [Largemouth bass virus] | 100 | 47 | 1.51E-20 | 7 |
| k141_110732_flag1_multi1.0000_len273 | W5-1 | 273 | QJE49128.1 hypothetical protein LMBV_065 [Largemouth bass virus] | 97.7 | 43 | 1.54E-20 | 0 |
| k141_136363_flag1_multi5.8358_len476 | W5-1 | 476 | AFD96401.1 major capsid protein, partial [Largemouth bass virus] | 100 | 43 | 1.55E-20 | 5 |
| k141_70833_flag0_multi1.0000_len278 | W5-1 | 278 | AIG51690.1 major capsid protein, partial [Koi ranavirus] | 100 | 46 | 1.55E-20 | 0 |
| k141_6194_flag1_multi3.8720_len680 | W1-1 | 680 | UUY86241.1 hypothetical protein [Largemouth bass virus] | 88.9 | 45 | 1.57E-20 | 7 |
| k141_62346_flag1_multi18.7124_len1518 | W4-1 | 1518 | AFD96401.1 major capsid protein, partial [Largemouth bass virus] | 100 | 47 | 1.64E-20 | 74 |
| k141_147424_flag1_multi3.7753_len715 | W5-1 | 715 | UUY86192.1 putative myristylated membrane protein [Largemouth bass virus] | 100 | 46 | 1.64E-20 | 14 |
| k141_118114_flag1_multi17.9084_len4411 | W5-1 | 4411 | QIZ30887.1 major capsid protein, partial [Largemouth bass virus] | 94 | 50 | 1.65E-20 | 261 |
| k141_39029_flag1_multi13.7760_len632 | W6-1 | 632 | WEI29006.1 putative 2-cysteine adaptor domain protein [Largemouth bass virus] | 100 | 46 | 1.66E-20 | 28 |
| k141_111999_flag1_multi11.0000_len2684 | W2-3 | 2684 | WAK75102.1 hypothetical protein [Mandarin fish ranavirus] | 95.9 | 49 | 1.69E-20 | 109 |
| k141_67036_flag1_multi3.8222_len529 | W1-1 | 529 | UUY86267.1 ribonucleotide reductase alpha subunit [Largemouth bass virus] | 94 | 50 | 1.72E-20 | 3 |
| k141_105676_flag1_multi14.5340_len832 | W2-1 | 832 | XRB52842.1 transcription factor TFIIB cyclin-like domain-containing protein, partial [Largemouth bass virus] | 100 | 46 | 1.76E-20 | 27 |
| k141_29002_flag1_multi3.4061_len993 | W5-1 | 993 | QJE49077.1 hypothetical protein LMBV_014 [Largemouth bass virus] | 97.7 | 43 | 1.76E-20 | 8 |
| k141_96093_flag1_multi1.0000_len367 | W2-1 | 367 | AIG51690.1 major capsid protein, partial [Koi ranavirus] | 100 | 46 | 1.78E-20 | 3.55 |
| k141_32100_flag1_multi1.9076_len390 | W6-1 | 390 | QJE49080.1 hypothetical protein LMBV_017 [Largemouth bass virus] | 97.8 | 46 | 1.80E-20 | 0 |
| k141_68596_flag1_multi3.8969_len810 | W1-1 | 810 | AYV88159.1 hypothetical protein [Mandarin fish ranavirus] | 100 | 46 | 1.82E-20 | 12 |
| k141_52126_flag1_multi3.6632_len622 | W5-1 | 622 | WEI28972.1 CTD-phosphotransferase [Largemouth bass virus] | 97.9 | 47 | 1.83E-20 | 5 |
| k141_66197_flag0_multi2.8079_len370 | W5-1 | 370 | AIG51690.1 major capsid protein, partial [Koi ranavirus] | 85 | 60 | 1.85E-20 | 2 |
| k141_51217_flag1_multi3.9590_len751 | W5-1 | 751 | WXI69525.1 caspase recruitment domain protein [Largemouth bass virus] | 100 | 46 | 1.90E-20 | 10 |
| k141_155458_flag1_multi1.8610_len472 | W5-1 | 472 | UUY86227.1 hypothetical protein [Largemouth bass virus] | 95.7 | 46 | 2.02E-20 | 0 |
| k141_57874_flag1_multi9.4887_len896 | W2-1 | 896 | WEI28972.1 CTD-phosphotransferase [Largemouth bass virus] | 97.9 | 47 | 2.05E-20 | 31 |
| k141_132215_flag0_multi14.0512_len395 | W1-1 | 395 | WEI28972.1 CTD-phosphotransferase [Largemouth bass virus] | 100 | 46 | 2.07E-20 | 1 |
| k141_114623_flag1_multi4.8533_len741 | W5-1 | 741 | QJE49104.1 hypothetical protein LMBV_041 [Largemouth bass virus] | 100 | 46 | 2.17E-20 | 9 |
| k141_72770_flag0_multi1.7993_len684 | W5-1 | 684 | ALR73092.1 major capsid protein, partial [Box turtle ranavirus] | 93.8 | 48 | 2.19E-20 | 9 |
| k141_72610_flag1_multi18.0000_len784 | W5-1 | 784 | WAK75102.1 hypothetical protein [Mandarin fish ranavirus] | 90.2 | 51 | 2.22E-20 | 99 |
| k141_19950_flag1_multi4.9588_len1014 | W5-1 | 1014 | UUY86199.1 hypothetical protein [Largemouth bass virus] | 83 | 53 | 2.35E-20 | 16 |
| k141_168185_flag1_multi5.5361_len764 | W5-1 | 764 | AYV88120.1 putative p31K protein [Mandarin fish ranavirus] | 97.8 | 46 | 2.37E-20 | 18 |
| k141_111838_flag1_multi1.8891_len745 | W1-1 | 745 | QJE49065.1 hypothetical protein LMBV_002 [Largemouth bass virus] | 100 | 43 | 2.50E-20 | 6 |
| k141_46050_flag1_multi17.0907_len1685 | W1-1 | 1685 | QIZ30887.1 major capsid protein, partial [Largemouth bass virus] | 100 | 46 | 2.55E-20 | 115 |
| k141_66267_flag1_multi4.8850_len1080 | W5-1 | 1080 | WXI69541.1 hypothetical protein [Largemouth bass virus] | 96 | 50 | 2.56E-20 | 26 |
| k141_19826_flag1_multi16.3984_len1233 | W6-1 | 1233 | AIZ70320.1 major capsid protein, partial [Chinese giant salamander virus] | 78.3 | 60 | 2.57E-20 | 33 |
| k141_64511_flag0_multi2.7427_len518 | W4-1 | 518 | WXI69548.1 hypothetical protein [Largemouth bass virus] | 100 | 45 | 2.59E-20 | 8 |
| k141_61248_flag1_multi2.5803_len851 | W5-1 | 851 | AYV88134.2 putative tyrosine kinase [Mandarin fish ranavirus] | 95.9 | 49 | 2.64E-20 | 27 |
| k141_127332_flag1_multi3.0000_len661 | W3-2 | 661 | AYV88172.2 hypothetical protein [Mandarin fish ranavirus] | 91.5 | 164 | 2.69E-20 | 4 |
| k141_104333_flag0_multi1.8889_len546 | W1-1 | 546 | AYV88159.1 hypothetical protein [Mandarin fish ranavirus] | 100 | 46 | 2.83E-20 | 11 |
| k141_47138_flag0_multi1.0000_len332 | W4-1 | 332 | AYV88191.1 hypothetical protein [Mandarin fish ranavirus] | 83.7 | 49 | 2.85E-20 | 0 |
| k141_47459_flag1_multi2.8817_len530 | W5-1 | 530 | QJE49137.1 hypothetical protein LMBV_074 [Largemouth bass virus] | 92.2 | 51 | 2.86E-20 | 5 |
| k141_41892_flag0_multi1.0000_len329 | W5-1 | 329 | AIG51690.1 major capsid protein, partial [Koi ranavirus] | 93.6 | 47 | 2.97E-20 | 0 |

| | | | | | | | |
|---|---|---|---|---|---|---|---|
| k141_16284_flag1_multi1.5706_len665 | W5-1 | 665 | UUY86235.1 hypothetical protein [Largemouth bass virus] | 100 | 46 | 3.05E-20 | 4 |
| k141_53993_flag1_multi1.0000_len376 | W5-1 | 376 | AYV88134.2 putative tyrosine kinase [Mandarin fish ranavirus] | 100 | 46 | 3.05E-20 | 1 |
| k141_85277_flag1_multi3.6469_len444 | W1-1 | 444 | AIG51690.1 major capsid protein, partial [Koi ranavirus] | 83.6 | 61 | 3.07E-20 | 1 |
| k141_163861_flag0_multi1.0886_len299 | W5-1 | 299 | QJE49215.1 putative orf58-like protein [Largemouth bass virus] | 100 | 46 | 3.08E-20 | 0 |
| k141_38384_flag1_multi4.5783_len767 | W5-1 | 767 | WEI29006.1 putative 2-cysteine adaptor domain protein [Largemouth bass virus] | 100 | 47 | 3.12E-20 | 6 |
| k141_6224_flag1_multi4.5440_len1106 | W4-1 | 1106 | WHA35503.1 putative myristylated membrane protein [Micropterus salmoides ranavirus] | 100 | 44 | 3.26E-20 | 24 |
| k141_27324_flag1_multi2.8847_len540 | W5-1 | 540 | QJE49137.1 hypothetical protein LMBV_074 [Largemouth bass virus] | 94 | 50 | 3.26E-20 | 4 |
| k141_145378_flag1_multi6.0000_len669 | W5-1 | 669 | AYV88167.1 hypothetical protein [Mandarin fish ranavirus] | 92.9 | 42 | 3.34E-20 | 11 |
| k141_153164_flag0_multi28.6273_len1869 | W5-1 | 1869 | WAK75073.1 hypothetical protein [Mandarin fish ranavirus] | 100 | 46 | 3.48E-20 | 119 |
| k141_21227_flag0_multi1.0000_len256 | W4-1 | 256 | QJE49077.1 hypothetical protein LMBV_014 [Largemouth bass virus] | 95 | 40 | 3.49E-20 | 0 |
| k141_146856_flag1_multi1.8767_len514 | W5-1 | 514 | AYV88120.1 putative p31K protein [Mandarin fish ranavirus] | 95.7 | 46 | 3.58E-20 | 1 |
| k141_28109_flag1_multi3.8990_len824 | W5-1 | 824 | AYV88191.1 hypothetical protein [Mandarin fish ranavirus] | 97.9 | 47 | 3.59E-20 | 8 |
| k141_97011_flag1_multi1.6868_len406 | W1-1 | 406 | AIG51690.1 major capsid protein, partial [Koi ranavirus] | 90.6 | 53 | 3.79E-20 | 1 |
| k141_146799_flag1_multi3.6465_len1049 | W2-3 | 1049 | AYV88136.2 hypothetical protein [Mandarin fish ranavirus] | 97.7 | 43 | 3.92E-20 | 14 |
| k141_112715_flag1_multi18.7039_len749 | W1-1 | 749 | QYU76034.1 putative myristylated membrane protein, partial [Koi ranavirus] | 100 | 46 | 3.96E-20 | 9 |
| k141_84634_flag0_multi171.7544_len1912 | W4-1 | 1912 | UUY86235.1 hypothetical protein [Largemouth bass virus] | 95.9 | 49 | 3.98E-20 | 1679 |
| k141_85586_flag1_multi11.9167_len561 | W4-1 | 561 | AIG51690.1 major capsid protein, partial [Koi ranavirus] | 92.2 | 51 | 3.99E-20 | 15 |
| k141_68589_flag1_multi1.9696_len503 | W5-1 | 503 | UUY86199.1 hypothetical protein [Largemouth bass virus] | 100 | 41 | 4.05E-20 | 3 |
| k141_22074_flag1_multi2.8509_len530 | W4-1 | 530 | AIG51690.1 major capsid protein, partial [Koi ranavirus] | 94.2 | 52 | 4.08E-20 | 4 |
| k141_145656_flag1_multi1.0000_len423 | W5-1 | 423 | AYV88191.1 hypothetical protein [Mandarin fish ranavirus] | 100 | 44 | 4.08E-20 | 0 |
| k141_72834_flag0_multi19.2727_len471 | W5-1 | 471 | AYV88142.2 hypothetical protein [Mandarin fish ranavirus] | 100 | 47 | 4.20E-20 | 0 |
| k141_3212_flag0_multi5.0947_len648 | W5-1 | 648 | QYU76034.1 putative myristylated membrane protein, partial [Koi ranavirus] | 89.6 | 48 | 4.33E-20 | 4 |
| k141_77542_flag1_multi1.0000_len568 | W1-1 | 568 | WEI28968.1 putative DNA dependent RNA polymerase a subunit [Largemouth bass virus] | 97.8 | 45 | 4.38E-20 | 8 |
| k141_6840_flag1_multi1.0000_len524 | W5-1 | 524 | UUY86235.1 hypothetical protein [Largemouth bass virus] | 100 | 46 | 4.38E-20 | 2 |
| k141_26388_flag0_multi4.9425_len1672 | W3-2 | 1672 | QJE49112.1 hypothetical protein LMBV_049 [Largemouth bass virus] | 100 | 46 | 4.72E-20 | 26 |
| k141_14187_flag0_multi1.0000_len278 | W4-1 | 278 | AYV88120.1 putative p31K protein [Mandarin fish ranavirus] | 100 | 46 | 4.82E-20 | 0 |
| k141_8124_flag0_multi3.6359_len1278 | W5-1 | 1278 | AYV88191.1 hypothetical protein [Mandarin fish ranavirus] | 100 | 45 | 4.84E-20 | 29 |
| k141_32091_flag1_multi10.4548_len561 | W4-1 | 561 | XRB52768.1 P31K protein, partial [Largemouth bass virus] | 97.8 | 46 | 4.92E-20 | 15 |
| k141_149795_flag0_multi1.0000_len322 | W5-1 | 322 | AIG51690.1 major capsid protein, partial [Koi ranavirus] | 98 | 50 | 5.31E-20 | 3.62 |
| k141_61217_flag1_multi1.0000_len280 | W5-1 | 280 | QJE49066.1 hypothetical protein LMBV_003 [Largemouth bass virus] | 100 | 46 | 5.58E-20 | 3 |
| k141_61217_flag1_multi1.0000_len280 | W5-1 | 280 | QJE49066.1 hypothetical protein LMBV_003 [Largemouth bass virus] | 100 | 46 | 5.58E-20 | 0 |
| k141_77977_flag0_multi1.0000_len281 | W5-1 | 281 | UVF58793.1 MAG: DNA polymerase [Halichoeres melanurus ranavirus] | 100 | 46 | 5.75E-20 | 0 |
| k141_88604_flag1_multi2.9624_len460 | W5-1 | 460 | UUY86241.1 hypothetical protein [Largemouth bass virus] | 100 | 38 | 5.77E-20 | 7 |
| k141_12550_flag1_multi1.0000_len537 | W3-1 | 537 | QJE49144.1 hypothetical protein LMBV_081 [Largemouth bass virus] | 100 | 46 | 6.05E-20 | 2 |
| k141_4045_flag1_multi4.8870_len495 | W4-1 | 495 | AYV88191.1 hypothetical protein [Mandarin fish ranavirus] | 95.2 | 42 | 6.27E-20 | 5 |
| k141_22321_flag0_multi84.6432_len1049 | W1-1 | 1049 | AYV88125.2 hypothetical protein [Mandarin fish ranavirus] | 93.5 | 62 | 6.33E-20 | 283 |
| k141_101051_flag1_multi1.0000_len331 | W4-1 | 331 | WEI28972.1 CTD-phosphotransferase [Largemouth bass virus] | 100 | 47 | 6.46E-20 | 0 |
| k141_153239_flag1_multi7.2500_len477 | W5-1 | 477 | UUY86241.1 hypothetical protein [Largemouth bass virus] | 97.4 | 38 | 6.77E-20 | 11 |

| | | | | | | | |
|---|---|---|---|---|---|---|---|
| k141_109571_flag1_multi3.0000_len342 | W3-2 | 342 | UUY86238.1 hypothetical protein [Largemouth bass virus] | 97.7 | 43 | 7.08E-20 | 2 |
| k141_114721_flag1_multi7.8000_len946 | W5-1 | 946 | WEI28972.1 CTD-phosphotransferase [Largemouth bass virus] | 100 | 46 | 7.21E-20 | 21 |
| k141_95128_flag1_multi4.6421_len711 | W1-1 | 711 | QJE49153.1 hypothetical protein LMBV_004 [Largemouth bass virus] | 73.4 | 64 | 7.22E-20 | 10 |
| k141_72402_flag1_multi1.6253_len560 | W4-1 | 560 | AYV88179.2 putative DNA dependent RNA polymerase II second largest subunit [Mandarin fish ranavirus] | 93.8 | 48 | 7.25E-20 | 3 |
| k141_121872_flag1_multi4.0775_len683 | W1-1 | 683 | XRB52842.1 transcription factor TFIIB cyclin-like domain-containing protein, partial [Largemouth bass virus] | 100 | 41 | 7.70E-20 | 9 |
| k141_72571_flag1_multi3.8874_len754 | W4-1 | 754 | WAK75107.1 hypothetical protein [Mandarin fish ranavirus] | 100 | 41 | 7.86E-20 | 8 |
| k141_82548_flag1_multi1.0000_len361 | W5-1 | 361 | UUY86235.1 hypothetical protein [Largemouth bass virus] | 100 | 46 | 8.02E-20 | 11 |
| k141_46066_flag1_multi2.9345_len843 | W5-1 | 843 | AYV88191.1 hypothetical protein [Mandarin fish ranavirus] | 97.8 | 46 | 8.15E-20 | 5 |
| k141_87014_flag1_multi2.1534_len519 | W2-3 | 519 | WEI28972.1 CTD-phosphotransferase [Largemouth bass virus] | 92.2 | 51 | 8.19E-20 | 4 |
| k141_97628_flag1_multi2.9017_len609 | W5-1 | 609 | WEI29006.1 putative 2-cysteine adaptor domain protein [Largemouth bass virus] | 100 | 46 | 8.30E-20 | 8 |
| k141_11997_flag0_multi3.6662_len797 | W5-1 | 797 | WEI28972.1 CTD-phosphotransferase [Largemouth bass virus] | 100 | 47 | 8.60E-20 | 5 |
| k141_90772_flag1_multi2.9218_len729 | W5-1 | 729 | QYU76034.1 putative myristylated membrane protein, partial [Koi ranavirus] | 86 | 57 | 8.62E-20 | 6 |
| k141_57634_flag1_multi4.8938_len970 | W5-1 | 970 | WHU98721.1 MCP, partial [Hybrid snakehead ranavirus] | 92 | 50 | 8.88E-20 | 860 |
| k141_99174_flag1_multi3.8028_len430 | W5-1 | 430 | AYV88159.1 hypothetical protein [Mandarin fish ranavirus] | 100 | 44 | 8.96E-20 | 2 |
| k141_125873_flag1_multi1.0000_len269 | W1-1 | 269 | WAK75103.1 hypothetical protein [Mandarin fish ranavirus] | 100 | 43 | 8.98E-20 | 0 |
| k141_166669_flag1_multi17.8368_len2059 | W5-1 | 2059 | UUY86192.1 putative myristylated membrane protein [Largemouth bass virus] | 100 | 47 | 9.30E-20 | 123 |
| k141_133567_flag1_multi9.6956_len821 | W5-1 | 821 | QYU76034.1 putative myristylated membrane protein, partial [Koi ranavirus] | 100 | 46 | 9.61E-20 | 24 |
| k141_94028_flag0_multi1.0000_len376 | W5-1 | 376 | WAK75136.1 hypothetical protein [Mandarin fish ranavirus] | 100 | 45 | 9.66E-20 | 5 |
| k141_43808_flag0_multi2.8537_len729 | W4-1 | 729 | AIG51690.1 major capsid protein, partial [Koi ranavirus] | 97.9 | 48 | 9.73E-20 | 2 |
| k141_23456_flag1_multi10.1534_len956 | W4-1 | 956 | UUY86192.1 putative myristylated membrane protein [Largemouth bass virus] | 100 | 46 | 9.86E-20 | 34 |
| k141_98852_flag1_multi11.6598_len2031 | W5-1 | 2031 | QJE49068.1 hypothetical protein LMBV_005 [Largemouth bass virus] | 97.9 | 47 | 9.88E-20 | 114.79 |
| k141_69565_flag0_multi7.9790_len1473 | W5-1 | 1473 | AFD96401.1 major capsid protein, partial [Largemouth bass virus] | 84.2 | 57 | 9.98E-20 | 52 |
| k141_26840_flag0_multi2.8693_len493 | W1-1 | 493 | AIG51690.1 major capsid protein, partial [Koi ranavirus] | 100 | 46 | 1.00E-19 | 6 |
| k141_34169_flag1_multi15.7945_len1820 | W1-1 | 1820 | QJE49112.1 hypothetical protein LMBV_049 [Largemouth bass virus] | 100 | 44 | 1.04E-19 | 78 |
| k141_57070_flag1_multi1.9395_len521 | W1-1 | 521 | AYV88211.1 hypothetical protein [Mandarin fish ranavirus] | 94 | 50 | 1.06E-19 | 1 |
| k141_76851_flag1_multi4.2332_len917 | W5-1 | 917 | XRB52797.1 Bcl-2 Bcl-2 homology region 1-3 domain-containing protein [Largemouth bass virus] | 97.9 | 47 | 1.12E-19 | 13 |
| k141_20970_flag1_multi1.8833_len501 | W1-1 | 501 | UUY86192.1 putative myristylated membrane protein [Largemouth bass virus] | 65.8 | 73 | 1.14E-19 | 7 |
| k141_109331_flag0_multi5.3570_len844 | W1-3 | 844 | XRL22821.1 hypothetical protein [Siniperca chuatsi ranavirus] | 100 | 46 | 1.14E-19 | 27 |
| k141_102609_flag1_multi2.9006_len604 | W4-1 | 604 | AIG51690.1 major capsid protein, partial [Koi ranavirus] | 100 | 46 | 1.15E-19 | 6 |
| k141_122495_flag1_multi1.0000_len280 | W5-1 | 280 | UUY86192.1 putative myristylated membrane protein [Largemouth bass virus] | 100 | 42 | 1.15E-19 | 0 |
| k141_147265_flag1_multi3.8565_len580 | W5-1 | 580 | QJE49104.1 hypothetical protein LMBV_041 [Largemouth bass virus] | 94 | 50 | 1.16E-19 | 5 |
| k141_164061_flag1_multi1.0000_len362 | W5-1 | 362 | AIG51690.1 major capsid protein, partial [Koi ranavirus] | 100 | 45 | 1.19E-19 | 6 |
| k141_94398_flag1_multi2.0000_len433 | W5-1 | 433 | UUY86241.1 hypothetical protein [Largemouth bass virus] | 92.7 | 41 | 1.20E-19 | 3 |
| k141_141168_flag1_multi18.8806_len3600 | W5-1 | 3600 | WEI28974.1 hypothetical protein [Largemouth bass virus] | 100 | 46 | 1.21E-19 | 244 |
| k141_106369_flag0_multi83.0000_len659 | W5-3 | 659 | XRB52842.1 transcription factor TFIIB cyclin-like domain-containing protein, partial [Largemouth bass virus] | 76.8 | 56 | 1.22E-19 | 213 |
| k141_6278_flag1_multi6.6231_len937 | W5-1 | 937 | QJE49077.1 hypothetical protein LMBV_014 [Largemouth bass virus] | 91.3 | 46 | 1.26E-19 | 32 |

| | | | | | | | |
|---|---|---|---|---|---|---|---|
| k141_26540_flag1_multi2.9548_len539 | W5-1 | 539 | AYV88120.1 putative p31K protein [Mandarin fish ranavirus] | 97.9 | 47 | 1.26E-19 | 5 |
| k141_60266_flag0_multi1.0000_len313 | W1-1 | 313 | AIG51690.1 major capsid protein, partial [Koi ranavirus] | 100 | 46 | 1.27E-19 | 90.65 |
| k141_105001_flag0_multi1.8284_len1586 | W5-1 | 1586 | WAK75136.1 hypothetical protein [Mandarin fish ranavirus] | 71.4 | 70 | 1.29E-19 | 40 |
| k141_99433_flag1_multi1.0000_len278 | W4-1 | 278 | AYV88211.1 hypothetical protein [Mandarin fish ranavirus] | 93.9 | 49 | 1.35E-19 | 0 |
| k141_95371_flag0_multi2.8845_len470 | W1-1 | 470 | AYV88159.1 hypothetical protein [Mandarin fish ranavirus] | 100 | 44 | 1.37E-19 | 0 |
| k141_69694_flag1_multi1.0000_len524 | W5-1 | 524 | UUY86258.1 hypothetical protein [Largemouth bass virus] | 100 | 47 | 1.44E-19 | 22 |
| k141_25061_flag1_multi3.8964_len807 | W5-1 | 807 | XRB52797.1 Bcl-2 Bcl-2 homology region 1-3 domain-containing protein [Largemouth bass virus] | 100 | 46 | 1.50E-19 | 8 |
| k141_141979_flag1_multi4.4437_len434 | W5-1 | 434 | QJE49104.1 hypothetical protein LMBV_041 [Largemouth bass virus] | 100 | 44 | 1.51E-19 | 26.63 |
| k141_74965_flag1_multi22.7678_len632 | W4-1 | 632 | QJE49137.1 hypothetical protein LMBV_074 [Largemouth bass virus] | 100 | 46 | 1.52E-19 | 65 |
| k141_88153_flag1_multi5.7709_len643 | W4-1 | 643 | XRB52768.1 P31K protein, partial [Largemouth bass virus] | 88.2 | 51 | 1.52E-19 | 8 |
| k141_60637_flag0_multi1.8593_len603 | W5-1 | 603 | AYV88120.1 putative p31K protein [Mandarin fish ranavirus] | 97.8 | 46 | 1.65E-19 | 3 |
| k141_31688_flag1_multi10.2433_len1004 | W4-1 | 1004 | AYV88120.1 putative p31K protein [Mandarin fish ranavirus] | 100 | 46 | 1.66E-19 | 34 |
| k141_89573_flag1_multi1.0000_len338 | W5-1 | 338 | AIG51690.1 major capsid protein, partial [Koi ranavirus] | 75.4 | 65 | 1.71E-19 | 0 |
| k141_49187_flag1_multi12.7811_len1402 | W1-1 | 1402 | AYV88191.1 hypothetical protein [Mandarin fish ranavirus] | 97.8 | 46 | 1.79E-19 | 50 |
| k141_53648_flag1_multi2.4714_len665 | W1-1 | 665 | QJE49215.1 putative orf58-like protein [Largemouth bass virus] | 92 | 50 | 1.80E-19 | 5 |
| k141_24489_flag1_multi12.4671_len567 | W4-1 | 567 | WEI28972.1 CTD-phosphotransferase [Largemouth bass virus] | 97.9 | 47 | 1.84E-19 | 29 |
| k141_62140_flag1_multi1.0000_len291 | W4-1 | 291 | AIG51690.1 major capsid protein, partial [Koi ranavirus] | 100 | 46 | 1.87E-19 | 61 |
| k141_39213_flag1_multi1.0000_len706 | W5-1 | 706 | UUY86221.1 hypothetical protein [Largemouth bass virus] | 100 | 46 | 1.87E-19 | 14 |
| k141_149313_flag1_multi2.5503_len817 | W5-1 | 817 | AYV88120.1 putative p31K protein [Mandarin fish ranavirus] | 97.8 | 46 | 1.87E-19 | 5 |
| k141_58220_flag1_multi2.8277_len408 | W5-1 | 408 | QJE49066.1 hypothetical protein LMBV_003 [Largemouth bass virus] | 84.5 | 58 | 1.89E-19 | 2 |
| k141_34050_flag1_multi3.3835_len1356 | W5-1 | 1356 | QYU76034.1 putative myristylated membrane protein, partial [Koi ranavirus] | 94.1 | 51 | 1.89E-19 | 11 |
| k141_37590_flag1_multi1.9347_len493 | W5-1 | 493 | AIG51690.1 major capsid protein, partial [Koi ranavirus] | 97.9 | 47 | 1.98E-19 | 2 |
| k141_79157_flag1_multi9.8745_len1790 | W1-1 | 1790 | AIZ70320.1 major capsid protein, partial [Chinese giant salamander virus] | 88.5 | 52 | 2.01E-19 | 68 |
| k141_91430_flag1_multi4.4578_len1172 | W5-1 | 1172 | WEI29002.1 hypothetical protein [Largemouth bass virus] | 100 | 45 | 2.07E-19 | 11 |
| k141_28965_flag1_multi4.4006_len463 | W4-1 | 463 | AYV88134.2 putative tyrosine kinase [Mandarin fish ranavirus] | 100 | 44 | 2.10E-19 | 3 |
| k141_54160_flag0_multi3.2873_len503 | W1-1 | 503 | AIG51690.1 major capsid protein, partial [Koi ranavirus] | 92.2 | 51 | 2.18E-19 | 98.35 |
| k141_165279_flag1_multi1.0000_len545 | W5-1 | 545 | QJE49065.1 hypothetical protein LMBV_002 [Largemouth bass virus] | 100 | 42 | 2.19E-19 | 6 |
| k141_101867_flag1_multi1.0000_len922 | W5-1 | 922 | WAK75114.1 hypothetical protein [Mandarin fish ranavirus] | 95.6 | 45 | 2.19E-19 | 31 |
| k141_82389_flag1_multi3.8597_len1445 | W5-1 | 1445 | QIZ30887.1 major capsid protein, partial [Largemouth bass virus] | 64.6 | 82 | 2.20E-19 | 25 |
| k141_93369_flag1_multi4.7500_len349 | W4-1 | 349 | AYV88120.1 putative p31K protein [Mandarin fish ranavirus] | 97.7 | 43 | 2.25E-19 | 5 |
| k141_150118_flag1_multi4.6893_len553 | W5-1 | 553 | UUY86194.1 hypothetical protein [Largemouth bass virus] | 97.9 | 47 | 2.53E-19 | 7 |
| k141_63554_flag1_multi6.3196_len773 | W5-1 | 773 | XPZ21261.1 hypothetical protein MRVORF002 [Mandarin fish ranavirus] | 97.8 | 46 | 2.63E-19 | 13 |
| k141_153331_flag0_multi9.8537_len2909 | W5-1 | 2909 | QJE49109.1 hypothetical protein LMBV_046 [Largemouth bass virus] | 100 | 46 | 2.66E-19 | 86 |
| k141_14507_flag1_multi1.0000_len277 | W2-1 | 277 | QJE49073.1 putative p31K protein [Largemouth bass virus] | 95.9 | 49 | 2.67E-19 | 4 |
| k141_159630_flag1_multi2.9374_len876 | W5-1 | 876 | AYV88179.2 putative DNA dependent RNA polymerase II second largest subunit [Mandarin fish ranavirus] | 100 | 46 | 2.73E-19 | 5 |
| k141_77097_flag0_multi1.0000_len270 | W4-1 | 270 | AIG51690.1 major capsid protein, partial [Koi ranavirus] | 100 | 43 | 2.75E-19 | 2 |
| k141_34373_flag1_multi1.9404_len527 | W5-1 | 527 | AIG51690.1 major capsid protein, partial [Koi ranavirus] | 100 | 47 | 2.80E-19 | 1 |
| k141_2511_flag1_multi3.8240_len533 | W5-1 | 533 | XRL22790.1 hypothetical protein [Siniperca chuatsi ranavirus] | 100 | 46 | 2.80E-19 | 5 |

| | | | | | | | |
|---|---|---|---|---|---|---|---|
| k141_21512_flag1_multi5.2164_len1033 | W4-1 | 1033 | QJE49090.1 hypothetical protein LMBV_027 [Largemouth bass virus] | 100 | 46 | 2.88E-19 | 15 |
| k141_13430_flag1_multi3.8282_len892 | W6-1 | 892 | QJE49114.1 hypothetical protein LMBV_051 [Largemouth bass virus] | 80.4 | 51 | 2.89E-19 | 27 |
| k141_89907_flag1_multi1.7493_len907 | W5-1 | 907 | XRL22790.1 hypothetical protein [Siniperca chuatsi ranavirus] | 83.9 | 56 | 2.96E-19 | 10 |
| k141_25877_flag1_multi1.0000_len293 | W6-1 | 293 | AYV88118.1 hypothetical protein [Mandarin fish ranavirus] | 100 | 37 | 3.01E-19 | 0 |
| k141_52117_flag1_multi3.8288_len544 | W4-1 | 544 | XRL22790.1 hypothetical protein [Siniperca chuatsi ranavirus] | 100 | 46 | 3.06E-19 | 9 |
| k141_2085_flag1_multi5.2237_len1035 | W5-1 | 1035 | AIG51690.1 major capsid protein, partial [Koi ranavirus] | 100 | 46 | 3.08E-19 | 20 |
| k141_109322_flag1_multi1.0000_len415 | W5-1 | 415 | AIG51690.1 major capsid protein, partial [Koi ranavirus] | 97.9 | 48 | 3.10E-19 | 5 |
| k141_13981_flag0_multi11.7067_len6141 | W1-1 | 6141 | XRL22821.1 hypothetical protein [Siniperca chuatsi ranavirus] | 90.4 | 52 | 3.14E-19 | 186 |
| k141_98498_flag1_multi1.0000_len573 | W5-1 | 573 | AIG51690.1 major capsid protein, partial [Koi ranavirus] | 100 | 45 | 3.17E-19 | 8 |
| k141_103397_flag1_multi5.8766_len1073 | W1-1 | 1073 | WAK75102.1 hypothetical protein [Mandarin fish ranavirus] | 93.8 | 48 | 3.31E-19 | 22 |
| k141_71435_flag1_multi15.8900_len3277 | W5-1 | 3277 | AYV88191.1 hypothetical protein [Mandarin fish ranavirus] | 90.4 | 52 | 3.41E-19 | 202 |
| k141_43786_flag1_multi1.0000_len280 | W1-1 | 280 | UUY86197.1 hypothetical protein [Largemouth bass virus] | 94 | 50 | 3.56E-19 | 0 |
| k141_120635_flag0_multi1.0000_len864 | W1-1 | 864 | AYV88120.1 putative p31K protein [Mandarin fish ranavirus] | 97.8 | 46 | 3.57E-19 | 23.88 |
| k141_35420_flag1_multi5.7876_len447 | W4-1 | 447 | AYV88120.1 putative p31K protein [Mandarin fish ranavirus] | 97.7 | 43 | 3.57E-19 | 11 |
| k141_155697_flag1_multi8.3645_len1864 | W5-1 | 1864 | QJE49125.1 hypothetical protein LMBV_062 [Largemouth bass virus] | 97.9 | 48 | 3.68E-19 | 45 |
| k141_92482_flag1_multi2.8340_len659 | W5-1 | 659 | AIG51690.1 major capsid protein, partial [Koi ranavirus] | 100 | 46 | 3.74E-19 | 4 |
| k141_96019_flag1_multi2.5806_len780 | W5-1 | 780 | QJE49066.1 hypothetical protein LMBV_003 [Largemouth bass virus] | 88.7 | 53 | 3.76E-19 | 9 |
| k141_58425_flag1_multi2.8540_len456 | W5-1 | 456 | AAF64582.1 capsid protein, partial [Guppyfish iridovirus] | 95.7 | 46 | 3.78E-19 | 0 |
| k141_16766_flag1_multi11.6241_len814 | W1-1 | 814 | QJE49149.1 putative immediate early protein ICP-46 [Largemouth bass virus] | 100 | 46 | 3.91E-19 | 32 |
| k141_132021_flag1_multi2.8919_len363 | W1-1 | 363 | AYV88120.1 putative p31K protein [Mandarin fish ranavirus] | 97.7 | 43 | 3.91E-19 | 4 |
| k141_158552_flag1_multi15.5898_len982 | W5-1 | 982 | WEI29006.1 putative 2-cysteine adaptor domain protein [Largemouth bass virus] | 100 | 47 | 3.95E-19 | 43 |
| k141_29073_flag1_multi2.9394_len405 | W5-1 | 405 | XRB52780.1 hypothetical protein LMBV_23 [Largemouth bass virus] | 100 | 41 | 3.97E-19 | 6 |
| k141_147941_flag1_multi2.8241_len357 | W5-1 | 357 | AIG51690.1 major capsid protein, partial [Koi ranavirus] | 100 | 45 | 4.10E-19 | 3 |
| k141_127257_flag1_multi1.0000_len279 | W1-1 | 279 | AIG51690.1 major capsid protein, partial [Koi ranavirus] | 92 | 50 | 4.13E-19 | 0 |
| k141_76451_flag1_multi2.5671_len931 | W4-1 | 931 | WEI28972.1 CTD-phosphotransferase [Largemouth bass virus] | 100 | 46 | 4.23E-19 | 19 |
| k141_34581_flag1_multi1.0000_len469 | W4-1 | 469 | QJE49104.1 hypothetical protein LMBV_041 [Largemouth bass virus] | 100 | 45 | 4.30E-19 | 96 |
| k141_92918_flag1_multi2.8198_len474 | W5-1 | 474 | UUY86221.1 hypothetical protein [Largemouth bass virus] | 100 | 46 | 4.56E-19 | 4 |
| k141_16493_flag1_multi4.0345_len721 | W5-1 | 721 | WXI69525.1 caspase recruitment domain protein [Largemouth bass virus] | 91.8 | 49 | 4.57E-19 | 19 |
| k141_70828_flag1_multi3.9006_len835 | W4-1 | 835 | XRL22821.1 hypothetical protein [Siniperca chuatsi ranavirus] | 100 | 46 | 4.72E-19 | 12 |
| k141_72897_flag0_multi1.0000_len516 | W5-1 | 516 | AIG51690.1 major capsid protein, partial [Koi ranavirus] | 100 | 44 | 4.89E-19 | 2 |
| k141_103229_flag1_multi1.9170_len418 | W5-1 | 418 | AAF64580.1 capsid protein, partial [Frog virus 3] | 95.7 | 46 | 5.09E-19 | 3 |
| k141_59837_flag1_multi1.0000_len742 | W5-1 | 742 | QJE49103.1 hypothetical protein LMBV_040 [Largemouth bass virus] | 100 | 46 | 5.20E-19 | 11 |
| k141_138996_flag1_multi1.5255_len455 | W5-1 | 455 | WAK75112.1 putative LITAF PIG7 possible membrane associated motif in LPS-induced tumor necrosis factor alpha factor [Mandarin fish ranavirus] | 100 | 39 | 5.37E-19 | 6 |
| k141_134716_flag1_multi1.0000_len294 | W5-1 | 294 | XRL22790.1 hypothetical protein [Siniperca chuatsi ranavirus] | 79.4 | 63 | 5.46E-19 | 0 |
| k141_143795_flag1_multi1.0000_len432 | W5-1 | 432 | XRB52831.1 hypothetical protein LMBV_74 [Largemouth bass virus] | 97.8 | 46 | 5.84E-19 | 1 |
| k141_124765_flag0_multi2.9173_len673 | W5-1 | 673 | AIG51690.1 major capsid protein, partial [Koi ranavirus] | 100 | 46 | 5.89E-19 | 11 |
| k141_132619_flag0_multi2.6436_len719 | W5-1 | 719 | QJE49153.1 hypothetical protein LMBV_004 [Largemouth bass virus] | 97.8 | 46 | 5.93E-19 | 166.38 |
| k141_170_flag1_multi1.0000_len648 | W5-1 | 648 | UUY86269.1 hypothetical protein [Largemouth bass virus] | 92 | 50 | 5.99E-19 | 0 |
| k141_25141_flag0_multi1.0000_len435 | W5-1 | 435 | WAK75139.1 putative myristylated membrane protein [Mandarin fish ranavirus] | 100 | 76 | 5.99E-19 | 486.05 |

| | | | | | | | |
|---|---|---|---|---|---|---|---|
| k141_97362_flag1_multi3.5245_len488 | W2-1 | 488 | UUY86241.1 hypothetical protein [Largemouth bass virus] | 92.5 | 40 | 6.01E-19 | 5 |
| k141_9809_flag1_multi3.1190_len620 | W5-1 | 620 | UUY86261.1 hypothetical protein [Largemouth bass virus] | 97.9 | 48 | 6.05E-19 | 5 |
| k141_133587_flag0_multi3.7122_len610 | W5-2 | 610 | UUY86221.1 hypothetical protein [Largemouth bass virus] | 100 | 46 | 6.30E-19 | 6 |
| k141_9346_flag1_multi1.9207_len431 | W1-1 | 431 | UVF58785.1 MAG: major capsid protein [Halichoeres melanurus ranavirus] | 95.9 | 49 | 6.69E-19 | 9 |
| k141_164548_flag1_multi3.7335_len670 | W5-1 | 670 | UUY86235.1 hypothetical protein [Largemouth bass virus] | 100 | 46 | 6.69E-19 | 12 |
| k141_33639_flag1_multi4.2129_len822 | W1-1 | 822 | WEI28972.1 CTD-phosphotransferase [Largemouth bass virus] | 100 | 45 | 6.82E-19 | 13 |
| k141_92034_flag1_multi7.6798_len1081 | W4-1 | 1081 | AYV88159.1 hypothetical protein [Mandarin fish ranavirus] | 100 | 46 | 6.83E-19 | 45 |
| k141_53237_flag1_multi4.9085_len1147 | W1-1 | 1147 | QYU76034.1 putative myristylated membrane protein, partial [Koi ranavirus] | 100 | 46 | 6.90E-19 | 20 |
| k141_121144_flag1_multi7.3063_len820 | W5-1 | 820 | AAC79508.1 viral core protein, partial [Ictalurus melas ranavirus] | 85.1 | 47 | 6.95E-19 | 20 |
| k141_179467_flag1_multi1.0000_len369 | W2-3 | 369 | UUY86256.1 putative orf58-like protein [Largemouth bass virus] | 100 | 46 | 7.05E-19 | 0 |
| k141_114528_flag1_multi2.8545_len622 | W5-1 | 622 | UUY86258.1 hypothetical protein [Largemouth bass virus] | 97.9 | 47 | 7.21E-19 | 4 |
| k141_142218_flag1_multi1.0000_len306 | W5-1 | 306 | QJE49096.1 putative D5 family NTPase/ATPase [Largemouth bass virus] | 73.1 | 67 | 7.24E-19 | 1 |
| k141_24931_flag1_multi1.0000_len634 | W1-1 | 634 | WAK75112.1 putative LITAF PIG7 possible membrane associated motif in LPS-induced tumor necrosis factor alpha factor [Mandarin fish ranavirus] | 100 | 41 | 7.32E-19 | 10 |
| k141_116507_flag0_multi6.4962_len403 | W1-1 | 403 | XRB52842.1 transcription factor TFIIB cyclin-like domain-containing protein, partial [Largemouth bass virus] | 49.1 | 108 | 7.49E-19 | 6 |
| k141_86011_flag1_multi5.4914_len662 | W1-1 | 662 | AYV88120.1 putative p31K protein [Mandarin fish ranavirus] | 90 | 50 | 7.73E-19 | 10 |
| k141_72844_flag1_multi12.0000_len1555 | W5-1 | 1555 | AIG51690.1 major capsid protein, partial [Koi ranavirus] | 100 | 49 | 8.02E-19 | 54 |
| k141_117925_flag1_multi6.9296_len1376 | W5-1 | 1376 | QJE49137.1 hypothetical protein LMBV_074 [Largemouth bass virus] | 90.4 | 52 | 8.19E-19 | 32 |
| k141_3124_flag1_multi8.8630_len1484 | W1-1 | 1484 | XRL22821.1 hypothetical protein [Siniperca chuatsi ranavirus] | 67.9 | 81 | 8.25E-19 | 52 |
| k141_83680_flag1_multi3.9585_len1298 | W2-1 | 1298 | UVF58787.1 MAG: DNA-dependent RNA polymerase largest subunit, partial [Halichoeres melanurus ranavirus] | 93.6 | 47 | 8.46E-19 | 24 |
| k141_104085_flag1_multi1.0000_len724 | W1-1 | 724 | AFD96401.1 major capsid protein, partial [Largemouth bass virus] | 88.2 | 51 | 8.50E-19 | 26 |
| k141_96314_flag1_multi6.7441_len563 | W4-1 | 563 | AYV88120.1 putative p31K protein [Mandarin fish ranavirus] | 97.7 | 44 | 8.52E-19 | 10 |
| k141_50895_flag1_multi5.4087_len1636 | W1-1 | 1636 | AFD96401.1 major capsid protein, partial [Largemouth bass virus] | 97.8 | 46 | 8.56E-19 | 30 |
| k141_4855_flag1_multi1.0000_len433 | W1-1 | 433 | QJE49153.1 hypothetical protein LMBV_004 [Largemouth bass virus] | 95.5 | 44 | 8.61E-19 | 1 |
| k141_149589_flag1_multi2.7455_len471 | W5-1 | 471 | UUY86261.1 hypothetical protein [Largemouth bass virus] | 81.4 | 59 | 8.94E-19 | 4 |
| k141_75314_flag1_multi1.8549_len458 | W1-1 | 458 | UVF58785.1 MAG: major capsid protein [Halichoeres melanurus ranavirus] | 100 | 47 | 9.05E-19 | 0 |
| k141_81887_flag1_multi1.0000_len435 | W5-1 | 435 | AYV88211.1 hypothetical protein [Mandarin fish ranavirus] | 100 | 45 | 9.11E-19 | 1 |
| k141_25007_flag0_multi14.6680_len647 | W4-1 | 647 | AYV88120.1 putative p31K protein [Mandarin fish ranavirus] | 95.7 | 47 | 9.47E-19 | 19 |
| k141_48992_flag1_multi52.1545_len510 | W4-1 | 510 | QJE49149.1 putative immediate early protein ICP-46 [Largemouth bass virus] | 75.9 | 58 | 9.49E-19 | 87 |
| k141_137480_flag1_multi3.7738_len446 | W4-3 | 446 | QJE49215.1 putative orf58-like protein [Largemouth bass virus] | 100 | 46 | 9.70E-19 | 4 |
| k141_90326_flag1_multi3.3163_len767 | W5-1 | 767 | UUY86211.1 hypothetical protein [Largemouth bass virus] | 100 | 46 | 9.79E-19 | 6 |
| k141_62551_flag1_multi4.8009_len784 | W5-1 | 784 | AIG51690.1 major capsid protein, partial [Koi ranavirus] | 76.9 | 65 | 1.03E-18 | 14 |
| k141_69653_flag1_multi4.5915_len682 | W5-1 | 682 | WEI29006.1 putative 2-cysteine adaptor domain protein [Largemouth bass virus] | 97.8 | 46 | 1.04E-18 | 11 |
| k141_74140_flag1_multi6.2659_len991 | W6-1 | 991 | QYU76034.1 putative myristylated membrane protein, partial [Koi ranavirus] | 100 | 46 | 1.04E-18 | 19 |
| k141_39031_flag0_multi1.7386_len229 | W5-1 | 229 | QJE49105.1 hypothetical protein LMBV_042 [Largemouth bass virus] | 97.9 | 47 | 1.05E-18 | 0 |
| k141_45773_flag1_multi1.0000_len426 | W5-1 | 426 | QJE49095.1 hypothetical protein LMBV_032 [Largemouth bass virus] | 97.7 | 43 | 1.06E-18 | 2 |
| k141_133428_flag1_multi1.9567_len672 | W5-1 | 672 | AIG51690.1 major capsid protein, partial [Koi ranavirus] | 94.1 | 51 | 1.09E-18 | 2 |
| k141_58964_flag1_multi18.2499_len2810 | W5-1 | 2810 | XRL22821.1 hypothetical protein [Siniperca chuatsi ranavirus] | 90.6 | 53 | 1.10E-18 | 160 |

| | | | | | | | |
|---|---|---|---|---|---|---|---|
| k141_66218_flag1_multi1.0000_len346 | W4-1 | 346 | AYV88211.1 hypothetical protein [Mandarin fish ranavirus] | 97.8 | 46 | 1.13E-18 | 0 |
| k141_27221_flag1_multi1.0000_len316 | W5-1 | 316 | QJE49066.1 hypothetical protein LMBV_003 [Largemouth bass virus] | 86.3 | 51 | 1.14E-18 | 0 |
| k141_68868_flag1_multi13.6729_len1055 | W5-1 | 1055 | QJE49104.1 hypothetical protein LMBV_041 [Largemouth bass virus] | 100 | 46 | 1.14E-18 | 35 |
| k141_144065_flag1_multi23.9241_len7114 | W5-1 | 7114 | UUY86235.1 hypothetical protein [Largemouth bass virus] | 97.9 | 47 | 1.16E-18 | 585 |
| k141_17202_flag1_multi2.5227_len405 | W2-1 | 405 | UUY86236.1 hypothetical protein [Largemouth bass virus] | 93.6 | 47 | 1.17E-18 | 2 |
| k141_36062_flag1_multi20.1139_len1177 | W4-1 | 1177 | AIG51690.1 major capsid protein, partial [Koi ranavirus] | 88.9 | 54 | 1.17E-18 | 77 |
| k141_103740_flag1_multi4.8897_len975 | W5-1 | 975 | AYV88122.1 putative 3-beta-hydroxy-delta-5-C27 steroid oxidoreductase-like protein [Mandarin fish ranavirus] | 100 | 46 | 1.17E-18 | 8 |
| k141_115784_flag1_multi10.8708_len1921 | W5-1 | 1921 | AYV88191.1 hypothetical protein [Mandarin fish ranavirus] | 97.8 | 46 | 1.20E-18 | 67 |
| k141_106082_flag1_multi7.9140_len1199 | W5-1 | 1199 | UUY86192.1 putative myristylated membrane protein [Largemouth bass virus] | 100 | 43 | 1.21E-18 | 30 |
| k141_143735_flag1_multi4.8435_len729 | W5-1 | 729 | AAF64582.1 capsid protein, partial [Guppyfish iridovirus] | 84.9 | 53 | 1.22E-18 | 32 |
| k141_124142_flag1_multi2.8700_len464 | W1-1 | 464 | UUY86199.1 hypothetical protein [Largemouth bass virus] | 100 | 39 | 1.23E-18 | 5 |
| k141_6340_flag0_multi12.2043_len1120 | W4-1 | 1120 | AIG51690.1 major capsid protein, partial [Koi ranavirus] | 97.9 | 47 | 1.23E-18 | 81 |
| k141_147360_flag0_multi43.1424_len1293 | W5-1 | 1293 | XRL22790.1 hypothetical protein [Siniperca chuatsi ranavirus] | 91.8 | 49 | 1.29E-18 | 180 |
| k141_61364_flag1_multi1.0000_len332 | W5-1 | 332 | AYV88191.1 hypothetical protein [Mandarin fish ranavirus] | 95 | 40 | 1.30E-18 | 1 |
| k141_77390_flag1_multi6.2881_len443 | W5-1 | 443 | QJE49066.1 hypothetical protein LMBV_003 [Largemouth bass virus] | 100 | 46 | 1.43E-18 | 5 |
| k141_38997_flag1_multi7.7811_len712 | W4-1 | 712 | AYV88134.2 putative tyrosine kinase [Mandarin fish ranavirus] | 95.8 | 48 | 1.45E-18 | 20 |
| k141_79395_flag1_multi1.0000_len820 | W5-1 | 820 | WEI28998.1 hypothetical protein [Largemouth bass virus] | 100 | 47 | 1.49E-18 | 8 |
| k141_60048_flag1_multi1.0000_len355 | W6-2 | 355 | AIG51690.1 major capsid protein, partial [Koi ranavirus] | 88 | 50 | 1.51E-18 | 5 |
| k141_108648_flag1_multi178.2810_len1027 | W5-1 | 1027 | QJE49101.1 hypothetical protein LMBV_038 [Largemouth bass virus] | 78.7 | 61 | 1.52E-18 | 13 |
| k141_26757_flag1_multi2.3990_len557 | W2-1 | 557 | QJE49144.1 hypothetical protein LMBV_081 [Largemouth bass virus] | 100 | 43 | 1.55E-18 | 7 |
| k141_99003_flag1_multi1.9463_len569 | W5-1 | 569 | AIG51690.1 major capsid protein, partial [Koi ranavirus] | 94 | 50 | 1.56E-18 | 1 |
| k141_130810_flag0_multi1.0000_len688 | W5-1 | 688 | UUY86197.1 hypothetical protein [Largemouth bass virus] | 100 | 48 | 1.57E-18 | 993.14 |
| k141_60288_flag1_multi2.9494_len1130 | W5-1 | 1130 | UUY86235.1 hypothetical protein [Largemouth bass virus] | 100 | 46 | 1.62E-18 | 36 |
| k141_15243_flag1_multi7.6239_len1872 | W5-1 | 1872 | AYV88120.1 putative p31K protein [Mandarin fish ranavirus] | 100 | 46 | 1.63E-18 | 50 |
| k141_57560_flag1_multi1.0000_len351 | W1-1 | 351 | XRB52785.1 hypothetical protein LMBV_28 [Largemouth bass virus] | 97.4 | 39 | 1.71E-18 | 0 |
| k141_97330_flag1_multi43.9239_len3044 | W1-1 | 3044 | QJE49149.1 putative immediate early protein ICP-46 [Largemouth bass virus] | 100 | 46 | 1.71E-18 | 429 |
| k141_51689_flag1_multi5.6327_len386 | W2-2 | 386 | UUY86223.1 hypothetical protein [Largemouth bass virus] | 100 | 45 | 1.72E-18 | 6 |
| k141_44542_flag1_multi1.0000_len425 | W5-1 | 425 | AIG51690.1 major capsid protein, partial [Koi ranavirus] | 100 | 45 | 1.76E-18 | 3 |
| k141_89673_flag1_multi2.9331_len829 | W5-1 | 829 | UUY86227.1 hypothetical protein [Largemouth bass virus] | 93.5 | 46 | 1.78E-18 | 9 |
| k141_152840_flag1_multi3.0000_len542 | W5-1 | 542 | QJE49077.1 hypothetical protein LMBV_014 [Largemouth bass virus] | 90.7 | 43 | 1.79E-18 | 11 |
| k141_82606_flag1_multi1.0000_len341 | W1-1 | 341 | UUY86235.1 hypothetical protein [Largemouth bass virus] | 95.2 | 42 | 1.91E-18 | 0 |
| k141_95175_flag1_multi10.7442_len1040 | W1-1 | 1040 | AYV88134.2 putative tyrosine kinase [Mandarin fish ranavirus] | 100 | 46 | 2.00E-18 | 30 |
| k141_44527_flag1_multi1.5561_len711 | W5-1 | 711 | QJE49066.1 hypothetical protein LMBV_003 [Largemouth bass virus] | 51.9 | 108 | 2.00E-18 | 5 |
| k141_211057_flag1_multi3.0000_len306 | W6-3 | 306 | YP_073558.1 RNA-dependent DNA polymerase [lymphocystis disease virus-China] | 52.3 | 86 | 2.01E-18 | 2 |
| k141_40141_flag1_multi6.7108_len390 | W4-1 | 390 | UUY86229.1 putative DNA dependent RNA polymerase a subunit [Largemouth bass virus] | 95.6 | 45 | 2.04E-18 | 8 |
| k141_31785_flag1_multi1.6401_len505 | W5-1 | 505 | AYV88169.1 hypothetical protein [Mandarin fish ranavirus] | 81.8 | 55 | 2.07E-18 | 4 |
| k141_37295_flag1_multi3.9992_len1411 | W5-1 | 1411 | AYV88159.1 hypothetical protein [Mandarin fish ranavirus] | 100 | 46 | 2.08E-18 | 29 |
| k141_90518_flag1_multi8.6290_len637 | W5-1 | 637 | AYV88211.1 hypothetical protein [Mandarin fish ranavirus] | 95.9 | 49 | 2.11E-18 | 16 |

| | | | | | | | |
|---|---|---|---|---|---|---|---|
| k141_44836_flag1_multi1.0000_len497 | W2-1 | 497 | AYV88176.1 putative tumor necrosis factor receptor [Mandarin fish ranavirus] | 100 | 38 | 2.14E-18 | 0 |
| k141_18062_flag1_multi11.7731_len1256 | W1-1 | 1256 | AIG51690.1 major capsid protein, partial [Koi ranavirus] | 100 | 47 | 2.16E-18 | 29 |
| k141_95761_flag1_multi11.8192_len993 | W5-1 | 993 | UUY86218.1 hypothetical protein [Largemouth bass virus] | 72.6 | 62 | 2.19E-18 | 53 |
| k141_32285_flag1_multi2.0000_len346 | W5-1 | 346 | AYV88191.1 hypothetical protein [Mandarin fish ranavirus] | 97.4 | 39 | 2.20E-18 | 1 |
| k141_117699_flag1_multi10.4369_len917 | W5-1 | 917 | UUY86221.1 hypothetical protein [Largemouth bass virus] | 75.8 | 62 | 2.22E-18 | 28 |
| k141_24372_flag0_multi76.6733_len594 | W4-1 | 594 | UUY86261.1 hypothetical protein [Largemouth bass virus] | 100 | 47 | 2.23E-18 | 214 |
| k141_139896_flag1_multi16.3333_len345 | W5-1 | 345 | AYV88120.1 putative p31K protein [Mandarin fish ranavirus] | 97.6 | 42 | 2.31E-18 | 4 |
| k141_141506_flag1_multi4.6654_len930 | W5-1 | 930 | AAF64582.1 capsid protein, partial [Guppyfish iridovirus] | 95.7 | 46 | 2.34E-18 | 16 |
| k141_40358_flag1_multi1.7080_len391 | W5-1 | 391 | AYV88211.1 hypothetical protein [Mandarin fish ranavirus] | 89.4 | 47 | 2.65E-18 | 0 |
| k141_99842_flag1_multi2.9024_len510 | W1-1 | 510 | AYV88120.1 putative p31K protein [Mandarin fish ranavirus] | 95.5 | 44 | 2.67E-18 | 5 |
| k141_16627_flag0_multi1.0000_len249 | W4-1 | 249 | AYV88191.1 hypothetical protein [Mandarin fish ranavirus] | 100 | 37 | 2.67E-18 | 0.61 |
| k141_1237_flag1_multi7.8530_len1046 | W4-1 | 1046 | AYV88120.1 putative p31K protein [Mandarin fish ranavirus] | 97.8 | 45 | 2.76E-18 | 25 |
| k141_180778_flag1_multi2.2715_len782 | W2-3 | 782 | WHA35533.1 putative DNA dependent RNA polymerase A subunit [Micropterus salmoides ranavirus] | 95.9 | 49 | 2.77E-18 | 15 |
| k141_24684_flag1_multi17.5278_len2765 | W5-1 | 2765 | AYV88173.2 putative ribonucleotide reductase beta subunit [Mandarin fish ranavirus] | 100 | 46 | 2.87E-18 | 153 |
| k141_150415_flag1_multi13.1874_len1235 | W5-1 | 1235 | XRB52768.1 P31K protein, partial [Largemouth bass virus] | 97.8 | 46 | 2.88E-18 | 66 |
| k141_33261_flag1_multi4.0000_len470 | W2-1 | 470 | WEI28972.1 CTD-phosphotransferase [Largemouth bass virus] | 66.2 | 68 | 2.95E-18 | 3 |
| k141_40603_flag0_multi1.0000_len334 | W2-1 | 334 | WEI28972.1 CTD-phosphotransferase [Largemouth bass virus] | 97.7 | 44 | 2.99E-18 | 4.87 |
| k141_46772_flag1_multi20.3106_len508 | W5-1 | 508 | UUY86261.1 hypothetical protein [Largemouth bass virus] | 97.9 | 48 | 3.36E-18 | 24 |
| k141_29676_flag1_multi5.8031_len725 | W5-1 | 725 | AYV88211.1 hypothetical protein [Mandarin fish ranavirus] | 97.9 | 47 | 3.44E-18 | 12 |
| k141_124028_flag1_multi4.7322_len1082 | W5-1 | 1082 | WEI28972.1 CTD-phosphotransferase [Largemouth bass virus] | 100 | 46 | 3.45E-18 | 14 |
| k141_62907_flag1_multi10.8653_len1744 | W5-1 | 1744 | UUY86235.1 hypothetical protein [Largemouth bass virus] | 100 | 43 | 3.47E-18 | 87 |
| k141_94855_flag1_multi13.0000_len369 | W2-1 | 369 | QIZ30887.1 major capsid protein, partial [Largemouth bass virus] | 100 | 39 | 3.57E-18 | 11 |
| k141_142375_flag1_multi3.2768_len701 | W5-1 | 701 | QJE49215.1 putative orf58-like protein [Largemouth bass virus] | 100 | 41 | 3.68E-18 | 11 |
| k141_34289_flag1_multi3.9053_len870 | W4-1 | 870 | AYV88120.1 putative p31K protein [Mandarin fish ranavirus] | 97.8 | 46 | 3.69E-18 | 11 |
| k141_37964_flag1_multi1.0000_len269 | W5-1 | 269 | QYU76034.1 putative myristylated membrane protein, partial [Koi ranavirus] | 95.9 | 49 | 3.72E-18 | 0 |
| k141_6474_flag1_multi8.8323_len1238 | W5-1 | 1238 | AIG51690.1 major capsid protein, partial [Koi ranavirus] | 100 | 46 | 3.74E-18 | 51 |
| k141_13785_flag0_multi14.1896_len1333 | W1-1 | 1333 | AIG51690.1 major capsid protein, partial [Koi ranavirus] | 92.2 | 51 | 3.82E-18 | 78 |
| k141_5481_flag1_multi1.7681_len624 | W1-1 | 624 | WAK75102.1 hypothetical protein [Mandarin fish ranavirus] | 97.7 | 44 | 4.09E-18 | 3 |
| k141_22894_flag1_multi5.8341_len834 | W4-1 | 834 | AYV88120.1 putative p31K protein [Mandarin fish ranavirus] | 97.8 | 46 | 4.12E-18 | 17 |
| k141_148391_flag1_multi9.0888_len614 | W5-1 | 614 | AYV88175.2 putative interleukin-1 beta convertase precursor [Mandarin fish ranavirus] | 100 | 44 | 4.12E-18 | 21 |
| k141_139912_flag1_multi79.1437_len2354 | W5-1 | 2354 | AYV88120.1 putative p31K protein [Mandarin fish ranavirus] | 97.8 | 46 | 4.16E-18 | 609 |
| k141_136222_flag1_multi2.8987_len595 | W1-1 | 595 | WAK75112.1 putative LITAF PIG7 possible membrane associated motif in LPS-induced tumor necrosis factor alpha factor [Mandarin fish ranavirus] | 97.4 | 39 | 4.21E-18 | 2 |
| k141_131438_flag0_multi9.7315_len357 | W5-1 | 357 | QYU76034.1 putative myristylated membrane protein, partial [Koi ranavirus] | 90 | 50 | 4.23E-18 | 7 |
| k141_126317_flag1_multi2.9147_len1619 | W5-1 | 1619 | QJE49125.1 hypothetical protein LMBV_062 [Largemouth bass virus] | 100 | 46 | 4.36E-18 | 12 |
| k141_57616_flag1_multi21.6398_len1482 | W5-1 | 1482 | AAC79508.1 viral core protein, partial [Ictalurus melas ranavirus] | 89.1 | 46 | 4.36E-18 | 78 |
| k141_150497_flag1_multi2.7284_len686 | W5-1 | 686 | AYV88169.1 hypothetical protein [Mandarin fish ranavirus] | 93.8 | 48 | 4.38E-18 | 10 |
| k141_61989_flag1_multi9.0000_len447 | W5-1 | 447 | UUY86241.1 hypothetical protein [Largemouth bass virus] | 100 | 35 | 4.39E-18 | 7 |

| | | | | | | | |
|---|---|---|---|---|---|---|---|
| k141_127794_flag1_multi10.1663_len1151 | W1-1 | 1151 | AYV88120.1 putative p31K protein [Mandarin fish ranavirus] | 97.8 | 45 | 4.43E-18 | 67 |
| k141_102706_flag1_multi2.4347_len723 | W4-1 | 723 | AIG51690.1 major capsid protein, partial [Koi ranavirus] | 97.9 | 47 | 4.52E-18 | 2 |
| k141_103088_flag1_multi1.0000_len519 | W5-1 | 519 | AYV88134.2 putative tyrosine kinase [Mandarin fish ranavirus] | 100 | 41 | 4.66E-18 | 11 |
| k141_143201_flag1_multi5.0000_len512 | W5-1 | 512 | AYV88191.1 hypothetical protein [Mandarin fish ranavirus] | 100 | 39 | 4.70E-18 | 7 |
| k141_46760_flag1_multi3.5974_len372 | W5-1 | 372 | UUY86261.1 hypothetical protein [Largemouth bass virus] | 100 | 46 | 4.74E-18 | 3 |
| k141_139442_flag0_multi12.9149_len6347 | W5-1 | 6347 | AYV88159.1 hypothetical protein [Mandarin fish ranavirus] | 100 | 46 | 4.75E-18 | 240 |
| k141_162201_flag0_multi16.9544_len1304 | W5-1 | 1304 | QJE49066.1 hypothetical protein LMBV_003 [Largemouth bass virus] | 78.5 | 65 | 4.76E-18 | 151 |
| k141_47848_flag1_multi1.0000_len279 | W1-1 | 279 | UVF58785.1 MAG: major capsid protein [Halichoeres melanurus ranavirus] | 97.9 | 47 | 4.95E-18 | 0 |
| k141_90310_flag1_multi4.5094_len622 | W4-1 | 622 | WAK75117.1 hypothetical protein [Mandarin fish ranavirus] | 90.5 | 42 | 5.14E-18 | 8 |
| k141_94001_flag1_multi1.0000_len330 | W5-1 | 330 | WAK75097.1 hypothetical protein [Mandarin fish ranavirus] | 100 | 44 | 5.17E-18 | 2 |
| k141_165022_flag0_multi10.3906_len781 | W5-1 | 781 | AYV88120.1 putative p31K protein [Mandarin fish ranavirus] | 97.7 | 44 | 5.32E-18 | 12 |
| k141_86545_flag1_multi2.5207_len358 | W5-1 | 358 | WEI28972.1 CTD-phosphotransferase [Largemouth bass virus] | 100 | 44 | 5.49E-18 | 0 |
| k141_139615_flag0_multi1.0000_len392 | W5-1 | 392 | XRB52831.1 hypothetical protein LMBV_74 [Largemouth bass virus] | 91.7 | 60 | 5.51E-18 | 1177.81 |
| k141_57817_flag0_multi1.0000_len302 | W5-1 | 302 | UUY86258.1 hypothetical protein [Largemouth bass virus] | 100 | 46 | 5.60E-18 | 3.35 |
| k141_154340_flag1_multi6.9395_len1033 | W5-1 | 1033 | AFD96401.1 major capsid protein, partial [Largemouth bass virus] | 97.7 | 43 | 5.76E-18 | 20 |
| k141_62718_flag1_multi47.8887_len4373 | W1-1 | 4373 | AIG51690.1 major capsid protein, partial [Koi ranavirus] | 80.6 | 62 | 5.93E-18 | 535 |
| k141_97305_flag1_multi5.9015_len801 | W1-1 | 801 | AIG51690.1 major capsid protein, partial [Koi ranavirus] | 100 | 44 | 5.97E-18 | 16 |
| k141_72208_flag1_multi2.9214_len421 | W5-1 | 421 | AIG51690.1 major capsid protein, partial [Koi ranavirus] | 100 | 43 | 6.04E-18 | 1 |
| k141_17063_flag1_multi1.0000_len428 | W5-1 | 428 | AYV88120.1 putative p31K protein [Mandarin fish ranavirus] | 97.6 | 42 | 6.09E-18 | 4.16 |
| k141_83280_flag1_multi4.8943_len898 | W2-1 | 898 | AIG51690.1 major capsid protein, partial [Koi ranavirus] | 95.8 | 48 | 6.11E-18 | 17 |
| k141_131931_flag0_multi9.7783_len1968 | W1-1 | 1968 | UUY86204.1 hypothetical protein [Largemouth bass virus] | 100 | 47 | 6.36E-18 | 137 |
| k141_128886_flag1_multi18.9667_len1223 | W5-1 | 1223 | QJE49077.1 hypothetical protein LMBV_014 [Largemouth bass virus] | 100 | 39 | 6.43E-18 | 61 |
| k141_81064_flag0_multi1.0000_len281 | W4-1 | 281 | UVF58785.1 MAG: major capsid protein [Halichoeres melanurus ranavirus] | 100 | 46 | 6.79E-18 | 0 |
| k141_158485_flag1_multi10.5150_len4904 | W5-1 | 4904 | AYV88179.2 putative DNA dependent RNA polymerase II second largest subunit [Mandarin fish ranavirus] | 78.8 | 66 | 6.83E-18 | 183 |
| k141_46668_flag1_multi53.7912_len4423 | W4-1 | 4423 | WAK75068.1 hypothetical protein [Mandarin fish ranavirus] | 90 | 50 | 6.90E-18 | 181 |
| k141_31549_flag1_multi4.0000_len687 | W1-1 | 687 | QYU76034.1 putative myristylated membrane protein, partial [Koi ranavirus] | 89.8 | 49 | 7.01E-18 | 5 |
| k141_84657_flag1_multi7.7402_len599 | W2-1 | 599 | QJE49065.1 hypothetical protein LMBV_002 [Largemouth bass virus] | 97.5 | 40 | 7.32E-18 | 13 |
| k141_113969_flag1_multi4.9725_len1051 | W5-1 | 1051 | QJE49146.1 putative thiol oxidoreductase [Largemouth bass virus] | 100 | 42 | 7.68E-18 | 10 |
| k141_112534_flag1_multi1.0000_len444 | W5-1 | 444 | AAC79876.1 viral core protein, partial [Labroides dimidatus ranavirus] | 100 | 39 | 7.71E-18 | 5.01 |
| k141_32600_flag1_multi1.9055_len416 | W4-2 | 416 | AAF64582.1 capsid protein, partial [Guppyfish iridovirus] | 97.6 | 42 | 7.76E-18 | 0 |
| k141_38993_flag0_multi1.0000_len279 | W1-1 | 279 | AIG51690.1 major capsid protein, partial [Koi ranavirus] | 97.9 | 47 | 7.80E-18 | 6 |
| k141_25207_flag1_multi3.6233_len1264 | W5-1 | 1264 | UUY86195.1 hypothetical protein [Largemouth bass virus] | 97.8 | 46 | 7.85E-18 | 10 |
| k141_9421_flag1_multi13.0000_len1334 | W4-1 | 1334 | QIZ30887.1 major capsid protein, partial [Largemouth bass virus] | 52.6 | 95 | 7.91E-18 | 47 |
| k141_20686_flag1_multi1.0000_len572 | W5-1 | 572 | AIG51690.1 major capsid protein, partial [Koi ranavirus] | 100 | 46 | 8.10E-18 | 4 |
| k141_13992_flag1_multi11.1116_len2158 | W5-1 | 2158 | XRB52797.1 Bcl-2 Bcl-2 homology region 1-3 domain-containing protein [Largemouth bass virus] | 97.8 | 46 | 8.16E-18 | 80 |
| k141_130371_flag1_multi8.6349_len897 | W1-1 | 897 | AIG51690.1 major capsid protein, partial [Koi ranavirus] | 97.8 | 46 | 8.22E-18 | 22 |
| k141_39772_flag1_multi10.7624_len1067 | W4-1 | 1067 | AIG51690.1 major capsid protein, partial [Koi ranavirus] | 97.9 | 47 | 8.42E-18 | 95.29 |
| k141_73391_flag0_multi8.5429_len526 | W4-1 | 526 | WAK75070.1 putative neurofilament triplet H1-like protein [Mandarin fish ranavirus] | 77.2 | 57 | 8.66E-18 | 10 |

| | | | | | | | |
|---|---|---|---|---|---|---|---|
| k141_12611_flag0_multi28.3574_len6283 | W5-1 | 6283 | UUY86235.1 hypothetical protein [Largemouth bass virus] | 100 | 46 | 8.68E-18 | 615 |
| k141_109160_flag1_multi8.9106_len2200 | W1-2 | 2200 | WEI29006.1 putative 2-cysteine adaptor domain protein [Largemouth bass virus] | 95.8 | 48 | 8.73E-18 | 143 |
| k141_65526_flag1_multi4.7254_len476 | W5-1 | 476 | UUY86258.1 hypothetical protein [Largemouth bass virus] | 100 | 46 | 9.15E-18 | 8 |
| k141_21644_flag1_multi2.1626_len547 | W5-1 | 547 | UUY86192.1 putative myristylated membrane protein [Largemouth bass virus] | 77.1 | 48 | 9.22E-18 | 6 |
| k141_164582_flag1_multi8.7875_len781 | W5-1 | 781 | QIZ30887.1 major capsid protein, partial [Largemouth bass virus] | 95.2 | 42 | 9.80E-18 | 23 |
| k141_86031_flag0_multi9.1831_len354 | W4-1 | 354 | AYV88198.1 hypothetical protein [Mandarin fish ranavirus] | 97.7 | 43 | 9.93E-18 | 5 |
| k141_53926_flag1_multi3.3996_len609 | W5-1 | 609 | QJE49153.1 hypothetical protein LMBV_004 [Largemouth bass virus] | 97.8 | 46 | 1.01E-17 | 5 |
| k141_20801_flag1_multi4.3269_len1019 | W6-1 | 1019 | QJE49077.1 hypothetical protein LMBV_014 [Largemouth bass virus] | 97.6 | 41 | 1.03E-17 | 90.04 |
| k141_132287_flag1_multi3.8763_len796 | W5-1 | 796 | AYV88121.1 hypothetical protein [Mandarin fish ranavirus] | 97.8 | 46 | 1.08E-17 | 4 |
| k141_35806_flag1_multi7.8591_len1284 | W5-1 | 1284 | WEI29006.1 putative 2-cysteine adaptor domain protein [Largemouth bass virus] | 93.9 | 49 | 1.14E-17 | 42 |
| k141_6300_flag1_multi42.0000_len1711 | W5-1 | 1711 | AYV88159.1 hypothetical protein [Mandarin fish ranavirus] | 100 | 44 | 1.15E-17 | 294 |
| k141_145305_flag1_multi12.9370_len760 | W6-2 | 760 | UUY86192.1 putative myristylated membrane protein [Largemouth bass virus] | 100 | 38 | 1.15E-17 | 24 |
| k141_14675_flag1_multi2.3534_len605 | W2-3 | 605 | AYV88179.2 putative DNA dependent RNA polymerase II second largest subunit [Mandarin fish ranavirus] | 100 | 43 | 1.16E-17 | 6 |
| k141_82705_flag1_multi8.9520_len3976 | W5-1 | 3976 | AAF64582.1 capsid protein, partial [Guppyfish iridovirus] | 93.8 | 48 | 1.16E-17 | 102 |
| k141_152249_flag1_multi4.9223_len450 | W5-1 | 450 | QIZ30887.1 major capsid protein, partial [Largemouth bass virus] | 92.9 | 42 | 1.19E-17 | 0 |
| k141_131316_flag1_multi2.9216_len728 | W5-1 | 728 | UVF58785.1 MAG: major capsid protein [Halichoeres melanurus ranavirus] | 85.5 | 55 | 1.19E-17 | 6 |
| k141_72809_flag0_multi1.0000_len289 | W1-1 | 289 | AIG51690.1 major capsid protein, partial [Koi ranavirus] | 100 | 45 | 1.21E-17 | 0 |
| k141_125819_flag1_multi4.0000_len455 | W5-1 | 455 | QIZ30887.1 major capsid protein, partial [Largemouth bass virus] | 100 | 39 | 1.27E-17 | 4 |
| k141_92906_flag1_multi4.0000_len802 | W4-3 | 802 | WEI29006.1 putative 2-cysteine adaptor domain protein [Largemouth bass virus] | 100 | 39 | 1.35E-17 | 10 |
| k141_96520_flag0_multi39.8919_len3804 | W2-3 | 3804 | AYV88120.1 putative p31K protein [Mandarin fish ranavirus] | 97.8 | 46 | 1.38E-17 | 569.21 |
| k141_60042_flag1_multi1.0000_len395 | W2-1 | 395 | QJE49101.1 hypothetical protein LMBV_038 [Largemouth bass virus] | 93.3 | 45 | 1.48E-17 | 3 |
| k141_19136_flag1_multi38.9071_len9545 | W1-1 | 9545 | AYV88120.1 putative p31K protein [Mandarin fish ranavirus] | 97.8 | 46 | 1.54E-17 | 1062 |
| k141_22565_flag1_multi9.9204_len2743 | W1-1 | 2743 | XRB52768.1 P31K protein, partial [Largemouth bass virus] | 95.7 | 47 | 1.57E-17 | 99 |
| k141_10215_flag1_multi5.3256_len1342 | W6-1 | 1342 | UVF58793.1 MAG: DNA polymerase [Halichoeres melanurus ranavirus] | 100 | 45 | 1.59E-17 | 20 |
| k141_71523_flag1_multi6.8311_len958 | W4-1 | 958 | AIG51690.1 major capsid protein, partial [Koi ranavirus] | 100 | 46 | 1.62E-17 | 18 |
| k141_158086_flag1_multi1.0000_len415 | W5-1 | 415 | WAK75112.1 putative LITAF PIG7 possible membrane associated motif in LPS-induced tumor necrosis factor alpha factor [Mandarin fish ranavirus] | 100 | 38 | 1.62E-17 | 1 |
| k141_138710_flag1_multi1.0000_len312 | W5-1 | 312 | AIG51690.1 major capsid protein, partial [Koi ranavirus] | 100 | 41 | 1.64E-17 | 0 |
| k141_46274_flag1_multi2.9117_len526 | W4-1 | 526 | UVF58785.1 MAG: major capsid protein [Halichoeres melanurus ranavirus] | 100 | 46 | 1.66E-17 | 5 |
| k141_11826_flag1_multi5.9123_len369 | W4-1 | 369 | AYV88120.1 putative p31K protein [Mandarin fish ranavirus] | 91.1 | 45 | 1.66E-17 | 0 |
| k141_165579_flag0_multi29.1505_len513 | W5-1 | 513 | UUY86261.1 hypothetical protein [Largemouth bass virus] | 100 | 46 | 1.66E-17 | 124 |
| k141_102827_flag1_multi4.8655_len825 | W5-1 | 825 | XRL22790.1 hypothetical protein [Siniperca chuatsi ranavirus] | 97.9 | 48 | 1.73E-17 | 12 |
| k141_25020_flag1_multi13.0000_len342 | W4-1 | 342 | AYV88120.1 putative p31K protein [Mandarin fish ranavirus] | 100 | 39 | 1.75E-17 | 8 |
| k141_33871_flag1_multi4.8874_len958 | W1-1 | 958 | QJE49088.1 hypothetical protein LMBV_025 [Largemouth bass virus] | 100 | 46 | 1.80E-17 | 9 |
| k141_159368_flag1_multi5.0633_len899 | W5-1 | 899 | WAK75112.1 putative LITAF PIG7 possible membrane associated motif in LPS-induced tumor necrosis factor alpha factor [Mandarin fish ranavirus] | 95 | 40 | 1.85E-17 | 12 |
| k141_57164_flag0_multi8.4210_len761 | W5-1 | 761 | QJE49137.1 hypothetical protein LMBV_074 [Largemouth bass virus] | 97.8 | 46 | 1.92E-17 | 16 |
| k141_102591_flag1_multi15.9692_len2313 | W1-1 | 2313 | WEI28972.1 CTD-phosphotransferase [Largemouth bass virus] | 95.8 | 48 | 1.93E-17 | 106 |
| k141_28809_flag1_multi3.7669_len437 | W5-1 | 437 | AIG51690.1 major capsid protein, partial [Koi ranavirus] | 69.4 | 62 | 1.95E-17 | 5 |

| | | | | | | | |
|---|---|---|---|---|---|---|---|
| k141_32212_flag1_multi1.0000_len269 | W5-1 | 269 | UVF58785.1 MAG: major capsid protein [Halichoeres melanurus ranavirus] | 100 | 42 | 2.07E-17 | 0 |
| k141_21777_flag0_multi7.6455_len1340 | W4-1 | 1340 | AYV88134.2 putative tyrosine kinase [Mandarin fish ranavirus] | 97.7 | 44 | 2.10E-17 | 30 |
| k141_120678_flag1_multi2.0000_len640 | W5-1 | 640 | AYV88118.1 hypothetical protein [Mandarin fish ranavirus] | 100 | 39 | 2.15E-17 | 6.1 |
| k141_30342_flag0_multi6.1799_len1992 | W2-1 | 1992 | UUY86241.1 hypothetical protein [Largemouth bass virus] | 84.1 | 44 | 2.18E-17 | 82.1 |
| k141_64205_flag1_multi5.3921_len544 | W5-1 | 544 | QJE49137.1 hypothetical protein LMBV_074 [Largemouth bass virus] | 100 | 41 | 2.18E-17 | 11 |
| k141_133439_flag1_multi1.0000_len511 | W5-1 | 511 | QIZ30887.1 major capsid protein, partial [Largemouth bass virus] | 97.4 | 39 | 2.19E-17 | 37 |
| k141_63973_flag0_multi1.0000_len281 | W5-1 | 281 | QJE49073.1 putative p31K protein [Largemouth bass virus] | 100 | 46 | 2.25E-17 | 0 |
| k141_130104_flag1_multi3.0000_len721 | W5-1 | 721 | AAC79876.1 viral core protein, partial [Labroides dimidatus ranavirus] | 100 | 39 | 2.28E-17 | 9 |
| k141_40111_flag1_multi1.0000_len313 | W1-1 | 313 | AYV88120.1 putative p31K protein [Mandarin fish ranavirus] | 97.6 | 42 | 2.30E-17 | 2 |
| k141_32063_flag1_multi3.2065_len417 | W4-1 | 417 | QJE49137.1 hypothetical protein LMBV_074 [Largemouth bass virus] | 79.2 | 53 | 2.45E-17 | 3 |
| k141_79270_flag1_multi1.0000_len842 | W5-1 | 842 | UUY86235.1 hypothetical protein [Largemouth bass virus] | 100 | 46 | 2.47E-17 | 6 |
| k141_166906_flag1_multi1.0000_len354 | W5-1 | 354 | QJE49110.1 hypothetical protein LMBV_047 [Largemouth bass virus] | 95.3 | 43 | 2.53E-17 | 4 |
| k141_125442_flag1_multi1.0000_len387 | W1-1 | 387 | UUY86197.1 hypothetical protein [Largemouth bass virus] | 100 | 41 | 2.56E-17 | 2 |
| k141_48830_flag1_multi5.9643_len2382 | W5-1 | 2382 | UUY86266.1 hypothetical protein [Largemouth bass virus] | 97.7 | 44 | 2.56E-17 | 54 |
| k141_33634_flag0_multi1.0000_len425 | W5-1 | 425 | WEI29006.1 putative 2-cysteine adaptor domain protein [Largemouth bass virus] | 100 | 40 | 2.63E-17 | 25 |
| k141_27192_flag1_multi1.0000_len790 | W5-1 | 790 | AIG51690.1 major capsid protein, partial [Koi ranavirus] | 97.6 | 42 | 2.69E-17 | 21 |
| k141_80145_flag1_multi9.7650_len1022 | W5-1 | 1022 | AYV88211.1 hypothetical protein [Mandarin fish ranavirus] | 100 | 46 | 2.74E-17 | 28 |
| k141_146674_flag1_multi3.9359_len1218 | W5-1 | 1218 | WEI29006.1 putative 2-cysteine adaptor domain protein [Largemouth bass virus] | 97.8 | 46 | 2.75E-17 | 14 |
| k141_135011_flag0_multi1.0000_len309 | W5-1 | 309 | UUY86235.1 hypothetical protein [Largemouth bass virus] | 100 | 43 | 2.77E-17 | 3 |
| k141_41495_flag1_multi1.0000_len599 | W5-1 | 599 | UUY86229.1 putative DNA dependent RNA polymerase a subunit [Largemouth bass virus] | 97.7 | 44 | 2.82E-17 | 6 |
| k141_66947_flag1_multi5.1094_len461 | W4-1 | 461 | XRL22821.1 hypothetical protein [Siniperca chuatsi ranavirus] | 100 | 40 | 2.96E-17 | 3 |
| k141_5690_flag1_multi5.0000_len662 | W5-1 | 662 | AFD96401.1 major capsid protein, partial [Largemouth bass virus] | 100 | 39 | 2.99E-17 | 6 |
| k141_137750_flag1_multi24.2398_len5128 | W1-1 | 5128 | QJE49100.1 hypothetical protein LMBV_037 [Largemouth bass virus] | 97.9 | 47 | 3.07E-17 | 266 |
| k141_145654_flag1_multi5.8594_len781 | W2-3 | 781 | QJE49100.1 hypothetical protein LMBV_037 [Largemouth bass virus] | 100 | 45 | 3.07E-17 | 14 |
| k141_115049_flag1_multi5.7444_len591 | W5-1 | 591 | UUY86197.1 hypothetical protein [Largemouth bass virus] | 100 | 46 | 3.10E-17 | 3 |
| k141_43274_flag1_multi32.8463_len798 | W5-1 | 798 | WEI28972.1 CTD-phosphotransferase [Largemouth bass virus] | 100 | 46 | 3.16E-17 | 91 |
| k141_128496_flag1_multi3.8428_len580 | W1-1 | 580 | UUY86261.1 hypothetical protein [Largemouth bass virus] | 100 | 46 | 3.30E-17 | 7 |
| k141_99631_flag1_multi17.4635_len868 | W4-1 | 868 | UUY86211.1 hypothetical protein [Largemouth bass virus] | 67.1 | 73 | 3.31E-17 | 35 |
| k141_103457_flag1_multi7.0000_len588 | W5-1 | 588 | QIZ30887.1 major capsid protein, partial [Largemouth bass virus] | 97.5 | 40 | 3.31E-17 | 12 |
| k141_163008_flag1_multi7.9148_len2689 | W5-1 | 2689 | UUY86238.1 hypothetical protein [Largemouth bass virus] | 100 | 47 | 3.44E-17 | 94 |
| k141_54192_flag1_multi1.9558_len594 | W5-1 | 594 | AYV88134.2 putative tyrosine kinase [Mandarin fish ranavirus] | 100 | 43 | 3.45E-17 | 3 |
| k141_28118_flag1_multi7.0000_len658 | W5-1 | 658 | AYV88191.1 hypothetical protein [Mandarin fish ranavirus] | 97.4 | 39 | 3.49E-17 | 19 |
| k141_146266_flag1_multi13.7506_len2270 | W5-1 | 2270 | QJE49101.1 hypothetical protein LMBV_038 [Largemouth bass virus] | 97.8 | 46 | 3.52E-17 | 107 |
| k141_113158_flag1_multi1.0000_len628 | W1-1 | 628 | AIG51690.1 major capsid protein, partial [Koi ranavirus] | 100 | 42 | 3.60E-17 | 20 |
| k141_94472_flag0_multi1.0000_len350 | W4-1 | 350 | AYV88191.1 hypothetical protein [Mandarin fish ranavirus] | 100 | 37 | 3.65E-17 | 7 |
| k141_32956_flag1_multi6.5382_len821 | W1-1 | 821 | WEI28972.1 CTD-phosphotransferase [Largemouth bass virus] | 70 | 60 | 3.69E-17 | 23 |
| k141_28448_flag1_multi9.9081_len2393 | W5-1 | 2393 | AIG51690.1 major capsid protein, partial [Koi ranavirus] | 100 | 46 | 3.74E-17 | 91 |
| k141_139266_flag1_multi1.0000_len352 | W2-3 | 352 | AYV88136.2 hypothetical protein [Mandarin fish ranavirus] | 83.7 | 43 | 3.86E-17 | 4 |
| k141_67328_flag1_multi3.3333_len1215 | W1-1 | 1215 | WXI69503.1 hypothetical protein [Largemouth bass virus] | 90 | 50 | 4.09E-17 | 48 |

| | | | | | | | |
|---|---|---|---|---|---|---|---|
| k141_90250_flag1_multi17.8899_len1222 | W1-1 | 1222 | XRL22821.1 hypothetical protein [Siniperca chuatsi ranavirus] | 97.7 | 43 | 4.13E-17 | 76 |
| k141_26547_flag1_multi1.0000_len483 | W5-1 | 483 | AYV88120.1 putative p31K protein [Mandarin fish ranavirus] | 78.3 | 60 | 4.15E-17 | 6.93 |
| k141_6950_flag1_multi6.0000_len361 | W4-1 | 361 | AYV88191.1 hypothetical protein [Mandarin fish ranavirus] | 100 | 35 | 4.20E-17 | 6 |
| k141_69043_flag1_multi68.0000_len756 | W1-1 | 756 | UUY86199.1 hypothetical protein [Largemouth bass virus] | 97.4 | 39 | 4.27E-17 | 162 |
| k141_3251_flag1_multi27.0000_len3550 | W5-1 | 3550 | QJE49114.1 hypothetical protein LMBV_051 [Largemouth bass virus] | 100 | 39 | 4.27E-17 | 254 |
| k141_90351_flag1_multi4.0312_len494 | W2-1 | 494 | AYV88191.1 hypothetical protein [Mandarin fish ranavirus] | 92.5 | 40 | 4.36E-17 | 5 |
| k141_112600_flag1_multi0.9596_len364 | W5-1 | 364 | AIG51690.1 major capsid protein, partial [Koi ranavirus] | 89.8 | 49 | 4.36E-17 | 1 |
| k141_29195_flag0_multi1.0000_len336 | W5-1 | 336 | UVF58793.1 MAG: DNA polymerase [Halichoeres melanurus ranavirus] | 75.9 | 58 | 4.43E-17 | 2 |
| k141_107110_flag1_multi2.2789_len539 | W1-1 | 539 | UUY86267.1 ribonucleotide reductase alpha subunit [Largemouth bass virus] | 100 | 40 | 4.46E-17 | 2 |
| k141_107627_flag1_multi8.3654_len1917 | W5-1 | 1917 | UUY86267.1 ribonucleotide reductase alpha subunit [Largemouth bass virus] | 100 | 42 | 4.46E-17 | 56 |
| k141_44620_flag1_multi7.9055_len2290 | W5-1 | 2290 | WEI28972.1 CTD-phosphotransferase [Largemouth bass virus] | 100 | 46 | 4.48E-17 | 51 |
| k141_57867_flag0_multi16.8760_len270 | W5-1 | 270 | XRL22821.1 hypothetical protein [Siniperca chuatsi ranavirus] | 100 | 39 | 4.61E-17 | 0 |
| k141_44665_flag1_multi5.0000_len821 | W5-1 | 821 | QIZ30887.1 major capsid protein, partial [Largemouth bass virus] | 85.4 | 48 | 4.87E-17 | 7 |
| k141_48159_flag0_multi56.2530_len1157 | W5-1 | 1157 | WEI29006.1 putative 2-cysteine adaptor domain protein [Largemouth bass virus] | 92.2 | 51 | 5.02E-17 | 223 |
| k141_130472_flag1_multi6.2425_len607 | W5-1 | 607 | UVF58785.1 MAG: major capsid protein [Halichoeres melanurus ranavirus] | 100 | 44 | 5.13E-17 | 108 |
| k141_87328_flag1_multi14.7718_len1552 | W2-1 | 1552 | AIG51690.1 major capsid protein, partial [Koi ranavirus] | 100 | 46 | 5.29E-17 | 69 |
| k141_129593_flag1_multi1.0000_len327 | W1-1 | 327 | AIG51690.1 major capsid protein, partial [Koi ranavirus] | 70.6 | 68 | 5.43E-17 | 3 |
| k141_29442_flag1_multi3.0000_len374 | W4-1 | 374 | AFD96401.1 major capsid protein, partial [Largemouth bass virus] | 97.4 | 39 | 5.53E-17 | 1 |
| k141_80451_flag1_multi2.8762_len545 | W2-1 | 545 | AYV88120.1 putative p31K protein [Mandarin fish ranavirus] | 100 | 46 | 5.55E-17 | 13 |
| k141_114671_flag1_multi18.0000_len867 | W5-1 | 867 | UUY86235.1 hypothetical protein [Largemouth bass virus] | 45 | 131 | 5.56E-17 | 57 |
| k141_94858_flag1_multi8.9366_len1276 | W4-1 | 1276 | AYV88191.1 hypothetical protein [Mandarin fish ranavirus] | 97.6 | 42 | 5.68E-17 | 34 |
| k141_29210_flag1_multi11.9377_len4024 | W5-1 | 4024 | UUY86199.1 hypothetical protein [Largemouth bass virus] | 97.4 | 39 | 5.71E-17 | 146 |
| k141_17083_flag1_multi2.9369_len870 | W5-1 | 870 | UUY86269.1 hypothetical protein [Largemouth bass virus] | 100 | 46 | 5.74E-17 | 8 |
| k141_29947_flag0_multi19.0692_len488 | W5-1 | 488 | QJE49066.1 hypothetical protein LMBV_003 [Largemouth bass virus] | 81.1 | 53 | 5.78E-17 | 10 |
| k141_49804_flag1_multi7.0000_len540 | W4-1 | 540 | QIZ30887.1 major capsid protein, partial [Largemouth bass virus] | 100 | 38 | 5.82E-17 | 9 |
| k141_80896_flag1_multi2.0000_len499 | W5-1 | 499 | UUY86235.1 hypothetical protein [Largemouth bass virus] | 93 | 43 | 6.03E-17 | 2 |
| k141_1454_flag0_multi12.7577_len595 | W4-1 | 595 | AYV88199.1 hypothetical protein [Mandarin fish ranavirus] | 80.4 | 51 | 6.55E-17 | 8 |
| k141_158190_flag1_multi1.0000_len701 | W5-1 | 701 | AIG51690.1 major capsid protein, partial [Koi ranavirus] | 100 | 47 | 6.64E-17 | 5 |
| k141_21114_flag1_multi1.0000_len400 | W1-1 | 400 | QJE49081.1 hypothetical protein LMBV_018 [Largemouth bass virus] | 100 | 39 | 7.02E-17 | 0 |
| k141_146365_flag1_multi18.1417_len2413 | W5-1 | 2413 | QJE49112.1 hypothetical protein LMBV_049 [Largemouth bass virus] | 100 | 39 | 7.07E-17 | 140 |
| k141_142252_flag1_multi3.1546_len665 | W5-1 | 665 | UUY86261.1 hypothetical protein [Largemouth bass virus] | 95.8 | 48 | 7.08E-17 | 7 |
| k141_1511_flag1_multi6.9478_len486 | W1-1 | 486 | UVF58793.1 MAG: DNA polymerase [Halichoeres melanurus ranavirus] | 95.5 | 44 | 7.17E-17 | 19 |
| k141_31219_flag1_multi49.6569_len3426 | W4-1 | 3426 | CBW45581.1 hypothetical protein, partial [Doctor fish virus] | 95.7 | 46 | 7.18E-17 | 234 |
| k141_108759_flag1_multi2.6060_len872 | W5-1 | 872 | UUY86251.1 hypothetical protein [Largemouth bass virus] | 100 | 40 | 7.22E-17 | 13 |
| k141_72955_flag0_multi1.0000_len281 | W2-3 | 281 | ABA41591.1 DNA-dependent DNA polymerase, partial [Largemouth bass virus] | 95.7 | 46 | 7.31E-17 | 0 |
| k141_145588_flag0_multi7.8519_len7752 | W5-1 | 7752 | XRL22821.1 hypothetical protein [Siniperca chuatsi ranavirus] | 100 | 46 | 7.35E-17 | 224 |
| k141_4266_flag1_multi5.0000_len847 | W5-1 | 847 | WAK75112.1 putative LITAF PIG7 possible membrane associated motif in LPS-induced tumor necrosis factor alpha factor [Mandarin fish ranavirus] | 94.7 | 38 | 7.39E-17 | 34 |
| k141_46904_flag1_multi3.7675_len799 | W1-2 | 799 | WAK75112.1 putative LITAF PIG7 possible membrane associated motif in LPS-induced tumor necrosis factor alpha factor [Mandarin fish ranavirus] | 100 | 36 | 7.72E-17 | 11 |

| | | | | | | | |
|---|---|---|---|---|---|---|---|
| k141_104030_flag1_multi4.0000_len1097 | W5-1 | 1097 | AYV88168.1 hypothetical protein [Mandarin fish ranavirus] | 79.6 | 54 | 7.83E-17 | 16 |
| k141_141326_flag1_multi1.0000_len679 | W5-1 | 679 | UUY86235.1 hypothetical protein [Largemouth bass virus] | 100 | 44 | 7.90E-17 | 23.86 |
| k141_98638_flag1_multi4.3280_len955 | W5-1 | 955 | QJE49137.1 hypothetical protein LMBV_074 [Largemouth bass virus] | 95.9 | 49 | 7.98E-17 | 14 |
| k141_20173_flag1_multi5.8591_len957 | W4-1 | 957 | AYV88211.1 hypothetical protein [Mandarin fish ranavirus] | 100 | 46 | 8.30E-17 | 17 |
| k141_105310_flag1_multi8.8218_len2565 | W5-1 | 2565 | WXI69548.1 hypothetical protein [Largemouth bass virus] | 97.8 | 45 | 8.48E-17 | 66 |
| k141_75013_flag1_multi1.6799_len419 | W5-1 | 419 | UUY86192.1 putative myristylated membrane protein [Largemouth bass virus] | 57.3 | 82 | 8.74E-17 | 1 |
| k141_33788_flag0_multi1.0000_len380 | W1-1 | 380 | QJE49100.1 hypothetical protein LMBV_037 [Largemouth bass virus] | 100 | 42 | 8.79E-17 | 0 |
| k141_39572_flag0_multi1.0000_len328 | W1-1 | 328 | QYU76036.1 helicase-like protein, partial [Koi ranavirus] | 97.7 | 44 | 8.86E-17 | 0 |
| k141_11156_flag1_multi4.9327_len587 | W5-1 | 587 | AIG51690.1 major capsid protein, partial [Koi ranavirus] | 89.6 | 48 | 8.91E-17 | 10 |
| k141_32255_flag0_multi1.0000_len259 | W1-1 | 259 | WEI28972.1 CTD-phosphotransferase [Largemouth bass virus] | 100 | 38 | 9.10E-17 | 0 |
| k141_75320_flag1_multi9.7875_len861 | W1-1 | 861 | QJE49101.1 hypothetical protein LMBV_038 [Largemouth bass virus] | 100 | 46 | 9.28E-17 | 18 |
| k141_145719_flag1_multi3.8642_len627 | W5-1 | 627 | UUY86266.1 hypothetical protein [Largemouth bass virus] | 100 | 40 | 9.53E-17 | 4 |
| k141_110897_flag1_multi2.0000_len636 | W5-1 | 636 | QIZ30887.1 major capsid protein, partial [Largemouth bass virus] | 100 | 39 | 9.85E-17 | 4 |
| k141_15468_flag1_multi1.0000_len498 | W5-1 | 498 | UUY86258.1 hypothetical protein [Largemouth bass virus] | 100 | 46 | 1.00E-16 | 11 |
| k141_2569_flag1_multi1.0000_len459 | W1-1 | 459 | XRB52768.1 P31K protein, partial [Largemouth bass virus] | 92.9 | 42 | 1.05E-16 | 0 |
| k141_21189_flag0_multi1.0000_len423 | W1-1 | 423 | XRL22790.1 hypothetical protein [Siniperca chuatsi ranavirus] | 97.7 | 43 | 1.06E-16 | 1 |
| k141_52319_flag1_multi6.4279_len1882 | W5-1 | 1882 | WHA35537.1 hypothetical protein MSRaV_49L [Micropterus salmoides ranavirus] | 100 | 41 | 1.09E-16 | 52 |
| k141_16876_flag1_multi8.8906_len1457 | W2-3 | 1457 | WHA35533.1 putative DNA dependent RNA polymerase A subunit [Micropterus salmoides ranavirus] | 100 | 45 | 1.12E-16 | 40 |
| k141_79075_flag1_multi9.9304_len9327 | W5-1 | 9327 | UUY86267.1 ribonucleotide reductase alpha subunit [Largemouth bass virus] | 100 | 46 | 1.12E-16 | 299 |
| k141_85909_flag1_multi1.0000_len506 | W5-1 | 506 | UUY86227.1 hypothetical protein [Largemouth bass virus] | 97.6 | 41 | 1.13E-16 | 17.96 |
| k141_100184_flag1_multi13.0000_len1264 | W1-1 | 1264 | QJE49215.1 putative orf58-like protein [Largemouth bass virus] | 100 | 39 | 1.16E-16 | 62 |
| k141_166990_flag1_multi4.8209_len610 | W5-1 | 610 | QJE49119.1 hypothetical protein LMBV_056 [Largemouth bass virus] | 100 | 40 | 1.19E-16 | 10 |
| k141_41563_flag1_multi1.0000_len767 | W5-1 | 767 | UUY86261.1 hypothetical protein [Largemouth bass virus] | 97.9 | 47 | 1.19E-16 | 9 |
| k141_16896_flag0_multi1.0000_len317 | W4-1 | 317 | UUY86235.1 hypothetical protein [Largemouth bass virus] | 90.9 | 44 | 1.21E-16 | 2 |
| k141_37862_flag1_multi4.9411_len1703 | W4-1 | 1703 | WEI29006.1 putative 2-cysteine adaptor domain protein [Largemouth bass virus] | 100 | 46 | 1.21E-16 | 51.45 |
| k141_65113_flag0_multi1.0000_len282 | W5-1 | 282 | UUY86218.1 hypothetical protein [Largemouth bass virus] | 97.3 | 37 | 1.25E-16 | 0 |
| k141_149610_flag0_multi59.1296_len8706 | W5-1 | 8706 | UUY86199.1 hypothetical protein [Largemouth bass virus] | 86.7 | 45 | 1.26E-16 | 1636 |
| k141_101341_flag1_multi2.0000_len708 | W6-1 | 708 | UUY86241.1 hypothetical protein [Largemouth bass virus] | 100 | 35 | 1.28E-16 | 5 |
| k141_28902_flag1_multi3.1193_len560 | W4-2 | 560 | AAC79876.1 viral core protein, partial [Labroides dimidatus ranavirus] | 97.4 | 38 | 1.32E-16 | 5 |
| k141_55374_flag1_multi3.4992_len1415 | W5-1 | 1415 | AIG51690.1 major capsid protein, partial [Koi ranavirus] | 100 | 46 | 1.33E-16 | 27 |
| k141_28670_flag1_multi8.6878_len756 | W5-1 | 756 | AYV88191.1 hypothetical protein [Mandarin fish ranavirus] | 97.5 | 40 | 1.41E-16 | 15 |
| k141_21924_flag1_multi2.9193_len364 | W5-1 | 364 | UUY86258.1 hypothetical protein [Largemouth bass virus] | 95.2 | 42 | 1.43E-16 | 0 |
| k141_89977_flag1_multi3.8313_len390 | W5-1 | 390 | UUY86258.1 hypothetical protein [Largemouth bass virus] | 100 | 43 | 1.44E-16 | 3 |
| k141_105718_flag0_multi1.0000_len464 | W5-1 | 464 | WAK75112.1 putative LITAF PIG7 possible membrane associated motif in LPS-induced tumor necrosis factor alpha factor [Mandarin fish ranavirus] | 97.2 | 36 | 1.52E-16 | 1 |
| k141_125792_flag1_multi3.0000_len620 | W5-1 | 620 | AYV88120.1 putative p31K protein [Mandarin fish ranavirus] | 92.9 | 42 | 1.53E-16 | 5 |
| k141_80241_flag1_multi20.5864_len2235 | W5-1 | 2235 | QJE49066.1 hypothetical protein LMBV_003 [Largemouth bass virus] | 100 | 46 | 1.54E-16 | 96 |
| k141_40241_flag1_multi13.8005_len1314 | W4-1 | 1314 | QJE49095.1 hypothetical protein LMBV_032 [Largemouth bass virus] | 100 | 45 | 1.57E-16 | 58 |

| | | | | | | | |
|---|---|---|---|---|---|---|---|
| k141_42207_flag1_multi10.6974_len802 | W4-1 | 802 | AYV88179.2 putative DNA dependent RNA polymerase II second largest subunit [Mandarin fish ranavirus] | 92 | 50 | 1.60E-16 | 27 |
| k141_43914_flag1_multi1.0000_len314 | W5-1 | 314 | QYU76034.1 putative myristylated membrane protein, partial [Koi ranavirus] | 100 | 44 | 1.61E-16 | 0 |
| k141_116521_flag1_multi1.5312_len1374 | W5-1 | 1374 | AIG51690.1 major capsid protein, partial [Koi ranavirus] | 100 | 44 | 1.62E-16 | 19 |
| k141_198576_flag1_multi1.0000_len335 | W5-3 | 335 | WEI28972.1 CTD-phosphotransferase [Largemouth bass virus] | 89.1 | 46 | 1.79E-16 | 0 |
| k141_83004_flag1_multi3.0000_len458 | W5-1 | 458 | WEI29006.1 putative 2-cysteine adaptor domain protein [Largemouth bass virus] | 90.9 | 44 | 1.80E-16 | 3 |
| k141_6841_flag1_multi38.5964_len1355 | W5-1 | 1355 | QJE49137.1 hypothetical protein LMBV_074 [Largemouth bass virus] | 92 | 50 | 1.82E-16 | 165 |
| k141_4697_flag1_multi18.8475_len3800 | W5-1 | 3800 | QYU76034.1 putative myristylated membrane protein, partial [Koi ranavirus] | 100 | 46 | 1.82E-16 | 226 |
| k141_117638_flag0_multi1.0000_len230 | W5-1 | 230 | UUY86218.1 hypothetical protein [Largemouth bass virus] | 97.3 | 37 | 1.83E-16 | 0 |
| k141_160227_flag1_multi1.0000_len420 | W5-1 | 420 | XPZ21261.1 hypothetical protein MRVORF002 [Mandarin fish ranavirus] | 95.3 | 43 | 1.89E-16 | 2 |
| k141_133_flag0_multi52.6393_len385 | W5-1 | 385 | UVF58793.1 MAG: DNA polymerase [Halichoeres melanurus ranavirus] | 97.6 | 42 | 2.02E-16 | 0 |
| k141_42937_flag1_multi1.0000_len517 | W5-1 | 517 | WEI28972.1 CTD-phosphotransferase [Largemouth bass virus] | 97.5 | 40 | 2.03E-16 | 0 |
| k141_12442_flag0_multi20.6203_len457 | W5-1 | 457 | UUY86235.1 hypothetical protein [Largemouth bass virus] | 100 | 41 | 2.14E-16 | 1 |
| k141_166405_flag1_multi2.1808_len600 | W5-1 | 600 | UUY86235.1 hypothetical protein [Largemouth bass virus] | 95.6 | 45 | 2.14E-16 | 7 |
| k141_109320_flag1_multi2.6777_len563 | W1-1 | 563 | UVF58785.1 MAG: major capsid protein [Halichoeres melanurus ranavirus] | 100 | 46 | 2.21E-16 | 6 |
| k141_25017_flag1_multi5.0229_len926 | W5-1 | 926 | UVF58785.1 MAG: major capsid protein [Halichoeres melanurus ranavirus] | 97.9 | 48 | 2.24E-16 | 16 |
| k141_30241_flag1_multi5.7862_len562 | W4-1 | 562 | WHA35533.1 putative DNA dependent RNA polymerase A subunit [Micropterus salmoides ranavirus] | 100 | 45 | 2.28E-16 | 12 |
| k141_108736_flag1_multi6.6396_len990 | W5-1 | 990 | QJE49121.1 hypothetical protein LMBV_058 [Largemouth bass virus] | 81.5 | 54 | 2.44E-16 | 11 |
| k141_65110_flag0_multi1.0000_len282 | W5-1 | 282 | UUY86218.1 hypothetical protein [Largemouth bass virus] | 90 | 40 | 2.50E-16 | 0 |
| k141_85666_flag1_multi7.0861_len2953 | W5-1 | 2953 | AAC79876.1 viral core protein, partial [Labroides dimidatus ranavirus] | 93 | 43 | 2.53E-16 | 68 |
| k141_22524_flag0_multi58.0999_len1623 | W6-1 | 1623 | XRL22821.1 hypothetical protein [Siniperca chuatsi ranavirus] | 100 | 46 | 2.69E-16 | 320 |
| k141_58615_flag1_multi5.5101_len488 | W5-3 | 488 | UUY86197.1 hypothetical protein [Largemouth bass virus] | 91.5 | 47 | 2.72E-16 | 27 |
| k141_47628_flag1_multi1.0000_len1001 | W5-1 | 1001 | AYV88120.1 putative p31K protein [Mandarin fish ranavirus] | 100 | 46 | 2.75E-16 | 84 |
| k141_153913_flag1_multi1.0000_len387 | W5-1 | 387 | WHA35533.1 putative DNA dependent RNA polymerase A subunit [Micropterus salmoides ranavirus] | 95.7 | 46 | 2.81E-16 | 1 |
| k141_78210_flag1_multi5.7970_len683 | W5-1 | 683 | AIG51690.1 major capsid protein, partial [Koi ranavirus] | 100 | 43 | 2.83E-16 | 11 |
| k141_167057_flag1_multi0.9750_len341 | W5-1 | 341 | QJE49096.1 putative D5 family NTPase/ATPase [Largemouth bass virus] | 89.1 | 46 | 2.97E-16 | 1 |
| k141_18651_flag1_multi5.8541_len614 | W2-1 | 614 | AIG51690.1 major capsid protein, partial [Koi ranavirus] | 58.8 | 85 | 2.98E-16 | 12 |
| k141_62962_flag1_multi6.8494_len1004 | W5-1 | 1004 | UUY86261.1 hypothetical protein [Largemouth bass virus] | 100 | 46 | 2.99E-16 | 31 |
| k141_100469_flag1_multi10.9806_len8382 | W5-1 | 8382 | UUY86258.1 hypothetical protein [Largemouth bass virus] | 100 | 44 | 3.11E-16 | 314 |
| k141_106834_flag1_multi1.0000_len453 | W5-1 | 453 | WEI29006.1 putative 2-cysteine adaptor domain protein [Largemouth bass virus] | 88.6 | 44 | 3.12E-16 | 0 |
| k141_158706_flag0_multi9.7062_len1424 | W5-1 | 1424 | QJE49125.1 hypothetical protein LMBV_062 [Largemouth bass virus] | 100 | 43 | 3.18E-16 | 35 |
| k141_154795_flag1_multi4.6090_len788 | W5-1 | 788 | QJE49101.1 hypothetical protein LMBV_038 [Largemouth bass virus] | 93.3 | 45 | 3.21E-16 | 6 |
| k141_94107_flag1_multi123.0177_len649 | W2-1 | 649 | UUY86263.1 hypothetical protein [Largemouth bass virus] | 97.6 | 41 | 3.22E-16 | 372 |
| k141_46797_flag1_multi4.8793_len870 | W5-1 | 870 | UUY86261.1 hypothetical protein [Largemouth bass virus] | 100 | 46 | 3.32E-16 | 9 |
| k141_86678_flag1_multi1.0000_len343 | W1-1 | 343 | WHU98721.1 MCP, partial [Hybrid snakehead ranavirus] | 100 | 39 | 3.39E-16 | 0 |
| k141_69031_flag1_multi11.6427_len849 | W4-1 | 849 | UVF58785.1 MAG: major capsid protein [Halichoeres melanurus ranavirus] | 100 | 46 | 3.42E-16 | 30 |
| k141_77870_flag1_multi2.5625_len781 | W1-1 | 781 | QYU76034.1 putative myristylated membrane protein, partial [Koi ranavirus] | 100 | 46 | 3.44E-16 | 5 |
| k141_64044_flag1_multi6.5225_len430 | W4-1 | 430 | QJE49073.1 putative p31K protein [Largemouth bass virus] | 75.8 | 62 | 3.59E-16 | 4 |

| | | | | | | | |
|---|---|---|---|---|---|---|---|
| k141_87942_flag1_multi4.1714_len1314 | W1-1 | 1314 | WAK75094.1 hypothetical protein [Mandarin fish ranavirus] | 100 | 38 | 3.62E-16 | 22 |
| k141_68702_flag1_multi5.2853_len1126 | W5-1 | 1126 | QJE49141.1 putative NTPase [Largemouth bass virus] | 94 | 50 | 3.66E-16 | 19 |
| k141_8868_flag0_multi18.2157_len1504 | W1-1 | 1504 | UUY86267.1 ribonucleotide reductase alpha subunit [Largemouth bass virus] | 93.5 | 46 | 3.81E-16 | 67 |
| k141_44496_flag1_multi2.9300_len798 | W5-1 | 798 | AYV88176.1 putative tumor necrosis factor receptor [Mandarin fish ranavirus] | 100 | 46 | 3.90E-16 | 7 |
| k141_43210_flag1_multi1.0000_len998 | W5-1 | 998 | UUY86197.1 hypothetical protein [Largemouth bass virus] | 100 | 46 | 4.11E-16 | 27.94 |
| k141_106802_flag1_multi2.9847_len926 | W2-1 | 926 | AYV88131.1 putative eIF-2 alpha-like protein [Mandarin fish ranavirus] | 100 | 41 | 4.14E-16 | 13 |
| k141_39063_flag0_multi52.4673_len860 | W2-1 | 860 | UUY86261.1 hypothetical protein [Largemouth bass virus] | 100 | 46 | 4.24E-16 | 99 |
| k141_75661_flag1_multi9.3790_len579 | W4-1 | 579 | UUY86258.1 hypothetical protein [Largemouth bass virus] | 90.2 | 51 | 4.30E-16 | 14 |
| k141_87876_flag1_multi4.0000_len616 | W1-1 | 616 | AFD96401.1 major capsid protein, partial [Largemouth bass virus] | 97.4 | 39 | 4.35E-16 | 7 |
| k141_38201_flag1_multi2.9198_len490 | W5-1 | 490 | UUY86258.1 hypothetical protein [Largemouth bass virus] | 97.7 | 44 | 4.43E-16 | 3 |
| k141_40314_flag1_multi5.9178_len1540 | W5-1 | 1540 | QJE49114.1 hypothetical protein LMBV_051 [Largemouth bass virus] | 100 | 37 | 4.45E-16 | 25 |
| k141_4680_flag1_multi3.9355_len1210 | W6-1 | 1210 | AIG51690.1 major capsid protein, partial [Koi ranavirus] | 100 | 46 | 4.55E-16 | 14 |
| k141_53383_flag0_multi1.0000_len403 | W4-1 | 403 | AIG51690.1 major capsid protein, partial [Koi ranavirus] | 97.8 | 45 | 4.61E-16 | 6.09 |
| k141_136490_flag0_multi1.0000_len548 | W1-1 | 548 | QJE49088.1 hypothetical protein LMBV_025 [Largemouth bass virus] | 97.9 | 47 | 4.75E-16 | 3.06 |
| k141_36075_flag0_multi87.6538_len2507 | W5-1 | 2507 | UUY86256.1 putative orf58-like protein [Largemouth bass virus] | 97.9 | 47 | 4.88E-16 | 745 |
| k141_161425_flag1_multi1.0000_len640 | W5-1 | 640 | AYV88120.1 putative p31K protein [Mandarin fish ranavirus] | 100 | 39 | 4.95E-16 | 14.78 |
| k141_104994_flag1_multi23.7164_len10848 | W2-3 | 10848 | QJE49112.1 hypothetical protein LMBV_049 [Largemouth bass virus] | 97.5 | 40 | 5.03E-16 | 768 |
| k141_38255_flag1_multi2.0000_len458 | W6-1 | 458 | AYV88118.1 hypothetical protein [Mandarin fish ranavirus] | 94.4 | 36 | 5.15E-16 | 3 |
| k141_75772_flag1_multi5.0000_len557 | W4-1 | 557 | WHA35533.1 putative DNA dependent RNA polymerase A subunit [Micropterus salmoides ranavirus] | 72.2 | 54 | 5.40E-16 | 13 |
| k141_24784_flag1_multi2.6675_len544 | W5-1 | 544 | UUY86193.1 hypothetical protein [Largemouth bass virus] | 100 | 40 | 6.11E-16 | 2 |
| k141_81114_flag0_multi2.7500_len365 | W2-1 | 365 | UUY86199.1 hypothetical protein [Largemouth bass virus] | 100 | 34 | 6.17E-16 | 4.85 |
| k141_66120_flag1_multi14.0000_len734 | W1-1 | 734 | QJE49149.1 putative immediate early protein ICP-46 [Largemouth bass virus] | 100 | 39 | 6.18E-16 | 34 |
| k141_81051_flag1_multi3.0000_len514 | W6-1 | 514 | QJE49112.1 hypothetical protein LMBV_049 [Largemouth bass virus] | 88.1 | 42 | 6.18E-16 | 9 |
| k141_164556_flag1_multi23.9760_len1183 | W5-1 | 1183 | UUY86269.1 hypothetical protein [Largemouth bass virus] | 88.2 | 51 | 6.33E-16 | 338.56 |
| k141_1926_flag1_multi4.0000_len589 | W6-1 | 589 | QYU76034.1 putative myristylated membrane protein, partial [Koi ranavirus] | 93 | 43 | 6.41E-16 | 6 |
| k141_23457_flag0_multi1.6759_len610 | W1-1 | 610 | QJE49097.1 hypothetical protein LMBV_034 [Largemouth bass virus] | 100 | 37 | 6.51E-16 | 5 |
| k141_74976_flag0_multi1.0000_len361 | W5-1 | 361 | QJE49103.1 hypothetical protein LMBV_040 [Largemouth bass virus] | 87 | 46 | 6.59E-16 | 0 |
| k141_66465_flag1_multi13.7521_len1347 | W1-1 | 1347 | XPZ21295.1 putative RNAseIII [Mandarin fish ranavirus] | 100 | 47 | 6.60E-16 | 50 |
| k141_168652_flag1_multi11.9914_len1426 | W5-1 | 1426 | WAK75107.1 hypothetical protein [Mandarin fish ranavirus] | 100 | 38 | 7.66E-16 | 53 |
| k141_34419_flag1_multi1.0000_len495 | W5-1 | 495 | AYV88154.1 putative helicase-like protein [Mandarin fish ranavirus] | 88.6 | 44 | 7.77E-16 | 3 |
| k141_143761_flag0_multi1.0000_len563 | W5-1 | 563 | UUY86197.1 hypothetical protein [Largemouth bass virus] | 73.2 | 71 | 7.83E-16 | 6 |
| k141_127950_flag1_multi1.8535_len414 | W5-1 | 414 | UVF58785.1 MAG: major capsid protein [Halichoeres melanurus ranavirus] | 100 | 39 | 7.86E-16 | 1 |
| k141_7341_flag1_multi4.0109_len508 | W5-1 | 508 | UUY86271.1 hypothetical protein [Largemouth bass virus] | 100 | 37 | 7.98E-16 | 6 |
| k141_13310_flag0_multi3.4399_len907 | W4-1 | 907 | QIZ30887.1 major capsid protein, partial [Largemouth bass virus] | 95.1 | 41 | 8.26E-16 | 14 |
| k141_100232_flag1_multi4.9078_len835 | W5-1 | 835 | QJE49101.1 hypothetical protein LMBV_038 [Largemouth bass virus] | 97.7 | 44 | 8.26E-16 | 11 |
| k141_57170_flag1_multi5.8940_len424 | W4-1 | 424 | UUY86229.1 putative DNA dependent RNA polymerase a subunit [Largemouth bass virus] | 92.9 | 42 | 8.31E-16 | 5 |
| k141_127803_flag1_multi3.9280_len683 | W5-1 | 683 | UUY86235.1 hypothetical protein [Largemouth bass virus] | 97.7 | 44 | 8.37E-16 | 6 |
| k141_24209_flag1_multi11.4000_len561 | W4-2 | 561 | UUY86266.1 hypothetical protein [Largemouth bass virus] | 83.3 | 48 | 8.42E-16 | 30 |

| | | | | | | | |
|---|---|---|---|---|---|---|---|
| k141_5784_flag0_multi3.0584_len603 | W1-1 | 603 | QJE49066.1 hypothetical protein LMBV_003 [Largemouth bass virus] | 97.6 | 41 | 8.56E-16 | 6 |
| k141_55999_flag1_multi3.0926_len670 | W5-1 | 670 | AYV88120.1 putative p31K protein [Mandarin fish ranavirus] | 100 | 46 | 8.59E-16 | 6 |
| k141_140227_flag1_multi1.0000_len330 | W6-2 | 330 | UUY86256.1 putative orf58-like protein [Largemouth bass virus] | 100 | 41 | 8.90E-16 | 4 |
| k141_46372_flag1_multi4.1533_len715 | W5-1 | 715 | QJE49096.1 putative D5 family NTPase/ATPase [Largemouth bass virus] | 100 | 40 | 8.97E-16 | 6 |
| k141_126455_flag0_multi1.0000_len281 | W1-1 | 281 | AYV88120.1 putative p31K protein [Mandarin fish ranavirus] | 95 | 40 | 9.01E-16 | 0 |
| k141_144289_flag1_multi1.0000_len455 | W5-1 | 455 | UUY86211.1 hypothetical protein [Largemouth bass virus] | 90.7 | 43 | 9.11E-16 | 1 |
| k141_75585_flag1_multi1.0000_len450 | W5-1 | 450 | UUY86197.1 hypothetical protein [Largemouth bass virus] | 91.3 | 46 | 9.24E-16 | 1 |
| k141_108207_flag1_multi5.8535_len926 | W5-1 | 926 | AYV88120.1 putative p31K protein [Mandarin fish ranavirus] | 100 | 46 | 9.36E-16 | 12 |
| k141_52472_flag1_multi2.0000_len488 | W5-1 | 488 | AYV88168.1 hypothetical protein [Mandarin fish ranavirus] | 100 | 40 | 9.61E-16 | 2 |
| k141_127690_flag0_multi4.1494_len837 | W5-1 | 837 | AYV88179.2 putative DNA dependent RNA polymerase II second largest subunit [Mandarin fish ranavirus] | 83.3 | 54 | 9.73E-16 | 11 |
| k141_125822_flag1_multi8.2604_len859 | W5-1 | 859 | AIG51690.1 major capsid protein, partial [Koi ranavirus] | 97.9 | 47 | 9.91E-16 | 25 |
| k141_115298_flag1_multi2.7186_len1136 | W5-1 | 1136 | UUY86261.1 hypothetical protein [Largemouth bass virus] | 100 | 46 | 1.02E-15 | 14 |
| k141_50361_flag1_multi20.4472_len2413 | W1-1 | 2413 | AYV88191.1 hypothetical protein [Mandarin fish ranavirus] | 95.1 | 41 | 1.04E-15 | 147 |
| k141_6834_flag1_multi2.6751_len1015 | W6-2 | 1015 | AYV88120.1 putative p31K protein [Mandarin fish ranavirus] | 100 | 42 | 1.06E-15 | 7 |
| k141_105571_flag1_multi8.1720_len1560 | W5-1 | 1560 | UUY86261.1 hypothetical protein [Largemouth bass virus] | 100 | 46 | 1.10E-15 | 50 |
| k141_6942_flag1_multi6.8635_len1767 | W5-1 | 1767 | UVF58790.1 MAG: myristylated membrane protein, partial [Halichoeres melanurus ranavirus] | 88.9 | 54 | 1.12E-15 | 34 |
| k141_122035_flag1_multi2.9197_len1212 | W1-1 | 1212 | AIG51690.1 major capsid protein, partial [Koi ranavirus] | 100 | 46 | 1.13E-15 | 19 |
| k141_33782_flag0_multi1.0000_len281 | W1-1 | 281 | QJE49100.1 hypothetical protein LMBV_037 [Largemouth bass virus] | 100 | 39 | 1.14E-15 | 0 |
| k141_54679_flag1_multi7.3305_len1300 | W1-1 | 1300 | UVF58785.1 MAG: major capsid protein [Halichoeres melanurus ranavirus] | 100 | 46 | 1.17E-15 | 39 |
| k141_66702_flag1_multi2.5219_len689 | W2-3 | 689 | UUY86241.1 hypothetical protein [Largemouth bass virus] | 94.6 | 37 | 1.17E-15 | 6.09 |
| k141_110257_flag1_multi4.0000_len497 | W5-1 | 497 | QJE49114.1 hypothetical protein LMBV_051 [Largemouth bass virus] | 100 | 33 | 1.21E-15 | 11 |
| k141_112428_flag1_multi10.0000_len495 | W6-2 | 495 | QJE49116.1 hypothetical protein LMBV_053 [Largemouth bass virus] | 100 | 40 | 1.27E-15 | 9 |
| k141_112_flag1_multi2.6594_len552 | W5-1 | 552 | WXI69548.1 hypothetical protein [Largemouth bass virus] | 100 | 39 | 1.30E-15 | 4 |
| k141_129064_flag0_multi15.5000_len207 | W5-1 | 207 | QYU76034.1 putative myristylated membrane protein, partial [Koi ranavirus] | 100 | 41 | 1.30E-15 | 0 |
| k141_17027_flag1_multi1.7265_len598 | W1-1 | 598 | UUY86238.1 hypothetical protein [Largemouth bass virus] | 97.6 | 42 | 1.32E-15 | 3 |
| k141_78995_flag1_multi4.0000_len1248 | W5-1 | 1248 | QJE49081.1 hypothetical protein LMBV_018 [Largemouth bass virus] | 100 | 39 | 1.38E-15 | 13 |
| k141_155921_flag1_multi3.0000_len442 | W5-1 | 442 | UUY86229.1 putative DNA dependent RNA polymerase a subunit [Largemouth bass virus] | 90.9 | 44 | 1.39E-15 | 2 |
| k141_41577_flag1_multi7.6785_len1012 | W5-1 | 1012 | AYV88120.1 putative p31K protein [Mandarin fish ranavirus] | 100 | 46 | 1.42E-15 | 23 |
| k141_138108_flag0_multi8.0000_len672 | W5-1 | 672 | WHA35678.1 hypothetical protein SCRaV_86R [Siniperca chuatsi ranavirus] | 100 | 39 | 1.50E-15 | 6 |
| k141_110057_flag0_multi26.0000_len393 | W5-1 | 393 | AYV88155.1 hypothetical protein [Mandarin fish ranavirus] | 97.4 | 39 | 1.53E-15 | 12 |
| k141_45913_flag1_multi1.9315_len477 | W5-1 | 477 | UUY86211.1 hypothetical protein [Largemouth bass virus] | 100 | 46 | 1.56E-15 | 2 |
| k141_180603_flag1_multi1.0000_len324 | W2-3 | 324 | QJE49090.1 hypothetical protein LMBV_027 [Largemouth bass virus] | 81.1 | 53 | 1.60E-15 | 2 |
| k141_55983_flag1_multi11.9138_len524 | W4-1 | 524 | CBW45581.1 hypothetical protein, partial [Doctor fish virus] | 92.7 | 41 | 1.60E-15 | 30 |
| k141_84771_flag1_multi5.0432_len604 | W4-1 | 604 | WAK75110.1 hypothetical protein [Mandarin fish ranavirus] | 87.8 | 49 | 1.68E-15 | 36 |
| k141_78658_flag1_multi11.6786_len897 | W6-2 | 897 | UUY86235.1 hypothetical protein [Largemouth bass virus] | 100 | 43 | 1.76E-15 | 37 |
| k141_48145_flag1_multi7.9842_len1220 | W2-1 | 1220 | UUY86199.1 hypothetical protein [Largemouth bass virus] | 87.2 | 39 | 1.78E-15 | 41 |
| k141_118315_flag0_multi147.0000_len313 | W1-1 | 313 | AYV88134.2 putative tyrosine kinase [Mandarin fish ranavirus] | 67.3 | 52 | 1.79E-15 | 2 |

| | | | | | | | |
|---|---|---|---|---|---|---|---|
| k141_157848_flag1_multi8.0000_len937 | W2-3 | 937 | AFD96401.1 major capsid protein, partial [Largemouth bass virus] | 88.9 | 45 | 1.82E-15 | 24 |
| k141_90551_flag0_multi31.0000_len591 | W1-1 | 591 | UVF58785.1 MAG: major capsid protein [Halichoeres melanurus ranavirus] | 81.6 | 49 | 1.85E-15 | 22.14 |
| k141_103137_flag1_multi8.0000_len2471 | W5-1 | 2471 | UUY86235.1 hypothetical protein [Largemouth bass virus] | 100 | 39 | 1.87E-15 | 74 |
| k141_99524_flag1_multi1.0000_len411 | W5-1 | 411 | AIG51690.1 major capsid protein, partial [Koi ranavirus] | 100 | 39 | 1.91E-15 | 10 |
| k141_92253_flag1_multi11.9085_len2906 | W1-1 | 2906 | UUY86269.1 hypothetical protein [Largemouth bass virus] | 92.2 | 51 | 1.92E-15 | 117 |
| k141_152238_flag1_multi3.4344_len850 | W5-1 | 850 | AYV88179.2 putative DNA dependent RNA polymerase II second largest subunit [Mandarin fish ranavirus] | 97.6 | 42 | 1.93E-15 | 12 |
| k141_130182_flag1_multi8.0000_len986 | W1-1 | 986 | UUY86211.1 hypothetical protein [Largemouth bass virus] | 79.2 | 53 | 1.95E-15 | 17 |
| k141_54460_flag1_multi2.9692_len660 | W5-1 | 660 | WEI28972.1 CTD-phosphotransferase [Largemouth bass virus] | 100 | 39 | 1.96E-15 | 1 |
| k141_136219_flag1_multi3.3630_len1097 | W5-1 | 1097 | WHA35534.1 hypothetical protein MSRaV_46R [Micropterus salmoides ranavirus] | 78.7 | 47 | 2.04E-15 | 10 |
| k141_22702_flag0_multi39.2678_len1403 | W4-2 | 1403 | UVF58785.1 MAG: major capsid protein [Halichoeres melanurus ranavirus] | 97.9 | 47 | 2.06E-15 | 69 |
| k141_163633_flag1_multi1.6573_len462 | W5-1 | 462 | UUY86267.1 ribonucleotide reductase alpha subunit [Largemouth bass virus] | 92.5 | 40 | 2.07E-15 | 1 |
| k141_70747_flag1_multi1.0000_len561 | W5-1 | 561 | WAK75112.1 putative LITAF PIG7 possible membrane associated motif in LPS-induced tumor necrosis factor alpha factor [Mandarin fish ranavirus] | 97 | 33 | 2.11E-15 | 5 |
| k141_129652_flag0_multi35.5711_len549 | W1-1 | 549 | WEI29006.1 putative 2-cysteine adaptor domain protein [Largemouth bass virus] | 90.5 | 42 | 2.16E-15 | 16 |
| k141_44485_flag1_multi1.0000_len359 | W5-1 | 359 | WAK75102.1 hypothetical protein [Mandarin fish ranavirus] | 97.4 | 38 | 2.30E-15 | 1 |
| k141_32197_flag1_multi8.5111_len910 | W5-1 | 910 | QJE49066.1 hypothetical protein LMBV_003 [Largemouth bass virus] | 100 | 42 | 2.31E-15 | 22 |
| k141_167803_flag1_multi18.7474_len1780 | W5-1 | 1780 | UUY86197.1 hypothetical protein [Largemouth bass virus] | 100 | 46 | 2.36E-15 | 113 |
| k141_10591_flag1_multi3.0000_len756 | W5-1 | 756 | WAK75112.1 putative LITAF PIG7 possible membrane associated motif in LPS-induced tumor necrosis factor alpha factor [Mandarin fish ranavirus] | 97.2 | 36 | 2.36E-15 | 8 |
| k141_146639_flag0_multi26.8824_len5004 | W5-1 | 5004 | WAK75075.1 hypothetical protein [Mandarin fish ranavirus] | 97.3 | 37 | 2.37E-15 | 368 |
| k141_43233_flag1_multi2.9547_len980 | W5-1 | 980 | UUY86197.1 hypothetical protein [Largemouth bass virus] | 100 | 45 | 2.38E-15 | 10 |
| k141_155617_flag1_multi17.8582_len4936 | W5-1 | 4936 | XRL22821.1 hypothetical protein [Siniperca chuatsi ranavirus] | 100 | 46 | 2.45E-15 | 232 |
| k141_181733_flag0_multi31.5222_len434 | W2-3 | 434 | QJE49116.1 hypothetical protein LMBV_053 [Largemouth bass virus] | 100 | 39 | 2.63E-15 | 12 |
| k141_89786_flag0_multi4.0331_len322 | W5-1 | 322 | QJE49153.1 hypothetical protein LMBV_004 [Largemouth bass virus] | 92.3 | 39 | 2.66E-15 | 2 |
| k141_146826_flag0_multi19.7231_len2709 | W5-1 | 2709 | UVF58785.1 MAG: major capsid protein [Halichoeres melanurus ranavirus] | 100 | 46 | 2.67E-15 | 184 |
| k141_84230_flag1_multi8.7401_len849 | W4-1 | 849 | XRL22790.1 hypothetical protein [Siniperca chuatsi ranavirus] | 100 | 46 | 2.69E-15 | 20 |
| k141_103216_flag1_multi3.6951_len1102 | W5-1 | 1102 | QJE49105.1 hypothetical protein LMBV_042 [Largemouth bass virus] | 97.8 | 46 | 2.77E-15 | 19 |
| k141_10876_flag1_multi12.5832_len760 | W5-1 | 760 | UUY86235.1 hypothetical protein [Largemouth bass virus] | 97.6 | 42 | 2.86E-15 | 35 |
| k141_32418_flag0_multi196.8737_len1028 | W6-1 | 1028 | QJE49101.1 hypothetical protein LMBV_038 [Largemouth bass virus] | 86 | 50 | 2.91E-15 | 444 |
| k141_5640_flag1_multi14.6767_len639 | W1-1 | 639 | UUY86261.1 hypothetical protein [Largemouth bass virus] | 100 | 44 | 3.11E-15 | 52 |
| k141_113678_flag1_multi27.7892_len3087 | W1-1 | 3087 | UUY86261.1 hypothetical protein [Largemouth bass virus] | 100 | 47 | 3.18E-15 | 328 |
| k141_28076_flag1_multi3.4578_len556 | W1-1 | 556 | WEI29006.1 putative 2-cysteine adaptor domain protein [Largemouth bass virus] | 100 | 38 | 3.21E-15 | 3 |
| k141_63035_flag0_multi21.1678_len576 | W4-1 | 576 | AAF64582.1 capsid protein, partial [Guppyfish iridovirus] | 95.2 | 42 | 3.27E-15 | 17 |
| k141_40059_flag1_multi11.7704_len1866 | W5-1 | 1866 | UUY86235.1 hypothetical protein [Largemouth bass virus] | 97.8 | 45 | 3.28E-15 | 68 |
| k141_9340_flag1_multi1.0000_len343 | W5-1 | 343 | AFD96401.1 major capsid protein, partial [Largemouth bass virus] | 94.6 | 37 | 3.33E-15 | 0 |
| k141_42036_flag1_multi3.7948_len1340 | W1-1 | 1340 | QIZ30887.1 major capsid protein, partial [Largemouth bass virus] | 100 | 39 | 3.34E-15 | 14 |
| k141_9435_flag0_multi1.0000_len378 | W2-1 | 378 | AIG51690.1 major capsid protein, partial [Koi ranavirus] | 100 | 40 | 3.35E-15 | 0 |
| k141_41886_flag1_multi2.3196_len554 | W5-1 | 554 | AYV88142.2 hypothetical protein [Mandarin fish ranavirus] | 97.4 | 39 | 3.36E-15 | 2 |
| k141_55668_flag1_multi3.0672_len1093 | W5-1 | 1093 | WAK75129.1 hypothetical protein [Mandarin fish ranavirus] | 97.2 | 36 | 3.43E-15 | 11 |

| | | | | | | | |
|---|---|---|---|---|---|---|---|
| k141_73199_flag1_multi24.3708_len586 | W5-1 | 586 | UUY86256.1 putative orf58-like protein [Largemouth bass virus] | 72.9 | 59 | 3.45E-15 | 43 |
| k141_86354_flag0_multi124.3179_len767 | W5-1 | 767 | UUY86261.1 hypothetical protein [Largemouth bass virus] | 100 | 46 | 3.46E-15 | 225 |
| k141_81185_flag1_multi2.8456_len536 | W4-1 | 536 | WEI29006.1 putative 2-cysteine adaptor domain protein [Largemouth bass virus] | 100 | 39 | 3.55E-15 | 104.68 |
| k141_114572_flag1_multi3.9189_len770 | W5-1 | 770 | AIG51690.1 major capsid protein, partial [Koi ranavirus] | 100 | 44 | 3.66E-15 | 12 |
| k141_166916_flag0_multi1.0000_len283 | W5-1 | 283 | UUY86256.1 putative orf58-like protein [Largemouth bass virus] | 88.4 | 43 | 3.68E-15 | 1 |
| k141_121140_flag1_multi13.6921_len1112 | W5-1 | 1112 | UUY86221.1 hypothetical protein [Largemouth bass virus] | 95.9 | 49 | 3.71E-15 | 61 |
| k141_156811_flag1_multi4.7617_len527 | W5-1 | 527 | UVF58790.1 MAG: myristylated membrane protein, partial [Halichoeres melanurus ranavirus] | 100 | 46 | 3.80E-15 | 8 |
| k141_116379_flag1_multi2.0000_len585 | W5-1 | 585 | UUY86235.1 hypothetical protein [Largemouth bass virus] | 97.5 | 40 | 3.85E-15 | 3 |
| k141_142285_flag0_multi2.2871_len545 | W5-1 | 545 | WEI29006.1 putative 2-cysteine adaptor domain protein [Largemouth bass virus] | 97.5 | 40 | 3.88E-15 | 2 |
| k141_102651_flag1_multi2.8614_len473 | W4-1 | 473 | AYV88120.1 putative p31K protein [Mandarin fish ranavirus] | 95.7 | 46 | 4.07E-15 | 2 |
| k141_114357_flag1_multi3.0000_len735 | W1-1 | 735 | UUY86192.1 putative myristylated membrane protein [Largemouth bass virus] | 97.5 | 40 | 4.15E-15 | 5 |
| k141_1439_flag1_multi4.9229_len1335 | W5-1 | 1335 | UVF58785.1 MAG: major capsid protein [Halichoeres melanurus ranavirus] | 85.5 | 55 | 4.16E-15 | 20 |
| k141_90880_flag1_multi1.8394_len527 | W6-1 | 527 | QJE49153.1 hypothetical protein LMBV_004 [Largemouth bass virus] | 97.5 | 40 | 4.28E-15 | 0 |
| k141_28945_flag1_multi69.4956_len825 | W5-1 | 825 | WAK75107.1 hypothetical protein [Mandarin fish ranavirus] | 66.7 | 60 | 4.45E-15 | 168 |
| k141_91450_flag1_multi2.0000_len683 | W6-1 | 683 | QJE49119.1 hypothetical protein LMBV_056 [Largemouth bass virus] | 92.9 | 42 | 4.47E-15 | 3 |
| k141_23621_flag1_multi4.0000_len534 | W3-1 | 534 | XRB52788.1 collagen-like protein [Largemouth bass virus] | 100 | 119 | 4.68E-15 | 4 |
| k141_69211_flag1_multi3.8623_len446 | W4-1 | 446 | QJE49137.1 hypothetical protein LMBV_074 [Largemouth bass virus] | 100 | 45 | 4.91E-15 | 3 |
| k141_94357_flag1_multi1.0000_len658 | W5-1 | 658 | UUY86261.1 hypothetical protein [Largemouth bass virus] | 100 | 42 | 4.93E-15 | 4 |
| k141_87463_flag1_multi20.0000_len5807 | W5-1 | 5807 | AYV88191.1 hypothetical protein [Mandarin fish ranavirus] | 78.2 | 55 | 4.99E-15 | 382 |
| k141_10514_flag1_multi27.3640_len2553 | W1-1 | 2553 | QJE49105.1 hypothetical protein LMBV_042 [Largemouth bass virus] | 93.9 | 49 | 5.20E-15 | 193 |
| k141_131396_flag1_multi2.3881_len543 | W5-1 | 543 | XRL22821.1 hypothetical protein [Siniperca chuatsi ranavirus] | 97.3 | 37 | 5.26E-15 | 6 |
| k141_28661_flag1_multi1.9482_len585 | W1-1 | 585 | AYV88211.1 hypothetical protein [Mandarin fish ranavirus] | 76.9 | 52 | 5.28E-15 | 8 |
| k141_6958_flag1_multi11.9226_len3696 | W5-1 | 3696 | UUY86235.1 hypothetical protein [Largemouth bass virus] | 65.3 | 72 | 5.52E-15 | 135 |
| k141_85767_flag1_multi25.2186_len992 | W5-1 | 992 | XRL22790.1 hypothetical protein [Siniperca chuatsi ranavirus] | 97.6 | 41 | 5.64E-15 | 123 |
| k141_122811_flag1_multi14.8862_len598 | W5-1 | 598 | WAK75112.1 putative LITAF PIG7 possible membrane associated motif in LPS-induced tumor necrosis factor alpha factor [Mandarin fish ranavirus] | 100 | 36 | 5.68E-15 | 14 |
| k141_7130_flag1_multi21.9464_len6019 | W1-1 | 6019 | QYU76034.1 putative myristylated membrane protein, partial [Koi ranavirus] | 100 | 43 | 5.80E-15 | 421 |
| k141_38405_flag1_multi4.7728_len546 | W2-1 | 546 | AYV88120.1 putative p31K protein [Mandarin fish ranavirus] | 97.8 | 46 | 5.85E-15 | 4 |
| k141_111796_flag1_multi10.8919_len955 | W2-1 | 955 | QJE49114.1 hypothetical protein LMBV_051 [Largemouth bass virus] | 89.7 | 39 | 5.89E-15 | 31 |
| k141_96171_flag1_multi7.9081_len1892 | W1-1 | 1892 | UVF58785.1 MAG: major capsid protein [Halichoeres melanurus ranavirus] | 100 | 46 | 5.99E-15 | 42 |
| k141_145880_flag1_multi2.0000_len456 | W5-1 | 456 | UUY86211.1 hypothetical protein [Largemouth bass virus] | 94.9 | 39 | 6.17E-15 | 3 |
| k141_46505_flag1_multi10.7105_len521 | W1-1 | 521 | UVF58785.1 MAG: major capsid protein [Halichoeres melanurus ranavirus] | 100 | 43 | 6.26E-15 | 94.01 |
| k141_5503_flag1_multi1.0000_len546 | W5-1 | 546 | AYV88125.2 hypothetical protein [Mandarin fish ranavirus] | 74 | 50 | 6.52E-15 | 1 |
| k141_62572_flag1_multi4.0000_len535 | W5-1 | 535 | AIG51690.1 major capsid protein, partial [Koi ranavirus] | 100 | 39 | 6.56E-15 | 10 |
| k141_148344_flag0_multi7.0000_len356 | W5-1 | 356 | AIG51690.1 major capsid protein, partial [Koi ranavirus] | 95 | 40 | 6.90E-15 | 2 |
| k141_148208_flag1_multi3.4785_len559 | W5-1 | 559 | QJE49105.1 hypothetical protein LMBV_042 [Largemouth bass virus] | 97.8 | 46 | 7.07E-15 | 2 |
| k141_30055_flag1_multi18.7668_len990 | W5-1 | 990 | AYV88191.1 hypothetical protein [Mandarin fish ranavirus] | 100 | 35 | 7.32E-15 | 59 |
| k141_80901_flag1_multi1.0000_len723 | W5-1 | 723 | WAK75138.1 hypothetical protein [Mandarin fish ranavirus] | 79.6 | 49 | 7.46E-15 | 7 |
| k141_101851_flag1_multi13.0000_len1861 | W5-1 | 1861 | QIZ30887.1 major capsid protein, partial [Largemouth bass virus] | 100 | 38 | 7.57E-15 | 121 |

| | | | | | | | |
|---|---|---|---|---|---|---|---|
| k141_126365_flag1_multi10.0000_len1303 | W1-1 | 1303 | QJE49107.1 hypothetical protein LMBV_044 [Largemouth bass virus] | 95.1 | 41 | 7.67E-15 | 32 |
| k141_102759_flag1_multi19.2896_len2544 | W4-1 | 2544 | UUY86197.1 hypothetical protein [Largemouth bass virus] | 100 | 46 | 7.72E-15 | 195 |
| k141_87333_flag1_multi8.7875_len1195 | W4-1 | 1195 | UUY86228.1 hypothetical protein [Largemouth bass virus] | 95.7 | 46 | 7.76E-15 | 30 |
| k141_35098_flag1_multi16.3749_len1240 | W4-1 | 1240 | AYV88120.1 putative p31K protein [Mandarin fish ranavirus] | 100 | 45 | 8.32E-15 | 43 |
| k141_126806_flag1_multi1.0000_len508 | W5-1 | 508 | AYV88191.1 hypothetical protein [Mandarin fish ranavirus] | 97.1 | 35 | 8.39E-15 | 2 |
| k141_62760_flag0_multi4.7784_len714 | W2-1 | 714 | QJE49077.1 hypothetical protein LMBV_014 [Largemouth bass virus] | 97.3 | 37 | 9.01E-15 | 11 |
| k141_161689_flag1_multi4.0000_len576 | W5-1 | 576 | UUY86258.1 hypothetical protein [Largemouth bass virus] | 100 | 39 | 9.08E-15 | 5 |
| k141_134357_flag1_multi2.5690_len800 | W5-1 | 800 | AYV88211.1 hypothetical protein [Mandarin fish ranavirus] | 78 | 59 | 9.09E-15 | 7 |
| k141_17857_flag1_multi9.7160_len870 | W5-1 | 870 | UVF58790.1 MAG: myristylated membrane protein, partial [Halichoeres melanurus ranavirus] | 95.8 | 48 | 9.36E-15 | 22 |
| k141_98350_flag1_multi1.0000_len328 | W5-1 | 328 | WAK75079.1 hypothetical protein [Mandarin fish ranavirus] | 88.9 | 36 | 9.36E-15 | 1 |
| k141_31954_flag1_multi16.0952_len939 | W4-1 | 939 | XPZ21261.1 hypothetical protein MRVORF002 [Mandarin fish ranavirus] | 95.3 | 43 | 9.57E-15 | 87 |
| k141_58459_flag0_multi1.0000_len372 | W2-1 | 372 | AYV88120.1 putative p31K protein [Mandarin fish ranavirus] | 86 | 43 | 1.01E-14 | 3 |
| k141_90533_flag0_multi24.0000_len542 | W5-1 | 542 | UUY86235.1 hypothetical protein [Largemouth bass virus] | 100 | 39 | 1.01E-14 | 12 |
| k141_43351_flag0_multi5.6113_len424 | W2-1 | 424 | AYV88191.1 hypothetical protein [Mandarin fish ranavirus] | 94.7 | 38 | 1.02E-14 | 0 |
| k141_95490_flag0_multi8.8854_len1467 | W5-1 | 1467 | UVF58785.1 MAG: major capsid protein [Halichoeres melanurus ranavirus] | 86.3 | 51 | 1.02E-14 | 36.03 |
| k141_28993_flag1_multi3.9434_len1465 | W1-1 | 1465 | QJE49090.1 hypothetical protein LMBV_027 [Largemouth bass virus] | 86.3 | 51 | 1.04E-14 | 25 |
| k141_3830_flag0_multi25.9719_len1491 | W1-1 | 1491 | WHA35503.1 putative myristylated membrane protein [Micropterus salmoides ranavirus] | 100 | 38 | 1.15E-14 | 116 |
| k141_130576_flag1_multi23.0000_len4857 | W5-1 | 4857 | QIZ30887.1 major capsid protein, partial [Largemouth bass virus] | 93.2 | 44 | 1.15E-14 | 441 |
| k141_166100_flag1_multi0.9947_len516 | W5-1 | 516 | AYV88120.1 putative p31K protein [Mandarin fish ranavirus] | 100 | 38 | 1.19E-14 | 0 |
| k141_122520_flag1_multi7.0647_len3943 | W1-1 | 3943 | AYV88143.1 hypothetical protein [Mandarin fish ranavirus] | 97.6 | 42 | 1.23E-14 | 430.45 |
| k141_56647_flag0_multi1.0000_len447 | W5-1 | 447 | WEI28966.1 hypothetical protein [Largemouth bass virus] | 97.2 | 36 | 1.23E-14 | 9.37 |
| k141_6468_flag1_multi7.9744_len960 | W4-1 | 960 | QJE49137.1 hypothetical protein LMBV_074 [Largemouth bass virus] | 100 | 39 | 1.31E-14 | 19 |
| k141_79280_flag0_multi1.0000_len258 | W2-1 | 258 | UUY86257.1 hypothetical protein [Largemouth bass virus] | 84.4 | 45 | 1.33E-14 | 1088.07 |
| k141_19841_flag1_multi3.0000_len575 | W5-1 | 575 | WAK75079.1 hypothetical protein [Mandarin fish ranavirus] | 88.9 | 36 | 1.37E-14 | 3 |
| k141_135122_flag0_multi69.6768_len1357 | W1-1 | 1357 | UVF58785.1 MAG: major capsid protein [Halichoeres melanurus ranavirus] | 100 | 46 | 1.43E-14 | 327 |
| k141_564_flag0_multi12.7120_len1273 | W4-1 | 1273 | AAF64582.1 capsid protein, partial [Guppyfish iridovirus] | 74.6 | 59 | 1.46E-14 | 44 |
| k141_7074_flag1_multi4.4321_len583 | W1-1 | 583 | AYV88130.2 putative ATPase-dependent protease [Mandarin fish ranavirus] | 67.8 | 59 | 1.51E-14 | 6 |
| k141_56386_flag0_multi1.0000_len307 | W5-1 | 307 | WEI29006.1 putative 2-cysteine adaptor domain protein [Largemouth bass virus] | 100 | 37 | 1.51E-14 | 0 |
| k141_91240_flag0_multi1.0000_len366 | W2-1 | 366 | AYV88191.1 hypothetical protein [Mandarin fish ranavirus] | 97.3 | 37 | 1.55E-14 | 5 |
| k141_78051_flag1_multi11.5831_len1242 | W5-1 | 1242 | AYV88120.1 putative p31K protein [Mandarin fish ranavirus] | 95.1 | 41 | 1.55E-14 | 50 |
| k141_31994_flag1_multi3.0024_len550 | W5-1 | 550 | XRB52763.1 hypothetical protein LMBV_6 [Largemouth bass virus] | 55.6 | 72 | 1.65E-14 | 1 |
| k141_105740_flag1_multi3.9326_len809 | W1-1 | 809 | AYV88120.1 putative p31K protein [Mandarin fish ranavirus] | 100 | 44 | 1.66E-14 | 12 |
| k141_30777_flag1_multi3.0000_len722 | W5-1 | 722 | AIG51690.1 major capsid protein, partial [Koi ranavirus] | 100 | 39 | 1.71E-14 | 3 |
| k141_96676_flag1_multi1.0000_len562 | W5-1 | 562 | QIZ30887.1 major capsid protein, partial [Largemouth bass virus] | 100 | 36 | 1.73E-14 | 5 |
| k141_139420_flag1_multi3.5446_len477 | W5-1 | 477 | QYU76034.1 putative myristylated membrane protein, partial [Koi ranavirus] | 94.9 | 39 | 1.74E-14 | 8 |
| k141_114329_flag1_multi7.2585_len965 | W2-3 | 965 | XRB52768.1 P31K protein, partial [Largemouth bass virus] | 84.4 | 45 | 1.76E-14 | 19 |
| k141_48765_flag1_multi7.4516_len2105 | W5-1 | 2105 | QJE49137.1 hypothetical protein LMBV_074 [Largemouth bass virus] | 95.3 | 43 | 1.79E-14 | 53 |
| k141_104557_flag1_multi3.0000_len473 | W1-1 | 473 | AIG51690.1 major capsid protein, partial [Koi ranavirus] | 100 | 38 | 1.80E-14 | 3 |

| | | | | | | | |
|---|---|---|---|---|---|---|---|
| k141_7929_flag1_multi1.0000_len619 | W5-1 | 619 | WAK75112.1 putative LITAF PIG7 possible membrane associated motif in LPS-induced tumor necrosis factor alpha factor [Mandarin fish ranavirus] | 100 | 34 | 1.87E-14 | 13 |
| k141_125133_flag1_multi1.0000_len365 | W1-1 | 365 | AIG51690.1 major capsid protein, partial [Koi ranavirus] | 90 | 40 | 2.01E-14 | 0 |
| k141_84644_flag1_multi7.5400_len1328 | W2-1 | 1328 | QJE49081.1 hypothetical protein LMBV_018 [Largemouth bass virus] | 97.4 | 38 | 2.05E-14 | 31 |
| k141_139099_flag1_multi89.8629_len15001 | W5-1 | 15001 | XRL22821.1 hypothetical protein [Siniperca chuatsi ranavirus] | 100 | 46 | 2.05E-14 | 4634.99 |
| k141_73739_flag0_multi1.0000_len391 | W2-1 | 391 | QJE49149.1 putative immediate early protein ICP-46 [Largemouth bass virus] | 88.6 | 44 | 2.07E-14 | 0 |
| k141_161790_flag0_multi20.3767_len510 | W5-1 | 510 | UUY86192.1 putative myristylated membrane protein [Largemouth bass virus] | 82.9 | 41 | 2.09E-14 | 5 |
| k141_93213_flag1_multi1.0000_len447 | W5-1 | 447 | QJE49153.1 hypothetical protein LMBV_004 [Largemouth bass virus] | 97.4 | 38 | 2.12E-14 | 16 |
| k141_48155_flag1_multi2.9276_len362 | W4-3 | 362 | XRL22821.1 hypothetical protein [Siniperca chuatsi ranavirus] | 100 | 35 | 2.18E-14 | 2 |
| k141_95393_flag1_multi3.8473_len593 | W2-3 | 593 | UUY86235.1 hypothetical protein [Largemouth bass virus] | 100 | 38 | 2.21E-14 | 6 |
| k141_117862_flag1_multi9.9451_len1617 | W1-1 | 1617 | WXI69525.1 caspase recruitment domain protein [Largemouth bass virus] | 97.6 | 42 | 2.22E-14 | 58 |
| k141_76232_flag1_multi2.8472_len704 | W1-1 | 704 | AYV88134.2 putative tyrosine kinase [Mandarin fish ranavirus] | 80.4 | 46 | 2.30E-14 | 4 |
| k141_65261_flag1_multi1.0000_len385 | W2-1 | 385 | QJE49125.1 hypothetical protein LMBV_062 [Largemouth bass virus] | 94.7 | 38 | 2.39E-14 | 0 |
| k141_151889_flag1_multi2.8060_len543 | W5-1 | 543 | WEI29006.1 putative 2-cysteine adaptor domain protein [Largemouth bass virus] | 97.3 | 37 | 2.39E-14 | 16 |
| k141_91173_flag0_multi1.0000_len453 | W5-1 | 453 | WAK75112.1 putative LITAF PIG7 possible membrane associated motif in LPS-induced tumor necrosis factor alpha factor [Mandarin fish ranavirus] | 94.1 | 34 | 2.43E-14 | 0 |
| k141_138955_flag1_multi5.0000_len653 | W5-1 | 653 | AYV88211.1 hypothetical protein [Mandarin fish ranavirus] | 92.9 | 42 | 2.44E-14 | 9 |
| k141_66705_flag1_multi4.0000_len606 | W5-1 | 606 | UUY86235.1 hypothetical protein [Largemouth bass virus] | 90.5 | 42 | 2.45E-14 | 8 |
| k141_63476_flag0_multi4.0000_len524 | W1-1 | 524 | AYV88120.1 putative p31K protein [Mandarin fish ranavirus] | 97.4 | 39 | 2.52E-14 | 7 |
| k141_75765_flag1_multi8.3839_len925 | W5-1 | 925 | WAK75093.1 hypothetical protein [Mandarin fish ranavirus] | 83 | 47 | 2.69E-14 | 16 |
| k141_2597_flag1_multi4.8595_len568 | W4-1 | 568 | AYV88120.1 putative p31K protein [Mandarin fish ranavirus] | 94.7 | 38 | 2.73E-14 | 7 |
| k141_61609_flag1_multi10.9160_len2283 | W5-1 | 2283 | QJE49068.1 hypothetical protein LMBV_005 [Largemouth bass virus] | 69.8 | 63 | 2.74E-14 | 66 |
| k141_77574_flag1_multi3.3590_len829 | W1-1 | 829 | QJE49149.1 putative immediate early protein ICP-46 [Largemouth bass virus] | 97.4 | 38 | 2.76E-14 | 11 |
| k141_46851_flag1_multi2.1571_len644 | W5-1 | 644 | WEI28972.1 CTD-phosphotransferase [Largemouth bass virus] | 76.4 | 55 | 2.77E-14 | 9 |
| k141_165999_flag1_multi1.0000_len323 | W5-1 | 323 | XRB52831.1 hypothetical protein LMBV_74 [Largemouth bass virus] | 97.4 | 38 | 2.81E-14 | 5 |
| k141_88214_flag1_multi4.0000_len699 | W5-1 | 699 | WHU98721.1 MCP, partial [Hybrid snakehead ranavirus] | 100 | 38 | 2.82E-14 | 4 |
| k141_34378_flag0_multi2.1779_len304 | W1-1 | 304 | ACT85833.1 major capsid protein, partial [Frog virus 3] | 82.1 | 39 | 2.84E-14 | 0 |
| k141_166452_flag1_multi2.7788_len349 | W5-1 | 349 | QJE49114.1 hypothetical protein LMBV_051 [Largemouth bass virus] | 100 | 31 | 2.93E-14 | 3 |
| k141_146467_flag1_multi1.0000_len557 | W5-1 | 557 | XRB52842.1 transcription factor TFIIB cyclin-like domain-containing protein, partial [Largemouth bass virus] | 80 | 50 | 3.03E-14 | 0 |
| k141_120443_flag1_multi2.5516_len538 | W1-1 | 538 | UUY86236.1 hypothetical protein [Largemouth bass virus] | 94.6 | 37 | 3.11E-14 | 7 |
| k141_21034_flag1_multi10.0000_len846 | W4-1 | 846 | WAK75102.1 hypothetical protein [Mandarin fish ranavirus] | 95 | 40 | 3.15E-14 | 25 |
| k141_72275_flag1_multi5.0000_len826 | W1-1 | 826 | QJE49105.1 hypothetical protein LMBV_042 [Largemouth bass virus] | 100 | 39 | 3.17E-14 | 10 |
| k141_37438_flag1_multi9.0000_len808 | W4-1 | 808 | AYV88211.1 hypothetical protein [Mandarin fish ranavirus] | 92.5 | 40 | 3.17E-14 | 25 |
| k141_76227_flag1_multi1.0000_len483 | W1-1 | 483 | UUY86241.1 hypothetical protein [Largemouth bass virus] | 97 | 33 | 3.21E-14 | 1 |
| k141_109368_flag1_multi5.2391_len601 | W5-1 | 601 | XRL22790.1 hypothetical protein [Siniperca chuatsi ranavirus] | 97.7 | 43 | 3.21E-14 | 14 |
| k141_771_flag0_multi1.0000_len518 | W5-1 | 518 | UUY86267.1 ribonucleotide reductase alpha subunit [Largemouth bass virus] | 75 | 52 | 3.23E-14 | 0 |
| k141_92596_flag1_multi3.5210_len617 | W4-1 | 617 | UUY86258.1 hypothetical protein [Largemouth bass virus] | 92.3 | 39 | 3.24E-14 | 9 |
| k141_10319_flag1_multi3.0000_len638 | W5-1 | 638 | WAK75072.1 hypothetical protein [Mandarin fish ranavirus] | 90.7 | 43 | 3.36E-14 | 4 |
| k141_21598_flag1_multi25.6235_len1602 | W4-1 | 1602 | AYV88176.1 putative tumor necrosis factor receptor [Mandarin fish ranavirus] | 97.4 | 38 | 3.39E-14 | 131 |

| | | | | | | | |
|---|---|---|---|---|---|---|---|
| k141_154422_flag0_multi1.0000_len458 | W5-1 | 458 | QJE49086.1 hypothetical protein LMBV_023 [Largemouth bass virus] | 87.2 | 39 | 3.52E-14 | 3 |
| k141_15707_flag1_multi2.9742_len1927 | W6-1 | 1927 | WAK75138.1 hypothetical protein [Mandarin fish ranavirus] | 80.9 | 47 | 3.92E-14 | 52 |
| k141_29951_flag0_multi3.4543_len480 | W5-1 | 480 | QJE49098.1 hypothetical protein LMBV_035 [Largemouth bass virus] | 91.7 | 36 | 3.94E-14 | 9 |
| k141_38722_flag1_multi16.6624_len4454 | W5-1 | 4454 | WAK75112.1 putative LITAF PIG7 possible membrane associated motif in LPS-induced tumor necrosis factor alpha factor [Mandarin fish ranavirus] | 64.1 | 64 | 3.98E-14 | 286 |
| k141_22876_flag1_multi9.0000_len2232 | W5-1 | 2232 | WAK75107.1 hypothetical protein [Mandarin fish ranavirus] | 100 | 34 | 4.00E-14 | 60 |
| k141_33978_flag1_multi6.9684_len711 | W1-1 | 711 | ALR73092.1 major capsid protein, partial [Box turtle ranavirus] | 86 | 43 | 4.07E-14 | 13 |
| k141_31057_flag0_multi67.6227_len520 | W1-1 | 520 | UUY86228.1 hypothetical protein [Largemouth bass virus] | 100 | 39 | 4.19E-14 | 250 |
| k141_113931_flag1_multi1.0000_len346 | W5-1 | 346 | UUY86258.1 hypothetical protein [Largemouth bass virus] | 94.6 | 37 | 4.40E-14 | 0 |
| k141_52130_flag1_multi4.0000_len488 | W5-1 | 488 | QJE49104.1 hypothetical protein LMBV_041 [Largemouth bass virus] | 100 | 35 | 4.55E-14 | 10 |
| k141_124547_flag1_multi1.0000_len829 | W5-1 | 829 | AYV88118.1 hypothetical protein [Mandarin fish ranavirus] | 94.3 | 35 | 4.68E-14 | 39 |
| k141_146850_flag1_multi16.8719_len4639 | W5-1 | 4639 | QIZ30887.1 major capsid protein, partial [Largemouth bass virus] | 100 | 38 | 5.15E-14 | 328 |
| k141_4542_flag1_multi2.9316_len1749 | W5-1 | 1749 | XRL22790.1 hypothetical protein [Siniperca chuatsi ranavirus] | 97.8 | 46 | 5.24E-14 | 43.21 |
| k141_40804_flag1_multi1.0000_len365 | W5-1 | 365 | AYV88210.2 putative proliferating cell nuclear antigen [Mandarin fish ranavirus] | 88.4 | 43 | 5.32E-14 | 0 |
| k141_71268_flag0_multi1.0000_len319 | W5-1 | 319 | UVF58785.1 MAG: major capsid protein [Halichoeres melanurus ranavirus] | 97.6 | 42 | 5.34E-14 | 0 |
| k141_64333_flag0_multi3.0000_len463 | W5-1 | 463 | UUY86256.1 putative orf58-like protein [Largemouth bass virus] | 100 | 37 | 5.45E-14 | 1 |
| k141_30423_flag1_multi7.0000_len729 | W2-1 | 729 | UVF58787.1 MAG: DNA-dependent RNA polymerase largest subunit, partial [Halichoeres melanurus ranavirus] | 100 | 38 | 5.51E-14 | 15 |
| k141_145426_flag1_multi1.9125_len404 | W5-1 | 404 | QYU76036.1 helicase-like protein, partial [Koi ranavirus] | 94.6 | 37 | 5.56E-14 | 3 |
| k141_6088_flag1_multi5.2128_len827 | W4-1 | 827 | QJE49080.1 hypothetical protein LMBV_017 [Largemouth bass virus] | 91.9 | 37 | 5.59E-14 | 8 |
| k141_153919_flag1_multi4.0000_len1095 | W5-1 | 1095 | QJE49203.1 hypothetical protein LMBV_054 [Largemouth bass virus] | 97.4 | 38 | 5.69E-14 | 11 |
| k141_119077_flag1_multi1.0000_len318 | W5-1 | 318 | AYV88211.1 hypothetical protein [Mandarin fish ranavirus] | 100 | 39 | 5.73E-14 | 0 |
| k141_148074_flag1_multi4.9582_len811 | W5-1 | 811 | AYV88191.1 hypothetical protein [Mandarin fish ranavirus] | 79.6 | 49 | 5.94E-14 | 9 |
| k141_116884_flag1_multi4.0000_len633 | W1-1 | 633 | QJE49102.1 putative DNA dependent RNA polymerase a subunit [Largemouth bass virus] | 100 | 39 | 5.95E-14 | 2 |
| k141_22012_flag1_multi24.2107_len1133 | W1-1 | 1133 | WHA35533.1 putative DNA dependent RNA polymerase A subunit [Micropterus salmoides ranavirus] | 100 | 44 | 5.98E-14 | 85 |
| k141_37158_flag1_multi2.9707_len550 | W6-3 | 550 | QJE49129.1 putative orf58-like protein [Largemouth bass virus] | 70.9 | 182 | 6.05E-14 | 13 |
| k141_1559_flag1_multi4.0000_len485 | W5-1 | 485 | UUY86235.1 hypothetical protein [Largemouth bass virus] | 100 | 38 | 6.22E-14 | 7 |
| k141_84241_flag1_multi2.0000_len494 | W5-1 | 494 | UUY86258.1 hypothetical protein [Largemouth bass virus] | 100 | 36 | 6.43E-14 | 1 |
| k141_155551_flag1_multi4.0000_len439 | W5-1 | 439 | QJE49105.1 hypothetical protein LMBV_042 [Largemouth bass virus] | 95.2 | 42 | 6.46E-14 | 4 |
| k141_88175_flag1_multi3.9704_len445 | W4-1 | 445 | UVF58785.1 MAG: major capsid protein [Halichoeres melanurus ranavirus] | 74.1 | 58 | 6.62E-14 | 1 |
| k141_60655_flag1_multi22.0000_len723 | W5-1 | 723 | AYV88191.1 hypothetical protein [Mandarin fish ranavirus] | 95 | 40 | 6.78E-14 | 62 |
| k141_56292_flag1_multi7.9757_len800 | W5-1 | 800 | QJE49137.1 hypothetical protein LMBV_074 [Largemouth bass virus] | 100 | 38 | 6.80E-14 | 11 |
| k141_85368_flag1_multi11.9602_len694 | W4-1 | 694 | QJE49149.1 putative immediate early protein ICP-46 [Largemouth bass virus] | 64.3 | 70 | 6.95E-14 | 19 |
| k141_127507_flag1_multi1.8324_len493 | W5-1 | 493 | AYV88186.1 hypothetical protein [Mandarin fish ranavirus] | 100 | 35 | 7.48E-14 | 6 |
| k141_10945_flag0_multi97.4330_len238 | W5-1 | 238 | QJE49065.1 hypothetical protein LMBV_002 [Largemouth bass virus] | 100 | 33 | 7.72E-14 | 1 |
| k141_19030_flag1_multi1.0000_len404 | W5-1 | 404 | UUY86241.1 hypothetical protein [Largemouth bass virus] | 97 | 33 | 7.73E-14 | 6 |
| k141_103611_flag1_multi17.8112_len8604 | W4-1 | 8604 | UUY86192.1 putative myristylated membrane protein [Largemouth bass virus] | 97.4 | 38 | 8.09E-14 | 537 |

| | | | | | | | |
|---|---|---|---|---|---|---|---|
| k141_168708_flag1_multi1.6146_len634 | W5-1 | 634 | WHA35533.1 putative DNA dependent RNA polymerase A subunit [Micropterus salmoides ranavirus] | 86 | 43 | 8.30E-14 | 3 |
| k141_54480_flag1_multi1.4397_len605 | W5-1 | 605 | WAK75107.1 hypothetical protein [Mandarin fish ranavirus] | 100 | 32 | 8.42E-14 | 3 |
| k141_25225_flag1_multi1.0000_len596 | W2-2 | 596 | UUY86235.1 hypothetical protein [Largemouth bass virus] | 57.1 | 84 | 8.62E-14 | 2 |
| k141_136392_flag1_multi2.3985_len550 | W5-1 | 550 | UUY86269.1 hypothetical protein [Largemouth bass virus] | 97.4 | 39 | 9.16E-14 | 5 |
| k141_132493_flag1_multi3.0000_len428 | W5-3 | 428 | AYV88122.1 putative 3-beta-hydroxy-delta-5-C27 steroid oxidoreductase-like protein [Mandarin fish ranavirus] | 97.1 | 34 | 9.36E-14 | 4 |
| k141_32213_flag1_multi2.3896_len390 | W1-1 | 390 | AYV88183.1 hypothetical protein [Mandarin fish ranavirus] | 100 | 33 | 9.62E-14 | 1 |
| k141_51468_flag1_multi4.0000_len655 | W4-1 | 655 | QJE49114.1 hypothetical protein LMBV_051 [Largemouth bass virus] | 52.5 | 80 | 9.68E-14 | 17 |
| k141_65254_flag1_multi7.5257_len1056 | W2-1 | 1056 | AYV88179.2 putative DNA dependent RNA polymerase II second largest subunit [Mandarin fish ranavirus] | 87.2 | 47 | 9.70E-14 | 30 |
| k141_105890_flag0_multi7.9299_len5734 | W5-1 | 5734 | QYU76034.1 putative myristylated membrane protein, partial [Koi ranavirus] | 88.9 | 45 | 9.75E-14 | 144 |
| k141_144546_flag1_multi5.0000_len2837 | W5-1 | 2837 | QYU76031.1 neurofilament triplet H1-like protein, partial [Koi ranavirus] | 100 | 39 | 9.92E-14 | 41 |
| k141_142796_flag1_multi1.0000_len327 | W5-1 | 327 | WAK75112.1 putative LITAF PIG7 possible membrane associated motif in LPS-induced tumor necrosis factor alpha factor [Mandarin fish ranavirus] | 100 | 31 | 1.02E-13 | 3 |
| k141_256390_flag1_multi1.0000_len669 | W5-3 | 669 | AIG51690.1 major capsid protein, partial [Koi ranavirus] | 100 | 38 | 1.03E-13 | 8 |
| k141_181161_flag1_multi4.0000_len713 | W2-3 | 713 | AIG51690.1 major capsid protein, partial [Koi ranavirus] | 65.2 | 66 | 1.06E-13 | 9 |
| k141_99680_flag1_multi2.0000_len645 | W5-1 | 645 | AYV88118.1 hypothetical protein [Mandarin fish ranavirus] | 97.1 | 35 | 1.13E-13 | 3 |
| k141_37910_flag1_multi3.0000_len894 | W5-1 | 894 | WEI29006.1 putative 2-cysteine adaptor domain protein [Largemouth bass virus] | 100 | 37 | 1.15E-13 | 10 |
| k141_38857_flag1_multi7.1707_len639 | W4-1 | 639 | UVF58785.1 MAG: major capsid protein [Halichoeres melanurus ranavirus] | 100 | 39 | 1.16E-13 | 18 |
| k141_26982_flag1_multi21.5900_len1319 | W1-1 | 1319 | AYV88120.1 putative p31K protein [Mandarin fish ranavirus] | 95.7 | 46 | 1.17E-13 | 82 |
| k141_15110_flag1_multi7.2468_len1405 | W5-1 | 1405 | UVF58785.1 MAG: major capsid protein [Halichoeres melanurus ranavirus] | 97.8 | 46 | 1.21E-13 | 34 |
| k141_20921_flag1_multi8.5950_len583 | W5-1 | 583 | QJE49109.1 hypothetical protein LMBV_046 [Largemouth bass virus] | 89.2 | 37 | 1.26E-13 | 19 |
| k141_79780_flag1_multi7.5296_len902 | W5-1 | 902 | UUY86261.1 hypothetical protein [Largemouth bass virus] | 95.8 | 48 | 1.26E-13 | 29 |
| k141_123331_flag1_multi4.2519_len788 | W5-1 | 788 | UUY86235.1 hypothetical protein [Largemouth bass virus] | 100 | 40 | 1.27E-13 | 9 |
| k141_96756_flag0_multi15.7823_len1657 | W5-1 | 1657 | QJE49104.1 hypothetical protein LMBV_041 [Largemouth bass virus] | 100 | 39 | 1.45E-13 | 40 |
| k141_37699_flag1_multi4.8442_len603 | W1-1 | 603 | UUY86229.1 putative DNA dependent RNA polymerase a subunit [Largemouth bass virus] | 100 | 38 | 1.55E-13 | 13 |
| k141_86939_flag1_multi13.7693_len995 | W5-1 | 995 | AIG51690.1 major capsid protein, partial [Koi ranavirus] | 100 | 42 | 1.59E-13 | 48 |
| k141_47503_flag1_multi5.0000_len1031 | W5-1 | 1031 | AYV88191.1 hypothetical protein [Mandarin fish ranavirus] | 97.2 | 36 | 1.59E-13 | 13 |
| k141_42311_flag1_multi1.0000_len382 | W5-1 | 382 | AYV88120.1 putative p31K protein [Mandarin fish ranavirus] | 79.6 | 49 | 1.61E-13 | 0 |
| k141_65913_flag1_multi4.9501_len1003 | W5-1 | 1003 | QJE49079.1 hypothetical protein LMBV_016 [Largemouth bass virus] | 97.3 | 37 | 1.64E-13 | 28 |
| k141_63350_flag1_multi10.4384_len871 | W1-1 | 871 | QJE49146.1 putative thiol oxidoreductase [Largemouth bass virus] | 75.5 | 49 | 1.67E-13 | 10 |
| k141_74649_flag1_multi6.1006_len1622 | W5-1 | 1622 | UUY86256.1 putative orf58-like protein [Largemouth bass virus] | 100 | 39 | 1.74E-13 | 49 |
| k141_978_flag0_multi3.9216_len830 | W5-1 | 830 | QJE49149.1 putative immediate early protein ICP-46 [Largemouth bass virus] | 88.6 | 44 | 1.78E-13 | 7 |
| k141_153818_flag1_multi1.0000_len467 | W2-3 | 467 | AAK54493.1 RSIV DNA polymerase-like protein, partial [Regina ranavirus] | 61.4 | 57 | 1.83E-13 | 3 |
| k141_35512_flag1_multi9.9201_len704 | W5-1 | 704 | UUY86269.1 hypothetical protein [Largemouth bass virus] | 97.5 | 40 | 1.93E-13 | 12 |
| k141_166952_flag1_multi1.0000_len775 | W5-1 | 775 | WHA35533.1 putative DNA dependent RNA polymerase A subunit [Micropterus salmoides ranavirus] | 97.6 | 42 | 1.93E-13 | 14 |
| k141_141808_flag1_multi1.0000_len914 | W5-1 | 914 | AIG51690.1 major capsid protein, partial [Koi ranavirus] | 100 | 38 | 1.98E-13 | 12 |

| | | | | | | | |
|---|---|---|---|---|---|---|---|
| k141_163940_flag0_multi4.8870_len318 | W5-1 | 318 | AYV88211.1 hypothetical protein [Mandarin fish ranavirus] | 83 | 47 | 1.99E-13 | 68 |
| k141_25768_flag1_multi1.0000_len462 | W5-1 | 462 | AYV88120.1 putative p31K protein [Mandarin fish ranavirus] | 53.5 | 86 | 2.00E-13 | 3 |
| k141_17680_flag1_multi8.0000_len632 | W4-1 | 632 | UUY86229.1 putative DNA dependent RNA polymerase a subunit [Largemouth bass virus] | 100 | 39 | 2.04E-13 | 18 |
| k141_97999_flag1_multi1.0000_len374 | W4-1 | 374 | QJE49155.1 hypothetical protein LMBV_006 [Largemouth bass virus] | 97.4 | 39 | 2.12E-13 | 3 |
| k141_63419_flag1_multi4.0000_len564 | W1-1 | 564 | AIG51690.1 major capsid protein, partial [Koi ranavirus] | 97.4 | 38 | 2.14E-13 | 2 |
| k141_180133_flag1_multi4.0000_len853 | W2-3 | 853 | QJE49112.1 hypothetical protein LMBV_049 [Largemouth bass virus] | 87.5 | 40 | 2.19E-13 | 13 |
| k141_31333_flag0_multi1.4741_len411 | W5-1 | 411 | AYV88120.1 putative p31K protein [Mandarin fish ranavirus] | 85.4 | 48 | 2.23E-13 | 1 |
| k141_77917_flag1_multi4.8535_len769 | W5-1 | 769 | AYV88149.2 hypothetical protein [Mandarin fish ranavirus] | 100 | 35 | 2.35E-13 | 8 |
| k141_161261_flag1_multi3.0000_len595 | W5-1 | 595 | AAC79876.1 viral core protein, partial [Labroides dimidatus ranavirus] | 100 | 33 | 2.38E-13 | 2 |
| k141_108628_flag1_multi8.7738_len698 | W5-1 | 698 | QJE49215.1 putative orf58-like protein [Largemouth bass virus] | 97.4 | 38 | 2.49E-13 | 18 |
| k141_35417_flag1_multi3.0000_len406 | W5-1 | 406 | UUY86192.1 putative myristylated membrane protein [Largemouth bass virus] | 100 | 35 | 2.55E-13 | 5 |
| k141_112524_flag0_multi7.6980_len651 | W6-2 | 651 | QJE49092.1 hypothetical protein LMBV_029 [Largemouth bass virus] | 100 | 37 | 2.59E-13 | 8 |
| k141_15078_flag0_multi1.8668_len877 | W5-1 | 877 | WEI28966.1 hypothetical protein [Largemouth bass virus] | 97.2 | 36 | 2.64E-13 | 10 |
| k141_145801_flag1_multi7.0000_len435 | W5-3 | 435 | QJE49092.1 hypothetical protein LMBV_029 [Largemouth bass virus] | 100 | 36 | 2.71E-13 | 7 |
| k141_102879_flag1_multi52.8629_len1279 | W4-1 | 1279 | AYV88120.1 putative p31K protein [Mandarin fish ranavirus] | 97.4 | 39 | 2.72E-13 | 188 |
| k141_128321_flag1_multi1.0000_len311 | W5-1 | 311 | WAK75072.1 hypothetical protein [Mandarin fish ranavirus] | 90 | 40 | 2.72E-13 | 3 |
| k141_138617_flag0_multi76.4762_len477 | W5-1 | 477 | XPZ21261.1 hypothetical protein MRVORF002 [Mandarin fish ranavirus] | 100 | 40 | 2.72E-13 | 95 |
| k141_121774_flag0_multi29.9036_len1687 | W5-1 | 1687 | AYV88211.1 hypothetical protein [Mandarin fish ranavirus] | 86.3 | 51 | 3.04E-13 | 83 |
| k141_16848_flag1_multi3.4282_len858 | W5-1 | 858 | UVF58790.1 MAG: myristylated membrane protein, partial [Halichoeres melanurus ranavirus] | 97.9 | 47 | 3.25E-13 | 15 |
| k141_73771_flag1_multi8.0000_len644 | W5-1 | 644 | QJE49100.1 hypothetical protein LMBV_037 [Largemouth bass virus] | 90.5 | 42 | 3.30E-13 | 14 |
| k141_71206_flag1_multi21.0000_len553 | W2-3 | 553 | QJE49090.1 hypothetical protein LMBV_027 [Largemouth bass virus] | 92.5 | 40 | 3.69E-13 | 23 |
| k141_17430_flag0_multi3.6190_len582 | W5-1 | 582 | UUY86197.1 hypothetical protein [Largemouth bass virus] | 83.9 | 56 | 3.73E-13 | 11 |
| k141_38925_flag1_multi14.0000_len415 | W5-1 | 415 | UUY86199.1 hypothetical protein [Largemouth bass virus] | 100 | 31 | 3.78E-13 | 8 |
| k141_54706_flag1_multi2.0000_len311 | W5-1 | 311 | UUY86235.1 hypothetical protein [Largemouth bass virus] | 97.1 | 35 | 3.91E-13 | 2 |
| k141_129561_flag1_multi2.9529_len1117 | W5-1 | 1117 | XRL22790.1 hypothetical protein [Siniperca chuatsi ranavirus] | 97.4 | 39 | 4.01E-13 | 114 |
| k141_91272_flag1_multi2.0000_len480 | W5-1 | 480 | XRB52842.1 transcription factor TFIIB cyclin-like domain-containing protein, partial [Largemouth bass virus] | 97.3 | 37 | 4.02E-13 | 3 |
| k141_42204_flag1_multi11.0000_len1899 | W5-1 | 1899 | AIG51690.1 major capsid protein, partial [Koi ranavirus] | 100 | 39 | 4.15E-13 | 77 |
| k141_40670_flag1_multi5.7067_len516 | W2-1 | 516 | QYU76034.1 putative myristylated membrane protein, partial [Koi ranavirus] | 88.1 | 42 | 4.42E-13 | 0 |
| k141_15043_flag1_multi2.0000_len394 | W5-1 | 394 | WHU98721.1 MCP, partial [Hybrid snakehead ranavirus] | 100 | 37 | 4.53E-13 | 0 |
| k141_30522_flag0_multi1.0000_len260 | W2-3 | 260 | AYV88122.1 putative 3-beta-hydroxy-delta-5-C27 steroid oxidoreductase-like protein [Mandarin fish ranavirus] | 100 | 32 | 4.55E-13 | 1 |
| k141_64292_flag1_multi2.5641_len788 | W6-1 | 788 | WAK75072.1 hypothetical protein [Mandarin fish ranavirus] | 100 | 40 | 4.72E-13 | 6 |
| k141_60220_flag1_multi1.0000_len334 | W1-2 | 334 | AYV88167.1 hypothetical protein [Mandarin fish ranavirus] | 100 | 31 | 4.73E-13 | 0 |
| k141_68243_flag1_multi2.0000_len510 | W5-1 | 510 | QJE49145.1 putative proliferating cell nuclear antigen [Largemouth bass virus] | 100 | 36 | 4.81E-13 | 2 |
| k141_40584_flag1_multi5.9201_len454 | W5-1 | 454 | QJE49076.1 putative myristylated membrane protein [Largemouth bass virus] | 89.4 | 47 | 4.93E-13 | 2 |
| k141_50746_flag1_multi1.0000_len416 | W5-1 | 416 | QJE49097.1 hypothetical protein LMBV_034 [Largemouth bass virus] | 64.4 | 59 | 5.00E-13 | 0 |
| k141_87159_flag0_multi1.0000_len581 | W4-1 | 581 | QJE49145.1 putative proliferating cell nuclear antigen [Largemouth bass virus] | 78.4 | 51 | 5.13E-13 | 124.3 |

| | | | | | | | |
|---|---|---|---|---|---|---|---|
| k141_67084_flag1_multi6.0000_len344 | W2-3 | 344 | WEI29006.1 putative 2-cysteine adaptor domain protein [Largemouth bass virus] | 87.5 | 40 | 5.28E-13 | 4 |
| k141_62978_flag1_multi5.3865_len630 | W4-1 | 630 | UUY86261.1 hypothetical protein [Largemouth bass virus] | 85.7 | 49 | 5.35E-13 | 14 |
| k141_167168_flag0_multi1.0000_len392 | W2-3 | 392 | QJE49097.1 hypothetical protein LMBV_034 [Largemouth bass virus] | 94.3 | 35 | 5.37E-13 | 0 |
| k141_163803_flag1_multi7.0000_len1038 | W5-1 | 1038 | UUY86251.1 hypothetical protein [Largemouth bass virus] | 97.2 | 36 | 5.50E-13 | 23 |
| k141_49160_flag1_multi4.1834_len719 | W1-1 | 719 | QJE49149.1 putative immediate early protein ICP-46 [Largemouth bass virus] | 100 | 35 | 5.53E-13 | 14 |
| k141_85625_flag1_multi4.0000_len324 | W1-1 | 324 | WAK75112.1 putative LITAF PIG7 possible membrane associated motif in LPS-induced tumor necrosis factor alpha factor [Mandarin fish ranavirus] | 100 | 30 | 5.62E-13 | 5 |
| k141_122634_flag1_multi10.0000_len2009 | W2-3 | 2009 | AYV88191.1 hypothetical protein [Mandarin fish ranavirus] | 100 | 35 | 5.66E-13 | 55 |
| k141_68503_flag1_multi1.0000_len348 | W4-1 | 348 | AYV88120.1 putative p31K protein [Mandarin fish ranavirus] | 91.3 | 46 | 5.85E-13 | 40.17 |
| k141_66186_flag1_multi3.0000_len700 | W4-1 | 700 | AYV88133.1 putative DNA-directed RNA polymerase II subunit [Mandarin fish ranavirus] | 92.5 | 40 | 6.10E-13 | 5 |
| k141_89166_flag1_multi2.6031_len398 | W4-1 | 398 | AYV88149.2 hypothetical protein [Mandarin fish ranavirus] | 89.5 | 38 | 6.29E-13 | 1 |
| k141_62711_flag0_multi15.9257_len5995 | W5-1 | 5995 | AYV88198.1 hypothetical protein [Mandarin fish ranavirus] | 100 | 41 | 6.81E-13 | 377 |
| k141_49567_flag1_multi18.8005_len6255 | W5-1 | 6255 | QJE49105.1 hypothetical protein LMBV_042 [Largemouth bass virus] | 97.8 | 45 | 6.88E-13 | 407 |
| k141_72194_flag1_multi1.9522_len810 | W5-1 | 810 | AIG51690.1 major capsid protein, partial [Koi ranavirus] | 100 | 37 | 6.98E-13 | 15 |
| k141_6850_flag1_multi1.0000_len373 | W5-1 | 373 | QJE49077.1 hypothetical protein LMBV_014 [Largemouth bass virus] | 100 | 32 | 7.00E-13 | 1 |
| k141_51870_flag1_multi6.0000_len584 | W5-1 | 584 | UUY86192.1 putative myristylated membrane protein [Largemouth bass virus] | 97.2 | 36 | 7.40E-13 | 11 |
| k141_43726_flag1_multi3.0000_len682 | W5-1 | 682 | UUY86269.1 hypothetical protein [Largemouth bass virus] | 93.2 | 44 | 7.43E-13 | 12 |
| k141_10505_flag1_multi6.0000_len1057 | W5-1 | 1057 | QIZ30887.1 major capsid protein, partial [Largemouth bass virus] | 94.6 | 37 | 7.46E-13 | 23 |
| k141_139723_flag1_multi9.0000_len1032 | W5-1 | 1032 | UUY86235.1 hypothetical protein [Largemouth bass virus] | 76 | 50 | 7.72E-13 | 38.19 |
| k141_83263_flag1_multi6.8997_len500 | W4-1 | 500 | AIG51690.1 major capsid protein, partial [Koi ranavirus] | 97.6 | 41 | 7.78E-13 | 5 |
| k141_77376_flag0_multi31.6562_len522 | W5-1 | 522 | WAK75072.1 hypothetical protein [Mandarin fish ranavirus] | 96.9 | 32 | 8.01E-13 | 3.12 |
| k141_76562_flag1_multi4.0000_len691 | W1-3 | 691 | UVF58785.1 MAG: major capsid protein [Halichoeres melanurus ranavirus] | 97.4 | 38 | 8.25E-13 | 6 |
| k141_35245_flag1_multi3.0000_len496 | W5-1 | 496 | UUY86199.1 hypothetical protein [Largemouth bass virus] | 96.9 | 32 | 8.48E-13 | 2 |
| k141_105726_flag3_multi293.7876_len513 | W4-1 | 513 | QJE49116.1 hypothetical protein LMBV_053 [Largemouth bass virus] | 100 | 37 | 8.56E-13 | 209 |
| k141_53393_flag1_multi9.0000_len1249 | W5-1 | 1249 | AYV88120.1 putative p31K protein [Mandarin fish ranavirus] | 94.9 | 39 | 8.66E-13 | 19 |
| k141_150796_flag1_multi1.9578_len544 | W5-1 | 544 | AYV88167.1 hypothetical protein [Mandarin fish ranavirus] | 100 | 31 | 8.98E-13 | 2 |
| k141_30215_flag0_multi2.4795_len579 | W5-1 | 579 | XRB52842.1 transcription factor TFIIB cyclin-like domain-containing protein, partial [Largemouth bass virus] | 100 | 41 | 1.01E-12 | 3 |
| k141_36372_flag1_multi1.8686_len491 | W5-1 | 491 | QJE49104.1 hypothetical protein LMBV_041 [Largemouth bass virus] | 92.3 | 39 | 1.02E-12 | 1 |
| k141_11222_flag1_multi7.0543_len785 | W4-1 | 785 | AYV88118.1 hypothetical protein [Mandarin fish ranavirus] | 100 | 33 | 1.04E-12 | 21 |
| k141_143749_flag0_multi3.8136_len495 | W5-1 | 495 | UUY86238.1 hypothetical protein [Largemouth bass virus] | 91.7 | 36 | 1.05E-12 | 4 |
| k141_84355_flag1_multi6.0000_len605 | W5-1 | 605 | AIG51690.1 major capsid protein, partial [Koi ranavirus] | 100 | 37 | 1.06E-12 | 13 |
| k141_120724_flag1_multi1.7979_len616 | W5-1 | 616 | AAC79876.1 viral core protein, partial [Labroides dimidatus ranavirus] | 100 | 32 | 1.09E-12 | 3 |
| k141_92277_flag0_multi32.0000_len757 | W1-1 | 757 | UUY86250.1 hypothetical protein [Largemouth bass virus] | 97.5 | 40 | 1.12E-12 | 63 |
| k141_52274_flag1_multi8.0000_len787 | W5-1 | 787 | UUY86211.1 hypothetical protein [Largemouth bass virus] | 97.4 | 39 | 1.19E-12 | 22 |
| k141_47593_flag1_multi6.0000_len848 | W2-1 | 848 | UUY86235.1 hypothetical protein [Largemouth bass virus] | 100 | 39 | 1.25E-12 | 10 |
| k141_76948_flag1_multi5.0000_len875 | W6-1 | 875 | QYU76034.1 putative myristylated membrane protein, partial [Koi ranavirus] | 100 | 33 | 1.34E-12 | 12 |
| k141_155255_flag1_multi6.8432_len983 | W5-1 | 983 | QJE49146.1 putative thiol oxidoreductase [Largemouth bass virus] | 73.5 | 49 | 1.39E-12 | 25 |
| k141_68069_flag1_multi5.6149_len1055 | W4-1 | 1055 | UVF58785.1 MAG: major capsid protein [Halichoeres melanurus ranavirus] | 100 | 40 | 1.43E-12 | 11 |

| | | | | | | | |
|---|---|---|---|---|---|---|---|
| k141_25167_flag1_multi2.8663_len485 | W2-3 | 485 | UUY86229.1 putative DNA dependent RNA polymerase a subunit [Largemouth bass virus] | 62.5 | 64 | 1.45E-12 | 4 |
| k141_119606_flag0_multi43.4816_len494 | W1-1 | 494 | WEI29006.1 putative 2-cysteine adaptor domain protein [Largemouth bass virus] | 64.8 | 54 | 1.50E-12 | 30 |
| k141_7246_flag1_multi8.5152_len504 | W5-1 | 504 | QJE49105.1 hypothetical protein LMBV_042 [Largemouth bass virus] | 100 | 40 | 1.51E-12 | 11 |
| k141_38910_flag1_multi3.0000_len405 | W5-1 | 405 | AYV88120.1 putative p31K protein [Mandarin fish ranavirus] | 91.7 | 36 | 1.53E-12 | 5 |
| k141_88323_flag1_multi7.0000_len1699 | W2-3 | 1699 | XRB52785.1 hypothetical protein LMBV_28 [Largemouth bass virus] | 91.7 | 36 | 1.64E-12 | 41 |
| k141_79260_flag1_multi12.9155_len851 | W4-1 | 851 | UUY86227.1 hypothetical protein [Largemouth bass virus] | 97.2 | 36 | 1.64E-12 | 25 |
| k141_94911_flag1_multi2.0000_len536 | W5-1 | 536 | WEI29006.1 putative 2-cysteine adaptor domain protein [Largemouth bass virus] | 82.9 | 41 | 1.67E-12 | 2 |
| k141_131124_flag0_multi2.3139_len552 | W5-1 | 552 | UVF58785.1 MAG: major capsid protein [Halichoeres melanurus ranavirus] | 92.3 | 39 | 1.68E-12 | 1 |
| k141_41888_flag0_multi25.2162_len548 | W2-2 | 548 | QJE49095.1 hypothetical protein LMBV_032 [Largemouth bass virus] | 100 | 34 | 1.71E-12 | 6 |
| k141_764_flag1_multi1.6734_len634 | W5-1 | 634 | AIG51690.1 major capsid protein, partial [Koi ranavirus] | 100 | 39 | 1.80E-12 | 12 |
| k141_18059_flag1_multi2.9034_len1487 | W5-1 | 1487 | UUY86235.1 hypothetical protein [Largemouth bass virus] | 93 | 43 | 1.96E-12 | 11 |
| k141_39887_flag1_multi17.9836_len810 | W5-1 | 810 | ACT85833.1 major capsid protein, partial [Frog virus 3] | 82.9 | 41 | 1.96E-12 | 6 |
| k141_61475_flag1_multi1.0000_len545 | W4-1 | 545 | AYV88191.1 hypothetical protein [Mandarin fish ranavirus] | 56.3 | 71 | 1.97E-12 | 34 |
| k141_161011_flag1_multi1.0000_len845 | W5-1 | 845 | UUY86258.1 hypothetical protein [Largemouth bass virus] | 97.5 | 40 | 1.98E-12 | 36.42 |
| k141_108819_flag1_multi4.9739_len447 | W1-1 | 447 | XRB52768.1 P31K protein, partial [Largemouth bass virus] | 97.1 | 34 | 1.99E-12 | 9 |
| k141_20653_flag1_multi1.0000_len355 | W5-1 | 355 | UVF58790.1 MAG: myristylated membrane protein, partial [Halichoeres melanurus ranavirus] | 100 | 41 | 1.99E-12 | 0 |
| k141_135433_flag0_multi380.8529_len515 | W5-1 | 515 | AYV88122.1 putative 3-beta-hydroxy-delta-5-C27 steroid oxidoreductase-like protein [Mandarin fish ranavirus] | 97.1 | 35 | 2.08E-12 | 90 |
| k141_39445_flag1_multi1.7154_len394 | W2-1 | 394 | QYU76034.1 putative myristylated membrane protein, partial [Koi ranavirus] | 58.5 | 65 | 2.10E-12 | 1 |
| k141_103996_flag0_multi81.1204_len789 | W4-1 | 789 | UVF58789.1 MAG: NIF/NLI interacting factor [Halichoeres melanurus ranavirus] | 91.9 | 37 | 2.16E-12 | 49 |
| k141_35161_flag1_multi1.0000_len334 | W2-1 | 334 | QJE49128.1 hypothetical protein LMBV_065 [Largemouth bass virus] | 96.9 | 32 | 2.20E-12 | 0 |
| k141_72857_flag1_multi6.9318_len2340 | W5-1 | 2340 | QJE49105.1 hypothetical protein LMBV_042 [Largemouth bass virus] | 81.1 | 53 | 2.30E-12 | 57 |
| k141_85489_flag1_multi3.0000_len851 | W4-1 | 851 | UVF58785.1 MAG: major capsid protein [Halichoeres melanurus ranavirus] | 100 | 39 | 2.36E-12 | 9 |
| k141_14530_flag1_multi3.0000_len708 | W5-1 | 708 | WEI28972.1 CTD-phosphotransferase [Largemouth bass virus] | 91.7 | 36 | 2.42E-12 | 12 |
| k141_162108_flag1_multi2.5621_len785 | W5-1 | 785 | XPZ21261.1 hypothetical protein MRVORF002 [Mandarin fish ranavirus] | 100 | 35 | 2.42E-12 | 8 |
| k141_99627_flag1_multi10.0000_len483 | W1-1 | 483 | AYV88173.2 putative ribonucleotide reductase beta subunit [Mandarin fish ranavirus] | 100 | 34 | 2.45E-12 | 14 |
| k141_124114_flag1_multi2.0000_len1406 | W5-1 | 1406 | UUY86192.1 putative myristylated membrane protein [Largemouth bass virus] | 90.5 | 42 | 2.46E-12 | 11 |
| k141_45509_flag1_multi10.0000_len2348 | W5-1 | 2348 | AIG51690.1 major capsid protein, partial [Koi ranavirus] | 100 | 39 | 2.49E-12 | 60 |
| k141_75964_flag1_multi6.0000_len1407 | W5-1 | 1407 | WHA35528.1 putative replicating factor [Micropterus salmoides ranavirus] | 100 | 38 | 2.52E-12 | 30 |
| k141_92236_flag1_multi6.9858_len565 | W4-1 | 565 | UVF58785.1 MAG: major capsid protein [Halichoeres melanurus ranavirus] | 100 | 39 | 2.54E-12 | 14 |
| k141_36377_flag1_multi1.8534_len523 | W5-1 | 523 | UUY86241.1 hypothetical protein [Largemouth bass virus] | 96.8 | 31 | 2.66E-12 | 3 |
| k141_137080_flag0_multi3.6413_len325 | W1-1 | 325 | XRL22821.1 hypothetical protein [Siniperca chuatsi ranavirus] | 100 | 33 | 2.67E-12 | 0 |
| k141_100188_flag0_multi6.8935_len1380 | W5-1 | 1380 | QJE49066.1 hypothetical protein LMBV_003 [Largemouth bass virus] | 94.7 | 38 | 2.77E-12 | 34 |
| k141_27649_flag1_multi4.0000_len620 | W5-1 | 620 | UUY86235.1 hypothetical protein [Largemouth bass virus] | 97.1 | 34 | 2.88E-12 | 9 |
| k141_104359_flag0_multi7.4439_len943 | W4-1 | 943 | AYV88120.1 putative p31K protein [Mandarin fish ranavirus] | 93.2 | 44 | 2.95E-12 | 24 |
| k141_73487_flag0_multi17.4096_len639 | W5-1 | 639 | QJE49066.1 hypothetical protein LMBV_003 [Largemouth bass virus] | 97.4 | 38 | 3.01E-12 | 34 |
| k141_86208_flag1_multi1.9245_len1664 | W1-1 | 1664 | UUY86269.1 hypothetical protein [Largemouth bass virus] | 84.6 | 52 | 3.04E-12 | 40 |
| k141_70720_flag1_multi14.9261_len2739 | W5-1 | 2739 | AYV88120.1 putative p31K protein [Mandarin fish ranavirus] | 72.3 | 65 | 3.04E-12 | 127 |

| | | | | | | | |
|---|---|---|---|---|---|---|---|
| k141_24906_flag1_multi9.0000_len632 | W2-1 | 632 | AYV88179.2 putative DNA dependent RNA polymerase II second largest subunit [Mandarin fish ranavirus] | 94.3 | 35 | 3.09E-12 | 7 |
| k141_75658_flag1_multi5.0000_len501 | W2-1 | 501 | UUY86229.1 putative DNA dependent RNA polymerase a subunit [Largemouth bass virus] | 97.2 | 36 | 3.13E-12 | 6 |
| k141_77885_flag1_multi1.7580_len608 | W1-1 | 608 | QJE49066.1 hypothetical protein LMBV_003 [Largemouth bass virus] | 97.1 | 35 | 3.15E-12 | 10 |
| k141_48327_flag1_multi1.4895_len521 | W1-1 | 521 | AYV88191.1 hypothetical protein [Mandarin fish ranavirus] | 100 | 32 | 3.17E-12 | 6 |
| k141_14028_flag1_multi1.7742_len606 | W1-1 | 606 | AYV88191.1 hypothetical protein [Mandarin fish ranavirus] | 80.9 | 47 | 3.19E-12 | 3 |
| k141_32228_flag1_multi6.0000_len408 | W4-1 | 408 | XPZ21261.1 hypothetical protein MRVORF002 [Mandarin fish ranavirus] | 94.7 | 38 | 3.20E-12 | 6 |
| k141_94683_flag1_multi2.0000_len335 | W2-2 | 335 | QJE49128.1 hypothetical protein LMBV_065 [Largemouth bass virus] | 96.9 | 32 | 3.22E-12 | 2 |
| k141_97829_flag1_multi27.7137_len864 | W4-1 | 864 | AIG51690.1 major capsid protein, partial [Koi ranavirus] | 100 | 39 | 3.32E-12 | 113 |
| k141_163602_flag1_multi1.0000_len421 | W5-1 | 421 | QJE49077.1 hypothetical protein LMBV_014 [Largemouth bass virus] | 93.9 | 33 | 3.32E-12 | 7.05 |
| k141_38480_flag1_multi1.9419_len468 | W5-1 | 468 | WHU98721.1 MCP, partial [Hybrid snakehead ranavirus] | 100 | 36 | 3.37E-12 | 3 |
| k141_128874_flag1_multi11.0853_len3376 | W5-1 | 3376 | WAK75107.1 hypothetical protein [Mandarin fish ranavirus] | 91.4 | 35 | 3.50E-12 | 127 |
| k141_133930_flag1_multi7.0000_len401 | W1-1 | 401 | WEI29006.1 putative 2-cysteine adaptor domain protein [Largemouth bass virus] | 100 | 33 | 3.55E-12 | 9 |
| k141_84344_flag1_multi5.1917_len981 | W1-1 | 981 | AYV88125.2 hypothetical protein [Mandarin fish ranavirus] | 100 | 46 | 3.63E-12 | 19 |
| k141_77511_flag1_multi5.0000_len1281 | W2-1 | 1281 | UUY86223.1 hypothetical protein [Largemouth bass virus] | 100 | 39 | 3.64E-12 | 21 |
| k141_78668_flag1_multi3.0000_len789 | W4-1 | 789 | QJE49101.1 hypothetical protein LMBV_038 [Largemouth bass virus] | 100 | 39 | 3.74E-12 | 2 |
| k141_166_flag1_multi8.0000_len414 | W5-1 | 414 | WEI28972.1 CTD-phosphotransferase [Largemouth bass virus] | 92.7 | 41 | 3.75E-12 | 12 |
| k141_143127_flag1_multi2.0000_len569 | W5-1 | 569 | UUY86193.1 hypothetical protein [Largemouth bass virus] | 94.4 | 36 | 3.81E-12 | 5 |
| k141_37689_flag1_multi2.6447_len535 | W2-3 | 535 | UUY86208.1 hypothetical protein [Largemouth bass virus] | 76 | 50 | 3.83E-12 | 5 |
| k141_1947_flag1_multi4.8312_len686 | W5-1 | 686 | UUY86258.1 hypothetical protein [Largemouth bass virus] | 94.9 | 39 | 4.09E-12 | 15 |
| k141_66643_flag1_multi16.9819_len4549 | W4-1 | 4549 | AYV88120.1 putative p31K protein [Mandarin fish ranavirus] | 97.4 | 38 | 4.11E-12 | 237 |
| k141_10451_flag1_multi2.3469_len657 | W6-1 | 657 | AIG51690.1 major capsid protein, partial [Koi ranavirus] | 50 | 96 | 4.14E-12 | 3 |
| k141_3900_flag1_multi1.9485_len588 | W2-1 | 588 | AYV88120.1 putative p31K protein [Mandarin fish ranavirus] | 92.1 | 38 | 4.36E-12 | 1 |
| k141_40629_flag0_multi5.8358_len811 | W2-3 | 811 | WEI29006.1 putative 2-cysteine adaptor domain protein [Largemouth bass virus] | 100 | 32 | 4.44E-12 | 6 |
| k141_39644_flag1_multi1.0000_len398 | W5-1 | 398 | UUY86267.1 ribonucleotide reductase alpha subunit [Largemouth bass virus] | 100 | 35 | 4.48E-12 | 2 |
| k141_37303_flag1_multi2.5531_len772 | W5-1 | 772 | QJE49101.1 hypothetical protein LMBV_038 [Largemouth bass virus] | 97.1 | 35 | 4.74E-12 | 4 |
| k141_39526_flag1_multi11.0000_len1002 | W1-1 | 1002 | AYV88194.1 hypothetical protein [Mandarin fish ranavirus] | 92.9 | 42 | 4.86E-12 | 32 |
| k141_27006_flag1_multi4.5598_len350 | W6-1 | 350 | UVF58793.1 MAG: DNA polymerase [Halichoeres melanurus ranavirus] | 100 | 33 | 5.07E-12 | 3 |
| k141_127194_flag1_multi2.8985_len594 | W5-1 | 594 | WAK75112.1 putative LITAF PIG7 possible membrane associated motif in LPS-induced tumor necrosis factor alpha factor [Mandarin fish ranavirus] | 96.7 | 30 | 5.11E-12 | 6 |
| k141_12555_flag1_multi4.0000_len979 | W5-1 | 979 | AYV88134.2 putative tyrosine kinase [Mandarin fish ranavirus] | 100 | 38 | 5.18E-12 | 10 |
| k141_13041_flag1_multi3.0000_len390 | W5-1 | 390 | UVF58785.1 MAG: major capsid protein [Halichoeres melanurus ranavirus] | 100 | 38 | 5.25E-12 | 0 |
| k141_94970_flag1_multi10.5455_len1263 | W5-1 | 1263 | QJE49137.1 hypothetical protein LMBV_074 [Largemouth bass virus] | 97.4 | 38 | 5.47E-12 | 40 |
| k141_144060_flag1_multi9.9412_len3200 | W5-1 | 3200 | UUY86229.1 putative DNA dependent RNA polymerase a subunit [Largemouth bass virus] | 97.5 | 40 | 5.54E-12 | 107 |
| k141_52114_flag1_multi8.5040_len512 | W1-1 | 512 | AYV88125.2 hypothetical protein [Mandarin fish ranavirus] | 100 | 46 | 5.57E-12 | 1 |
| k141_29945_flag1_multi2.0000_len338 | W6-2 | 338 | AYV88172.2 hypothetical protein [Mandarin fish ranavirus] | 75 | 44 | 6.00E-12 | 2 |
| k141_145012_flag1_multi13.0000_len1087 | W5-1 | 1087 | QYU76034.1 putative myristylated membrane protein, partial [Koi ranavirus] | 100 | 36 | 6.33E-12 | 62 |
| k141_104202_flag0_multi17.9454_len8855 | W5-1 | 8855 | UUY86235.1 hypothetical protein [Largemouth bass virus] | 100 | 41 | 6.38E-12 | 569 |

| | | | | | | | |
|---|---|---|---|---|---|---|---|
| k141_58375_flag1_multi1.0000_len400 | W2-1 | 400 | AIG51690.1 major capsid protein, partial [Koi ranavirus] | 82.6 | 46 | 6.57E-12 | 0 |
| k141_73151_flag1_multi8.6850_len395 | W5-1 | 395 | WHU98721.1 MCP, partial [Hybrid snakehead ranavirus] | 100 | 35 | 7.54E-12 | 10 |
| k141_101456_flag1_multi1.0000_len566 | W4-1 | 566 | AYV88134.2 putative tyrosine kinase [Mandarin fish ranavirus] | 100 | 33 | 7.76E-12 | 12 |
| k141_99131_flag0_multi26.8247_len586 | W5-1 | 586 | UUY86261.1 hypothetical protein [Largemouth bass virus] | 100 | 40 | 8.04E-12 | 37 |
| k141_63767_flag0_multi34.8992_len7562 | W1-1 | 7562 | UUY86267.1 ribonucleotide reductase alpha subunit [Largemouth bass virus] | 90.5 | 42 | 8.16E-12 | 963 |
| k141_101736_flag1_multi3.0000_len492 | W4-1 | 492 | XRB52768.1 P31K protein, partial [Largemouth bass virus] | 72.2 | 54 | 8.20E-12 | 5 |
| k141_143975_flag0_multi6.0000_len340 | W5-1 | 340 | UUY86235.1 hypothetical protein [Largemouth bass virus] | 100 | 32 | 8.23E-12 | 2 |
| k141_43248_flag1_multi25.0000_len1452 | W2-3 | 1452 | WEI28972.1 CTD-phosphotransferase [Largemouth bass virus] | 76.6 | 47 | 8.38E-12 | 98 |
| k141_47800_flag1_multi6.3241_len1508 | W5-1 | 1508 | UUY86193.1 hypothetical protein [Largemouth bass virus] | 95 | 40 | 9.06E-12 | 29 |
| k141_40667_flag1_multi5.0000_len511 | W1-2 | 511 | AYV88172.2 hypothetical protein [Mandarin fish ranavirus] | 97.7 | 44 | 9.08E-12 | 6 |
| k141_6559_flag1_multi4.0000_len1087 | W5-1 | 1087 | WHU98721.1 MCP, partial [Hybrid snakehead ranavirus] | 100 | 37 | 9.08E-12 | 26 |
| k141_101194_flag1_multi20.4903_len1789 | W4-1 | 1789 | AIG51690.1 major capsid protein, partial [Koi ranavirus] | 89.1 | 46 | 9.24E-12 | 143 |
| k141_95037_flag1_multi2.5216_len605 | W5-1 | 605 | QYU76036.1 helicase-like protein, partial [Koi ranavirus] | 100 | 39 | 9.30E-12 | 7 |
| k141_77171_flag1_multi2.5385_len687 | W6-1 | 687 | AIG51690.1 major capsid protein, partial [Koi ranavirus] | 92.5 | 40 | 9.40E-12 | 5 |
| k141_124087_flag1_multi4.0000_len831 | W5-1 | 831 | UVF58785.1 MAG: major capsid protein [Halichoeres melanurus ranavirus] | 100 | 39 | 9.43E-12 | 8 |
| k141_78598_flag1_multi3.0000_len406 | W4-1 | 406 | QJE49141.1 putative NTPase [Largemouth bass virus] | 75 | 48 | 9.46E-12 | 2 |
| k141_83139_flag1_multi10.0000_len967 | W5-1 | 967 | XPZ21295.1 putative RNAseIII [Mandarin fish ranavirus] | 94.9 | 39 | 9.58E-12 | 14 |
| k141_24940_flag1_multi3.0000_len650 | W4-1 | 650 | AYV88176.1 putative tumor necrosis factor receptor [Mandarin fish ranavirus] | 100 | 31 | 9.73E-12 | 9 |
| k141_64207_flag1_multi3.0000_len736 | W5-1 | 736 | WHA35533.1 putative DNA dependent RNA polymerase A subunit [Micropterus salmoides ranavirus] | 86.7 | 45 | 9.87E-12 | 8 |
| k141_53066_flag1_multi2.0000_len494 | W5-1 | 494 | QYU76031.1 neurofilament triplet H1-like protein, partial [Koi ranavirus] | 100 | 33 | 9.99E-12 | 2 |
| k141_108665_flag1_multi1.5772_len562 | W1-1 | 562 | QJE49104.1 hypothetical protein LMBV_041 [Largemouth bass virus] | 97.4 | 38 | 1.03E-11 | 1 |
| k141_11552_flag1_multi12.0000_len334 | W5-1 | 334 | UUY86235.1 hypothetical protein [Largemouth bass virus] | 81.4 | 43 | 1.08E-11 | 0 |
| k141_148763_flag1_multi4.4625_len900 | W5-1 | 900 | UUY86197.1 hypothetical protein [Largemouth bass virus] | 100 | 46 | 1.10E-11 | 11 |
| k141_92485_flag1_multi20.1911_len680 | W5-1 | 680 | WHA35533.1 putative DNA dependent RNA polymerase A subunit [Micropterus salmoides ranavirus] | 92.1 | 38 | 1.17E-11 | 62 |
| k141_18056_flag1_multi8.7480_len522 | W5-1 | 522 | UUY86267.1 ribonucleotide reductase alpha subunit [Largemouth bass virus] | 100 | 36 | 1.19E-11 | 9 |
| k141_10455_flag0_multi1.0000_len347 | W5-1 | 347 | UUY86229.1 putative DNA dependent RNA polymerase a subunit [Largemouth bass virus] | 97.1 | 35 | 1.27E-11 | 0 |
| k141_76614_flag1_multi2.2732_len668 | W5-1 | 668 | WAK75072.1 hypothetical protein [Mandarin fish ranavirus] | 92.3 | 39 | 1.32E-11 | 6 |
| k141_148538_flag0_multi12.5550_len950 | W5-1 | 950 | UUY86252.1 hypothetical protein [Largemouth bass virus] | 98.1 | 52 | 1.43E-11 | 379.56 |
| k141_24984_flag1_multi8.6213_len2972 | W5-1 | 2972 | AYV88120.1 putative p31K protein [Mandarin fish ranavirus] | 100 | 37 | 1.51E-11 | 81 |
| k141_111035_flag1_multi2.5054_len606 | W5-1 | 606 | AYV88179.2 putative DNA dependent RNA polymerase II second largest subunit [Mandarin fish ranavirus] | 86 | 43 | 1.52E-11 | 7 |
| k141_87367_flag1_multi2.0000_len446 | W5-1 | 446 | QYU76036.1 helicase-like protein, partial [Koi ranavirus] | 93.9 | 33 | 1.53E-11 | 0 |
| k141_55671_flag1_multi35.0000_len1678 | W4-1 | 1678 | WEI28972.1 CTD-phosphotransferase [Largemouth bass virus] | 100 | 39 | 1.58E-11 | 181 |
| k141_114731_flag1_multi5.9419_len1174 | W5-1 | 1174 | QJE49149.1 putative immediate early protein ICP-46 [Largemouth bass virus] | 100 | 33 | 1.58E-11 | 21 |
| k141_80555_flag1_multi11.9086_len1946 | W5-1 | 1946 | UUY86228.1 hypothetical protein [Largemouth bass virus] | 92.7 | 41 | 1.71E-11 | 94 |
| k141_81263_flag1_multi13.7152_len1191 | W5-1 | 1191 | AYV88183.1 hypothetical protein [Mandarin fish ranavirus] | 100 | 34 | 1.71E-11 | 55 |
| k141_107592_flag1_multi5.0000_len676 | W5-1 | 676 | QJE49119.1 hypothetical protein LMBV_056 [Largemouth bass virus] | 100 | 32 | 1.85E-11 | 13 |

| | | | | | | | |
|---|---|---|---|---|---|---|---|
| k141_164124_flag1_multi1.0000_len423 | W5-1 | 423 | UUY86263.1 hypothetical protein [Largemouth bass virus] | 97.1 | 34 | 1.85E-11 | 3 |
| k141_121533_flag1_multi9.0000_len513 | W1-1 | 513 | WAK75107.1 hypothetical protein [Mandarin fish ranavirus] | 100 | 28 | 1.90E-11 | 14 |
| k141_126568_flag1_multi1.9630_len546 | W1-1 | 546 | UUY86195.1 hypothetical protein [Largemouth bass virus] | 97.1 | 34 | 1.97E-11 | 8 |
| k141_32581_flag1_multi1.0000_len481 | W4-1 | 481 | UUY86227.1 hypothetical protein [Largemouth bass virus] | 97 | 33 | 1.97E-11 | 1 |
| k141_158737_flag1_multi1.8139_len501 | W5-1 | 501 | QJE49066.1 hypothetical protein LMBV_003 [Largemouth bass virus] | 100 | 33 | 2.02E-11 | 0 |
| k141_78596_flag1_multi3.4959_len385 | W4-1 | 385 | AIG51690.1 major capsid protein, partial [Koi ranavirus] | 100 | 35 | 2.05E-11 | 6 |
| k141_55390_flag1_multi3.0000_len754 | W1-1 | 754 | UUY86258.1 hypothetical protein [Largemouth bass virus] | 88.4 | 43 | 2.24E-11 | 8 |
| k141_24470_flag1_multi4.0000_len860 | W5-1 | 860 | WHA35533.1 putative DNA dependent RNA polymerase A subunit [Micropterus salmoides ranavirus] | 100 | 34 | 2.25E-11 | 10 |
| k141_29839_flag1_multi9.0000_len817 | W5-1 | 817 | AYV88191.1 hypothetical protein [Mandarin fish ranavirus] | 100 | 32 | 2.32E-11 | 14 |
| k141_49239_flag1_multi4.0000_len831 | W6-1 | 831 | AYV88203.1 putative insulin-like growth factor [Mandarin fish ranavirus] | 97.4 | 38 | 2.52E-11 | 10 |
| k141_88907_flag1_multi1.6529_len383 | W1-1 | 383 | WEI29006.1 putative 2-cysteine adaptor domain protein [Largemouth bass virus] | 96.9 | 32 | 2.54E-11 | 0 |
| k141_168239_flag1_multi4.0000_len323 | W5-1 | 323 | WAK75112.1 putative LITAF PIG7 possible membrane associated motif in LPS-induced tumor necrosis factor alpha factor [Mandarin fish ranavirus] | 93.3 | 30 | 2.59E-11 | 2 |
| k141_57194_flag1_multi19.0000_len1506 | W5-1 | 1506 | QJE49066.1 hypothetical protein LMBV_003 [Largemouth bass virus] | 92.1 | 38 | 2.61E-11 | 93 |
| k141_101469_flag1_multi3.4428_len543 | W5-1 | 543 | QJE49145.1 putative proliferating cell nuclear antigen [Largemouth bass virus] | 100 | 35 | 2.64E-11 | 3 |
| k141_116568_flag1_multi6.9662_len496 | W5-1 | 496 | ABA41591.1 DNA-dependent DNA polymerase, partial [Largemouth bass virus] | 70.2 | 47 | 2.65E-11 | 13 |
| k141_3665_flag1_multi5.0852_len458 | W5-1 | 458 | QYU76034.1 putative myristylated membrane protein, partial [Koi ranavirus] | 96.8 | 31 | 2.65E-11 | 5 |
| k141_92921_flag1_multi1.0000_len500 | W2-1 | 500 | ALR73092.1 major capsid protein, partial [Box turtle ranavirus] | 85.3 | 34 | 2.66E-11 | 16 |
| k141_103475_flag1_multi7.6598_len1314 | W4-1 | 1314 | WHA35533.1 putative DNA dependent RNA polymerase A subunit [Micropterus salmoides ranavirus] | 100 | 40 | 2.71E-11 | 25 |
| k141_135449_flag1_multi11.8443_len1695 | W1-1 | 1695 | UUY86238.1 hypothetical protein [Largemouth bass virus] | 85 | 40 | 2.75E-11 | 68 |
| k141_78530_flag1_multi9.2727_len493 | W5-1 | 493 | ACT85834.1 major capsid protein, partial [Frog virus 3] | 79.5 | 39 | 2.78E-11 | 21 |
| k141_26913_flag1_multi2.0000_len372 | W2-1 | 372 | UUY86211.1 hypothetical protein [Largemouth bass virus] | 100 | 32 | 2.79E-11 | 1 |
| k141_25579_flag1_multi4.5509_len524 | W2-1 | 524 | WXI69541.1 hypothetical protein [Largemouth bass virus] | 91.4 | 35 | 2.81E-11 | 6 |
| k141_67825_flag1_multi4.0000_len530 | W4-1 | 530 | QYU76034.1 putative myristylated membrane protein, partial [Koi ranavirus] | 100 | 33 | 2.81E-11 | 9 |
| k141_18703_flag1_multi1.0000_len459 | W5-1 | 459 | ABB92283.1 hypothetical protein [Tiger frog virus] | 80 | 45 | 2.84E-11 | 1 |
| k141_76944_flag1_multi4.0000_len572 | W5-1 | 572 | QJE49097.1 hypothetical protein LMBV_034 [Largemouth bass virus] | 100 | 33 | 2.89E-11 | 5 |
| k141_116432_flag1_multi5.0000_len664 | W5-1 | 664 | QJE49149.1 putative immediate early protein ICP-46 [Largemouth bass virus] | 97.4 | 38 | 2.90E-11 | 10 |
| k141_138995_flag1_multi5.0000_len1102 | W5-1 | 1102 | WAK75106.1 hypothetical protein [Mandarin fish ranavirus] | 100 | 31 | 3.04E-11 | 12 |
| k141_8381_flag1_multi7.0000_len940 | W5-1 | 940 | WEI29006.1 putative 2-cysteine adaptor domain protein [Largemouth bass virus] | 53.2 | 94 | 3.15E-11 | 19 |
| k141_39359_flag1_multi1.0000_len596 | W4-1 | 596 | WEI29006.1 putative 2-cysteine adaptor domain protein [Largemouth bass virus] | 100 | 32 | 3.35E-11 | 16.07 |
| k141_154672_flag1_multi10.5958_len710 | W5-1 | 710 | AYV88125.2 hypothetical protein [Mandarin fish ranavirus] | 100 | 46 | 3.44E-11 | 26 |
| k141_60892_flag1_multi1.0000_len523 | W4-1 | 523 | UUY86266.1 hypothetical protein [Largemouth bass virus] | 91.7 | 36 | 3.46E-11 | 0 |
| k141_79049_flag1_multi2.0000_len800 | W2-3 | 800 | WHA35533.1 putative DNA dependent RNA polymerase A subunit [Micropterus salmoides ranavirus] | 100 | 35 | 3.77E-11 | 13 |
| k141_91165_flag1_multi2.8933_len572 | W5-1 | 572 | QJE49137.1 hypothetical protein LMBV_074 [Largemouth bass virus] | 100 | 35 | 3.89E-11 | 2 |
| k141_47747_flag1_multi8.1945_len794 | W4-1 | 794 | AYV88211.1 hypothetical protein [Mandarin fish ranavirus] | 82.2 | 45 | 3.98E-11 | 14 |
| k141_107348_flag1_multi4.0000_len825 | W1-1 | 825 | AIG51690.1 major capsid protein, partial [Koi ranavirus] | 81.4 | 43 | 4.22E-11 | 14 |
| k141_63664_flag1_multi1.0000_len471 | W5-1 | 471 | UUY86261.1 hypothetical protein [Largemouth bass virus] | 94.7 | 38 | 4.37E-11 | 4 |

| | | | | | | | |
|---|---|---|---|---|---|---|---|
| k141_54510_flag1_multi3.9623_len936 | W1-1 | 936 | XRB52771.1 hypothetical protein LMBV_14 [Largemouth bass virus] | 100 | 31 | 4.81E-11 | 11 |
| k141_24992_flag1_multi18.8254_len2512 | W5-1 | 2512 | AIG51690.1 major capsid protein, partial [Koi ranavirus] | 100 | 39 | 4.99E-11 | 128 |
| k141_86417_flag1_multi4.9093_len659 | W5-1 | 659 | UUY86258.1 hypothetical protein [Largemouth bass virus] | 97.3 | 37 | 5.11E-11 | 12 |
| k141_82722_flag1_multi7.0000_len608 | W5-1 | 608 | QYU76034.1 putative myristylated membrane protein, partial [Koi ranavirus] | 96.8 | 31 | 5.46E-11 | 13 |
| k141_36831_flag1_multi8.6829_len532 | W5-1 | 532 | UUY86211.1 hypothetical protein [Largemouth bass virus] | 85.4 | 41 | 5.63E-11 | 15 |
| k141_43402_flag1_multi3.2061_len568 | W6-1 | 568 | UUY86267.1 ribonucleotide reductase alpha subunit [Largemouth bass virus] | 94.4 | 36 | 6.11E-11 | 8 |
| k141_106100_flag0_multi1.6141_len566 | W5-1 | 566 | WEI29006.1 putative 2-cysteine adaptor domain protein [Largemouth bass virus] | 97.1 | 35 | 6.44E-11 | 16 |
| k141_104464_flag1_multi12.9739_len600 | W4-1 | 600 | AFD96401.1 major capsid protein, partial [Largemouth bass virus] | 100 | 31 | 7.09E-11 | 24 |
| k141_70993_flag0_multi14.9085_len447 | W5-1 | 447 | QJE49101.1 hypothetical protein LMBV_038 [Largemouth bass virus] | 97 | 33 | 7.21E-11 | 7 |
| k141_19260_flag1_multi17.7864_len3366 | W4-2 | 3366 | AYV88191.1 hypothetical protein [Mandarin fish ranavirus] | 94.1 | 34 | 7.63E-11 | 222 |
| k141_81508_flag1_multi8.0000_len1212 | W5-1 | 1212 | WHA35533.1 putative DNA dependent RNA polymerase A subunit [Micropterus salmoides ranavirus] | 100 | 39 | 8.32E-11 | 64 |
| k141_117325_flag1_multi4.0000_len436 | W4-2 | 436 | AYV88134.2 putative tyrosine kinase [Mandarin fish ranavirus] | 100 | 30 | 8.70E-11 | 4 |
| k141_27226_flag1_multi3.0000_len469 | W6-1 | 469 | AIG51690.1 major capsid protein, partial [Koi ranavirus] | 94.1 | 34 | 8.82E-11 | 5 |
| k141_63299_flag1_multi3.5613_len1119 | W5-1 | 1119 | UVF58788.1 MAG: helicase, partial [Halichoeres melanurus ranavirus] | 87.8 | 41 | 8.92E-11 | 21 |
| k141_15325_flag1_multi7.1082_len1278 | W4-1 | 1278 | UUY86192.1 putative myristylated membrane protein [Largemouth bass virus] | 85 | 40 | 9.09E-11 | 21 |
| k141_127255_flag1_multi6.0000_len339 | W5-1 | 339 | UVF58793.1 MAG: DNA polymerase [Halichoeres melanurus ranavirus] | 100 | 33 | 9.83E-11 | 3 |
| k141_137961_flag1_multi3.3183_len760 | W1-1 | 760 | UUY86261.1 hypothetical protein [Largemouth bass virus] | 94.7 | 38 | 9.90E-11 | 8 |
| k141_97415_flag0_multi43.9813_len781 | W1-1 | 781 | AYV88133.1 putative DNA-directed RNA polymerase II subunit [Mandarin fish ranavirus] | 100 | 30 | 1.00E-10 | 301 |
| k141_96034_flag1_multi11.0000_len984 | W5-1 | 984 | QIZ30887.1 major capsid protein, partial [Largemouth bass virus] | 94.1 | 34 | 1.02E-10 | 33 |
| k141_11708_flag1_multi6.7607_len1796 | W6-1 | 1796 | QJE49123.1 putative DNA repair protein RAD2 [Largemouth bass virus] | 95 | 40 | 1.09E-10 | 41 |
| k141_69803_flag0_multi77.5781_len506 | W4-1 | 506 | AYV88133.1 putative DNA-directed RNA polymerase II subunit [Mandarin fish ranavirus] | 100 | 30 | 1.14E-10 | 121 |
| k141_93861_flag1_multi2.0000_len552 | W5-1 | 552 | AYV88191.1 hypothetical protein [Mandarin fish ranavirus] | 100 | 30 | 1.20E-10 | 8 |
| k141_29612_flag1_multi3.0000_len608 | W1-1 | 608 | WEI29006.1 putative 2-cysteine adaptor domain protein [Largemouth bass virus] | 62.9 | 62 | 1.26E-10 | 3 |
| k141_123994_flag1_multi35.0000_len858 | W1-1 | 858 | UUY86199.1 hypothetical protein [Largemouth bass virus] | 100 | 28 | 1.27E-10 | 76 |
| k141_57435_flag1_multi3.2834_len568 | W4-1 | 568 | AYV88120.1 putative p31K protein [Mandarin fish ranavirus] | 93.9 | 33 | 1.35E-10 | 10 |
| k141_127492_flag0_multi7.9562_len3975 | W5-1 | 3975 | UUY86235.1 hypothetical protein [Largemouth bass virus] | 94.7 | 38 | 1.40E-10 | 103 |
| k141_48195_flag1_multi1.0000_len557 | W6-1 | 557 | UUY86241.1 hypothetical protein [Largemouth bass virus] | 100 | 28 | 1.40E-10 | 1 |
| k141_76918_flag1_multi14.1547_len1634 | W1-1 | 1634 | AIG51690.1 major capsid protein, partial [Koi ranavirus] | 100 | 38 | 1.43E-10 | 58 |
| k141_12396_flag0_multi1.0000_len485 | W5-1 | 485 | AYV88179.2 putative DNA dependent RNA polymerase II second largest subunit [Mandarin fish ranavirus] | 100 | 32 | 1.45E-10 | 4 |
| k141_76556_flag1_multi4.6014_len427 | W4-1 | 427 | AIG51690.1 major capsid protein, partial [Koi ranavirus] | 72 | 50 | 1.49E-10 | 1 |
| k141_43246_flag1_multi3.2083_len674 | W5-1 | 674 | QJE49101.1 hypothetical protein LMBV_038 [Largemouth bass virus] | 88.6 | 35 | 1.50E-10 | 4 |
| k141_154196_flag1_multi4.0000_len652 | W5-1 | 652 | QJE49107.1 hypothetical protein LMBV_044 [Largemouth bass virus] | 56.3 | 64 | 1.51E-10 | 5 |
| k141_33460_flag1_multi14.0000_len1742 | W1-1 | 1742 | WEI29006.1 putative 2-cysteine adaptor domain protein [Largemouth bass virus] | 97.2 | 36 | 1.52E-10 | 97 |
| k141_110062_flag1_multi3.9739_len486 | W5-1 | 486 | QYU76036.1 helicase-like protein, partial [Koi ranavirus] | 100 | 35 | 1.56E-10 | 7 |
| k141_53168_flag1_multi7.0000_len532 | W4-1 | 532 | AIG51690.1 major capsid protein, partial [Koi ranavirus] | 100 | 36 | 1.57E-10 | 16 |
| k141_38750_flag1_multi2.5582_len906 | W5-1 | 906 | QJE49072.1 hypothetical protein LMBV_009 [Largemouth bass virus] | 93.9 | 33 | 1.62E-10 | 13.2 |

| | | | | | | | |
|---|---|---|---|---|---|---|---|
| k141_20843_flag1_multi4.7920_len516 | W1-1 | 516 | UUY86261.1 hypothetical protein [Largemouth bass virus] | 100 | 37 | 1.69E-10 | 9 |
| k141_18345_flag1_multi3.0000_len411 | W2-1 | 411 | UUY86235.1 hypothetical protein [Largemouth bass virus] | 91.9 | 37 | 1.79E-10 | 3 |
| k141_101323_flag0_multi60.7082_len398 | W5-1 | 398 | QJE49101.1 hypothetical protein LMBV_038 [Largemouth bass virus] | 97 | 33 | 1.80E-10 | 3 |
| k141_166788_flag1_multi1.5876_len512 | W5-1 | 512 | QJE49101.1 hypothetical protein LMBV_038 [Largemouth bass virus] | 97.1 | 35 | 1.81E-10 | 1 |
| k141_1796_flag1_multi4.0000_len332 | W5-1 | 332 | AIG51690.1 major capsid protein, partial [Koi ranavirus] | 69.6 | 56 | 1.93E-10 | 3 |
| k141_44673_flag1_multi6.0000_len1273 | W5-1 | 1273 | AIG51690.1 major capsid protein, partial [Koi ranavirus] | 100 | 35 | 2.00E-10 | 20 |
| k141_59542_flag0_multi14.8758_len2621 | W5-1 | 2621 | AYV88191.1 hypothetical protein [Mandarin fish ranavirus] | 97 | 33 | 2.00E-10 | 84 |
| k141_106505_flag1_multi1.0000_len492 | W2-3 | 492 | AIG51690.1 major capsid protein, partial [Koi ranavirus] | 75 | 48 | 2.07E-10 | 0 |
| k141_158443_flag1_multi11.0000_len2521 | W5-1 | 2521 | QIZ30887.1 major capsid protein, partial [Largemouth bass virus] | 88.2 | 34 | 2.18E-10 | 101 |
| k141_56601_flag1_multi0.9780_len414 | W5-1 | 414 | UUY86267.1 ribonucleotide reductase alpha subunit [Largemouth bass virus] | 96.8 | 31 | 2.24E-10 | 0 |
| k141_17275_flag1_multi7.3052_len2084 | W5-1 | 2084 | AYV88211.1 hypothetical protein [Mandarin fish ranavirus] | 90.5 | 42 | 2.24E-10 | 51 |
| k141_3693_flag1_multi1.0000_len771 | W6-1 | 771 | ACT85833.1 major capsid protein, partial [Frog virus 3] | 90.9 | 33 | 2.32E-10 | 22 |
| k141_77359_flag1_multi5.9970_len811 | W4-1 | 811 | XRB52801.1 hypothetical protein LMBV_44 [Largemouth bass virus] | 78.9 | 38 | 2.44E-10 | 20 |
| k141_160075_flag1_multi4.7631_len2032 | W5-1 | 2032 | QJE49137.1 hypothetical protein LMBV_074 [Largemouth bass virus] | 81.8 | 44 | 2.50E-10 | 31 |
| k141_137468_flag1_multi1.0000_len605 | W2-3 | 605 | UUY86235.1 hypothetical protein [Largemouth bass virus] | 88.9 | 36 | 2.59E-10 | 9 |
| k141_138326_flag1_multi4.9004_len1065 | W5-1 | 1065 | UVF58789.1 MAG: NIF/NLI interacting factor [Halichoeres melanurus ranavirus] | 100 | 30 | 2.84E-10 | 21 |
| k141_101460_flag1_multi4.0000_len498 | W2-2 | 498 | WAK75102.1 hypothetical protein [Mandarin fish ranavirus] | 97 | 33 | 2.88E-10 | 4 |
| k141_89416_flag1_multi2.7789_len521 | W1-1 | 521 | WAK75097.1 hypothetical protein [Mandarin fish ranavirus] | 94.4 | 36 | 2.89E-10 | 2 |
| k141_118087_flag1_multi4.0746_len677 | W5-1 | 677 | QJE49072.1 hypothetical protein LMBV_009 [Largemouth bass virus] | 93.8 | 32 | 2.90E-10 | 8 |
| k141_87378_flag1_multi1.9954_len575 | W5-1 | 575 | UUY86223.1 hypothetical protein [Largemouth bass virus] | 100 | 35 | 2.93E-10 | 4 |
| k141_30388_flag1_multi5.0000_len465 | W6-1 | 465 | QIZ30887.1 major capsid protein, partial [Largemouth bass virus] | 78 | 41 | 2.94E-10 | 8 |
| k141_49005_flag1_multi20.0000_len1067 | W5-1 | 1067 | UVF58785.1 MAG: major capsid protein [Halichoeres melanurus ranavirus] | 100 | 39 | 2.97E-10 | 66 |
| k141_50718_flag1_multi4.0000_len321 | W5-1 | 321 | AYV88167.1 hypothetical protein [Mandarin fish ranavirus] | 100 | 26 | 3.00E-10 | 4 |
| k141_38266_flag1_multi17.0000_len1699 | W5-1 | 1699 | WHA35528.1 putative replicating factor [Micropterus salmoides ranavirus] | 100 | 32 | 3.17E-10 | 89 |
| k141_8292_flag1_multi2.9864_len583 | W2-1 | 583 | AYV88169.1 hypothetical protein [Mandarin fish ranavirus] | 97 | 33 | 3.34E-10 | 4 |
| k141_136803_flag1_multi5.0000_len832 | W1-1 | 832 | XRL22821.1 hypothetical protein [Siniperca chuatsi ranavirus] | 88.9 | 36 | 3.47E-10 | 12 |
| k141_115202_flag1_multi19.0000_len3819 | W1-1 | 3819 | AYV88143.1 hypothetical protein [Mandarin fish ranavirus] | 100 | 35 | 3.52E-10 | 178 |
| k141_74620_flag1_multi4.2239_len476 | W5-1 | 476 | UVF58788.1 MAG: helicase, partial [Halichoeres melanurus ranavirus] | 91.9 | 37 | 3.62E-10 | 8 |
| k141_159823_flag1_multi3.0000_len957 | W5-1 | 957 | QYU76034.1 putative myristylated membrane protein, partial [Koi ranavirus] | 95.1 | 41 | 4.34E-10 | 15 |
| k141_43662_flag1_multi5.0000_len308 | W4-1 | 308 | AXB27397.1 major capsid protein, partial [Ranavirus sp.] | 88.6 | 35 | 4.47E-10 | 5 |
| k141_869_flag1_multi1.7486_len491 | W5-1 | 491 | WXI69541.1 hypothetical protein [Largemouth bass virus] | 97 | 33 | 4.68E-10 | 3 |
| k141_16898_flag1_multi6.8099_len1372 | W5-1 | 1372 | AYV88125.2 hypothetical protein [Mandarin fish ranavirus] | 77 | 61 | 4.84E-10 | 30 |
| k141_80605_flag0_multi1.0000_len223 | W4-1 | 223 | WXI69513.1 hypothetical protein [Largemouth bass virus] | 70.8 | 72 | 5.07E-10 | 204 |
| k141_61315_flag1_multi4.9642_len1258 | W1-1 | 1258 | AYV88191.1 hypothetical protein [Mandarin fish ranavirus] | 100 | 33 | 5.24E-10 | 16 |
| k141_110331_flag1_multi3.1725_len628 | W1-1 | 628 | AYV88179.2 putative DNA dependent RNA polymerase II second largest subunit [Mandarin fish ranavirus] | 97.1 | 35 | 5.32E-10 | 6 |
| k141_16031_flag1_multi4.0000_len629 | W4-1 | 629 | AYV88120.1 putative p31K protein [Mandarin fish ranavirus] | 74.5 | 47 | 5.58E-10 | 7 |
| k141_38047_flag1_multi4.0000_len651 | W4-1 | 651 | AYV88211.1 hypothetical protein [Mandarin fish ranavirus] | 100 | 36 | 5.63E-10 | 1 |
| k141_122957_flag0_multi3.0110_len414 | W5-1 | 414 | WEI29006.1 putative 2-cysteine adaptor domain protein [Largemouth bass virus] | 100 | 33 | 5.76E-10 | 5 |
| k141_168723_flag1_multi1.9413_len533 | W5-1 | 533 | QJE49137.1 hypothetical protein LMBV_074 [Largemouth bass virus] | 100 | 32 | 5.81E-10 | 4 |

| | | | | | | | |
|---|---|---|---|---|---|---|---|
| k141_119478_flag0_multi10.7101_len3094 | W5-1 | 3094 | WAK75072.1 hypothetical protein [Mandarin fish ranavirus] | 100 | 36 | 5.83E-10 | 98.69 |
| k141_26549_flag1_multi4.4757_len1110 | W2-1 | 1110 | WEI28966.1 hypothetical protein [Largemouth bass virus] | 88.1 | 42 | 6.37E-10 | 23 |
| k141_128060_flag1_multi4.9222_len861 | W1-1 | 861 | XRL22790.1 hypothetical protein [Siniperca chuatsi ranavirus] | 97.3 | 37 | 7.24E-10 | 14 |
| k141_14651_flag1_multi6.0000_len571 | W5-1 | 571 | UUY86199.1 hypothetical protein [Largemouth bass virus] | 100 | 27 | 7.25E-10 | 8 |
| k141_114117_flag1_multi6.0000_len447 | W5-1 | 447 | AYV88191.1 hypothetical protein [Mandarin fish ranavirus] | 91.2 | 34 | 7.34E-10 | 10 |
| k141_98925_flag1_multi3.0000_len707 | W5-1 | 707 | UUY86258.1 hypothetical protein [Largemouth bass virus] | 82.5 | 40 | 8.15E-10 | 5 |
| k141_41694_flag1_multi4.0000_len1010 | W5-1 | 1010 | AYV88191.1 hypothetical protein [Mandarin fish ranavirus] | 100 | 32 | 8.51E-10 | 24 |
| k141_62496_flag1_multi38.4805_len2528 | W1-1 | 2528 | WHA35533.1 putative DNA dependent RNA polymerase A subunit [Micropterus salmoides ranavirus] | 100 | 34 | 8.63E-10 | 332 |
| k141_59986_flag1_multi5.0457_len754 | W5-1 | 754 | UUY86257.1 hypothetical protein [Largemouth bass virus] | 100 | 30 | 9.46E-10 | 12 |
| k141_91600_flag1_multi4.0000_len309 | W4-1 | 309 | QJE49144.1 hypothetical protein LMBV_081 [Largemouth bass virus] | 67.3 | 52 | 1.71E-09 | 2 |
| k141_50675_flag1_multi13.7958_len905 | W1-1 | 905 | UUY86261.1 hypothetical protein [Largemouth bass virus] | 100 | 36 | 1.75E-09 | 36 |
| k141_150797_flag1_multi3.0000_len938 | W5-1 | 938 | QJE49104.1 hypothetical protein LMBV_041 [Largemouth bass virus] | 97.1 | 34 | 2.07E-09 | 7 |
| k141_15034_flag1_multi8.0000_len1086 | W5-1 | 1086 | UUY86261.1 hypothetical protein [Largemouth bass virus] | 100 | 39 | 2.62E-09 | 25 |

**Table S2. Top 1000 expressed host genes, their abundance, and functions.**

| W1.1_spleen | W1.2_liver | W1.3_brain | W2.1_spleen | W2.2_liver | W2.3_brain | W3.1_spleen | W3.2_liver | W3.3_brain | W4.1_spleen | W4.2_liver | W4.3_brain | W5.1_spleen | W5.2_liver | W5.3_brain | W6.1_spleen | W6.2_liver | W6.3_brain | Total counts | Protein names | Gene names | Gene Ontology (Biological Process) |
|---|---|---|---|---|---|---|---|---|---|---|---|---|---|---|---|---|---|---|---|---|---|
| 396696.4 | 409326.69 | 0 | 394744.67 | 237728.71 | 1691.56 | 2033.97 | 290103.14 | 0 | 690273.55 | 753442.86 | 0 | 208038.05 | 246538.5 | 0 | 355145.83 | 605680.27 | 16780.83 | 4608225.03 | Nucleoredoxin (EC 1.8.1.8) | nxn zgc:110449 | cell differentiation [GO:0030154]; circulatory system development [GO:0072359]; negative regulation of protein ubiquitination [GO:0031397]; negative regulation of Wnt signaling pathway [GO:0030178]; Wnt signaling pathway [GO:0016055] |
| 188775.06 | 254248.56 | 120008.44 | 444064.98 | 207158.42 | 752312.47 | 1634.61 | 160837.75 | 59854.22 | 397073.85 | 249766.89 | 1191591.13 | 249225.44 | 86684.59 | 75011.34 | 22533.79 | 74381.69 | 67234.25 | 4602397.48 | Receptor-type tyrosine-protein phosphatase eta (Protein-tyrosine phosphatase eta) (R-PTP-eta) (EC 3.1.3.48) (HPTP eta) (Protein-tyrosine phosphatase receptor type J) (R-PTP-J) (Supporting-cell antigen) | PTPRJ | negative regulation of epidermal growth factor receptor signaling pathway [GO:0042059]; negative regulation of MAP kinase activity [GO:0043407]; negative regulation of phosphatidylinositol 3-kinase/protein kinase B signal transduction [GO:0051898]; negative regulation of T cell receptor signaling pathway [GO:0050860]; platelet formation [GO:0030220]; platelet-derived growth factor receptor signaling pathway [GO:0048008]; positive regulation of cell adhesion [GO:0045785]; positive regulation of focal adhesion assembly [GO:0051894] |

| | | | | | | | | | | | | | | | | | | | | | |
|---|---|---|---|---|---|---|---|---|---|---|---|---|---|---|---|---|---|---|---|---|---|
| 3129.37 | 2.35 | 71.93 | 83626.61 | 2.93 | 571.54 | 1038674.27 | 1.8 | 131.8 | 6129.62 | 14181.99 | 51.92 | 300.37 | 7.48 | 197.6 | 602658.55 | 15099.23 | 261.65 | 1765101.01 | Eukaryotic translation initiation factor 2-alpha kinase 3 (EC 2.7.11.1) (PRKR-like endoplasmic reticulum kinase) (Pancreatic eIF2-alpha kinase) (Protein tyrosine kinase EIF2AK3) (EC 2.7.10.2) | Eif2ak3 Pek Perk | angiogenesis [GO:0001525]; bone mineralization [GO:0030282]; calcium-mediated signaling [GO:0019722]; cellular response to amino acid starvation [GO:0034198]; cellular response to cold [GO:0070417]; cellular response to glucose starvation [GO:0042149]; chondrocyte development [GO:0002063]; endocrine pancreas development [GO:0031018]; endoplasmic reticulum organization [GO:0007029]; endoplasmic reticulum unfolded protein response [GO:0030968]; ER overload response [GO:0006983]; ERAD pathway [GO:0036503]; fat cell differentiation [GO:0045444]; insulin-like growth factor receptor signaling pathway [GO:0048009]; intrinsic apoptotic signaling pathway in response to endoplasmic reticulum stress [GO:0070059]; lactation [GO:0007595]; negative regulation of apoptotic process [GO:0043066]; negative regulation of gene expression [GO:0010629]; negative regulation of myelination [GO:0031642]; |
| 376417.3 | 4169.88 | 2186.3 | 102710.03 | 28628.32 | 52729.12 | 1580.48 | 2527.13 | 16.5 | 234002.85 | 20185.64 | 10985.18 | 769111.54 | 2977.48 | 3184.9 | 18004.34 | 5263.68 | 220.69 | 1634901.36 | Histone H1 [Cleaved into: Oncorhyncin II] | LOC141806056 | defense response to bacterium [GO:0042742]; nucleosome assembly [GO:0006334] |
| 79168.92 | 61476.22 | 68384.67 | 26432.59 | 66918.9 | 158670.16 | 10623.24 | 29755.86 | 115301.45 | 45097.35 | 23062.29 | 54267.14 | 126804.59 | 144889.86 | 169443.96 | 49958.84 | 74016.03 | 184482.15 | 1488754.22 | Tumor necrosis factor receptor superfamily member 1B (Tumor necrosis factor receptor 2) (TNF-R2) (Tumor necrosis factor receptor type II) (TNF-RII) (TNFR-II) (p75) (p80 TNF-alpha receptor) (CD antigen CD120b) (Etanercept) [Cleaved into: Tumor necrosis factor receptor superfamily member 1b, membrane form;Tumor necrosis factor-binding protein 2 (TBP-2) (TBPII)] | TNFRSF1B TNFBR TNFR2 | aortic valve development [GO:0003176]; cellular response to lipopolysaccharide [GO:0071222]; extrinsic apoptotic signaling pathway [GO:0097191]; glial cell-neuron signaling [GO:0150098]; inflammatory response [GO:0006954]; intrinsic apoptotic signaling pathway in response to DNA damage [GO:0008630]; negative regulation of cardiac muscle hypertrophy [GO:0010614]; negative regulation of extracellular matrix constituent secretion [GO:0003332]; negative regulation of neuroinflammatory response [GO:0150079]; positive regulation of apoptotic process involved in morphogenesis [GO:1902339]; positive regulation of membrane protein ectodomain proteolysis [GO:0051044]; positive regulation of myelination [GO:0031643]; positive regulation of oligodendrocyte differentiation [GO:0048714]; pulmonary valve development [GO:0003177]; regulation of cytokine production involved in |

| | | | | | | | | | | | | | | | | | | | | | |
|---|---|---|---|---|---|---|---|---|---|---|---|---|---|---|---|---|---|---|---|---|---|
| 1891.59 | 2.29 | 16.03 | 61180.55 | 0 | 622.23 | 733032.4 | 0 | 0 | 3243.7 | 7843.86 | 5.97 | 192.43 | 0 | 0 | 507281.15 | 11733.32 | 197.01 | 1327242.53 | Nuclear receptor subfamily 1 group D member 2 (Orphan nuclear hormone receptor BD73) (Rev-erb alpha-related receptor) (RVR) (Rev-erb-beta) (V-erbA-related protein 1-related) (EAR-1R) | NR1D2 | cell differentiation [GO:0030154]; circadian behavior [GO:0048512]; energy homeostasis [GO:0097009]; hormone-mediated signaling pathway [GO:0009755]; intracellular receptor signaling pathway [GO:0030522]; lipid homeostasis [GO:0055088]; negative regulation of DNA-templated transcription [GO:0045892]; negative regulation of inflammatory response [GO:0050728]; negative regulation of transcription by RNA polymerase II [GO:0000122]; positive regulation of DNA-templated transcription [GO:0045893]; positive regulation of transcription by RNA polymerase II [GO:0045944]; regulation of circadian rhythm [GO:0042752]; regulation of DNA-templated transcription [GO:0006355]; regulation of inflammatory response [GO:0050727]; regulation of lipid metabolic process [GO:0019216]; regulation of skeletal muscle cell differentiation [GO:2001014] |
| 171973.47 | 2446.59 | 1307.72 | 89095.66 | 17463.73 | 22240.65 | 1560.67 | 1652.63 | 45.32 | 486158.8 | 11032.74 | 9872.92 | 384390.13 | 1498.7 | 1446.39 | 14378.14 | 3039.5 | 126.97 | 1219730.73 | General transcription factor II-I repeat domain-containing protein 2 (GTF2I repeat domain-containing protein 2) (Transcription factor GTF2IRD2) | GTF2IRD2 | transition between fast and slow fiber [GO:0014883] |
| 52087.46 | 57349.11 | 156292.68 | 30038.46 | 73490.84 | 104232.21 | 643.13 | 72278.55 | 87798.38 | 56910.39 | 63829.75 | 180356.45 | 18780.13 | 29511.83 | 55413.17 | 3564.24 | 19552.83 | 55092.9 | 1117222.51 | N-acylglucosamine 2-epimerase (AGE) (EC 5.1.3.8) (GlcNAc 2-epimerase) (N-acetyl-D-glucosamine 2-epimerase) (Renin-binding protein) (RnBP) | RENBP | carbohydrate metabolic process [GO:0005975]; N-acetylglucosamine metabolic process [GO:0006044]; N-acetylmannosamine metabolic process [GO:0006051]; N-acetylneuraminate catabolic process [GO:0019262]; regulation of blood pressure [GO:0008217] |

| | | | | | | | | | | | | | | | | | | | | | |
|---|---|---|---|---|---|---|---|---|---|---|---|---|---|---|---|---|---|---|---|---|---|
| 35873.73 | 60619.86 | 27748.84 | 71686.5 | 55455.05 | 168778.11 | 197.37 | 48075.94 | 25398.11 | 42988.84 | 38832.78 | 215865.61 | 70513.59 | 45005.97 | 29814.76 | 9549.62 | 24950.51 | 30370.83 | 1001726.02 | Nuclear pore complex protein Nup160 (160 kDa nucleoporin) (Gene trap locus 1-13 protein) (GTL-13) (Nucleoporin Nup160) | Nup160 Gtl1-13 Kiaa0197 | mRNA export from nucleus [GO:0006406]; nucleocytoplasmic transport [GO:0006913]; protein transport [GO:0015031] |
| 40071.19 | 60971.45 | 11220.57 | 85612.54 | 22118.77 | 98743.48 | 437.8 | 27008.14 | 8344.56 | 98628 | 57965.83 | 101873.15 | 19260.16 | 7358.7 | 6912.99 | 1945.33 | 4682.56 | 7623.48 | 660778.7 | Tuftelin-interacting protein 11 (Septin and tuftelin-interacting protein 1) (STIP-1) | tfip11 stip zgc:86644 | spliceosomal complex disassembly [GO:0000390] |
| 25865.79 | 36614.35 | 17977.1 | 57614.89 | 34291.86 | 104493.92 | 169.48 | 20601.61 | 9689.2 | 44098.41 | 33538.6 | 149458.42 | 36155.26 | 16430.65 | 17722.88 | 4376.83 | 12076.87 | 11989.58 | 633165.7 | Dynein axonemal heavy chain 12 (Bm259) | Dnah12 | cilium movement involved in cell motility [GO:0060294] |
| 137350.44 | 1514.17 | 780.7 | 28905.85 | 6242.65 | 20003.95 | 342.68 | 697.53 | 21.39 | 77241.27 | 4189.84 | 6242.6 | 316725.69 | 845.83 | 1281.13 | 7369.48 | 1436.98 | 107.49 | 611299.67 | U3 small nucleolar RNA-associated protein 18 homolog (WD repeat-containing protein 50) | UTP18 WDR50 CDABP0061 CGI-48 | ribosomal small subunit biogenesis [GO:0042274]; rRNA processing [GO:0006364] |
| 1264.27 | 10.62 | 13.26 | 15401.15 | 21.73 | 343.37 | 332056.54 | 2.21 | 3.28 | 3310.84 | 7894.86 | 42.21 | 453.83 | 23.92 | 31.26 | 233689.16 | 5884.87 | 115.05 | 600562.43 | Transcriptional regulator Myc (c-Myc) | myc | |
| 32606.24 | 31014.44 | 59489.18 | 14453.25 | 42907.98 | 61770.06 | 236.64 | 38146.19 | 59246.94 | 29962.83 | 24713.07 | 84550.28 | 14583.44 | 17933.87 | 37634.88 | 2135.94 | 11997.6 | 35774.9 | 599157.73 | Complement C3 [Cleaved into: Complement C3 beta chain;Complement C3 alpha chain;C3a anaphylatoxin;Complement C3b (Complement C3b-alpha' chain)] | c3 | complement activation, alternative pathway [GO:0006957]; complement activation, classical pathway [GO:0006958]; defense response to bacterium [GO:0042742]; inflammatory response [GO:0006954] |

| | | | | | | | | | | | | | | | | | | | | | |
|---|---|---|---|---|---|---|---|---|---|---|---|---|---|---|---|---|---|---|---|---|---|
| 127332.8 | 1761.45 | 1185.15 | 27242.71 | 3420.46 | 8606.8 | 367.23 | 503.05 | 218.79 | 100439.32 | 3003.21 | 6009.27 | 290847.44 | 794.61 | 1225.05 | 8365.43 | 1435.83 | 329.33 | 583087.93 | Polycomb protein eed | eed zgc:112509 | digestive tract development [GO:0048565]; exocrine pancreas development [GO:0031017]; heterochromatin formation [GO:0031507]; liver development [GO:0001889]; negative regulation of transcription by RNA polymerase II [GO:0000122]; regulation of glial cell differentiation [GO:0045685]; regulation of neuron differentiation [GO:0045664] |
| 14789.4 | 15559.83 | 36916.71 | 10797.3 | 35845 | 60055.25 | 1060.12 | 23289.26 | 62584.49 | 13598.39 | 13661 | 72109.13 | 27372.61 | 40053.7 | 81772.3 | 3586.89 | 17696.27 | 38316.17 | 569063.82 | Arginine-glutamic acid dipeptide repeats protein (Atrophin-2) | Rere Atr2 Kiaa0458 | branching morphogenesis of a nerve [GO:0048755]; cerebellar granule cell precursor proliferation [GO:0021930]; cerebellar Purkinje cell layer maturation [GO:0021691]; cerebellum development [GO:0021549]; chromatin remodeling [GO:0006338]; dendrite morphogenesis [GO:0048813]; radial glia guided migration of Purkinje cell [GO:0021942] |
| 25917.17 | 39463.73 | 34.72 | 106781.18 | 68634.76 | 743.75 | 295.62 | 39118.7 | 6.39 | 72041.14 | 105981.72 | 7.2 | 11655.37 | 15909.43 | 7.1 | 22299.39 | 50553.02 | 523.15 | 559973.54 | Enkurin domain-containing protein 1 | ENKD1 | establishment of mitotic spindle orientation [GO:0000132]; motile cilium assembly [GO:0044458]; non-motile cilium assembly [GO:1905515] |

| | | | | | | | | | | | | | | | | | | | | | |
|---|---|---|---|---|---|---|---|---|---|---|---|---|---|---|---|---|---|---|---|---|---|
| 35587.75 | 76927.31 | 11475.43 | 8823.94 | 32617.48 | 21397.28 | 9345.36 | 28196.4 | 7713.04 | 6376.19 | 16803.06 | 4833.65 | 50546.9 | 107044.11 | 7420.56 | 6398.97 | 48864.24 | 10064.82 | 490436.49 | U5 small nuclear ribonucleoprotein 200 kDa helicase (EC 3.6.4.13) (Activating signal cointegrator 1 complex subunit 3-like 1) (BRR2 homolog) (U5 snRNP-specific 200 kDa protein) (U5-200KD) | SNRNP200 ASCC3L1 BRR2 HELIC2 KIAA0788 | cis assembly of pre-catalytic spliceosome [GO:0000354]; mRNA splicing, via spliceosome [GO:0000398]; osteoblast differentiation [GO:0001649]; spliceosome conformational change to release U4 (or U4atac) and U1 (or U11) [GO:0000388] |
| 26356.48 | 15368.97 | 43895.18 | 9511.3 | 14117.8 | 32112.95 | 60632.54 | 3893.38 | 12187.89 | 7906.3 | 5589.84 | 65581.83 | 51632.4 | 31413.72 | 26396.14 | 33568.32 | 26382.22 | 21664.5 | 488211.76 | mRNA export factor GLE1 (GLE1 RNA export mediator) (GLE1-like protein) (Nucleoporin GLE1) | GLE1 GLE1L | poly(A)+ mRNA export from nucleus [GO:0016973]; protein transport [GO:0015031] |
| 81.66 | 166150.38 | 151.95 | 223.38 | 49289.06 | 152.32 | 17.43 | 63383.53 | 96.75 | 185.35 | 17917.86 | 97.9 | 117.28 | 85341.43 | 146.89 | 53.64 | 99474.71 | 806.5 | 483688.02 | Cell cycle control protein 50A (P4-ATPase flippase complex beta subunit TMEM30A) (Transmembrane protein 30A) | TMEM30A CDC50A | phospholipid translocation [GO:0045332] |
| 1269.06 | 13.31 | 18.81 | 8929.2 | 18.6 | 215.62 | 267315.7 | 5.81 | 1.92 | 2142.51 | 4604.14 | 35.44 | 565.65 | 30.43 | 39.52 | 174972.33 | 4038.35 | 102.06 | 464318.46 | Transmembrane protein 268 | TMEM268 | |

| | | | | | | | | | | | | | | | | | | | | | |
|---|---|---|---|---|---|---|---|---|---|---|---|---|---|---|---|---|---|---|---|---|---|
| 17587.51 | 29300.38 | 9088.47 | 35978.08 | 27206.71 | 79426.52 | 63.66 | 19079.74 | 9112.17 | 16930.08 | 15680.98 | 101869.59 | 31085.96 | 16194.8 | 9557.01 | 3464.7 | 10975.68 | 11609.55 | 444211.59 | Phospholipid phosphatase 3 (EC 3.1.3.-) (EC 3.1.3.4) (Lipid phosphate phosphohydrolase 3) (PAP2-beta) (Phosphatidate phosphohydrolase type 2b) (Phosphatidic acid phosphatase 2b) (PAP-2b) (PAP2b) (Vascular endothelial growth factor and type I collagen-inducible protein) (VCIP) | PLPP3 LPP3 PPAP2B | Bergmann glial cell differentiation [GO:0060020]; blood vessel development [GO:0001568]; cell-cell adhesion [GO:0098609]; cell-cell adhesion mediated by integrin [GO:0033631]; ceramide metabolic process [GO:0006672]; gastrulation with mouth forming second [GO:0001702]; integrin-mediated signaling pathway [GO:0007229]; phospholipid dephosphorylation [GO:0046839]; phospholipid metabolic process [GO:0006644]; positive regulation of endothelial cell migration [GO:0010595]; positive regulation of endothelial cell-matrix adhesion via fibronectin [GO:1904906]; positive regulation of homotypic cell-cell adhesion [GO:0034112]; positive regulation of intracellular signal transduction [GO:1902533]; positive regulation of transcription by RNA polymerase II [GO:0045944]; protein stabilization [GO:0050821]; regulation of sphingolipid mediated signaling pathway [GO:1902068]; regulation of |
| 37754.23 | 51475.86 | 39.19 | 82775.22 | 41970.5 | 442.66 | 223.63 | 32131.35 | 55.86 | 27345.15 | 38108.82 | 24.39 | 22796.77 | 30192.78 | 52.9 | 23887.61 | 53947.81 | 467.22 | 443691.95 | Zinc-binding protein A33 | | |
| 87831.81 | 1526.18 | 690.32 | 24373.2 | 13681.36 | 24203.41 | 185.11 | 1046.97 | 31.63 | 43637.77 | 8391.75 | 3287.88 | 215223.85 | 1032.87 | 1288.89 | 5322.94 | 2178.11 | 76.36 | 434010.41 | Histone-lysine N-methyltransferase SETD5 (EC 2.1.1.359) (EC 2.1.1.367) (SET domain-containing protein 5) | Setd5 Kiaa1757 | cognition [GO:0050890]; methylation [GO:0032259]; negative regulation of transcription by RNA polymerase III [GO:0016480]; regulation of chromatin organization [GO:1902275]; regulation of DNA-templated transcription elongation [GO:0032784]; regulation of synapse assembly [GO:0051963] |

| | | | | | | | | | | | | | | | | | | | | | |
|---|---|---|---|---|---|---|---|---|---|---|---|---|---|---|---|---|---|---|---|---|---|
| 0 | 0 | 184417.91 | 0 | 0 | 225891.12 | 0 | 0 | 0 | 0 | 0 | 0 | 0 | 0 | 11771.17 | 0 | 0 | 0 | 422080.2 | Ras-related protein Rab-39B (EC 3.6.5.2) | RAB39B | autophagy [GO:0006914]; protein transport [GO:0015031]; Rab protein signal transduction [GO:0032482]; regulation of autophagy [GO:0010506]; synapse organization [GO:0050808]; vesicle-mediated transport [GO:0016192] |
| 15935.91 | 20444.27 | 21949.87 | 17198.21 | 24894.66 | 31660.63 | 33853.08 | 9338.02 | 6016.79 | 9414.8 | 11626.37 | 23189.42 | 28810.85 | 44007.52 | 11855.57 | 34071.65 | 23523.21 | 9607.92 | 377398.75 | Phospholipid scramblase 1 (PL scramblase 1) (Ca(2+)-dependent phospholipid scramblase 1) (Erythrocyte phospholipid scramblase) (Mg(2+)-dependent nuclease) (EC 3.1.-.-) (MmTRA1b) | PLSCR1 | acute-phase response [GO:0006953]; apoptotic process [GO:0006915]; defense response to virus [GO:0051607]; negative regulation of phagocytosis [GO:0050765]; negative regulation of viral genome replication [GO:0045071]; phosphatidylserine biosynthetic process [GO:0006659]; phosphatidylserine exposure on apoptotic cell surface [GO:0070782]; plasma membrane phospholipid scrambling [GO:0017121]; platelet activation [GO:0030168]; positive regulation of chromosome separation [GO:1905820]; positive regulation of DNA topoisomerase (ATP-hydrolyzing) activity [GO:2000373]; positive regulation of gene expression [GO:0010628]; positive regulation of innate immune response [GO:0045089]; positive regulation of transcription by RNA polymerase II [GO:0045944]; regulation of Fc receptor mediated stimulatory signaling pathway [GO:0060368]; regulation of mast cell activation [GO:0033003]; response to |
| 69547.43 | 801.91 | 604.46 | 17637.8 | 3836.56 | 15592.51 | 263.47 | 365.11 | 3.43 | 53547.35 | 2164.57 | 3361.47 | 202445.45 | 411.53 | 1014.88 | 4070.98 | 828.23 | 24.53 | 376521.67 | SREBP regulating gene protein | spring zgc:110063 | intestinal lipid absorption [GO:0098856]; positive regulation of SREBP signaling pathway [GO:2000640] |

| | | | | | | | | | | | | | | | | | | | | | |
|---|---|---|---|---|---|---|---|---|---|---|---|---|---|---|---|---|---|---|---|---|---|
| 16.47 | 39.94 | 72376.02 | 5.73 | 0.97 | 8777 | 47.81 | 0.76 | 316.57 | 6.19 | 3.39 | 286527.42 | 4.63 | 15.72 | 1392.47 | 39.53 | 14.86 | 569.92 | 370155.4 | CD48 antigen (BCM1 surface antigen) (BLAST-1) (HM48-1) (MRC OX-45 surface antigen) (SLAM family member 2) (SLAMF2) (Signaling lymphocytic activation molecule 2) (sgp-60) (CD antigen CD48) | Cd48 Bcm-1 | signal transduction [GO:0007165]; T cell activation [GO:0042110] |
| 11921.65 | 12066.12 | 16668.62 | 8634.84 | 12641.18 | 22381.49 | 97479.05 | 3790.75 | 11703.73 | 6240.73 | 8163.46 | 13598.11 | 25056.77 | 14957.16 | 10724.18 | 52934.21 | 14403.75 | 14156.28 | 357522.08 | V-type immunoglobulin domain-containing suppressor of T-cell activation (Platelet receptor Gi24) (V-set domain-containing immunoregulatory receptor) (V-set immunoregulatory receptor) | Vsir Dies1 PD-1H VISTA | BMP signaling pathway [GO:0030509]; negative regulation of CD4-positive, alpha-beta T cell proliferation [GO:2000562]; negative regulation of T cell cytokine production [GO:0002725]; positive regulation of BMP signaling pathway [GO:0030513]; positive regulation of stem cell differentiation [GO:2000738]; stem cell differentiation [GO:0048863] |

| | | | | | | | | | | | | | | | | | | | | | |
|---|---|---|---|---|---|---|---|---|---|---|---|---|---|---|---|---|---|---|---|---|---|
| 14550.77 | 20528.65 | 10776.81 | 18338.85 | 19827.88 | 61196.6 | 199.62 | 16524.88 | 11156.66 | 15469.87 | 17040.83 | 40623.49 | 20001.89 | 17389.16 | 11852.45 | 3696.59 | 8743.15 | 11036.86 | 318955.01 | Fos-related antigen 2 (FRA-2) | Fosl2 Fra-2 Fra2 | alveolar secondary septum development [GO:0061144]; B cell differentiation [GO:0030183]; B cell proliferation [GO:0042100]; bone development [GO:0060348]; bone mineralization [GO:0030282]; cartilage development [GO:0051216]; cell morphogenesis [GO:0000902]; chondrocyte differentiation [GO:0002062]; chondrocyte proliferation [GO:0035988]; collagen biosynthetic process [GO:0032964]; collagen metabolic process [GO:0032963]; fat cell apoptotic process [GO:1904606]; fat cell differentiation [GO:0045444]; fat pad development [GO:0060613]; gene expression [GO:0010467]; glucose homeostasis [GO:0042593]; growth plate cartilage development [GO:0003417]; homeostasis of number of cells [GO:0048872]; homeostasis of number of cells within a tissue [GO:0048873]; inflammatory response [GO:0006954]; inflammatory response to antigenic stimulus [GO:0002437]; |
| 7652.26 | 12549.91 | 16197.39 | 10724.51 | 20960.88 | 25833.53 | 21153.09 | 8345.99 | 4171.41 | 6199.58 | 9466.68 | 25866.46 | 19464.79 | 43806.11 | 8133.99 | 20963.54 | 20592.4 | 6357.34 | 288439.86 | Ephrin type-A receptor 4 (EC 2.7.10.1) (Tyrosine-protein kinase receptor MPK-3) (Tyrosine-protein kinase receptor SEK-1) | Epha4 Sek Sek1 | adherens junction organization [GO:0034332]; adult walking behavior [GO:0007628]; axon guidance [GO:0007411]; cell adhesion [GO:0007155]; cellular response to amyloid-beta [GO:1904646]; cochlea development [GO:0090102]; corticospinal tract morphogenesis [GO:0021957]; ephrin receptor signaling pathway [GO:0048013]; fasciculation of motor neuron axon [GO:0097156]; fasciculation of sensory neuron axon [GO:0097155]; glial cell migration [GO:0008347]; innervation [GO:0060384]; motor neuron axon guidance [GO:0008045]; negative regulation of axon regeneration [GO:0048681]; negative regulation of cell adhesion [GO:0007162]; negative regulation of cell migration [GO:0030336]; negative regulation of cellular response to hypoxia [GO:1900038]; negative regulation of epithelial to mesenchymal transition [GO:0010719]; negative regulation of ERK1 and ERK2 cascade [GO:0070373]; |

| | | | | | | | | | | | | | | | | | | | | | | |
|---|---|---|---|---|---|---|---|---|---|---|---|---|---|---|---|---|---|---|---|---|---|---|
| 35793.12 | | 15581.68 | 26238.23 | 16953 | 13465.77 | 24183.97 | 209.62 | 37579.85 | 31498.28 | 37216.83 | 7301.47 | 3181.8 | 6913.92 | 10613.06 | 3105.83 | 1201.21 | 11230.85 | 5588.68 | 287857.17 | Dedicator of cytokinesis protein 2 | DOCK2 KIAA0209 | actin cytoskeleton organization [GO:0030036]; alpha-beta T cell proliferation [GO:0046633]; cell migration [GO:0016477]; chemotaxis [GO:0006935]; establishment of T cell polarity [GO:0001768]; immunological synapse formation [GO:0001771]; macropinocytosis [GO:0044351]; membrane raft polarization [GO:0001766]; myeloid dendritic cell activation involved in immune response [GO:0002277]; myoblast fusion [GO:0007520]; negative thymic T cell selection [GO:0045060]; positive regulation of phagocytosis [GO:0050766]; positive thymic T cell selection [GO:0045059]; regulation of small GTPase mediated signal transduction [GO:0051056]; small GTPase-mediated signal transduction [GO:0007264] |
| 10250.2 | | 16905.24 | 14107.92 | 8973.67 | 20668.03 | 12535.47 | 41635.1 | 8404.35 | 3906.19 | 6576.67 | 11215.86 | 12537.05 | 14729.88 | 28300.72 | 6520.07 | 36666.34 | 17174.8 | 4849.71 | 275957.27 | Retinal dehydrogenase 2 (RALDH 2) (RalDH2) (EC 1.2.1.36) (Aldehyde dehydrogenase family 1 member A2) (ALDH1A2) (Retinaldehyde-specific dehydrogenase type 2) (RALDH(II)) | ALDH1A2 RALDH2 | heart looping [GO:0001947]; heart morphogenesis [GO:0003007]; positive regulation of gene expression [GO:0010628]; protein homotetramerization [GO:0051289]; response to retinoic acid [GO:0032526]; retinoic acid biosynthetic process [GO:0002138]; retinoic acid metabolic process [GO:0042573]; retinol metabolic process [GO:0042572]; spinal cord motor neuron differentiation [GO:0021522] |

| | | | | | | | | | | | | | | | | | | | | | |
|---|---|---|---|---|---|---|---|---|---|---|---|---|---|---|---|---|---|---|---|---|---|
| 13723.53 | 13283.7 | 15056.64 | 11760.6 | 19082.97 | 21761.15 | 24165.56 | 5229.64 | 6037.08 | 7133.45 | 8355.93 | 17580.39 | 20399.69 | 29922.72 | 7666.65 | 23507.67 | 20246.93 | 9280.85 | 274195.15 | BCL2/adenovirus E1B 19 kDa protein-interacting protein 3-like (Adenovirus E1B19K-binding protein B5) (BCL2/adenovirus E1B 19 kDa protein-interacting protein 3A) (NIP3-like protein X) (NIP3L) | BNIP3L BNIP3A BNIP3H NIX | cellular response to hypoxia [GO:0071456]; defense response to virus [GO:0051607]; mitochondrial outer membrane permeabilization [GO:0097345]; mitochondrial protein catabolic process [GO:0035694]; negative regulation of apoptotic process [GO:0043066]; negative regulation of mitochondrial membrane potential [GO:0010917]; negative regulation of programmed cell death [GO:0043069]; positive regulation of apoptotic process [GO:0043065]; positive regulation of macroautophagy [GO:0016239]; regulation of mitophagy [GO:1901524]; regulation of programmed cell death [GO:0043067]; regulation of protein targeting to mitochondrion [GO:1903214] |
| 62624.26 | 421.11 | 300.39 | 12575.19 | 1305.33 | 3125.76 | 176.84 | 91.24 | 1.69 | 44861.05 | 1075.02 | 2708.43 | 137038.48 | 225.6 | 438.41 | 3742.69 | 511.78 | 47.08 | 271270.35 | Tripartite motif-containing protein 16 (EC 2.3.2.27) (E3 ubiquitin-protein ligase TRIM16) (Estrogen-responsive B box protein) | TRIM16 EBBP | positive regulation of DNA-templated transcription [GO:0045893]; positive regulation of interleukin-1 beta production [GO:0032731]; positive regulation of keratinocyte differentiation [GO:0045618]; positive regulation of retinoic acid receptor signaling pathway [GO:0048386]; response to growth hormone [GO:0060416]; response to retinoic acid [GO:0032526] |
| 69201.46 | 617.2 | 344.77 | 14923.83 | 4055.06 | 8470.63 | 129.92 | 304.53 | 9.63 | 30302 | 2244.1 | 1904.95 | 125693.15 | 319.05 | 428.81 | 4232.38 | 743.33 | 22.22 | 263947.02 | NADH dehydrogenase [ubiquinone] 1 alpha subcomplex subunit 11 (Complex I-B14.7) (CI-B14.7) (NADH-ubiquinone oxidoreductase subunit B14.7) | NDUFA11 | mitochondrial electron transport, NADH to ubiquinone [GO:0006120] |

| | | | | | | | | | | | | | | | | | | | | | |
|---|---|---|---|---|---|---|---|---|---|---|---|---|---|---|---|---|---|---|---|---|---|
| 11575.38 | 12305.18 | 13632.98 | 8617.53 | 16218.03 | 23703.77 | 20374.88 | 6573.8 | 6006.83 | 5787.07 | 8215.14 | 15966.81 | 21447.84 | 27852.47 | 9203.05 | 20162.86 | 15244.21 | 7974.75 | 250862.58 | Sialidase-1 (EC 3.2.1.18) (Acetylneuraminyl hydrolase) (Lysosomal sialidase) (N-acetyl-alpha-neuraminidase 1) | NEU1 | ganglioside catabolic process [GO:0006689]; oligosaccharide catabolic process [GO:0009313] |
| 10635.66 | 20874.03 | 16024.13 | 8788.44 | 9637.72 | 43643.84 | 124.66 | 22166.21 | 9541.56 | 9379.01 | 6135.69 | 20480.72 | 12832.68 | 8306.28 | 9383.75 | 3111.08 | 14603.78 | 10353.42 | 236022.66 | Leukocyte antigen CD37 (Tetraspanin-26) (Tspan-26) (CD antigen CD37) | CD37 TSPAN26 | defense response to protozoan [GO:0042832]; negative regulation of myeloid dendritic cell activation [GO:0030886]; negative regulation of T cell proliferation [GO:0042130]; positive regulation of immunoglobulin production [GO:0002639]; regulation of defense response to virus [GO:0050688]; regulation of humoral immune response [GO:0002920]; T cell proliferation [GO:0042098] |
| 7.87 | 72009.1 | 23.49 | 6.91 | 36714.15 | 125.19 | 0.22 | 22718.53 | 22.36 | 4.52 | 14929.32 | 16.98 | 25.26 | 47700.4 | 43.97 | 4.67 | 31537.82 | 96.2 | 225986.96 | N-myc-interactor (Nmi) (N-myc and STAT interactor) | NMI | cell surface receptor signaling pathway via JAK-STAT [GO:0007259]; innate immune response [GO:0045087]; macrophage activation involved in immune response [GO:0002281]; negative regulation of cell population proliferation [GO:0008285]; negative regulation of innate immune response [GO:0045824]; negative regulation of interferon-alpha production [GO:0032687]; negative regulation of interferon-beta production [GO:0032688]; negative regulation of non-canonical NF-kappaB signal transduction [GO:1901223]; positive regulation of inflammatory response [GO:0050729]; positive regulation of innate immune response [GO:0045089]; positive regulation of non-canonical NF-kappaB signal transduction [GO:1901224]; positive regulation of protein K48-linked ubiquitination [GO:1902524]; protein K48-linked ubiquitination [GO:0070936]; response to virus [GO:0009615]; toll-like receptor 4 signaling pathway [GO:0034142] |

| | | | | | | | | | | | | | | | | | | | | | |
|---|---|---|---|---|---|---|---|---|---|---|---|---|---|---|---|---|---|---|---|---|---|
| 1160.03 | 4.97 | 0 | 3460.11 | 0 | 94.45 | 144058.5 | 6.87 | 5.49 | 1147.52 | 3219.8 | 7.56 | 146.8 | 0 | 0 | 64446.61 | 2339.43 | 63.46 | 220161.6 | Secretogranin-2 (Chromogranin-C) (Secretogranin II) (SgII) [Cleaved into: Secretoneurin (SN);Manserin] | SCG2 CHGC | angiogenesis [GO:0001525]; eosinophil chemotaxis [GO:0048245]; induction of positive chemotaxis [GO:0050930]; intracellular signal transduction [GO:0035556]; MAPK cascade [GO:0000165]; negative regulation of endothelial cell apoptotic process [GO:2000352]; negative regulation of extrinsic apoptotic signaling pathway [GO:2001237]; positive chemotaxis [GO:0050918]; positive regulation of endothelial cell proliferation [GO:0001938] |
| 561.64 | 4.64 | 5.9 | 9030.59 | 8.49 | 172.25 | 114392.87 | 3.08 | 4.81 | 1165.99 | 2018.9 | 22.98 | 164.12 | 39.98 | 5.76 | 86299.09 | 2431.43 | 67.47 | 216399.99 | Exosome complex component RRP43 (Exosome component 8) (Opa-interacting protein 2) (OIP-2) (Ribosomal RNA-processing protein 43) (p9) | EXOSC8 OIP2 RRP43 | exonucleolytic trimming to generate mature 3'-end of 5.8S rRNA from tricistronic rRNA transcript (SSU-rRNA, 5.8S rRNA, LSU-rRNA) [GO:0000467]; nuclear mRNA surveillance [GO:0071028]; nuclear polyadenylation-dependent rRNA catabolic process [GO:0071035]; RNA catabolic process [GO:0006401]; RNA processing [GO:0006396]; rRNA catabolic process [GO:0016075]; TRAMP-dependent tRNA surveillance pathway [GO:0071038]; U1 snRNA 3'-end processing [GO:0034473]; U4 snRNA 3'-end processing [GO:0034475]; U5 snRNA 3'-end processing [GO:0034476] |

| | | | | | | | | | | | | | | | | | | | | | |
|---|---|---|---|---|---|---|---|---|---|---|---|---|---|---|---|---|---|---|---|---|---|
| 13539.44 | 12970.11 | 23134.41 | 17680.12 | 19363.8 | 16096.13 | 1741.28 | 9779.45 | 5228.46 | 8554.36 | 9400.64 | 16160.26 | 9115.55 | 15588.93 | 7413.27 | 6094.08 | 13780.17 | 7206.85 | 212847.31 | Coronin-1C (Coronin-3) (hCRNN4) | CORO1C CRN2 CRNN4 | actin filament organization [GO:0007015]; activation of GTPase activity [GO:0090630]; corpus callosum development [GO:0022038]; endosomal transport [GO:0016197]; endosome fission [GO:0140285]; endosome membrane tubulation [GO:0097750]; establishment of protein localization [GO:0045184]; membrane fission [GO:0090148]; negative regulation of epithelial cell migration [GO:0010633]; negative regulation of focal adhesion assembly [GO:0051895]; negative regulation of protein kinase activity by regulation of protein phosphorylation [GO:0044387]; negative regulation of protein phosphorylation [GO:0001933]; negative regulation of substrate adhesion-dependent cell spreading [GO:1900025]; neural crest cell migration [GO:0001755]; phagocytosis [GO:0006909]; positive regulation of lamellipodium morphogenesis [GO:2000394]; regulation of epithelial cell migration [GO:0010632]; regulation of fibroblast migration [GO:0010762]; |
| 9772.55 | 8151.35 | 10689.8 | 9027.91 | 10507.61 | 17101.31 | 12387.19 | 5057.59 | 6272.47 | 15052.77 | 7325.95 | 26654.33 | 15956.85 | 9547.95 | 16749.11 | 13300.19 | 8181.69 | 10206.38 | 211943 | Bcl-2-like protein 11 (Bcl2-L-11) (Bcl-2-related ovarian death protein) (Bcl2-interacting mediator of cell death) | Bcl2l11 Bim Bod | apoptotic process involved in embryonic digit morphogenesis [GO:1902263]; B cell homeostasis [GO:0001782]; cell-matrix adhesion [GO:0007160]; cellular response to amyloid-beta [GO:1904646]; cellular response to estradiol stimulus [GO:0071392]; cellular response to nerve growth factor stimulus [GO:1990090]; developmental pigmentation [GO:0048066]; ear development [GO:0043583]; extrinsic apoptotic signaling pathway in absence of ligand [GO:0097192]; in utero embryonic development [GO:0001701]; intrinsic apoptotic signaling pathway in response to DNA damage [GO:0008630]; kidney development [GO:0001822]; leukocyte homeostasis [GO:0001776]; lymphocyte apoptotic process [GO:0070227]; lymphocyte homeostasis [GO:0002260]; male gonad development [GO:0008584]; mammary gland development [GO:0030879]; meiosis I [GO:0007127]; myeloid cell homeostasis [GO:0002262]; |
| 7759.07 | 5409.11 | 8049.52 | 7637.21 | 9769.67 | 28367.27 | 10359.42 | 5429.73 | 5448.85 | 12750.69 | 7831.23 | 9759.1 | 9746.7 | 19927.75 | 11641.88 | 22755.94 | 10110.83 | 11499.89 | 204253.86 | Protein-methionine sulfoxide oxidase mical3b (EC 1.14.13.225) (Molecule interacting with CasL protein 3B) (MICAL-3B) | mical3b | actin filament depolymerization [GO:0030042]; cytoskeleton organization [GO:0007010]; exocytosis [GO:0006887] |

| | | | | | | | | | | | | | | | | | | | | | |
|---|---|---|---|---|---|---|---|---|---|---|---|---|---|---|---|---|---|---|---|---|---|
| 294.85 | 0 | 0 | 941.17 | 0 | 63.15 | 164025.9 | 0 | 2.69 | 310.7 | 1273.17 | 0 | 43.75 | 0 | 0 | 34190.78 | 688.56 | 30.28 | 201865 | Peroxisomal membrane protein PMP34 (34 kDa peroxisomal membrane protein) (Solute carrier family 25 member 17) | SLC25A17 PMP34 | ATP transport [GO:0015867]; fatty acid alpha-oxidation [GO:0001561]; fatty acid beta-oxidation [GO:0006635]; fatty acid transport [GO:0015908]; nucleotide transmembrane transport [GO:1901679] |
| 8713.05 | 15841.11 | 5802.24 | 11300.47 | 14938.52 | 28042.64 | 760.03 | 16301.94 | 6131.81 | 14225.15 | 16038.28 | 10094.83 | 6896.88 | 23028.77 | 6159.11 | 2625.46 | 6780.56 | 5227.77 | 198908.62 | Inward rectifier potassium channel 2 (BIK) (Inward rectifier K(+) channel Kir2.1) (IRK-1) (Potassium channel, inwardly rectifying subfamily J member 2) | KCNJ2 IRK1 | cardiac muscle cell action potential involved in contraction [GO:0086002]; magnesium ion transport [GO:0015693]; potassium ion import across plasma membrane [GO:1990573]; potassium ion transport [GO:0006813]; protein homotetramerization [GO:0051289]; regulation of heart rate by cardiac conduction [GO:0086091]; regulation of membrane repolarization [GO:0060306]; regulation of monoatomic ion transmembrane transport [GO:0034765]; regulation of skeletal muscle contraction via regulation of action potential [GO:0014861]; relaxation of cardiac muscle [GO:0055119]; relaxation of skeletal muscle [GO:0090076] |
| 21530.71 | 20165.89 | 0 | 27998.52 | 13656.16 | 190.94 | 145.17 | 19958.58 | 0 | 23986.22 | 27757.05 | 21.46 | 10586.49 | 12820.09 | 20.83 | 5611.75 | 14199.64 | 54.22 | 198703.72 | Glycylpeptide N-tetradecanoyltransferase 2 (EC 2.3.1.97) (Myristoyl-CoA:protein N-myristoyltransferase 2) (NMT 2) (Peptide N-myristoyltransferase 2) (Type II N-myristoyltransferase) | nmt2 | N-terminal peptidyl-glycine N-myristoylation [GO:0018008]; protein localization to membrane [GO:0072657] |

| | | | | | | | | | | | | | | | | | | | | | | |
|---|---|---|---|---|---|---|---|---|---|---|---|---|---|---|---|---|---|---|---|---|---|---|
| 1540.93 | | 40847.28 | 9693.3 | 184.89 | 44136.59 | 15890.91 | 134.07 | 16498.81 | 643.01 | 677.56 | 6982.46 | 5513.39 | 613.75 | 35431.42 | 857.72 | 578.85 | 16873.89 | 757.57 | 197856.4 | Nucleosome-remodeling factor subunit BPTF (Bromodomain and PHD finger-containing transcription factor) (Fetal Alz-50 clone 1 protein) (Fetal Alzheimer antigen) | BPTF FAC1 FALZ | anterior/posterior pattern specification [GO:0009952]; brain development [GO:0007420]; cellular response to nerve growth factor stimulus [GO:1990090]; chromatin remodeling [GO:0006338]; embryonic placenta development [GO:0001892]; endoderm development [GO:0007492]; negative regulation of transcription by RNA polymerase II [GO:0000122]; positive regulation of transcription by RNA polymerase II [GO:0045944]; regulation of DNA-templated transcription [GO:0006355]; regulation of transcription by RNA polymerase II [GO:0006357] |
| 9971.97 | | 9593.77 | 20376.39 | 5262.64 | 7318.68 | 15460.78 | 11722.75 | 1277.06 | 2410.93 | 6561.8 | 6287.73 | 17240.5 | 19751.79 | 24616.62 | 8939.88 | 8733.62 | 8085.96 | 5728.08 | 189340.95 | Microtubule-associated serine/threonine-protein kinase 1 (EC 2.7.11.1) (Syntrophin-associated serine/threonine-protein kinase) | Mast1 Sast | brain development [GO:0007420]; cytoskeleton organization [GO:0007010]; intracellular signal transduction [GO:0035556]; protein phosphorylation [GO:0006468] |

| | | | | | | | | | | | | | | | | | | | | | |
|---|---|---|---|---|---|---|---|---|---|---|---|---|---|---|---|---|---|---|---|---|---|
| 0 | 53751.07 | 0 | 4.15 | 14273.16 | 60.48 | 0 | 25221.95 | 0 | 0.38 | 9668.01 | 1.63 | 1.68 | 37120.2 | 0 | 2.6 | 47457.36 | 190.3 | 187752.97 | Potassium voltage-gated channel subfamily KQT member 2 (KQT-like 2) (Potassium channel subunit alpha KvLQT2) (Voltage-gated potassium channel subunit Kv7.2) | Kcnq2 Kqt2 | action potential [GO:0001508]; action potential initiation [GO:0099610]; apoptosome assembly [GO:0097314]; brain development [GO:0007420]; cellular response to calcium ion [GO:0071277]; cellular response to xenobiotic stimulus [GO:0071466]; cognition [GO:0050890]; dentate gyrus development [GO:0021542]; determination of adult lifespan [GO:0008340]; endocytosis [GO:0006897]; establishment of cell polarity [GO:0030010]; excitatory chemical synaptic transmission [GO:0098976]; exocytosis [GO:0006887]; exploration behavior [GO:0035640]; gene expression [GO:0010467]; grooming behavior [GO:0007625]; hippocampal pyramidal neuron differentiation [GO:0097432]; hippocampus development [GO:0021766]; inhibitory chemical synaptic transmission [GO:0098977]; learning [GO:0007612]; memory [GO:0007613]; mitochondrial depolarization [GO:0051882]; |
| 7321.45 | 12132.91 | 6484.89 | 6929.43 | 14718.01 | 12371.48 | 47105.56 | 6542.52 | 2407.2 | 3906.01 | 6196.62 | 5048.71 | 8256.81 | 16778.11 | 2945.47 | 16623.3 | 8550.71 | 2782.57 | 187101.76 | Metabotropic glycine receptor (mGlyR) (G-protein coupled receptor 158) | Gpr158 | brain development [GO:0007420]; cognition [GO:0050890]; G protein-coupled receptor signaling pathway [GO:0007186]; positive regulation of neurotransmitter secretion [GO:0001956]; protein localization to plasma membrane [GO:0072659]; regulation of G protein-coupled receptor signaling pathway [GO:0008277]; regulation of synapse organization [GO:0050807] |

| | | | | | | | | | | | | | | | | | | | | | | |
|---|---|---|---|---|---|---|---|---|---|---|---|---|---|---|---|---|---|---|---|---|---|---|
| 633.36 | | 7553.21 | 2289.83 | 2512.41 | 20801.75 | 1685.11 | 28127.82 | 11517.87 | 16.91 | 716.46 | 5974.21 | 33656.37 | 328.23 | 29100.63 | 310.75 | 20012.93 | 16379.18 | 162.28 | 181779.31 | Mismatch repair endonuclease PMS2 (EC 3.1.-.-) (DNA mismatch repair protein PMS2) (PMS1 protein homolog 2) | PMS2 PMSL2 | mismatch repair [GO:0006298]; positive regulation of isotype switching to IgA isotypes [GO:0048298]; positive regulation of isotype switching to IgG isotypes [GO:0048304]; response to xenobiotic stimulus [GO:0009410]; somatic hypermutation of immunoglobulin genes [GO:0016446]; somatic recombination of immunoglobulin gene segments [GO:0016447] |
| 3421.58 | | 5833.38 | 0 | 26145.31 | 14145.25 | 140.99 | 110.07 | 14930.12 | 0 | 20261.65 | 30968.03 | 0 | 6551.63 | 10797.79 | 0 | 14389.43 | 32609.39 | 278.6 | 180583.22 | EF-hand domain-containing protein D2 (Swiprosin-1) | EFHD2 SWS1 | |
| 11.06 | | 84687.17 | 2.89 | 244.65 | 10652.1 | 34.47 | 0.45 | 16358.52 | 0 | 175.78 | 5591.79 | 2 | 4.79 | 44267.37 | 1.29 | 2.09 | 17606.8 | 30.33 | 179673.55 | Transcription initiation factor TFIID subunit 2 (TBP-associated factor 150 kDa) (Transcription initiation factor TFIID 150 kDa subunit) (TAF(II)150) (TAFII-150) (TAFII150) | taf2 | transcription initiation at RNA polymerase II promoter [GO:0006367] |

| | | | | | | | | | | | | | | | | | | | | | |
|---|---|---|---|---|---|---|---|---|---|---|---|---|---|---|---|---|---|---|---|---|---|
| 239.78 | 32.73 | 10204.4 | 89.78 | 3.38 | 4963.39 | 0.5 | 2.66 | 55.26 | 70.22 | 8.42 | 161352.68 | 513.66 | 2.99 | 782.81 | 15.12 | 10.48 | 359.26 | 178707.52 | Peroxisomal membrane protein PEX14 (PTS1 receptor-docking protein) (Peroxin-14) (Peroxisomal membrane anchor protein PEX14) | Pex14 | cellular response to reactive oxygen species [GO:0034614]; microtubule anchoring [GO:0034453]; peroxisome transport along microtubule [GO:0036250]; protein import into peroxisome matrix, docking [GO:0016560]; protein import into peroxisome matrix, substrate release [GO:0044721]; protein import into peroxisome matrix, translocation [GO:0016561]; protein-containing complex assembly [GO:0065003] |
| 37440.75 | 272.84 | 276.69 | 9089.22 | 1469.34 | 7159.96 | 162.8 | 100.87 | 4.77 | 35451.69 | 970.53 | 1824.03 | 76019.8 | 162.81 | 331.36 | 1893.29 | 325.52 | 17.26 | 172973.53 | 26S proteasome regulatory subunit 6A (26S proteasome AAA-ATPase subunit RPT5) (Proteasome 26S subunit ATPase 3) (Spermatogenic cell/sperm-associated Tat-binding protein homolog SATA) (Tat-binding protein 1) (TBP-1) | Psmc3 Tbp1 | blastocyst development [GO:0001824]; modulation by host of viral transcription [GO:0043921]; positive regulation of transcription by RNA polymerase II [GO:0045944]; proteasome-mediated ubiquitin-dependent protein catabolic process [GO:0043161] |

| | | | | | | | | | | | | | | | | | | | | | | |
|---|---|---|---|---|---|---|---|---|---|---|---|---|---|---|---|---|---|---|---|---|---|---|
| 4540.6 | | 6354.63 | 8516.53 | 5380.27 | 6041.8 | 8130.98 | 42587.88 | 2457.53 | 2766.09 | 2597.61 | 3227.68 | 7986.18 | 7138.66 | 17106.85 | 3746.37 | 28541.34 | 10519.12 | 4027.36 | 171667.48 | Activity-regulated cytoskeleton-associated protein (hArc) (Activity-regulated gene 3.1 protein homolog) (ARC/ARG3.1) (Arg3.1) | ARC KIAA0278 | anterior/posterior pattern specification [GO:0009952]; cell migration [GO:0016477]; cytoskeleton organization [GO:0007010]; dendritic spine morphogenesis [GO:0060997]; endocytosis [GO:0006897]; endoderm development [GO:0007492]; long-term memory [GO:0007616]; long-term synaptic potentiation [GO:0060291]; modulation of chemical synaptic transmission [GO:0050804]; mRNA transport [GO:0051028]; protein homooligomerization [GO:0051260]; regulation of cell morphogenesis [GO:0022604]; regulation of dendritic spine morphogenesis [GO:0061001]; regulation of long-term synaptic depression [GO:1900452]; regulation of long-term synaptic potentiation [GO:1900271]; regulation of neuronal synaptic plasticity [GO:0048168]; regulation of postsynaptic neurotransmitter receptor internalization [GO:0099149]; vesicle-mediated intercellular transport [GO:0110077] |
| 3718.61 | | 8252.91 | 7471.32 | 6362.35 | 11662.97 | 12002.25 | 25020.64 | 5756.09 | 3430.16 | 3757.27 | 5976.02 | 6554.83 | 7240.19 | 17491.49 | 4844.67 | 23967.08 | 10676.89 | 2868.66 | 167054.4 | Immediate early response gene 2 protein (Protein ETR101) | IER2 ETR101 PIP92 | cell motility [GO:0048870]; neuron differentiation [GO:0030182]; positive regulation of transcription by RNA polymerase II [GO:0045944]; response to fibroblast growth factor [GO:0071774] |

| | | | | | | | | | | | | | | | | | | | | | |
|---|---|---|---|---|---|---|---|---|---|---|---|---|---|---|---|---|---|---|---|---|---|
| 420.5 | 486.88 | 27653.55 | 276.96 | 214.12 | 52737.51 | 34.76 | 219.87 | 622.23 | 1024.55 | 832.48 | 70601.95 | 270.97 | 378.38 | 7959.09 | 939.82 | 368.65 | 771.72 | 165813.99 | Dihydrolipoyllysine-residue succinyltransferase component of 2-oxoglutarate dehydrogenase complex, mitochondrial (EC 2.3.1.61) (2-oxoglutarate dehydrogenase complex component E2) (OGDC-E2) (Dihydrolipoamide succinyltransferase component of 2-oxoglutarate dehydrogenase complex) (E2K) | dlst | 2-oxoglutarate metabolic process [GO:0006103]; L-lysine catabolic process to acetyl-CoA via saccharopine [GO:0033512]; succinyl-CoA metabolic process [GO:0006104]; tricarboxylic acid cycle [GO:0006099] |
| 8809.37 | 8025.54 | 8619.89 | 7215.68 | 8516.51 | 15337 | 12111.43 | 2806.82 | 3359.36 | 5147.11 | 5173.19 | 12795.92 | 12954.57 | 16716.12 | 5105.12 | 12050.64 | 10261.87 | 4724.47 | 159730.61 | GDNF family receptor alpha-4 (GDNF receptor alpha-4) (GDNFR-alpha-4) (GFR-alpha-4) | GFRA4 | glial cell-derived neurotrophic factor receptor signaling pathway [GO:0035860]; nervous system development [GO:0007399] |
| 7847.49 | 7831.37 | 11697.75 | 2971.88 | 14345.28 | 19640.92 | 55.12 | 6566.38 | 1120.45 | 3841.71 | 5066.01 | 6263.86 | 23923.11 | 18125.07 | 11921.23 | 1692.88 | 8838.52 | 6600.94 | 158349.97 | Intermembrane lipid transfer protein VPS13B (Cohen syndrome protein 1) (Vacuolar protein sorting-associated protein 13B) | VPS13B CHS1 COH1 KIAA0532 | acrosome assembly [GO:0001675]; adipose tissue development [GO:0060612]; central nervous system development [GO:0007417]; dentate gyrus development [GO:0021542]; Golgi organization [GO:0007030]; Golgi reassembly [GO:0090168]; head morphogenesis [GO:0060323]; lipid transport [GO:0006869]; maintenance of lens transparency [GO:0036438]; memory [GO:0007613]; multicellular organism growth [GO:0035264]; muscle organ development [GO:0007517]; nervous system development [GO:0007399]; neuron projection development [GO:0031175]; slow endocytic recycling [GO:0032458]; social behavior [GO:0035176]; vesicle-mediated transport [GO:0016192] |

| | | | | | | | | | | | | | | | | | | | | | |
|---|---|---|---|---|---|---|---|---|---|---|---|---|---|---|---|---|---|---|---|---|---|
| 305.09 | 0 | 0 | 4705.98 | 0 | 100.2 | 111199.19 | 0 | 0 | 428.26 | 1326.06 | 0 | 41.4 | 0 | 4.35 | 38751.64 | 848.25 | 5.34 | 157715.76 | Centrosomal protein of 44 kDa (Cep44) | cep44 | centriole replication [GO:0007099]; centriole-centriole cohesion [GO:0010457]; centrosome cycle [GO:0007098] |
| 0 | 19628.17 | 0 | 11.45 | 36534.32 | 230.16 | 0 | 16348.73 | 0 | 5.5 | 9679.05 | 0 | 0 | 54505.83 | 0 | 0 | 16074.52 | 56.32 | 153074.05 | Neurabin-2 (Neurabin-II) (Protein phosphatase 1 regulatory subunit 9B) (Spinophilin) | PPP1R9B PPP1R6 | actin filament depolymerization [GO:0030042]; actin filament organization [GO:0007015]; calcium-mediated signaling [GO:0019722]; cell migration [GO:0016477]; cellular response to epidermal growth factor stimulus [GO:0071364]; cellular response to estradiol stimulus [GO:0071392]; cellular response to morphine [GO:0071315]; cellular response to peptide [GO:1901653]; cellular response to xenobiotic stimulus [GO:0071466]; cerebral cortex development [GO:0021987]; dendrite development [GO:0016358]; developmental process involved in reproduction [GO:0003006]; filopodium assembly [GO:0046847]; hippocampus development [GO:0021766]; learning [GO:0007612]; male mating behavior [GO:0060179]; negative regulation of cell growth [GO:0030308]; neuron projection development [GO:0031175]; positive regulation of protein localization to actin cortical patch [illegible] |
| 34732.03 | 277.79 | 234.54 | 6066.06 | 887.6 | 2616.96 | 74.15 | 65.44 | 0 | 19186.18 | 708.67 | 1202.11 | 83986.2 | 206.58 | 260.84 | 1808.71 | 324.58 | 16.66 | 152655.1 | Dixin-A (Coiled-coil protein DIX1-A) (Coiled-coil-DIX1-A) (DIX domain-containing protein 1-A) | dixdc1a ccd1 dixdc1 | camera-type eye development [GO:0043010]; canonical Wnt signaling pathway [GO:0060070]; forebrain development [GO:0030900] |

| | | | | | | | | | | | | | | | | | | | | |
|---|---|---|---|---|---|---|---|---|---|---|---|---|---|---|---|---|---|---|---|---|
| 118.97 | 0 | 0 | 1884.31 | 0 | 69.92 | 100441.18 | 0 | 0 | 469.61 | 1192.51 | 0 | 30.76 | 0 | 0 | 42461.72 | 1117.19 | 52.71 | 147838.88 | Collectin-12 (Collectin placenta protein 1) (CL-P1) (Nurse cell scavenger receptor 2) | Colec12 Clp1 Nsr2 | cellular response to exogenous dsRNA [GO:0071360]; defense response to bacterium [GO:0042742]; immune response [GO:0006955]; phagocytosis, recognition [GO:0006910]; plasma membrane raft organization [GO:0044857]; toll-like receptor 3 signaling pathway [GO:0034138] |
| 897.99 | 627.93 | 311.25 | 1838.32 | 629.48 | 915.97 | 77748.53 | 160.9 | 120.58 | 1066.78 | 1730.58 | 233.43 | 3737.61 | 716.48 | 590.86 | 54358.18 | 1603.15 | 282.19 | 147570.21 | Protein Aster-B (GRAM domain-containing protein 1B) | GRAMD1B KIAA1201 UNQ3032/PRO9834 | cellular response to cholesterol [GO:0071397]; cholesterol homeostasis [GO:0042632]; intracellular sterol transport [GO:0032366] |
| 119.26 | 3.59 | 19.45 | 914.61 | 3.79 | 62.9 | 103940.76 | 0 | 0 | 583.66 | 1204.11 | 57.37 | 22.9 | 4.29 | 8.68 | 38359.76 | 780.53 | 47.49 | 146133.15 | Mitogen-activated protein kinase-binding protein 1 (JNK-binding protein 1) (JNKBP-1) | MAPKBP1 JNKBP1 KIAA0596 | negative regulation of canonical NF-kappaB signal transduction [GO:0043124]; negative regulation of defense response to bacterium [GO:1900425]; negative regulation of interleukin-8 production [GO:0032717]; positive regulation of JNK cascade [GO:0046330] |

| | | | | | | | | | | | | | | | | | | | | | |
|---|---|---|---|---|---|---|---|---|---|---|---|---|---|---|---|---|---|---|---|---|---|
| 5264.08 | 4443.26 | 4799.07 | 3707.05 | 4309.01 | 10311.73 | 29.22 | 71147.62 | 10018.04 | 4264.36 | 5328.3 | 1263.97 | 1779.42 | 9922.26 | 1606.35 | 537 | 3486.8 | 2162.03 | 144379.57 | Y+L amino acid transporter 1 (Monocyte amino acid permease 2) (MOP-2) (Solute carrier family 7 member 7) (y(+)L-type amino acid transporter 1) (Y+LAT1) (y+LAT-1) | SLC7A7 | amino acid transmembrane transport [GO:0003333]; basic amino acid transmembrane transport [GO:1990822]; L-arginine transmembrane transport [GO:1903826]; L-leucine transport [GO:0015820]; regulation of arginine metabolic process [GO:0000821] |
| 9214.37 | 14775 | 10673.02 | 2793.44 | 16485.84 | 20125.96 | 76.61 | 5237.93 | 626.52 | 1936.12 | 5600.04 | 6630.22 | 7871.71 | 19875.89 | 6582.05 | 1673.11 | 7836.3 | 4816.21 | 142830.34 | Vasoactive intestinal polypeptide receptor 2 (VIP-R-2) (Pituitary adenylate cyclase-activating polypeptide type III receptor) (PACAP type III receptor) (PACAP-R-3) (PACAP-R3) | Vipr2 | adenylate cyclase-activating G protein-coupled receptor signaling pathway [GO:0007189]; adenylate cyclase-modulating G protein-coupled receptor signaling pathway [GO:0007188]; cell surface receptor signaling pathway [GO:0007166]; negative regulation of smooth muscle cell proliferation [GO:0048662] |

| | | | | | | | | | | | | | | | | | | | | | |
|---|---|---|---|---|---|---|---|---|---|---|---|---|---|---|---|---|---|---|---|---|---|
| 12507.29 | 7561.08 | 6948.51 | 6006.73 | 11712.51 | 10849.59 | 17209.39 | 4381.3 | 3478.31 | 4906.62 | 4494.42 | 5045.73 | 8355.07 | 12784.57 | 2453.95 | 13646.93 | 6942 | 3022.92 | 142306.92 | FACT complex subunit SPT16 (Chromatin-specific transcription elongation factor 140 kDa subunit) (FACT 140 kDa subunit) (FACTp140) (Facilitates chromatin transcription complex subunit SPT16) | Supt16h Fact140 Factp140 Supt16 | DNA repair [GO:0006281]; DNA replication [GO:0006260]; nucleosome assembly [GO:0006334]; nucleosome disassembly [GO:0006337] |
| 6423.65 | 11939.6 | 15447.46 | 2031.27 | 10835.97 | 14727.01 | 496.82 | 11248.19 | 1376.13 | 2019.14 | 5109.7 | 8328.89 | 12120.65 | 16713.2 | 5993.38 | 1527.38 | 11032.12 | 4100.92 | 141471.48 | Androgen-induced gene 1 protein (AIG-1) (Fatty acid esters of hydroxy fatty acids hydrolase AIG1) (FAHFA hydrolase AIG1) (EC 3.1.-.-) | AIG1 CGI-103 | long-chain fatty acid catabolic process [GO:0042758] |
| 36406.51 | 295.12 | 117.51 | 6778.04 | 923.75 | 2045.08 | 77.73 | 119.07 | 3.06 | 22923.94 | 843.69 | 1156.38 | 66292.2 | 139.14 | 158.42 | 1622.45 | 264.15 | 30.67 | 140196.91 | SEC14 domain and spectrin repeat-containing protein 1 (Protein Solo) | sestd1 sol solo si:dz106a20.1 si:dz42h5.2 | blood vessel morphogenesis [GO:0048514] |
| 3305.5 | 1545.84 | 3261.51 | 2650.96 | 1737.4 | 5533.36 | 51164.88 | 1072.72 | 5095.18 | 1948.31 | 1546.1 | 3500.64 | 8672.51 | 2265.52 | 4406.86 | 33448.2 | 3117.67 | 4985.25 | 139258.41 | LIM domain-binding protein 3 (Protein cypher) (Protein oracle) (Z-band alternatively spliced PDZ-motif protein) | Ldb3 Kiaa0613 | sarcomere organization [GO:0045214] |

| | | | | | | | | | | | | | | | | | | | | |
|---|---|---|---|---|---|---|---|---|---|---|---|---|---|---|---|---|---|---|---|---|
| 13823.23 | 11780.55 | 8068.18 | 7141.56 | 9390.69 | 16274.44 | 136 | 6258.6 | 11800.31 | 12622.51 | 12220.97 | 9670.16 | 2842.68 | 5292.27 | 6191.57 | 150.95 | 750.93 | 2342.72 | 136758.32 | Integrin alpha-7 (H36-alpha7) [Cleaved into: Integrin alpha-7 heavy chain;Integrin alpha-7 light chain;Integrin alpha-7 70 kDa form] | Itga7 | blood vessel morphogenesis [GO:0048514]; cell adhesion [GO:0007155]; cell adhesion mediated by integrin [GO:0033627]; cell migration [GO:0016477]; cell-cell adhesion [GO:0098609]; endodermal cell differentiation [GO:0035987]; heterotypic cell-cell adhesion [GO:0034113]; integrin-mediated signaling pathway [GO:0007229]; leukocyte migration [GO:0050900]; regulation of cell shape [GO:0008360]; skeletal muscle tissue development [GO:0007519] |
| 30330.28 | 255.87 | 141.28 | 5749.25 | 1066.42 | 1491.99 | 67.93 | 98.53 | 3.31 | 17070.6 | 668.3 | 1011.85 | 74633.7 | 130.05 | 185.82 | 2157.41 | 363.26 | 17.71 | 135443.56 | High affinity cGMP-specific 3',5'-cyclic phosphodiesterase 9A (EC 3.1.4.35) | Pde9a | cAMP-mediated signaling [GO:0019933]; cGMP catabolic process [GO:0046069]; cGMP metabolic process [GO:0046068]; negative regulation of neural precursor cell proliferation [GO:2000178]; positive regulation of cardiac muscle hypertrophy [GO:0010613]; positive regulation of long-term synaptic potentiation [GO:1900273] |

| | | | | | | | | | | | | | | | | | | | | | |
|---|---|---|---|---|---|---|---|---|---|---|---|---|---|---|---|---|---|---|---|---|---|
| 33142.19 | 261.41 | 140.4 | 6149.95 | 656.81 | 1108.94 | 79.51 | 57.87 | 0.96 | 28794.37 | 588.37 | 1218.06 | 60653.06 | 104.32 | 105.58 | 1607.5 | 251.42 | 16.12 | 134936.84 | Sodium/hydrogen exchanger 6 (Na(+)/H(+) exchanger 6) (NHE-6) (Sodium/hydrogen exchanger) (Solute carrier family 9 member 6) | Slc9a6 | axon extension [GO:0048675]; brain-derived neurotrophic factor receptor signaling pathway [GO:0031547]; dendrite extension [GO:0097484]; dendritic spine development [GO:0060996]; establishment of cell polarity [GO:0030010]; glial cell activation [GO:0061900]; neuron projection morphogenesis [GO:0048812]; potassium ion transmembrane transport [GO:0071805]; proton transmembrane transport [GO:1902600]; regulation of intracellular pH [GO:0051453]; regulation of neurotrophin TRK receptor signaling pathway [GO:0051386]; regulation of postsynaptic membrane neurotransmitter receptor levels [GO:0099072]; regulation of synaptic vesicle lumen acidification [GO:1901546]; sodium ion import across plasma membrane [GO:0098719]; synapse organization [GO:0050808] |
| 4391.63 | 5926.03 | 6850.55 | 5130.31 | 8543.07 | 6922.81 | 20832.55 | 3311.11 | 7023.06 | 2416.85 | 4057.15 | 5375.76 | 5880.94 | 12226.27 | 5637.61 | 13756.82 | 7446.41 | 6995.88 | 132724.81 | Transmembrane protein 101 | TMEM101 | |
| 48412.44 | 1406.11 | 10793.09 | 9409.36 | 1799.48 | 20543.36 | 2014.44 | 890.03 | 2703.78 | 8917.91 | 740.17 | 4745.39 | 7225.03 | 1110.9 | 2454.71 | 2790.59 | 1793.34 | 4539.77 | 132289.9 | Dapper homolog 2 (Frodo 2) | dact2 dpr2 frd2 zgc:152832 | mesoderm morphogenesis [GO:0048332]; negative regulation of nodal signaling pathway [GO:1900108]; Wnt signaling pathway [GO:0016055] |

| | | | | | | | | | | | | | | | | | | | | | |
|---|---|---|---|---|---|---|---|---|---|---|---|---|---|---|---|---|---|---|---|---|---|
| 102.47 | 60270.24 | 22.87 | 261.93 | 28905.15 | 162.83 | 1.11 | 13427.36 | 6.59 | 12.66 | 2717.54 | 0 | 33.58 | 20589.13 | 4.93 | 5.87 | 5452.84 | 40.27 | 132017.37 | Cadherin-related family member 2 | Cdhr2 | cell-cell adhesion mediated by cadherin [GO:0044331]; epithelial cell differentiation [GO:0030855]; homophilic cell adhesion via plasma membrane adhesion molecules [GO:0007156]; intermicrovillar adhesion [GO:0090675]; negative regulation of cell growth involved in contact inhibition [GO:0060243]; regulation of microvillus length [GO:0032532] |
| 207.99 | 0 | 0 | 2949.51 | 0 | 29.04 | 83448.17 | 0 | 0 | 440.78 | 1063.2 | 0 | 0 | 0 | 0 | 42797.51 | 935 | 0 | 131871.2 | Transient receptor potential cation channel subfamily A member 1 (Ankyrin-like with transmembrane domains protein 1) (Transformation-sensitive protein p120) (p120) (Wasabi receptor) | TRPA1 ANKTM1 | calcium ion transmembrane import into cytosol [GO:0097553]; calcium ion transmembrane transport [GO:0070588]; cell surface receptor signaling pathway [GO:0007166]; cellular response to caffeine [GO:0071313]; cellular response to carbon dioxide [GO:0071244]; cellular response to cold [GO:0070417]; cellular response to food [GO:0071240]; cellular response to heat [GO:0034605]; cellular response to hydrogen peroxide [GO:0070301]; cellular response to toxic substance [GO:0097237]; detection of chemical stimulus involved in sensory perception of pain [GO:0050968]; detection of mechanical stimulus involved in sensory perception of pain [GO:0050966]; intracellular calcium ion homeostasis [GO:0006874]; monoatomic ion transport [GO:0006811]; positive regulation of insulin secretion involved in cellular response to glucose stimulus [GO:0035774]; positive regulation of monoatomic anion transport |

| | | | | | | | | | | | | | | | | | | | | | |
|---|---|---|---|---|---|---|---|---|---|---|---|---|---|---|---|---|---|---|---|---|---|
| 10235.64 | 8950.65 | 6777.07 | 7687.92 | 9749.53 | 13635 | 38.77 | 6950.39 | 9995.83 | 10204.12 | 9931.42 | 12600.82 | 3655.04 | 3600.6 | 5266.12 | 536.74 | 2783.98 | 9175.49 | 131775.13 | Nuclear receptor subfamily 1 group D member 1 (Rev-erbA-alpha) (V-erbA-related protein 1) (EAR-1) | Nr1d1 | cell differentiation [GO:0030154]; cellular response to interleukin-1 [GO:0071347]; cellular response to lipopolysaccharide [GO:0071222]; cellular response to tumor necrosis factor [GO:0071356]; cholesterol homeostasis [GO:0042632]; circadian regulation of gene expression [GO:0032922]; circadian rhythm [GO:0007623]; circadian temperature homeostasis [GO:0060086]; glycogen biosynthetic process [GO:0005978]; hormone-mediated signaling pathway [GO:0009755]; intracellular glucose homeostasis [GO:0001678]; intracellular receptor signaling pathway [GO:0030522]; negative regulation of astrocyte activation [GO:0061889]; negative regulation of canonical NF-kappaB signal transduction [GO:0043124]; negative regulation of cold-induced thermogenesis [GO:0120163]; negative regulation of DNA-templated transcription [GO:0045892]; negative regulation of inflammatory response [GO:0050728]; |
| 5795.02 | 10843.2 | 3524.82 | 7636.47 | 10458.52 | 20495.68 | 503.69 | 9728.28 | 3053.39 | 9329.81 | 11661.69 | 6552.98 | 3826.93 | 11942.73 | 3347.42 | 1880.67 | 4067.8 | 2725.44 | 127374.54 | TRIO and F-actin-binding protein (Protein Tara) (Trio-associated repeat on actin) | Triobp Kiaa1662 Tara | actin filament bundle assembly [GO:0051017]; auditory receptor cell stereocilium organization [GO:0060088]; cell division [GO:0051301]; positive regulation of substrate adhesion-dependent cell spreading [GO:1900026]; sensory perception of sound [GO:0007605] |
| 32548.95 | 291.08 | 99.5 | 6455.29 | 844.07 | 2172.91 | 88.01 | 63.02 | 0 | 20021.05 | 674.63 | 1345.04 | 60447.15 | 163.41 | 144.68 | 1340.42 | 248.04 | 2.89 | 126950.14 | RNA-binding protein PNO1 | pno1 | ribosome biogenesis [GO:0042254] |

| | | | | | | | | | | | | | | | | | | | | | |
|---|---|---|---|---|---|---|---|---|---|---|---|---|---|---|---|---|---|---|---|---|---|
| 26124.58 | 229.81 | 152.69 | 6032.92 | 612.28 | 2376.43 | 76.98 | 49.52 | 0 | 22919.54 | 515.11 | 1332.7 | 58950.21 | 121.49 | 140.64 | 1588.1 | 182.26 | 8.29 | 121413.55 | NMDA receptor synaptonuclear signaling and neuronal migration factor (Nasal embryonic luteinizing hormone-releasing hormone factor) (Nasal embryonic LHRH factor) | nsmf nelf nelfb | positive regulation of axon extension [GO:0045773]; positive regulation of neuron migration [GO:2001224]; regulation of neuronal synaptic plasticity [GO:0048168] |
| 9544.12 | 4177.58 | 11220.12 | 2471.54 | 15183.77 | 13790.7 | 62.96 | 7572.6 | 739.15 | 3783.62 | 7688.33 | 11624.39 | 6483.19 | 9305.14 | 3074.81 | 2725.18 | 9950.04 | 1655.49 | 121052.73 | Core histone macro-H2A.1 (Histone macroH2A1) (mH2A1) (H2A.y) (H2A/y) | Macroh2a1 H2afy | DNA repair [GO:0006281]; epigenetic regulation of gene expression [GO:0040029]; establishment of protein localization to chromatin [GO:0071169]; heterochromatin formation [GO:0031507]; negative regulation of cell cycle G2/M phase transition [GO:1902750]; negative regulation of gene expression, epigenetic [GO:0045814]; negative regulation of protein localization to chromosome, telomeric region [GO:1904815]; negative regulation of transcription by RNA polymerase II [GO:0000122]; negative regulation of transcription of nucleolar large rRNA by RNA polymerase I [GO:1901837]; nucleosome assembly [GO:0006334]; positive regulation of endodermal cell differentiation [GO:1903226]; positive regulation of keratinocyte differentiation [GO:0045618]; positive regulation of maintenance of mitotic sister chromatid cohesion [GO:0034184]; regulation of lipid metabolic process [GO:0019216]; |
| 205.13 | 0 | 0 | 452.97 | 0 | 4.84 | 68627.87 | 0 | 0 | 1008.64 | 1870.9 | 1.89 | 47.33 | 0 | 0 | 46851.35 | 1317 | 29.56 | 120417.48 | E3 ubiquitin-protein ligase KCMF1 (EC 2.3.2.27) | kcmf1 si:ch211-220b11.7 | negative regulation of HRI-mediated signaling [GO:0141191]; protein K63-linked ubiquitination [GO:0070534]; synaptic signaling [GO:0099536] |

| | | | | | | | | | | | | | | | | | | | | | |
|---|---|---|---|---|---|---|---|---|---|---|---|---|---|---|---|---|---|---|---|---|---|
| 4770.12 | 5490.42 | 4608.59 | 4563.79 | 8234.69 | 9001.56 | 13618.86 | 3056.34 | 2711.24 | 4270.7 | 4627.02 | 8376.91 | 7512.74 | 12552.8 | 3217.42 | 10541.14 | 7913.88 | 3583.13 | 118651.35 | Proline-serine-threonine phosphatase-interacting protein 2 (PEST phosphatase-interacting protein 2) | PSTPIP2 | |
| 8108.34 | 13653.02 | 8409.23 | 1625.16 | 17017.82 | 10801.29 | 187.43 | 3690.93 | 738.92 | 2950.53 | 8242.3 | 10954.42 | 6095.55 | 11968.69 | 2827.85 | 1501.38 | 7876 | 1948.9 | 118597.76 | Phosphatidylinositol 3,4,5-trisphosphate-dependent Rac exchanger 2 protein (P-Rex2) (PtdIns(3,4,5)-dependent Rac exchanger 2) (DEP domain-containing protein 2) | PREX2 DEPDC2 | adult locomotory behavior [GO:0008344]; dendrite morphogenesis [GO:0048813]; G protein-coupled receptor signaling pathway [GO:0007186]; negative regulation of TOR signaling [GO:0032007]; phosphatidylinositol 3-kinase/protein kinase B signal transduction [GO:0043491]; regulation of small GTPase mediated signal transduction [GO:0051056] |
| 6050.13 | 5838.48 | 8460.75 | 4815.17 | 6690.05 | 11256.99 | 4873.26 | 2282.61 | 3588.76 | 3364.45 | 3763.57 | 10482.81 | 9284.61 | 12637.83 | 4083.92 | 7468.06 | 7901.27 | 4365.77 | 117208.49 | 4-galactosyl-N-acetylglucosaminide 3-alpha-L-fucosyltransferase 9 (EC 2.4.1.152) (Fucosyltransferase 9) (Fucosyltransferase IX) (Fuc-TIX) (FucT-IX) (Galactoside 3-L-fucosyltransferase) | Fut9 | fucosylation [GO:0036065]; glycosphingolipid biosynthetic process [GO:0006688]; Lewis x epitope biosynthetic process [GO:0106402]; N-glycan fucosylation [GO:0036071]; neuron differentiation [GO:0030182]; neuronal stem cell division [GO:0036445]; oligosaccharide biosynthetic process [GO:0009312]; polysaccharide biosynthetic process [GO:0000271]; positive regulation of neuron projection development [GO:0010976]; protein glycosylation [GO:0006486]; protein N-linked glycosylation [GO:0006487]; protein O-linked glycosylation [GO:0006493]; regulation of leukocyte cell-cell adhesion [GO:1903037]; regulation of leukocyte tethering or rolling [GO:1903236] |

| | | | | | | | | | | | | | | | | | | | | | |
|---|---|---|---|---|---|---|---|---|---|---|---|---|---|---|---|---|---|---|---|---|---|
| 3650.58 | 1245.06 | 1898.71 | 2189.73 | 762.92 | 1999 | 45802.35 | 495.29 | 3691.59 | 1306.95 | 1183.51 | 1620.9 | 4422.85 | 3385.83 | 2671.77 | 32723.59 | 2150.83 | 2956.43 | 114157.89 | Ecto-ADP-ribosyltransferase 4 (EC 2.4.2.31) (ADP-ribosyltransferase C2 and C3 toxin-like 4) (ARTC4) (Mono(ADP-ribosyl)transferase 4) (NAD(P)(+)--arginine ADP-ribosyltransferase 4) (CD antigen CD297) | Art4 | |
| 405.43 | 150.56 | 247.47 | 1566.68 | 145.05 | 316.69 | 51411.74 | 58.4 | 215.32 | 492.77 | 1320.07 | 194.58 | 374.57 | 201.7 | 276.3 | 54973.38 | 1384.76 | 287.68 | 114023.15 | Erythroid differentiation-related factor 1 | EDRF1 C10orf137 | positive regulation of DNA-templated transcription [GO:0045893] |
| 3820.22 | 8157.78 | 5187.65 | 3827.21 | 11062.63 | 5824.31 | 10342.04 | 4763.61 | 2616.56 | 3741.96 | 6539.3 | 6009.81 | 3952.8 | 12751.12 | 2536.33 | 8291.71 | 9840.78 | 3878.22 | 113144.04 | Uncharacterized protein K02A2.6 | K02A2.6 | DNA integration [GO:0015074] |
| 23422.9 | 326.7 | 158.46 | 11149.24 | 1805.04 | 6706.04 | 166.82 | 412.75 | 17.25 | 24410.36 | 783.59 | 699.6 | 39222.64 | 165.83 | 214.87 | 2913.54 | 416.26 | 88.48 | 113080.37 | Probable methyltransferase-like protein 25 (EC 2.1.1.-) | METTL25 C12orf26 | methylation [GO:0032259] |
| 120.71 | 0 | 0 | 635.19 | 0 | 3.06 | 61055.81 | 0 | 0 | 510.5 | 1339.18 | 0 | 10.62 | 0 | 0 | 46990.29 | 1205.09 | 29.55 | 111900 | Nck-associated protein 5-like (NCKAP5-like) (Centrosomal protein of 169 kDa) (Cep169) | NCKAP5L CEP169 KIAA1602 FP1193 | microtubule bundle formation [GO:0001578]; microtubule depolymerization [GO:0007019] |

| | | | | | | | | | | | | | | | | | | | | | |
|---|---|---|---|---|---|---|---|---|---|---|---|---|---|---|---|---|---|---|---|---|---|
| 5016.76 | 3745.33 | 4945.83 | 4644.77 | 5017.03 | 11054.76 | 14103.28 | 1670.75 | 2879.71 | 3346.87 | 3132.83 | 7707.26 | 7075.63 | 8968.99 | 3552.93 | 13448.12 | 6136.27 | 3766.01 | 110213.13 | ATP-binding cassette sub-family C member 8 (Sulfonylurea receptor 1) | Abcc8 Sur Sur1 | action potential [GO:0001508]; cellular response to nutrient levels [GO:0031669]; female pregnancy [GO:0007565]; glutamate secretion, neurotransmission [GO:0061535]; intracellular glucose homeostasis [GO:0001678]; memory [GO:0007613]; negative regulation of angiogenesis [GO:0016525]; negative regulation of blood-brain barrier permeability [GO:1905604]; negative regulation of glial cell proliferation [GO:0060253]; negative regulation of insulin secretion [GO:0046676]; negative regulation of low-density lipoprotein particle clearance [GO:0010989]; negative regulation of neuroblast migration [GO:0061855]; negative regulation of neurogenesis [GO:0050768]; neuromuscular process [GO:0050905]; positive regulation of insulin secretion involved in cellular response to glucose stimulus [GO:0035774]; positive regulation of potassium ion transport [GO:0043268]; positive regulation of tight junction disassembly |
| 94.57 | 1.01 | 45.72 | 1356.03 | 0 | 40.34 | 83330.33 | 0 | 3.05 | 265.97 | 661.03 | 25.85 | 14.99 | 0 | 3.49 | 21558.52 | 461.44 | 6.57 | 107868.91 | NAD-dependent protein deacylase sirtuin-6 (EC 2.3.1.-) (NAD-dependent protein deacetylase sirtuin-6) (EC 2.3.1.286) (Protein mono-ADP-ribosyltransferase sirtuin-6) (EC 2.4.2.-) | SIRT6 | cardiac muscle cell differentiation [GO:0055007]; circadian regulation of gene expression [GO:0032922]; determination of adult lifespan [GO:0008340]; double-strand break repair [GO:0006302]; ketone biosynthetic process [GO:0042181]; negative regulation of gluconeogenesis [GO:0045721]; negative regulation of glycolytic process [GO:0045820]; negative regulation of protein import into nucleus [GO:0042308]; negative regulation of transcription by RNA polymerase II [GO:0000122]; negative regulation of transcription elongation by RNA polymerase II [GO:0034244]; pericentric heterochromatin formation [GO:0031508]; positive regulation of cold-induced thermogenesis [GO:0120162]; positive regulation of double-strand break repair [GO:2000781]; positive regulation of fat cell differentiation [GO:0045600]; positive regulation of insulin secretion [GO:0032024]; positive regulation of protein export from nucleus |
| 8447.03 | 5803.85 | 8686.98 | 4423.87 | 11347.97 | 12019.37 | 959.35 | 5493.47 | 1848.64 | 4409.08 | 4258.9 | 9691.62 | 5670.03 | 7538.88 | 2626.69 | 2544.38 | 6649.89 | 2825.71 | 105245.71 | Spermatid perinuclear RNA-binding protein | STRBP | cell differentiation [GO:0030154]; spermatogenesis [GO:0007283] |

| | | | | | | | | | | | | | | | | | | | | | |
|---|---|---|---|---|---|---|---|---|---|---|---|---|---|---|---|---|---|---|---|---|---|
| 0 | 34703.25 | 0 | 0 | 6644.64 | 0 | 0 | 7921.53 | 0 | 0 | 4268.15 | 0 | 0 | 36161.07 | 0 | 0 | 13000.05 | 92.85 | 102791.54 | Transcription factor IIIB 90 kDa subunit (TFIIIB90) (mTFIIIB90) (B-related factor 1) (BRF-1) | Brf1 | regulation of mRNA stability [GO:0043488]; transcription preinitiation complex assembly [GO:0070897] |
| 10857.91 | 4666.7 | 9630.43 | 3295.54 | 4083.01 | 7615.27 | 5479.99 | 1263.44 | 3194.5 | 2315.77 | 1078.18 | 3950.57 | 17475.3 | 4686.02 | 5123.19 | 5215.63 | 7172.47 | 4334.02 | 101437.94 | Phospholipid phosphatase-related protein type 1 (Inactive 2-lysophosphatidate phosphatase PLPPR1) (Lipid phosphate phosphatase-related protein type 1) | plppr1 lppr1 zgc:86759 | nervous system development [GO:0007399]; phospholipid dephosphorylation [GO:0046839]; phospholipid metabolic process [GO:0006644]; signal transduction [GO:0007165] |
| 5866.19 | 4926.82 | 6408.47 | 4988.08 | 4874.23 | 9198.39 | 7998.51 | 1998.58 | 2053.7 | 2791.3 | 2842.3 | 7204.62 | 7917.24 | 11923.78 | 2811.34 | 7803.72 | 6802.35 | 3022 | 101431.62 | E3 SUMO-protein ligase ZBED1 (EC 2.3.2.-) (DNA replication-related element-binding factor) (Putative Ac-like transposable element) (Zinc finger BED domain-containing protein 1) (dREF homolog) | ZBED1 ALTE DREF hDREF KIAA0785 TRAMP | negative regulation by host of viral genome replication [GO:0044828]; positive regulation of transcription by RNA polymerase II [GO:0045944]; protein autosumoylation [GO:1990466]; protein sumoylation [GO:0016925]; regulation of DNA-templated transcription [GO:0006355]; regulation of transcription by RNA polymerase II [GO:0006357] |

| | | | | | | | | | | | | | | | | | | | | | |
|---|---|---|---|---|---|---|---|---|---|---|---|---|---|---|---|---|---|---|---|---|---|
| 32.68 | 45306.35 | 68.78 | 32.22 | 13654.7 | 210.85 | 94.08 | 2971.23 | 38.41 | 105.26 | 3618.5 | 45.61 | 71.61 | 17474.45 | 182.7 | 62.1 | 16770.37 | 418.62 | 101158.52 | Lysine-specific demethylase 2A (EC 1.14.11.27) (CXXC-type zinc finger protein 8) (F-box and leucine-rich repeat protein 11) (F-box protein FBL7) (F-box protein Lilina) (F-box/LRR-repeat protein 11) (JmjC domain-containing histone demethylation protein 1A) ([Histone-H3]-lysine-36 demethylase 1A) | KDM2A CXXC8 FBL11 FBL7 FBXL11 JHDM1A KIAA1004 | chromatin remodeling [GO:0006338]; circadian regulation of gene expression [GO:0032922]; double-strand break repair via nonhomologous end joining [GO:0006303]; negative regulation of transcription by competitive promoter binding [GO:0010944]; regulation of circadian rhythm [GO:0042752]; regulation of transcription by RNA polymerase II [GO:0006357] |
| 20537.3 | 1478.47 | 5363.49 | 10565.53 | 1812.94 | 3354.71 | 311.71 | 664.72 | 904.89 | 19037.65 | 1684.07 | 631.16 | 29707.97 | 416.03 | 1066.72 | 1726.55 | 687.02 | 943.47 | 100894.4 | Serine/threonine-protein phosphatase 2A 55 kDa regulatory subunit B beta isoform (PP2A subunit B isoform B55-beta) (PP2A subunit B isoform PR55-beta) (PP2A subunit B isoform R2-beta) (PP2A subunit B isoform beta) | PPP2R2B | |

| | | | | | | | | | | | | | | | | | | | | | |
|---|---|---|---|---|---|---|---|---|---|---|---|---|---|---|---|---|---|---|---|---|---|
| 9311.15 | 7892.76 | 6912.07 | 3787.75 | 7099.19 | 8627.86 | 3351.29 | 3432.7 | 3602.47 | 3017.21 | 2636.88 | 4939.01 | 5654.33 | 10335.42 | 4389.66 | 3748.17 | 6526.81 | 5624.67 | 100889.4 | Ras-related protein Rab-6A (Rab-6) (EC 3.6.5.2) | RAB6A | intra-Golgi vesicle-mediated transport [GO:0006891]; intracellular protein transport [GO:0006886]; protein localization to Golgi apparatus [GO:0034067]; protein localization to Golgi membrane [GO:1903292]; retrograde transport, endosome to Golgi [GO:0042147]; retrograde vesicle-mediated transport, Golgi to endoplasmic reticulum [GO:0006890] |
| 4986.25 | 4343.43 | 5028.94 | 2008.51 | 4631.98 | 4862.9 | 19959.18 | 2151.29 | 3782.05 | 2096.88 | 2227.91 | 3635.19 | 5848.22 | 6656.46 | 3756.2 | 16325.31 | 3674.11 | 4059.44 | 100034.25 | Disks large-associated protein 1 (DAP-1) (Guanylate kinase-associated protein) | dlgap1 si:zc142h2.3 | modulation of chemical synaptic transmission [GO:0050804]; signaling [GO:0023052] |
| 4629.07 | 15241.06 | 7200.02 | 756.85 | 14863.78 | 7194.74 | 3226.21 | 10449.96 | 633.56 | 1096.16 | 3930.98 | 4437.43 | 2510.64 | 8472 | 2124.21 | 2415.25 | 7195.74 | 1752.49 | 98130.15 | Tax1-binding protein 3 (Glutaminase-interacting protein 3) (Tax interaction protein 1) (TIP-1) (Tax-interacting protein 1) | TAX1BP3 TIP1 | negative regulation of cell population proliferation [GO:0008285]; negative regulation of protein localization to cell surface [GO:2000009]; negative regulation of Wnt signaling pathway [GO:0030178]; regulation of Cdc42 protein signal transduction [GO:0032489]; Rho protein signal transduction [GO:0007266]; Wnt signaling pathway [GO:0016055] |

| | | | | | | | | | | | | | | | | | | | | | |
|---|---|---|---|---|---|---|---|---|---|---|---|---|---|---|---|---|---|---|---|---|---|
| 6906.53 | 5443.29 | 5064.27 | 5105.93 | 6415.27 | 8785.75 | 73.63 | 4050.52 | 4571.53 | 6190.36 | 6778.53 | 16426.41 | 3248.98 | 867.65 | 8514.62 | 551.74 | 1749.53 | 7057.33 | 97801.87 | Chymotrypsin-like elastase family member 2A (EC 3.4.21.71) (Elastase-2A) | CELA2A ELA2A | insulin catabolic process [GO:1901143]; proteolysis [GO:0006508]; regulation of insulin secretion [GO:0050796]; regulation of platelet aggregation [GO:0090330]; response to insulin [GO:0032868] |
| 18032.89 | 309.23 | 70.37 | 6207.36 | 1874.81 | 7054.6 | 72.34 | 285.18 | 3.42 | 10996.11 | 842.84 | 284.29 | 42180.62 | 185.2 | 163.89 | 3260.6 | 411.35 | 66.3 | 92301.4 | Transcription factor SOX-10 (cSOX10) | SOX10 | cardiac neuron development [GO:0060959]; central nervous system myelination [GO:0022010]; enteric nervous system development [GO:0048484]; morphogenesis of an epithelium [GO:0002009]; negative regulation of transcription by RNA polymerase II [GO:0000122]; neural crest cell migration [GO:0001755]; oligodendrocyte development [GO:0014003]; oligodendrocyte differentiation [GO:0048709]; peripheral nervous system development [GO:0007422]; positive regulation of DNA-templated transcription [GO:0045893] |

| | | | | | | | | | | | | | | | | | | | | | |
|---|---|---|---|---|---|---|---|---|---|---|---|---|---|---|---|---|---|---|---|---|---|
| 4892.82 | 7809.3 | 0 | 19934.82 | 12243.44 | 137.14 | 40.91 | 7146.81 | 0 | 4923.07 | 10160.45 | 0 | 2244.91 | 2352.76 | 1.03 | 5149.25 | 13310.83 | 126.25 | 90473.79 | Actin-binding LIM protein 3 (abLIM-3) (Actin-binding LIM protein family member 3) | ABLIM3 KIAA0843 HMFN1661 | cilium assembly [GO:0060271]; cytoskeleton organization [GO:0007010]; lamellipodium assembly [GO:0030032]; positive regulation of protein targeting to mitochondrion [GO:1903955]; positive regulation of transcription by RNA polymerase II [GO:0045944]; transcription by RNA polymerase II [GO:0006366] |
| 1269.82 | 16154.95 | 0 | 294.18 | 9139.03 | 232.98 | 0 | 12221.49 | 0 | 328.57 | 4682.61 | 31.84 | 945.21 | 25757.24 | 138.35 | 119.44 | 17024.69 | 134.82 | 88475.22 | Chaperone protein DnaK (HSP70) (Heat shock 70 kDa protein) (Heat shock protein 70) | dnaK VIBHAR_01134 | |
| 4266.3 | 6238.78 | 5870.5 | 1347.17 | 5240.41 | 11657.81 | 2227.76 | 2588.31 | 6954.09 | 2362.23 | 2809.94 | 7656.38 | 3477.2 | 6816.99 | 6100.17 | 1577.27 | 2827.09 | 6037.48 | 86055.88 | Opsin-5 (G-protein coupled receptor 136) (G-protein coupled receptor PGR12) (Neuropsin) (Transmembrane protein 13) | OPN5 GPR136 PGR12 TMEM13 | cellular response to light stimulus [GO:0071482]; cellular response to UV-A [GO:0071492]; entrainment of circadian clock by photoperiod [GO:0043153]; G protein-coupled receptor signaling pathway [GO:0007186]; hyaloid vascular plexus regression [GO:1990384]; phototransduction [GO:0007602]; phototransduction, UV [GO:0007604]; visual perception [GO:0007601] |

| | | | | | | | | | | | | | | | | | | | | | |
|---|---|---|---|---|---|---|---|---|---|---|---|---|---|---|---|---|---|---|---|---|---|
| 2853.94 | 11671.76 | 2217.81 | 408.98 | 19041.53 | 6730.28 | 32.87 | 3448.56 | 541.8 | 646.2 | 7200.62 | 1810.01 | 1614.2 | 20848.27 | 1366.77 | 222.61 | 4661.16 | 673.22 | 85990.59 | Protein phosphatase 1 regulatory subunit 37 (Leucine-rich repeat-containing protein 68) | PPP1R37 LRRC68 | |
| 6370.35 | 8303.54 | 4326.71 | 2547.16 | 9859.65 | 7759.91 | 1363.37 | 5205.68 | 439.38 | 3433.71 | 4313.84 | 3270.95 | 6878.77 | 8713.66 | 1938.73 | 2715.55 | 6546.24 | 1843.31 | 85830.51 | DNA replication licensing factor mcm7-B (EC 3.6.4.12) (CDC47 homolog B) (CDC47-2p) (Minichromosome maintenance protein 7-B) (xMCM7-B) | mcm7-b cdc47-2 | DNA replication [GO:0006260]; DNA replication initiation [GO:0006270]; DNA strand elongation involved in DNA replication [GO:0006271]; double-strand break repair via break-induced replication [GO:0000727]; regulation of DNA-templated DNA replication initiation [GO:0030174] |
| 6083.27 | 8510.62 | 5.43 | 14302.26 | 8745.7 | 73.02 | 56.94 | 7868.84 | 1.33 | 7355.98 | 11257.23 | 1.21 | 1839.81 | 2891.37 | 0 | 4352.71 | 11875.19 | 67.68 | 85288.59 | Evolutionarily conserved signaling intermediate in Toll pathway, mitochondrial | ecsit zgc:152999 | cell surface receptor protein serine/threonine kinase signaling pathway [GO:0007178]; innate immune response [GO:0045087]; regulation of oxidoreductase activity [GO:0051341]; regulation of protein complex stability [GO:0061635] |
| 2833.33 | 7229.91 | 6973.58 | 1958.22 | 5485.32 | 8696.91 | 82.78 | 4782.64 | 2857.03 | 1316.37 | 2409.54 | 8118.01 | 4486.93 | 10440.84 | 4918.39 | 1062.35 | 4731.36 | 6330.86 | 84714.37 | Pre-mRNA-splicing factor SPF27 (Protein BCAS2 homolog) | bcas2 zgc:101730 | mRNA splicing, via spliceosome [GO:0000398] |

| | | | | | | | | | | | | | | | | | | | | | |
|---|---|---|---|---|---|---|---|---|---|---|---|---|---|---|---|---|---|---|---|---|---|
| 1845.27 | 15348.25 | 2461.19 | 2007.69 | 11564.72 | 3195.6 | 2246.57 | 7978.1 | 726.22 | 866.62 | 3631.29 | 2842.56 | 3288.07 | 9035.35 | 1667.87 | 2237.93 | 11791.03 | 1331.27 | 84065.6 | CUB and sushi domain-containing protein 1 (CUB and sushi multiple domains protein 1) | CSMD1 KIAA1890 UNQ5952/PRO19863 | conditioned place preference [GO:1990708]; female gonad development [GO:0008585]; gene expression [GO:0010467]; glucose homeostasis [GO:0042593]; male gonad development [GO:0008584]; mammary gland branching involved in pregnancy [GO:0060745]; memory [GO:0007613]; oviduct epithelium development [GO:0035846]; startle response [GO:0001964] |
| 4088.92 | 4959.28 | 4190.17 | 3201.6 | 5290.11 | 8440.29 | 9341.55 | 2036.44 | 1973.77 | 2064.15 | 2767.44 | 5539.05 | 5576.66 | 7218.8 | 2415.06 | 7874.88 | 4577.67 | 2193.4 | 83749.24 | Gamma-crystallin M3 (Gamma-M3) | | lens development in camera-type eye [GO:0002088]; visual perception [GO:0007601] |
| 15147.6 | 173.24 | 93.95 | 3269.81 | 689.33 | 1512.66 | 61.24 | 38.9 | 3.43 | 10976 | 431.33 | 760.99 | 48475.8 | 119.66 | 137.79 | 994.79 | 171.76 | 11.63 | 83069.91 | Carbonyl reductase [NADPH] 1 (EC 1.1.1.184) (15-hydroxyprostaglandin dehydrogenase [NADP(+)]) (EC 1.1.1.196, EC 1.1.1.197) (20-beta-hydroxysteroid dehydrogenase) (Alcohol dehydrogenase [NAD(P)+] CBR1) (EC 1.1.1.71) (NADPH-dependent carbonyl reductase 1) (Prostaglandin 9-ketoreductase) (PG-9-KR) (Prostaglandin-E(2) 9-reductase) | CBR1 | lipid metabolic process [GO:0006629]; vitamin K metabolic process [GO:0042373]; xenobiotic metabolic process [GO:0006805] |

| | | | | | | | | | | | | | | | | | | | | | |
|---|---|---|---|---|---|---|---|---|---|---|---|---|---|---|---|---|---|---|---|---|---|
| 272.42 | 121.18 | 154.93 | 132.41 | 209.93 | 133.74 | 3.59 | 89.37 | 50.75 | 125.24 | 93.94 | 99.31 | 78075.32 | 1952.84 | 235.73 | 168.65 | 666.07 | 74.81 | 82660.23 | Signal transducing adapter molecule 2 (STAM-2) (Hrs-binding protein) | Stam2 Hbp | protein transport [GO:0015031] |
| 6324.42 | 8120.24 | 4.75 | 9114.35 | 5327.9 | 45.2 | 49.4 | 7262.63 | 0 | 8977.84 | 11536.63 | 3.03 | 3717.29 | 4797.11 | 1.89 | 4477.28 | 11861.52 | 119.18 | 81740.66 | Peroxisomal N(1)-acetyl-spermine/spermidine oxidase (EC 1.5.3.13) (Polyamine oxidase) | Paox Pao | polyamine catabolic process [GO:0006598]; positive regulation of spermidine biosynthetic process [GO:1901307]; putrescine biosynthetic process [GO:0009446]; putrescine catabolic process [GO:0009447]; spermidine catabolic process [GO:0046203]; spermine catabolic process [GO:0046208] |
| 7957.77 | 21324.2 | 4.78 | 9283.75 | 9495.32 | 54.93 | 48.82 | 1234.54 | 0 | 3412.19 | 7321.14 | 0 | 3733.69 | 6723.97 | 0 | 2747.47 | 8024.95 | 61.85 | 81429.37 | Stathmin-2 (Superior cervical ganglion-10 protein) (Protein SCG10) | Stmn2 Scg10 Scgn10 Stmb2 | cellular response to nerve growth factor stimulus [GO:1990090]; negative regulation of microtubule depolymerization [GO:0007026]; negative regulation of microtubule polymerization [GO:0031115]; negative regulation of neuron projection development [GO:0010977]; positive regulation of microtubule depolymerization [GO:0031117]; positive regulation of neuron projection development [GO:0010976] |

| | | | | | | | | | | | | | | | | | | | | | |
|---|---|---|---|---|---|---|---|---|---|---|---|---|---|---|---|---|---|---|---|---|---|
| 59.02 | 26857.79 | 71.79 | 42.63 | 11476.75 | 141.32 | 0 | 7163.48 | 63.55 | 49.96 | 2611.37 | 98.68 | 26.76 | 22195.16 | 105.03 | 116.39 | 9945.25 | 154.45 | 81179.38 | Mitogen-activated protein kinase kinase kinase 7-interacting protein 3 homolog | map3k7ip3 | positive regulation of canonical NF-kappaB signal transduction [GO:0043123] |
| 953.77 | 2613.68 | 5970.3 | 2245.1 | 2003.72 | 2246.79 | 38.97 | 2094.78 | 721.85 | 1142.22 | 1849.1 | 8583.33 | 1150.84 | 28142.04 | 2266.18 | 7403.59 | 9618.35 | 1397.84 | 80442.45 | NF-kappa-B essential modulator (NEMO) (IkB kinase-associated protein 1) (IKKAP1) (Inhibitor of nuclear factor kappa-B kinase subunit gamma) (I-kappa-B kinase subunit gamma) (IKK-gamma) (IKKG) (IkB kinase subunit gamma) (NF-kappa-B essential modifier) | IKBKG NEMO | DNA damage response [GO:0006974]; positive regulation of canonical NF-kappaB signal transduction [GO:0043123]; positive regulation of NF-kappaB transcription factor activity [GO:0051092]; positive regulation of T cell receptor signaling pathway [GO:0050862]; positive regulation of transcription by RNA polymerase II [GO:0045944] |
| 7392.65 | 5278.38 | 4309.79 | 4590.79 | 3942.37 | 8956.72 | 14.82 | 21384.27 | 5820.7 | 4177.29 | 2575.31 | 1500.43 | 1859.49 | 2774.39 | 1122.65 | 309.18 | 2775.4 | 1655.93 | 80440.56 | Gamma-aminobutyric acid receptor subunit beta-3 (GABA(A) receptor subunit beta-3) (GABAAR subunit beta-3) | GABRB3 | chloride transmembrane transport [GO:1902476]; gamma-aminobutyric acid signaling pathway [GO:0007214]; inhibitory synapse assembly [GO:1904862] |

| | | | | | | | | | | | | | | | | | | | | | |
|---|---|---|---|---|---|---|---|---|---|---|---|---|---|---|---|---|---|---|---|---|---|
| 380.73 | 496.54 | 260 | 157.76 | 346.29 | 429.9 | 37.37 | 364.6 | 33.63 | 129.98 | 629.62 | 259.72 | 671.97 | 50789.28 | 303.51 | 3339.23 | 21302.87 | 428.18 | 80361.18 | Probable JmjC domain-containing histone demethylation protein 2C (EC 1.14.11.-) (Jumonji domain-containing protein 1C) (Thyroid receptor-interacting protein 8) (TR-interacting protein 8) (TRIP-8) | JMJD1C JHDM2C KIAA1380 TRIP8 | blood coagulation [GO:0007596]; regulation of DNA-templated transcription [GO:0006355]; regulation of transcription by RNA polymerase II [GO:0006357] |
| 250.01 | 6.63 | 0.96 | 1073.48 | 6.64 | 26.78 | 41861.69 | 8.48 | 36.84 | 587.12 | 1591.07 | 28.91 | 20.94 | 9.1 | 12.61 | 33050.37 | 704.7 | 86.39 | 79362.72 | Sodium/potassium/calcium exchanger 3 (Na(+)/K(+)/Ca(2+)-exchange protein 3) (Solute carrier family 24 member 3) | Slc24a3 Nckx3 | bone mineralization [GO:0030282]; calcium ion transmembrane transport [GO:0070588]; intracellular calcium ion homeostasis [GO:0006874]; monoatomic cation transport [GO:0006812]; negative regulation of gene expression [GO:0010629]; positive regulation of gene expression [GO:0010628]; potassium ion transmembrane transport [GO:0071805]; sodium ion transmembrane transport [GO:0035725] |
| 12999.38 | 104.52 | 70.43 | 3846.37 | 645.74 | 2289.8 | 46.14 | 42.78 | 1.44 | 13520.42 | 392.36 | 600.84 | 42183.8 | 72.79 | 155.71 | 1098.63 | 161.34 | 6.48 | 78238.97 | Homeobox protein HMX1 (GH6) (Homeobox protein H6) | HMX1 H6 | negative regulation of DNA-templated transcription [GO:0045892]; regulation of transcription by RNA polymerase II [GO:0006357] |

| | | | | | | | | | | | | | | | | | | | | | |
|---|---|---|---|---|---|---|---|---|---|---|---|---|---|---|---|---|---|---|---|---|---|
| 3852.69 | 6577.51 | 5807.96 | 2090.26 | 7087.93 | 5427.64 | 6544.23 | 3745.31 | 2020.65 | 1435.34 | 3503.2 | 3927.22 | 4672.91 | 7060.3 | 2392.58 | 4865.39 | 4942.05 | 2219.64 | 78172.81 | Elongation factor 1-alpha (EF-1-alpha) (42Sp50) (Thesaurin A) | | translation [GO:0006412]; translational elongation [GO:0006414] |
| 22565.62 | 994.12 | 10777.31 | 2481.73 | 579.04 | 9587.16 | 255.65 | 286.82 | 516.22 | 4924.79 | 662.72 | 3735.16 | 9607.48 | 2067.59 | 3972.27 | 1959.07 | 1521.75 | 1098.28 | 77592.78 | Kelch-like protein 29 (Kelch repeat and BTB domain-containing protein 9) | KLHL29 KBTBD9 KIAA1921 | proteasome-mediated ubiquitin-dependent protein catabolic process [GO:0043161] |
| 3266.85 | 4570.72 | 5132.26 | 3065.45 | 5999.57 | 5879.78 | 6677.58 | 2544.06 | 1676.63 | 1856.1 | 3323.95 | 5963.51 | 4181.61 | 7162.05 | 2019.21 | 7419.12 | 4368.09 | 2053.46 | 77160 | Centromere-associated protein E (Centromere protein E) (CENP-E) (Kinesin-7) (Kinesin-related protein CENPE) | CENPE | attachment of mitotic spindle microtubules to kinetochore [GO:0051315]; cell division [GO:0051301]; chromosome segregation [GO:0007059]; kinetochore assembly [GO:0051382]; lateral attachment of mitotic spindle microtubules to kinetochore [GO:0099607]; metaphase chromosome alignment [GO:0051310]; microtubule plus-end directed mitotic chromosome migration [GO:0099606]; microtubule-based movement [GO:0007018]; mitotic cell cycle [GO:0000278]; mitotic chromosome movement towards spindle pole [GO:0007079]; mitotic metaphase chromosome alignment [GO:0007080]; mitotic spindle organization [GO:0007052]; positive regulation of protein kinase activity [GO:0045860]; regulation of mitotic metaphase/anaphase transition [GO:0030071] |

| | | | | | | | | | | | | | | | | | | | | | |
|---|---|---|---|---|---|---|---|---|---|---|---|---|---|---|---|---|---|---|---|---|---|
| 2803.63 | 4066.36 | 4543.09 | 3241.68 | 5087.45 | 6637.22 | 7356.73 | 1834.65 | 1666.16 | 2028.69 | 2761.29 | 5520.77 | 4934.06 | 7772.14 | 2350.14 | 6318.54 | 4538.3 | 1959.21 | 75420.11 | Voltage-gated purine nucleotide uniporter SLC17A9 (Solute carrier family 17 member 9) (Vesicular nucleotide transporter) (VNUT) | Slc17a9 | ADP transport [GO:0015866]; ATP export [GO:1904669]; ATP transport [GO:0015867]; guanine nucleotide transmembrane transport [GO:1903790]; lysosomal protein catabolic process [GO:1905146]; purine nucleotide import into lysosome [GO:0141013] |
| 4019.57 | 3423.64 | 5280.85 | 4931.13 | 3563.36 | 7081.98 | 5293.71 | 1777.75 | 1509.33 | 2736.85 | 2585.57 | 5610.65 | 5692.06 | 6713.06 | 2111.98 | 5709.75 | 4885.61 | 2204.5 | 75131.35 | Phosphatidylinositol transfer protein beta isoform (PI-TP-beta) (PtdIns transfer protein beta) (PtdInsTP beta) (Phosphatidylinositol-transfer protein 36 kDa isoform) (PI-TP 36 kda isoform) | PITPNB | retrograde vesicle-mediated transport, Golgi to endoplasmic reticulum [GO:0006890] |

| | | | | | | | | | | | | | | | | | | | | | |
|---|---|---|---|---|---|---|---|---|---|---|---|---|---|---|---|---|---|---|---|---|---|
| 3677.32 | 3856.42 | 6755.25 | 2007.59 | 5761.29 | 5323.75 | 6020.99 | 2279.91 | 3580.44 | 1520.27 | 2259.72 | 4270.8 | 3925.93 | 6715.16 | 3693.63 | 4854.74 | 3716.69 | 4506.97 | 74726.87 | Beta-hexosaminidase subunit beta (EC 3.2.1.52) (Beta-N-acetylhexosaminidase subunit beta) (Hexosaminidase subunit B) (N-acetyl-beta-glucosaminidase subunit beta) | HEXB | carbohydrate metabolic process [GO:0005975]; ganglioside catabolic process [GO:0006689]; glycosaminoglycan metabolic process [GO:0030203]; N-glycan processing [GO:0006491]; single fertilization [GO:0007338] |
| 0 | 0 | 0 | 0 | 0 | 74535.57 | 18.96 | 0 | 0 | 0 | 0 | 13.86 | 14.35 | 0 | 0 | 11.69 | 4.3 | 8.39 | 74607.12 | Glycine receptor subunit beta (Glycine receptor 58 kDa subunit) | GLRB | acrosome reaction [GO:0007340]; adult walking behavior [GO:0007628]; chemical synaptic transmission [GO:0007268]; chloride transmembrane transport [GO:1902476]; gamma-aminobutyric acid receptor clustering [GO:0097112]; monoatomic ion transport [GO:0006811]; nervous system development [GO:0007399]; neuropeptide signaling pathway [GO:0007218]; righting reflex [GO:0060013]; startle response [GO:0001964]; synaptic transmission, glycinergic [GO:0060012]; visual perception [GO:0007601] |

| | | | | | | | | | | | | | | | | | | | | | | |
|---|---|---|---|---|---|---|---|---|---|---|---|---|---|---|---|---|---|---|---|---|---|---|
| 2834.98 | | 1494.72 | 3941.11 | 1998.44 | 1548.26 | 10454.62 | 83.21 | 714.24 | 278.12 | 1625.34 | 1001.35 | 2079.76 | 17726.99 | 9556.25 | 11289.75 | 2315.05 | 4548.24 | 932.76 | 74423.19 | Prostaglandin E2 receptor EP1 subtype (PGE receptor EP1 subtype) (PGE2 receptor EP1 subtype) (Prostanoid EP1 receptor) | Ptger1 Ptgerep1 | adenylate cyclase-activating dopamine receptor signaling pathway [GO:0007191]; response to lipopolysaccharide [GO:0032496]; response to prostaglandin E [GO:0034695] |
| 2734.36 | | 5809.45 | 3464.35 | 3939.37 | 5545.08 | 8503.16 | 39.86 | 3043.95 | 2446.97 | 6502.6 | 5902.54 | 4131.57 | 3799.68 | 8308.52 | 3112.06 | 1015.99 | 2842.02 | 2688.23 | 73829.76 | Guanine nucleotide-binding protein G(i) subunit alpha-2 (Adenylate cyclase-inhibiting G alpha protein) | GNAI2 | adenylate cyclase-inhibiting G protein-coupled receptor signaling pathway [GO:0007193]; cell division [GO:0051301]; cell population proliferation [GO:0008283]; G protein-coupled acetylcholine receptor signaling pathway [GO:0007213]; G protein-coupled adenosine receptor signaling pathway [GO:0001973]; gamma-aminobutyric acid signaling pathway [GO:0007214]; modulation of chemical synaptic transmission [GO:0050804]; positive regulation of cell population proliferation [GO:0008284] |

| | | | | | | | | | | | | | | | | | | | | | |
|---|---|---|---|---|---|---|---|---|---|---|---|---|---|---|---|---|---|---|---|---|---|
| 2523.4 | 4705.31 | 3632.85 | 1370.25 | 8159.96 | 6650.74 | 1202.89 | 3695.14 | 717.91 | 1307.5 | 2673.08 | 3177.68 | 5465.69 | 9552.57 | 5021.77 | 1113.08 | 6030.28 | 5881.65 | 72881.75 | NLR family CARD domain-containing protein 3 (CARD15-like protein) (Caterpiller protein 16.2) (CLR16.2) (NACHT, LRR and CARD domains-containing protein 3) (Nucleotide-binding oligomerization domain protein 3) | NLRC3 NOD3 | canonical NF-kappaB signal transduction [GO:0007249]; negative regulation of canonical NF-kappaB signal transduction [GO:0043124]; negative regulation of cytokine production involved in inflammatory response [GO:1900016]; negative regulation of epithelial cell proliferation [GO:0050680]; negative regulation of fibroblast proliferation [GO:0048147]; negative regulation of inflammatory response [GO:0050728]; negative regulation of innate immune response [GO:0045824]; negative regulation of interferon-alpha production [GO:0032687]; negative regulation of interferon-beta production [GO:0032688]; negative regulation of interleukin-6 production [GO:0032715]; negative regulation of NF-kappaB transcription factor activity [GO:0032088]; negative regulation of NLRP3 inflammasome complex assembly [GO:1900226]; negative regulation of non-canonical NF-kappaB signal transduction [GO:1901223]; |
| 126.63 | 0 | 0 | 876.9 | 0 | 35.59 | 54268.62 | 1.98 | 0 | 431.81 | 795.54 | 16.58 | 36.27 | 0 | 13.67 | 14877.61 | 682.17 | 22.08 | 72185.45 | Serine/threonine-protein phosphatase 2A 55 kDa regulatory subunit B delta isoform (PP2A subunit B isoform B55-delta) (PP2A subunit B isoform PR55-delta) (PP2A subunit B isoform R2-delta) (PP2A subunit B isoform delta) | ppp2r2d zgc:76887 | cell division [GO:0051301]; exit from mitosis [GO:0010458]; mitotic cell cycle [GO:0000278]; myoblast fusion [GO:0007520]; regulation of chromosome segregation [GO:0051983] |

| | | | | | | | | | | | | | | | | | | | | | |
|---|---|---|---|---|---|---|---|---|---|---|---|---|---|---|---|---|---|---|---|---|---|
| 0 | 0 | 71924.65 | 0 | 0 | 0 | 0 | 0 | 1.28 | 0 | 0 | 0 | 0 | 0 | 0 | 0 | 0 | 2.87 | 71928.8 | Regulating synaptic membrane exocytosis protein 1 (Rab-3-interacting molecule 1) (RIM 1) (Rab-3-interacting protein 2) | RIMS1 KIAA0340 RAB3IP2 RIM1 Nbla00761 | acrosomal vesicle exocytosis [GO:0060478]; calcium-ion regulated exocytosis [GO:0017156]; cell differentiation [GO:0030154]; intracellular protein transport [GO:0006886]; membrane fusion [GO:0061025]; positive regulation of dendrite extension [GO:1903861]; positive regulation of excitatory postsynaptic potential [GO:2000463]; positive regulation of gene expression [GO:0010628]; positive regulation of inhibitory postsynaptic potential [GO:0097151]; protein-containing complex assembly [GO:0065003]; regulated exocytosis [GO:0045055]; regulation of neurotransmitter secretion [GO:0046928]; regulation of synaptic vesicle exocytosis [GO:2000300]; secretion [GO:0046903]; synaptic vesicle docking [GO:0016081]; synaptic vesicle exocytosis [GO:0016079]; synaptic vesicle priming [GO:0016082]; visual perception [GO:0007601] |
| 2.88 | 24198.55 | 34.65 | 4.08 | 7589.49 | 165.3 | 0.81 | 5361.8 | 9.77 | 7.88 | 4301.09 | 176.73 | 3.09 | 20825.75 | 19.14 | 2.99 | 8889.26 | 40.93 | 71634.19 | Glucoside xylosyltransferase 1 (EC 2.4.2.42) (Glycosyltransferase 8 domain-containing protein 3) | Gxylt1 Glt8d3 Gm87 | O-glycan processing [GO:0016266] |
| 2139.68 | 3317.83 | 7423.47 | 1715.84 | 3285.85 | 4810.33 | 7913.82 | 1775.14 | 2027.24 | 1012.34 | 1366.5 | 11972.96 | 2852.57 | 5353.27 | 2359.28 | 5717.76 | 3037.23 | 2141.55 | 70222.66 | Dehydrogenase/reductase SDR family member 11 (17-beta-hydroxysteroid dehydrogenase) (3-beta-hydroxysteroid 3-dehydrogenase) (EC 1.1.1.270) (Estradiol 17-beta-dehydrogenase) (EC 1.1.1.62) (Short-chain dehydrogenase/reductase family 24C member 1) | DHRS11 SDR24C1 | estrogen biosynthetic process [GO:0006703] |

| | | | | | | | | | | | | | | | | | | | | | |
|---|---|---|---|---|---|---|---|---|---|---|---|---|---|---|---|---|---|---|---|---|---|
| 208.35 | 0 | 0 | 1320.85 | 1.82 | 46.75 | 49937.05 | 0 | 0 | 184.02 | 258.92 | 17.15 | 63.49 | 0 | 3.39 | 17453.35 | 407.66 | 11.97 | 69914.77 | Actin-binding protein IPP (Intracisternal A particle-promoted polypeptide) (IPP) (Kelch-like protein 27) | IPP KLHL27 | proteasome-mediated ubiquitin-dependent protein catabolic process [GO:0043161] |
| 188.28 | 0 | 0 | 503.66 | 0 | 10.77 | 6695.57 | 0 | 0 | 155.06 | 480.91 | 0 | 0 | 0 | 0 | 59402.27 | 1081.71 | 0 | 68518.23 | Retinol dehydrogenase 13 (EC 1.1.1.300) (Short chain dehydrogenase/reductase family 7C member 3) | RDH13 SDR7C3 PSEC0082 UNQ736/PRO1430 | eye photoreceptor cell development [GO:0042462]; response to high light intensity [GO:0009644]; retina layer formation [GO:0010842]; retinal metabolic process [GO:0042574] |
| 0 | 0 | 9.93 | 0 | 0 | 68374.09 | 0 | 0 | 0 | 7.68 | 0 | 0 | 13.43 | 0 | 22.5 | 20.61 | 0 | 44.82 | 68493.06 | Rho guanine nucleotide exchange factor TIAM1 (T-lymphoma invasion and metastasis-inducing protein 1) (TIAM-1) | Tiam1 Tiam-1 | activation of GTPase activity [GO:0090630]; brain-derived neurotrophic factor receptor signaling pathway [GO:0031547]; cell migration [GO:0016477]; cell-matrix adhesion [GO:0007160]; ephrin receptor signaling pathway [GO:0048013]; positive regulation of axonogenesis [GO:0050772]; positive regulation of neuron projection development [GO:0010976]; Rac protein signal transduction [GO:0016601]; regulation of dopaminergic neuron differentiation [GO:1904338]; regulation of GTPase activity [GO:0043087]; regulation of non-canonical Wnt signaling pathway [GO:2000050]; small GTPase-mediated signal transduction [GO:0007264] |

| | | | | | | | | | | | | | | | | | | | | | |
|---|---|---|---|---|---|---|---|---|---|---|---|---|---|---|---|---|---|---|---|---|---|
| 1938.54 | 1524.7 | 3964.68 | 1170.56 | 1713.6 | 2356.58 | 15853.6 | 543.84 | 3957.78 | 712.04 | 965.64 | 6363.93 | 2851.42 | 3112.89 | 3385.25 | 10638.8 | 1895.95 | 4530.11 | 67479.91 | Poly(rC)-binding protein 2 (Alpha-CP2) (CTBP) (CBP) (Putative heterogeneous nuclear ribonucleoprotein X) (hnRNP X) | Pcbp2 Cbp Hnrnpx Hnrpx | defense response to virus [GO:0051607]; innate immune response [GO:0045087]; negative regulation of defense response to virus [GO:0050687]; positive regulation of transcription by RNA polymerase II [GO:0045944]; proteasome-mediated ubiquitin-dependent protein catabolic process [GO:0043161]; protein maturation [GO:0051604] |
| 0.78 | 22924.69 | 0.56 | 6.23 | 7871.54 | 40.49 | 0.61 | 7936.12 | 1.79 | 1.04 | 4262.98 | 3.4 | 2.41 | 13048.16 | 0 | 3.29 | 10204.75 | 60.32 | 66369.16 | Peroxidasin (EC 1.11.2.-) [Cleaved into: PXDN active fragment] | pxdn pxn | angiogenesis [GO:0001525]; basement membrane assembly [GO:0070831]; basement membrane organization [GO:0071711]; cell adhesion [GO:0007155]; hydrogen peroxide catabolic process [GO:0042744]; protein homooligomerization [GO:0051260]; protein homotrimerization [GO:0070207]; response to oxidative stress [GO:0006979] |
| 79.89 | 10.89 | 49.67 | 391.36 | 19.39 | 46.04 | 33754.48 | 4.89 | 12.3 | 455.17 | 846.85 | 23.19 | 55.12 | 25.53 | 50.3 | 29211.51 | 759.8 | 62.04 | 65858.42 | cGMP-specific 3',5'-cyclic phosphodiesterase (EC 3.1.4.35) (cGMP-binding cGMP-specific phosphodiesterase) (CGB-PDE) | PDE5A PDE5 | cAMP-mediated signaling [GO:0019933]; cGMP catabolic process [GO:0046069] |

| | | | | | | | | | | | | | | | | | | | | | |
|---|---|---|---|---|---|---|---|---|---|---|---|---|---|---|---|---|---|---|---|---|---|
| 2154.35 | 3991.76 | 7902.9 | 1874 | 5872.76 | 6769.14 | 1595.18 | 2093.25 | 4278.36 | 3669.91 | 3284.15 | 1316.57 | 3568.75 | 4916.09 | 3919.3 | 2392.27 | 3626.57 | 2587.07 | 65812.38 | Retrovirus-related Pol polyprotein from transposon 17.6 [Includes: Protease (EC 3.4.23.-);Reverse transcriptase (EC 2.7.7.49);Endonuclease] | pol | DNA integration [GO:0015074]; proteolysis [GO:0006508] |
| 1077.16 | 11950.28 | 1897.85 | 1067.1 | 11052.05 | 1477.86 | 374.9 | 3411.62 | 1076.57 | 412.5 | 3003.1 | 1721.97 | 951.51 | 16021.34 | 1373.94 | 618.3 | 6603.82 | 1694.37 | 65786.24 | Serine/threonine-protein kinase 26 (EC 2.7.11.1) (MST3 and SOK1-related kinase) (Mammalian STE20-like protein kinase 4) (MST-4) (STE20-like kinase MST4) (Serine/threonine-protein kinase MASK) | STK26 MASK MST4 | apoptotic process [GO:0006915]; cellular response to oxidative stress [GO:0034599]; cellular response to starvation [GO:0009267]; intracellular signal transduction [GO:0035556]; microvillus assembly [GO:0030033]; negative regulation of cell migration [GO:0030336]; protein autophosphorylation [GO:0046777]; protein phosphorylation [GO:0006468]; regulation of apoptotic process [GO:0042981] |
| 3125.63 | 1866.83 | 6909.87 | 1657.41 | 1519.44 | 9441.62 | 18.94 | 2373.55 | 7279.69 | 3242.63 | 1200.82 | 12135.83 | 1936.58 | 1375.43 | 4936.96 | 156.86 | 991.18 | 5098.95 | 65268.22 | Lanosterol synthase (EC 5.4.99.7) (2,3-epoxysqualene--lanosterol cyclase) (Oxidosqualene--lanosterol cyclase) (OSC) | Lss Osc | cholesterol biosynthetic process [GO:0006695]; regulation of protein stability [GO:0031647]; steroid biosynthetic process [GO:0006694]; sterol metabolic process [GO:0016125]; triterpenoid biosynthetic process [GO:0016104] |

| | | | | | | | | | | | | | | | | | | | | | |
|---|---|---|---|---|---|---|---|---|---|---|---|---|---|---|---|---|---|---|---|---|---|
| 2648.87 | 3920.67 | 5213.47 | 2382.25 | 4095.14 | 5517.28 | 2918.18 | 1862.47 | 2151.03 | 1477.12 | 2040.83 | 5066.63 | 5098.41 | 7879.89 | 2876.73 | 3611.1 | 3796.14 | 2202.71 | 64758.92 | Transcription factor IIIB 50 kDa subunit (B-related factor 2) (BRF-2) | brf2 zgc:100856 | cellular response to oxidative stress [GO:0034599]; regulation of transcription by RNA polymerase III [GO:0006359]; transcription preinitiation complex assembly [GO:0070897] |
| 5520.66 | 1530.39 | 3785.16 | 1975.6 | 1104.63 | 4613.12 | 411.68 | 322.67 | 6622.32 | 1347.08 | 549.98 | 3147.92 | 10363.23 | 3719.16 | 7446.34 | 1583.46 | 3097.35 | 7451.84 | 64592.59 | Probable E3 ubiquitin-protein ligase HERC3 (EC 2.3.2.26) (HECT domain and RCC1-like domain-containing protein 3) (HECT-type E3 ubiquitin transferase HERC3) | HERC3 KIAA0032 | protein ubiquitination [GO:0016567]; ubiquitin-dependent protein catabolic process [GO:0006511] |
| 2743.9 | 4925.89 | 3873.15 | 1806.84 | 4317.54 | 5874.08 | 136.32 | 6325.96 | 4994.27 | 2880.27 | 3453.41 | 1961.95 | 2270.31 | 9764.57 | 2927.56 | 1023.27 | 2868.44 | 2217.96 | 64365.69 | Protein phosphatase 1 regulatory subunit 12A (Myosin phosphatase-targeting subunit 1) (Myosin phosphatase target subunit 1) (Protein phosphatase myosin-binding subunit) | ppp1r12a mbs mypt1 si:dkey-28j4.1 zgc:110448 | actin filament bundle assembly [GO:0051017]; actin filament bundle distribution [GO:0070650]; axonogenesis [GO:0007409]; convergent extension involved in axis elongation [GO:0060028]; exocrine pancreas development [GO:0031017]; liver development [GO:0001889]; midbrain-hindbrain boundary morphogenesis [GO:0021555]; motor neuron axon guidance [GO:0008045]; neuron projection morphogenesis [GO:0048812]; regulation of BMP signaling pathway [GO:0030510]; regulation of cell adhesion [GO:0030155]; regulation of cell shape [GO:0008360]; signal transduction [GO:0007165] |

| | | | | | | | | | | | | | | | | | | | | | | |
|---|---|---|---|---|---|---|---|---|---|---|---|---|---|---|---|---|---|---|---|---|---|---|
| 3418.07 | | 3199.99 | 3676.95 | 1323.67 | 2909.71 | 3650.35 | 4392 | 1255.54 | 7703.2 | 999.28 | 1029.52 | 4006.94 | 4478.3 | 3465.52 | 7377.12 | 2939.73 | 1907.77 | 6525.3 | 64258.96 | Axin-1 (Axis inhibition protein 1) (Protein Fused) | Axin1 Axin Fu | apoptotic process [GO:0006915]; axial mesoderm development [GO:0048318]; axial mesoderm formation [GO:0048320]; beta-catenin destruction complex assembly [GO:1904885]; canonical Wnt signaling pathway [GO:0060070]; cytoplasmic microtubule organization [GO:0031122]; dorsal/ventral axis specification [GO:0009950]; dorsal/ventral pattern formation [GO:0009953]; epigenetic programming in the zygotic pronuclei [GO:0044725]; head development [GO:0060322]; in utero embryonic development [GO:0001701]; negative regulation of canonical Wnt signaling pathway [GO:0090090]; negative regulation of fat cell differentiation [GO:0045599]; negative regulation of gene expression [GO:0010629]; negative regulation of protein metabolic process [GO:0051248]; negative regulation of transcription elongation by RNA polymerase II [GO:0034244]; negative regulation of Wnt signaling pathway |
| 1228.85 | | 3162.27 | 4356.19 | 1495.14 | 3643.94 | 3073.75 | 131.95 | 1435.64 | 4847.82 | 1126.6 | 1519.68 | 7420.59 | 1467.96 | 8462.5 | 8912.07 | 1793.07 | 3904.64 | 6225.35 | 64208.01 | Properdin (Complement factor P) | Cfp Pfc | complement activation [GO:0006956]; complement activation, alternative pathway [GO:0006957]; positive regulation of immune response [GO:0050778]; positive regulation of opsonization [GO:1903028] |
| 3194.51 | | 4226.07 | 6273.25 | 1445.97 | 2518.58 | 3228.35 | 4545.47 | 1797.01 | 1965.61 | 2066.78 | 2708.98 | 6307.6 | 5815.06 | 9054.97 | 2668.96 | 1989.05 | 2809.36 | 1459.92 | 64075.5 | Mesoderm induction early response protein 3 (Mi-er3) | Mier3 | |

| | | | | | | | | | | | | | | | | | | | | | |
|---|---|---|---|---|---|---|---|---|---|---|---|---|---|---|---|---|---|---|---|---|---|
| 2230.85 | 2949.14 | 2926.1 | 1963.84 | 3522.18 | 4632.98 | 9710.72 | 1254.94 | 1328.36 | 1332.3 | 1918.27 | 3485.48 | 4122.16 | 8301.82 | 2148.35 | 6449.62 | 4238.28 | 1463.96 | 63979.35 | Cadherin EGF LAG seven-pass G-type receptor 3 | Celsr3 | axonal fasciculation [GO:0007413]; cilium assembly [GO:0060271]; dopaminergic neuron axon guidance [GO:0036514]; homophilic cell adhesion via plasma membrane adhesion molecules [GO:0007156]; motor neuron migration [GO:0097475]; regulation of protein localization [GO:0032880]; serotonergic neuron axon guidance [GO:0036515]; Wnt signaling pathway, planar cell polarity pathway [GO:0060071] |
| 3954.89 | 6305.99 | 1673.61 | 1341.41 | 6225.43 | 2791.94 | 240.86 | 2921.13 | 3073.75 | 852.68 | 2170.93 | 1608.36 | 1911.4 | 13716.71 | 1353.36 | 1088.45 | 10165.42 | 2539.17 | 63935.49 | Prolyl 3-hydroxylase OGFOD1 (EC 1.14.11.-) (2-oxoglutarate and iron-dependent oxygenase domain-containing protein 1) (uS12 prolyl 3-hydroxylase) | OGFOD1 | cell population proliferation [GO:0008283]; protein hydroxylation [GO:0018126]; regulation of translational termination [GO:0006449]; stress granule assembly [GO:0034063] |

| | | | | | | | | | | | | | | | | | | | | | |
|---|---|---|---|---|---|---|---|---|---|---|---|---|---|---|---|---|---|---|---|---|---|
| 14541.9 | 117.89 | 75.29 | 2640.72 | 421.52 | 645.94 | 30.39 | 36.68 | 1.58 | 14688.32 | 302.22 | 489.72 | 28719.04 | 47.7 | 69.04 | 966.36 | 114.49 | 5.31 | 63914.11 | Rab11 family-interacting protein 3 (FIP3) (FIP3-Rab11) (Rab11-FIP3) (Arfophilin-1) (EF hands-containing Rab-interacting protein) (Eferin) (MU-MB-17.148) | RAB11FIP3 ARFO1 KIAA0665 | cell division [GO:0051301]; early endosome to recycling endosome transport [GO:0061502]; endocytic recycling [GO:0032456]; Golgi to plasma membrane protein transport [GO:0043001]; negative regulation of adiponectin secretion [GO:0070164]; positive regulation of cilium assembly [GO:0045724]; positive regulation of mitotic cytokinetic process [GO:1903438]; protein localization to cilium [GO:0061512]; protein localization to cleavage furrow [GO:1905345]; regulation of cilium assembly [GO:1902017]; regulation of cytokinesis [GO:0032465]; regulation of early endosome to recycling endosome transport [GO:1902954]; regulation of endocytic recycling [GO:2001135]; regulation of protein localization to centrosome [GO:1904779]; regulation of vesicle-mediated transport [GO:0060627]; vesicle-mediated transport [GO:0016192] |
| 6.55 | 0 | 62785.31 | 0 | 3.34 | 0 | 0 | 0 | 0 | 0 | 0 | 0 | 0 | 0 | 0 | 0 | 0 | 0 | 62795.2 | Trafficking regulator of GLUT4 1 (Dispanin subfamily B member 1) (DSPB1) (Tumor suppressor candidate 5 homolog) | trarg1 tusc5 | cellular response to insulin stimulus [GO:0032869]; endosome to plasma membrane protein transport [GO:0099638]; glucose import in response to insulin stimulus [GO:0044381]; protein localization to plasma membrane [GO:0072659] |
| 3415.58 | 3251.9 | 2981.53 | 2596.28 | 3704.17 | 5846.35 | 5785.24 | 1418.88 | 1796.84 | 1676.79 | 1759.11 | 3231.85 | 4855.69 | 7768.39 | 1966.06 | 5625.85 | 3416.23 | 1665.11 | 62761.85 | NF-kappa-B inhibitor-interacting Ras-like protein 1 (I-kappa-B-interacting Ras-like protein 1) (Kappa B-Ras protein 1) (KappaB-Ras1) | nkiras1 si:dkey-126a1.3 zgc:92823 | negative regulation of canonical NF-kappaB signal transduction [GO:0043124]; Ral protein signal transduction [GO:0032484] |

| | | | | | | | | | | | | | | | | | | | | | |
|---|---|---|---|---|---|---|---|---|---|---|---|---|---|---|---|---|---|---|---|---|---|
| 5038.22 | 5457.24 | 5249.88 | 917.67 | 4199.96 | 3540.74 | 124.65 | 4397.81 | 749.2 | 1598.25 | 3563.36 | 2818.86 | 6837.62 | 9725.51 | 2647.05 | 523.84 | 3468.23 | 1570.77 | 62428.86 | Enhancer of polycomb homolog 1 | Epc1 | double-strand break repair via homologous recombination [GO:0000724]; negative regulation of gene expression, epigenetic [GO:0045814]; negative regulation of transcription by RNA polymerase II [GO:0000122]; positive regulation of DNA-templated transcription [GO:0045893]; positive regulation of double-strand break repair via homologous recombination [GO:1905168]; positive regulation of striated muscle cell differentiation [GO:0051155]; positive regulation of transcription by RNA polymerase II [GO:0045944]; regulation of apoptotic process [GO:0042981]; regulation of cell cycle [GO:0051726]; regulation of chromatin organization [GO:1902275]; regulation of double-strand break repair [GO:2000779]; sperm DNA condensation [GO:0035092]; vascular associated smooth muscle cell differentiation [GO:0035886] |
| 13392.29 | 192.14 | 150.36 | 5510.57 | 2046.72 | 4520.62 | 56.78 | 171.49 | 7.66 | 9312.85 | 1126.37 | 410.51 | 24193.77 | 128.82 | 127.64 | 685.53 | 256.53 | 17.25 | 62307.9 | Vasopressin V2 receptor (V2R) (AVPR V2) (Antidiuretic hormone receptor) (Renal-type arginine vasopressin receptor) | AVPR2 | cellular response to hormone stimulus [GO:0032870]; G protein-coupled receptor signaling pathway [GO:0007186]; positive regulation of gene expression [GO:0010628]; positive regulation of vasoconstriction [GO:0045907]; regulation of systemic arterial blood pressure by vasopressin [GO:0001992] |

| | | | | | | | | | | | | | | | | | | | | | |
|---|---|---|---|---|---|---|---|---|---|---|---|---|---|---|---|---|---|---|---|---|---|
| 9075.41 | 934.86 | 3620.09 | 3191.23 | 3558.09 | 12790.13 | 16.72 | 652.4 | 123.76 | 3584.13 | 1169.73 | 2043.65 | 9557.84 | 1797.56 | 4094.66 | 1645.94 | 3538.84 | 793 | 62188.04 | N-acyl-phosphatidylethanolamine-hydrolyzing phospholipase D (N-acyl phosphatidylethanolamine phospholipase D) (NAPE-PLD) (NAPE-hydrolyzing phospholipase D) (EC 3.1.4.54) | Napepld | host-mediated regulation of intestinal microbiota composition [GO:0048874]; N-acylethanolamine metabolic process [GO:0070291]; N-acylphosphatidylethanolamine metabolic process [GO:0070292]; negative regulation of eating behavior [GO:1903999]; phospholipid catabolic process [GO:0009395]; phospholipid metabolic process [GO:0006644]; positive regulation of brown fat cell differentiation [GO:0090336]; positive regulation of inflammatory response [GO:0050729]; response to isolation stress [GO:0035900]; temperature homeostasis [GO:0001659] |
| 1867.69 | 563.1 | 1643.01 | 646.98 | 452.53 | 2052.55 | 9.66 | 98.78 | 13370.39 | 400.68 | 223.14 | 6100.01 | 2552.35 | 862.26 | 12564.38 | 442.65 | 654.72 | 17193.71 | 61698.59 | Retinal homeobox protein Rx3 | rx3 | adenohypophysis development [GO:0021984]; camera-type eye morphogenesis [GO:0048593]; cell fate specification [GO:0001708]; embryonic camera-type eye morphogenesis [GO:0048596]; eye development [GO:0001654]; eye field cell fate commitment involved in camera-type eye formation [GO:0060898]; forebrain development [GO:0030900]; hypothalamus development [GO:0021854]; neural crest cell migration [GO:0001755]; positive regulation of transcription by RNA polymerase II [GO:0045944]; regulation of transcription by RNA polymerase II [GO:0006357]; retina development in camera-type eye [GO:0060041] |
| 31965.08 | 454.18 | 299.73 | 9207.66 | 1875.56 | 1574.94 | 68.55 | 651.9 | 205.03 | 4666.14 | 862.16 | 477.74 | 5809.85 | 602.41 | 179.48 | 618.82 | 1450.45 | 135.31 | 61104.99 | Transposable element Tcb1 transposase (Transposable element Barney transposase) | | DNA integration [GO:0015074]; DNA transposition [GO:0006313] |

| | | | | | | | | | | | | | | | | | | | | | |
|---|---|---|---|---|---|---|---|---|---|---|---|---|---|---|---|---|---|---|---|---|---|
| 5900.41 | 7596.72 | 3142.38 | 1264.52 | 4374.77 | 7877.94 | 80.46 | 2093.33 | 872.72 | 1164.62 | 1565.71 | 5080.22 | 5871.21 | 5159.03 | 2842.56 | 513.12 | 3543.47 | 2053.66 | 60996.85 | Mitochondrial inner membrane protein OXA1L (Oxidase assembly 1-like protein) (OXA1-like protein) | OXA1L | mitochondrial protein quality control [GO:0141164]; mitochondrial respiratory chain complex I assembly [GO:0032981]; protein insertion into mitochondrial inner membrane from matrix [GO:0032979]; protein tetramerization [GO:0051262] |
| 1422.69 | 466.68 | 3286.05 | 1166.71 | 1137.7 | 3666.31 | 30682.54 | 533.06 | 653.9 | 1108.05 | 988.64 | 3109.59 | 4774.8 | 1550.93 | 2614.17 | 2869.8 | 336.9 | 242.55 | 60611.07 | Protein FAM13B (GAP-like protein N61) | FAM13B C5orf5 FAM13B1 | regulation of small GTPase mediated signal transduction [GO:0051056]; signal transduction [GO:0007165] |
| 1696.96 | 1653.11 | 1549.35 | 1661.66 | 2204.51 | 5331.15 | 92 | 28489.81 | 4239.05 | 2200.3 | 1230.81 | 1504.68 | 990.91 | 2336.37 | 1702.79 | 195.88 | 1413.68 | 2051.59 | 60544.61 | Semaphorin-6A (Semaphorin VIA) (Sema VIA) (Semaphorin-6A-1) (SEMA6A-1) | SEMA6A KIAA1368 SEMAQ | animal organ morphogenesis [GO:0009887]; apoptotic process [GO:0006915]; axon guidance [GO:0007411]; cell surface receptor signaling pathway [GO:0007166]; cellular response to vascular endothelial growth factor stimulus [GO:0035924]; cytoskeleton organization [GO:0007010]; negative regulation of angiogenesis [GO:0016525]; negative regulation of cell adhesion involved in sprouting angiogenesis [GO:0106089]; negative regulation of ERK1 and ERK2 cascade [GO:0070373]; negative regulation of sprouting angiogenesis [GO:1903671]; negative regulation of vascular endothelial growth factor signaling pathway [GO:1900747]; nervous system development [GO:0007399]; neural crest cell migration [GO:0001755]; positive regulation of neuron migration [GO:2001224]; semaphorin-plexin signaling pathway [GO:0071526] |

| | | | | | | | | | | | | | | | | | | | | | |
|---|---|---|---|---|---|---|---|---|---|---|---|---|---|---|---|---|---|---|---|---|---|
| 3008.53 | 4713.48 | 5.67 | 18310.24 | 9936.47 | 63.79 | 65.22 | 5970.57 | 0 | 1591.95 | 2659.04 | 0 | 1683.51 | 2126.27 | 0 | 2697.83 | 7529.08 | 85.03 | 60446.68 | Pro-opiomelanocortin B (POMC-B) (Corticotropin-lipotropin B) [Cleaved into: NPP 2;Corticotropin (Adrenocorticotropic hormone) (ACTH);Melanocyte-stimulating hormone alpha 2 (Alpha-MSH 2) (Melanotropin alpha 2);Corticotropin-like intermediary peptide 2 (CLIP-2);Lipotropin beta (Beta-LPH);Lipotropin gamma (Gamma-LPH);Melanocyte-stimulating hormone beta 2 | pomcb | neuropeptide signaling pathway [GO:0007218] |
| 2661.47 | 2262.49 | 1955.15 | 1573.28 | 1832.79 | 3295.3 | 12516.65 | 985.11 | 1855.47 | 1321 | 1672.09 | 3170.2 | 3890.44 | 6835.56 | 2046.48 | 8508.02 | 2603.23 | 1368.81 | 60353.54 | Golgi reassembly-stacking protein 2 (GRS2) (Golgi phosphoprotein 6) (GOLPH6) (Golgi reassembly-stacking protein of 55 kDa) (GRASP55) (p59) | GORASP2 GOLPH6 | cell differentiation [GO:0030154]; establishment of protein localization to plasma membrane [GO:0061951]; Golgi organization [GO:0007030]; organelle assembly [GO:0070925]; organelle organization [GO:0006996]; response to endoplasmic reticulum stress [GO:0034976]; response to unfolded protein [GO:0006986]; spermatogenesis [GO:0007283] |

| | | | | | | | | | | | | | | | | | | | | | |
|---|---|---|---|---|---|---|---|---|---|---|---|---|---|---|---|---|---|---|---|---|---|
| 2985.52 | 4125.3 | 2938.37 | 1432.39 | 4644.24 | 6380.98 | 510.4 | 2608.58 | 650.59 | 1529.94 | 1638.24 | 2797.24 | 6742.19 | 8445.29 | 4014.61 | 1243.42 | 4163.56 | 3434.23 | 60285.09 | Death domain-associated protein 6 (Daxx) | DAXX | androgen receptor signaling pathway [GO:0030521]; cellular response to cadmium ion [GO:0071276]; cellular response to copper ion [GO:0071280]; cellular response to diamide [GO:0072738]; cellular response to heat [GO:0034605]; cellular response to sodium arsenite [GO:1903936]; cellular response to unfolded protein [GO:0034620]; extrinsic apoptotic signaling pathway via death domain receptors [GO:0008625]; JNK cascade [GO:0007254]; negative regulation of DNA-templated transcription [GO:0045892]; negative regulation of gene expression [GO:0010629]; neuron intrinsic apoptotic signaling pathway in response to oxidative stress [GO:0036480]; nucleosome assembly [GO:0006334]; regulation of apoptotic process [GO:0042981]; regulation of protein ubiquitination [GO:0031396] |
| 0 | 1.22 | 4563.96 | 0 | 0 | 423.55 | 0 | 0 | 14995.26 | 0 | 7.24 | 14310.01 | 3.37 | 0 | 7428.94 | 1.49 | 12.03 | 18272.6 | 60019.67 | Integrin beta-7 (Gut homing receptor beta subunit) | ITGB7 | cell adhesion [GO:0007155]; cell adhesion mediated by integrin [GO:0033627]; cell-cell adhesion [GO:0098609]; cell-matrix adhesion [GO:0007160]; cell-matrix adhesion involved in ameboidal cell migration [GO:0003366]; heterotypic cell-cell adhesion [GO:0034113]; immune response in gut-associated lymphoid tissue [GO:0002387]; integrin-mediated signaling pathway [GO:0007229]; leukocyte migration [GO:0050900]; leukocyte tethering or rolling [GO:0050901]; receptor clustering [GO:0043113]; substrate adhesion-dependent cell spreading [GO:0034446]; T cell migration [GO:0072678] |
| 2649.68 | 3245.62 | 4363 | 2011.49 | 3780.85 | 6481.72 | 3165.04 | 1514.92 | 2088.41 | 1688.65 | 1749.14 | 4874.88 | 4591.94 | 5897.69 | 2317.17 | 3536.67 | 3542.1 | 2321.79 | 59820.76 | Peroxisome proliferator-activated receptor gamma coactivator-related protein 1 (PGC-1-related coactivator) (PRC) | PPRC1 KIAA0595 | energy homeostasis [GO:0097009]; positive regulation of transcription by RNA polymerase II [GO:0045944] |

| | | | | | | | | | | | | | | | | | | | | | |
|---|---|---|---|---|---|---|---|---|---|---|---|---|---|---|---|---|---|---|---|---|---|
| 215.95 | 0 | 0 | 996.07 | 0 | 39.68 | 28681.18 | 7.27 | 0 | 281.89 | 933.21 | 4.72 | 31.28 | 0 | 0 | 27790.46 | 651.98 | 11.45 | 59645.14 | Myelin transcription factor 1-like protein (MyT1-L) (MyT1L) (Neural zinc finger factor 1) (NZF-1) (Postmitotic neural gene 1 protein) (Zinc finger protein Png-1) | Myt1l Kiaa1106 Nzf1 Png1 | negative regulation of transcription by RNA polymerase II [GO:0000122]; nervous system development [GO:0007399]; neuron development [GO:0048666]; neuron differentiation [GO:0030182]; neuron fate commitment [GO:0048663]; neuron fate specification [GO:0048665]; regulation of transcription by RNA polymerase II [GO:0006357] |
| 4592.57 | 5146.47 | 0 | 8800.46 | 4993.18 | 34.08 | 37.1 | 2901.84 | 0 | 5674.12 | 7805.09 | 0 | 4828.74 | 2020.16 | 0 | 5257.89 | 7216.02 | 111.18 | 59418.9 | LINE-1 retrotransposable element ORF2 protein (ORF2p) [Includes: Reverse transcriptase (EC 2.7.7.49);Endonuclease (EC 3.1.21.-)] | | DNA recombination [GO:0006310]; nucleic acid metabolic process [GO:0090304]; retrotransposition [GO:0032197] |
| 2120.86 | 2903.3 | 3241.72 | 2927.5 | 7386.68 | 2607.53 | 15.11 | 2292.67 | 4356.52 | 983.87 | 1961.29 | 2367.48 | 2445.26 | 8772.51 | 3390.84 | 797.33 | 5647.36 | 4943.01 | 59160.84 | AP-4 complex accessory subunit RUSC1 (New molecule containing SH3 at the carboxy-terminus) (Nesca) (RUN and SH3 domain-containing protein 1) | RUSC1 NESCA | protein polyubiquitination [GO:0000209] |
| 4833.38 | 7063.27 | 9045.57 | 2218.87 | 2994.28 | 3763.52 | 4351.37 | 1228.6 | 853.74 | 20.79 | 29.61 | 0 | 7625.14 | 10955.05 | 3182.88 | 813.56 | 28 | 0 | 59007.63 | Translationally-controlled tumor protein homolog (TCTP) | tpt1 tct1 tctp | |

| | | | | | | | | | | | | | | | | | | | | | |
|---|---|---|---|---|---|---|---|---|---|---|---|---|---|---|---|---|---|---|---|---|---|
| 51.83 | 69.48 | 17391.29 | 51.8 | 92.11 | 2094.47 | 21.19 | 52.04 | 166.65 | 35.18 | 36.69 | 37127.51 | 80.07 | 175.47 | 369.96 | 49.68 | 65.02 | 116.89 | 58047.33 | Dedicator of cytokinesis protein 8 | Dock8 | cellular response to chemokine [GO:1990869]; dendritic cell migration [GO:0036336]; immunological synapse formation [GO:0001771]; memory T cell proliferation [GO:0061485]; negative regulation of T cell apoptotic process [GO:0070233]; positive regulation of establishment of T cell polarity [GO:1903905]; positive regulation of T cell migration [GO:2000406]; small GTPase-mediated signal transduction [GO:0007264] |
| 2473.33 | 1678.92 | 6693.87 | 1238.74 | 2060.32 | 3177.05 | 2629.35 | 1314.16 | 5591.48 | 963.19 | 756.67 | 3214.76 | 2275.95 | 4715.59 | 6538.08 | 2207.22 | 3227.67 | 6441.88 | 57198.23 | Condensin-2 complex subunit H2 (Non-SMC condensin II complex subunit H2) | ncaph2 si:dkey-202b22.2 | meiotic chromosome condensation [GO:0010032]; mitotic sister chromatid separation [GO:0051306] |
| 4981.7 | 9171.96 | 0 | 19293.27 | 10581.94 | 68.05 | 44.04 | 5456.27 | 9.96 | 45.18 | 0 | 0 | 463.83 | 506.43 | 0 | 1488.98 | 4616.78 | 57.65 | 56786.04 | SLAM family member 9 | Slamf9 | defense response to bacterium [GO:0042742]; plasmacytoid dendritic cell chemotaxis [GO:0002410]; plasmacytoid dendritic cell differentiation [GO:0002273] |
| 116.84 | 4068.87 | 614.26 | 195.76 | 4307.29 | 1832.25 | 5715.66 | 1746.17 | 3166.04 | 90.17 | 2220.37 | 2282.12 | 94.43 | 10135.99 | 2540.94 | 6352.45 | 8736.79 | 2563.38 | 56779.78 | Solute carrier family 25 member 48 | slc25a48 zgc:92090 | |

| | | | | | | | | | | | | | | | | | | | | | |
|---|---|---|---|---|---|---|---|---|---|---|---|---|---|---|---|---|---|---|---|---|---|
| 3439.74 | 2789.03 | 3928.81 | 1446.88 | 2760.68 | 3281.12 | 7414.88 | 1341.78 | 1921.6 | 755.89 | 1191.75 | 5106.22 | 1794.96 | 4750.52 | 1871.99 | 6149.1 | 3923.87 | 2740.63 | 56609.45 | Serine/threonine-protein phosphatase 6 catalytic subunit (PP6C) (EC 3.1.3.16) (Protein phosphatase V) (PP-V) | Ppp6c Ppv | G1/S transition of mitotic cell cycle [GO:0000082]; innate immune response [GO:0045087]; negative regulation of cGAS/STING signaling pathway [GO:0160049] |
| 1426.63 | 1095.51 | 4667.77 | 715.66 | 1258.84 | 2004.47 | 12650.91 | 377.86 | 2450.57 | 329.33 | 604.88 | 7295.66 | 2253.35 | 2464.42 | 3111.96 | 7096.7 | 1814.27 | 4256.23 | 55875.02 | THAP domain-containing protein 5 | THAP5 | negative regulation of cell cycle [GO:0045786] |
| 3649.21 | 4611.2 | 0 | 12258.25 | 6942.32 | 77.02 | 31.63 | 3020.7 | 0 | 4269.3 | 4784.61 | 0 | 2377.43 | 3950.02 | 0 | 3107.23 | 6690.81 | 88.68 | 55858.41 | DNA repair protein XRCC3 (X-ray repair cross-complementing protein 3) | Xrcc3 | double-strand break repair via homologous recombination [GO:0000724]; interstrand cross-link repair [GO:0036297]; positive regulation of mitotic cell cycle spindle assembly checkpoint [GO:0090267]; regulation of centrosome duplication [GO:0010824]; resolution of mitotic recombination intermediates [GO:0071140]; t-circle formation [GO:0090656]; telomere maintenance via recombination [GO:0000722] |
| 2343.32 | 2510.85 | 2869.63 | 1842.51 | 3652.5 | 4917.05 | 5674.7 | 989.06 | 1819.81 | 1315.28 | 1707.15 | 3314.39 | 3874 | 7336.64 | 1755.87 | 5496.14 | 3131.83 | 1282.17 | 55832.9 | Thiosulfate:glutathione sulfurtransferase (TST) (EC 2.8.1.3) | TSTD1 KAT | sulfide oxidation, using sulfide:quinone oxidoreductase [GO:0070221] |

| | | | | | | | | | | | | | | | | | | | | | |
|---|---|---|---|---|---|---|---|---|---|---|---|---|---|---|---|---|---|---|---|---|---|
| 12.17 | 3.23 | 879.56 | 4.09 | 15.78 | 165.34 | 1.27 | 5.09 | 13077.72 | 0 | 23.96 | 4574.75 | 41.13 | 3.2 | 13463.1 | 59.12 | 68.89 | 23361.4 | 55759.8 | LINE-1 retrotransposable element ORF2 protein (ORF2p) (Long interspersed element-1) (L1) (Retrovirus-related Pol polyprotein LINE-1) [Includes: Reverse transcriptase (EC 2.7.7.49);Endonuclease (EC 3.1.21.-)] | Pol Gm17492 | DNA recombination [GO:0006310] |
| 711.99 | 2703.27 | 4192.81 | 752.51 | 5793.72 | 1794.35 | 2652.36 | 3466.47 | 4835.28 | 372.13 | 2148.3 | 4094.49 | 1262.84 | 6468.74 | 3648.13 | 1664.65 | 4346.92 | 4760.55 | 55669.51 | Protein YIF1B (YIP1-interacting factor homolog B) | yif1b zgc:103562 | cilium assembly [GO:0060271]; endoplasmic reticulum to Golgi vesicle-mediated transport [GO:0006888]; protein transport [GO:0015031]; sperm flagellum assembly [GO:0120316] |
| 135.67 | 0 | 0 | 756.68 | 0 | 17.94 | 32700.05 | 0.88 | 2.88 | 233.2 | 629.36 | 3.39 | 21.71 | 0 | 0 | 20095.23 | 714.18 | 21.29 | 55332.46 | Junctional adhesion molecule A (JAM-A) (Junctional adhesion molecule 1) (JAM-1) (CD antigen CD321) | F11R JAM1 | cell adhesion [GO:0007155]; establishment of endothelial intestinal barrier [GO:0090557]; intestinal absorption [GO:0050892]; regulation of membrane permeability [GO:0090559] |

| | | | | | | | | | | | | | | | | | | | | | |
|---|---|---|---|---|---|---|---|---|---|---|---|---|---|---|---|---|---|---|---|---|---|
| 5.63 | 4.75 | 20233.55 | 5.95 | 0.21 | 1922.01 | 70.23 | 0.09 | 212.17 | 3.59 | 1.02 | 32134.71 | 9.32 | 8.87 | 217.3 | 69.93 | 3.49 | 51.97 | 54954.79 | Titin (EC 2.7.11.1) (Connectin) (Rhabdomyosarcoma antigen MU-RMS-40.14) | TTN | cardiac muscle cell development [GO:0055013]; cardiac muscle contraction [GO:0060048]; cardiac muscle hypertrophy [GO:0003300]; cardiac muscle tissue morphogenesis [GO:0055008]; cardiac myofibril assembly [GO:0055003]; detection of muscle stretch [GO:0035995]; mitotic chromosome condensation [GO:0007076]; muscle contraction [GO:0006936]; muscle filament sliding [GO:0030049]; positive regulation of gene expression [GO:0010628]; positive regulation of protein secretion [GO:0050714]; protein kinase A signaling [GO:0010737]; response to calcium ion [GO:0051592]; sarcomere organization [GO:0045214]; sarcomerogenesis [GO:0048769]; skeletal muscle contraction [GO:0003009]; skeletal muscle myosin thick filament assembly [GO:0030241]; skeletal muscle thin filament assembly [GO:0030240]; striated muscle contraction [GO:0006941] |
| 0 | 0 | 5.81 | 0 | 0 | 54886 | 0 | 0 | 0 | 0 | 0 | 0 | 0 | 0 | 6.71 | 5.31 | 3.38 | 13.24 | 54920.45 | Collagen alpha-1(XIV) chain (Undulin) | COL14A1 | cell adhesion [GO:0007155] |
| 2368.05 | 2316.68 | 2875.23 | 1039 | 2393.25 | 2726.76 | 7654.45 | 1193.76 | 4275.77 | 723.41 | 1197.74 | 2271.23 | 2287.74 | 5751.51 | 2671.95 | 5927.16 | 3576.46 | 3301.27 | 54551.42 | Arf-GAP with dual PH domain-containing protein 2 (Centaurin-alpha-2) (Cnt-a2) | Adap2 Centa2 | heart development [GO:0007507] |
| 87.13 | 0 | 4.63 | 337.13 | 0 | 5.63 | 38597.51 | 0 | 0 | 180.3 | 570.99 | 0 | 29.67 | 0 | 0 | 14208.85 | 371.59 | 2.71 | 54396.14 | Large ribosomal subunit protein eL6 (60S ribosomal protein L6) (TAX-responsive enhancer element-binding protein 107) (TAXREB107) | Rpl6 | cytoplasmic translation [GO:0002181]; translation at postsynapse [GO:0140242]; translation at presynapse [GO:0140236] |

| | | | | | | | | | | | | | | | | | | | | | |
|---|---|---|---|---|---|---|---|---|---|---|---|---|---|---|---|---|---|---|---|---|---|
| 834.29 | 9229.19 | 447.35 | 189.61 | 10088.04 | 1394.99 | 18.78 | 3956.4 | 8.47 | 306.99 | 4158.76 | 576.42 | 889.49 | 16682.12 | 401.74 | 210.47 | 4883.27 | 75.78 | 54352.16 | Rho GTPase-activating protein SYDE2 (Synapse defective protein 1 homolog 2) (Protein syd-1 homolog 2) | SYDE2 | activation of GTPase activity [GO:0090630]; cell migration [GO:0016477]; regulation of Ras protein signal transduction [GO:0046578]; regulation of small GTPase mediated signal transduction [GO:0051056]; signal transduction [GO:0007165] |
| 1405.74 | 6107.07 | 3665.35 | 607.45 | 4390.43 | 4022.17 | 7494.21 | 1490.2 | 250.49 | 761.98 | 2074.38 | 2165.39 | 2640.26 | 5677.86 | 2480.52 | 4496.44 | 3185.42 | 956.26 | 53871.62 | AP-4 complex accessory subunit RUSC2 (Interacting protein of Rab1) (Iporin) (RUN and SH3 domain-containing protein 2) | RUSC2 KIAA0375 | |
| 1131.48 | 924.26 | 1003.08 | 966.58 | 708.26 | 1522.26 | 25741.73 | 406.72 | 1926.6 | 550.81 | 550.23 | 998.05 | 1450.33 | 1416.36 | 1477.76 | 10460.7 | 1104.27 | 1440.41 | 53779.89 | Small ribosomal subunit protein uS5m (28S ribosomal protein S5, mitochondrial) (MRP-S5) (S5mt) | Mrps5 | mitochondrial translation [GO:0032543] |
| 335.32 | 9724.19 | 682.22 | 52.97 | 14871.63 | 2583.33 | 0.17 | 5094.05 | 6.26 | 166.29 | 2896.69 | 1036.89 | 720.03 | 10464.26 | 404.76 | 56.29 | 4385.37 | 53.09 | 53533.81 | HEAT repeat-containing protein 6 | heatr6 zgc:172359 | |
| 3551.64 | 4133.67 | 1334.39 | 928.21 | 4696.28 | 1528.61 | 484.19 | 1410.01 | 1285.71 | 665.75 | 1126.26 | 982.32 | 4808.01 | 18050.36 | 1346.63 | 2485.61 | 3218.59 | 1082.06 | 53118.3 | Glutaminyl-peptide cyclotransferase (EC 2.3.2.5) (Glutaminyl cyclase) (QC) (Glutaminyl-tRNA cyclotransferase) | QPCT | peptidyl-pyroglutamic acid biosynthetic process, using glutaminyl-peptide cyclotransferase [GO:0017186] |

| | | | | | | | | | | | | | | | | | | | | | |
|---|---|---|---|---|---|---|---|---|---|---|---|---|---|---|---|---|---|---|---|---|---|
| 1872.48 | 3818.87 | 3880.15 | 904.43 | 9025.46 | 4442.69 | 122.46 | 3105.55 | 316.47 | 756.55 | 2143.91 | 1712.57 | 3814.88 | 7812.79 | 2122.94 | 665.85 | 5021.01 | 1052.25 | 52591.31 | Ankyrin repeat domain-containing protein 33B | Ankrd33b | |
| 9562.49 | 128.59 | 88.84 | 2837.95 | 890.23 | 2100.73 | 39.86 | 110.4 | 1.08 | 6002.96 | 436.96 | 394.64 | 28496.5 | 95.86 | 140.69 | 844.62 | 182.44 | 17.69 | 52372.53 | Histone-lysine N-methyltransferase SMYD3 (EC 2.1.1.354) (SET and MYND domain-containing protein 3) (Zinc finger MYND domain-containing protein 1) | Smyd3 Zmynd1 | cellular response to dexamethasone stimulus [GO:0071549]; establishment of protein localization [GO:0045184]; methylation [GO:0032259]; myotube cell development [GO:0014904]; nucleosome assembly [GO:0006334]; positive regulation of transcription by RNA polymerase II [GO:0045944] |
| 1543.17 | 315.96 | 1021.96 | 796.39 | 342.33 | 3982.16 | 14404.25 | 308.52 | 391.75 | 866 | 900.1 | 743.96 | 6733.92 | 2112.52 | 2080.02 | 13784.33 | 884.72 | 819.46 | 52031.52 | Protein unc-80 homolog (mUNC-80) | Unc80 Kiaa1843 | |
| 29.78 | 0 | 3299.68 | 0 | 0 | 35624.84 | 11.75 | 0 | 0 | 6.71 | 3.36 | 64.48 | 6.81 | 3.5 | 12097.88 | 9.7 | 0 | 0 | 51158.49 | CD209 antigen-like protein A (Dendritic cell-specific ICAM-3-grabbing non-integrin) (DC-SIGN) (CD antigen CD209) | Cd209a Cire | endocytosis [GO:0006897]; regulation of T cell proliferation [GO:0042129] |

| | | | | | | | | | | | | | | | | | | | | | |
|---|---|---|---|---|---|---|---|---|---|---|---|---|---|---|---|---|---|---|---|---|---|
| 4367.66 | 2542.63 | 1161.74 | 532.95 | 1579.74 | 2455.41 | 27.46 | 1085.59 | 242.25 | 0.7 | 0 | 0 | 11252.26 | 15907.05 | 5904.38 | 905.62 | 2530.25 | 596.02 | 51091.71 | Ceramide-1-phosphate transfer protein (CPTP) (Glycolipid transfer protein domain-containing protein 1) (GLTP domain-containing protein 1) | CPTP GLTPD1 | ceramide 1-phosphate transport [GO:1902389]; ceramide transport [GO:0035627]; intermembrane lipid transfer [GO:0120009]; negative regulation of autophagy [GO:0010507]; negative regulation of interleukin-1 beta production [GO:0032691]; negative regulation of NLRP3 inflammasome complex assembly [GO:1900226] |
| 4.27 | 0 | 6.93 | 0 | 0 | 50277.55 | 7.47 | 0 | 3.54 | 0 | 0 | 9.86 | 4.87 | 0 | 8.14 | 3.79 | 4.06 | 16.02 | 50346.5 | Lumican (Keratan sulfate proteoglycan lumican) (KSPG lumican) | LUM LDC | |
| 1513.77 | 4278.34 | 1149.37 | 3834.22 | 5405.26 | 1053.29 | 5124.8 | 5243.95 | 439.36 | 2013.16 | 4194.34 | 1063.89 | 1680.74 | 3606.6 | 615.57 | 3092.73 | 4931.09 | 818.83 | 50059.31 | CCN family member 1 (Cellular communication network factor 1) (Protein CEF-10) (Protein CYR61) | CCN1 CYR61 | apoptotic process involved in heart morphogenesis [GO:0003278]; atrial septum morphogenesis [GO:0060413]; atrioventricular valve morphogenesis [GO:0003181]; cell adhesion [GO:0007155]; cell-cell adhesion [GO:0098609]; chondroblast differentiation [GO:0060591]; extracellular matrix organization [GO:0030198]; integrin-mediated signaling pathway [GO:0007229]; intussusceptive angiogenesis [GO:0002041]; negative regulation of apoptotic process [GO:0043066]; osteoblast differentiation [GO:0001649]; positive regulation of apoptotic process [GO:0043065]; positive regulation of BMP signaling pathway [GO:0030513]; positive regulation of bone mineralization [GO:0030501]; positive regulation of cartilage development [GO:0061036]; positive regulation of cell differentiation [GO:0045597]; positive regulation of cell migration [GO:0030335]; positive regulation of cell-substrate adhesion [illegible] |
| 15431.23 | 824.19 | 1175.33 | 4799.67 | 1068.08 | 3721.22 | 41.79 | 717.6 | 328.86 | 3381.39 | 1033.33 | 1448.52 | 10666.81 | 1469.56 | 718.03 | 562.2 | 1850.15 | 348.64 | 49586.6 | Protein MON2 homolog (Protein SF21) | MON2 KIAA1040 SF21 | Golgi to endosome transport [GO:0006895]; protein transport [GO:0015031] |

| | | | | | | | | | | | | | | | | | | | | | |
|---|---|---|---|---|---|---|---|---|---|---|---|---|---|---|---|---|---|---|---|---|---|
| 72.98 | 145.45 | 9423.4 | 27 | 553.18 | 3923.89 | 156.94 | 112.44 | 107.43 | 41.18 | 260.26 | 31807.77 | 21.47 | 771.66 | 482.59 | 252.62 | 444.28 | 113.13 | 48717.67 | Hemicentin-1 (Fibulin-6) (FIBL-6) | HMCN1 FIBL6 | actin cytoskeleton organization [GO:0030036]; basement membrane organization [GO:0071711]; cell division [GO:0051301]; heterophilic cell-cell adhesion via plasma membrane cell adhesion molecules [GO:0007157]; homophilic cell adhesion via plasma membrane adhesion molecules [GO:0007156]; response to bacterium [GO:0009617]; visual perception [GO:0007601] |
| 2300.56 | 2645 | 1903.21 | 1947.13 | 2988.15 | 3527.53 | 5317.49 | 985.63 | 1229.66 | 1369.42 | 1654.34 | 3199.43 | 3394.22 | 5709.05 | 1368.94 | 4126.41 | 3108.99 | 1368.56 | 48143.72 | Echinoderm microtubule-associated protein-like 6 (EMAP-6) (Echinoderm microtubule-associated protein-like 5-like) | EML6 EML5L | |
| 18.02 | 9.59 | 962.01 | 9.69 | 14.23 | 109.56 | 0 | 0 | 16668.25 | 0 | 0 | 3647.06 | 114.35 | 43.99 | 11423.98 | 3.26 | 6.87 | 15070.94 | 48101.8 | Poly(rC)-binding protein 4 (Alpha-CP4) | PCBP4 | negative regulation of mRNA splicing, via spliceosome [GO:0048025]; regulation of transcription by RNA polymerase II [GO:0006357] |

| | | | | | | | | | | | | | | | | | | | | | |
|---|---|---|---|---|---|---|---|---|---|---|---|---|---|---|---|---|---|---|---|---|---|
| 658.42 | 289.3 | 239.53 | 709.91 | 564.81 | 586.8 | 20572.93 | 338.37 | 374.78 | 432.52 | 591.03 | 227.41 | 2095.12 | 776.35 | 494.9 | 17546.86 | 921.12 | 494.31 | 47914.47 | SAP domain-containing ribonucleoprotein (Nuclear protein Hcc-1) | SARNP HCC1 | mRNA export from nucleus [GO:0006406]; poly(A)+ mRNA export from nucleus [GO:0016973]; regulation of translation [GO:0006417] |
| 1490.97 | 2912.93 | 2907.95 | 1035.08 | 3478.34 | 3061.62 | 5890.94 | 1156.63 | 2075.37 | 751.8 | 1345.37 | 3312.8 | 2077.45 | 4139.46 | 2179.07 | 3552.13 | 2201.45 | 3024.1 | 46593.46 | Phosphatase and actin regulator 2 | Phactr2 | actin cytoskeleton organization [GO:0030036] |
| 3717.54 | 2875.16 | 5346.97 | 1211.81 | 3246.28 | 3350.9 | 1345.08 | 1503.81 | 2675.75 | 932.58 | 802.31 | 2784.84 | 2436.82 | 3719.81 | 3284.2 | 1362.32 | 2305.76 | 3390.52 | 46292.46 | Protein phosphatase 1 regulatory subunit 12A (MBSP) (Myosin phosphatase-targeting subunit 1) (Myosin phosphatase target subunit 1) (Protein phosphatase myosin-binding subunit) (Protein phosphatase subunit 1M) (PP-1M) (Serine/threonine protein phosphatase PP1 smooth muscle regulatory subunit M110) | Ppp1r12a Mbs Mypt1 | cellular response to xenobiotic stimulus [GO:0071466]; centrosome cycle [GO:0007098]; mitotic cell cycle [GO:0000278]; negative regulation of catalytic activity [GO:0043086]; neuron projection morphogenesis [GO:0048812]; positive regulation of transcription by RNA polymerase II [GO:0045944]; protein dephosphorylation [GO:0006470]; regulation of cell adhesion [GO:0030155]; regulation of nucleocytoplasmic transport [GO:0046822]; signal transduction [GO:0007165] |

| | | | | | | | | | | | | | | | | | | | | | |
|---|---|---|---|---|---|---|---|---|---|---|---|---|---|---|---|---|---|---|---|---|---|
| 1277.07 | 1129.93 | 4625.09 | 863.06 | 2214.83 | 4136.39 | 56.07 | 564.27 | 2849.2 | 814.41 | 1055.77 | 2118.6 | 1235.09 | 10016.85 | 2321.86 | 2282.26 | 5575.54 | 3008.16 | 46144.45 | Golgi resident protein GCP60 (Acyl-CoA-binding domain-containing protein 3) (Golgi complex-associated protein 1) (GOCAP1) (Golgi phosphoprotein 1) (GOLPH1) (PBR- and PKA-associated protein 7) (Peripheral benzodiazepine receptor-associated protein PAP7) [Cleaved into: Golgi resident protein GCP60, N-terminally processed] | ACBD3 GCP60 GOCAP1 GOLPH1 | steroid biosynthetic process [GO:0006694] |
| 212.29 | 8518.72 | 2419.1 | 458.94 | 3956.69 | 2595 | 43.14 | 2541.15 | 3455.38 | 353.25 | 1060.48 | 4102.52 | 88.83 | 7066.52 | 2994.64 | 113.05 | 1730.48 | 4217.74 | 45927.92 | Probable ATP-dependent RNA helicase DDX59 (EC 3.6.4.13) (DEAD box protein 59) (Zinc finger HIT domain-containing protein 5) | Ddx59 Znhit5 | |
| 714.49 | 7813.2 | 4880.15 | 213.12 | 5483.24 | 6301.05 | 60.61 | 2879.85 | 230.31 | 220.6 | 1453.76 | 563.4 | 687.62 | 6712.39 | 2051.38 | 178.14 | 4369.84 | 837.35 | 45650.5 | Rho GTPase-activating protein 17 (Rho-type GTPase-activating protein 17) (RhoGAP interacting with CIP4 homologs protein 1) (RICH-1) | ARHGAP17 RICH1 MSTP066 MSTP110 | negative regulation of small GTPase mediated signal transduction [GO:0051058]; regulation of actin cytoskeleton organization [GO:0032956]; regulation of Rac protein signal transduction [GO:0035020]; signal transduction [GO:0007165] |

| | | | | | | | | | | | | | | | | | | | | | |
|---|---|---|---|---|---|---|---|---|---|---|---|---|---|---|---|---|---|---|---|---|---|
| 9868.91 | 123.19 | 61.33 | 2697.08 | 422.73 | 1413.78 | 33.39 | 71.17 | 1.71 | 10852.18 | 244.9 | 489.44 | 18183.79 | 51.77 | 69.34 | 916.96 | 100.93 | 16.41 | 45619.01 | Syntaxin-binding protein 4 (Syntaxin 4-interacting protein) (STX4-interacting protein) (Synip) | STXBP4 | cellular response to type II interferon [GO:0071346]; DNA damage response [GO:0006974]; insulin receptor signaling pathway [GO:0008286]; positive regulation of cell cycle G1/S phase transition [GO:1902808]; positive regulation of keratinocyte proliferation [GO:0010838]; protein stabilization [GO:0050821]; protein targeting [GO:0006605]; regulation of D-glucose transmembrane transport [GO:0010827]; regulation of insulin secretion involved in cellular response to glucose stimulus [GO:0061178] |
| 6368.83 | 54.16 | 122.64 | 1877.43 | 106.58 | 512.1 | 0 | 0 | 5.24 | 11992.24 | 181.02 | 564.81 | 22971 | 0 | 78.99 | 632.25 | 109.45 | 16.95 | 45593.69 | DNA fragmentation factor subunit beta (EC 3.-.-.-) (Caspase-activated deoxyribonuclease) (CAD) (Caspase-activated DNase) (DNA fragmentation factor 40 kDa subunit) (DFF-40) | Dffb Cad | apoptotic chromosome condensation [GO:0030263]; apoptotic DNA fragmentation [GO:0006309]; negative regulation of apoptotic DNA fragmentation [GO:1902511] |

| | | | | | | | | | | | | | | | | | | | | | |
|---|---|---|---|---|---|---|---|---|---|---|---|---|---|---|---|---|---|---|---|---|---|
| 313.75 | 16517.95 | 106.25 | 739.23 | 8229.16 | 211.79 | 52.11 | 5302.59 | 55.44 | 209.17 | 1791.64 | 169.54 | 130.05 | 5325.81 | 51.75 | 106.06 | 5870.7 | 109 | 45291.99 | Phosphatidylinositol 3,4,5-trisphosphate 3-phosphatase and dual-specificity protein phosphatase PTEN (EC 3.1.3.16) (EC 3.1.3.48) (EC 3.1.3.67) (Inositol polyphosphate 3-phosphatase) (EC 3.1.3.-) (Phosphatase and tensin homolog) | Pten rCG_47874 | adult behavior [GO:0030534]; angiogenesis [GO:0001525]; apoptotic process [GO:0006915]; B cell proliferation [GO:0042100]; brain morphogenesis [GO:0048854]; cardiac muscle tissue development [GO:0048738]; cell migration [GO:0016477]; cell motility [GO:0048870]; cell population proliferation [GO:0008283]; cellular response to decreased oxygen levels [GO:0036294]; cellular response to electrical stimulus [GO:0071257]; cellular response to ethanol [GO:0071361]; cellular response to hypoxia [GO:0071456]; cellular response to insulin stimulus [GO:0032869]; cellular response to insulin-like growth factor stimulus [GO:1990314]; cellular response to leptin stimulus [GO:0044320]; cellular response to nerve growth factor stimulus [GO:1990090]; cellular response to sorbitol [GO:0072709]; central nervous system development [GO:0007417]; central nervous system |
| 9697.19 | 105.81 | 64.44 | 2360.68 | 638.35 | 1174.66 | 0 | 60.46 | 0.91 | 8492.83 | 461.67 | 372.3 | 21081.54 | 56.58 | 54.42 | 458.66 | 111.32 | 7.13 | 45198.95 | BCL-6 corepressor-like protein 1 (BCoR-L1) (BCoR-like protein 1) | BCORL1 | chromatin organization [GO:0006325]; negative regulation of transcription by RNA polymerase II [GO:0000122] |
| 5149.4 | 2148.85 | 6319.24 | 1342.31 | 2341.38 | 4333.61 | 191.5 | 856.09 | 3103.51 | 3675.08 | 1195.13 | 4489.2 | 2823.44 | 1569.01 | 1534.33 | 459.55 | 580.53 | 2348.26 | 44460.42 | L-amino-acid oxidase (EC 1.4.3.2) (Antiparasitic protein) (APP) (Serum L-amino-acid oxidase) (SR-LAAO) | | amino acid catabolic process [GO:0009063]; defense response to bacterium [GO:0042742]; innate immune response [GO:0045087] |

| | | | | | | | | | | | | | | | | | | | | | |
|---|---|---|---|---|---|---|---|---|---|---|---|---|---|---|---|---|---|---|---|---|---|
| 110.97 | 0 | 9.83 | 100.47 | 0 | 0 | 21047.45 | 0 | 0 | 349.7 | 836.15 | 5.99 | 16.75 | 0 | 1.35 | 21261.03 | 576.68 | 14.3 | 44330.67 | 72 kDa type IV collagenase (EC 3.4.24.24) (72 kDa gelatinase) (Gelatinase A) (Matrix metalloproteinase-2) (MMP-2) (TBE-1) [Cleaved into: PEX] | MMP2 CLG4A | angiogenesis [GO:0001525]; blood vessel maturation [GO:0001955]; bone trabecula formation [GO:0060346]; cell migration [GO:0016477]; cellular response to amino acid stimulus [GO:0071230]; cellular response to estradiol stimulus [GO:0071392]; cellular response to fluid shear stress [GO:0071498]; cellular response to interleukin-1 [GO:0071347]; cellular response to reactive oxygen species [GO:0034614]; cellular response to UV-A [GO:0071492]; collagen catabolic process [GO:0030574]; endodermal cell differentiation [GO:0035987]; ephrin receptor signaling pathway [GO:0048013]; extracellular matrix disassembly [GO:0022617]; extracellular matrix organization [GO:0030198]; face morphogenesis [GO:0060325]; heart development [GO:0007507]; intramembranous ossification [GO:0001957]; luteinization [GO:0001553]; macrophage chemotaxis [GO:0048246]; negative regulation of cell adhesion [GO:0007162]; negative [illegible] |
| 4670.87 | 6479.71 | 4.77 | 6267.26 | 3870 | 43.94 | 22.72 | 2718.75 | 0 | 3785.2 | 4785.14 | 0 | 1835.12 | 2549.17 | 2.86 | 2214.74 | 5036.84 | 39.44 | 44326.53 | Anoctamin-3 (Transmembrane protein 16C) | Ano3 Tmem16c | calcium activated galactosylceramide scrambling [GO:0061591]; calcium activated phosphatidylcholine scrambling [GO:0061590]; detection of mechanical stimulus [GO:0050982]; detection of temperature stimulus [GO:0016048]; establishment of localization in cell [GO:0051649] |
| 718.6 | 1174.21 | 4189.44 | 460.44 | 1454.41 | 1922.98 | 2346.79 | 583.12 | 6385.92 | 354.23 | 467.88 | 6122.9 | 851.23 | 1526.63 | 5759.56 | 1351.83 | 712.06 | 7895.52 | 44277.75 | DBF4-type zinc finger-containing protein 2 | ZDBF2 KIAA1571 | genomic imprinting [GO:0071514] |

| | | | | | | | | | | | | | | | | | | | | | |
|---|---|---|---|---|---|---|---|---|---|---|---|---|---|---|---|---|---|---|---|---|---|
| 2431.11 | 2651.01 | 2576.53 | 588.26 | 6409.98 | 1819.83 | 788.19 | 1468.65 | 176.5 | 528.94 | 5330.85 | 1786.81 | 707.51 | 9899.09 | 464.11 | 427.09 | 5456.64 | 270.97 | 43782.07 | Peroxisome biogenesis factor 10 (EC 2.3.2.27) (Peroxin-10) (Peroxisomal biogenesis factor 10) (Peroxisome assembly protein 10) | Pex10 | cellular response to reactive oxygen species [GO:0034614]; protein import into peroxisome matrix, receptor recycling [GO:0016562]; protein polyubiquitination [GO:0000209] |
| 4607.57 | 7304.6 | 3518.23 | 791.21 | 7223.24 | 2501.33 | 0 | 32.81 | 43.11 | 239.21 | 325.3 | 60.17 | 44.47 | 76.01 | 12.01 | 3415.97 | 9839.43 | 3698.3 | 43732.97 | Protocadherin-10 | PCDH10 KIAA1400 | cell adhesion [GO:0007155]; homophilic cell adhesion via plasma membrane adhesion molecules [GO:0007156]; nervous system development [GO:0007399] |
| 208.38 | 58.05 | 3444.8 | 16.76 | 49.03 | 658.82 | 1697.66 | 3.19 | 12063.67 | 60.53 | 23.65 | 4033.93 | 192.19 | 64.08 | 8841.22 | 1044.63 | 63.61 | 11195.07 | 43719.27 | Myosin-7 (Myosin heavy chain 7) (Myosin heavy chain slow isoform) (MyHC-slow) (Myosin heavy chain, cardiac muscle beta isoform) (MyHC-beta) | Myh7 | adult heart development [GO:0007512]; ATP metabolic process [GO:0046034]; cardiac muscle contraction [GO:0060048]; cardiac muscle hypertrophy in response to stress [GO:0014898]; muscle filament sliding [GO:0030049]; regulation of heart rate [GO:0002027]; regulation of slow-twitch skeletal muscle fiber contraction [GO:0031449]; regulation of the force of heart contraction [GO:0002026]; regulation of the force of skeletal muscle contraction [GO:0014728]; skeletal muscle contraction [GO:0003009]; transition between fast and slow fiber [GO:0014883]; ventricular cardiac muscle tissue morphogenesis [GO:0055010] |

| | | | | | | | | | | | | | | | | | | | | | |
|---|---|---|---|---|---|---|---|---|---|---|---|---|---|---|---|---|---|---|---|---|---|
| 3.66 | 14077.21 | 4.2 | 0 | 4648.05 | 54.79 | 0 | 4592.95 | 14.16 | 0.51 | 2104.27 | 9.25 | 11.11 | 10947.8 | 46.24 | 2.47 | 6889.04 | 45.59 | 43451.3 | DnaJ homolog subfamily C member 2 | dnajc2 zgc:85671 | 'de novo' cotranslational protein folding [GO:0051083]; positive regulation of DNA-templated transcription [GO:0045893]; regulation of translational fidelity [GO:0006450] |
| 504.66 | 5633.15 | 1474.06 | 471.23 | 6413.9 | 1458.32 | 2398.44 | 3713.06 | 347.35 | 263.99 | 2326.48 | 1148.47 | 1034.15 | 8329.21 | 1036.39 | 1551.75 | 3839.05 | 719.78 | 42663.44 | V-set domain-containing T-cell activation inhibitor 1 | Vtcn1 | adaptive immune response [GO:0002250]; negative regulation of apoptotic process [GO:0043066]; negative regulation of T cell activation [GO:0050868]; negative regulation of T cell proliferation [GO:0042130]; positive regulation of interleukin-2 production [GO:0032743]; positive regulation of T cell proliferation [GO:0042102]; regulation of cytokine production [GO:0001817]; response to protozoan [GO:0001562]; T cell receptor signaling pathway [GO:0050852] |
| 191.61 | 4509.55 | 272.22 | 91.36 | 12995.7 | 663.39 | 3.58 | 8350.04 | 77.75 | 197.32 | 3754.7 | 299.36 | 440.99 | 6964.16 | 133.93 | 55.54 | 3127.17 | 512.47 | 42640.84 | Phosphatidylinositol N-acetylglucosaminyltransferase subunit C (Phosphatidylinositol-glycan biosynthesis class C protein) (PIG-C) | PIGC GPI2 | GPI anchor biosynthetic process [GO:0006506] |

| | | | | | | | | | | | | | | | | | | | | | |
|---|---|---|---|---|---|---|---|---|---|---|---|---|---|---|---|---|---|---|---|---|---|
| 1294.39 | 3998.92 | 3104.03 | 841.64 | 4733.15 | 3043.01 | 2596.65 | 1999.95 | 927.67 | 697.84 | 1199.3 | 3525.11 | 1245.5 | 6053.07 | 1532.64 | 2543.06 | 1939.71 | 1252.31 | 42527.95 | Semaphorin-3F (Sema III/F) (Semaphorin IV) (Sema IV) | SEMA3F | axon extension involved in axon guidance [GO:0048846]; axon guidance [GO:0007411]; branchiomotor neuron axon guidance [GO:0021785]; facial nerve structural organization [GO:0021612]; negative chemotaxis [GO:0050919]; negative regulation of axon extension involved in axon guidance [GO:0048843]; nerve development [GO:0021675]; neural crest cell migration [GO:0001755]; neural crest cell migration involved in autonomic nervous system development [GO:1901166]; positive regulation of cell migration [GO:0030335]; regulation of postsynapse organization [GO:0099175]; semaphorin-plexin signaling pathway [GO:0071526]; sympathetic ganglion development [GO:0061549]; sympathetic neuron projection extension [GO:0097490]; sympathetic neuron projection guidance [GO:0097491]; trigeminal nerve structural organization [GO:0021637]; ventral trunk neural |
| 1703.8 | 2802.14 | 4181.27 | 1156.11 | 1476.58 | 5791.83 | 13.63 | 3412.41 | 3749.77 | 2372.39 | 1140.13 | 2194.29 | 2151.85 | 2059.8 | 2651.36 | 634.24 | 2523.94 | 2304.74 | 42320.28 | Xylulose kinase (Xylulokinase) (EC 2.7.1.17) | Xylb | D-xylose metabolic process [GO:0042732]; xylulose metabolic process [GO:0005997] |
| 1493.01 | 4540.89 | 1723.82 | 2476.6 | 3933.24 | 1963.93 | 2146.74 | 2770.53 | 471.67 | 1098.33 | 1859.49 | 1632.1 | 2672.74 | 4407.37 | 934.4 | 2331.54 | 4665.58 | 1030.68 | 42152.66 | Ankyrin repeat domain-containing protein 34C | Ankrd34c | |
| 57.74 | 0 | 0 | 340.23 | 0 | 6.67 | 25233.48 | 0 | 0 | 242.42 | 682.72 | 1.12 | 24.56 | 0 | 0 | 15122.52 | 400.37 | 24.89 | 42136.72 | UDP-glucuronosyltransferase 2B15 (UDPGT 2B15) (UGT2B15) (EC 2.4.1.17) (HLUG4) (UDP-glucuronosyltransferase 2B8) (UDPGT 2B8) (UDPGTh-3) | UGT2B15 UGT2B8 | estrogen metabolic process [GO:0008210]; steroid metabolic process [GO:0008202]; xenobiotic metabolic process [GO:0006805] |

| | | | | | | | | | | | | | | | | | | | | | |
|---|---|---|---|---|---|---|---|---|---|---|---|---|---|---|---|---|---|---|---|---|---|
| 3968.45 | 2667.92 | 1383.86 | 1020.76 | 4157.82 | 4108.44 | 41.39 | 2759.6 | 184.38 | 1672.51 | 1899.71 | 1263.62 | 5988.75 | 5576.3 | 1556.79 | 609.79 | 2718.6 | 530.03 | 42108.72 | E3 ubiquitin-protein ligase TRAF7 (EC 2.3.2.-) (EC 2.3.2.27) (RING finger and WD repeat-containing protein 1) (RING finger protein 119) (RING-type E3 ubiquitin transferase TRAF7) (TNF receptor-associated factor 7) | TRAF7 RFWD1 RNF119 | apoptotic process [GO:0006915]; positive regulation of apoptotic signaling pathway [GO:2001235]; positive regulation of MAPK cascade [GO:0043410]; positive regulation of neuron apoptotic process [GO:0043525]; positive regulation of ubiquitin-dependent protein catabolic process [GO:2000060]; protein K29-linked ubiquitination [GO:0035519]; protein ubiquitination [GO:0016567]; regulation of ERK1 and ERK2 cascade [GO:0070372] |
| 939.61 | 1997.46 | 2429.29 | 702.19 | 1498.25 | 6765.95 | 795.2 | 1050.98 | 1112.59 | 1413.11 | 1953.83 | 4406.17 | 703.92 | 5357.11 | 2408.28 | 2419.11 | 3947.27 | 1843.95 | 41744.27 | Voltage-gated potassium channel subunit beta-1 (EC 1.1.1.-) (K(+) channel subunit beta-1) (Kv-beta-1) | KCNAB1 KVB1.1 | regulation of potassium ion transmembrane transport [GO:1901379] |
| 4051.49 | 3621.95 | 1230 | 790.15 | 6187.44 | 1721.71 | 1004.68 | 871.95 | 561.52 | 554.42 | 1949.66 | 824.01 | 775.73 | 683.6 | 155.96 | 922.38 | 15237.21 | 567.86 | 41711.72 | Diacylglycerol kinase theta (DAG kinase theta) (EC 2.7.1.107) (EC 2.7.1.93) (Diglyceride kinase theta) (DGK-theta) | Dgkq | adenylate cyclase-activating G protein-coupled receptor signaling pathway [GO:0007189]; cellular response to bile acid [GO:1903413]; diacylglycerol metabolic process [GO:0046339]; epidermal growth factor receptor signaling pathway [GO:0007173]; lipid phosphorylation [GO:0046834]; negative regulation of gene expression [GO:0010629]; phosphatidic acid biosynthetic process [GO:0006654]; phospholipase C-activating G protein-coupled receptor signaling pathway [GO:0007200]; positive regulation of gene expression [GO:0010628]; positive regulation of protein kinase C signaling [GO:0090037]; regulation of cholesterol metabolic process [GO:0090181]; regulation of cortisol biosynthetic process [GO:2000064]; regulation of gluconeogenesis [GO:0006111]; regulation of progesterone biosynthetic process [GO:2000182]; regulation of synaptic vesicle endocytosis [GO:1900242]; regulation of TORC1 signaling [GO:1903432]; regulation of |
| 2742.98 | 6429.34 | 3257.54 | 830.65 | 2810.27 | 4606.71 | 32.88 | 1884.59 | 1601.52 | 960.23 | 898.64 | 1501.43 | 1674.14 | 4901.81 | 2000.44 | 321.54 | 2643.28 | 2378.33 | 41476.32 | Transmembrane protein 134 | TMEM134 | |

| | | | | | | | | | | | | | | | | | | | | | |
|---|---|---|---|---|---|---|---|---|---|---|---|---|---|---|---|---|---|---|---|---|---|
| 121.53 | 5722.65 | 526.55 | 102.96 | 9895.61 | 255.5 | 1841.28 | 3953.83 | 45.87 | 48.86 | 3527.6 | 1975.06 | 80.34 | 9080.94 | 86.98 | 321.85 | 3474.25 | 82.14 | 41143.8 | F-box/LRR-repeat protein 3 (F-box and leucine-rich repeat protein 3A) (F-box/LRR-repeat protein 3A) | FBXL3 FBL3A FBXL3A | entrainment of circadian clock by photoperiod [GO:0043153]; protein destabilization [GO:0031648]; protein ubiquitination [GO:0016567]; regulation of circadian rhythm [GO:0042752]; rhythmic process [GO:0048511]; SCF-dependent proteasomal ubiquitin-dependent protein catabolic process [GO:0031146] |
| 12.81 | 18538.75 | 249.66 | 31.24 | 7413.93 | 52.02 | 11.06 | 1936.08 | 4.75 | 29.75 | 2001.23 | 13 | 4.01 | 7227.71 | 1.09 | 10.53 | 3576.79 | 19.91 | 41134.32 | TBC1 domain family member 9 (TBC1 domain family member 9A) | TBC1D9 KIAA0882 TBC1D9A | |
| 754.07 | 65.12 | 1991.32 | 325.07 | 86.08 | 22531.89 | 14.4 | 12.95 | 18.78 | 104.88 | 20.99 | 88.56 | 298.02 | 82.3 | 14277.5 | 45.81 | 40.33 | 7.04 | 40765.11 | HLA class II histocompatibility antigen, DR alpha chain (MHC class II antigen DRA) | HLA-DRA HLA-DRA1 | adaptive immune response [GO:0002250]; antigen processing and presentation of endogenous peptide antigen via MHC class II [GO:0002491]; antigen processing and presentation of exogenous peptide antigen via MHC class II [GO:0019886]; antigen processing and presentation of peptide or polysaccharide antigen via MHC class II [GO:0002504]; cognition [GO:0050890]; immune response [GO:0006955]; myeloid dendritic cell antigen processing and presentation [GO:0002469]; peptide antigen assembly with MHC class II protein complex [GO:0002503]; positive regulation of CD4-positive, alpha-beta T cell activation [GO:2000516]; positive regulation of CD4-positive, CD25-positive, alpha-beta regulatory T cell differentiation [GO:0032831]; positive regulation of immune response [GO:0050778]; positive regulation of memory T cell differentiation [GO:0043382]; positive regulation of T cell activation [GO:0050870]; positive |

| | | | | | | | | | | | | | | | | | | | | | |
|---|---|---|---|---|---|---|---|---|---|---|---|---|---|---|---|---|---|---|---|---|---|
| 0.65 | 1.68 | 38396.1 | 1.3 | 1.05 | 1158.58 | 13.32 | 0 | 6.55 | 3.55 | 0.48 | 94.65 | 1.74 | 29.67 | 920 | 11.93 | 0.18 | 9.34 | 40650.77 | Death-inducer obliterator 1 (DIO-1) (Death-associated transcription factor 1) (DATF-1) | Dido1 Datf1 Dio1 | apoptotic signaling pathway [GO:0097190]; DNA-templated transcription [GO:0006351] |
| 2084.06 | 2808 | 2127.48 | 3011.61 | 2636.97 | 4833.17 | 420.49 | 1372.65 | 1342.14 | 3130.12 | 2574.73 | 5236.48 | 2502.94 | 1889.17 | 1474.28 | 738.15 | 1216.68 | 960.42 | 40359.54 | E3 ubiquitin-protein ligase RNF6 (EC 2.3.2.27) | RNF6 SPG2 | axon extension [GO:0048675]; negative regulation of axon extension [GO:0030517]; positive regulation of DNA-templated transcription [GO:0045893]; protein K27-linked ubiquitination [GO:0044314]; protein K48-linked ubiquitination [GO:0070936]; protein K6-linked ubiquitination [GO:0085020]; protein ubiquitination [GO:0016567]; regulation of androgen receptor signaling pathway [GO:0060765]; regulation of DNA-templated transcription [GO:0006355]; ubiquitin-dependent protein catabolic process [GO:0006511] |
| 194.4 | 129.69 | 103.41 | 183 | 114.69 | 240.57 | 19803.94 | 42.95 | 187.72 | 154.17 | 458.68 | 78.5 | 262.64 | 156.77 | 185.1 | 17355.78 | 472.53 | 152.45 | 40276.99 | Carboxypeptidase E (CPE) (EC 3.4.17.10) (Carboxypeptidase H) (CPH) (Enkephalin convertase) (Prohormone-processing carboxypeptidase) | cpe | peptide metabolic process [GO:0006518]; protein processing [GO:0016485] |

| | | | | | | | | | | | | | | | | | | | | | |
|---|---|---|---|---|---|---|---|---|---|---|---|---|---|---|---|---|---|---|---|---|---|
| 9099.12 | 70.89 | 34.98 | 2371.4 | 214.82 | 1329.56 | 57.6 | 17.35 | 0 | 6788.49 | 140.11 | 369.25 | 18846.9 | 54.07 | 45.97 | 673.1 | 58.8 | 1.27 | 40173.68 | Charged multivesicular body protein 4c (Chromatin-modifying protein 4c) (CHMP4c) | chmp4c chmp4b zgc:55566 | cilium assembly [GO:0060271]; late endosome to vacuole transport via multivesicular body sorting pathway [GO:0032511]; midbody abscission [GO:0061952]; mitotic cytokinesis checkpoint signaling [GO:0044878]; negative regulation of cytokinesis [GO:0032466]; protein transport [GO:0015031]; vesicle budding from membrane [GO:0006900] |
| 172.6 | 11250.65 | 2352.45 | 246.75 | 3079.19 | 1199.01 | 73.21 | 1748.52 | 169.34 | 270.65 | 1237.51 | 1506.09 | 251.44 | 12906.27 | 430.88 | 59.93 | 2991.95 | 219 | 40165.44 | Zinc finger protein 512B | ZNF512B KIAA1196 | negative regulation of miRNA transcription [GO:1902894] |
| 0 | 1.16 | 23214.82 | 0 | 0 | 1129.7 | 14.17 | 0 | 191.78 | 0.36 | 0 | 15341.55 | 0 | 0 | 81.81 | 31.35 | 0 | 41.3 | 40048 | Kinesin-1 heavy chain (Conventional kinesin heavy chain) (Ubiquitous kinesin heavy chain) (UKHC) | Kif5b Khcs Kns1 | anterograde axonal protein transport [GO:0099641]; cellular response to type II interferon [GO:0071346]; centrosome localization [GO:0051642]; cytoplasm organization [GO:0007028]; lysosome localization [GO:0032418]; microtubule-based process [GO:0007017]; mitochondrial transport [GO:0006839]; mitochondrion transport along microtubule [GO:0047497]; mitocytosis [GO:0160040]; natural killer cell mediated cytotoxicity [GO:0042267]; plus-end-directed vesicle transport along microtubule [GO:0072383]; positive regulation of potassium ion transport [GO:0043268]; positive regulation of protein localization to plasma membrane [GO:1903078]; positive regulation of synaptic transmission, GABAergic [GO:0032230]; regulation of membrane potential [GO:0042391]; regulation of modification of synapse structure, modulating synaptic transmission [GO:0098987]; stress granule disassembly [GO:0035617]; vesicle [illegible] |

| | | | | | | | | | | | | | | | | | | | | | | |
|---|---|---|---|---|---|---|---|---|---|---|---|---|---|---|---|---|---|---|---|---|---|---|
| 3882.32 | | 4883.45 | 616.72 | 906.77 | 3532.67 | 1221.21 | 1287.96 | 4996.12 | 500.2 | 844.48 | 1230.25 | 2645.02 | 2208.93 | 1545.57 | 601.59 | 985.72 | 7401.89 | 565.52 | 39856.39 | WD repeat domain phosphoinositide-interacting protein 1 (WIPI-1) (Atg18 protein homolog) (WD40 repeat protein interacting with phosphoinositides of 49 kDa) (WIPI 49 kDa) | WIPI1 WIPI49 | autophagosome assembly [GO:0000045]; autophagy [GO:0006914]; autophagy of mitochondrion [GO:0000422]; cellular response to starvation [GO:0009267]; glycophagy [GO:0061723]; nucleophagy [GO:0044804]; pexophagy [GO:0000425]; positive regulation of autophagosome assembly [GO:2000786]; protein localization to phagophore assembly site [GO:0034497]; vesicle targeting, trans-Golgi to endosome [GO:0048203] |
| 3937.06 | | 274.5 | 6265.08 | 1906.08 | 896.03 | 1615.25 | 568.17 | 296.87 | 3900.01 | 1796.06 | 511.47 | 4031.78 | 1987.42 | 199.83 | 3565.16 | 791.41 | 807.95 | 6442.73 | 39792.86 | Cytospin-B (Sperm antigen with calponin homology and coiled-coil domains 1) | Specc1 Cytsb Kiaa4061 | associative learning [GO:0008306]; blastocyst development [GO:0001824] |
| 3231.72 | | 3922.62 | 2837.73 | 708.72 | 2359.53 | 3230.09 | 396.6 | 1136.39 | 376.88 | 726.2 | 978.03 | 1792.48 | 2979.05 | 6267.76 | 2378.48 | 1665.98 | 3767.41 | 1000.61 | 39756.28 | A-kinase anchor protein 9 (AKAP-9) (Protein kinase A-anchoring protein 9) (PRKA9) | Akap9 Kiaa0803 | maintenance of centrosome location [GO:0051661]; microtubule nucleation [GO:0007020]; protein-containing complex localization [GO:0031503]; regulation of Golgi organization [GO:1903358]; Sertoli cell development [GO:0060009]; signal transduction [GO:0007165]; spermatogenesis [GO:0007283] |

| | | | | | | | | | | | | | | | | | | | | | |
|---|---|---|---|---|---|---|---|---|---|---|---|---|---|---|---|---|---|---|---|---|---|
| 3079.43 | 2187.24 | 3379.41 | 1182.26 | 2550.94 | 5226.97 | 2471.71 | 1379.19 | 1247.12 | 925.53 | 795.55 | 2658.72 | 2302.92 | 2806.26 | 1716.62 | 1976.91 | 1991.86 | 1774.09 | 39652.73 | Calpain-3 (EC 3.4.22.54) (Calcium-activated neutral proteinase 3) (CANP 3) (Calpain L3) (Calpain p94) (Muscle-specific calcium-activated neutral protease 3) | Capn3 | calcium-dependent self proteolysis [GO:1990092]; cellular response to calcium ion [GO:0071277]; cellular response to salt stress [GO:0071472]; G1 to G0 transition involved in cell differentiation [GO:0070315]; muscle structure development [GO:0061061]; myofibril assembly [GO:0030239]; negative regulation of apoptotic process [GO:0043066]; negative regulation of DNA-templated transcription [GO:0045892]; negative regulation of protein sumoylation [GO:0033234]; negative regulation of skeletal muscle cell differentiation [GO:2001015]; positive regulation of DNA-templated transcription [GO:0045893]; positive regulation of proteolysis [GO:0045862]; positive regulation of release of sequestered calcium ion into cytosol [GO:0051281]; positive regulation of satellite cell activation involved in skeletal muscle regeneration [GO:0014718]; programmed cell death [GO:0012501]; protein catabolic process [GO:0030163]; protein |
| 0 | 0 | 618.82 | 2.48 | 0 | 12.92 | 0 | 0 | 13006.49 | 1.99 | 0 | 5226.53 | 0 | 4.66 | 5814.69 | 0.52 | 13.08 | 14609.27 | 39311.45 | Transmembrane protein 198 | tmem198 | Wnt signaling pathway [GO:0016055] |
| 1.34 | 73.03 | 3787.85 | 0 | 136.18 | 1054.48 | 19.96 | 91.07 | 18.96 | 67.41 | 144.86 | 33376.35 | 44.39 | 49.91 | 198.57 | 38.88 | 0 | 151.61 | 39254.85 | Acyl-coenzyme A thioesterase 8 (Acyl-CoA thioesterase 8) (EC 3.1.2.1) (EC 3.1.2.11) (EC 3.1.2.2) (EC 3.1.2.3) (EC 3.1.2.5) (Choloyl-coenzyme A thioesterase) (EC 3.1.2.27) (Peroxisomal acyl-coenzyme A thioester hydrolase 1) (PTE-1) (Peroxisomal long-chain acyl-CoA thioesterase 1) | Acot8 Pte1 | acyl-CoA metabolic process [GO:0006637]; dicarboxylic acid catabolic process [GO:0043649]; fatty acid catabolic process [GO:0009062]; negative regulation of glycoprotein biosynthetic process [GO:0010561]; peroxisome fission [GO:0016559] |

| | | | | | | | | | | | | | | | | | | | | | |
|---|---|---|---|---|---|---|---|---|---|---|---|---|---|---|---|---|---|---|---|---|---|
| 59.68 | 7603.33 | 0 | 112.34 | 7942.99 | 24.68 | 18.4 | 2955.25 | 1.36 | 417.98 | 2981.37 | 0 | 1.97 | 11786.64 | 2.19 | 153.3 | 5037.83 | 21.93 | 39121.24 | LIM and senescent cell antigen-like-containing domain protein 1 (Particularly interesting new Cys-His protein 1) (PINCH-1) (Renal carcinoma antigen NY-REN-48) | LIMS1 PINCH PINCH1 | cell-cell adhesion [GO:0098609]; cell-cell junction organization [GO:0045216]; cellular response to transforming growth factor beta stimulus [GO:0071560]; establishment of protein localization [GO:0045184]; negative regulation of DNA-templated transcription [GO:0045892]; positive regulation of canonical NF-kappaB signal transduction [GO:0043123]; positive regulation of cell-substrate adhesion [GO:0010811]; positive regulation of focal adhesion assembly [GO:0051894]; positive regulation of gene expression [GO:0010628]; positive regulation of GTPase activity [GO:0043547]; positive regulation of integrin-mediated signaling pathway [GO:2001046]; positive regulation of substrate adhesion-dependent cell spreading [GO:1900026]; tumor necrosis factor-mediated signaling pathway [GO:0033209] |
| 4143.53 | 1225.59 | 3341.62 | 1347.44 | 3384.66 | 3638.64 | 49.16 | 1283.94 | 869.26 | 4015.87 | 1698.3 | 2605.64 | 2853.12 | 2920.06 | 1819.7 | 682.87 | 1791.89 | 1330.49 | 39001.78 | Serine/threonine-protein phosphatase with EF-hands 2 (PPEF-2) (EC 3.1.3.16) | PPEF2 | cellular response to hydrogen peroxide [GO:0070301]; detection of stimulus involved in sensory perception [GO:0050906]; negative regulation of apoptotic signaling pathway [GO:2001234]; negative regulation of MAPK cascade [GO:0043409]; protein dephosphorylation [GO:0006470]; visual perception [GO:0007601] |
| 488.9 | 8391.3 | 46.69 | 1201.24 | 6049.49 | 60.86 | 9.69 | 2154.29 | 5.03 | 571.33 | 3459.7 | 512.56 | 233.92 | 9957.87 | 32.71 | 402.42 | 5255.4 | 55.44 | 38888.84 | Carnosine N-methyltransferase (EC 2.1.1.22) | CARNMT1 C9orf41 | carnosine metabolic process [GO:0035498]; L-histidine catabolic process [GO:0006548]; methylation [GO:0032259] |

| | | | | | | | | | | | | | | | | | | | | | |
|---|---|---|---|---|---|---|---|---|---|---|---|---|---|---|---|---|---|---|---|---|---|
| 12.15 | 1.1 | 1929.2 | 1.86 | 0.81 | 29.04 | 1.07 | 0.95 | 11686.07 | 2.9 | 2.63 | 3463.97 | 10.22 | 7.85 | 9941.94 | 4.69 | 7.49 | 11645.62 | 38749.56 | CUGBP Elav-like family member 2 (CELF-2) (Bruno-like protein 3) (CUG triplet repeat RNA-binding protein 2) (CUG-BP2) (CUG-BP- and ETR-3-like factor 2) (ELAV-type RNA-binding protein 3) (ETR-3) (RNA-binding protein BRUNOL-3) | celf2 brunol3 cugbp2 etr3 | mRNA processing [GO:0006397] |
| 19.03 | 23.7 | 13890.74 | 2.72 | 3.93 | 20869.96 | 79.15 | 3.33 | 11.13 | 0 | 2.41 | 3160.6 | 4.38 | 109.58 | 283.57 | 46.98 | 16.04 | 16.63 | 38543.88 | F-actin-uncapping protein LRRC16A (CARMIL homolog) (Capping protein regulator and myosin 1 linker protein 1) (Capping protein, Arp2/3 and myosin-I linker homolog 1) (Capping protein, Arp2/3 and myosin-I linker protein 1) (Leucine-rich repeat-containing protein 16A) | CARMIL1 CARMIL LRRC16 LRRC16A | actin filament network formation [GO:0051639]; actin filament organization [GO:0007015]; barbed-end actin filament uncapping [GO:0051638]; cell migration [GO:0016477]; lamellipodium assembly [GO:0030032]; macropinocytosis [GO:0044351]; negative regulation of barbed-end actin filament capping [GO:2000813]; positive regulation of actin filament polymerization [GO:0030838]; positive regulation of cell migration [GO:0030335]; positive regulation of lamellipodium organization [GO:1902745]; positive regulation of stress fiber assembly [GO:0051496]; positive regulation of substrate adhesion-dependent cell spreading [GO:1900026]; regulation of Arp2/3 complex-mediated actin nucleation [GO:0034315]; ruffle organization [GO:0031529]; urate metabolic process [GO:0046415] |

| | | | | | | | | | | | | | | | | | | | | | |
|---|---|---|---|---|---|---|---|---|---|---|---|---|---|---|---|---|---|---|---|---|---|
| 507.3 | 1660.31 | 4561.71 | 733.63 | 2238.23 | 16390.68 | 18.66 | 617.95 | 64.25 | 630.92 | 1286.28 | 2979.03 | 1769.42 | 1504.01 | 1840.9 | 493.03 | 1153.44 | 63.76 | 38513.51 | Claudin-5 | Cldn5 | bicellular tight junction assembly [GO:0070830]; calcium-independent cell-cell adhesion via plasma membrane cell-adhesion molecules [GO:0016338]; cell adhesion [GO:0007155]; cell-cell junction assembly [GO:0007043]; establishment of blood-retinal barrier [GO:1990963]; establishment of endothelial barrier [GO:0061028]; maintenance of blood-brain barrier [GO:0035633]; myelination [GO:0042552]; negative regulation of angiogenesis [GO:0016525]; negative regulation of cell migration [GO:0030336]; negative regulation of complement-dependent cytotoxicity [GO:1903660]; negative regulation of gene expression [GO:0010629]; negative regulation of vascular permeability [GO:0043116]; positive regulation of bicellular tight junction assembly [GO:1903348]; positive regulation of cell population proliferation [GO:0008284]; positive regulation of establishment of endothelial barrier [GO:1903142]; positive regulation of gene expression |
| 0 | 6157.87 | 0 | 0 | 1004.74 | 6.86 | 0 | 3266.59 | 0 | 0 | 4503.92 | 0 | 0 | 19653.91 | 0.58 | 0 | 3462.74 | 10.05 | 38067.26 | P2Y purinoceptor 14 (P2Y14) (UDP-glucose receptor) | P2RY14 | G protein-coupled receptor signaling pathway [GO:0007186] |
| 1625.76 | 1438.58 | 713.23 | 5685.6 | 2814.99 | 1286.48 | 7.85 | 1089.1 | 84 | 5960.42 | 7227.37 | 362.62 | 1383.13 | 1286.98 | 102.42 | 1542 | 4994.08 | 373.23 | 37977.84 | TIMELESS-interacting protein | tipin zgc:91928 | cell cycle phase transition [GO:0044770]; cell division [GO:0051301]; DNA replication checkpoint signaling [GO:0000076]; mitotic intra-S DNA damage checkpoint signaling [GO:0031573]; positive regulation of cell population proliferation [GO:0008284]; replication fork arrest [GO:0043111]; replication fork processing [GO:0031297] |

| | | | | | | | | | | | | | | | | | | | | | | |
|---|---|---|---|---|---|---|---|---|---|---|---|---|---|---|---|---|---|---|---|---|---|---|
| 1801.28 | | 2386.13 | 2715.07 | 1869.61 | 3813.83 | 4351.57 | 63.98 | 1407.82 | 340.12 | 797.87 | 878.11 | 1319.44 | 5028.53 | 4535.86 | 2678.46 | 492.93 | 1478.93 | 1724.04 | 37683.58 | Cyclin-dependent kinase 14 (EC 2.7.11.22) (Cell division protein kinase 14) (Serine/threonine-protein kinase PFTAIRE-1) | Cdk14 Kiaa0834 Pftk1 | cell division [GO:0051301]; G2/M transition of mitotic cell cycle [GO:0000086]; regulation of canonical Wnt signaling pathway [GO:0060828]; Wnt signaling pathway [GO:0016055] |
| 2254.42 | | 3787.75 | 263.92 | 7136.05 | 6515.18 | 106.19 | 3.18 | 2838.78 | 18.55 | 2081.77 | 3325.06 | 768.45 | 1158.84 | 2517.01 | 2.19 | 1339.62 | 3496.3 | 45.74 | 37659 | Tropomodulin-3 (Ubiquitous tropomodulin) (U-Tmod) | TMOD3 | actin filament organization [GO:0007015]; erythrocyte development [GO:0048821]; mitotic cell cycle phase transition [GO:0044772]; muscle contraction [GO:0006936]; myofibril assembly [GO:0030239]; pointed-end actin filament capping [GO:0051694]; positive regulation of mitotic cell cycle phase transition [GO:1901992] |

| | | | | | | | | | | | | | | | | | | | | | |
|---|---|---|---|---|---|---|---|---|---|---|---|---|---|---|---|---|---|---|---|---|---|
| 1096.14 | 3473.72 | 995.6 | 5110.63 | 5975.21 | 2068.2 | 800.33 | 2430.65 | 140.4 | 857.88 | 1541.47 | 929.2 | 1131.35 | 3620.54 | 289.19 | 1724.8 | 5135.07 | 326.64 | 37647.02 | LINE-1 retrotransposable element ORF1 protein (L1-ORF1p) (LINE retrotransposable element 1) (LINE1 retrotransposable element 1) (Transposase element L1Md-A101/L1Md-A102/L1Md-A2) | Lire1 | retrotransposition [GO:0032197] |
| 465.73 | 9977.13 | 287.69 | 166.1 | 5773 | 307.54 | 6.43 | 3652.95 | 175.3 | 112.24 | 1706.52 | 195.12 | 773.63 | 8870.22 | 277.98 | 134.86 | 4409.23 | 355.25 | 37646.92 | NK-tumor recognition protein (NK-TR protein) (Natural-killer cells cyclophilin-related protein) (Peptidyl-prolyl cis-trans isomerase NKTR) (PPIase) (EC 5.2.1.8) (Rotamase) | NKTR | protein folding [GO:0006457] |
| 1635.72 | 1995.31 | 2107.26 | 1504.26 | 2326.26 | 3528.53 | 3170.13 | 996.69 | 692.51 | 1006.43 | 1189.88 | 2716.41 | 2889.18 | 3979.8 | 1147.38 | 3185.89 | 2462.98 | 1021.5 | 37556.12 | TBC1 domain family member 14 | TBC1D14 KIAA1322 | autophagy [GO:0006914]; negative regulation of autophagy [GO:0010507]; recycling endosome to Golgi transport [GO:0071955]; regulation of autophagosome assembly [GO:2000785] |

| | | | | | | | | | | | | | | | | | | | | | |
|---|---|---|---|---|---|---|---|---|---|---|---|---|---|---|---|---|---|---|---|---|---|
| 3807.67 | 692.45 | 3262.8 | 717.86 | 514.92 | 1579.14 | 64.82 | 172.26 | 5432.55 | 1343.36 | 460.34 | 4416.88 | 3502.06 | 722.99 | 5728.91 | 370.38 | 472.87 | 4260.64 | 37522.9 | Ras association domain-containing protein 1 (Protein 123F2) | Rassf1 | DNA damage response [GO:0006974]; positive regulation of protein ubiquitination [GO:0031398]; protein stabilization [GO:0050821]; Ras protein signal transduction [GO:0007265]; regulation of cell cycle [GO:0051726]; regulation of cell cycle G1/S phase transition [GO:1902806] |
| 0 | 0 | 1226.54 | 1.87 | 0 | 15.35 | 0 | 0 | 11130.96 | 0 | 0 | 2938.8 | 3 | 0.59 | 10375.69 | 2.35 | 1.64 | 11596.68 | 37293.47 | Dynamin-1 (EC 3.6.5.5) (B-dynamin) (D100) (Dynamin I) (Dynamin, brain) | Dnm1 Dnm | clathrin coat assembly involved in endocytosis [GO:0099049]; endocytosis [GO:0006897]; endosome organization [GO:0007032]; G protein-coupled receptor internalization [GO:0002031]; modulation of chemical synaptic transmission [GO:0050804]; positive regulation of synaptic vesicle endocytosis [GO:1900244]; positive regulation of synaptic vesicle recycling [GO:1903423]; protein homooligomerization [GO:0051260]; protein homotetramerization [GO:0051289]; receptor internalization [GO:0031623]; receptor-mediated endocytosis [GO:0006898]; regulation of synaptic vesicle endocytosis [GO:1900242]; regulation of vesicle size [GO:0097494]; response to amyloid-beta [GO:1904645]; synaptic vesicle budding from presynaptic endocytic zone membrane [GO:0016185]; synaptic vesicle endocytosis [GO:0048488]; vesicle scission [GO:0099050] |
| 2019.34 | 1390.4 | 2250.26 | 915.03 | 1328.73 | 2704.64 | 6591.32 | 598.02 | 2599.63 | 998.99 | 777.01 | 1954.16 | 2082.31 | 1842.89 | 2450.84 | 3927.87 | 1010.25 | 1835.97 | 37277.66 | UPF0606 protein KIAA1549L | KIAA1549L C11orf41 C11orf69 | |

| | | | | | | | | | | | | | | | | | | | | | |
|---|---|---|---|---|---|---|---|---|---|---|---|---|---|---|---|---|---|---|---|---|---|
| 95.98 | 7099.95 | 587.33 | 179.84 | 4258.91 | 354.28 | 260.38 | 4218.57 | 650.15 | 43.2 | 1478.57 | 364.23 | 101.28 | 9231.26 | 729.69 | 571.46 | 5955.13 | 978.97 | 37159.18 | Ubiquitin carboxyl-terminal hydrolase 38 (EC 3.4.19.12) (Deubiquitinating enzyme 38) (HP43.8KD) (Ubiquitin thioesterase 38) (Ubiquitin-specific-processing protease 38) | USP38 KIAA1891 | negative regulation of innate immune response [GO:0045824]; negative regulation of proteasomal ubiquitin-dependent protein catabolic process [GO:0032435]; protein K33-linked deubiquitination [GO:1990168]; regulation of protein stability [GO:0031647]; ubiquitin-dependent protein catabolic process [GO:0006511] |
| 0 | 0 | 913.15 | 3.19 | 1.23 | 17.39 | 0 | 2.37 | 9551.33 | 1.53 | 0 | 3624.34 | 5.95 | 4.98 | 10360.52 | 0.67 | 7.01 | 12652.54 | 37146.2 | Ubiquitin-conjugating enzyme E2 K (EC 2.3.2.23) (E2 ubiquitin-conjugating enzyme K) (Huntingtin-interacting protein 2) (HIP-2) (Ubiquitin carrier protein) (Ubiquitin-conjugating enzyme E2-25 kDa) (Ubiquitin-conjugating enzyme E2(25K)) (Ubiquitin-conjugating enzyme E2-25K) (Ubiquitin-protein ligase) | UBE2K HIP2 | protein K48-linked ubiquitination [GO:0070936]; protein polyubiquitination [GO:0000209] |
| 1103.47 | 1137.4 | 1383.79 | 2133.86 | 2977.26 | 3354.32 | 1682.14 | 966.45 | 1448.33 | 1050.27 | 1032.96 | 2645.34 | 2788.21 | 4712.71 | 1394.18 | 2243.2 | 3049.47 | 2028.52 | 37131.88 | Nebulette (Actin-binding Z-disk protein) | Nebl Lnebl | |

| | | | | | | | | | | | | | | | | | | | | | | |
|---|---|---|---|---|---|---|---|---|---|---|---|---|---|---|---|---|---|---|---|---|---|---|
| 520.52 | | 298.84 | 137.62 | 215.13 | 29.07 | 48.99 | 17354.85 | 22.43 | 3.9 | 299.9 | 735.18 | 18.12 | 532.5 | 46.52 | 62.76 | 15396.15 | 369.28 | 27.58 | 36119.34 | 2',3'-cyclic-nucleotide 3'-phosphodiesterase (CNP) (CNPase) (EC 3.1.4.37) | Cnp Cnp1 | adult locomotory behavior [GO:0008344]; axonogenesis [GO:0007409]; cyclic nucleotide catabolic process [GO:0009214]; oligodendrocyte differentiation [GO:0048709]; response to toxic substance [GO:0009636] |
| 2947.68 | | 1974.16 | 7933.7 | 469.73 | 1320.88 | 1946.76 | 141.26 | 1275.85 | 4160.67 | 1295.8 | 1359.63 | 5258.21 | 995.98 | 1436.03 | 1083.44 | 590.5 | 802.92 | 1112 | 36105.2 | U5 small nuclear ribonucleoprotein 40 kDa protein (U5 snRNP 40 kDa protein) (WD repeat-containing protein 57) | Snrnp40 Wdr57 | mRNA processing [GO:0006397]; RNA splicing [GO:0008380] |
| 6860.09 | | 163.5 | 96.23 | 2066.72 | 1460.45 | 2690.94 | 27.68 | 130.9 | 2.8 | 5482.81 | 780.75 | 310.87 | 15277.66 | 70.85 | 65.81 | 307.75 | 201.74 | 14.17 | 36011.72 | O-phosphoseryl-tRNA(Sec) selenium transferase (EC 2.9.1.2) (Selenocysteine synthase) (Sec synthase) (Selenocysteinyl-tRNA(Sec) synthase) (Sep-tRNA:Sec-tRNA synthase) (SepSecS) (Soluble liver antigen/liver pancreas antigen-like) (UGA suppressor tRNA-associated protein homolog) | sepsecs sla/lpl zgc:55980 | conversion of seryl-tRNAsec to selenocys-tRNAsec [GO:0001717]; selenocysteine incorporation [GO:0001514] |

| | | | | | | | | | | | | | | | | | | | | | |
|---|---|---|---|---|---|---|---|---|---|---|---|---|---|---|---|---|---|---|---|---|---|
| 1174.74 | 330.45 | 2142.86 | 369.12 | 686.25 | 2891.92 | 8766.11 | 121.47 | 3468.02 | 555.98 | 444.06 | 1461.12 | 1285.05 | 1185.42 | 3115.71 | 4351.34 | 685.57 | 2576.02 | 35611.21 | RING finger protein 122 | Rnf122 | negative regulation of mitochondrial membrane potential [GO:0010917]; positive regulation of apoptotic process [GO:0043065]; proteasome-mediated ubiquitin-dependent protein catabolic process [GO:0043161]; protein autoubiquitination [GO:0051865] |
| 7805 | 83.29 | 129.42 | 1772.43 | 384.2 | 636.88 | 18.79 | 46.57 | 70.94 | 8374.57 | 185.71 | 321.62 | 14995.54 | 39.78 | 72.93 | 451.92 | 60.9 | 42.77 | 35493.26 | Dedicator of cytokinesis protein 3 (Modifier of cell adhesion) (Presenilin-binding protein) (PBP) | DOCK3 KIAA0299 MOCA | neurotrophin TRK receptor signaling pathway [GO:0048011]; regulation of small GTPase mediated signal transduction [GO:0051056]; small GTPase-mediated signal transduction [GO:0007264] |

| | | | | | | | | | | | | | | | | | | | | |
|---|---|---|---|---|---|---|---|---|---|---|---|---|---|---|---|---|---|---|---|---|
| 2644.06 | 1939.09 | 3053.09 | 811.04 | 2501.18 | 5381.2 | 63.1 | 1611.81 | 329.09 | 1251.86 | 930.82 | 1653.26 | 4600.79 | 2724.65 | 2354.86 | 540.02 | 1558.55 | 1465.85 | 35414.32 | E3 ubiquitin-protein ligase rnf213-beta (EC 2.3.2.27) (EC 3.6.4.-) (E3 ubiquitin-lipopolysaccharide ligase rnf213-beta) (EC 2.3.2.-) (Mysterin-B) (Mysterin-beta) (RING finger protein 213-B) (RING finger protein 213-beta) | rnf213b | angiogenesis [GO:0001525]; defense response to bacterium [GO:0042742]; immune system process [GO:0002376]; lipid droplet formation [GO:0140042]; lipid ubiquitination [GO:0120323]; protein K63-linked ubiquitination [GO:0070534]; regulation of lipid metabolic process [GO:0019216]; xenophagy [GO:0098792] |
| 4111.73 | 1170.75 | 2668.67 | 1042.86 | 1853.41 | 4283.73 | 382.68 | 899.51 | 413.14 | 986.38 | 725.44 | 1966.9 | 4017.84 | 4350.2 | 2127.92 | 1332.87 | 1924.87 | 1074.37 | 35333.27 | 1-acyl-sn-glycerol-3-phosphate acyltransferase gamma (EC 2.3.1.51) (1-acylglycerol-3-phosphate O-acyltransferase 3) (1-AGP acyltransferase 3) (1-AGPAT 3) (Lysophosphatidic acid acyltransferase gamma) (LPAAT-gamma) | Agpat3 Lpaat3 | CDP-diacylglycerol biosynthetic process [GO:0016024] |
| 1360.07 | 3204.8 | 2763.28 | 978.12 | 4293.46 | 2952.63 | 47.55 | 1522.76 | 1542.21 | 688.87 | 1866.86 | 2215.89 | 1210.14 | 3268.43 | 1451.44 | 404.82 | 4390.6 | 1078.17 | 35240.1 | Transcription factor COE3 (Early B-cell factor 3) (EBF-3) (Olf-1/EBF-like 2) (O/E-2) (OE-2) | Ebf3 Coe3 | positive regulation of DNA-templated transcription [GO:0045893]; positive regulation of transcription by RNA polymerase II [GO:0045944] |

| | | | | | | | | | | | | | | | | | | | | | |
|---|---|---|---|---|---|---|---|---|---|---|---|---|---|---|---|---|---|---|---|---|---|
| 50.7 | 0.73 | 5.32 | 179.71 | 1.36 | 7.6 | 20334.33 | 1.07 | 7.61 | 101.23 | 378.94 | 0 | 24.97 | 0.87 | 2.56 | 13527.58 | 309.95 | 9.41 | 34943.94 | Mothers against decapentaplegic homolog 1 (MAD homolog 1) (Mothers against DPP homolog 1) (SMAD family member 1) (SMAD 1) (Smad1) | smad1 madh1 | anatomical structure morphogenesis [GO:0009653]; BMP signaling pathway [GO:0030509]; cell differentiation [GO:0030154]; dorsal/ventral pattern formation [GO:0009953]; embryonic pattern specification [GO:0009880]; myeloid cell development [GO:0061515]; negative regulation of DNA-templated transcription [GO:0045892]; regulation of DNA-templated transcription [GO:0006355]; regulation of transcription by RNA polymerase II [GO:0006357]; SMAD protein signal transduction [GO:0060395]; transforming growth factor beta receptor signaling pathway [GO:0007179] |
| 4943.89 | 144.95 | 78.77 | 2027.64 | 906.86 | 3036.99 | 11.06 | 268.74 | 0 | 5037.02 | 241.1 | 205.24 | 16695.28 | 48.19 | 76.96 | 817.57 | 221.73 | 47.02 | 34809.01 | Ubiquitin-conjugating enzyme E2Q-like protein 1 (EC 2.3.2.23) (E2Q-like ubiquitin-conjugating enzyme 1) | Ube2ql1 | protein ubiquitination [GO:0016567] |
| 70.19 | 9.78 | 18.39 | 162.01 | 2.73 | 14.52 | 23341.36 | 15.7 | 13.93 | 180.57 | 340.28 | 2.35 | 30.12 | 0 | 13.86 | 10134.84 | 283.66 | 16.6 | 34650.89 | NudC domain-containing protein 1 | nudcd1 zgc:110705 | |
| 8158.78 | 91.97 | 3538 | 397.93 | 77.08 | 5351.21 | 16.88 | 79.31 | 99.56 | 625.68 | 38.41 | 12419.92 | 3007.15 | 87.87 | 172.35 | 169.6 | 143.02 | 140.67 | 34615.39 | Sodium/potassium-transporting ATPase subunit alpha-1 (Na(+)/K(+) ATPase alpha-1 subunit) (EC 7.2.2.13) (Sodium pump subunit alpha-1) | atp1a1 | intracellular potassium ion homeostasis [GO:0030007]; intracellular sodium ion homeostasis [GO:0006883]; potassium ion import across plasma membrane [GO:1990573]; proton transmembrane transport [GO:1902600]; sodium ion export across plasma membrane [GO:0036376] |

| | | | | | | | | | | | | | | | | | | | | | |
|---|---|---|---|---|---|---|---|---|---|---|---|---|---|---|---|---|---|---|---|---|---|
| 105.56 | 103.1 | 31.45 | 247.58 | 233.16 | 148.33 | 2.31 | 109.39 | 0 | 331.42 | 180.35 | 18 | 31715.88 | 342.24 | 505.8 | 278.5 | 221.52 | 14.72 | 34589.31 | Zinc finger BED domain-containing protein DAYSLEEPER (Transposase-like protein DAYSLEEPER) | HAT At3g42170 T27B3.40 | post-embryonic development [GO:0009791] |
| 1890.73 | 1705.23 | 2532.85 | 320.18 | 1092.47 | 3715.74 | 468.87 | 383.61 | 5294.43 | 442.88 | 420.12 | 3304.81 | 889.49 | 1503.29 | 3907.53 | 401.36 | 550.28 | 5707.86 | 34531.73 | Glycerophosphocholine phosphodiesterase GPCPD1 (EC 3.1.4.2) (Glycerophosphodiester phosphodiesterase 5) | GPCPD1 GDE5 KIAA1434 | glycerophospholipid catabolic process [GO:0046475]; skeletal muscle tissue development [GO:0007519] |
| 11.93 | 19.77 | 34261.39 | 2.86 | 10.28 | 26.31 | 0.23 | 18.28 | 35.08 | 5.85 | 6.2 | 23.62 | 7.68 | 31.14 | 32.46 | 0.45 | 6.54 | 10.96 | 34511.03 | Next to BRCA1 gene 1 protein (Neighbor of BRCA1 gene 1 protein) | NBR1 | macroautophagy [GO:0016236] |
| 2360.54 | 4157.16 | 1403.05 | 709.58 | 3254.32 | 1888.59 | 1571.02 | 1842.17 | 779.66 | 704.08 | 1069.24 | 1765.09 | 3256.82 | 2483.35 | 1202.86 | 1477.48 | 3635.75 | 937.1 | 34497.86 | PDZ domain-containing protein 7 | PDZD7 PDZK7 | auditory receptor cell development [GO:0060117]; auditory receptor cell stereocilium organization [GO:0060088]; detection of mechanical stimulus involved in sensory perception of sound [GO:0050910]; establishment of protein localization [GO:0045184] |

| | | | | | | | | | | | | | | | | | | | | | |
|---|---|---|---|---|---|---|---|---|---|---|---|---|---|---|---|---|---|---|---|---|---|
| 183.96 | 171.36 | 364 | 472.42 | 133.41 | 213.77 | 15260.02 | 44.41 | 336.41 | 157.34 | 273.75 | 559.89 | 1076.14 | 735 | 1176.31 | 12386.34 | 510.33 | 405.98 | 34460.84 | Phosphoribosyl pyrophosphate synthase-associated protein 1 (PRPP synthase-associated protein 1) (39 kDa phosphoribosypyrophosphate synthase-associated protein) (PAP39) | PRPSAP1 | 5-phosphoribose 1-diphosphate biosynthetic process [GO:0006015]; nucleobase-containing compound metabolic process [GO:0006139]; purine nucleotide biosynthetic process [GO:0006164] |
| 2473.62 | 1308.27 | 3737.47 | 1132.99 | 1493.86 | 3489.67 | 37.68 | 852.13 | 4337.15 | 750.19 | 473.32 | 2033.66 | 621.06 | 1592.63 | 3578.92 | 118.09 | 994.21 | 5362.63 | 34387.55 | Neuronal membrane glycoprotein M6-a (M6a) | Gpm6a M6a | neural retina development [GO:0003407]; neuron migration [GO:0001764]; neuron projection morphogenesis [GO:0048812]; positive regulation of filopodium assembly [GO:0051491]; regulation of synapse organization [GO:0050807]; response to bacterium [GO:0009617]; stem cell differentiation [GO:0048863]; synapse assembly [GO:0007416] |
| 2.28 | 11206.72 | 19.09 | 1.29 | 5583.52 | 7.77 | 1.35 | 1660.81 | 7.01 | 11.27 | 2755.26 | 49.36 | 4.46 | 8872.24 | 9.02 | 6.81 | 4138.2 | 12.45 | 34348.91 | Protein DENND6A (DENN domain-containing protein 6A) | DENND6A FAM116A | positive regulation of cell-cell adhesion mediated by cadherin [GO:2000049] |

| | | | | | | | | | | | | | | | | | | | | | |
|---|---|---|---|---|---|---|---|---|---|---|---|---|---|---|---|---|---|---|---|---|---|
| 173.73 | 10161.19 | 203.54 | 21.46 | 4796.5 | 272.35 | 7.64 | 2369.11 | 54.27 | 68.37 | 788.18 | 158.51 | 93.81 | 13098.61 | 168.36 | 57.3 | 1414.87 | 361.06 | 34268.86 | CCN family member 2 (Cellular communication network factor 2) (Connective tissue growth factor) (Connective tissue growth-related protein) | Ccn2 Ctgf | angiogenesis [GO:0001525]; calcium ion transmembrane import into cytosol [GO:0097553]; cartilage condensation [GO:0001502]; cell adhesion [GO:0007155]; cell migration [GO:0016477]; cell-matrix adhesion [GO:0007160]; cellular response to fatty acid [GO:0071398]; cellular response to glucose stimulus [GO:0071333]; cellular response to interleukin-13 [GO:0035963]; chondrocyte differentiation [GO:0002062]; chondrocyte proliferation [GO:0035988]; connective tissue development [GO:0061448]; DNA biosynthetic process [GO:0071897]; extracellular matrix constituent secretion [GO:0070278]; fibroblast growth factor receptor signaling pathway [GO:0008543]; integrin-mediated signaling pathway [GO:0007229]; intracellular signal transduction [GO:0035556]; lung development [GO:0030324]; microtubule depolymerization [GO:0007019]; negative regulation of |
| 2017.88 | 3542.26 | 1284.56 | 716.1 | 3966.44 | 3672.21 | 26.05 | 2252.16 | 239.05 | 905.5 | 1400.75 | 828.83 | 3215.79 | 4351.08 | 1346.01 | 410.63 | 3101.55 | 863.44 | 34140.29 | Talin-1 | TLN1 TLN | cell-cell adhesion [GO:0098609] |
| 2.27 | 12195.2 | 0 | 0 | 2560.94 | 2.46 | 0 | 4668.7 | 0 | 0 | 2047.35 | 0.67 | 0 | 5285.45 | 0 | 0 | 7334.44 | 1.27 | 34098.75 | B-cell receptor CD22 (Sialic acid-binding Ig-like lectin 2) (Siglec-2) (CD antigen CD22) | CD22 SIGLEC2 | B cell activation [GO:0042113]; cell adhesion [GO:0007155]; negative regulation of B cell receptor signaling pathway [GO:0050859]; negative regulation of calcium-mediated signaling [GO:0050849]; negative regulation of immunoglobulin production [GO:0002638]; regulation of B cell proliferation [GO:0030888]; regulation of endocytosis [GO:0030100] |

| | | | | | | | | | | | | | | | | | | | | | |
|---|---|---|---|---|---|---|---|---|---|---|---|---|---|---|---|---|---|---|---|---|---|
| 0.19 | 6855.41 | 0.3 | 0 | 1836 | 8.44 | 0 | 1895.11 | 0 | 0.8 | 1575.56 | 1.35 | 0.79 | 16044.74 | 0 | 0 | 5813.89 | 22.07 | 34054.65 | Neural-cadherin (Cadherin-N) (dN-cadherin) | CadN CG7100 | axon extension [GO:0048675]; axon extension involved in axon guidance [GO:0048846]; axon guidance [GO:0007411]; axon target recognition [GO:0007412]; axonal fasciculation [GO:0007413]; calcium-dependent cell-cell adhesion via plasma membrane cell adhesion molecules [GO:0016339]; cell-cell adhesion [GO:0098609]; cell-cell adhesion mediated by cadherin [GO:0044331]; homophilic cell adhesion via plasma membrane adhesion molecules [GO:0007156]; negative regulation of dendrite morphogenesis [GO:0050774]; ommatidial rotation [GO:0016318]; R7 cell development [GO:0045467]; R8 cell development [GO:0045463]; regulation of axon extension involved in axon guidance [GO:0048841]; regulation of dendrite morphogenesis [GO:0048814]; retinal ganglion cell axon guidance [GO:0031290] |
| 2670.59 | 3700.98 | 1249.79 | 862.57 | 3155.24 | 1331.79 | 380.18 | 1468.16 | 1717.65 | 809.45 | 1111.99 | 1540.5 | 2548.87 | 4014.34 | 1287.61 | 867.08 | 3498.95 | 1637.26 | 33853 | Gamma-adducin (Adducin-like protein 70) (Protein kinase C-binding protein 35H) | Add3 | barbed-end actin filament capping [GO:0051016]; positive regulation of cytoskeleton organization [GO:0051495]; positive regulation of vasoconstriction [GO:0045907]; response to xenobiotic stimulus [GO:0009410] |
| 884.91 | 627.48 | 750.05 | 812.19 | 725.51 | 1049.65 | 11540.8 | 438.31 | 207.92 | 433.91 | 470.09 | 477.44 | 2645.91 | 1446.33 | 784.23 | 8903.97 | 940.18 | 677.63 | 33816.51 | G protein-coupled receptor kinase 3 (EC 2.7.11.15) (Beta-adrenergic receptor kinase 2) (Beta-ARK-2) | GRK3 ADRBK2 | desensitization of G protein-coupled receptor signaling pathway [GO:0002029]; G protein-coupled receptor signaling pathway [GO:0007186] |

| | | | | | | | | | | | | | | | | | | | | | |
|---|---|---|---|---|---|---|---|---|---|---|---|---|---|---|---|---|---|---|---|---|---|
| 6077.93 | 69.56 | 999.29 | 4091.7 | 275.68 | 11714.63 | 15.17 | 65.41 | 478.94 | 3383.77 | 162.38 | 2137.47 | 712.06 | 84.5 | 336.07 | 355.2 | 583.26 | 2258.21 | 33801.23 | Folliculin (BHD skin lesion fibrofolliculoma protein) (Birt-Hogg-Dube syndrome protein) | FLCN BHD | cell proliferation involved in kidney development [GO:0072111]; cell-cell junction assembly [GO:0007043]; cellular response to amino acid starvation [GO:0034198]; cellular response to starvation [GO:0009267]; energy homeostasis [GO:0097009]; epithelial cell proliferation [GO:0050673]; ERK1 and ERK2 cascade [GO:0070371]; hemopoiesis [GO:0030097]; in utero embryonic development [GO:0001701]; intracellular signal transduction [GO:0035556]; intrinsic apoptotic signaling pathway [GO:0097193]; lysosome localization [GO:0032418]; negative regulation of brown fat cell differentiation [GO:1903444]; negative regulation of cell proliferation involved in kidney development [GO:1901723]; negative regulation of cold-induced thermogenesis [GO:0120163]; negative regulation of epithelial cell proliferation [GO:0050680]; negative regulation of ERK1 and ERK2 cascade [GO:0070373]; negative regulation of glycolytic process |
| 2406.84 | 1219.7 | 4227.53 | 796.65 | 1070.82 | 2530.51 | 674.34 | 467.77 | 2948.32 | 781.52 | 540.23 | 2122.44 | 2318.72 | 1784.27 | 3602.14 | 991.38 | 1183.35 | 4114.72 | 33781.25 | Relaxin-3 (Insulin-like peptide INSL7) (Insulin-like peptide 7) (Prorelaxin R3) [Cleaved into: Relaxin-3 B chain;Relaxin-3 A chain] | Rln3 Insl7 | |
| 6.85 | 6203.02 | 4.3 | 10.22 | 6440.38 | 36.1 | 0.36 | 2817.35 | 0.94 | 7.59 | 2887.97 | 1.74 | 2.84 | 11069.18 | 2.6 | 2.04 | 4002.84 | 24.67 | 33520.99 | Sterile alpha motif domain-containing protein 9 (SAM domain-containing protein 9) | SAMD9 C7orf5 DRIF1 KIAA2004 OEF1 | endosomal vesicle fusion [GO:0034058]; innate immune response [GO:0045087] |

| | | | | | | | | | | | | | | | | | | | | | |
|---|---|---|---|---|---|---|---|---|---|---|---|---|---|---|---|---|---|---|---|---|---|
| 2128.17 | 1633.07 | 1811.26 | 880.91 | 1621.41 | 2616.16 | 4640.96 | 1074.58 | 552.78 | 940.79 | 825.89 | 1407.67 | 2150.58 | 1666.33 | 702.47 | 5431.84 | 2517.08 | 841.71 | 33443.66 | Conserved oligomeric Golgi complex subunit 3 (COG complex subunit 3) (Component of oligomeric Golgi complex 3) | Cog3 | endoplasmic reticulum to Golgi vesicle-mediated transport [GO:0006888]; intra-Golgi vesicle-mediated transport [GO:0006891]; intracellular protein transport [GO:0006886]; retrograde vesicle-mediated transport, Golgi to endoplasmic reticulum [GO:0006890] |
| 1252.03 | 824.68 | 475.52 | 495.45 | 954.1 | 962.57 | 12427.29 | 361.18 | 307.47 | 311.26 | 436.4 | 188.67 | 1738.17 | 2422.72 | 522.43 | 8235.32 | 1056.87 | 413.62 | 33385.75 | WD repeat and FYVE domain-containing protein 1 (WD40- and FYVE domain-containing protein 1) | Wdfy1 Kiaa1435 | positive regulation of toll-like receptor 3 signaling pathway [GO:0034141]; positive regulation of toll-like receptor 4 signaling pathway [GO:0034145] |
| 0 | 8742.07 | 7.51 | 0.96 | 7267.66 | 41.92 | 0.7 | 2791.43 | 0 | 0.54 | 2693.61 | 32.39 | 0.9 | 4716.85 | 0 | 0 | 6696.33 | 20.66 | 33013.53 | Nuclear body protein SP140 (Lymphoid-restricted homolog of Sp100) (LYSp100) (Nuclear autoantigen Sp-140) (Speckled 140 kDa) | SP140 LYSP100 | defense response [GO:0006952]; regulation of transcription by RNA polymerase II [GO:0006357] |

| | | | | | | | | | | | | | | | | | | | | | |
|---|---|---|---|---|---|---|---|---|---|---|---|---|---|---|---|---|---|---|---|---|---|
| 548.58 | 979.17 | 2355.16 | 76.86 | 575.43 | 1140.94 | 245.45 | 435.81 | 16283.59 | 322.39 | 268.43 | 4914.35 | 208.39 | 251.71 | 1285 | 132.51 | 192.15 | 2581.43 | 32797.35 | Polyadenylate-binding protein 2 (PABP-2) (Poly(A)-binding protein 2) (Nuclear poly(A)-binding protein 1) (Poly(A)-binding protein II) (PABII) (Polyadenylate-binding nuclear protein 1) | Pabpn1 Pab2 Pabp2 | cellular response to lipopolysaccharide [GO:0071222]; MAPK cascade [GO:0000165]; mRNA processing [GO:0006397]; positive regulation of polynucleotide adenylyltransferase activity [GO:1904247]; regulation of mRNA 3'-end processing [GO:0031440] |
| 114.89 | 620.7 | 1357.34 | 17.14 | 431.82 | 714.44 | 4.28 | 218.38 | 9273.26 | 54.76 | 214.03 | 2870.62 | 51.19 | 1299.66 | 7448.19 | 205.75 | 784.84 | 7066.98 | 32748.27 | Serine/threonine-protein kinase 10 (EC 2.7.11.1) | stk10 zgc:63495 | intracellular signal transduction [GO:0035556]; protein autophosphorylation [GO:0046777]; regulation of lymphocyte migration [GO:2000401] |
| 1677.35 | 1543.87 | 195.65 | 63.64 | 189.06 | 309.96 | 7080.32 | 308.37 | 2360.37 | 1237.12 | 810.24 | 2136.47 | 3451.02 | 9341.51 | 1652.64 | 34.47 | 154.95 | 83.14 | 32630.15 | TBC1 domain family member 17 | TBC1D17 | autophagy [GO:0006914]; protein transport [GO:0015031]; retrograde transport, endosome to Golgi [GO:0042147] |
| 1217.93 | 5928.39 | 608 | 443.71 | 4155.53 | 1760.65 | 808.29 | 2253.43 | 1071.04 | 359.86 | 1366.16 | 1278.95 | 1403.73 | 2858.83 | 1542.09 | 730.95 | 3217.1 | 1562.68 | 32567.32 | Amphiphysin | AMPH | synaptic vesicle endocytosis [GO:0048488] |

| | | | | | | | | | | | | | | | | | | | | | |
|---|---|---|---|---|---|---|---|---|---|---|---|---|---|---|---|---|---|---|---|---|---|
| 105.29 | 139.54 | 143.27 | 49.66 | 167.82 | 212.44 | 17283.63 | 60.19 | 111.43 | 130.47 | 309.07 | 72.45 | 110.74 | 167.52 | 172.34 | 12862.86 | 324.54 | 142.7 | 32565.96 | Protein dispatched homolog 1 (Protein chameleon) | disp1 con | adenohypophysis development [GO:0021984]; animal organ morphogenesis [GO:0009887]; anterior/posterior pattern specification [GO:0009952]; blood circulation [GO:0008015]; embryonic cranial skeleton morphogenesis [GO:0048701]; inner ear development [GO:0048839]; muscle cell fate specification [GO:0042694]; retinal ganglion cell axon guidance [GO:0031290]; skeletal muscle tissue development [GO:0007519]; smoothened signaling pathway [GO:0007224] |
| 1183.48 | 792.36 | 1003.68 | 792.71 | 663.85 | 1665.87 | 9544.96 | 268.65 | 1022.27 | 517.2 | 388.8 | 944.89 | 2080.27 | 1065.42 | 1096.55 | 7479.42 | 774.32 | 1181.75 | 32466.45 | Neuronal growth regulator 1 (IgLON family member 4) | NEGR1 IGLON4 UNQ2433/PRO 4993 | brain development [GO:0007420]; cell-cell adhesion [GO:0098609]; cholesterol homeostasis [GO:0042632]; fat cell differentiation [GO:0045444]; feeding behavior [GO:0007631]; lipid droplet formation [GO:0140042]; locomotory behavior [GO:0007626]; neuron projection morphogenesis [GO:0048812]; positive regulation of calcium-mediated signaling [GO:0050850]; positive regulation of neuron projection development [GO:0010976]; positive regulation of saliva secretion [GO:0046878]; regulation of synapse assembly [GO:0051963]; skeletal muscle organ development [GO:0060538]; social behavior [GO:0035176] |

| | | | | | | | | | | | | | | | | | | | | | |
|---|---|---|---|---|---|---|---|---|---|---|---|---|---|---|---|---|---|---|---|---|---|
| 3994.56 | 1985.28 | 2068.19 | 2071.72 | 1330.87 | 2566.8 | 402.02 | 4282.18 | 4489.01 | 3056.09 | 878.95 | 564.36 | 920.97 | 1131.36 | 388.98 | 337 | 1167.97 | 645.21 | 32281.52 | Period circadian protein homolog 2 (cPER2) (Circadian clock protein PERIOD 2) | PER2 | circadian regulation of gene expression [GO:0032922]; circadian rhythm [GO:0007623]; entrainment of circadian clock by photoperiod [GO:0043153]; negative regulation of transcription by RNA polymerase II [GO:0000122] |
| 2374.31 | 1822.14 | 3728.44 | 672.64 | 1679.08 | 2487.12 | 1271.84 | 862.98 | 2879.39 | 772.25 | 953.88 | 2119.25 | 1405.84 | 2016.28 | 1528.44 | 1154.89 | 2253.55 | 2229.41 | 32211.73 | Growth arrest-specific protein 6 (GAS-6) (AXL receptor tyrosine kinase ligand) | GAS6 AXLLG | activation of protein kinase B activity [GO:0032148]; animal organ regeneration [GO:0031100]; apoptotic cell clearance [GO:0043277]; B cell chemotaxis [GO:0035754]; blood coagulation [GO:0007596]; calcium ion transmembrane transport [GO:0070588]; cell surface receptor signaling pathway [GO:0007166]; cell-substrate adhesion [GO:0031589]; cellular response to glucose stimulus [GO:0071333]; cellular response to growth factor stimulus [GO:0071363]; cellular response to interferon-alpha [GO:0035457]; cellular response to starvation [GO:0009267]; cellular response to vitamin K [GO:0071307]; cellular response to xenobiotic stimulus [GO:0071466]; dendritic cell differentiation [GO:0097028]; enzyme-linked receptor protein signaling pathway [GO:0007167]; extracellular matrix assembly [GO:0085029]; fibroblast apoptotic process [GO:0044346]; fusion of virus membrane with host plasma membrane [illegible] |

| | | | | | | | | | | | | | | | | | | | | | |
|---|---|---|---|---|---|---|---|---|---|---|---|---|---|---|---|---|---|---|---|---|---|
| 787.26 | 3792.89 | 1091 | 383.62 | 4084.01 | 1592.34 | 93.08 | 2215.07 | 574.66 | 300.42 | 1374.2 | 946.74 | 1818.3 | 7062.78 | 1367.26 | 335.82 | 3101.56 | 1196 | 32117.01 | Junction-mediating and -regulatory protein | jmy zgc:77377 | 'de novo' actin filament nucleation [GO:0070060]; actin polymerization-dependent cell motility [GO:0070358]; Arp2/3 complex-mediated actin nucleation [GO:0034314]; cellular response to starvation [GO:0009267]; DNA repair [GO:0006281]; intrinsic apoptotic signaling pathway by p53 class mediator [GO:0072332]; positive regulation of apoptotic process [GO:0043065] |
| 1109.44 | 586.01 | 10536.25 | 309.57 | 500.02 | 8145.07 | 27.9 | 177.18 | 232.8 | 346.35 | 238.54 | 4592.41 | 2250.87 | 355.73 | 1616.63 | 370.96 | 402.3 | 269.65 | 32067.68 | Arf-GAP with GTPase, ANK repeat and PH domain-containing protein 1 (AGAP-1) (Centaurin-gamma-2) (Cnt-g2) (GTP-binding and GTPase-activating protein 1) (GGAP1) | AGAP1 CENTG2 KIAA1099 | protein transport [GO:0015031]; regulation of modification of postsynaptic structure [GO:0099159] |

| | | | | | | | | | | | | | | | | | | | | | |
|---|---|---|---|---|---|---|---|---|---|---|---|---|---|---|---|---|---|---|---|---|---|
| 0 | 2.04 | 7722.95 | 5.65 | 0 | 738.36 | 0 | 0 | 38.46 | 0 | 0 | 23385.57 | 0 | 1.23 | 35.16 | 0 | 0 | 43.64 | 31973.06 | Glutathione-specific gamma-glutamylcyclotransferase 1 (Gamma-GCG 1) (EC 4.3.2.7) (Cation transport regulator-like protein 1) | chac1 | apoptotic process [GO:0006915]; glutathione catabolic process [GO:0006751]; negative regulation of Notch signaling pathway [GO:0045746]; negative regulation of protein processing [GO:0010955]; neurogenesis [GO:0022008]; Notch signaling pathway [GO:0007219]; response to unfolded protein [GO:0006986] |
| 1424.56 | 1361.49 | 2921.47 | 495.75 | 1485.47 | 1912.75 | 78.11 | 1019.79 | 5016.59 | 920.75 | 821.13 | 2508.97 | 1986.69 | 2406.33 | 2196.5 | 518.17 | 1572.46 | 3277.94 | 31924.92 | Galectin-3 (Gal-3) (35 kDa lectin) (Carbohydrate-binding protein 35) (CBP 35) (Galactose-specific lectin 3) (IgE-binding protein) (L-34 galactoside-binding lectin) (Laminin-binding protein) (Lectin L-29) (Mac-2 antigen) | Lgals3 | cell differentiation [GO:0030154]; extracellular matrix organization [GO:0030198]; innate immune response [GO:0045087]; mRNA processing [GO:0006397]; negative regulation of endocytosis [GO:0045806]; negative regulation of immunological synapse formation [GO:2000521]; negative regulation of T cell activation via T cell receptor contact with antigen bound to MHC molecule on antigen presenting cell [GO:2001189]; negative regulation of T cell receptor signaling pathway [GO:0050860]; RNA splicing [GO:0008380]; skeletal system development [GO:0001501] |

| | | | | | | | | | | | | | | | | | | | | | |
|---|---|---|---|---|---|---|---|---|---|---|---|---|---|---|---|---|---|---|---|---|---|
| 1454.91 | 1814.38 | 2575.82 | 703.4 | 2158.05 | 3560.02 | 110.53 | 1267.1 | 647.89 | 694.06 | 860.52 | 2322.49 | 2722.74 | 3275.58 | 2889.2 | 347.58 | 1621.98 | 2855.9 | 31882.15 | Transmembrane 9 superfamily member 2 | TM9SF2 | ceramide metabolic process [GO:0006672]; glycosphingolipid biosynthetic process [GO:0006688]; protein localization to membrane [GO:0072657]; regulation of heparan sulfate proteoglycan biosynthetic process [GO:0010908] |
| 1625.71 | 1752.78 | 1589.1 | 1195.34 | 1703.72 | 2900.73 | 2974.44 | 693.79 | 764.53 | 947.23 | 1005.83 | 2015.77 | 2453.42 | 3059.36 | 1048.09 | 3453.49 | 1723.46 | 912.6 | 31819.39 | Retinoschisin (X-linked juvenile retinoschisis protein homolog) | xlrs1 | cell adhesion [GO:0007155]; visual perception [GO:0007601] |
| 3.82 | 2048.62 | 27.06 | 11.35 | 4102.14 | 70.24 | 251.48 | 1707.29 | 10.2 | 8.06 | 3026.84 | 21.33 | 6.43 | 16917.82 | 8.05 | 109.97 | 3231.29 | 30.11 | 31592.1 | Insulin-like growth factor 2 mRNA-binding protein 2 (IGF2 mRNA-binding protein 2) (IMP-2) (Hepatocellular carcinoma autoantigen p62) (IGF-II mRNA-binding protein 2) (VICKZ family member 2) | IGF2BP2 IMP2 VICKZ2 | anatomical structure morphogenesis [GO:0009653]; cold-induced thermogenesis [GO:0106106]; CRD-mediated mRNA stabilization [GO:0070934]; energy homeostasis [GO:0097009]; mRNA transport [GO:0051028]; negative regulation of translation [GO:0017148]; nervous system development [GO:0007399]; regulation of cytokine production [GO:0001817]; RNA stabilization [GO:0043489] |

| | | | | | | | | | | | | | | | | | | | | | |
|---|---|---|---|---|---|---|---|---|---|---|---|---|---|---|---|---|---|---|---|---|---|
| 24.48 | 0 | 54.34 | 0.73 | 0 | 0.78 | 16039.35 | 0.25 | 12.85 | 68.43 | 321.74 | 31.05 | 10.77 | 9.37 | 429.71 | 14263.18 | 238.71 | 11.85 | 31517.59 | Caspase-6 (CASP-6) (EC 3.4.22.59) [Cleaved into: Caspase-6 subunit p18;Caspase-6 subunit p11] | CASP6 | activation of innate immune response [GO:0002218]; apoptotic chromosome condensation [GO:0030263]; apoptotic DNA fragmentation [GO:0006309]; apoptotic nuclear changes [GO:0030262]; apoptotic process [GO:0006915]; hepatocyte apoptotic process [GO:0097284]; intrinsic apoptotic signaling pathway by p53 class mediator [GO:0072332]; lens fiber cell differentiation [GO:0070306]; positive regulation of apoptotic process [GO:0043065]; positive regulation of necroptotic process [GO:0060545]; positive regulation of neuron apoptotic process [GO:0043525]; protein autoprocessing [GO:0016540]; proteolysis [GO:0006508]; pyroptotic inflammatory response [GO:0070269] |
| 1327.04 | 1479.81 | 1778.32 | 1387.06 | 1751.62 | 2174.67 | 5046.65 | 635.84 | 896.98 | 662.94 | 882.59 | 1387.96 | 2009.2 | 3052.21 | 888.1 | 3733.33 | 1470.81 | 772.87 | 31338 | Inhibin beta B chain (Activin beta-B chain) | INHBB | activin receptor signaling pathway [GO:0032924]; animal organ development [GO:0048513]; cellular response to insulin stimulus [GO:0032869]; cellular response to starvation [GO:0009267]; fat cell differentiation [GO:0045444]; negative regulation of follicle-stimulating hormone secretion [GO:0046882]; negative regulation of hepatocyte growth factor production [GO:0032686]; negative regulation of insulin secretion [GO:0046676]; positive regulation of follicle-stimulating hormone secretion [GO:0046881]; positive regulation of ovulation [GO:0060279]; response to wounding [GO:0009611] |

| | | | | | | | | | | | | | | | | | | | | | |
|---|---|---|---|---|---|---|---|---|---|---|---|---|---|---|---|---|---|---|---|---|---|
| 0.46 | 0 | 31219.36 | 0 | 0 | 0 | 23.79 | 0 | 3.98 | 0 | 1.16 | 2.71 | 0 | 0 | 3.41 | 32.82 | 0 | 2.23 | 31289.92 | 5-aminolevulinate synthase, non-specific, mitochondrial (ALAS-H) (EC 2.3.1.37) (5-aminolevulinic acid synthase 1) (Delta-ALA synthase 1) (Delta-aminolevulinate synthase 1) | alas1 | erythrocyte development [GO:0048821]; hemoglobin biosynthetic process [GO:0042541]; protoporphyrinogen IX biosynthetic process [GO:0006782]; response to bile acid [GO:1903412] |
| 1667.3 | 3349.8 | 1562.52 | 241.23 | 2156.62 | 1369.34 | 16.07 | 1714.33 | 3411.66 | 541.86 | 1137.66 | 1570.33 | 1501.08 | 3271.61 | 1887.39 | 331.22 | 2438.72 | 3098.89 | 31267.63 | Protein DOP1A | DOP1A DOPEY1 KIAA1117 | Golgi to endosome transport [GO:0006895]; protein transport [GO:0015031] |
| 1686.86 | 1720.26 | 1446.57 | 582.88 | 2838.65 | 4615.52 | 22.43 | 1488.25 | 148.83 | 1032.88 | 945.94 | 1270.97 | 3812.43 | 3654.7 | 2295.05 | 421.16 | 2197.57 | 1036.16 | 31217.11 | E3 ubiquitin-protein ligase UBR4 (EC 2.3.2.27) (600 kDa retinoblastoma protein-associated factor) (p600) (N-recognin-4) (Retinoblastoma-associated factor of 600 kDa) (RBAF600) | UBR4 KIAA0462 KIAA1307 RBAF600 | cytoplasm protein quality control [GO:0140455]; cytoplasm protein quality control by the ubiquitin-proteasome system [GO:0071629]; endosome organization [GO:0007032]; negative regulation of fatty acid biosynthetic process [GO:0045717]; negative regulation of HRI-mediated signaling [GO:0141191]; positive regulation of autophagy [GO:0010508]; proteasome-mediated ubiquitin-dependent protein catabolic process [GO:0043161]; protein branched polyubiquitination [GO:0141198]; protein K11-linked ubiquitination [GO:0070979]; protein K27-linked ubiquitination [GO:0044314]; protein K48-linked ubiquitination [GO:0070936]; protein stabilization [GO:0050821]; response to oxidative stress [GO:0006979]; ubiquitin-dependent protein catabolic process [GO:0006511]; ubiquitin-dependent protein catabolic process via the N-end rule pathway [GO:0071596] |

| | | | | | | | | | | | | | | | | | | | | | |
|---|---|---|---|---|---|---|---|---|---|---|---|---|---|---|---|---|---|---|---|---|---|
| 3346.56 | 3059.99 | 2104.71 | 727.43 | 3179.92 | 2801.09 | 36.12 | 1519.06 | 582.81 | 839.97 | 993.67 | 1541.45 | 2796.85 | 3281.8 | 1162.92 | 262.63 | 1906.91 | 1065.11 | 31209 | Nucleoprotein TPR (Megator) (NPC-associated intranuclear protein) (Translocated promoter region protein) | TPR | cell division [GO:0051301]; cellular response to heat [GO:0034605]; cellular response to interferon-alpha [GO:0035457]; mitotic spindle assembly checkpoint signaling [GO:0007094]; mRNA export from nucleus [GO:0006406]; mRNA export from nucleus in response to heat stress [GO:0031990]; negative regulation of RNA export from nucleus [GO:0046832]; negative regulation of transcription by RNA polymerase II [GO:0000122]; negative regulation of translational initiation [GO:0045947]; nuclear pore organization [GO:0006999]; nucleocytoplasmic transport [GO:0006913]; positive regulation of heterochromatin formation [GO:0031453]; positive regulation of intracellular protein transport [GO:0090316]; positive regulation of mitotic cell cycle spindle assembly checkpoint [GO:0090267]; positive regulation of protein export from nucleus [GO:0046827]; positive regulation of protein import into nucleus |
| 1722.08 | 890.34 | 731.11 | 791.29 | 1250.83 | 2551.68 | 8938.06 | 642.75 | 408.08 | 1078.63 | 923.37 | 1061.23 | 1821.17 | 1043.78 | 608.11 | 5016.66 | 1128.09 | 550.75 | 31158.01 | Gamma-aminobutyric acid receptor subunit pi (GABA(A) receptor subunit pi) (GABAAR subunit pi) | Gabrp | |
| 1187.18 | 1612.2 | 5116.35 | 844.01 | 1335.04 | 4710.22 | 55.31 | 421.89 | 195.05 | 1356.34 | 1192.06 | 8107.07 | 709.1 | 1364.03 | 545.4 | 772.35 | 1275.01 | 253.3 | 31051.91 | von Willebrand factor D and EGF domain-containing protein | VWDE | |

| | | | | | | | | | | | | | | | | | | | | | |
|---|---|---|---|---|---|---|---|---|---|---|---|---|---|---|---|---|---|---|---|---|---|
| 1024.98 | 1108.5 | 476.04 | 594.4 | 1010.44 | 1921.45 | 11909.32 | 465.75 | 388.75 | 511.09 | 625.8 | 452.17 | 916.47 | 1194.66 | 574.78 | 6924.01 | 556.14 | 266.65 | 30921.4 | cAMP-dependent protein kinase catalytic subunit beta (PKA C-beta) (EC 2.7.11.11) | PRKACB | adenylate cyclase-modulating G protein-coupled receptor signaling pathway [GO:0007188]; high-density lipoprotein particle assembly [GO:0034380]; negative regulation of smoothened signaling pathway [GO:0045879]; negative regulation of TORC1 signaling [GO:1904262]; neural tube closure [GO:0001843]; protein phosphorylation [GO:0006468]; regulation of protein processing [GO:0070613]; renal water homeostasis [GO:0003091]; signal transduction [GO:0007165]; vascular endothelial cell response to laminar fluid shear stress [GO:0097700] |
| 14.67 | 2.52 | 0 | 207.37 | 0 | 21.59 | 18869.62 | 0 | 0 | 172.7 | 542.79 | 42 | 0 | 0 | 5.74 | 10413.93 | 392.77 | 18.91 | 30704.61 | Solute carrier family 12 member 2 (Bumetanide-sensitive sodium-(potassium)-chloride cotransporter 2) (BSC2) (Na-K-Cl cotransporter 1) (NKCC1) | slc12a2 nkcc1 | ammonium transmembrane transport [GO:0072488]; cell volume homeostasis [GO:0006884]; chloride ion homeostasis [GO:0055064]; chloride transmembrane transport [GO:1902476]; ear development [GO:0043583]; inner ear morphogenesis [GO:0042472]; potassium ion homeostasis [GO:0055075]; potassium ion import across plasma membrane [GO:1990573]; sodium ion homeostasis [GO:0055078]; sodium ion transmembrane transport [GO:0035725]; swim bladder inflation [GO:0048798] |

| | | | | | | | | | | | | | | | | | | | | | |
|---|---|---|---|---|---|---|---|---|---|---|---|---|---|---|---|---|---|---|---|---|---|
| 1940.34 | 1787.56 | 6628.43 | 326.17 | 1506.66 | 2329.3 | 177.09 | 391.78 | 5256.29 | 973.14 | 455.47 | 4333.39 | 519.08 | 95.97 | 1072.51 | 383.38 | 434.79 | 1833.11 | 30444.46 | Rho GTPase-activating protein 7 (Deleted in liver cancer 1 protein) (DLC-1) (HP protein) (Rho-type GTPase-activating protein 7) (START domain-containing protein 12) (StARD12) (StAR-related lipid transfer protein 12) | DLC1 ARHGAP7 KIAA1723 STARD12 | actin cytoskeleton organization [GO:0030036]; focal adhesion assembly [GO:0048041]; forebrain development [GO:0030900]; heart morphogenesis [GO:0003007]; hindbrain morphogenesis [GO:0021575]; intracellular signal transduction [GO:0035556]; negative regulation of cell migration [GO:0030336]; negative regulation of cell population proliferation [GO:0008285]; negative regulation of focal adhesion assembly [GO:0051895]; negative regulation of Rho protein signal transduction [GO:0035024]; negative regulation of stress fiber assembly [GO:0051497]; neural tube closure [GO:0001843]; positive regulation of execution phase of apoptosis [GO:1900119]; regulation of Rho protein signal transduction [GO:0035023]; regulation of small GTPase mediated signal transduction [GO:0051056] |
| 646.92 | 1369.43 | 361.3 | 1186.05 | 4076.55 | 3476.17 | 1315.8 | 1176.03 | 870.09 | 303.51 | 601.19 | 1201.78 | 2439.14 | 4390.81 | 1838.69 | 1656.97 | 2170.54 | 1269.71 | 30350.68 | Hexokinase-1 (EC 2.7.1.1) (Brain form hexokinase) (Hexokinase type I) (HK I) (Hexokinase-A) | HK1 | canonical glycolysis [GO:0061621]; carbohydrate phosphorylation [GO:0046835]; establishment of protein localization to mitochondrion [GO:0072655]; fructose 6-phosphate metabolic process [GO:0006002]; GDP-mannose biosynthetic process [GO:0009298]; GDP-mannose biosynthetic process from mannose [GO:0061728]; glucose 6-phosphate metabolic process [GO:0051156]; glucose metabolic process [GO:0006006]; glycolytic process [GO:0006096]; inflammatory response [GO:0006954]; innate immune response [GO:0045087]; intracellular glucose homeostasis [GO:0001678]; maintenance of protein location in mitochondrion [GO:0072656]; mannose metabolic process [GO:0006013]; positive regulation of cytokine production involved in immune response [GO:0002720]; positive regulation of interleukin-1 beta production [GO:0032731] |
| 3802.45 | 369.01 | 5908.25 | 1070.41 | 400.42 | 1502.04 | 436.46 | 136.75 | 3535.83 | 681.5 | 151.86 | 1558.66 | 2409.69 | 858.82 | 2293.7 | 642.8 | 954.39 | 3626.03 | 30339.07 | Cytochrome P450 20A1 (EC 1.14.-.-) | CYP20A1 UNQ667/PRO1301 | |
| 3099.03 | 583.99 | 373.05 | 1793.3 | 2044.26 | 2349.8 | 389.88 | 1797.55 | 793.35 | 1204.77 | 544 | 996.76 | 6301.83 | 2092.81 | 2041.08 | 587.16 | 1166.36 | 2102.77 | 30261.75 | Uncharacterized protein 045L | FV3-045L | |

| | | | | | | | | | | | | | | | | | | | | | |
|---|---|---|---|---|---|---|---|---|---|---|---|---|---|---|---|---|---|---|---|---|---|
| 1930.76 | 2598.79 | 2839.02 | 821.83 | 1746.6 | 5029.99 | 65.46 | 629.44 | 1183.57 | 723.72 | 501.59 | 3688.73 | 1344.79 | 2331.55 | 1168.01 | 534.66 | 1537.38 | 1469.03 | 30144.92 | Serine/threonine-protein phosphatase PP1-beta catalytic subunit (PP-1B) (EC 3.1.3.16) | ppp1cb TEgg049h05.1 | cell division [GO:0051301]; glycogen metabolic process [GO:0005977]; regulation of cell adhesion [GO:0030155] |
| 320.56 | 520.69 | 918.56 | 130.37 | 653.46 | 325.55 | 102.13 | 636.7 | 5.78 | 241.69 | 937.15 | 20003.77 | 737.71 | 2814.29 | 35.72 | 404.82 | 1255.62 | 40.39 | 30084.96 | DENN domain-containing protein 5B (Rab6IP1-like protein) | dennd5b si:ch211-11c20.2 zgc:77218 | |
| 1133.38 | 1597.39 | 3091.19 | 344.34 | 1730.36 | 2021.96 | 5660.31 | 800.1 | 1018.94 | 348.09 | 628.68 | 2171.29 | 959.33 | 1969.52 | 1297.71 | 3041.49 | 1093.49 | 1078.55 | 29986.12 | Protein FAM193A (Protein IT14) | FAM193A C4orf8 RES4-22 | |
| 0 | 0 | 29937.28 | 0 | 0 | 0 | 0 | 0 | 0 | 0 | 0.59 | 2.77 | 0 | 1.22 | 3.47 | 0 | 5.14 | 3.68 | 29954.15 | Rieske domain-containing protein | Rfesd | |
| 1194.78 | 1714.34 | 2027.06 | 561.31 | 1213.39 | 1393.5 | 1834.36 | 617.01 | 3133.98 | 401.63 | 584.44 | 2286.89 | 1416.16 | 2441.89 | 3318.9 | 1642.52 | 1183.42 | 2980.58 | 29946.16 | Protein jagged-1 (Jagged1) (hJ1) (CD antigen CD339) | JAG1 JAGL1 | angiogenesis [GO:0001525]; aorta morphogenesis [GO:0035909]; aortic valve morphogenesis [GO:0003180]; blood vessel remodeling [GO:0001974]; cardiac neural crest cell development involved in outflow tract morphogenesis [GO:0061309]; cardiac right ventricle morphogenesis [GO:0003215]; cardiac septum morphogenesis [GO:0060411]; cell fate determination [GO:0001709]; ciliary body morphogenesis [GO:0061073]; distal tubule development [GO:0072017]; endocardial cushion cell development [GO:0061444]; endothelial cell differentiation [GO:0045446]; hemopoiesis [GO:0030097]; inhibition of neuroepithelial cell differentiation [GO:0002085]; inner ear auditory receptor cell differentiation [GO:0042491]; keratinocyte differentiation [GO:0030216]; loop of Henle development [GO:0072070]; morphogenesis of an epithelial sheet [GO:0002011]; myoblast differentiation [GO:0045445]; negative regulation of |
| 145.27 | 5687.75 | 603.69 | 429.25 | 3467.46 | 160.08 | 3.08 | 2390.63 | 3560.96 | 224.57 | 1182.85 | 2170.22 | 61.86 | 4033.34 | 1764.38 | 56.32 | 1449.64 | 2522.62 | 29913.97 | Cerebellar degeneration-related protein 2 | CDR2 RCJMB04_19l3 | |

| | | | | | | | | | | | | | | | | | | | | | |
|---|---|---|---|---|---|---|---|---|---|---|---|---|---|---|---|---|---|---|---|---|---|
| 7225.13 | 69.49 | 33.13 | 1357.44 | 129.64 | 354.83 | 24.85 | 11.45 | 0.83 | 6161.85 | 104.55 | 275.1 | 13492.52 | 26.24 | 29.09 | 403.77 | 54.09 | 6.28 | 29760.28 | Transcription initiation factor TFIID subunit 4 (RNA polymerase II TBP-associated factor subunit C) (TBP-associated factor 4) (Transcription initiation factor TFIID 130 kDa subunit) (TAF(II)130) (TAFII-130) (TAFII130) (Transcription initiation factor TFIID 135 kDa subunit) (TAF(II)135) (TAFII-135) (TAFII135) | TAF4 TAF2C TAF2C1 TAF4A TAFII130 TAFII135 | DNA-templated transcription initiation [GO:0006352]; mRNA transcription by RNA polymerase II [GO:0042789]; ovarian follicle development [GO:0001541]; positive regulation of DNA-templated transcription [GO:0045893]; positive regulation of transcription initiation by RNA polymerase II [GO:0060261]; regulation of DNA repair [GO:0006282]; regulation of transcription by RNA polymerase II [GO:0006357]; RNA polymerase II preinitiation complex assembly [GO:0051123]; transcription by RNA polymerase II [GO:0006366]; transcription initiation at RNA polymerase II promoter [GO:0006367] |
| 0 | 0 | 0 | 0 | 0 | 29716.85 | 5.83 | 0 | 11.18 | 0 | 0 | 0 | 0 | 4.42 | 0 | 8.93 | 6.49 | 6.36 | 29760.06 | Procollagen-lysine,2-oxoglutarate 5-dioxygenase 1 (EC 1.14.11.4) (Lysyl hydroxylase 1) (LH1) | Plod1 Plod | cellular response to hormone stimulus [GO:0032870]; collagen fibril organization [GO:0030199]; epidermis development [GO:0008544]; peptidyl-lysine hydroxylation [GO:0017185]; response to hypoxia [GO:0001666] |

| | | | | | | | | | | | | | | | | | | | | | |
|---|---|---|---|---|---|---|---|---|---|---|---|---|---|---|---|---|---|---|---|---|---|
| 5211.44 | 43.22 | 57.09 | 1099.62 | 215.41 | 996.1 | 14.82 | 22.21 | 5.59 | 3902.45 | 161.7 | 216.51 | 17279.49 | 41.74 | 66.02 | 304.49 | 57.01 | 2.09 | 29697 | E3 ubiquitin-protein ligase pellino homolog 1 (Pellino-1) (EC 2.3.2.27) (Pellino-related intracellular-signaling molecule) (RING-type E3 ubiquitin transferase pellino homolog 1) | PELI1 PRISM | DNA repair [GO:0006281]; negative regulation of necroptotic process [GO:0060546]; negative regulation of T cell proliferation [GO:0042130]; negative regulation of TORC1 signaling [GO:1904262]; positive regulation of B cell proliferation [GO:0030890]; positive regulation of canonical NF-kappaB signal transduction [GO:0043123]; positive regulation of cytokine production [GO:0001819]; positive regulation of double-strand break repair via homologous recombination [GO:1905168]; positive regulation of protein ubiquitination [GO:0031398]; positive regulation of toll-like receptor 3 signaling pathway [GO:0034141]; positive regulation of toll-like receptor 4 signaling pathway [GO:0034145]; proteasome-mediated ubiquitin-dependent protein catabolic process [GO:0043161]; protein K48-linked ubiquitination [GO:0070936]; protein K63-linked ubiquitination [GO:0070534]; regulation of necroptotic process [GO:0060544]; regulation of Toll [illegible] |
| 33.39 | 6268.79 | 355.37 | 28.84 | 5752.32 | 648.4 | 248.23 | 3251.16 | 59.92 | 29.17 | 2514.06 | 192.81 | 4.13 | 7263.67 | 282.7 | 87.74 | 2544.57 | 116.69 | 29681.96 | Transmembrane channel-like protein 6 | Tmc6 Ever1 | intracellular zinc ion homeostasis [GO:0006882]; protein stabilization [GO:0050821] |
| 2231.65 | 1524.98 | 3171.14 | 759.42 | 1847.53 | 5302.43 | 120.08 | 373.26 | 758.6 | 585.61 | 596.36 | 1771.3 | 3066.88 | 1865.23 | 2061.55 | 508.87 | 1166.25 | 1957.75 | 29668.89 | E3 ubiquitin-protein ligase RNF114 (EC 2.3.2.27) (RING finger protein 114) (RING-type E3 ubiquitin transferase RNF114) (Zinc finger protein 313) | Rnf114 Zfp313 Znf313 | cell differentiation [GO:0030154]; protein polyubiquitination [GO:0000209]; spermatogenesis [GO:0007283]; ubiquitin-dependent protein catabolic process [GO:0006511] |

| | | | | | | | | | | | | | | | | | | | | | |
|---|---|---|---|---|---|---|---|---|---|---|---|---|---|---|---|---|---|---|---|---|---|
| 0.95 | 11730.72 | 25.96 | 1.83 | 1304.78 | 25.46 | 1.21 | 2881.45 | 8.26 | 0.51 | 1585.32 | 113.1 | 7.98 | 2855.33 | 8.09 | 0.67 | 9046.78 | 29.61 | 29628.01 | Solute carrier family 25 member 47-A (Hepatocellular carcinoma down-regulated mitochondrial carrier homolog A) | slc25a47a hdmcpa zgc:136752 | |
| 503.17 | 1078.66 | 7006.96 | 212.08 | 874.69 | 2424.7 | 1008.01 | 583.6 | 2253.99 | 161.08 | 387.25 | 4798.87 | 704.63 | 972.24 | 2309.94 | 698.02 | 964.54 | 2640.91 | 29583.34 | Phosphofurin acidic cluster sorting protein 1 (PACS-1) | PACS1 KIAA1175 | lymphocyte homeostasis [GO:0002260]; protein localization to Golgi apparatus [GO:0034067]; protein localization to plasma membrane [GO:0072659] |
| 1402.11 | 1504.15 | 965.8 | 883.44 | 1688.08 | 2929.19 | 6279.43 | 603.8 | 399.84 | 666.08 | 595.07 | 1058.98 | 2026.61 | 1056.17 | 975.46 | 4660.26 | 929.04 | 885.55 | 29509.06 | F-box only protein 50 (NCC receptor protein 1) (NCCRP-1) (Non-specific cytotoxic cell receptor protein 1) | nccrp1 si:dkey-85K8.6 wu:fa93e02 wu:fb12f07 zgc:122987 | ERAD pathway [GO:0036503]; glycoprotein catabolic process [GO:0006516]; SCF-dependent proteasomal ubiquitin-dependent protein catabolic process [GO:0031146] |

| | | | | | | | | | | | | | | | | | | | | | | |
|---|---|---|---|---|---|---|---|---|---|---|---|---|---|---|---|---|---|---|---|---|---|---|
| 1443.78 | | 5244.74 | 385.36 | 682.53 | 2515.04 | 878.34 | 2139.33 | 2472.37 | 383.1 | 380.51 | 808.67 | 767.56 | 2084.98 | 1772.94 | 457.66 | 1619.75 | 4716.75 | 555.45 | 29308.86 | Cadherin EGF LAG seven-pass G-type receptor 1 (Cadherin family member 9) (Flamingo homolog 2) (hFmi2) | CELSR1 CDHF9 FMI2 | apical protein localization [GO:0045176]; cell-cell adhesion [GO:0098609]; central nervous system development [GO:0007417]; establishment of body hair planar orientation [GO:0048105]; establishment of planar polarity [GO:0001736]; establishment of planar polarity of embryonic epithelium [GO:0042249]; homophilic cell adhesion via plasma membrane adhesion molecules [GO:0007156]; lateral sprouting involved in lung morphogenesis [GO:0060490]; neural tube closure [GO:0001843]; neuron migration [GO:0001764]; orthogonal dichotomous subdivision of terminal units involved in lung branching morphogenesis [GO:0060488]; planar dichotomous subdivision of terminal units involved in lung branching morphogenesis [GO:0060489]; protein localization involved in establishment of planar polarity [GO:0090251]; regulation of actin cytoskeleton organization [GO:0032956]; Rho protein |
| 1375.69 | | 1646.97 | 2387.02 | 1047.21 | 1605.49 | 1984.63 | 1972.19 | 801.09 | 1390.36 | 744.24 | 1011.59 | 2053.54 | 1993.87 | 2744.45 | 1371.74 | 1640.22 | 1787.91 | 1693.05 | 29251.26 | Receptor-type tyrosine-protein phosphatase zeta (R-PTP-zeta) (EC 3.1.3.48) (3F8 chondroitin sulfate proteoglycan) (3H1 keratan sulfate proteoglycan) (Phosphacan) | Ptprz1 Ptprz Ptpz | axonal fasciculation [GO:0007413]; axonogenesis [GO:0007409]; hematopoietic progenitor cell differentiation [GO:0002244]; hippocampus development [GO:0021766]; learning or memory [GO:0007611]; negative regulation of cell population proliferation [GO:0008285]; negative regulation of cell-substrate adhesion [GO:0010812]; negative regulation of dendrite development [GO:2000171]; negative regulation of neuron apoptotic process [GO:0043524]; neuron development [GO:0048666]; neuron projection development [GO:0031175]; oligodendrocyte differentiation [GO:0048709]; peptidyl-tyrosine dephosphorylation [GO:0035335]; positive regulation of cell migration [GO:0030335]; positive regulation of dendrite development [GO:1900006]; positive regulation of fibroblast proliferation [GO:0048146]; positive regulation of neuron migration [GO:2001224]; positive regulation of |

| | | | | | | | | | | | | | | | | | | | | | |
|---|---|---|---|---|---|---|---|---|---|---|---|---|---|---|---|---|---|---|---|---|---|
| 5599.53 | 100.98 | 105.96 | 3410.42 | 470.8 | 2730.63 | 23.24 | 196.86 | 0 | 5682.47 | 151.39 | 160.51 | 9012.62 | 55.52 | 47 | 1179.17 | 196.66 | 69.64 | 29193.4 | Lipopolysaccharide-induced tumor necrosis factor-alpha factor homolog (LPS-induced TNF-alpha factor homolog) (Small integral membrane protein of lysosome/late endosome) | LITAF SIMPLE | cytokine production [GO:0001816]; regulation of cytokine production [GO:0001817]; sexual reproduction [GO:0019953] |
| 1421.54 | 783.93 | 1737.98 | 567.52 | 2361.02 | 1994.14 | 44.65 | 565.58 | 1835.83 | 605.1 | 280.35 | 955.33 | 4697.97 | 4046.09 | 3826.83 | 285.61 | 808.03 | 2221.17 | 29038.67 | Leukocyte receptor cluster member 8 homolog | leng8 zgc:158262 | |
| 2078.22 | 1886.66 | 1641.51 | 651.39 | 1641.44 | 2278.36 | 1594.28 | 761.82 | 2267.95 | 751.39 | 774.65 | 1746.19 | 2036.7 | 1848.24 | 2065.23 | 1149.33 | 1292.72 | 2529.93 | 28996.01 | snRNA-activating protein complex subunit 4 (SNAPc subunit 4) (snRNA-activating protein complex 190 kDa subunit) (SNAPc 190 kDa subunit) | Snapc4 | snRNA transcription by RNA polymerase II [GO:0042795]; snRNA transcription by RNA polymerase III [GO:0042796] |
| 4.12 | 0 | 739.06 | 0.86 | 0.47 | 0 | 0 | 0 | 7188.83 | 0 | 0.3 | 2692.5 | 9.66 | 0.63 | 7689.85 | 0.84 | 2.65 | 10634.01 | 28963.78 | Leucine-rich repeat transmembrane neuronal protein 4 | LRRTM4 UNQ3075/PRO9907 | |

| | | | | | | | | | | | | | | | | | | | | | |
|---|---|---|---|---|---|---|---|---|---|---|---|---|---|---|---|---|---|---|---|---|---|
| 1356.72 | 1472.7 | 1936.46 | 433.75 | 1098.17 | 1539.33 | 3301.83 | 641.75 | 2590.63 | 367.52 | 451.78 | 1334.58 | 1604.42 | 1973.55 | 2536.81 | 2292.53 | 1093.34 | 2812.62 | 28838.49 | BAR/IMD domain-containing adapter protein 2 (Brain-specific angiogenesis inhibitor 1-associated protein 2) (BAI-associated protein 2) (BAI1-associated protein 2) (Insulin receptor substrate protein of 53 kDa) (IRSp53) (Insulin receptor substrate p53) (Insulin receptor tyrosine kinase substrate protein p53) | Baiap2 | actin crosslink formation [GO:0051764]; actin filament bundle assembly [GO:0051017]; cellular response to epidermal growth factor stimulus [GO:0071364]; cellular response to L-glutamate [GO:1905232]; dendrite development [GO:0016358]; modulation of chemical synaptic transmission [GO:0050804]; neuron differentiation [GO:0030182]; plasma membrane organization [GO:0007009]; positive regulation of actin filament polymerization [GO:0030838]; positive regulation of dendritic spine morphogenesis [GO:0061003]; positive regulation of excitatory postsynaptic potential [GO:2000463]; positive regulation of neuron projection development [GO:0010976]; protein localization to synapse [GO:0035418]; regulation of actin cytoskeleton organization [GO:0032956]; regulation of cell shape [GO:0008360]; regulation of modification of postsynaptic actin cytoskeleton [GO:1905274]; regulation of postsynapse organization |
| 6.67 | 9528.29 | 60.91 | 9.63 | 7430.47 | 80.67 | 1.38 | 2658.59 | 7.28 | 8.95 | 999.22 | 20.6 | 7.65 | 6409.84 | 10.32 | 101.92 | 1471.44 | 14.29 | 28828.12 | tRNA (guanosine(18)-2'-O)-methyltransferase TARBP1 (EC 2.1.1.34) (TAR RNA-binding protein 1) (TAR RNA-binding protein of 185 kDa) (TRP-185) | TARBP1 TRM3 TRP185 | regulation of transcription by RNA polymerase II [GO:0006357]; tRNA methylation [GO:0030488] |
| 493.22 | 3623.18 | 1197.98 | 316.49 | 5203.59 | 928.8 | 628.08 | 2046.42 | 792.65 | 153.28 | 1382.33 | 1718.6 | 1091.92 | 5100.07 | 670.77 | 467.88 | 2126.61 | 782.31 | 28724.18 | Protein NLRC3 | Nlrc3 | canonical NF-kappaB signal transduction [GO:0007249]; negative regulation of canonical NF-kappaB signal transduction [GO:0043124]; negative regulation of cytokine production involved in inflammatory response [GO:1900016]; negative regulation of epithelial cell proliferation [GO:0050680]; negative regulation of fibroblast proliferation [GO:0048147]; negative regulation of inflammatory response [GO:0050728]; negative regulation of innate immune response [GO:0045824]; negative regulation of interferon-alpha production [GO:0032687]; negative regulation of interferon-beta production [GO:0032688]; negative regulation of interleukin-1 beta production [GO:0032691]; negative regulation of interleukin-12 production [GO:0032695]; negative regulation of interleukin-6 production [GO:0032715]; negative regulation of NLRP3 inflammasome complex assembly [GO:1900226]; negative regulation of |

| | | | | | | | | | | | | | | | | | | | | | |
|---|---|---|---|---|---|---|---|---|---|---|---|---|---|---|---|---|---|---|---|---|---|
| 2719.29 | 3581.3 | 2135.89 | 544 | 2999.55 | 1379.91 | 450.58 | 1330.3 | 724.44 | 909.03 | 1262.39 | 999.69 | 2242.84 | 2667.4 | 1466.96 | 484.16 | 1896.77 | 868.41 | 28662.91 | Acid sphingomyelinase-like phosphodiesterase 3b (ASM-like phosphodiesterase 3b) (EC 3.1.4.-) | SMPDL3B ASML3B ASMLPD | inflammatory response [GO:0006954]; innate immune response [GO:0045087]; membrane lipid catabolic process [GO:0046466]; negative regulation of inflammatory response [GO:0050728]; negative regulation of toll-like receptor signaling pathway [GO:0034122]; sphingomyelin catabolic process [GO:0006685] |
| 54.01 | 6.55 | 0 | 216.68 | 1.2 | 5.39 | 15152.56 | 0.98 | 3.97 | 103.41 | 303.51 | 2.7 | 6.11 | 2599.26 | 1.17 | 8608.53 | 1565.55 | 24.34 | 28655.92 | RNA-directed RNA polymerase (EC 2.7.7.48) (p91) | | |
| 4698.02 | 60.83 | 64.43 | 2123.54 | 329.59 | 1758.5 | 15.26 | 74.94 | 2.3 | 6358.26 | 171.14 | 239.46 | 11933.26 | 37.81 | 51.12 | 513.47 | 76.61 | 16.74 | 28525.28 | Regucalcin (RC) (Gluconolactonase) (GNL) (EC 3.1.1.17) (Senescence marker protein 30) (SMP-30) (xSMP-30) | rgn | intracellular calcium ion homeostasis [GO:0006874]; L-ascorbic acid biosynthetic process [GO:0019853]; positive regulation of ATP-dependent activity [GO:0032781]; regulation of calcium-mediated signaling [GO:0050848] |

| | | | | | | | | | | | | | | | | | | | | | |
|---|---|---|---|---|---|---|---|---|---|---|---|---|---|---|---|---|---|---|---|---|---|
| 3034.41 | 924.66 | 2461.37 | 920.48 | 792.45 | 2530.15 | 672.47 | 324.17 | 1988.36 | 888.3 | 401.09 | 3078.99 | 2600.97 | 1133.53 | 1894.71 | 972.11 | 765.16 | 3072.21 | 28455.59 | Importin-5 (Imp5) (Importin subunit beta-3) (Karyopherin beta-3) (Ran-binding protein 5) (RanBP5) | Ipo5 Kpnb3 Ranbp5 | cellular response to amino acid stimulus [GO:0071230]; cytoplasmic pattern recognition receptor signaling pathway [GO:0002753]; NLS-bearing protein import into nucleus [GO:0006607]; positive regulation of protein import into nucleus [GO:0042307]; ribosomal protein import into nucleus [GO:0006610] |
| 688.45 | 674.92 | 6059.19 | 634.41 | 1410.11 | 1619.12 | 3893.12 | 430.78 | 1400.35 | 434.13 | 578.6 | 1372.36 | 1302.71 | 1442.53 | 1537.67 | 2524.23 | 754.06 | 1662.74 | 28419.48 | PDZ domain-containing protein 4 (PDZ domain-containing RING finger protein 4-like protein) | PDZD4 KIAA1444 PDZK4 PDZRN4L | |
| 5153.46 | 128.12 | 29.02 | 1114.69 | 916.87 | 1218.72 | 4.58 | 108.97 | 0 | 2638.25 | 567.65 | 188.4 | 15640.38 | 93.55 | 91.32 | 307.55 | 155 | 9.67 | 28366.2 | Replication factor C subunit 5 (Activator 1 36 kDa subunit) (A1 36 kDa subunit) (Activator 1 subunit 5) (Replication factor C 36 kDa subunit) (RF-C 36 kDa subunit) (RFC36) | Rfc5 | DNA-templated DNA replication [GO:0006261]; positive regulation of DNA-directed DNA polymerase activity [GO:1900264] |

| | | | | | | | | | | | | | | | | | | | | |
|---|---|---|---|---|---|---|---|---|---|---|---|---|---|---|---|---|---|---|---|---|
| 1732.34 | 1937.45 | 2758.15 | 838.86 | 2126.33 | 2954.85 | 1390.74 | 823.56 | 719.4 | 582.88 | 715.03 | 2692.82 | 1902.21 | 2405.63 | 1202.96 | 1188.86 | 1232.45 | 967.33 | 28171.85 | Colorectal mutant cancer protein (Protein MCC) | MCC | establishment of protein localization [GO:0045184]; negative regulation of canonical Wnt signaling pathway [GO:0090090]; negative regulation of epithelial cell migration [GO:0010633]; negative regulation of epithelial cell proliferation [GO:0050680]; signal transduction [GO:0007165]; Wnt signaling pathway [GO:0016055] |
| 13.1 | 7889.98 | 4.19 | 0 | 5944.83 | 39.88 | 3.87 | 2466.17 | 2.18 | 1.78 | 864.29 | 0 | 3.56 | 8430.57 | 1.67 | 2.36 | 2467.94 | 6.58 | 28142.95 | Cell adhesion molecule DSCAM (Down syndrome cell adhesion molecule homolog) | DSCAM | axon guidance [GO:0007411]; camera-type eye photoreceptor cell differentiation [GO:0060219]; dendrite morphogenesis [GO:0048813]; dendrite self-avoidance [GO:0070593]; dendritic spine development [GO:0060996]; homophilic cell adhesion via plasma membrane adhesion molecules [GO:0007156]; locomotory behavior [GO:0007626]; negative regulation of cell adhesion [GO:0007162]; positive regulation of axon extension involved in axon guidance [GO:0048842]; positive regulation of phosphorylation [GO:0042327]; post-embryonic retina morphogenesis in camera-type eye [GO:0060060]; retina layer formation [GO:0010842]; social behavior [GO:0035176]; synapse assembly [GO:0007416]; synaptic transmission, glutamatergic [GO:0035249] |

| | | | | | | | | | | | | | | | | | | | | | |
|---|---|---|---|---|---|---|---|---|---|---|---|---|---|---|---|---|---|---|---|---|---|
| 4388.6 | 1255.98 | 634.12 | 1277.14 | 1821.16 | 2942.17 | 10.97 | 460.66 | 79.9 | 3301.83 | 941.42 | 1329.27 | 7666.53 | 312.06 | 928.81 | 308.17 | 331.91 | 151.98 | 28142.68 | Teneurin-4 (Ten-4) (Protein Odd Oz/ten-m homolog 4) (Tenascin-M4) (Ten-m4) (Teneurin transmembrane protein 4) | tenm4 odz4 tnm4 | axon guidance [GO:0007411]; central nervous system myelin formation [GO:0032289]; homophilic cell adhesion via plasma membrane adhesion molecules [GO:0007156]; neuron development [GO:0048666]; positive regulation of gastrulation [GO:2000543]; positive regulation of myelination [GO:0031643]; positive regulation of neuron projection development [GO:0010976]; positive regulation of oligodendrocyte differentiation [GO:0048714]; regulation of myelination [GO:0031641]; signal transduction [GO:0007165] |
| 37.71 | 56.43 | 0 | 22.73 | 68.72 | 0 | 1.4 | 28.46 | 33.2 | 3.93 | 358.34 | 25.67 | 0 | 16544.49 | 34.18 | 2371.61 | 8353.24 | 72.64 | 28012.75 | Zinc fingers and homeoboxes protein 3 (Triple homeobox protein 1) (Zinc finger and homeodomain protein 3) | Zhx3 Kiaa0395 Tix1 | cell differentiation [GO:0030154]; negative regulation of DNA-templated transcription [GO:0045892]; negative regulation of transcription by RNA polymerase II [GO:0000122]; positive regulation of osteoblast differentiation [GO:0045669] |

| | | | | | | | | | | | | | | | | | | | | | |
|---|---|---|---|---|---|---|---|---|---|---|---|---|---|---|---|---|---|---|---|---|---|
| 108.22 | | 4.01 | 32.69 | 135.22 | 3.88 | 54.81 | 11749.63 | 2.37 | 11.48 | 113.28 | 509.19 | 19.89 | 66.64 | 4.83 | 47.61 | 14674.02 | 335.97 | 30.27 | 27904.01 | Sodium/potassium-transporting ATPase subunit beta-1-interacting protein 1 (Na(+)/K(+)-transporting ATPase subunit beta-1-interacting protein 1) (Protein FAM77C) | Nkain1 Fam77c | |
| 1160.74 | | 140.86 | 4440 | 161.78 | 113.25 | 449.18 | 1305.49 | 28.7 | 3958.42 | 203.96 | 49.37 | 1177.01 | 1339.04 | 422.32 | 6310.67 | 809 | 265.62 | 5544.76 | 27880.17 | Tetraspanin-9 (Tspan-9) (Tetraspan NET-5) | TSPAN9 NET5 | |
| 532.07 | | 3290.34 | 2316.16 | 431.8 | 3851.15 | 2781.41 | 58.48 | 1936.44 | 527.77 | 445.12 | 930.74 | 1243.18 | 691.29 | 5014.05 | 780.81 | 201.28 | 1403.04 | 1398.33 | 27833.46 | Ras-related protein R-Ras2 (EC 3.6.5.2) (Ras-like protein TC21) (Teratocarcinoma oncogene) | RRAS2 TC21 | osteoblast differentiation [GO:0001649]; positive regulation of Schwann cell migration [GO:1900149]; Ras protein signal transduction [GO:0007265]; Schwann cell migration [GO:0036135] |

| | | | | | | | | | | | | | | | | | | | | | |
|---|---|---|---|---|---|---|---|---|---|---|---|---|---|---|---|---|---|---|---|---|---|
| 0 | 7949.88 | 4.64 | 5 | 4293.76 | 19.29 | 1.32 | 2235.9 | 1.85 | 0.26 | 1063.87 | 21 | 0 | 7913.1 | 0.7 | 4.38 | 4276.68 | 11.95 | 27803.58 | Epidermal growth factor receptor kinase substrate 8-like protein 2 (EPS8-like protein 2) (Epidermal growth factor receptor pathway substrate 8-related protein 2) (EPS8-related protein 2) | Eps8l2 Eps8r2 | positive regulation of ruffle assembly [GO:1900029]; Rac protein signal transduction [GO:0016601]; regulation of Rho protein signal transduction [GO:0035023]; Rho protein signal transduction [GO:0007266]; sensory perception of sound [GO:0007605] |
| 4030.71 | 828.94 | 1645.94 | 1261.95 | 2382.09 | 2347.05 | 146.93 | 1018.94 | 314.77 | 2563.5 | 1488.79 | 803.15 | 3370.05 | 2057 | 552.29 | 574.15 | 1852.79 | 545.52 | 27784.56 | Inositol monophosphatase 1 (IMP 1) (IMPase 1) (EC 3.1.3.25) (D-galactose 1-phosphate phosphatase) (EC 3.1.3.94) (Inositol-1(or 4)-monophosphatase 1) (Lithium-sensitive myo-inositol monophosphatase A1) | IMPA1 IMPA | inositol biosynthetic process [GO:0006021]; inositol metabolic process [GO:0006020]; phosphatidylinositol phosphate biosynthetic process [GO:0046854]; signal transduction [GO:0007165] |
| 2475.05 | 1776.98 | 372.18 | 1181.15 | 3060.23 | 1664.54 | 127.11 | 1792.81 | 273.84 | 1951.79 | 2191.12 | 1173.05 | 1565.04 | 2621.8 | 422.22 | 661.15 | 3826.96 | 478.52 | 27615.54 | VPS10 domain-containing receptor SorCS1 (hSorCS) | SORCS1 SORCS | neuropeptide signaling pathway [GO:0007218]; post-Golgi vesicle-mediated transport [GO:0006892] |

| | | | | | | | | | | | | | | | | | | | | | |
|---|---|---|---|---|---|---|---|---|---|---|---|---|---|---|---|---|---|---|---|---|---|
| 403.01 | 10.1 | 307.79 | 37.29 | 31.98 | 83.55 | 11.13 | 9.49 | 7374.64 | 20.8 | 15.36 | 3548.31 | 39.69 | 21.58 | 7927.75 | 3.46 | 9.07 | 7744.64 | 27599.64 | Lysosome-associated membrane glycoprotein 1 (LAMP-1) (Lysosome-associated membrane protein 1) (CD107 antigen-like family member A) (CD antigen CD107a) | LAMP1 | establishment of protein localization to organelle [GO:0072594]; Golgi to lysosome transport [GO:0090160]; granzyme-mediated programmed cell death signaling pathway [GO:0140507]; lysosomal lumen acidification [GO:0007042]; positive regulation of natural killer cell degranulation [GO:0043323]; positive regulation of natural killer cell mediated cytotoxicity [GO:0045954]; protein stabilization [GO:0050821]; regulation of organelle transport along microtubule [GO:1902513] |
| 5905.71 | 537.7 | 1954.3 | 1620.48 | 1043.98 | 5669.41 | 29.87 | 483.79 | 548.44 | 1895.87 | 434.19 | 1167.4 | 3015.12 | 1087.49 | 755.98 | 306.43 | 765.11 | 375.72 | 27596.99 | Ceramide kinase (hCERK) (EC 2.7.1.138) (Acylsphingosine kinase) (Lipid kinase 4) (LK4) | CERK KIAA1646 | ceramide metabolic process [GO:0006672]; glycosphingolipid biosynthetic process [GO:0006688] |
| 284.08 | 6998.31 | 306.96 | 109.45 | 5041.67 | 332.9 | 86.61 | 3990.81 | 147.85 | 112.82 | 1270.73 | 253.3 | 507.54 | 3633.67 | 268.44 | 185.91 | 3729.38 | 218.56 | 27478.99 | Apoptosis-stimulating of p53 protein 2 (Bcl2-binding protein) (Bbp) (Renal carcinoma antigen NY-REN-51) (Tumor suppressor p53-binding protein 2) (53BP2) (p53-binding protein 2) (p53BP2) | TP53BP2 ASPP2 BBP | intrinsic apoptotic signaling pathway by p53 class mediator [GO:0072332]; negative regulation of cell cycle [GO:0045786]; positive regulation of execution phase of apoptosis [GO:1900119]; signal transduction [GO:0007165] |
| 578.98 | 792.07 | 1827.12 | 309.84 | 990.92 | 3016.24 | 3222.98 | 409.85 | 2969.47 | 410.86 | 512.04 | 1600.27 | 1081.52 | 903.1 | 2388.11 | 2316.26 | 848.56 | 3258.03 | 27436.22 | Anoctamin-10 (Transmembrane protein 16K) | ano10 tmem16k zgc:114140 | chloride transmembrane transport [GO:1902476] |

| | | | | | | | | | | | | | | | | | | | | | |
|---|---|---|---|---|---|---|---|---|---|---|---|---|---|---|---|---|---|---|---|---|---|
| 2070.43 | 1614.88 | 2360.09 | 767.48 | 1554.88 | 1986.58 | 2486.42 | 719.22 | 809.35 | 768.16 | 755.31 | 1373.08 | 2488.75 | 2372.16 | 1365.12 | 1814.12 | 1207.41 | 864.99 | 27378.43 | S-adenosylmethionine synthase isoform type-1 (AdoMet synthase 1) (EC 2.5.1.6) (Methionine adenosyltransferase 1) (MAT 1) | MAT1A | methionine catabolic process [GO:0009087]; one-carbon metabolic process [GO:0006730]; S-adenosylmethionine biosynthetic process [GO:0006556] |
| 1343.55 | 1488.06 | 1885.42 | 784.51 | 1827.31 | 2065.04 | 2255.76 | 795.97 | 1753.02 | 600.49 | 1021.52 | 2026.5 | 1121.91 | 2474.67 | 1156.5 | 1570.75 | 1556.84 | 1530.99 | 27258.81 | Secretogranin-3 (Secretogranin III) (SgIII) | scg3 si:ch211-12a1.3 zgc:73236 | protein localization to secretory granule [GO:0033366] |
| 1087.65 | 1573.28 | 541.91 | 1022.17 | 1856.85 | 5221.21 | 417.25 | 1755.75 | 847.25 | 1789.64 | 1990.05 | 529.2 | 1038.76 | 3716.09 | 1608.49 | 587.37 | 831.21 | 824.04 | 27238.17 | Triple functional domain protein (EC 2.7.11.1) | Trio | negative regulation of fat cell differentiation [GO:0045599]; neuron projection morphogenesis [GO:0048812]; postsynaptic modulation of chemical synaptic transmission [GO:0099170] |

| | | | | | | | | | | | | | | | | | | | | | |
|---|---|---|---|---|---|---|---|---|---|---|---|---|---|---|---|---|---|---|---|---|---|
| 0 | 0 | 1016.07 | 0 | 0.97 | 0 | 0 | 0.33 | 7941.45 | 0 | 1.07 | 2305.76 | 0 | 0 | 6940.99 | 0.57 | 6.29 | 8949.98 | 27163.48 | Catenin delta-2 (Delta-catenin) (GT24) (Neural plakophilin-related ARM-repeat protein) (NPRAP) (Neurojungin) | CTNND2 NPRAP | cell adhesion [GO:0007155]; cell-cell adhesion [GO:0098609]; dendritic spine morphogenesis [GO:0060997]; regulation of canonical Wnt signaling pathway [GO:0060828]; signal transduction [GO:0007165]; synapse organization [GO:0050808]; Wnt signaling pathway [GO:0016055] |
| 1299.99 | 1860.93 | 7169.47 | 2023.25 | 2103.72 | 250.99 | 457.15 | 1423.04 | 33.26 | 451.8 | 595.34 | 3944.2 | 798.8 | 1831.25 | 46.97 | 1164.64 | 1434.92 | 65.24 | 26954.96 | Intraflagellar transport protein 20 homolog | ift20 zgc:103674 | cell differentiation [GO:0030154]; cilium assembly [GO:0060271]; intraciliary transport [GO:0042073]; protein localization to cilium [GO:0061512]; regulation of platelet-derived growth factor receptor-alpha signaling pathway [GO:2000583]; spermatogenesis [GO:0007283] |

| | | | | | | | | | | | | | | | | | | | | | |
|---|---|---|---|---|---|---|---|---|---|---|---|---|---|---|---|---|---|---|---|---|---|
| 1972.64 | 1625.59 | 2228.95 | 588.65 | 2022.63 | 1959.48 | 884.27 | 673.64 | 2063.88 | 924.55 | 816.5 | 1670.25 | 1334.01 | 2028.45 | 1709.28 | 886.29 | 1283.14 | 2261.77 | 26933.97 | Mediator of RNA polymerase II transcription subunit 7 (Cofactor required for Sp1 transcriptional activation subunit 9) (CRSP complex subunit 9) (Mediator complex subunit 7) | med7 crsp9 | regulation of transcription by RNA polymerase II [GO:0006357] |
| 21.06 | 5291.32 | 224.63 | 22.26 | 4674.44 | 160.88 | 1.13 | 5352.09 | 124.61 | 50.53 | 1902.6 | 148.65 | 14 | 4636.67 | 88.91 | 46.38 | 3914.59 | 207.35 | 26882.1 | CCR4-NOT transcription complex subunit 10 | CNOT10 | mRNA catabolic process [GO:0006402]; negative regulation of translation [GO:0017148] |
| 930.98 | 3424.5 | 1590.56 | 297.61 | 3293.05 | 2049.37 | 661.6 | 1851.7 | 999.62 | 295.67 | 740.02 | 1649.91 | 1115.41 | 3249.52 | 973.67 | 570.36 | 1575.42 | 1585.29 | 26854.26 | IQ motif and SEC7 domain-containing protein 3 (Potential synaptic guanine nucleotide exchange factor for Arf) (SynArfGEF-Po) | Iqsec3 Sag | actin cytoskeleton organization [GO:0030036]; activation of GTPase activity [GO:0090630]; postsynapse organization [GO:0099173]; regulation of ARF protein signal transduction [GO:0032012]; regulation of small GTPase mediated signal transduction [GO:0051056] |

| | | | | | | | | | | | | | | | | | | | | | |
|---|---|---|---|---|---|---|---|---|---|---|---|---|---|---|---|---|---|---|---|---|---|
| 48.98 | 0 | 0 | 264.05 | 0 | 1.41 | 14037.13 | 0 | 0 | 156.61 | 444.87 | 0 | 6 | 0 | 1.81 | 11604.28 | 285.78 | 2.09 | 26853.01 | Lamin-B1 | LMNB1 LMN2 LMNB | heterochromatin formation [GO:0031507]; nuclear envelope organization [GO:0006998]; nuclear migration [GO:0007097]; nuclear pore localization [GO:0051664]; protein localization to nuclear envelope [GO:0090435] |
| 2807.16 | 2493.12 | 2162.38 | 519.19 | 1753.37 | 2221.05 | 154.95 | 1080.73 | 323.54 | 1047.33 | 562.51 | 1051.29 | 3305.89 | 3476.28 | 1116.58 | 364.64 | 1567.48 | 834.97 | 26842.46 | Sodium/iodide cotransporter (Na(+)/I(-) cotransporter) (Sodium-iodide symporter) (Na(+)/I(-) symporter) (Solute carrier family 5 member 5) | Slc5a5 Nis | thyroid hormone generation [GO:0006590] |
| 1521.93 | 2010.67 | 622.43 | 3483.68 | 1836.79 | 82.56 | 2451.82 | 746.51 | 37.61 | 1849.27 | 2210.56 | 1199.37 | 903.21 | 1421.55 | 26.92 | 2889.08 | 3458.77 | 47.64 | 26800.37 | Protocadherin-19 | Pcdh19 Kiaa1313 | homophilic cell adhesion via plasma membrane adhesion molecules [GO:0007156] |

| | | | | | | | | | | | | | | | | | | | | | |
|---|---|---|---|---|---|---|---|---|---|---|---|---|---|---|---|---|---|---|---|---|---|
| 7.67 | 19.88 | 1533.12 | 11.97 | 24.73 | 85.93 | 28.36 | 14.52 | 6389.11 | 9.99 | 23.93 | 2554.08 | 180.93 | 69.16 | 6373.29 | 38.54 | 39.66 | 9272.26 | 26677.13 | ER degradation-enhancing alpha-mannosidase-like protein 2 | Edem2 | carbohydrate metabolic process [GO:0005975]; endoplasmic reticulum mannose trimming [GO:1904380]; ERAD pathway [GO:0036503]; mannoprotein catabolic process [GO:0006058]; positive regulation of retrograde protein transport, ER to cytosol [GO:1904154]; response to unfolded protein [GO:0006986] |
| 427.55 | 609.24 | 1765.98 | 105.65 | 333.33 | 1118.3 | 150.6 | 168.37 | 5966.37 | 138.6 | 148.14 | 2886.27 | 520.87 | 816.62 | 5677.13 | 199.29 | 282.87 | 5341.39 | 26656.57 | SH3 domain and tetratricopeptide repeat-containing protein 1 | SH3TC1 | |
| 1763.63 | 1075.18 | 963.13 | 816.88 | 1195.89 | 1480.74 | 4759.04 | 484.87 | 1745.54 | 686.43 | 626.86 | 1350.9 | 1578.34 | 1486.93 | 1296.3 | 3273.96 | 825.84 | 1239.68 | 26650.14 | Alanine--glyoxylate aminotransferase (AGT) (EC 2.6.1.44) (Serine--pyruvate aminotransferase, mitochondrial) (SPT) (EC 2.6.1.51) | AGXT AGT1 | glycine biosynthetic process, by transamination of glyoxylate [GO:0019265]; glyoxylate metabolic process [GO:0046487] |

| | | | | | | | | | | | | | | | | | | | | | |
|---|---|---|---|---|---|---|---|---|---|---|---|---|---|---|---|---|---|---|---|---|---|
| 35.74 | 0 | 0 | 209.61 | 0 | 7.65 | 12744.23 | 0 | 2.19 | 117.18 | 350.71 | 0 | 8.68 | 0.56 | 0 | 12846.09 | 300.8 | 8.58 | 26632.02 | Lysophosphatidic acid receptor 1-A (LPA receptor 1-A) (LPA-1-A) (Lysophosphatidic acid receptor LPA1 homolog 1) (xLPA1-1) | lpar1-a lpa1r lpa1r1 | adenylate cyclase-activating G protein-coupled receptor signaling pathway [GO:0007189]; adenylate cyclase-inhibiting G protein-coupled receptor signaling pathway [GO:0007193]; negative regulation of neuron projection development [GO:0010977]; neurogenesis [GO:0022008]; positive regulation of MAPK cascade [GO:0043410]; positive regulation of Rho protein signal transduction [GO:0035025]; positive regulation of stress fiber assembly [GO:0051496]; regulation of cell shape [GO:0008360]; regulation of metabolic process [GO:0019222] |
| 0.8 | 2235.48 | 184.6 | 1.5 | 318.87 | 35.59 | 0.73 | 657.6 | 64.57 | 1.68 | 1933.75 | 123.3 | 0.93 | 2352.4 | 1997.09 | 1.18 | 4877.41 | 11788.89 | 26576.37 | Serine/threonine-protein kinase 17A (EC 2.7.11.1) (DAP kinase-related apoptosis-inducing protein kinase 1) | STK17A DRAK1 | apoptotic process [GO:0006915]; intracellular signal transduction [GO:0035556]; positive regulation of apoptotic process [GO:0043065]; positive regulation of fibroblast apoptotic process [GO:2000271]; protein phosphorylation [GO:0006468]; regulation of reactive oxygen species metabolic process [GO:2000377] |

| | | | | | | | | | | | | | | | | | | | | | |
|---|---|---|---|---|---|---|---|---|---|---|---|---|---|---|---|---|---|---|---|---|---|
| 1250.54 | 1902.76 | 1234.51 | 883.92 | 2039.32 | 1940.8 | 3924.89 | 475.1 | 412.17 | 412.08 | 714.12 | 938.29 | 1680.25 | 2796.69 | 620.51 | 3057.13 | 1477.73 | 668.86 | 26429.67 | Relaxin receptor 1 (Leucine-rich repeat-containing G-protein coupled receptor 7) (Relaxin family peptide receptor 1) | RXFP1 LGR7 | adenylate cyclase-activating G protein-coupled receptor signaling pathway [GO:0007189]; extracellular matrix organization [GO:0030198]; hormone-mediated signaling pathway [GO:0009755]; lung connective tissue development [GO:0060427]; myofibroblast differentiation [GO:0036446]; nipple morphogenesis [GO:0060658]; parturition [GO:0007567] |
| 1792.63 | 1678.57 | 2420.02 | 550.59 | 1367.72 | 2726.02 | 2594.78 | 784.93 | 293.46 | 618.25 | 615.3 | 1152.14 | 1741.64 | 2121.48 | 1235.3 | 2673.1 | 1513.61 | 527.95 | 26407.49 | Single-strand DNA endonuclease ASTE1 (EC 3.1.-.-) (Protein asteroid homolog 1) | aste1a si:dkey-11p23.5 | |
| 473.77 | 1201.05 | 1702.64 | 127.94 | 729.28 | 936.44 | 99.44 | 413.51 | 10688.78 | 153.42 | 250.9 | 3610.14 | 156.26 | 310.94 | 729.77 | 231.84 | 372.56 | 4088.42 | 26277.1 | Chromodomain-helicase-DNA-binding protein 3 (CHD-3) (EC 3.6.4.-) (ATP-dependent helicase CHD3) (Mi-2 autoantigen 240 kDa protein) (Mi2-alpha) (Zinc finger helicase) (hZFH) | CHD3 | centrosome cycle [GO:0007098]; chromatin remodeling [GO:0006338]; negative regulation of DNA-templated transcription [GO:0045892]; negative regulation of transcription by RNA polymerase II [GO:0000122]; positive regulation of DNA-templated transcription [GO:0045893]; regulation of cell fate specification [GO:0042659]; regulation of DNA-templated transcription [GO:0006355]; regulation of stem cell differentiation [GO:2000736]; spindle organization [GO:0007051] |

| | | | | | | | | | | | | | | | | | | | | | |
|---|---|---|---|---|---|---|---|---|---|---|---|---|---|---|---|---|---|---|---|---|---|
| 0 | 0 | 0 | 181.27 | 0 | 0 | 12649.19 | 0 | 0 | 42.76 | 447.78 | 0 | 0 | 0 | 0 | 12689.02 | 193.75 | 0 | 26203.77 | Trans-1,2-dihydrobenzene-1,2-diol dehydrogenase (EC 1.3.1.20) (D-xylose 1-dehydrogenase) (D-xylose-NADP dehydrogenase) (EC 1.1.1.179) (Dimeric dihydrodiol dehydrogenase) | dhdh ch211-203b17.3 zgc:101723 | |
| 1339.46 | 2576.42 | 2148.29 | 572.73 | 2586.29 | 3150.74 | 87.73 | 1183.48 | 1038.56 | 615.29 | 691.81 | 1890.5 | 1288.35 | 2518.82 | 1253.03 | 311.67 | 1401.33 | 1398.29 | 26052.79 | Mitogen-activated protein kinase kinase kinase kinase 3 (EC 2.7.11.1) (Germinal center kinase-related protein kinase) (GLK) (MAPK/ERK kinase kinase kinase 3) (MEK kinase kinase 3) (MEKKK 3) | Map4k3 | intracellular signal transduction [GO:0035556]; protein phosphorylation [GO:0006468]; response to tumor necrosis factor [GO:0034612]; response to UV [GO:0009411] |
| 364.25 | 597.68 | 2518.2 | 75.2 | 401.68 | 769.35 | 8.66 | 234.84 | 5603.32 | 145.67 | 312.24 | 2610.33 | 224.9 | 401.56 | 5011.73 | 57.69 | 183.39 | 6481.63 | 26002.32 | Sodium-dependent phosphate transporter 2 (Solute carrier family 20 member 2) | slc20a2 TEgg117l18.1 | phosphate ion transport [GO:0006817] |
| 0 | 0 | 25850.12 | 0 | 0 | 0 | 10.12 | 0 | 0 | 0 | 0 | 0 | 0 | 0 | 2.42 | 3.75 | 0 | 2.38 | 25868.79 | RNA polymerase II subunit A C-terminal domain phosphatase SSU72 (CTD phosphatase SSU72) (EC 3.1.3.16) | ssu72 zgc:73143 | mRNA processing [GO:0006397]; termination of RNA polymerase II transcription [GO:0006369] |

| | | | | | | | | | | | | | | | | | | | | | |
|---|---|---|---|---|---|---|---|---|---|---|---|---|---|---|---|---|---|---|---|---|---|
| 2.93 | 3.22 | 1754.94 | 0 | 5.98 | 18.86 | 2.51 | 0 | 6972.69 | 7.73 | 9.23 | 2554.79 | 3.3 | 0 | 6208 | 10.22 | 8.31 | 8274.44 | 25837.15 | Serologically defined colon cancer antigen 8 (Antigen NY-CO-8) (Centrosomal colon cancer autoantigen protein) (hCCCAP) | SDCCAG8 CCCAP NPHP10 HSPC085 | cell projection organization [GO:0030030]; centrosome cycle [GO:0007098]; establishment of cell polarity [GO:0030010]; microtubule organizing center organization [GO:0031023]; neuron migration [GO:0001764]; regulation of cilium assembly [GO:1902017]; tube formation [GO:0035148] |
| 1092.04 | 6289.01 | 753.99 | 770.83 | 2463.93 | 2093.79 | 172.37 | 1357.37 | 293.69 | 401.3 | 848.54 | 747.46 | 716.11 | 4064.5 | 612.79 | 385.97 | 2359.27 | 338.07 | 25761.03 | Testis-expressed protein 264 (Putative secreted protein Zsig11) | TEX264 ZSIG11 UNQ337/PRO536 | protein-DNA covalent cross-linking repair [GO:0106300]; reticulophagy [GO:0061709] |
| 967.24 | 724.39 | 3499.21 | 715.45 | 1164.84 | 1722.67 | 1703.67 | 509.53 | 987.68 | 348.95 | 388.33 | 5059.83 | 881.54 | 1658.61 | 1108.72 | 1634.39 | 1093.84 | 1560.57 | 25729.46 | Transmembrane protein 200A | tmem200a si:ch211-45m15.5 | |
| 3944.71 | 1646.9 | 846.63 | 1410.45 | 1801.37 | 465.54 | 358.79 | 948.38 | 265.41 | 2981.01 | 837.53 | 499.24 | 4593.98 | 2223.34 | 503.61 | 284.04 | 1363.68 | 675.31 | 25649.92 | E3 ubiquitin-protein ligase RNF180 (EC 2.3.2.27) (RING finger protein 180) (RING-type E3 ubiquitin transferase RNF180) | RNF180 | adult behavior [GO:0030534]; norepinephrine metabolic process [GO:0042415]; positive regulation of proteasomal ubiquitin-dependent protein catabolic process [GO:0032436]; positive regulation of protein ubiquitination [GO:0031398]; protein polyubiquitination [GO:0000209]; serotonin metabolic process [GO:0042428] |

| | | | | | | | | | | | | | | | | | | | | | |
|---|---|---|---|---|---|---|---|---|---|---|---|---|---|---|---|---|---|---|---|---|---|
| 2987.92 | 1250.02 | 3065.49 | 807.71 | 1391.34 | 2717.53 | 219.28 | 505.69 | 1386.41 | 753.3 | 512.97 | 1455.27 | 1810.39 | 1553.52 | 1793.52 | 566.63 | 1046.89 | 1626.67 | 25450.55 | Transposable element Tcb2 transposase | | DNA integration [GO:0015074]; DNA transposition [GO:0006313] |
| 290.53 | 183.94 | 870.58 | 109.46 | 404.12 | 467.01 | 107.06 | 179.84 | 4254.35 | 169.26 | 130.23 | 1734.5 | 558.17 | 394.76 | 8068.35 | 136.44 | 253.31 | 7030.33 | 25342.24 | Semaphorin-7A (CDw108) (JMH blood group antigen) (John-Milton-Hargen human blood group Ag) (Semaphorin-K1) (Sema K1) (Semaphorin-L) (Sema L) (CD antigen CD108) | SEMA7A CD108 SEMAL | axon extension [GO:0048675]; axon guidance [GO:0007411]; immune response [GO:0006955]; inflammatory response [GO:0006954]; integrin-mediated signaling pathway [GO:0007229]; negative chemotaxis [GO:0050919]; neural crest cell migration [GO:0001755]; olfactory lobe development [GO:0021988]; osteoblast differentiation [GO:0001649]; positive regulation of axon extension [GO:0045773]; positive regulation of cell migration [GO:0030335]; positive regulation of ERK1 and ERK2 cascade [GO:0070374]; positive regulation of macrophage cytokine production [GO:0060907]; regulation of inflammatory response [GO:0050727]; regulation of synapse maturation [GO:0090128]; semaphorin-plexin signaling pathway [GO:0071526] |
| 1038.14 | 1346.37 | 1371.13 | 694.76 | 1703.54 | 1785.47 | 4088.43 | 754.69 | 957.18 | 652.93 | 622.44 | 1209.28 | 1127.31 | 2181.69 | 889.45 | 2808.6 | 957.33 | 1142.22 | 25330.96 | E3 ubiquitin-protein ligase znrf2 (EC 2.3.2.27) (RING-type E3 ubiquitin transferase znrf2) (Zinc/RING finger protein 2) | znrf2 zgc:152865 | proteasome-mediated ubiquitin-dependent protein catabolic process [GO:0043161]; protein K48-linked ubiquitination [GO:0070936] |

| | | | | | | | | | | | | | | | | | | | | |
|---|---|---|---|---|---|---|---|---|---|---|---|---|---|---|---|---|---|---|---|---|
| 1355 | 1755.73 | 2097.79 | 958.35 | 1418.56 | 1670.98 | 2544.98 | 726.79 | 467.56 | 538.53 | 832.05 | 1265.7 | 1715.06 | 3167.07 | 724.01 | 1982.03 | 1575.81 | 509.24 | 25305.24 | Single-stranded DNA-binding protein 3 (Sequence-specific single-stranded-DNA-binding protein) | SSBP3 SSDP SSDP1 | head morphogenesis [GO:0060323]; hematopoietic progenitor cell differentiation [GO:0002244]; mesendoderm development [GO:0048382]; midbrain-hindbrain boundary initiation [GO:0021547]; positive regulation of anterior head development [GO:2000744]; positive regulation of cell population proliferation [GO:0008284]; positive regulation of transcription by RNA polymerase II [GO:0045944]; prechordal plate formation [GO:0021501]; protein-containing complex assembly [GO:0065003] |
| 2515.95 | 1671.14 | 1649.92 | 622.73 | 2519.69 | 3401.63 | 1174.51 | 1080.2 | 482.52 | 558.59 | 617.14 | 1376.46 | 1956.77 | 1447.77 | 1138.43 | 656.71 | 1665.14 | 739.55 | 25274.85 | Transcription factor SOX-4 | Sox4 Sox-4 | ascending aorta morphogenesis [GO:0035910]; atrial septum primum morphogenesis [GO:0003289]; cardiac right ventricle morphogenesis [GO:0003215]; cellular response to glucose stimulus [GO:0071333]; endocrine pancreas development [GO:0031018]; gene expression [GO:0010467]; glial cell development [GO:0021782]; glial cell proliferation [GO:0014009]; glucose homeostasis [GO:0042593]; heart development [GO:0007507]; hematopoietic stem cell homeostasis [GO:0061484]; kidney morphogenesis [GO:0060993]; mesenchyme development [GO:0060485]; mitral valve morphogenesis [GO:0003183]; negative regulation of myoblast differentiation [GO:0045662]; negative regulation of transcription by RNA polymerase II [GO:0000122]; nervous system development [GO:0007399]; neuroepithelial cell differentiation [GO:0060563]; noradrenergic neuron [illegible] |
| 1093.25 | 2836.3 | 1284.11 | 278.31 | 2925.69 | 1918.88 | 1556.21 | 1604.64 | 1299.3 | 482.04 | 1238.7 | 1236.25 | 586.65 | 2560.59 | 1415.35 | 586.69 | 1209.35 | 1023.84 | 25136.15 | Myosin-binding protein C, fast-type (Fast MyBP-C) (C-protein, skeletal muscle fast isoform) | MYBPC2 MYBPCF | cell adhesion [GO:0007155]; sarcomere organization [GO:0045214] |

| | | | | | | | | | | | | | | | | | | | | | |
|---|---|---|---|---|---|---|---|---|---|---|---|---|---|---|---|---|---|---|---|---|---|
| 1755.36 | 1685.83 | 2939.65 | 495.58 | 1278.38 | 2528.05 | 2363.06 | 483 | 547.05 | 456.85 | 486.21 | 1366.46 | 1645.63 | 1736.26 | 704.6 | 2348.68 | 1499.48 | 733.99 | 25054.12 | Tyrosine-protein phosphatase non-receptor type 11 (EC 3.1.3.48) (SH-PTP2) (cSH-PTP2) | PTPN11 | cellular response to epidermal growth factor stimulus [GO:0071364]; ERBB signaling pathway [GO:0038127]; negative regulation of chondrocyte differentiation [GO:0032331]; positive regulation of ERK1 and ERK2 cascade [GO:0070374]; positive regulation of ossification [GO:0045778] |
| 284.56 | 81.78 | 74.83 | 394.66 | 95.49 | 165.63 | 15228.26 | 68.17 | 62.86 | 159.33 | 511.93 | 29.54 | 50.26 | 127.53 | 35.56 | 7327.29 | 231.48 | 47.89 | 24977.05 | Transmembrane protein 43 (Protein LUMA) | Tmem43 | innate immune response [GO:0045087]; nuclear membrane organization [GO:0071763] |
| 3291.27 | 1214.22 | 4483.49 | 781.38 | 1298.79 | 1581.49 | 46.92 | 558.44 | 1255.27 | 842.41 | 470.12 | 1788.24 | 2508.87 | 927.66 | 1316.45 | 302.71 | 709.27 | 1524.45 | 24901.45 | Leucine-rich repeat and calponin homology domain-containing protein 4 (Leucine-rich repeat neuronal protein 4) (Leucine-rich neuronal protein) | LRCH4 LRN LRRN1 LRRN4 | membrane raft assembly [GO:0001765]; nervous system development [GO:0007399]; positive regulation of toll-like receptor signaling pathway [GO:0034123] |
| 59.98 | 1.35 | 0 | 366.33 | 0 | 6.37 | 15344.79 | 0 | 2.04 | 131.32 | 250.76 | 3.79 | 2.8 | 0 | 0 | 8459.54 | 227.35 | 6.85 | 24863.27 | Glucoside xylosyltransferase 2 (EC 2.4.2.42) (Glycosyltransferase 8 domain-containing protein 4) | GXYLT2 GLT8D4 | O-glycan processing [GO:0016266] |

| | | | | | | | | | | | | | | | | | | | | | |
|---|---|---|---|---|---|---|---|---|---|---|---|---|---|---|---|---|---|---|---|---|---|
| 111.9 | 6548.73 | 4.8 | 74.55 | 5592.95 | 9.03 | 2.14 | 3430.01 | 2.84 | 24.25 | 1002.48 | 0 | 58 | 5424.37 | 7.87 | 6.42 | 2492.12 | 26.27 | 24818.73 | DNA polymerase eta (EC 2.7.7.7) (RAD30 homolog A) (Xeroderma pigmentosum variant type protein homolog) | Polh Rad30a Xpv | cellular response to UV-C [GO:0071494]; DNA replication [GO:0006260]; DNA synthesis involved in DNA repair [GO:0000731]; postreplication repair [GO:0006301]; pyrimidine dimer repair [GO:0006290] |
| 279.65 | 290.94 | 519.5 | 131.13 | 263.16 | 403.7 | 11554.07 | 115.6 | 453.24 | 142.05 | 437.04 | 272.48 | 380.06 | 317.37 | 453.95 | 7860.44 | 375.21 | 414.02 | 24663.61 | Aftiphilin | Aftph Afth | intracellular transport [GO:0046907]; protein transport [GO:0015031] |
| 482.78 | 703.68 | 1377.04 | 289.94 | 620.73 | 1546.43 | 147.33 | 291.35 | 3936.25 | 265.98 | 476.42 | 1740.76 | 800.34 | 1153.51 | 5344.45 | 322.61 | 493.09 | 4434.96 | 24427.65 | Ran-binding protein 3 (RanBP3) | RANBP3 QccE-19613 | protein export from nucleus [GO:0006611] |
| 132.88 | 667.95 | 1639.03 | 78.26 | 662.43 | 2062.61 | 100.72 | 248.04 | 4050.53 | 81.65 | 232.77 | 1981.76 | 43.29 | 1358.38 | 4488.36 | 491.5 | 827.22 | 5031.24 | 24178.62 | Copper-transporting ATPase 1 (EC 7.2.2.8) (Copper pump 1) (Menkes disease-associated protein) | ATP7A MC1 MNK | ATP metabolic process [GO:0046034]; blood vessel development [GO:0001568]; blood vessel remodeling [GO:0001974]; cartilage development [GO:0051216]; catecholamine metabolic process [GO:0006584]; central nervous system neuron development [GO:0021954]; cerebellar Purkinje cell differentiation [GO:0021702]; collagen fibril organization [GO:0030199]; copper ion export [GO:0060003]; copper ion import [GO:0015677]; copper ion transport [GO:0006825]; dendrite morphogenesis [GO:0048813]; detoxification of copper ion [GO:0010273]; dopamine metabolic process [GO:0042417]; elastic fiber assembly [GO:0048251]; epinephrine metabolic process [GO:0042414]; establishment of localization in cell [GO:0051649]; extracellular matrix organization [GO:0030198]; glycoprotein biosynthetic process [GO:0009101]; hair follicle morphogenesis [GO:0031069]; intracellular copper ion homeostasis |

| | | | | | | | | | | | | | | | | | | | | | |
|---|---|---|---|---|---|---|---|---|---|---|---|---|---|---|---|---|---|---|---|---|---|
| 497.09 | 510.14 | 367.7 | 366.12 | 561.55 | 1004.88 | 8585 | 139.14 | 272.37 | 308.62 | 274.29 | 521.13 | 1447.76 | 657.4 | 467.31 | 7166.6 | 620.85 | 337.04 | 24104.99 | Protein KRI1 homolog | kri1 kri1l | definitive hemopoiesis [GO:0060216]; endonucleolytic cleavage in ITS1 to separate SSU-rRNA from 5.8S rRNA and LSU-rRNA from tricistronic rRNA transcript (SSU-rRNA, 5.8S rRNA, LSU-rRNA) [GO:0000447]; ribosome biogenesis [GO:0042254] |
| 900.03 | 467.8 | 3054.06 | 539.23 | 678.11 | 1579.08 | 1684.24 | 282.02 | 1194.47 | 384.97 | 285.6 | 6545.11 | 989.3 | 1227.36 | 1009.95 | 1286.01 | 802.75 | 1180.97 | 24091.06 | Sphingosine 1-phosphate receptor 3 (S1P receptor 3) (S1P3) (Sphingosine 1-phosphate receptor Edg-3) (S1P receptor Edg-3) | s1pr3 edg3 | |
| 1167.04 | 924.44 | 1624.39 | 365.67 | 750.84 | 1499.13 | 4388.97 | 353.37 | 1899.9 | 367.01 | 425.27 | 1063.31 | 994.84 | 1580.47 | 1653.82 | 2309.22 | 844.15 | 1853.12 | 24064.96 | Opsin-5 (G-protein coupled receptor 136) (G-protein coupled receptor PGR12) (Neuropsin) | Opn5 Gpr136 Pgr12 | cellular response to light stimulus [GO:0071482]; cellular response to UV-A [GO:0071492]; entrainment of circadian clock by photoperiod [GO:0043153]; hyaloid vascular plexus regression [GO:1990384]; phototransduction [GO:0007602]; phototransduction, UV [GO:0007604]; visual perception [GO:0007601] |

| | | | | | | | | | | | | | | | | | | | | | |
|---|---|---|---|---|---|---|---|---|---|---|---|---|---|---|---|---|---|---|---|---|---|
| 458.58 | 2057.03 | 1964.82 | 361.79 | 2356.19 | 2959.04 | 6.24 | 724.73 | 1183.98 | 260.91 | 867.97 | 2003.71 | 737.78 | 3689.08 | 1550.62 | 251.81 | 1329.02 | 1301.28 | 24064.58 | Neural cell adhesion molecule L1 (N-CAM-L1) (NCAM-L1) (L1-CAM) | l1cam | axon guidance [GO:0007411]; brain development [GO:0007420]; positive regulation of axon extension [GO:0045773] |
| 1005.78 | 618.29 | 1324.6 | 285.01 | 333.35 | 1706.31 | 6855.9 | 234.21 | 563.38 | 257.05 | 255.6 | 1179.56 | 1396.31 | 696.19 | 1191.71 | 4320.54 | 631.98 | 1178.84 | 24034.61 | General transcription factor IIE subunit 1 (Transcription initiation factor IIE subunit alpha) (TFIIE-alpha) | GTF2E1 | transcription initiation at RNA polymerase II promoter [GO:0006367] |
| 472.6 | 166.26 | 378.92 | 86.43 | 112.28 | 250.56 | 102.75 | 73.31 | 99.01 | 111.85 | 51.03 | 280.48 | 72.72 | 85.11 | 233.01 | 1510.08 | 4931.89 | 14976.48 | 23994.77 | Lipase member H (EC 3.1.1.-) | liph zgc:91985 | lipid catabolic process [GO:0016042] |
| 1832.59 | 123.3 | 4609.23 | 383.07 | 140.86 | 7051.52 | 5.53 | 77.43 | 1196.89 | 399.69 | 89.66 | 700.66 | 820.89 | 534.31 | 4152.9 | 39.07 | 251.53 | 1542.91 | 23952.04 | Chromosome-associated kinesin KIF4 (Chromokinesin) (Chromosome-associated kinesin KLP1) | kif4 kif4a klp1 | microtubule-based movement [GO:0007018]; mitotic spindle organization [GO:0007052]; spindle elongation [GO:0051231] |
| 56.97 | 3.14 | 3.37 | 530.08 | 13.44 | 11.73 | 12432.87 | 5.27 | 3.52 | 120.89 | 294.88 | 0 | 13.13 | 49.35 | 12.09 | 10143.57 | 210.39 | 7.36 | 23912.05 | Zinc finger protein 181 | ZNF181 | regulation of transcription by RNA polymerase II [GO:0006357] |

| | | | | | | | | | | | | | | | | | | | | | |
|---|---|---|---|---|---|---|---|---|---|---|---|---|---|---|---|---|---|---|---|---|---|
| 49.54 | 61.98 | 3272.92 | 128.94 | 20.03 | 1033.43 | 10493.86 | 11.97 | 84.91 | 117.95 | 221.47 | 2911.38 | 51.95 | 70.5 | 212.15 | 4686.82 | 217.02 | 243.93 | 23890.75 | Histone-lysine N-methyltransferase SETD7 (EC 2.1.1.364) (SET domain-containing protein 7) | setd7 zgc:92330 | heart morphogenesis [GO:0003007]; heterochromatin organization [GO:0070828]; peptidyl-lysine dimethylation [GO:0018027]; peptidyl-lysine monomethylation [GO:0018026]; positive regulation of DNA-templated transcription [GO:0045893] |
| 1865.79 | 1144.15 | 2413.54 | 612.78 | 1961.59 | 3078.63 | 24.12 | 935.89 | 596.89 | 647.3 | 640.97 | 1193.33 | 3117.32 | 2062.19 | 1377.43 | 249.96 | 1064.82 | 883.51 | 23870.21 | G-protein coupled receptor 39 | GPR39 | G protein-coupled receptor signaling pathway [GO:0007186] |
| 851.02 | 2576.39 | 1651.54 | 173.67 | 2518.96 | 1881.8 | 85.36 | 1984.21 | 224.57 | 689.15 | 1522.28 | 1558.35 | 1349.25 | 2959.64 | 1355.1 | 151.44 | 1215.22 | 1059.82 | 23807.77 | Xin actin-binding repeat-containing protein 2 (Beta-xin) (Cardiomyopathy-associated protein 3) (L-NAME-induced actin cytoskeletal protein) | Xirp2 Cmya3 Xin2 | actin filament organization [GO:0007015]; cardiac muscle tissue morphogenesis [GO:0055008]; cell-cell junction organization [GO:0045216]; positive regulation of protein localization [GO:1903829]; regulation of actin filament organization [GO:0110053]; ventricular septum development [GO:0003281] |

| | | | | | | | | | | | | | | | | | | | | | |
|---|---|---|---|---|---|---|---|---|---|---|---|---|---|---|---|---|---|---|---|---|---|
| 1362.03 | 1788.6 | 2041.3 | 510.88 | 1753.09 | 1676.04 | 1923.15 | 895.76 | 755.6 | 385.09 | 725.2 | 1129.85 | 1773.42 | 1872.76 | 1548.79 | 1117.06 | 1283.05 | 1125.71 | 23667.38 | Puratrophin-1 (Pleckstrin homology domain-containing family G member 4) (PH domain-containing family G member 4) (Purkinje cell atrophy-associated protein 1) | PLEKHG4 PRTPHN1 | axon guidance [GO:0007411]; regulation of small GTPase mediated signal transduction [GO:0051056] |
| 1407.27 | 2909.99 | 1102.78 | 557.41 | 1826.83 | 995.29 | 2589.75 | 839.79 | 666.61 | 362.99 | 666.35 | 922.4 | 1848.59 | 1171.73 | 877.02 | 1815.28 | 2241.48 | 767.53 | 23569.09 | Apoptotic chromatin condensation inducer in the nucleus (Acinus) | Acin1 Acinus | apoptotic chromosome condensation [GO:0030263]; apoptotic process [GO:0006915]; erythrocyte differentiation [GO:0030218]; mRNA processing [GO:0006397]; negative regulation of mRNA splicing, via spliceosome [GO:0048025]; positive regulation of apoptotic process [GO:0043065]; positive regulation of monocyte differentiation [GO:0045657]; RNA splicing [GO:0008380] |
| 4.15 | 1.87 | 439.01 | 1.9 | 0 | 14.4 | 0 | 0 | 6365.11 | 1.73 | 1.64 | 2231.18 | 4.7 | 0 | 5886.39 | 3.15 | 7.5 | 8601.94 | 23564.67 | Glycine N-methyltransferase (EC 2.1.1.20) (Folate-binding protein) | Gnmt Fbp-cII | glycine metabolic process [GO:0006544]; glycogen metabolic process [GO:0005977]; methionine metabolic process [GO:0006555]; methylation [GO:0032259]; one-carbon metabolic process [GO:0006730]; protein homotetramerization [GO:0051289]; regulation of gluconeogenesis [GO:0006111]; S-adenosylhomocysteine metabolic process [GO:0046498]; S-adenosylmethionine metabolic process [GO:0046500]; sarcosine metabolic process [GO:1901052] |

| | | | | | | | | | | | | | | | | | | | | | |
|---|---|---|---|---|---|---|---|---|---|---|---|---|---|---|---|---|---|---|---|---|---|
| 1184.72 | | 1618.16 | 1743.41 | 336.29 | 1766.74 | 1626.57 | 1627.58 | 1060.06 | 965.23 | 328.09 | 611.02 | 1172.39 | 2102.96 | 2045.47 | 1367.93 | 1386.1 | 1223.1 | 1396.33 | 23562.15 | Large ribosomal subunit protein mL39 (39S ribosomal protein L39, mitochondrial) (L39mt) (MRP-L39) (39S ribosomal protein L5, mitochondrial) (L5mt) (MRP-L5) | MRPL39 C21orf92 MRPL5 RPML5 MSTP003 PRED22 | mitochondrial translation [GO:0032543] |
| 857.13 | | 4109.91 | 443.71 | 1163.19 | 2726.57 | 536.9 | 1002.06 | 2342.36 | 158.42 | 577.73 | 1306.61 | 425.58 | 1110.83 | 1484.34 | 254.72 | 1037.74 | 3759.97 | 247.59 | 23545.36 | Regulating synaptic membrane exocytosis protein 2 (Rab-3-interacting molecule 2) (RIM 2) (Rab-3-interacting protein 2) | Rims2 Rab3ip2 Rim2 | adenylate cyclase-modulating G protein-coupled receptor signaling pathway [GO:0007188]; calcium ion-regulated exocytosis of neurotransmitter [GO:0048791]; calcium-ion regulated exocytosis [GO:0017156]; cell differentiation [GO:0030154]; establishment of localization in cell [GO:0051649]; insulin secretion [GO:0030073]; intracellular protein transport [GO:0006886]; neurotransmitter secretion [GO:0007269]; positive regulation of excitatory postsynaptic potential [GO:2000463]; positive regulation of gene expression [GO:0010628]; positive regulation of inhibitory postsynaptic potential [GO:0097151]; regulation of calcium-dependent activation of synaptic vesicle fusion [GO:0150037]; regulation of exocytosis [GO:0017157]; regulation of membrane potential [GO:0042391]; spontaneous neurotransmitter secretion [GO:0061669]; synaptic vesicle docking [GO:0016081]; synaptic vesicle priming |
| 717.47 | | 2751.41 | 187.4 | 150.93 | 2041.83 | 229.38 | 11.74 | 2593.06 | 334.62 | 274.13 | 1521.4 | 373.05 | 1332.93 | 5631.61 | 311.96 | 128.15 | 4497.02 | 320.07 | 23408.16 | RNA-binding protein FXR1 (FMR1 autosomal homolog 1) | fxr1 | animal organ development [GO:0048513]; dentate gyrus development [GO:0021542]; heart development [GO:0007507]; membraneless organelle assembly [GO:0140694]; mRNA destabilization [GO:0061157]; mRNA transport [GO:0051028]; muscle cell development [GO:0055001]; negative regulation of inflammatory response [GO:0050728]; negative regulation of long-term synaptic potentiation [GO:1900272]; negative regulation of translation [GO:0017148]; negative regulation of tumor necrosis factor production [GO:0032720]; positive regulation of long-term neuronal synaptic plasticity [GO:0048170]; positive regulation of translation [GO:0045727]; post-transcriptional regulation of gene expression [GO:0010608]; regulation of mRNA stability [GO:0043488]; regulation of neurogenesis [GO:0050767]; regulation of synaptic transmission, glutamatergic [GO:0051966]; regulation of translation |

| | | | | | | | | | | | | | | | | | | | | | |
|---|---|---|---|---|---|---|---|---|---|---|---|---|---|---|---|---|---|---|---|---|---|
| 716.16 | 631.98 | 1341.46 | 512.65 | 515.08 | 943.21 | 3194.24 | 219.41 | 1570.29 | 287.1 | 299.92 | 2560.61 | 1857.37 | 1517.35 | 1319.54 | 3067.13 | 973.2 | 1847.25 | 23373.95 | F-box and WD repeat domain-containing 11-B | fbxw11b | protein ubiquitination [GO:0016567]; sprouting angiogenesis [GO:0002040]; Wnt signaling pathway [GO:0016055] |
| 1826.14 | 1213.23 | 1500.59 | 504.87 | 1034.5 | 1396.61 | 3130.81 | 531.65 | 943.56 | 639.75 | 633.28 | 935.35 | 1172.63 | 1958.57 | 969.37 | 2863.09 | 1443.82 | 644.43 | 23342.25 | Ubiquitin-like modifier-activating enzyme 1 (EC 6.2.1.45) (Ubiquitin-activating enzyme E1) | UBA1 UBE1 | DNA damage response [GO:0006974]; protein ubiquitination [GO:0016567]; ubiquitin-dependent protein catabolic process [GO:0006511] |
| 777.53 | 1190.11 | 2742.81 | 472.1 | 2163.59 | 1379.8 | 3347.73 | 870.83 | 481.56 | 287.96 | 867.53 | 771.86 | 1107.51 | 2353.4 | 710.58 | 2008.71 | 1219.46 | 574.02 | 23327.09 | Semaphorin-4G | Sema4g | cell differentiation [GO:0030154]; nervous system development [GO:0007399]; semaphorin-plexin signaling pathway [GO:0071526] |
| 1.08 | 6234.39 | 12.24 | 2.31 | 5809.41 | 30.39 | 0 | 2346.01 | 0 | 0.76 | 809.21 | 10.6 | 2.18 | 6875.15 | 0 | 0.32 | 1161.86 | 13.29 | 23309.2 | Choline dehydrogenase, mitochondrial (CDH) (CHD) (EC 1.1.99.1) | Chdh | glycine betaine biosynthetic process from choline [GO:0019285] |

| | | | | | | | | | | | | | | | | | | | | | |
|---|---|---|---|---|---|---|---|---|---|---|---|---|---|---|---|---|---|---|---|---|---|
| 2057.56 | 1156.76 | 1194.66 | 602.98 | 1750.79 | 948.91 | 1100.68 | 387.44 | 2188.12 | 552.63 | 568.13 | 954.09 | 2051.61 | 1425.01 | 2400.5 | 848.59 | 884.28 | 2227.08 | 23299.82 | Replication factor C subunit 3 (Activator 1 38 kDa subunit) (A1 38 kDa subunit) (Activator 1 subunit 3) (Replication factor C 38 kDa subunit) (RF-C 38 kDa subunit) (RFC38) | RFC3 | DNA repair [GO:0006281]; DNA-templated DNA replication [GO:0006261]; positive regulation of DNA-directed DNA polymerase activity [GO:1900264] |
| 1632.64 | 1016.04 | 1614.85 | 475.92 | 1085.88 | 1608.67 | 668.34 | 475.08 | 2641.98 | 497.98 | 413.19 | 1773.65 | 1125.29 | 1453.91 | 2234.66 | 675.94 | 791.88 | 3109.84 | 23295.74 | Alpha-mannosidase 2 (EC 3.2.1.114) (Golgi alpha-mannosidase II) (AMan II) (Man II) (Mannosidase alpha class 2A member 1) (Mannosyl-oligosaccharide 1,3-1,6-alpha-mannosidase) | Man2a1 Mana2 | in utero embryonic development [GO:0001701]; liver development [GO:0001889]; lung alveolus development [GO:0048286]; mannose metabolic process [GO:0006013]; mitochondrion organization [GO:0007005]; N-glycan processing [GO:0006491]; positive regulation of neurogenesis [GO:0050769]; protein glycosylation [GO:0006486]; respiratory gaseous exchange by respiratory system [GO:0007585]; retina morphogenesis in camera-type eye [GO:0060042]; vacuole organization [GO:0007033] |

| | | | | | | | | | | | | | | | | | | | | | | |
|---|---|---|---|---|---|---|---|---|---|---|---|---|---|---|---|---|---|---|---|---|---|---|
| 635.28 | | 1994.95 | 1355.75 | 442.73 | 2628.41 | 3204.45 | 8.53 | 951.91 | 1560.33 | 292.94 | 498.04 | 1357.84 | 438.78 | 3466.16 | 1633.94 | 81.6 | 714.85 | 2024.29 | 23290.78 | E3 ubiquitin/ISG15 ligase TRIM25 (EC 6.3.2.n3) (Estrogen-responsive finger protein) (RING finger protein 147) (RING-type E3 ubiquitin transferase) (EC 2.3.2.27) (RING-type E3 ubiquitin transferase TRIM25) (Tripartite motif-containing protein 25) (Ubiquitin/ISG15-conjugating enzyme TRIM25) (Zinc finger protein 147) | TRIM25 EFP RNF147 ZNF147 | antiviral innate immune response [GO:0140374]; cellular response to leukemia inhibitory factor [GO:1990830]; cytoplasmic pattern recognition receptor signaling pathway [GO:0002753]; ERAD pathway [GO:0036503]; host-mediated suppression of symbiont invasion [GO:0046597]; innate immune response [GO:0045087]; positive regulation of canonical NF-kappaB signal transduction [GO:0043123]; positive regulation of DNA-binding transcription factor activity [GO:0051091]; positive regulation of NF-kappaB transcription factor activity [GO:0051092]; protein K48-linked ubiquitination [GO:0070936]; protein monoubiquitination [GO:0006513]; regulation of protein localization [GO:0032880]; regulation of viral entry into host cell [GO:0046596]; response to estrogen [GO:0043627]; response to oxidative stress [GO:0006979]; response to vitamin D [GO:0033280]; suppression of viral release by host |
| 1342.62 | | 1797.27 | 3111.7 | 369.85 | 1490.26 | 1664.13 | 208.53 | 854.62 | 1463.24 | 411.98 | 592.77 | 1527.06 | 2109.11 | 1940.92 | 1089.1 | 411.97 | 1082.7 | 1813.4 | 23281.23 | Sodium channel protein type 3 subunit alpha (Sodium channel protein brain III subunit alpha) (Sodium channel protein type III subunit alpha) (Voltage-gated sodium channel subtype III) (Voltage-gated sodium channel subunit alpha Nav1.3) | SCN3A KIAA1356 NAC3 | behavioral response to pain [GO:0048266]; cardiac muscle cell action potential involved in contraction [GO:0086002]; membrane depolarization during action potential [GO:0086010]; sodium ion transmembrane transport [GO:0035725]; sodium ion transport [GO:0006814] |

| | | | | | | | | | | | | | | | | | | | | | |
|---|---|---|---|---|---|---|---|---|---|---|---|---|---|---|---|---|---|---|---|---|---|
| 1440.95 | 1533.18 | 1034.89 | 550.08 | 1274.97 | 1679.66 | 2633.04 | 667.58 | 1282.55 | 480.13 | 561.57 | 1142.71 | 1419.94 | 1483.91 | 1211.57 | 2085.93 | 1173.63 | 1601.73 | 23258.02 | Calmodulin-binding transcription activator 1 | Camta1 Kiaa0833 | neuromuscular process controlling balance [GO:0050885]; positive regulation of calcineurin-NFAT signaling cascade [GO:0070886]; positive regulation of transcription by RNA polymerase II [GO:0045944] |
| 1309.98 | 2859.83 | 523.95 | 433.91 | 2881.46 | 1001.14 | 14.45 | 1826.42 | 73.24 | 328.24 | 600.66 | 330.31 | 2802.05 | 4693.01 | 882.47 | 289.26 | 1825.67 | 569.2 | 23245.25 | RelA-associated inhibitor (Inhibitor of ASPP protein) (Protein iASPP) (NFkB-interacting protein 1) (PPP1R13B-like protein) | PPP1R13L IASPP NKIP1 PPP1R13BL RAI | apoptotic process [GO:0006915]; cardiac muscle contraction [GO:0060048]; cardiac right ventricle morphogenesis [GO:0003215]; embryonic camera-type eye development [GO:0031076]; hair cycle [GO:0042633]; multicellular organism growth [GO:0035264]; multicellular organismal-level homeostasis [GO:0048871]; negative regulation of inflammatory response [GO:0050728]; negative regulation of transcription by RNA polymerase II [GO:0000122]; positive regulation of cell differentiation [GO:0045597]; post-embryonic development [GO:0009791]; regulation of transcription by RNA polymerase II [GO:0006357]; ventricular cardiac muscle tissue development [GO:0003229] |

| | | | | | | | | | | | | | | | | | | | | | |
|---|---|---|---|---|---|---|---|---|---|---|---|---|---|---|---|---|---|---|---|---|---|
| 676.5 | 3225.23 | 1355.6 | 732.09 | 2351.09 | 1467.51 | 487.91 | 1572.88 | 451.8 | 400.96 | 1604.91 | 2878.85 | 1260.42 | 1115.65 | 504.68 | 355.47 | 2229.47 | 569.2 | 23240.22 | Inward rectifier potassium channel 2 (Inward rectifier K(+) channel Kir2.1) (IRK-1) (RBL-IRK1) (Potassium channel, inwardly rectifying subfamily J member 2) | Kcnj2 Irk1 | cardiac muscle cell action potential [GO:0086001]; cardiac muscle cell action potential involved in contraction [GO:0086002]; cellular response to mechanical stimulus [GO:0071260]; magnesium ion transport [GO:0015693]; membrane repolarization during action potential [GO:0086011]; membrane repolarization during cardiac muscle cell action potential [GO:0086013]; positive regulation of potassium ion transmembrane transport [GO:1901381]; potassium ion import across plasma membrane [GO:1990573]; potassium ion transmembrane transport [GO:0071805]; potassium ion transport [GO:0006813]; protein homotetramerization [GO:0051289]; regulation of cardiac muscle cell contraction [GO:0086004]; regulation of heart rate by cardiac conduction [GO:0086091]; regulation of membrane repolarization [GO:0060306]; regulation of monoatomic ion transmembrane transport [GO:0034765]; regulation of [illegible] |
| 104.36 | 4234.91 | 307.69 | 281.03 | 3633.68 | 265.84 | 6.62 | 3074.88 | 628.08 | 174.45 | 1203.72 | 624.87 | 92.25 | 3756.68 | 631.96 | 76.28 | 2970.99 | 996.31 | 23064.6 | TBC1 domain family member 25 | Tbc1d25 | autophagy [GO:0006914]; regulation of autophagosome maturation [GO:1901096] |
| 245.01 | 397.14 | 1673.25 | 563.73 | 540.34 | 123.22 | 3.82 | 375.44 | 5453.35 | 391.57 | 529.98 | 2401.37 | 247.12 | 377.96 | 3820.67 | 160.53 | 543.04 | 5195.38 | 23042.92 | MICAL-like protein 2 (Junctional Rab13-binding protein) (Molecule interacting with CasL-like 2) (MICAL-L2) | MICALL2 JRAB | actin cytoskeleton organization [GO:0030036]; actin filament polymerization [GO:0030041]; bicellular tight junction assembly [GO:0070830]; endocytic recycling [GO:0032456]; endosome membrane tubulation [GO:0097750]; neuron projection development [GO:0031175]; positive regulation of protein targeting to mitochondrion [GO:1903955]; Rab protein signal transduction [GO:0032482]; substrate adhesion-dependent cell spreading [GO:0034446] |

| | | | | | | | | | | | | | | | | | | | | | |
|---|---|---|---|---|---|---|---|---|---|---|---|---|---|---|---|---|---|---|---|---|---|
| 1082.16 | 1959.09 | 39.27 | 1677.95 | 981.2 | 13.54 | 127.45 | 913.07 | 3.87 | 2412.01 | 3360.78 | 1.8 | 1538.08 | 1919.6 | 8.84 | 2041.16 | 4866.78 | 96.24 | 23042.89 | Tumor protein p63-regulated gene 1-like protein (Mossy fiber terminal-associated vertebrate-specific presynaptic protein) (Protein FAM79A) | TPRG1L FAM79A MOVER | calcineurin-NFAT signaling cascade [GO:0033173]; negative regulation of synaptic transmission [GO:0050805]; regulation of synaptic transmission, glutamatergic [GO:0051966]; synaptic vesicle docking [GO:0016081] |
| 1165.41 | 1647.55 | 1282.92 | 921.55 | 2538.87 | 1446.15 | 1790.55 | 920.42 | 674.87 | 537.51 | 1214.79 | 1028.81 | 1153.48 | 2486.17 | 650.37 | 1225.04 | 1632.19 | 706.02 | 23022.67 | Unconventional myosin-If (Myosin-Ie) | MYO1F | actin filament organization [GO:0007015]; endocytosis [GO:0006897] |
| 1093.83 | 1000.67 | 1785.8 | 329.55 | 1673.61 | 2380.22 | 19.86 | 830.85 | 1647.82 | 445.51 | 557.13 | 1364.76 | 1552.17 | 2473.53 | 2493.41 | 150.48 | 566.7 | 2434.52 | 22800.42 | Intersectin-1 | itsn1 | endocytosis [GO:0006897]; exocytosis [GO:0006887]; intracellular signal transduction [GO:0035556]; positive regulation of Rho protein signal transduction [GO:0035025]; protein transport [GO:0015031] |

| | | | | | | | | | | | | | | | | | | | | | |
|---|---|---|---|---|---|---|---|---|---|---|---|---|---|---|---|---|---|---|---|---|---|
| 1238.09 | 1551.78 | 1413.12 | 667.65 | 1664.41 | 1926.53 | 1919.89 | 525.19 | 1038.17 | 356.64 | 508.71 | 1586.01 | 1673.24 | 2138.28 | 963.52 | 1401.39 | 1135.64 | 1089.53 | 22797.79 | Rap1 GTPase-GDP dissociation stimulator 1-A (Rap1gds1-A protein) (RalB-binding protein A) (XsmgGDS-A) (smgGDS-A) | rap1gds1-a | positive regulation of GTPase activity [GO:0043547]; small GTPase-mediated signal transduction [GO:0007264] |
| 573.73 | 109.66 | 1289.02 | 174.42 | 168.41 | 1171.32 | 7171.7 | 105.82 | 602.92 | 254.01 | 300.19 | 684.11 | 735.6 | 1268.97 | 760.22 | 6252.63 | 561.57 | 542.16 | 22726.46 | Inner nuclear membrane protein Man1 (LEM domain-containing protein 3) | Lemd3 Man1 | angiogenesis [GO:0001525]; blood vessel endothelial cell migration involved in intussusceptive angiogenesis [GO:0002044]; nucleus organization [GO:0006997]; regulation of cell cycle [GO:0051726]; regulation of extracellular matrix organization [GO:1903053]; skeletal muscle cell differentiation [GO:0035914] |
| 1221.56 | 1657.89 | 1003.15 | 574.76 | 2055.25 | 2099.66 | 39.8 | 1426.41 | 123.45 | 635.54 | 981.51 | 898.89 | 3546.76 | 2654.69 | 1231.83 | 303.75 | 1517.39 | 638.04 | 22610.33 | Protein phosphatase 1A (EC 3.1.3.16) (Protein phosphatase 2C isoform alpha) (PP2C-alpha) (Protein phosphatase IA) | PPM1A PPPM1A | cellular response to transforming growth factor beta stimulus [GO:0071560]; dephosphorylation [GO:0016311]; N-terminal protein myristoylation [GO:0006499]; negative regulation of BMP signaling pathway [GO:0030514]; negative regulation of canonical NF-kappaB signal transduction [GO:0043124]; negative regulation of non-canonical NF-kappaB signal transduction [GO:1901223]; negative regulation of transcription by RNA polymerase II [GO:0000122]; negative regulation of transforming growth factor beta receptor signaling pathway [GO:0030512]; positive regulation of canonical NF-kappaB signal transduction [GO:0043123]; positive regulation of canonical Wnt signaling pathway [GO:0090263]; positive regulation of DNA-templated transcription [GO:0045893]; positive regulation of protein export from nucleus [GO:0046827]; protein dephosphorylation [GO:0006470]; protein export from nucleus [illegible] |

| | | | | | | | | | | | | | | | | | | | | | |
|---|---|---|---|---|---|---|---|---|---|---|---|---|---|---|---|---|---|---|---|---|---|
| 215.54 | 165.17 | 1755.94 | 65.75 | 114.78 | 768 | 258.93 | 19.48 | 5729.87 | 68.71 | 57.46 | 1734.46 | 491.57 | 459.35 | 5386.94 | 287.37 | 419.57 | 4475.94 | 22474.83 | tRNA (34-2'-O)-methyltransferase regulator WDR6 (WD repeat-containing protein 6) | WDR6 | G1 to G0 transition [GO:0070314]; negative regulation of autophagy [GO:0010507]; negative regulation of cell population proliferation [GO:0008285]; tRNA methylation [GO:0030488]; wobble position ribose methylation [GO:0002130] |
| 1257.64 | 1801.07 | 1737.28 | 300.65 | 1378.33 | 2721.52 | 446.92 | 679.09 | 1150 | 923.37 | 606.71 | 1661.05 | 1192.79 | 1804.94 | 1805.84 | 401.49 | 988.36 | 1606.19 | 22463.24 | Ankyrin-2 (ANK-2) (Ankyrin-B) (Brain ankyrin) | Ank2 AnkB | atrial cardiac muscle cell action potential [GO:0086014]; atrial cardiac muscle cell to AV node cell communication [GO:0086066]; cardiac muscle contraction [GO:0060048]; endocytosis [GO:0006897]; intracellular calcium ion homeostasis [GO:0006874]; positive regulation of calcium ion transport [GO:0051928]; positive regulation of gene expression [GO:0010628]; positive regulation of potassium ion import across plasma membrane [GO:1903288]; protein localization [GO:0008104]; protein localization to cell surface [GO:0034394]; protein localization to endoplasmic reticulum [GO:0070972]; protein localization to M-band [GO:0036309]; protein localization to plasma membrane [GO:0072659]; protein localization to T-tubule [GO:0036371]; protein stabilization [GO:0050821]; protein transport [GO:0015031]; regulation of atrial cardiac muscle cell action potential [GO:0098910]; regulation of AV node cell |
| 1981.57 | 1883.35 | 1363.31 | 308.97 | 1293.14 | 1526.78 | 428.71 | 838.06 | 104.83 | 566.34 | 928.53 | 787.23 | 2299.65 | 3305.47 | 1117.6 | 1261.99 | 2071.57 | 375.88 | 22442.98 | Palmitoyltransferase ZDHHC5-A (EC 2.3.1.225) (Zinc finger DHHC domain-containing protein 5-A) (DHHC-5A) | zdhhc5a | positive regulation of pattern recognition receptor signaling pathway [GO:0062208] |

| | | | | | | | | | | | | | | | | | | | | | |
|---|---|---|---|---|---|---|---|---|---|---|---|---|---|---|---|---|---|---|---|---|---|
| 2395.24 | 1547.28 | 1892.76 | 553.62 | 1699.68 | 2730.31 | 179.49 | 802.08 | 456.38 | 595.79 | 665.25 | 1013.36 | 2332.12 | 1965.03 | 1021.67 | 374.06 | 1318.65 | 850.98 | 22393.75 | Alpha-2-macroglobulin-like protein 1 (C3 and PZP-like alpha-2-macroglobulin domain-containing protein 9) | A2ML1 CPAMD9 | regulation of endopeptidase activity [GO:0052548] |
| 0 | 0 | 25.41 | 0 | 0 | 0 | 0 | 1.02 | 6825.49 | 0 | 0 | 1854.17 | 0 | 0 | 7372.85 | 0 | 2.43 | 6281.48 | 22362.85 | Growth hormone-regulated TBC protein 1-A | grtp1a grtp1 | |
| 4440.43 | 55.2 | 39.11 | 723.49 | 261.41 | 355.9 | 9.8 | 24.37 | 0 | 2774.14 | 159.35 | 137.77 | 12904.27 | 39.18 | 35.67 | 299.41 | 47.68 | 3.5 | 22310.68 | Oxysterol-binding protein-related protein 10 (ORP-10) (OSBP-related protein 10) | Osbpl10 Orp10 | lipid metabolic process [GO:0006629]; lipid transport [GO:0006869] |
| 1217.59 | 2080.5 | 1403.08 | 532.68 | 2038.35 | 2095.55 | 28.79 | 1051.66 | 283.55 | 924.67 | 1057.93 | 1100.86 | 2450.3 | 2753.92 | 1045.93 | 185.61 | 1440.5 | 583.5 | 22274.97 | Opioid growth factor receptor (OGFr) (Zeta-type opioid receptor) | Ogfr | |
| 3541.04 | 857.19 | 1184.37 | 897.42 | 1440.17 | 2763.67 | 223.93 | 999.63 | 1625.44 | 1293.4 | 694.35 | 1258.88 | 1059.8 | 1256.19 | 542.11 | 221.44 | 1410.58 | 1003.3 | 22272.91 | A disintegrin and metalloproteinase with thrombospondin motifs 6 (ADAM-TS 6) (ADAM-TS6) (ADAMTS-6) (EC 3.4.24.-) | ADAMTS6 | aorta development [GO:0035904]; cardiac septum development [GO:0003279]; coronary vasculature development [GO:0060976]; extracellular matrix organization [GO:0030198]; kidney development [GO:0001822]; proteolysis [GO:0006508] |

| | | | | | | | | | | | | | | | | | | | | | |
|---|---|---|---|---|---|---|---|---|---|---|---|---|---|---|---|---|---|---|---|---|---|
| 313.72 | 1439.13 | 2105.24 | 222.46 | 1155.4 | 1292.95 | 115.22 | 807.72 | 2553.62 | 172.07 | 373.69 | 1695.64 | 427.04 | 2642.97 | 2622.87 | 321.53 | 727.48 | 3056.96 | 22045.71 | Transposon Tf2-9 polyprotein (Retrotransposable element Tf2 155 kDa protein) | Tf2-9 SPBC9B6.02c | DNA integration [GO:0015074]; DNA recombination [GO:0006310]; proteolysis [GO:0006508] |
| 629.12 | 852.42 | 1945.81 | 239.41 | 945.39 | 1474.65 | 32.2 | 531.37 | 2671.43 | 310.07 | 333.06 | 1955 | 677.02 | 1348.21 | 3406.1 | 212.89 | 967.95 | 3494.1 | 22026.2 | Son of sevenless homolog 1 (SOS-1) | SOS1 | axon guidance [GO:0007411]; B cell homeostasis [GO:0001782]; B cell receptor signaling pathway [GO:0050853]; blood vessel morphogenesis [GO:0048514]; cardiac atrium morphogenesis [GO:0003209]; cytokine-mediated signaling pathway [GO:0019221]; epidermal growth factor receptor signaling pathway [GO:0007173]; eyelid development in camera-type eye [GO:0061029]; Fc-epsilon receptor signaling pathway [GO:0038095]; fibroblast growth factor receptor signaling pathway [GO:0008543]; hair follicle development [GO:0001942]; heart trabecula morphogenesis [GO:0061384]; insulin receptor signaling pathway [GO:0008286]; insulin-like growth factor receptor signaling pathway [GO:0048009]; leukocyte migration [GO:0050900]; midbrain morphogenesis [GO:1904693]; multicellular organism growth [GO:0035264]; myelination [GO:0042552]; neurotrophin TRK receptor signaling pathway |
| 643.65 | 882.82 | 531 | 542.9 | 1404.56 | 1415.98 | 4857.54 | 504.09 | 991.21 | 647.55 | 627.94 | 1363.42 | 1076.96 | 1639.6 | 662.76 | 1462.07 | 1454.45 | 1314.66 | 22023.16 | MAP kinase-activated protein kinase 5 (MAPK-activated protein kinase 5) (MAPKAP kinase 5) (MAPKAPK-5) (EC 2.7.11.1) | Mapkapk5 | negative regulation of TOR signaling [GO:0032007]; positive regulation of dendritic spine development [GO:0060999]; positive regulation of telomere maintenance [GO:0032206]; positive regulation of transcription by RNA polymerase II [GO:0045944]; protein autophosphorylation [GO:0046777]; Ras protein signal transduction [GO:0007265]; regulation of translation [GO:0006417]; stress-induced premature senescence [GO:0090400] |

| | | | | | | | | | | | | | | | | | | | | | |
|---|---|---|---|---|---|---|---|---|---|---|---|---|---|---|---|---|---|---|---|---|---|
| 68.08 | 0 | 2.09 | 304.64 | 0 | 10.15 | 14395.45 | 0 | 0 | 156.3 | 420.96 | 0 | 0 | 0 | 0 | 6469.82 | 131.88 | 0 | 21959.37 | Trimeric intracellular cation channel type A (TRIC-A) (TRICA) (Transmembrane protein 38A) | tmem38a zgc:77831 | regulation of release of sequestered calcium ion into cytosol [GO:0051279] |
| 3208.36 | 330.41 | 1141.02 | 953.32 | 239.78 | 1866.54 | 2901.42 | 165.58 | 66.61 | 1046.72 | 466.45 | 445.23 | 2371.62 | 411.01 | 1730.06 | 3858.23 | 675.89 | 58.14 | 21936.39 | Transposable element Tc1 transposase | tc1a tca T07D3.8 | DNA integration [GO:0015074]; DNA transposition [GO:0006313] |
| 1050.73 | 709.72 | 1497.08 | 271.19 | 651.89 | 580.42 | 1108.07 | 294.76 | 4114.53 | 276.05 | 187.93 | 1363.32 | 1010.79 | 1257.24 | 2691.36 | 724.12 | 542.41 | 3587.92 | 21919.53 | Apolipoprotein F (Apo-F) (Liver regeneration-related protein LRRG151) | Apof Ba1-666 | cholesterol efflux [GO:0033344]; cholesterol metabolic process [GO:0008203]; triglyceride metabolic process [GO:0006641] |
| 424.94 | 225.96 | 3626 | 216.54 | 160.08 | 2674.71 | 3.63 | 111.14 | 2927.77 | 253.66 | 125.67 | 1323.4 | 1360.66 | 560.18 | 2910.03 | 238.79 | 425.1 | 4265.9 | 21834.16 | E3 ubiquitin-protein ligase RING2 (EC 2.3.2.27) (RING finger protein 1B) (RING1b) (RING finger protein 2) (RING-type E3 ubiquitin transferase RING2) | rnf2 ring1b | cardiac conduction system development [GO:0003161]; central nervous system development [GO:0007417]; embryonic cranial skeleton morphogenesis [GO:0048701]; enteric nervous system development [GO:0048484]; epigenetic regulation of gene expression [GO:0040029]; negative regulation of intrinsic apoptotic signaling pathway by p53 class mediator [GO:1902254]; negative regulation of transcription by RNA polymerase II [GO:0000122]; pectoral fin development [GO:0033339]; protein ubiquitination [GO:0016567]; regulation of cardiac myofibril assembly [GO:1905304]; regulation of chondrocyte differentiation [GO:0032330]; regulation of neural crest cell differentiation [GO:1905292]; regulation of neural crest cell fate specification [GO:1905295]; retina morphogenesis in camera-type eye [GO:0060042] |

| | | | | | | | | | | | | | | | | | | | | | |
|---|---|---|---|---|---|---|---|---|---|---|---|---|---|---|---|---|---|---|---|---|---|
| 8145.76 | 522.97 | 273.52 | 1658.55 | 234.78 | 384.69 | 14.23 | 215.88 | 103.17 | 3070.7 | 610.94 | 318.89 | 3778.48 | 505.67 | 183.83 | 331.05 | 1120.33 | 249.47 | 21722.91 | Spindlin-1 (Spindlin1) | SPIN1 SPIN | gamete generation [GO:0007276]; meiotic cell cycle [GO:0051321]; positive regulation of DNA-templated transcription [GO:0045893]; positive regulation of Wnt signaling pathway [GO:0030177]; rRNA transcription [GO:0009303]; Wnt signaling pathway [GO:0016055] |
| 795.52 | 3154.72 | 623.02 | 476.01 | 1760.94 | 856.31 | 1951.02 | 1088.04 | 779.54 | 293.99 | 608.89 | 709.89 | 1578.76 | 1493.41 | 961.08 | 1315.36 | 2419.96 | 840.73 | 21707.19 | TBC1 domain family member 4 (Akt substrate of 160 kDa) (AS160) | TBC1D4 AS160 KIAA0603 | cellular response to insulin stimulus [GO:0032869]; negative regulation of vesicle fusion [GO:0031339]; vesicle-mediated transport [GO:0016192] |
| 547.32 | 504.13 | 1393.26 | 153.71 | 376.27 | 533.45 | 291.98 | 174.06 | 4022.63 | 137.89 | 219.04 | 2085 | 506.34 | 518.36 | 3721.64 | 449.06 | 312.1 | 5700.32 | 21646.56 | Carboxypeptidase O (EC 3.4.17.-) | cpo | proteolysis [GO:0006508] |
| 1346.58 | 864.91 | 1731.73 | 527.37 | 625.38 | 1650.52 | 2726.01 | 543.83 | 1201.34 | 437.65 | 384.61 | 1883.44 | 749.62 | 801.28 | 903.21 | 2276.23 | 923.3 | 2035.11 | 21612.12 | Epoxide hydrolase 4 (EC 3.3.-.-) (Abhydrolase domain-containing protein 7) (Epoxide hydrolase-related protein) | Ephx4 Abhd7 Ephxrp | |

| | | | | | | | | | | | | | | | | | | | | | |
|---|---|---|---|---|---|---|---|---|---|---|---|---|---|---|---|---|---|---|---|---|---|
| 873.11 | 841.61 | 1347.45 | 235.94 | 404.93 | 851.94 | 5348.72 | 485.93 | 803.84 | 82.13 | 139.02 | 446.19 | 1361.59 | 700.2 | 938.79 | 4799.93 | 760.75 | 1183.78 | 21605.85 | Storkhead-box protein 2 | STOX2 QmoA-11510 | embryo development ending in birth or egg hatching [GO:0009792]; maternal placenta development [GO:0001893]; regulation of transcription by RNA polymerase II [GO:0006357] |
| 894.46 | 515.93 | 1393.17 | 468.95 | 460.85 | 783.37 | 3596.3 | 212.33 | 1453.88 | 415.72 | 425.44 | 1936.18 | 1218.15 | 1625.42 | 1437.97 | 2935.25 | 746.87 | 1007.79 | 21528.03 | Glutaredoxin 1 | grxC1 grx RP204 | cell redox homeostasis [GO:0045454] |
| 887.38 | 938.2 | 1158.24 | 961.47 | 1188.53 | 2152.38 | 0 | 5.65 | 127.4 | 1247.84 | 1117.36 | 2990.97 | 1543.91 | 1866.76 | 883.27 | 1792.88 | 1544.75 | 1084.07 | 21491.06 | Spliceosome RNA helicase DDX39B (EC 3.6.4.13) (56 kDa U2AF65-associated protein) (ATP-dependent RNA helicase p47) (DEAD box protein UAP56) (HLA-B-associated transcript 1 protein) | DDX39B BAT1 UAP56 | mRNA export from nucleus [GO:0006406]; mRNA splicing, via spliceosome [GO:0000398]; RNA export from nucleus [GO:0006405]; RNA splicing [GO:0008380]; spliceosomal complex assembly [GO:0000245] |

| | | | | | | | | | | | | | | | | | | | | | |
|---|---|---|---|---|---|---|---|---|---|---|---|---|---|---|---|---|---|---|---|---|---|
| 2037.13 | 1116.22 | 2548.76 | 612.69 | 898.79 | 1476.29 | 321.62 | 287.88 | 824.07 | 509.65 | 326.72 | 2738.57 | 2836.08 | 1413.88 | 1508.42 | 508.9 | 635.48 | 812.86 | 21414.01 | DNA-directed RNA polymerase I subunit RPA34 (A34.5) (Antisense to ERCC-1 protein) (ASE-1) (CD3-epsilon-associated protein) (CD3E-associated protein) (DNA-directed RNA polymerase I subunit G) (RNA polymerase I-associated factor PAF49) | POLR1G ASE1 CAST CD3EAP PAF49 | cell surface receptor protein tyrosine kinase signaling pathway [GO:0007169]; rRNA transcription [GO:0009303]; transcription initiation at RNA polymerase I promoter [GO:0006361] |
| 145.56 | 11.84 | 3122.18 | 27.59 | 8.68 | 44.63 | 0.96 | 2.99 | 5976.62 | 19.53 | 12.87 | 1669.35 | 223.66 | 47.06 | 4067.56 | 12.56 | 36.85 | 5971.14 | 21401.63 | Putative E3 ubiquitin-protein ligase UBR7 (EC 2.3.2.27) (N-recognin-7) (RING-type E3 ubiquitin transferase UBR7) | Ubr7 | protein ubiquitination [GO:0016567] |
| 1132.27 | 847.75 | 1621.95 | 294.08 | 566.23 | 1033.88 | 2570.31 | 284.86 | 1680.76 | 369.68 | 245 | 1043.87 | 1542.92 | 1337.07 | 2093.62 | 2100.86 | 667.08 | 1954.24 | 21386.43 | Polymerase delta-interacting protein 2 (38 kDa DNA polymerase delta interaction protein) (p38) | POLDIP2 PDIP38 POLD4 HSPC017 | error-free translesion synthesis [GO:0070987]; mitochondrion organization [GO:0007005]; mitotic spindle assembly [GO:0090307]; negative regulation of macroautophagy [GO:0016242]; positive regulation of focal adhesion assembly [GO:0051894]; positive regulation of mitotic cell cycle [GO:0045931]; positive regulation of mitotic cytokinesis [GO:1903490]; vascular associated smooth muscle cell proliferation [GO:1990874] |

| | | | | | | | | | | | | | | | | | | | | | |
|---|---|---|---|---|---|---|---|---|---|---|---|---|---|---|---|---|---|---|---|---|---|
| 701.6 | 2628.43 | 1804.18 | 311.72 | 1941.73 | 1326.06 | 21.37 | 1566.11 | 837.76 | 355.46 | 994.14 | 1024.84 | 834.61 | 2944.71 | 752.77 | 148.95 | 2003.85 | 1128.98 | 21327.27 | Zyxin | ZYX | cell adhesion [GO:0007155]; integrin-mediated signaling pathway [GO:0007229]; transforming growth factor beta receptor signaling pathway [GO:0007179] |
| 1464.53 | 1643.82 | 3143.06 | 173.57 | 701.81 | 522.06 | 15.38 | 598.77 | 1576.68 | 531.37 | 467.84 | 2120.7 | 1095.49 | 1514.25 | 1198.71 | 426.24 | 1508.96 | 2601.13 | 21304.37 | Tissue factor (TF) (Coagulation factor III) (CD antigen CD142) | F3 Cf-3 Cf3 | blood coagulation [GO:0007596] |
| 656.23 | 834.73 | 695.39 | 139.87 | 1178.81 | 1314.01 | 4509.63 | 567.67 | 1031.44 | 247.44 | 391.7 | 1001.79 | 1164.94 | 1252.29 | 1816.54 | 2277.44 | 555.57 | 1643.46 | 21278.95 | DNA (cytosine-5)-methyltransferase 3C (Dnmt3c) (EC 2.1.1.37) | Dnmt3c Gm14490 | cell differentiation [GO:0030154]; gene silencing by piRNA-directed DNA methylation [GO:0141176]; homologous chromosome pairing at meiosis [GO:0007129]; male meiosis I [GO:0007141]; methylation [GO:0032259]; spermatogenesis [GO:0007283]; transposable element silencing by heterochromatin formation [GO:0141005]; transposable element silencing by piRNA-mediated DNA methylation [GO:0141196]; transposable element silencing by piRNA-mediated heterochromatin formation [GO:0141006] |

| | | | | | | | | | | | | | | | | | | | | | |
|---|---|---|---|---|---|---|---|---|---|---|---|---|---|---|---|---|---|---|---|---|---|
| 783 | 1717.77 | 1347.34 | 1286.27 | 1559.1 | 2443.75 | 26.73 | 943.49 | 692.49 | 1317.68 | 1413.46 | 715.37 | 1109.61 | 2380.36 | 907.83 | 341.86 | 1265.4 | 1007.29 | 21258.8 | Cadherin EGF LAG seven-pass G-type receptor 3 (Cadherin family member 11) (Epidermal growth factor-like protein 1) (EGF-like protein 1) (Flamingo homolog 1) (hFmi1) (Multiple epidermal growth factor-like domains protein 2) (Multiple EGF-like domains protein 2) | CELSR3 CDHF11 EGFL1 FMI1 KIAA0812 MEGF2 | cell-cell adhesion [GO:0098609]; dopaminergic neuron axon guidance [GO:0036514]; G protein-coupled receptor signaling pathway [GO:0007186]; homophilic cell adhesion via plasma membrane adhesion molecules [GO:0007156]; serotonergic neuron axon guidance [GO:0036515]; Wnt signaling pathway, planar cell polarity pathway [GO:0060071] |
| 2089.55 | 1042.35 | 1176.7 | 883.28 | 1051.84 | 1715.71 | 2840.76 | 384.31 | 466.32 | 558.03 | 460.87 | 1018.26 | 1599.93 | 1683.56 | 548.66 | 2098.47 | 1043.12 | 567.4 | 21229.12 | T-box transcription factor TBX1 (T-box protein 1) | tbx1 zgc:136724 | cardiac muscle cell differentiation [GO:0055007]; cartilage morphogenesis [GO:0060536]; cell fate specification [GO:0001708]; chordate pharyngeal muscle development [GO:0043282]; ear development [GO:0043583]; ear morphogenesis [GO:0042471]; embryonic heart tube morphogenesis [GO:0003143]; embryonic viscerocranium morphogenesis [GO:0048703]; heart development [GO:0007507]; heart looping [GO:0001947]; neural crest cell development [GO:0014032]; neural crest cell migration [GO:0001755]; parathyroid gland development [GO:0060017]; pharyngeal system development [GO:0060037]; positive regulation of DNA-templated transcription [GO:0045893]; positive regulation of secondary heart field cardioblast proliferation [GO:0072513]; regulation of transcription by RNA polymerase II [GO:0006357]; semicircular canal morphogenesis [GO:0048752]; soft palate development [illegible] |

| | | | | | | | | | | | | | | | | | | | | | |
|---|---|---|---|---|---|---|---|---|---|---|---|---|---|---|---|---|---|---|---|---|---|
| 56.55 | 2607.81 | 164.82 | 32.19 | 2708.3 | 458.22 | 1.39 | 2541.74 | 1.73 | 35.52 | 1039.22 | 82.1 | 63.29 | 5988.07 | 102.57 | 141.39 | 5070.66 | 62.09 | 21157.66 | 5'-3' exonuclease PLD3 (EC 3.1.16.1) (Choline phosphatase 3) (Phosphatidylcholine-hydrolyzing phospholipase D3) (Phospholipase D3) (PLD 3) | pld3 | immune system process [GO:0002376]; inflammatory response [GO:0006954]; myotube differentiation [GO:0014902]; regulation of cytokine production involved in inflammatory response [GO:1900015] |
| 1360.68 | 1271.11 | 1286.44 | 309.63 | 1694.03 | 2053.03 | 1200.96 | 808.11 | 1013.01 | 512.24 | 600.55 | 1134.2 | 1539.21 | 1812.99 | 1568.18 | 753.3 | 935.58 | 1290.76 | 21144.01 | Probable E3 ubiquitin-protein ligase HECTD4 (EC 2.3.2.26) (HECT domain-containing protein 4) (HECT-type E3 ubiquitin transferase HECTD4) | HECTD4 C12orf51 KIAA0614 | glucose homeostasis [GO:0042593]; glucose metabolic process [GO:0006006]; protein ubiquitination [GO:0016567] |
| 38.37 | 1662.84 | 73.92 | 80.6 | 5430.48 | 433.19 | 0 | 2016.12 | 0 | 48.99 | 5181.19 | 465.97 | 0 | 4030.65 | 148.79 | 196.11 | 1286.35 | 0 | 21093.57 | MAP7 domain-containing protein 1 | Map7d1 Mtap7d1 | microtubule cytoskeleton organization [GO:0000226] |
| 4171.69 | 52.44 | 35.02 | 913.85 | 386.32 | 1152.34 | 0 | 62.94 | 1.31 | 2306.32 | 215.07 | 144.57 | 11222.37 | 46.73 | 70.63 | 145.16 | 78 | 3.43 | 21008.19 | WD repeat-containing protein 13 | WDR13 | |

| | | | | | | | | | | | | | | | | | | | | | | |
|---|---|---|---|---|---|---|---|---|---|---|---|---|---|---|---|---|---|---|---|---|---|---|
| 790.69 | | 1513.66 | 3346.76 | 420.81 | 869.9 | 5907.76 | 153.33 | 441.65 | 303.11 | 389.17 | 594.51 | 2579.03 | 596.54 | 753.33 | 936.99 | 349.37 | 699.97 | 349.57 | 20996.15 | Netrin-G2 (Laminet-2) | Ntng2 Lmnt2 | axonogenesis [GO:0007409]; modulation of chemical synaptic transmission [GO:0050804]; postsynaptic specialization assembly [GO:0098698]; regulation of neuron migration [GO:2001222]; regulation of neuron projection arborization [GO:0150011]; regulation of neuron projection development [GO:0010975]; regulation of presynapse assembly [GO:1905606]; synaptic membrane adhesion [GO:0099560] |
| 483.9 | | 1650.93 | 297 | 257 | 1608.28 | 423.61 | 1335.14 | 344.6 | 403.46 | 135.44 | 431.86 | 269.65 | 644.13 | 7720.62 | 307.19 | 2269.36 | 2097.85 | 293.29 | 20973.31 | Protein FAM110B | fam110b zgc:114139 | |
| 278.07 | | 1559.14 | 239.98 | 763.12 | 2034.63 | 1375.04 | 19.55 | 1249.83 | 427.61 | 110.55 | 286.17 | 326.26 | 480.12 | 6459.5 | 778.86 | 294.61 | 3396.31 | 877.82 | 20957.17 | Sorting nexin-2 | SNX2 | early endosome to Golgi transport [GO:0034498]; intracellular protein transport [GO:0006886]; lamellipodium morphogenesis [GO:0072673]; retrograde transport, endosome to Golgi [GO:0042147] |

| | | | | | | | | | | | | | | | | | | | | | |
|---|---|---|---|---|---|---|---|---|---|---|---|---|---|---|---|---|---|---|---|---|---|
| 234.85 | 3677.8 | 2517.78 | 147.3 | 1570.95 | 823.5 | 5.06 | 1427.86 | 1068.79 | 88.23 | 626.54 | 670.12 | 221.1 | 4785.34 | 393.3 | 44.86 | 1721.27 | 926.5 | 20951.15 | Mitochondrial tRNA methylthiotransferase CDK5RAP1 (EC 2.8.4.3) (CDK5 activator-binding protein C42) (CDK5 regulatory subunit-associated protein 1) (mt-tRNA-2-methylthio-N6-dimethylallyladenosine synthase) (mt-tRNA-N6-(dimethylallyl)adenosine(37) methylthiotransferase) | Cdk5rap1 | mitochondrial tRNA modification [GO:0070900]; negative regulation of cyclin-dependent protein serine/threonine kinase activity [GO:0045736]; positive regulation of mitochondrial translation [GO:0070131]; positive regulation of translational fidelity [GO:0045903]; regulation of cyclin-dependent protein serine/threonine kinase activity [GO:0000079]; RNA modification [GO:0009451] |
| 1144.8 | 1273.93 | 1570.37 | 478.27 | 1562.3 | 3680.46 | 687.99 | 307.97 | 130.58 | 616.14 | 666.09 | 1346.44 | 1635.53 | 1788.26 | 1507.18 | 622.21 | 878 | 1025.59 | 20922.11 | Paired amphipathic helix protein Sin3b (Histone deacetylase complex subunit Sin3b) (Transcriptional corepressor Sin3b) | Sin3b Kiaa0700 | cardiac muscle tissue development [GO:0048738]; negative regulation of cell cycle [GO:0045786]; negative regulation of cell migration [GO:0030336]; negative regulation of DNA-templated transcription [GO:0045892]; negative regulation of transcription by RNA polymerase II [GO:0000122]; skeletal muscle tissue development [GO:0007519] |

| | | | | | | | | | | | | | | | | | | | | | |
|---|---|---|---|---|---|---|---|---|---|---|---|---|---|---|---|---|---|---|---|---|---|
| 133.41 | 3314.19 | 73.9 | 166.67 | 3636.77 | 32.73 | 34.24 | 2162.37 | 13.46 | 53.6 | 1274.99 | 78.41 | 45.26 | 7861.29 | 7.88 | 188.61 | 1703.51 | 28.04 | 20809.33 | LON peptidase N-terminal domain and RING finger protein 2 (Neuroblastoma apoptosis-related protease) (RING finger protein 192) | LONRF2 RNF192 | motor behavior [GO:0061744]; negative regulation of neuron apoptotic process [GO:0043524]; neuromuscular process [GO:0050905]; neuron apoptotic process [GO:0051402]; neuron projection development [GO:0031175]; protein quality control for misfolded or incompletely synthesized proteins [GO:0006515]; spinal cord motor neuron differentiation [GO:0021522] |
| 123.74 | 136.81 | 186.1 | 158.25 | 122.46 | 122.83 | 12023.56 | 56.53 | 210.35 | 139.12 | 398.34 | 377.9 | 317.63 | 195.52 | 306.27 | 5176.45 | 265.15 | 439.79 | 20756.8 | F-box/LRR-repeat protein 19 (F-box and leucine-rich repeat protein 19) | Fbxl19 | proteasome-mediated ubiquitin-dependent protein catabolic process [GO:0043161] |
| 1105.55 | 1582.33 | 789.55 | 468.47 | 1714.56 | 2784.28 | 1203.14 | 979.03 | 774.51 | 397.74 | 602.04 | 1448.86 | 1633.34 | 1570.93 | 1367.24 | 667.8 | 734.86 | 818.89 | 20643.12 | Caspase-8 (CASP-8) (EC 3.4.22.61) [Cleaved into: Caspase-8 subunit p18;Caspase-8 subunit p10] | Casp8 | angiogenesis [GO:0001525]; apoptotic process [GO:0006915]; apoptotic signaling pathway [GO:0097190]; cardiac muscle tissue development [GO:0048738]; execution phase of apoptosis [GO:0097194]; extrinsic apoptotic signaling pathway [GO:0097191]; extrinsic apoptotic signaling pathway via death domain receptors [GO:0008625]; heart development [GO:0007507]; hepatocyte apoptotic process [GO:0097284]; macrophage differentiation [GO:0030225]; necroptotic process [GO:0070266]; negative regulation of necroptotic process [GO:0060546]; neural tube formation [GO:0001841]; positive regulation of apoptotic process [GO:0043065]; positive regulation of execution phase of apoptosis [GO:1900119]; positive regulation of extrinsic apoptotic signaling pathway [GO:2001238]; protein processing [GO:0016485]; proteolysis involved in protein catabolic process [illegible] |

| | | | | | | | | | | | | | | | | | | | | | | |
|---|---|---|---|---|---|---|---|---|---|---|---|---|---|---|---|---|---|---|---|---|---|---|
| 501.89 | | 388.8 | 3770.06 | 134.64 | 486.81 | 1396.82 | 327.14 | 187.73 | 3055.41 | 259.66 | 228.43 | 1940.47 | 600.64 | 1020.98 | 2859.2 | 295.39 | 326.88 | 2859.76 | 20640.71 | Serine palmitoyltransferase 2 (EC 2.3.1.50) (Long chain base biosynthesis protein 2) (LCB 2) (Long chain base biosynthesis protein 2a) (LCB2a) (Serine-palmitoyl-CoA transferase 2) (SPT 2) | Sptlc2 Lcb2 | adipose tissue development [GO:0060612]; ceramide biosynthetic process [GO:0046513]; lipophagy [GO:0061724]; positive regulation of lipophagy [GO:1904504]; sphinganine biosynthetic process [GO:0046511]; sphingolipid biosynthetic process [GO:0030148]; sphingomyelin biosynthetic process [GO:0006686]; sphingosine biosynthetic process [GO:0046512] |
| 1161.58 | | 973.86 | 4345.19 | 229.41 | 740.31 | 610.56 | 45.43 | 607.59 | 2194.65 | 441.19 | 521.52 | 986.35 | 352.02 | 2687.12 | 1562.06 | 121.76 | 638.92 | 2417.58 | 20637.1 | Cadherin-2 (Neural cadherin) (N-cadherin) (Parachute) (ZNCAD) | cdh2 pac | adherens junction organization [GO:0034332]; auditory receptor cell stereocilium organization [GO:0060088]; axial mesoderm structural organization [GO:0048331]; axon guidance [GO:0007411]; axonal fasciculation [GO:0007413]; brain development [GO:0007420]; calcium-dependent cell-cell adhesion via plasma membrane cell adhesion molecules [GO:0016339]; cardioblast differentiation [GO:0010002]; cartilage condensation [GO:0001502]; cell migration [GO:0016477]; cell migration in hindbrain [GO:0021535]; cell migration involved in somitogenic axis elongation [GO:0090248]; cell morphogenesis [GO:0000902]; cell-cell adhesion [GO:0098609]; cell-cell adhesion mediated by cadherin [GO:0044331]; cell-cell adhesion via plasma-membrane adhesion molecules [GO:0098742]; cell-cell junction assembly [GO:0007043]; cilium assembly [GO:0060271]; commissural neuron axon guidance [GO:0071679]; |

| | | | | | | | | | | | | | | | | | | | | | |
|---|---|---|---|---|---|---|---|---|---|---|---|---|---|---|---|---|---|---|---|---|---|
| 1034.86 | 604.32 | 7773.22 | 328.95 | 168.5 | 1856.67 | 5.54 | 73.77 | 75.49 | 364.3 | 198.01 | 1923.98 | 2605.66 | 659.34 | 698.18 | 1135.63 | 883.05 | 238.88 | 20628.35 | Protein kinase C-binding protein NELL1 (NEL-like protein 1) | Nell1 | cell differentiation [GO:0030154]; negative regulation of osteoblast proliferation [GO:0033689]; negative regulation of protein catabolic process [GO:0042177]; positive regulation of bone mineralization [GO:0030501]; positive regulation of osteoblast differentiation [GO:0045669]; regulation of gene expression [GO:0010468] |
| 78.2 | 482.02 | 300.03 | 280.99 | 187.05 | 949.85 | 9746.59 | 70.88 | 56.58 | 149.56 | 284.11 | 205.34 | 18.43 | 160.89 | 837.21 | 6354.93 | 395.02 | 15.56 | 20573.24 | Uridine-cytidine kinase 2-B (UCK 2-B) (EC 2.7.1.48) (Cytidine monophosphokinase 2-B) (Uridine monophosphokinase 2-B) | uck2b si:ch211-284b7.4 zgc:56174 | CTP salvage [GO:0044211]; UMP salvage [GO:0044206] |
| 0.76 | 1.72 | 14207.02 | 0.96 | 0 | 313.91 | 0.23 | 0.21 | 83.71 | 1.27 | 0.51 | 5879.5 | 3.27 | 1.05 | 36.13 | 2.1 | 0.49 | 2.44 | 20535.28 | Protein piccolo (Aczonin) | PCLO ACZ | cytoskeleton organization [GO:0007010]; presynaptic active zone assembly [GO:1904071]; protein localization to synapse [GO:0035418] |

| | | | | | | | | | | | | | | | | | | | | | |
|---|---|---|---|---|---|---|---|---|---|---|---|---|---|---|---|---|---|---|---|---|---|
| 5103.26 | 71.12 | 953.6 | 2155.88 | 114.3 | 4077.33 | 7.07 | 40.06 | 264.66 | 2142.35 | 58.97 | 1157.44 | 2502.06 | 67.74 | 373.67 | 122.51 | 206.84 | 1097.14 | 20516 | Ubiquitin-like protein 4A | ubl4a | post-translational protein targeting to endoplasmic reticulum membrane [GO:0006620]; tail-anchored membrane protein insertion into ER membrane [GO:0071816] |
| 2688.7 | 828.13 | 1424.66 | 527.18 | 1692.24 | 2084.4 | 75.2 | 847.73 | 717.34 | 597.74 | 733.22 | 1017.73 | 2216.58 | 1335.26 | 1223.4 | 179.3 | 1256.61 | 1064.98 | 20510.4 | Plexin-C1 (Virus-encoded semaphorin protein receptor) (CD antigen CD232) | PLXNC1 VESPR | cell adhesion [GO:0007155]; negative regulation of cell adhesion [GO:0007162]; positive regulation of axonogenesis [GO:0050772]; regulation of cell migration [GO:0030334]; regulation of cell shape [GO:0008360]; regulation of synapse pruning [GO:1905806]; semaphorin-plexin signaling pathway [GO:0071526]; synapse assembly [GO:0007416] |
| 3330.86 | 578.11 | 199.76 | 713.44 | 763.58 | 268.67 | 10.2 | 235.54 | 6.39 | 620.61 | 248.39 | 176.44 | 10032.85 | 1336.33 | 960.87 | 414.54 | 379.46 | 162.31 | 20438.35 | Protein phosphatase 1 regulatory subunit 16A (Myosin phosphatase-targeting subunit 3) | PPP1R16A MYPT3 | |
| 349.68 | 910.09 | 483.21 | 32.27 | 902.26 | 1590.2 | 0 | 926.32 | 3896.89 | 259.2 | 395.39 | 1011.41 | 242.99 | 1207.87 | 2618.51 | 255.3 | 1239.87 | 4110.03 | 20431.49 | Signal peptidase complex subunit 3 (Microsomal signal peptidase 22/23 kDa subunit) (SPC22/23) (SPase 22/23 kDa subunit) | SPCS3 SPC22 | signal peptide processing [GO:0006465]; viral protein processing [GO:0019082] |

| | | | | | | | | | | | | | | | | | | | | | | |
|---|---|---|---|---|---|---|---|---|---|---|---|---|---|---|---|---|---|---|---|---|---|---|
| 1200.72 | | 1285.73 | 1505.36 | 418.42 | 1640.93 | 1901.3 | 16.14 | 1330.42 | 303.7 | 454.28 | 518.44 | 1189.56 | 2013.69 | 1918.24 | 1690.04 | 203.39 | 1099.21 | 1571.53 | 20261.1 | Guanine nucleotide exchange factor VAV3 (VAV-3) | Vav3 | angiogenesis [GO:0001525]; B cell receptor signaling pathway [GO:0050853]; cell migration [GO:0016477]; cell projection assembly [GO:0030031]; DNA damage response [GO:0006974]; integrin-mediated signaling pathway [GO:0007229]; lamellipodium assembly [GO:0030032]; neutrophil chemotaxis [GO:0030593]; positive regulation of B cell proliferation [GO:0030890]; positive regulation of cell adhesion [GO:0045785]; positive regulation of phosphatidylinositol 3-kinase/protein kinase B signal transduction [GO:0051897]; regulation of cell size [GO:0008361]; response to xenobiotic stimulus [GO:0009410]; small GTPase-mediated signal transduction [GO:0007264]; vesicle fusion [GO:0006906] |
| 492.72 | | 397.21 | 657.74 | 148.39 | 332.03 | 458.85 | 449.21 | 191.18 | 4871.76 | 194.35 | 145.97 | 1481.67 | 513.18 | 413.96 | 3622.35 | 533.05 | 242.7 | 4927.19 | 20073.51 | Pleckstrin homology domain-containing family A member 5 (PH domain-containing family A member 5) (Phosphoinositol 3-phosphate-binding protein 2) (PEPP-2) | PLEKHA5 KIAA1686 PEPP2 | reproductive system development [GO:0061458] |
| 1030.17 | | 1095.57 | 2102.02 | 329.72 | 1096.87 | 1607.52 | 17.78 | 543.49 | 1470.58 | 393.11 | 491.27 | 1714.53 | 1588.05 | 2034.67 | 1504.38 | 264.59 | 1014.53 | 1754.08 | 20052.93 | Neuropilin-1a (znrp1) | nrp1a np-1 nrp1 | angiogenesis [GO:0001525]; angiogenesis involved in wound healing [GO:0060055]; axon extension [GO:0048675]; axon guidance [GO:0007411]; axonal fasciculation [GO:0007413]; cartilage development [GO:0051216]; cell activation [GO:0001775]; forebrain development [GO:0030900]; larval heart development [GO:0007508]; motor neuron axon guidance [GO:0008045]; neural crest cell migration [GO:0001755]; olfactory bulb axon guidance [GO:0071678]; positive regulation of cell migration involved in sprouting angiogenesis [GO:0090050]; positive regulation of endothelial cell migration [GO:0010595]; positive regulation of filopodium assembly [GO:0051491]; regulation of angiogenesis [GO:0045765]; regulation of axonogenesis [GO:0050770]; regulation of photoreceptor cell axon guidance [GO:2000289]; regulation of retinal ganglion cell axon guidance [GO:0090259]; regulation of vascular [illegible] |

| | | | | | | | | | | | | | | | | | | | | | |
|---|---|---|---|---|---|---|---|---|---|---|---|---|---|---|---|---|---|---|---|---|---|
| 1074.64 | 1025.85 | 331.78 | 25.42 | 94.38 | 0 | 27.71 | 1229.38 | 4392.15 | 1385.67 | 1242.61 | 302.71 | 317.22 | 3127.22 | 1361.19 | 67.02 | 741.45 | 3306.02 | 20052.42 | Mitochondrial transcription rescue factor 1 | Mtres1 | regulation of mitochondrial transcription [GO:1903108]; rescue of stalled ribosome [GO:0072344] |
| 970.63 | 1262.74 | 2163.66 | 307.24 | 1463.2 | 1457.97 | 460.3 | 557.81 | 1369.06 | 371.37 | 455.42 | 1287.63 | 1036.36 | 1617.25 | 1533.09 | 541.27 | 1098.13 | 2098.51 | 20051.64 | Coiled-coil domain-containing protein 120 | CCDC120 JM11 | microtubule anchoring at centrosome [GO:0034454]; protein localization [GO:0008104] |
| 38.04 | 0 | 10.15 | 114.23 | 0 | 14.12 | 11506.1 | 0 | 0 | 75.26 | 214.38 | 4.76 | 10.7 | 0 | 734.16 | 7108.47 | 179.16 | 3.24 | 20012.77 | Pre-mRNA-processing factor 19 (EC 2.3.2.27) (PRP19/PSO4 homolog) (RING-type E3 ubiquitin transferase PRP19) | PRPF19 | DNA damage checkpoint signaling [GO:0000077]; double-strand break repair via nonhomologous end joining [GO:0006303]; inner cell mass cell proliferation [GO:0001833]; lipid biosynthetic process [GO:0008610]; mRNA splicing, via spliceosome [GO:0000398]; positive regulation of mRNA splicing, via spliceosome [GO:0048026]; proteasomal protein catabolic process [GO:0010498]; protein K63-linked ubiquitination [GO:0070534]; protein localization [GO:0008104]; spliceosomal complex assembly [GO:0000245]; spliceosomal tri-snRNP complex assembly [GO:0000244] |

| | | | | | | | | | | | | | | | | | | | | | |
|---|---|---|---|---|---|---|---|---|---|---|---|---|---|---|---|---|---|---|---|---|---|
| 337.17 | 1230.38 | 655.59 | 313.1 | 2271.71 | 1443.54 | 1932.72 | 786.6 | 1110.98 | 341.22 | 580.44 | 960.36 | 1070.37 | 3011.28 | 1405.07 | 780.47 | 741.18 | 1020.76 | 19992.94 | Gag-Pol polyprotein [Cleaved into: Matrix protein p10 (MA);p20;Capsid protein p25 (CA);Nucleocapsid protein p14 (NC-pol);Protease p15 (PR) (EC 3.4.23.-);Reverse transcriptase/ribonuclease H p90 (RT) (EC 2.7.7.49) (EC 2.7.7.7) (EC 3.1.26.4);Integrase p46 (IN) (EC 2.7.7.-) (EC 3.1.-.-)] | gag-pol | DNA integration [GO:0015074]; DNA recombination [GO:0006310]; proteolysis [GO:0006508] |
| 4.72 | 1.05 | 548.85 | 1.33 | 3.04 | 11.07 | 1.7 | 3.98 | 6877.84 | 1.53 | 2.94 | 1905.53 | 0.53 | 0.6 | 4603.94 | 5.81 | 5.77 | 5976.05 | 19956.28 | Elongation factor 2b (EF-2) (EC 3.6.5.-) | eef2b zgc:63584 | cellular response to xenobiotic stimulus [GO:0071466]; chordate embryonic development [GO:0043009]; translational elongation [GO:0006414] |
| 43.09 | 332.44 | 56.29 | 33.28 | 1129.34 | 68.54 | 2.18 | 808.59 | 53.09 | 10.98 | 371.54 | 41.78 | 52.42 | 16129.38 | 60.56 | 16.92 | 658.75 | 58.42 | 19927.59 | Vacuolar protein sorting-associated protein 37A (hVps37A) (ESCRT-I complex subunit VPS37A) (Hepatocellular carcinoma-related protein 1) | VPS37A HCRP1 | macroautophagy [GO:0016236]; membrane fission [GO:0090148]; multivesicular body assembly [GO:0036258]; protein targeting to membrane [GO:0006612]; protein targeting to vacuole [GO:0006623]; protein transport to vacuole involved in ubiquitin-dependent protein catabolic process via the multivesicular body sorting pathway [GO:0043328]; ubiquitin-dependent protein catabolic process via the multivesicular body sorting pathway [GO:0043162]; viral budding via host ESCRT complex [GO:0039702] |

| | | | | | | | | | | | | | | | | | | | | | |
|---|---|---|---|---|---|---|---|---|---|---|---|---|---|---|---|---|---|---|---|---|---|
| 294.89 | 2339.63 | 168.27 | 835.15 | 2223.83 | 2870.4 | 54.79 | 816.47 | 103.02 | 10.13 | 333.38 | 37.56 | 3463.71 | 1883.82 | 1095.73 | 560.18 | 2548.68 | 272.77 | 19912.41 | Aminopeptidase N (AP-N) (bAPN) (EC 3.4.11.2) (Alanyl aminopeptidase) (Aminopeptidase M) (AP-M) (Microsomal aminopeptidase) (CD antigen CD13) | ANPEP APN | angiogenesis [GO:0001525]; cell differentiation [GO:0030154]; peptide catabolic process [GO:0043171]; proteolysis [GO:0006508] |
| 2907.78 | 927.25 | 1375.68 | 259.26 | 882.66 | 1252.95 | 24.07 | 1346.83 | 95.95 | 1047.65 | 1176.51 | 590.92 | 4264.57 | 1703.8 | 677.49 | 90.89 | 888.45 | 383.42 | 19896.13 | Putative fibroblast growth factor 1 (FGF-1) (Acidic fibroblast growth factor) (aFGF) (Heparin-binding growth factor 1) (HBGF-1) | fgf1 fgf-1 zgc:73249 | angiogenesis [GO:0001525]; branch elongation involved in ureteric bud branching [GO:0060681]; fibroblast growth factor receptor signaling pathway [GO:0008543]; hemopoiesis [GO:0030097]; mesonephric epithelium development [GO:0072163]; neurogenesis [GO:0022008]; positive regulation of angiogenesis [GO:0045766]; positive regulation of cell division [GO:0051781]; positive regulation of cell migration [GO:0030335]; positive regulation of cell population proliferation [GO:0008284]; positive regulation of cholesterol biosynthetic process [GO:0045542]; positive regulation of intracellular signal transduction [GO:1902533]; positive regulation of MAPK cascade [GO:0043410]; positive regulation of transcription by RNA polymerase II [GO:0045944]; regulation of cell migration [GO:0030334] |
| 130.2 | 43.02 | 93.7 | 615.89 | 26 | 122.88 | 10700.27 | 18.23 | 65.05 | 97.78 | 222.02 | 73.39 | 97.48 | 75.68 | 58.29 | 7181.53 | 196.4 | 70.55 | 19888.36 | Myosin-binding protein H (MyBP-H) (86 kDa protein) (H-protein) | MYBPH | cell adhesion [GO:0007155]; sarcomere organization [GO:0045214] |
| 17 | 4340.35 | 6.21 | 26.95 | 3629.28 | 34.79 | 0.85 | 3401.48 | 11.15 | 2.6 | 905.73 | 4.78 | 4.52 | 4290.07 | 4.13 | 5.79 | 3154.84 | 21.32 | 19861.84 | DISP complex protein LRCH3 (Leucine-rich repeat and calponin homology domain-containing protein 3) | Lrch3 | septin cytoskeleton organization [GO:0032185] |

| | | | | | | | | | | | | | | | | | | | | | |
|---|---|---|---|---|---|---|---|---|---|---|---|---|---|---|---|---|---|---|---|---|---|
| 149.16 | 4604.46 | 234.64 | 504.06 | 4214.92 | 835.13 | 21.04 | 1451.32 | 93.05 | 64.13 | 1089.15 | 425.84 | 84.89 | 3662.7 | 121.46 | 121.54 | 2015.17 | 114.51 | 19807.17 | Embryonal Fyn-associated substrate (hEFS) (Cas scaffolding protein family member 3) | EFS CASS3 | cell adhesion [GO:0007155]; cell migration [GO:0016477]; cell surface receptor protein tyrosine kinase signaling pathway [GO:0007169]; intracellular signal transduction [GO:0035556] |
| 1264.68 | 1591.59 | 1235.75 | 375.23 | 2058.08 | 2465.09 | 23.98 | 1005.1 | 475.39 | 503.79 | 817.47 | 943.29 | 1026.73 | 2468.77 | 825.96 | 194.96 | 1468.36 | 1060.48 | 19804.7 | Translation initiation factor IF-2, mitochondrial (IF-2(Mt)) (IF-2Mt) (IF2(mt)) | Mtif2 | mitochondrial translational initiation [GO:0070124]; ribosome disassembly [GO:0032790] |
| 2541.4 | 745 | 2239.4 | 638.23 | 756.02 | 2143.81 | 1077.15 | 255.29 | 993.86 | 877.68 | 236.96 | 1522.3 | 1342.47 | 1000.1 | 1055.19 | 732.09 | 647.73 | 998.73 | 19803.41 | Acyl-CoA:lysophosphatidylglycerol acyltransferase 1 (2-acylglycerophosphocholine O-acyltransferase) (EC 2.3.1.62) (Acyl-CoA:monoacylglycerol acyltransferase LPGAT1) (EC 2.3.1.22) (Lysophospholipid acyltransferase 7) (LPLAT7) (EC 2.3.1.-) (Stearoyl-CoA:1-lyso-2-acyl-PE acyltransferase) | LPGAT1 FAM34A KIAA0205 | phosphatidylethanolamine acyl-chain remodeling [GO:0036152]; phosphatidylglycerol acyl-chain remodeling [GO:0036148]; phosphatidylinositol acyl-chain remodeling [GO:0036149]; phospholipid biosynthetic process [GO:0008654]; positive regulation of fatty acid biosynthetic process [GO:0045723]; triglyceride biosynthetic process [GO:0019432] |

| | | | | | | | | | | | | | | | | | | | | | | |
|---|---|---|---|---|---|---|---|---|---|---|---|---|---|---|---|---|---|---|---|---|---|---|
| 903.11 | | 866.29 | 1304.29 | 385.39 | 554.46 | 1090.13 | 2445.71 | 329.73 | 1574.07 | 627.48 | 415.21 | 1083.17 | 1522.96 | 849.62 | 1476.2 | 2260.87 | 549.95 | 1559.92 | 19798.56 | Formin-like protein 2 (Formin homology 2 domain-containing protein 2) | FMNL2 FHOD2 KIAA1902 | cell migration [GO:0016477]; cortical actin cytoskeleton organization [GO:0030866]; cytoskeleton organization [GO:0007010]; regulation of cell morphogenesis [GO:0022604]; regulation of cell shape [GO:0008360] |
| 665.16 | | 735.84 | 609.05 | 699.4 | 2922.44 | 3075.14 | 100.63 | 808.2 | 80.52 | 358.49 | 507.94 | 366.08 | 2729.52 | 3414.83 | 1255.39 | 185.56 | 1002.87 | 262.5 | 19779.56 | SLIT and NTRK-like protein 4 | Slitrk4 | axonogenesis [GO:0007409]; behavioral fear response [GO:0001662]; cell population proliferation [GO:0008283]; long-term synaptic potentiation [GO:0060291]; memory [GO:0007613]; positive regulation of synapse assembly [GO:0051965]; smoothened signaling pathway [GO:0007224] |

| | | | | | | | | | | | | | | | | | | | | | |
|---|---|---|---|---|---|---|---|---|---|---|---|---|---|---|---|---|---|---|---|---|---|
| 34.47 | 0 | 0 | 202.82 | 0 | 4.82 | 11000.4 | 0 | 0 | 90.9 | 252.05 | 0 | 7.27 | 0 | 2.02 | 7945.8 | 210.73 | 6.14 | 19757.42 | Anoctamin-1 (Transmembrane protein 16A) | Ano1 Tmem16a | cellular response to heat [GO:0034605]; chloride transmembrane transport [GO:1902476]; chloride transport [GO:0006821]; detection of temperature stimulus involved in sensory perception of pain [GO:0050965]; establishment of localization in cell [GO:0051649]; glial cell projection elongation [GO:0106091]; mucus secretion [GO:0070254]; phospholipase C-activating G protein-coupled receptor signaling pathway [GO:0007200]; positive regulation of insulin secretion involved in cellular response to glucose stimulus [GO:0035774]; regulation of membrane potential [GO:0042391]; trachea development [GO:0060438] |
| 1225.55 | 864.02 | 1844.21 | 814.85 | 726.63 | 2018.74 | 1133.09 | 448.79 | 898.65 | 1150.88 | 721.58 | 1950 | 1297.65 | 842.62 | 898.78 | 1096.94 | 829.93 | 992.68 | 19755.59 | Sphingosine-1-phosphate phosphatase 1 (SPPase1) (Spp1) (EC 3.1.3.-) (Sphingosine-1-phosphatase 1) | Sgpp1 | ER to Golgi ceramide transport [GO:0035621]; extrinsic apoptotic signaling pathway [GO:0097191]; intrinsic apoptotic signaling pathway [GO:0097193]; phospholipid dephosphorylation [GO:0046839]; regulation of epidermis development [GO:0045682]; regulation of keratinocyte differentiation [GO:0045616]; sphinganine-1-phosphate metabolic process [GO:0006668]; sphingolipid metabolic process [GO:0006665]; sphingosine metabolic process [GO:0006670] |
| 416.62 | 357.22 | 1612.83 | 145.13 | 333.35 | 731.76 | 613.4 | 140.16 | 4282.34 | 136.48 | 124.3 | 1499.17 | 410.19 | 484.38 | 2977.45 | 560.11 | 346.72 | 4527.71 | 19699.32 | Regulator of G-protein signaling 3 (RGS3) (SRB-RGS) | Rgs3 | negative regulation of signal transduction [GO:0009968] |

| | | | | | | | | | | | | | | | | | | | | | |
|---|---|---|---|---|---|---|---|---|---|---|---|---|---|---|---|---|---|---|---|---|---|
| 523.27 | 626.96 | 727.34 | 373.44 | 767.1 | 1317.11 | 6477.3 | 281.9 | 274.2 | 272.37 | 375.34 | 526.72 | 970.67 | 640.29 | 727.5 | 3931.12 | 443.63 | 439.56 | 19695.82 | PHD finger protein 20 (Hepatocellular carcinoma-associated antigen 58 homolog) | Phf20 Hca58 | chromatin organization [GO:0006325]; positive regulation of DNA-templated transcription [GO:0045893] |
| 770.58 | 729.95 | 1120.06 | 256.09 | 745.72 | 1075.94 | 2576.7 | 322.28 | 2917.72 | 414.52 | 357.23 | 1055.84 | 751.68 | 823.61 | 1872.34 | 1247.49 | 516.39 | 2121.18 | 19675.32 | Acetyl-CoA carboxylase (ACC) (EC 6.4.1.2) [Includes: Biotin carboxylase (EC 6.3.4.14)] | ACAC | epigenetic regulation of gene expression [GO:0040029]; fatty acid biosynthetic process [GO:0006633]; malonyl-CoA biosynthetic process [GO:2001295]; positive regulation of DNA-templated transcription [GO:0045893]; positive regulation of gene expression [GO:0010628]; regulation of biological quality [GO:0065008]; response to carbohydrate [GO:0009743]; response to fatty acid [GO:0070542]; response to thyroid hormone [GO:0097066]; transport [GO:0006810] |
| 6.83 | 1.59 | 6.94 | 53.8 | 0 | 4.24 | 12544.12 | 0 | 0 | 60.16 | 278.29 | 0 | 3.31 | 0 | 0 | 6579.44 | 132.93 | 0 | 19671.65 | Protein O-mannose kinase (POMK) (EC 2.7.1.183) (Protein kinase-like protein SgK196) (Sugen kinase 196) | pomk sgk196 zgc:101572 | carbohydrate phosphorylation [GO:0046835]; muscle structure development [GO:0061061]; protein O-linked glycosylation [GO:0006493]; swimming [GO:0036268] |

| | | | | | | | | | | | | | | | | | | | | | |
|---|---|---|---|---|---|---|---|---|---|---|---|---|---|---|---|---|---|---|---|---|---|
| 883.55 | 603.78 | 831.36 | 230.92 | 339.49 | 893.05 | 6083.85 | 212.75 | 1145.92 | 306.48 | 259.55 | 596.97 | 895.7 | 719.62 | 1079.83 | 3056.24 | 446.28 | 1070.51 | 19655.85 | DDB1- and CUL4-associated factor 7 (WD repeat-containing protein 68) (WD repeat-containing protein An11 homolog) | DCAF7 HAN11 WDR68 | protein ubiquitination [GO:0016567] |
| 2629.58 | 964.4 | 2906.23 | 467.65 | 500.59 | 1362.47 | 19.51 | 622.95 | 1687.59 | 395.18 | 269.75 | 1135.71 | 1318.42 | 1331.12 | 1592.66 | 256.84 | 645.89 | 1529.15 | 19635.69 | Centriolar coiled-coil protein of 110 kDa (Centrosomal protein of 110 kDa) (CP110) (Cep110) | CCP110 CEP110 CP110 KIAA0419 | centriole replication [GO:0007099]; centrosome duplication [GO:0051298]; ciliary basal body organization [GO:0032053]; negative regulation of cilium assembly [GO:1902018]; positive regulation of cilium assembly [GO:0045724]; regulation of cytokinesis [GO:0032465] |
| 1190.91 | 1919.95 | 1085.71 | 277.34 | 593.49 | 2824.92 | 207.68 | 451.56 | 1884.7 | 437.98 | 425.43 | 1016.35 | 810.19 | 1557.22 | 1880.36 | 210.6 | 742.89 | 2112.1 | 19629.38 | Chloride anion exchanger (Down-regulated in adenoma) (Protein DRA) (Solute carrier family 26 member 3) | Slc26a3 Dra | cellular response to cAMP [GO:0071320]; intracellular pH elevation [GO:0051454]; membrane hyperpolarization [GO:0060081]; sperm capacitation [GO:0048240] |

| | | | | | | | | | | | | | | | | | | | | | | |
|---|---|---|---|---|---|---|---|---|---|---|---|---|---|---|---|---|---|---|---|---|---|---|
| 696.45 | | 442.37 | 893.91 | 262.24 | 455.97 | 780.75 | 5516.84 | 229.04 | 619.23 | 263.46 | 328.07 | 550.51 | 872.04 | 544.56 | 739.83 | 5226.24 | 469.09 | 694.79 | 19585.39 | Protein RCC2 homolog | rcc2 zgc:77115 | cell division [GO:0051301]; chromosome passenger complex localization to kinetochore [GO:0072356]; neural crest cell migration [GO:0001755]; positive regulation of attachment of spindle microtubules to kinetochore [GO:0051987] |
| 481.76 | | 425.03 | 3239.56 | 127.75 | 397.74 | 846.25 | 4630.8 | 216.25 | 1763.89 | 204.97 | 231.89 | 759.94 | 214.65 | 229.5 | 1609.23 | 2098.79 | 277.89 | 1821.42 | 19577.31 | Ribonucleoside-diphosphate reductase large subunit (EC 1.17.4.1) (Ribonucleoside-diphosphate reductase subunit M1) (Ribonucleotide reductase large subunit) (Ribonucleotide reductase protein R1 class I) | rrm1 | deoxyribonucleotide biosynthetic process [GO:0009263] |

| | | | | | | | | | | | | | | | | | | | | | | |
|---|---|---|---|---|---|---|---|---|---|---|---|---|---|---|---|---|---|---|---|---|---|---|
| 476.81 | | 111.3 | 3488.18 | 212.75 | 196.35 | 815.85 | 1721.79 | 55 | 1733.8 | 234.37 | 145.18 | 4132.9 | 799.76 | 253.63 | 1144.32 | 1850.37 | 265.59 | 1922.84 | 19560.79 | Transcription initiation factor TFIID subunit 5 (Transcription initiation factor TFIID 100 kDa subunit) (TAF(II)100) (TAFII-100) (TAFII100) | TAF5 TAF2D | DNA-templated transcription initiation [GO:0006352]; mRNA transcription by RNA polymerase II [GO:0042789]; positive regulation of DNA-templated transcription [GO:0045893]; positive regulation of transcription initiation by RNA polymerase II [GO:0060261]; regulation of DNA repair [GO:0006282]; regulation of transcription by RNA polymerase II [GO:0006357]; RNA polymerase II preinitiation complex assembly [GO:0051123]; transcription initiation at RNA polymerase II promoter [GO:0006367] |
| 775.89 | | 820.87 | 655.54 | 402.67 | 675.26 | 1221.53 | 4998.09 | 301.18 | 648.32 | 238.52 | 299.63 | 473.97 | 1373.73 | 922.37 | 1039.74 | 3351.59 | 565.45 | 795.83 | 19560.18 | Protein GREB1 | Greb1 Kiaa0575 | |
| 827.29 | | 1296.95 | 460.88 | 2612.76 | 1591.35 | 462.33 | 1025.77 | 998.6 | 305.17 | 1296.56 | 1699.98 | 493.65 | 1408.74 | 1482.8 | 351.87 | 1230.66 | 1650.61 | 362.4 | 19558.37 | Rho-related GTP-binding protein Rho6 (Rho family GTPase 1) (Rnd1) | RND1 RHO6 | actin filament organization [GO:0007015]; negative regulation of cell adhesion [GO:0007162]; neuron remodeling [GO:0016322]; regulation of actin cytoskeleton organization [GO:0032956]; signal transduction [GO:0007165]; small GTPase-mediated signal transduction [GO:0007264] |

| | | | | | | | | | | | | | | | | | | | | | |
|---|---|---|---|---|---|---|---|---|---|---|---|---|---|---|---|---|---|---|---|---|---|
| 975.48 | 810.77 | 1065.9 | 494.45 | 865.81 | 1704.7 | 2302.37 | 377.71 | 824.26 | 408.95 | 540.85 | 1356.45 | 1438.64 | 2434.98 | 744.95 | 1561.57 | 884.93 | 760.39 | 19553.16 | Ubiquitin-conjugating enzyme E2 G1 (EC 2.3.2.23) (E2 ubiquitin-conjugating enzyme G1) (E217K) (UBC7) (Ubiquitin carrier protein G1) (Ubiquitin-protein ligase G1) [Cleaved into: Ubiquitin-conjugating enzyme E2 G1, N-terminally processed] | UBE2G1 UBE2G | proteasome-mediated ubiquitin-dependent protein catabolic process [GO:0043161]; protein K48-linked ubiquitination [GO:0070936]; protein K63-linked ubiquitination [GO:0070534]; protein polyubiquitination [GO:0000209]; ubiquitin-dependent protein catabolic process [GO:0006511] |
| 414.46 | 3624.78 | 570.53 | 166.71 | 3669.52 | 844.55 | 673.4 | 2562.86 | 468.23 | 119.78 | 661.05 | 1237.05 | 419.7 | 1306.74 | 821.67 | 394.6 | 829.52 | 757.5 | 19542.65 | Insulin receptor substrate 2-B (IRS-2-B) | irs2-b | insulin receptor signaling pathway [GO:0008286] |
| 2725.04 | 659.36 | 2579.36 | 523.2 | 471.26 | 4093.54 | 218.79 | 268.64 | 235.02 | 836.66 | 236.15 | 1049.23 | 3370.32 | 284.99 | 920.99 | 262.28 | 408.53 | 356.62 | 19499.98 | Histone-lysine N-methyltransferase EZH1 (EC 2.1.1.356) (Enhancer of zeste homolog 1) | EZH1 | heterochromatin formation [GO:0031507]; methylation [GO:0032259] |
| 843.17 | 1306.95 | 1017.36 | 702.76 | 1153.7 | 1769.33 | 5067.2 | 1192.99 | 2419.22 | 7.89 | 44.96 | 178.54 | 3.48 | 30.07 | 31.54 | 1864.73 | 1167.68 | 673.89 | 19475.46 | 6-phosphogluconate dehydrogenase, decarboxylating (EC 1.1.1.44) | Pgd | D-gluconate metabolic process [GO:0019521]; NADP metabolic process [GO:0006739]; pentose biosynthetic process [GO:0019322]; pentose-phosphate shunt [GO:0006098]; pentose-phosphate shunt, oxidative branch [GO:0009051] |

| | | | | | | | | | | | | | | | | | | | | | |
|---|---|---|---|---|---|---|---|---|---|---|---|---|---|---|---|---|---|---|---|---|---|
| 38.31 | 58.97 | 1662.62 | 54.38 | 35.57 | 253.73 | 165.65 | 15.71 | 5998.94 | 15.06 | 21.98 | 1757.8 | 32.52 | 369.5 | 3521.91 | 206.43 | 147.9 | 5080.25 | 19437.23 | Lysine-specific histone demethylase 1A (EC 1.14.99.66) (BRAF35-HDAC complex protein BHC110) (Flavin-containing amine oxidase domain-containing protein 2) ([histone H3]-dimethyl-L-lysine(4) FAD-dependent demethylase 1A) | KDM1A AOF2 KDM1 KIAA0601 LSD1 | cellular response to cAMP [GO:0071320]; cellular response to gamma radiation [GO:0071480]; cellular response to UV [GO:0034644]; cerebral cortex development [GO:0021987]; DNA repair-dependent chromatin remodeling [GO:0140861]; epigenetic regulation of gene expression [GO:0040029]; guanine metabolic process [GO:0046098]; muscle cell development [GO:0055001]; negative regulation of DNA damage response, signal transduction by p53 class mediator [GO:0043518]; negative regulation of intrinsic apoptotic signaling pathway by p53 class mediator [GO:1902254]; negative regulation of intrinsic apoptotic signaling pathway in response to DNA damage by p53 class mediator [GO:1902166]; negative regulation of transcription by RNA polymerase II [GO:0000122]; negative regulation of transcription initiation-coupled chromatin remodeling [GO:0160217]; neuron maturation [GO:0042551]; positive regulation of cell |
| 1521 | 2202.29 | 97 | 2227.65 | 1425.52 | 875.98 | 36.06 | 1015.3 | 6.16 | 1575.67 | 2027.73 | 32.62 | 713.18 | 1687.28 | 440.09 | 1314.94 | 2186.32 | 42.64 | 19427.43 | Focal adhesion kinase 1 (FADK 1) (EC 2.7.10.2) (Focal adhesion kinase-related nonkinase) (FRNK) (p41/p43FRNK) (Protein-tyrosine kinase 2) (p125FAK) (pp125FAK) | PTK2 FAK FAK1 | actin filament organization [GO:0007015]; angiogenesis involved in wound healing [GO:0060055]; epidermal growth factor receptor signaling pathway [GO:0007173]; negative regulation of anoikis [GO:2000811]; negative regulation of cell-substrate adhesion [GO:0010812]; negative regulation of protein autophosphorylation [GO:0031953]; positive regulation of cell migration [GO:0030335]; positive regulation of cell population proliferation [GO:0008284]; positive regulation of focal adhesion assembly [GO:0051894]; positive regulation of protein binding [GO:0032092]; positive regulation of protein tyrosine kinase activity [GO:0061098]; positive regulation of substrate-dependent cell migration, cell attachment to substrate [GO:1904237]; protein autophosphorylation [GO:0046777]; radial glia-guided pyramidal neuron migration [GO:0140650]; regulation of cell adhesion [GO:0030155]; response |

| | | | | | | | | | | | | | | | | | | | | | |
|---|---|---|---|---|---|---|---|---|---|---|---|---|---|---|---|---|---|---|---|---|---|
| 1389.05 | 688.22 | 2870.84 | 381.98 | 460.35 | 2186.53 | 601.59 | 150.13 | 3030.87 | 339.12 | 139.24 | 1539.38 | 907.1 | 470.1 | 1256.18 | 400.6 | 448.06 | 2166.2 | 19425.54 | Glutamate decarboxylase 1 (EC 4.1.1.15) (67 kDa glutamic acid decarboxylase) (GAD-67) (Glutamate decarboxylase 67 kDa isoform) | GAD1 GAD67 | gamma-aminobutyric acid biosynthetic process [GO:0009449]; glutamate catabolic process [GO:0006538]; locomotory exploration behavior [GO:0035641]; social behavior [GO:0035176] |
| 1366.27 | 1059.17 | 1086.89 | 354.73 | 1230.64 | 1285.01 | 394.3 | 1498.74 | 1341.51 | 434.51 | 884.87 | 860.77 | 935.6 | 2313.65 | 1157.12 | 425.81 | 1399.39 | 1285.81 | 19314.79 | Protein Niban 2 (Meg-3) (Niban-like protein 1) (Protein FAM129B) | Niban2 Fam129b | gonadotropin secretion [GO:0032274]; negative regulation of apoptotic process [GO:0043066]; negative regulation of cell population proliferation [GO:0008285]; negative regulation of DNA biosynthetic process [GO:2000279]; negative regulation of DNA-templated transcription [GO:0045892]; positive regulation of DNA-templated transcription [GO:0045893] |

| | | | | | | | | | | | | | | | | | | | | | |
|---|---|---|---|---|---|---|---|---|---|---|---|---|---|---|---|---|---|---|---|---|---|
| 3401.55 | 758.34 | 1003.09 | 832.7 | 692.58 | 1314.28 | 693.46 | 246.55 | 669.06 | 814.02 | 334.2 | 804.5 | 3591.51 | 1041.71 | 1007.18 | 707.84 | 707 | 656.43 | 19276 | DNA topoisomerase 3-alpha (EC 5.6.2.1) (DNA topoisomerase III alpha) | TOP3A TOP3 | chromosome separation [GO:0051304]; DNA recombination [GO:0006310]; DNA repair [GO:0006281]; DNA topological change [GO:0006265]; double-strand break repair via homologous recombination [GO:0000724]; meiotic cell cycle [GO:0051321]; mitochondrial DNA metabolic process [GO:0032042]; resolution of DNA recombination intermediates [GO:0071139] |
| 51.4 | 256.22 | 4.5 | 89.22 | 194.15 | 1.89 | 0.8 | 50.06 | 0 | 119.13 | 888.04 | 36.44 | 16.39 | 4759.15 | 1.32 | 9593.89 | 3162.78 | 42.88 | 19268.26 | Interleukin-17 receptor E-like protein (IL-17 receptor E-like) (IL-17RE-like) | IL17REL | |
| 4302.52 | 638.36 | 1331.94 | 737.88 | 717.85 | 1554.44 | 37.67 | 227.14 | 1186.98 | 1120.99 | 274.91 | 749.28 | 2307.81 | 971.97 | 1358.07 | 204.74 | 457.45 | 1068.3 | 19248.3 | F-BAR and double SH3 domains protein 2 (Carom) (Protein nervous wreck 1) (NWK1) (SH3 multiple domains protein 3) | FCHSD2 KIAA0769 SH3MD3 | clathrin-dependent endocytosis [GO:0072583]; membrane organization [GO:0061024]; neuromuscular synaptic transmission [GO:0007274]; positive regulation of actin filament polymerization [GO:0030838]; positive regulation of Arp2/3 complex-mediated actin nucleation [GO:2000601]; protein transport [GO:0015031]; regulation of actin filament polymerization [GO:0030833] |

| | | | | | | | | | | | | | | | | | | | | | |
|---|---|---|---|---|---|---|---|---|---|---|---|---|---|---|---|---|---|---|---|---|---|
| 1476.53 | 4520.22 | 173.51 | 174.34 | 2140.05 | 310.04 | 0.86 | 1403.73 | 23.47 | 223.27 | 474.43 | 109.29 | 225.22 | 4842.15 | 60.75 | 70.91 | 2935.79 | 75.41 | 19239.97 | EH domain-containing protein 4 (PAST homolog 2) (mPAST2) | Ehd4 Past2 | cell migration involved in sprouting angiogenesis [GO:0002042]; cellular response to growth factor stimulus [GO:0071363]; endocytic recycling [GO:0032456]; pinocytosis [GO:0006907]; protein homooligomerization [GO:0051260]; regulation of endocytosis [GO:0030100] |
| 426.27 | 252.99 | 350.33 | 232.22 | 256.05 | 526.49 | 6550.43 | 134.4 | 266.5 | 197.99 | 319.91 | 230.04 | 494.58 | 329.09 | 329.82 | 7621.82 | 413.87 | 299.98 | 19232.78 | Huntingtin-interacting protein 1-related protein (HIP1-related protein) | Hip1r | clathrin coat assembly [GO:0048268]; digestive system development [GO:0055123]; intrinsic apoptotic signaling pathway [GO:0097193]; membrane organization [GO:0061024]; negative regulation of actin filament polymerization [GO:0030837]; negative regulation of apoptotic process [GO:0043066]; negative regulation of Arp2/3 complex-mediated actin nucleation [GO:0034316]; positive regulation of clathrin coat assembly [GO:1905445]; positive regulation of clathrin-dependent endocytosis [GO:2000370]; positive regulation of epidermal growth factor receptor signaling pathway [GO:0045742]; positive regulation of mitochondrial outer membrane permeabilization involved in apoptotic signaling pathway [GO:1901030]; positive regulation of platelet-derived growth factor receptor-beta signaling pathway [GO:2000588]; postsynapse organization [GO:0099173]; protein stabilization [GO:0050821]; receptor-mediated [illegible] |

| | | | | | | | | | | | | | | | | | | | | | | |
|---|---|---|---|---|---|---|---|---|---|---|---|---|---|---|---|---|---|---|---|---|---|---|
| 100.65 | | 5224.48 | 512.22 | 272.72 | 3094.58 | 152.34 | 2.38 | 1847.08 | 153.94 | 116.38 | 1078.5 | 201.92 | 135.35 | 3776.1 | 247.14 | 24.63 | 2032.2 | 260.15 | 19232.76 | Frizzled-7-A (Fz-7-A) (Xfz7-A) | fzd7-a fz7-a | canonical Wnt signaling pathway [GO:0060070]; convergent extension involved in gastrulation [GO:0060027]; dorsal/ventral axis specification [GO:0009950]; mesoderm formation [GO:0001707]; non-canonical Wnt signaling pathway [GO:0035567]; positive regulation of transcription by RNA polymerase II [GO:0045944]; protein localization [GO:0008104]; tissue morphogenesis [GO:0048729]; Wnt signaling pathway [GO:0016055] |
| 1785.1 | | 788.21 | 1039.87 | 522.61 | 501.43 | 1148.15 | 2328.98 | 227 | 495.7 | 684.53 | 259.68 | 1064.87 | 1438 | 731.87 | 678.08 | 3310.55 | 1355.79 | 823.32 | 19183.74 | Solute carrier family 2, facilitated glucose transporter member 5 (Fructose transporter) (Glucose transporter type 5, small intestine) (GLUT-5) | Slc2a5 Glut5 | cellular response to fructose stimulus [GO:0071332]; D-glucose import [GO:0046323]; D-glucose transmembrane transport [GO:1904659]; dehydroascorbic acid transport [GO:0070837]; fructose import across plasma membrane [GO:1990539]; fructose transmembrane transport [GO:0015755]; metanephric proximal tubule development [GO:0072237]; regulation of systemic arterial blood pressure mediated by a chemical signal [GO:0003044]; response to fructose [GO:0009750] |
| 1237.16 | | 1242.05 | 1408.54 | 470.91 | 1207.85 | 1628.32 | 1499.76 | 680.97 | 807.28 | 373.54 | 467.91 | 1087.18 | 1043.35 | 1940 | 763.43 | 1349.7 | 1090.95 | 825.28 | 19124.18 | Fibulin-2 (FIBL-2) | FBLN2 | positive regulation of cell-substrate adhesion [GO:0010811] |

| | | | | | | | | | | | | | | | | | | | | | |
|---|---|---|---|---|---|---|---|---|---|---|---|---|---|---|---|---|---|---|---|---|---|
| 2084.51 | 1064.26 | 1441.67 | 964.79 | 1485.92 | 1575.98 | 1936.2 | 598.84 | 330.34 | 557.27 | 616.77 | 654.07 | 1202.74 | 1335.07 | 428.8 | 1346.46 | 1074.18 | 356.44 | 19054.31 | CXXC-type zinc finger protein 4 | cxxc4 | Wnt signaling pathway [GO:0016055]; zygotic specification of dorsal/ventral axis [GO:0007352] |
| 2218.33 | 750.24 | 2039.35 | 530.98 | 1405.65 | 1627.48 | 29.66 | 443.32 | 826.85 | 1027.7 | 647.5 | 1585.51 | 1155.31 | 568.04 | 927.58 | 285.2 | 2139.01 | 833.78 | 19041.49 | tRNA N(3)-cytidine methyltransferase METTL6 (EC 2.1.1.-) (Methyltransferase-like protein 6) (hMETTL6) | METTL6 | tRNA methylation [GO:0030488]; tRNA modification [GO:0006400] |
| 89.22 | 16.21 | 11.63 | 376.33 | 19.8 | 26.72 | 10394.01 | 5.85 | 3.28 | 127.38 | 234.26 | 14.15 | 275.7 | 22.37 | 24.31 | 7159.69 | 185.38 | 4.59 | 18990.88 | Carboxyl-terminal PDZ ligand of neuronal nitric oxide synthase protein (C-terminal PDZ ligand of neuronal nitric oxide synthase protein) (Nitric oxide synthase 1 adaptor protein) | Nos1ap Capon Kiaa0464 | postsynaptic actin cytoskeleton organization [GO:0098974]; regulation of heart rate by chemical signal [GO:0003062]; regulation of ventricular cardiac muscle cell membrane repolarization [GO:0060307] |
| 812.79 | 349.02 | 1720.27 | 355.1 | 2543.76 | 5801.21 | 6.26 | 446.89 | 230.15 | 285.25 | 300.38 | 417.95 | 1212.17 | 670.44 | 488.85 | 316.65 | 1358.94 | 1631.4 | 18947.48 | Transient receptor potential cation channel subfamily M member 5 | trpm5 | |

| | | | | | | | | | | | | | | | | | | | | | |
|---|---|---|---|---|---|---|---|---|---|---|---|---|---|---|---|---|---|---|---|---|---|
| 2041.72 | 1329.56 | 583.28 | 331.4 | 1528.44 | 629.48 | 393.24 | 895.3 | 237.5 | 429.58 | 594.26 | 302.32 | 2638.38 | 3100.99 | 776.15 | 538.51 | 2066.37 | 496.82 | 18913.3 | Multiple C2 and transmembrane domain-containing protein 1 | Mctp1 | negative regulation of cell migration [GO:0030336]; negative regulation of endocytosis [GO:0045806]; negative regulation of response to oxidative stress [GO:1902883] |
| 568.21 | 519.72 | 1099.66 | 263.06 | 477.47 | 1066.13 | 1755.65 | 149.15 | 2538.32 | 182.05 | 261.05 | 1327.47 | 1268.32 | 858.35 | 2273.33 | 980.7 | 475.63 | 2814.66 | 18878.93 | Receptor-type tyrosine-protein phosphatase C (EC 3.1.3.48) (Leukocyte common antigen) (L-CA) (T200) (CD antigen CD45) | PTPRC CD45 | alpha-beta T cell proliferation [GO:0046633]; B cell differentiation [GO:0030183]; B cell proliferation [GO:0042100]; B cell receptor signaling pathway [GO:0050853]; bone marrow development [GO:0048539]; cell cycle phase transition [GO:0044770]; cell surface receptor signaling pathway [GO:0007166]; defense response to virus [GO:0051607]; dephosphorylation [GO:0016311]; DN2 thymocyte differentiation [GO:1904155]; extrinsic apoptotic signaling pathway [GO:0097191]; gamma-delta T cell differentiation [GO:0042492]; hematopoietic progenitor cell differentiation [GO:0002244]; heterotypic cell-cell adhesion [GO:0034113]; leukocyte cell-cell adhesion [GO:0007159]; MAPK cascade [GO:0000165]; natural killer cell differentiation [GO:0001779]; negative regulation of cell adhesion involved in substrate-bound cell migration [GO:0006933]; negative regulation of cytokine-mediated signaling pathway |

| | | | | | | | | | | | | | | | | | | | | | |
|---|---|---|---|---|---|---|---|---|---|---|---|---|---|---|---|---|---|---|---|---|---|
| 1068.08 | 868.1 | 1013.05 | 420.5 | 931.34 | 1423.66 | 2946.74 | 387.12 | 1288.78 | 368.62 | 275.86 | 969.91 | 1386.85 | 941.42 | 1252.07 | 1201.38 | 623 | 1461.72 | 18828.2 | Mediator of RNA polymerase II transcription subunit 13-like (Mediator complex subunit 13-like) (Thyroid hormone receptor-associated protein 2) (Thyroid hormone receptor-associated protein complex 240 kDa component-like) | MED13L KIAA1025 PROSIT240 THRAP2 TRAP240L | positive regulation of transcription by RNA polymerase II [GO:0045944] |
| 0 | 0 | 0 | 0 | 0 | 0 | 12322.49 | 0.79 | 0 | 42.32 | 0 | 4.57 | 0 | 0 | 0 | 6211.07 | 210.93 | 16.38 | 18808.55 | RNA-directed DNA polymerase homolog (EC 2.7.7.48) (Reverse transcriptase homolog) | | |
| 1758.11 | 1430.91 | 1279.68 | 545.76 | 1809.46 | 2999.89 | 69.91 | 805.17 | 109.12 | 656.93 | 741.36 | 851.07 | 1513.01 | 1468.19 | 660.74 | 263.78 | 1270.86 | 548.44 | 18782.39 | Protein capicua homolog | Cic Kiaa0306 | brain development [GO:0007420]; learning [GO:0007612]; lung alveolus development [GO:0048286]; memory [GO:0007613]; negative regulation of DNA-templated transcription [GO:0045892]; negative regulation of transcription by RNA polymerase II [GO:0000122]; social behavior [GO:0035176]; transcription by RNA polymerase II [GO:0006366] |

| | | | | | | | | | | | | | | | | | | | | | |
|---|---|---|---|---|---|---|---|---|---|---|---|---|---|---|---|---|---|---|---|---|---|
| 1.79 | 0.23 | 925.89 | 0.88 | 1.31 | 13.1 | 9.98 | 0 | 4361.11 | 0.71 | 1.48 | 1805.01 | 5.66 | 3.89 | 5140.83 | 4.37 | 4.17 | 6499.36 | 18779.77 | Intraflagellar transport protein 140 homolog (WD and tetratricopeptide repeats protein 2) | Ift140 WDTC2 | cilium assembly [GO:0060271]; determination of left/right symmetry [GO:0007368]; embryonic brain development [GO:1990403]; embryonic camera-type eye development [GO:0031076]; embryonic cranial skeleton morphogenesis [GO:0048701]; embryonic digit morphogenesis [GO:0042733]; heart development [GO:0007507]; intraciliary retrograde transport [GO:0035721]; intraciliary transport [GO:0042073]; limb morphogenesis [GO:0035108]; neural tube patterning [GO:0021532]; non-motile cilium assembly [GO:1905515]; photoreceptor cell outer segment organization [GO:0035845]; protein localization to cilium [GO:0061512]; regulation of cilium assembly [GO:1902017]; regulation of smoothened signaling pathway [GO:0008589] |
| 591.86 | 1207.81 | 543.34 | 957.59 | 1217.9 | 486.51 | 274.52 | 1985.53 | 361.11 | 864.03 | 1405.61 | 1202.88 | 1724.32 | 3430.18 | 579.01 | 365.55 | 1224.49 | 329.29 | 18751.53 | Transmembrane protein 222 | TMEM222 C1orf160 | |
| 660.5 | 1745.94 | 903.56 | 234.42 | 1775.86 | 1484.97 | 603.33 | 1165.74 | 79.17 | 307.57 | 929.06 | 828.95 | 565.31 | 3205.6 | 723.97 | 871.21 | 2422.42 | 219.77 | 18727.35 | Transcription initiation factor TFIID subunit 8 (Protein taube nuss-like) (TBP-associated factor 8) | taf8 tbnl | cell differentiation [GO:0030154]; transcription initiation at RNA polymerase II promoter [GO:0006367] |
| 930.98 | 404.4 | 977.49 | 420.1 | 2236.51 | 1198.86 | 534.1 | 440.9 | 621.42 | 440.61 | 274.3 | 569.61 | 2241.06 | 3203.77 | 1438.31 | 528.28 | 1177.58 | 1084.76 | 18723.04 | WD repeat-containing protein 19 (Intraflagellar transport 144 homolog) | Wdr19 Ift144 Kiaa1638 | cell morphogenesis [GO:0000902]; cilium assembly [GO:0060271]; digestive system development [GO:0055123]; ear morphogenesis [GO:0042471]; embryonic camera-type eye development [GO:0031076]; embryonic cranial skeleton morphogenesis [GO:0048701]; embryonic limb morphogenesis [GO:0030326]; gonad development [GO:0008406]; in utero embryonic development [GO:0001701]; intraciliary retrograde transport [GO:0035721]; myotome development [GO:0061055]; nervous system process [GO:0050877]; protein localization to ciliary membrane [GO:1903441]; protein localization to cilium [GO:0061512]; protein-containing complex assembly [GO:0065003]; receptor clustering [GO:0043113]; smoothened signaling pathway [GO:0007224]; smoothened signaling pathway involved in dorsal/ventral neural tube patterning [GO:0060831] |

| | | | | | | | | | | | | | | | | | | | | | |
|---|---|---|---|---|---|---|---|---|---|---|---|---|---|---|---|---|---|---|---|---|---|
| 2.06 | 10.8 | 1962.67 | 3 | 0.71 | 14.55 | 0 | 0.56 | 6318.07 | 3.84 | 1.36 | 1768.5 | 12.01 | 8.45 | 2928.96 | 2.51 | 3.3 | 5673.53 | 18714.88 | BMP/retinoic acid-inducible neural-specific protein 3 | Brinp3 Fam5c | cellular response to retinoic acid [GO:0071300]; central nervous system neuron differentiation [GO:0021953]; exploration behavior [GO:0035640]; multicellular organism growth [GO:0035264]; negative regulation of mitotic cell cycle [GO:0045930]; nervous system development [GO:0007399]; positive regulation of neuron differentiation [GO:0045666]; social behavior [GO:0035176] |
| 1389.16 | 1214.37 | 1160.67 | 579.14 | 1338.59 | 1657.74 | 2143.33 | 325.72 | 411.79 | 328.33 | 492.31 | 1211.4 | 1320.86 | 1574.16 | 570.25 | 1474.35 | 970.9 | 524.17 | 18687.24 | Armadillo repeat-containing protein 6 | Armc6 | hematopoietic progenitor cell differentiation [GO:0002244] |
| 25.19 | 0 | 0 | 456.77 | 2.68 | 30.5 | 8220.72 | 4.14 | 0 | 45.38 | 115.48 | 0 | 108.38 | 17.14 | 0 | 9355.4 | 263.85 | 5.33 | 18650.96 | Iduronate 2-sulfatase (EC 3.1.6.13) (Alpha-L-iduronate sulfate sulfatase) (Idursulfase) [Cleaved into: Iduronate 2-sulfatase 42 kDa chain;Iduronate 2-sulfatase 14 kDa chain] | IDS SIDS | dermatan sulfate proteoglycan catabolic process [GO:0030209]; glycosaminoglycan catabolic process [GO:0006027]; heparan sulfate proteoglycan catabolic process [GO:0030200] |
| 0.66 | 0.74 | 2139.82 | 0.41 | 0.68 | 424.6 | 6.59 | 0 | 31.43 | 0.46 | 0 | 15925.83 | 0.77 | 0 | 64.55 | 12.04 | 4.13 | 18.77 | 18631.48 | Nuclear prelamin A recognition factor | narf zgc:92186 | |

| | | | | | | | | | | | | | | | | | | | | | |
|---|---|---|---|---|---|---|---|---|---|---|---|---|---|---|---|---|---|---|---|---|---|
| 31.97 | 84.68 | 2721.11 | 45.14 | 43.97 | 3448.11 | 65.16 | 20.16 | 38.12 | 120.21 | 118.13 | 11301.23 | 16.22 | 52.65 | 76.44 | 87.32 | 105.15 | 234.65 | 18610.42 | Transcription factor HIVEP3 (Human immunodeficiency virus type I enhancer-binding protein 3 homolog) (KB-binding and recognition component) (Kappa-B and V(D)J recombination signal sequences-binding protein) (Kappa-binding protein 1) (KBP-1) (Recombinant component) (Schnurri-3) (Zinc finger protein ZAS3) | Hivep3 KBP1 Kiaa1555 Krc Rc shn3 Zas3 | positive regulation of DNA-templated transcription [GO:0045893]; skeletal muscle cell differentiation [GO:0035914] |
| 511.87 | 1807.12 | 847.6 | 324.31 | 1669.33 | 625.41 | 1549.98 | 762.91 | 552.58 | 232.1 | 488.94 | 707.85 | 587.59 | 3577.02 | 747.47 | 1590.49 | 1193.24 | 822.61 | 18598.42 | Fatty acid desaturase 6 (EC 1.14.19.-) | Fads6 | fatty acid biosynthetic process [GO:0006633] |
| 1053.66 | 771.75 | 1652.69 | 251.75 | 716.37 | 1356.94 | 432.81 | 438.77 | 2031.47 | 528.31 | 466.87 | 1340.97 | 865.87 | 1326.15 | 2327.54 | 312.67 | 639.83 | 2081.71 | 18596.13 | Serine/threonine-protein kinase SMG1 (smg-1) (EC 2.7.11.1) (Nonsense-mediated mRNA decay-associated PI3K-related kinase SMG1) | smg1 atx | nuclear-transcribed mRNA catabolic process, nonsense-mediated decay [GO:0000184] |
| 513.79 | 2510 | 1124.29 | 557.84 | 1632.17 | 886.47 | 820.35 | 1423.12 | 317.17 | 287.74 | 749.3 | 1421.62 | 775.82 | 1787.29 | 407.43 | 856.17 | 2152.29 | 363.79 | 18586.65 | GPI ethanolamine phosphate transferase 3, catalytic subunit (EC 2.-.-.-) (Phosphatidylinositol-glycan biosynthesis class O protein) (PIG-O) | Pigo | GPI anchor biosynthetic process [GO:0006506] |

| | | | | | | | | | | | | | | | | | | | | | |
|---|---|---|---|---|---|---|---|---|---|---|---|---|---|---|---|---|---|---|---|---|---|
| 1412.67 | 690.66 | 1437.93 | 369 | 918.57 | 1049.97 | 213.5 | 341.58 | 1906.23 | 411.65 | 244.66 | 1094.47 | 1560.5 | 1149.69 | 1663.3 | 603.56 | 739.46 | 2755.16 | 18562.56 | SAM domain-containing protein SAMSN-1 (SAM domain, SH3 domain and nuclear localization signals protein 1) (SH3 protein expressed in lymphocytes 2) (SH3-lymphocyte protein 2) (SLy2) | Samsn1 | negative regulation of adaptive immune response [GO:0002820]; negative regulation of B cell activation [GO:0050869]; negative regulation of peptidyl-tyrosine phosphorylation [GO:0050732] |
| 681.96 | 731.76 | 1454.9 | 416.23 | 847.98 | 1206.84 | 3700.26 | 239.46 | 938.35 | 218.11 | 330.05 | 823.04 | 992.35 | 1025.67 | 1178.44 | 2150.93 | 569.32 | 1051.73 | 18557.38 | AN1-type zinc finger protein 2B (Arsenite-inducible RNA-associated protein-like protein) (AIRAP-like protein) | ZFAND2B AIRAPL | proteasome-mediated ubiquitin-dependent protein catabolic process [GO:0043161]; protein targeting to ER [GO:0045047]; regulation of insulin-like growth factor receptor signaling pathway [GO:0043567]; SRP-dependent cotranslational protein targeting to membrane, translocation [GO:0006616] |

| | | | | | | | | | | | | | | | | | | | | | |
|---|---|---|---|---|---|---|---|---|---|---|---|---|---|---|---|---|---|---|---|---|---|
| 1053.95 | 1152.9 | 927.7 | 452.05 | 1278.33 | 1629.5 | 3225.84 | 532.26 | 617.95 | 346.37 | 425.34 | 766.3 | 1503 | 1225.24 | 903.31 | 1464.1 | 408.76 | 632.14 | 18545.04 | E3 ubiquitin-protein ligase MSL2 (EC 2.3.2.27) (Male-specific lethal-2 homolog) (MSL-2) | MSL2 | DNA damage response [GO:0006974]; epigenetic regulation of gene expression [GO:0040029]; positive regulation of DNA-templated transcription [GO:0045893]; protein monoubiquitination [GO:0006513]; protein ubiquitination [GO:0016567] |
| 1400.84 | 849.81 | 1557.29 | 276.68 | 741.43 | 1328.06 | 1867.31 | 332.21 | 1498.92 | 311.65 | 256.34 | 949.45 | 1029.77 | 1104.06 | 1492.06 | 1367.96 | 606.65 | 1574.3 | 18544.79 | Probable RNA-directed DNA polymerase from transposon BS (EC 2.7.7.49) (Reverse transcriptase) | RTase | DNA transposition [GO:0006313] |
| 21.58 | 24.65 | 488.59 | 10.86 | 12.4 | 5.61 | 44.28 | 1.72 | 6146.97 | 4.81 | 1.28 | 2004.95 | 60.09 | 22.86 | 3900.76 | 93.92 | 11.82 | 5674.73 | 18531.88 | Alkaline phosphatase, tissue-nonspecific isozyme (AP-TNAP) (TNS-ALP) (TNSALP) (EC 3.1.3.1) (Alkaline phosphatase liver/bone/kidney isozyme) (Phosphoamidase) (Phosphocreatine phosphatase) (EC 3.9.1.1) | ALPL | biomineral tissue development [GO:0031214]; bone mineralization [GO:0030282]; calcium ion homeostasis [GO:0055074]; cellular homeostasis [GO:0019725]; cementum mineralization [GO:0071529]; developmental process involved in reproduction [GO:0003006]; endochondral ossification [GO:0001958]; futile creatine cycle [GO:0140651]; inhibition of non-skeletal tissue mineralization [GO:0140928]; osteoblast differentiation [GO:0001649]; phosphate ion homeostasis [GO:0055062]; positive regulation of cold-induced thermogenesis [GO:0120162]; pyridoxal phosphate metabolic process [GO:0042822]; response to antibiotic [GO:0046677]; response to glucocorticoid [GO:0051384]; response to insulin [GO:0032868]; response to lipopolysaccharide [GO:0032496]; response to macrophage colony-stimulating factor [GO:0036005]; response to sodium phosphate [illegible] |

| | | | | | | | | | | | | | | | | | | | | | |
|---|---|---|---|---|---|---|---|---|---|---|---|---|---|---|---|---|---|---|---|---|---|
| 847.39 | 662.74 | 1000.88 | 292.07 | 854.99 | 689.77 | 398.59 | 287.32 | 2332.9 | 231.52 | 228.95 | 1159.5 | 1249.73 | 1608.73 | 2564.63 | 681.5 | 729.73 | 2709.49 | 18530.43 | Retrovirus-related Pol polyprotein from type-2 retrotransposable element R2DM (Retrovirus-related Pol polyprotein from type II retrotransposable element R2DM) [Includes: Protease (EC 3.4.23.-);Reverse transcriptase (EC 2.7.7.49);Endonuclease] | pol | proteolysis [GO:0006508] |
| 106.13 | 12.73 | 16.55 | 310.04 | 13.68 | 40.93 | 9070.8 | 0 | 137.2 | 145.01 | 145.82 | 97.29 | 1051.29 | 215.47 | 809.83 | 5790.69 | 227.69 | 331.96 | 18523.11 | Endophilin-A2 (Endophilin-2) (SH3 domain protein 2B) (SH3 domain-containing GRB2-like protein 1) (SH3p8) | Sh3gl1 Sh3p8 | endocytosis [GO:0006897]; modulation of excitatory postsynaptic potential [GO:0098815]; positive regulation of synaptic vesicle endocytosis [GO:1900244]; regulation of synaptic vesicle endocytosis [GO:1900242] |

| | | | | | | | | | | | | | | | | | | | | | |
|---|---|---|---|---|---|---|---|---|---|---|---|---|---|---|---|---|---|---|---|---|---|
| 1793.63 | 648.42 | 1836.97 | 436.03 | 578.48 | 980.08 | 1066.66 | 249.02 | 2057.31 | 516.11 | 304.89 | 812.48 | 1016.05 | 1545.7 | 1545.5 | 812.78 | 509.92 | 1776.82 | 18486.85 | Hypoxanthine-guanine phosphoribosyltransferase (HGPRT) (HGPRTase) (EC 2.4.2.8) | HPRT1 HPRT | GMP catabolic process [GO:0046038]; guanine salvage [GO:0006178]; hypoxanthine metabolic process [GO:0046100]; hypoxanthine salvage [GO:0043103]; IMP metabolic process [GO:0046040]; IMP salvage [GO:0032264]; positive regulation of dopamine metabolic process [GO:0045964]; purine nucleotide biosynthetic process [GO:0006164]; purine ribonucleoside salvage [GO:0006166] |
| 970.98 | 1050.22 | 1366.67 | 331.29 | 1102.1 | 1491.97 | 1129.23 | 1730.19 | 715.66 | 427.5 | 683.37 | 596.63 | 1128.97 | 1797.09 | 1197.62 | 968.89 | 826.84 | 936.23 | 18451.45 | Aldo-keto reductase family 1 member D1 (EC 1.3.1.3) (3-oxo-5-beta-steroid 4-dehydrogenase) (Delta(4)-3-ketosteroid 5-beta-reductase) (Delta(4)-3-oxosteroid 5-beta-reductase) | AKR1D1 SRD5B1 | androgen metabolic process [GO:0008209]; bile acid biosynthetic process [GO:0006699]; bile acid catabolic process [GO:0030573]; C21-steroid hormone metabolic process [GO:0008207]; cholesterol catabolic process [GO:0006707]; digestion [GO:0007586] |
| 722.8 | 789.84 | 654.6 | 172 | 774.77 | 1087.69 | 628.03 | 145.62 | 2532.41 | 214.7 | 141.1 | 1427.91 | 583.92 | 543.36 | 3560.05 | 378.69 | 282.07 | 3790.43 | 18429.99 | YLP motif-containing protein 1 (Nuclear protein ZAP3) (ZAP113) | YLPM1 C14orf170 ZAP3 | regulation of telomere maintenance [GO:0032204] |

| | | | | | | | | | | | | | | | | | | | | | |
|---|---|---|---|---|---|---|---|---|---|---|---|---|---|---|---|---|---|---|---|---|---|
| 497.43 | 1661.39 | 621.76 | 297 | 1523.27 | 1492.63 | 1529.85 | 684.28 | 1060.01 | 169.49 | 348.63 | 2033.06 | 786.79 | 1402.76 | 890.06 | 918.68 | 1160.47 | 1311.37 | 18388.93 | Diacylglycerol kinase epsilon (DAG kinase epsilon) (EC 2.7.1.107) (Diglyceride kinase epsilon) (DGK-epsilon) | DGKE DAGK5 | diacylglycerol metabolic process [GO:0046339]; glycerolipid metabolic process [GO:0046486]; intracellular signal transduction [GO:0035556]; lipid phosphorylation [GO:0046834]; modulation of chemical synaptic transmission [GO:0050804]; phosphatidic acid biosynthetic process [GO:0006654]; phosphatidylinositol biosynthetic process [GO:0006661]; phospholipase C-activating G protein-coupled receptor signaling pathway [GO:0007200]; platelet activation [GO:0030168] |
| 4170.23 | 1518.99 | 2025.16 | 446.87 | 2374.22 | 821.26 | 7.93 | 527.92 | 143.65 | 620.69 | 937.02 | 139.63 | 417.01 | 2225.2 | 89.96 | 290.74 | 1420.92 | 120.63 | 18298.03 | Platelet endothelial cell adhesion molecule (PECAM-1) (CD antigen CD31) | Pecam1 Pecam Pecam-1 | angiogenesis [GO:0001525]; bicellular tight junction assembly [GO:0070830]; cell adhesion [GO:0007155]; cell surface receptor signaling pathway [GO:0007166]; cellular response to mechanical stimulus [GO:0071260]; detection of mechanical stimulus [GO:0050982]; diapedesis [GO:0050904]; endothelial cell migration [GO:0043542]; endothelial cell morphogenesis [GO:0001886]; endothelial cell-matrix adhesion [GO:0090673]; establishment of endothelial barrier [GO:0061028]; glomerular endothelium development [GO:0072011]; homophilic cell adhesion via plasma membrane adhesion molecules [GO:0007156]; leukocyte cell-cell adhesion [GO:0007159]; monocyte extravasation [GO:0035696]; neutrophil extravasation [GO:0072672]; phagocytosis [GO:0006909]; positive regulation of cell migration [GO:0030335]; positive regulation of MAPK cascade [GO:0043410]; positive regulation of phosphatidylinositol 3- |

| | | | | | | | | | | | | | | | | | | | | | |
|---|---|---|---|---|---|---|---|---|---|---|---|---|---|---|---|---|---|---|---|---|---|
| 667.12 | 713.86 | 2365.84 | 301.31 | 577.49 | 899.77 | 1706.31 | 253.82 | 1598.41 | 149.5 | 263.49 | 1463.66 | 1002.71 | 1108.19 | 1549.74 | 1432.46 | 558.01 | 1680.28 | 18291.97 | Dual specificity protein phosphatase 26 (EC 3.1.3.16) (EC 3.1.3.48) | DUSP26 | negative regulation of ERK1 and ERK2 cascade [GO:0070373]; negative regulation of transcription by RNA polymerase II [GO:0000122]; positive regulation of cell adhesion [GO:0045785] |
| 149.41 | 3239.44 | 221.99 | 133.21 | 2624.67 | 341.38 | 11.34 | 2750.67 | 309.04 | 115.18 | 1126.64 | 242.49 | 190.7 | 1928.44 | 444.93 | 79.92 | 3709.81 | 661.21 | 18280.47 | Melanopsin-A (Opsin-4A) | opn4a | phototransduction [GO:0007602]; visual perception [GO:0007601] |
| 70.48 | 3.82 | 8.98 | 122.94 | 3.49 | 31.8 | 9865.24 | 1.38 | 5.39 | 77.72 | 333.73 | 5.25 | 28.88 | 3.33 | 13.22 | 7462.85 | 202.96 | 15.28 | 18256.74 | RIMS-binding protein 2 (RIM-BP2) | RIMBP2 RBP2 | neuromuscular synaptic transmission [GO:0007274] |
| 2944.61 | 767.42 | 855.13 | 511.73 | 590.2 | 1348.82 | 374.3 | 373.88 | 543.06 | 671.27 | 282.11 | 645.03 | 667.96 | 3521.14 | 1006.05 | 605.78 | 1578.99 | 960.89 | 18248.37 | Clusterin (51.5 kDa protein) [Cleaved into: Clusterin beta chain;Clusterin alpha chain] | CLUT64 | positive regulation of apoptotic process [GO:0043065]; positive regulation of proteasomal ubiquitin-dependent protein catabolic process [GO:0032436]; positive regulation of receptor-mediated endocytosis [GO:0048260]; protein stabilization [GO:0050821]; regulation of cell population proliferation [GO:0042127] |

| | | | | | | | | | | | | | | | | | | | | | |
|---|---|---|---|---|---|---|---|---|---|---|---|---|---|---|---|---|---|---|---|---|---|
| 1588.63 | 648.63 | 1559.06 | 443.68 | 666.35 | 884.16 | 153.27 | 380.47 | 2524.33 | 451.45 | 235.03 | 953.27 | 1550.72 | 1237.85 | 1791.83 | 329.64 | 616.95 | 2226.22 | 18241.54 | Polycystin-2-like protein 1 (Polycystin-2L1) (Polycystic kidney disease 2-like 1 protein) (Polycystin-2 homolog) | Pkd2l1 Trpp3 | cellular response to acidic pH [GO:0071468]; cellular response to pH [GO:0071467]; detection of chemical stimulus involved in sensory perception of sour taste [GO:0001581]; detection of chemical stimulus involved in sensory perception of taste [GO:0050912]; inorganic cation transmembrane transport [GO:0098662]; monoatomic cation transport [GO:0006812]; protein homotetramerization [GO:0051289]; protein tetramerization [GO:0051262]; response to water [GO:0009415]; smoothened signaling pathway [GO:0007224] |
| 1045.61 | 916.21 | 1285.56 | 347.07 | 1049.18 | 1229.76 | 2093.99 | 482.35 | 1189.47 | 386.36 | 341.23 | 870.22 | 1088.58 | 1038.37 | 1200.98 | 1493.6 | 716.32 | 1420.98 | 18195.84 | E3 ubiquitin-protein ligase TRIP12 (EC 2.3.2.26) (HECT-type E3 ubiquitin transferase TRIP12) (Thyroid receptor-interacting protein 12) (TR-interacting protein 12) (TRIP-12) | trip12 si:ch211-272f3.4 | DNA damage response [GO:0006974]; DNA repair [GO:0006281]; DNA repair-dependent chromatin remodeling [GO:0140861]; heterochromatin boundary formation [GO:0033696]; proteasome-mediated ubiquitin-dependent protein catabolic process [GO:0043161]; protein polyubiquitination [GO:0000209]; regulation of embryonic development [GO:0045995]; ubiquitin-dependent protein catabolic process [GO:0006511] |

| | | | | | | | | | | | | | | | | | | | | | |
|---|---|---|---|---|---|---|---|---|---|---|---|---|---|---|---|---|---|---|---|---|---|
| 0 | 22.36 | 4.4 | 4.51 | 67.21 | 1.19 | 4.09 | 1.46 | 4.61 | 0 | 18.43 | 0 | 0 | 14899 | 1.75 | 3.23 | 3144.65 | 15.36 | 18192.25 | Neurosecretory protein VGF [Cleaved into: Neuroendocrine regulatory peptide-1 (NERP-1);Neuroendocrine regulatory peptide-2 (NERP-2);VGF-derived peptide TLQP-21;VGF-derived peptide TLQP-62;Antimicrobial peptide VGF[554-577]] | VGF | carbohydrate homeostasis [GO:0033500]; defense response to bacterium [GO:0042742]; generation of precursor metabolites and energy [GO:0006091]; glucose homeostasis [GO:0042593]; insulin secretion [GO:0030073]; ovarian follicle development [GO:0001541]; regulation of synaptic plasticity [GO:0048167]; response to cAMP [GO:0051591]; response to cold [GO:0009409]; response to dietary excess [GO:0002021]; response to insulin [GO:0032868]; sexual reproduction [GO:0019953] |
| 1199.93 | 198.05 | 5381.95 | 308.55 | 238.62 | 1879.92 | 340.2 | 95.92 | 830.23 | 406.19 | 149.71 | 1062.92 | 958.22 | 247.05 | 1801.3 | 323.18 | 352.95 | 2396.1 | 18170.99 | Low-density lipoprotein receptor-related protein 1B (LRP-1B) (Low-density lipoprotein receptor-related protein-deleted in tumor) (LRP-DIT) | LRP1B LRPDIT | protein transport [GO:0015031]; receptor-mediated endocytosis [GO:0006898] |
| 33.34 | 2.25 | 321.47 | 5.86 | 2.81 | 230.63 | 0.33 | 3.05 | 4402.09 | 2.05 | 1.69 | 1695.81 | 52.52 | 2.75 | 4880.95 | 6.71 | 9.64 | 6516.9 | 18170.85 | Tetratricopeptide repeat protein 7B (TPR repeat protein 7B) (Tetratricopeptide repeat protein 7-like-1) (TPR repeat protein 7-like-1) | TTC7B TTC7L1 | phosphatidylinositol phosphate biosynthetic process [GO:0046854]; protein localization to plasma membrane [GO:0072659] |

| | | | | | | | | | | | | | | | | | | | | | |
|---|---|---|---|---|---|---|---|---|---|---|---|---|---|---|---|---|---|---|---|---|---|
| 16.89 | 0 | 0 | 484.16 | 0 | 7.09 | 4259.7 | 0 | 1.1 | 89.14 | 182.15 | 0 | 8.38 | 0 | 7.5 | 12783.15 | 308.4 | 18.07 | 18165.73 | E3 SUMO-protein ligase RanBP2 (EC 2.3.2.-) (358 kDa nucleoporin) (Nuclear pore complex protein Nup358) (Nucleoporin Nup358) (Ran-binding protein 2) (RanBP2) (p270) | RANBP2 | intracellular transport [GO:0046907]; mRNA transport [GO:0051028]; protein folding [GO:0006457]; protein sumoylation [GO:0016925]; protein transport [GO:0015031] |
| 969.19 | 2784.88 | 2674.24 | 772.84 | 1130.1 | 814.37 | 70.52 | 600.95 | 409.48 | 411.77 | 303.96 | 445.02 | 574.37 | 4423.39 | 724.93 | 234.05 | 391.63 | 417.43 | 18153.12 | Leucine-rich repeat-containing protein 58 | lrrc58 | |
| 248.98 | 96.26 | 1019.13 | 271.17 | 165.26 | 403.22 | 6957.28 | 58.45 | 464.86 | 145.56 | 286.36 | 530.54 | 268.49 | 263.01 | 373.86 | 5597.85 | 380.19 | 596.52 | 18126.99 | Ryanodine receptor 1 (RYR-1) (RyR1) (Skeletal muscle calcium release channel) (Skeletal muscle ryanodine receptor) (Skeletal muscle-type ryanodine receptor) (Type 1 ryanodine receptor) | RYR1 | calcium ion transmembrane transport [GO:0070588]; cellular response to caffeine [GO:0071313]; cellular response to calcium ion [GO:0071277]; intracellular calcium ion homeostasis [GO:0006874]; muscle contraction [GO:0006936]; ossification involved in bone maturation [GO:0043931]; outflow tract morphogenesis [GO:0003151]; protein homotetramerization [GO:0051289]; release of sequestered calcium ion into cytosol [GO:0051209]; release of sequestered calcium ion into cytosol by sarcoplasmic reticulum [GO:0014808]; skeletal muscle fiber development [GO:0048741]; skin development [GO:0043588]; striated muscle contraction [GO:0006941] |
| 124.34 | 172.78 | 1412.92 | 122.99 | 136.7 | 162.21 | 218.13 | 62.98 | 4395.77 | 79.82 | 81.35 | 1506.47 | 465.4 | 422.34 | 3573.08 | 188.91 | 160.84 | 4839.94 | 18126.97 | MOB kinase activator 1A (Mob1 alpha) (Mob1A) (Mob1 homolog 1B) (Mps one binder kinase activator-like 1B) | MOB1A C2orf6 MOB4B MOBK1B MOBKL1B | hippo signaling [GO:0035329] |

| | | | | | | | | | | | | | | | | | | | | | |
|---|---|---|---|---|---|---|---|---|---|---|---|---|---|---|---|---|---|---|---|---|---|
| 2312.22 | 801.28 | 1116.54 | 501.47 | 609.98 | 986.41 | 2064.9 | 230.41 | 164.8 | 499.69 | 317.7 | 421.06 | 3801.41 | 866.47 | 772.5 | 1653.39 | 602.45 | 379.55 | 18102.23 | Collectin-12 (Collectin placenta protein 1) (CL-P1) | COLEC12 CLP1 | phagocytosis, recognition [GO:0006910] |
| 59.74 | 44.11 | 1140.97 | 93.15 | 45.1 | 223.65 | 4961.04 | 17.99 | 2398.18 | 53.95 | 152.98 | 538.07 | 15.55 | 239.88 | 1258.69 | 4285.65 | 493.69 | 2069.05 | 18091.44 | Oxysterol-binding protein-related protein 3 (ORP-3) (OSBP-related protein 3) | OSBPL3 KIAA0704 ORP3 OSBP3 | bile acid biosynthetic process [GO:0006699] |
| 281.56 | 5266.68 | 458.07 | 372.76 | 2429.57 | 1406.92 | 2.01 | 986.2 | 52.45 | 130.08 | 438.64 | 432.74 | 252.01 | 3768.93 | 429.75 | 49.32 | 1248.34 | 71.5 | 18077.53 | V-type proton ATPase subunit H (V-ATPase subunit H) (V-ATPase 50/57 kDa subunits) (Vacuolar proton pump subunit H) (Vacuolar proton pump subunit SFD) | ATP6V1H | endocytosis [GO:0006897]; pH reduction [GO:0045851]; proton transmembrane transport [GO:1902600] |
| 638.15 | 522.44 | 2854.63 | 357.28 | 658.16 | 1000.63 | 848.62 | 311.84 | 1071.55 | 242.79 | 279.04 | 485.46 | 1045.13 | 2496.93 | 1336.95 | 833.05 | 1996.61 | 1090.91 | 18070.17 | cGMP-dependent protein kinase 2 (cGK 2) (cGK2) (EC 2.7.11.12) (cGMP-dependent protein kinase II) (cGKII) | PRKG2 PRKGR2 | negative regulation of chloride transport [GO:2001226]; positive regulation of chondrocyte differentiation [GO:0032332]; positive regulation of protein localization [GO:1903829]; protein localization to plasma membrane [GO:0072659]; protein phosphorylation [GO:0006468]; signal transduction [GO:0007165]; tetrahydrobiopterin metabolic process [GO:0046146] |

| | | | | | | | | | | | | | | | | | | | | | |
|---|---|---|---|---|---|---|---|---|---|---|---|---|---|---|---|---|---|---|---|---|---|
| 1.52 | 2852.04 | 3 | 1.96 | 2065.27 | 2.79 | 0 | 1640.75 | 7.53 | 0 | 527.35 | 32.77 | 0.87 | 9100.19 | 58.52 | 0.68 | 1639.64 | 96.01 | 18030.89 | Microtubule-associated protein tau (Neurofibrillary tangle protein) (Paired helical filament-tau) (PHF-tau) | MAPT TAU | microtubule cytoskeleton organization [GO:0000226]; neuron projection development [GO:0031175] |
| 680.51 | 625.56 | 730.05 | 295.61 | 717.29 | 881.24 | 1471.77 | 270.38 | 2045.91 | 273.39 | 402.54 | 768.43 | 827.19 | 1907.69 | 1787.02 | 1048.37 | 993.48 | 2284.95 | 18011.38 | Interferon alpha/beta receptor 1a (Membrane-associated type I interferon receptor) (mIFNAR1) (Type I interferon receptor 1a) | ifnar1a | type I interferon-mediated signaling pathway [GO:0060337] |
| 553.08 | 1149.54 | 1124.04 | 526.69 | 1825.63 | 1871.79 | 927.98 | 764.45 | 997.59 | 892.28 | 501.89 | 764.46 | 748.74 | 1452.74 | 1221.82 | 780.8 | 718.35 | 1183.03 | 18004.9 | Cytochrome P450 2D6 (EC 1.14.14.-) (CYPIID6) (Cholesterol 25-hydroxylase) (Cytochrome P450-DB1) (Debrisoquine 4-hydroxylase) | CYP2D6 CYP2DL1 | alkaloid catabolic process [GO:0009822]; alkaloid metabolic process [GO:0009820]; arachidonate metabolic process [GO:0019369]; cholesterol metabolic process [GO:0008203]; coumarin metabolic process [GO:0009804]; estrogen metabolic process [GO:0008210]; isoquinoline alkaloid metabolic process [GO:0033076]; long-chain fatty acid biosynthetic process [GO:0042759]; monoterpenoid metabolic process [GO:0016098]; negative regulation of organofluorine metabolic process [GO:0090350]; oxidative demethylation [GO:0070989]; retinol metabolic process [GO:0042572]; steroid metabolic process [GO:0008202]; xenobiotic catabolic process [GO:0042178]; xenobiotic metabolic process [GO:0006805] |

| | | | | | | | | | | | | | | | | | | | | | | |
|---|---|---|---|---|---|---|---|---|---|---|---|---|---|---|---|---|---|---|---|---|---|---|
| 921.95 | | 1286.09 | 1587.11 | 385.48 | 1209.63 | 1172.39 | 67.22 | 616.93 | 1500.73 | 330.47 | 411.13 | 1130.05 | 992.09 | 1773.71 | 1604.29 | 228.94 | 908.15 | 1868.43 | 17994.79 | High affinity cationic amino acid transporter 1 (CAT-1) (CAT1) (Ecotropic retroviral leukemia receptor homolog) (Ecotropic retrovirus receptor homolog) (Solute carrier family 7 member 1) (System Y+ basic amino acid transporter) | SLC7A1 ATRC1 ERR REC1L | amino acid import across plasma membrane [GO:0089718]; amino acid transport [GO:0006865]; L-amino acid transport [GO:0015807]; L-arginine import across plasma membrane [GO:0097638]; L-arginine transmembrane transport [GO:1903826]; L-histidine import across plasma membrane [GO:1903810]; L-ornithine transmembrane transport [GO:1903352]; lysine transport [GO:0015819]; ornithine transport [GO:0015822]; positive regulation of T cell proliferation [GO:0042102]; transport across blood-brain barrier [GO:0150104] |
| 711.09 | | 646.69 | 1988.13 | 201.72 | 562.6 | 947.57 | 376.87 | 313.41 | 2210.1 | 306.42 | 234.48 | 1244.85 | 1175.57 | 1066.09 | 2227.46 | 472.11 | 513.82 | 2761.29 | 17960.27 | Sodium-dependent neutral amino acid transporter SLC6A17 (Sodium-dependent neurotransmitter transporter NTT4) (Solute carrier family 6 member 17) | Slc6a17 Ntt4 | alanine transport [GO:0032328]; brain development [GO:0007420]; glycine transport [GO:0015816]; L-leucine transport [GO:0015820]; neurotransmitter transport [GO:0006836]; neutral amino acid transport [GO:0015804]; proline transport [GO:0015824]; protein catabolic process [GO:0030163]; sodium ion transport [GO:0006814] |

| | | | | | | | | | | | | | | | | | | | | | |
|---|---|---|---|---|---|---|---|---|---|---|---|---|---|---|---|---|---|---|---|---|---|
| 2291.54 | 1039.19 | 1365.83 | 875.19 | 839.95 | 1629.98 | 1303.41 | 269.47 | 685.37 | 326.96 | 118.36 | 429.34 | 1996.39 | 787.04 | 676.72 | 1244.69 | 1116.87 | 957.69 | 17953.99 | Pituitary adenylate cyclase-activating polypeptide type I receptor (PACAP type I receptor) (PACAP-R-1) (PACAP-R1) | Adcyap1r1 | adenylate cyclase-activating G protein-coupled receptor signaling pathway [GO:0007189]; cell differentiation [GO:0030154]; cell surface receptor signaling pathway [GO:0007166]; G protein-coupled receptor signaling pathway [GO:0007186]; spermatogenesis [GO:0007283] |
| 171.04 | 4001.63 | 5.25 | 296.01 | 2277.67 | 21.27 | 1.8 | 1271.84 | 14.52 | 130.25 | 1347.84 | 19.54 | 153.99 | 4193 | 10.47 | 247.84 | 3718.36 | 58.03 | 17940.35 | WD repeat-containing protein 74 | WDR74 | blastocyst formation [GO:0001825]; ribosomal large subunit biogenesis [GO:0042273]; RNA metabolic process [GO:0016070]; rRNA processing [GO:0006364] |

| | | | | | | | | | | | | | | | | | | | | | |
|---|---|---|---|---|---|---|---|---|---|---|---|---|---|---|---|---|---|---|---|---|---|
| 74.51 | 0 | 18.78 | 237.77 | 1.87 | 4.09 | 11955.43 | 0.74 | 4.35 | 93.46 | 311.35 | 3.87 | 21.65 | 0 | 1.85 | 4925.46 | 242.24 | 33.14 | 17930.56 | Fumarate hydratase, mitochondrial (Fumarase) (EC 4.2.1.2) | fh zgc:66253 | DNA damage response [GO:0006974]; DNA repair [GO:0006281]; fumarate metabolic process [GO:0006106]; malate metabolic process [GO:0006108]; positive regulation of double-strand break repair via nonhomologous end joining [GO:2001034]; regulation of arginine metabolic process [GO:0000821]; tricarboxylic acid cycle [GO:0006099]; urea cycle [GO:0000050] |
| 1274.94 | 3061.73 | 57.75 | 301.43 | 1354.33 | 83.81 | 5.17 | 3837.89 | 12.4 | 381.55 | 1102.09 | 82.37 | 775.6 | 3441.29 | 40.08 | 68.67 | 2001.21 | 39.61 | 17921.92 | Drebrin-like protein | DBNL | adaptive immune response [GO:0002250]; endocytosis [GO:0006897]; neuron projection morphogenesis [GO:0048812]; podosome assembly [GO:0071800]; positive regulation of axon extension [GO:0045773]; positive regulation of dendritic spine morphogenesis [GO:0061003]; postsynaptic actin cytoskeleton organization [GO:0098974]; regulation of actin filament polymerization [GO:0030833]; synapse assembly [GO:0007416] |
| 414.44 | 345.15 | 1450.12 | 143.2 | 347.8 | 1013 | 229.43 | 120.45 | 3405.35 | 140.55 | 142.51 | 1734.9 | 955.44 | 432.44 | 3032.49 | 168.78 | 210.73 | 3617.37 | 17904.15 | Vitamin K-dependent protein S | PROS1 PROS | blood coagulation [GO:0007596]; fibrinolysis [GO:0042730] |

| | | | | | | | | | | | | | | | | | | | | |
|---|---|---|---|---|---|---|---|---|---|---|---|---|---|---|---|---|---|---|---|---|
| 6.35 | 6272.76 | 9.16 | 1.93 | 2365.4 | 8.49 | 0.64 | 2065.9 | 10.93 | 1.28 | 1489.68 | 7.08 | 6.79 | 3405.94 | 12.84 | 1.68 | 2209.14 | 21.9 | 17897.89 | SH2 domain-containing protein 3C (Cas/HEF1-associated signal transducer) (SH2 domain-containing Eph receptor-binding protein 1) | Sh2d3c Chat Shep1 | signal transduction [GO:0007165]; small GTPase-mediated signal transduction [GO:0007264] |
| 6737.68 | 107.43 | 1499.54 | 1006.92 | 89.95 | 3803.38 | 14.17 | 55.46 | 351.95 | 1436.09 | 66.94 | 428.23 | 732.85 | 424.52 | 238.84 | 49.1 | 308.17 | 517.71 | 17868.93 | Heparan-sulfate 6-O-sulfotransferase 1-B (HS 6-OST-1B) (EC 2.8.2.-) | hs6st1b | |
| 901.99 | 1446.99 | 999.37 | 379.88 | 2085.4 | 1925.25 | 14.79 | 1125.86 | 68.18 | 357.76 | 616.88 | 509.85 | 2323.57 | 2416.61 | 1016.13 | 139.37 | 1119.39 | 419.19 | 17866.46 | Interphotoreceptor matrix proteoglycan 2 (Interphotoreceptor matrix proteoglycan of 200 kDa) (IPM 200) (Sialoprotein associated with cones and rods proteoglycan) (Spacrcan) | IMPG2 IPM200 | extracellular matrix organization [GO:0030198]; protein localization [GO:0008104]; retina morphogenesis in camera-type eye [GO:0060042]; visual perception [GO:0007601] |

| | | | | | | | | | | | | | | | | | | | | | |
|---|---|---|---|---|---|---|---|---|---|---|---|---|---|---|---|---|---|---|---|---|---|
| 632.84 | 1117.7 | 393.78 | 622.73 | 1254.21 | 3085.62 | 46.83 | 1154.82 | 765.41 | 658.19 | 1034.03 | 329.31 | 730.65 | 2628.71 | 1418.49 | 397.88 | 627.81 | 950.02 | 17849.03 | Probable ATP-dependent RNA helicase DDX5 (EC 3.6.4.13) (DEAD box protein 5) (RNA helicase p68) | DDX5 G17P1 HELR HLR1 | alternative mRNA splicing, via spliceosome [GO:0000380]; androgen receptor signaling pathway [GO:0030521]; BMP signaling pathway [GO:0030509]; epithelial to mesenchymal transition [GO:0001837]; estrogen receptor signaling pathway [GO:0030520]; intrinsic apoptotic signaling pathway by p53 class mediator [GO:0072332]; miRNA transcription [GO:0061614]; mRNA splicing, via spliceosome [GO:0000398]; mRNA transcription [GO:0009299]; myoblast differentiation [GO:0045445]; negative regulation of transcription by RNA polymerase II [GO:0000122]; nuclear-transcribed mRNA catabolic process [GO:0000956]; positive regulation of DNA damage response, signal transduction by p53 class mediator [GO:0043517]; primary miRNA processing [GO:0031053]; regulation of alternative mRNA splicing, via spliceosome [GO:0000381]; regulation of androgen receptor signaling pathway [GO:0060765]; regulation of osteoblast |
| 23.98 | 0 | 1366.81 | 93.24 | 2.23 | 72.94 | 7834.62 | 4.14 | 824.71 | 41.18 | 175.91 | 267.29 | 10.08 | 4.14 | 357.45 | 5806.39 | 141.1 | 781.95 | 17808.16 | RILP-like protein 2 (Rab-interacting lysosomal-like protein 2) | rilpl2 si:ch211-275j6.7 zgc:162589 | cilium assembly [GO:0060271]; epithelial cell morphogenesis [GO:0003382]; protein transport from ciliary membrane to plasma membrane [GO:1903445] |

| | | | | | | | | | | | | | | | | | | | | | |
|---|---|---|---|---|---|---|---|---|---|---|---|---|---|---|---|---|---|---|---|---|---|
| 1015.72 | 1205.2 | 1060.35 | 334.65 | 1050.03 | 890.97 | 364.07 | 470.99 | 1788.23 | 307.88 | 408.78 | 1058.35 | 938.02 | 1493.9 | 1829.93 | 483.24 | 840.27 | 2255.99 | 17796.57 | Histone acetyltransferase KAT6A (EC 2.3.1.48) (MOZ, YBF2/SAS3, SAS2 and TIP60 protein 3) (MYST-3) (Monocytic leukemia zinc finger protein) (Runt-related transcription factor-binding protein 2) (Zinc finger protein 220) | KAT6A MOZ MYST3 RUNXBP2 ZNF220 | cellular senescence [GO:0090398]; chromosome organization [GO:0051276]; myeloid cell differentiation [GO:0030099]; negative regulation of DNA-templated transcription [GO:0045892]; nucleosome assembly [GO:0006334]; positive regulation of DNA-templated transcription [GO:0045893]; positive regulation of gene expression [GO:0010628]; protein acetylation [GO:0006473]; regulation of developmental process [GO:0050793]; regulation of DNA-templated transcription [GO:0006355]; regulation of hemopoiesis [GO:1903706]; regulation of signal transduction by p53 class mediator [GO:1901796]; regulation of transcription by RNA polymerase II [GO:0006357] |
| 1264.73 | 1681.95 | 1677.2 | 97.62 | 548.97 | 465.98 | 36.52 | 517.31 | 2930.88 | 714.72 | 588.1 | 1280.56 | 141.13 | 546.95 | 657.18 | 265.35 | 592.81 | 3760.88 | 17768.84 | Myosin regulatory light chain 2, ventricular/cardiac muscle isoform (MLC-2) (MLC-2v) (Cardiac myosin light chain 2) (Myosin light chain 2, slow skeletal/ventricular muscle isoform) (MLC-2s/v) (Ventricular myosin light chain 2) | MYL2 MLC2 | cardiac muscle contraction [GO:0060048]; cardiac myofibril assembly [GO:0055003]; heart contraction [GO:0060047]; heart development [GO:0007507]; muscle cell fate specification [GO:0042694]; negative regulation of cell growth [GO:0030308]; positive regulation of the force of heart contraction [GO:0098735]; post-embryonic development [GO:0009791]; regulation of striated muscle contraction [GO:0006942]; regulation of the force of heart contraction [GO:0002026]; ventricular cardiac muscle tissue morphogenesis [GO:0055010] |

| | | | | | | | | | | | | | | | | | | | | | |
|---|---|---|---|---|---|---|---|---|---|---|---|---|---|---|---|---|---|---|---|---|---|
| 2662.62 | 659.31 | 973.08 | 568.75 | 590.29 | 1517.12 | 426.77 | 309.98 | 1010.18 | 897.19 | 354.59 | 811.67 | 2391.02 | 776.84 | 1173.87 | 502.26 | 684.48 | 1448.08 | 17758.1 | Laminin subunit alpha-3 (Epiligrin 170 kDa subunit) (E170) (Epiligrin subunit alpha) (Kalinin subunit alpha) (Laminin-5 subunit alpha) (Laminin-6 subunit alpha) (Laminin-7 subunit alpha) (Nicein subunit alpha) | LAMA3 LAMNA | cell-cell adhesion [GO:0098609]; endodermal cell differentiation [GO:0035987]; epidermis development [GO:0008544]; hemidesmosome assembly [GO:0031581]; regulation of cell adhesion [GO:0030155]; regulation of cell migration [GO:0030334]; regulation of embryonic development [GO:0045995] |
| 563.24 | 1410.22 | 1071.97 | 188.45 | 1224.23 | 1332.8 | 1026.9 | 492.67 | 1603.41 | 311.99 | 438.76 | 1447.88 | 716.4 | 1202.11 | 1518.06 | 611.09 | 785.55 | 1776.81 | 17722.54 | Intermembrane lipid transfer protein VPS13C (Vacuolar protein sorting-associated protein 13C) | VPS13C KIAA1421 | Golgi to endosome transport [GO:0006895]; lipid transport [GO:0006869]; mitochondrion organization [GO:0007005]; negative regulation of type 2 mitophagy [GO:1905090]; protein retention in Golgi apparatus [GO:0045053]; protein targeting to vacuole [GO:0006623]; response to insulin [GO:0032868] |

| | | | | | | | | | | | | | | | | | | | | | |
|---|---|---|---|---|---|---|---|---|---|---|---|---|---|---|---|---|---|---|---|---|---|
| 779.41 | 759.1 | 1219.42 | 277.73 | 888.61 | 1203.85 | 1373.78 | 412.47 | 1378.57 | 288.23 | 307.77 | 1077.76 | 849.67 | 986.08 | 1409.3 | 1716.22 | 660.72 | 2128.83 | 17717.52 | Nuclear mitotic apparatus protein 1 (Nuclear matrix protein-22) (NMP-22) (Nuclear mitotic apparatus protein) (NuMA protein) (SP-H antigen) | NUMA1 NMP22 NUMA | anastral spindle assembly [GO:0055048]; astral microtubule organization [GO:0030953]; cell division [GO:0051301]; establishment of mitotic spindle orientation [GO:0000132]; meiotic cell cycle [GO:0051321]; microtubule bundle formation [GO:0001578]; nucleus organization [GO:0006997]; positive regulation of BMP signaling pathway [GO:0030513]; positive regulation of chromosome segregation [GO:0051984]; positive regulation of chromosome separation [GO:1905820]; positive regulation of hair follicle development [GO:0051798]; positive regulation of intracellular transport [GO:0032388]; positive regulation of keratinocyte differentiation [GO:0045618]; positive regulation of microtubule polymerization [GO:0031116]; positive regulation of mitotic spindle elongation [GO:1902846]; positive regulation of protein localization to cell cortex [GO:1904778]; positive regulation of protein localization to spindle pole body |
| 2346.59 | 933.9 | 2424.17 | 518.45 | 768.09 | 1314.42 | 34.51 | 368.85 | 1243.44 | 633.28 | 354.05 | 1693.19 | 1785.62 | 591.19 | 796.46 | 227.49 | 442.17 | 1233.39 | 17709.26 | Annexin A1 (Annexin I) (Annexin-1) (Calpactin II) (Calpactin-2) (Chromobindin-9) (Lipocortin I) (Phospholipase A2 inhibitory protein) (p35) [Cleaved into: Annexin Ac2-26] | ANXA1 ANX1 LPC1 | actin cytoskeleton organization [GO:0030036]; adaptive immune response [GO:0002250]; alpha-beta T cell differentiation [GO:0046632]; arachidonate secretion [GO:0050482]; cell surface receptor signaling pathway [GO:0007166]; cellular response to glucocorticoid stimulus [GO:0071385]; cellular response to vascular endothelial growth factor stimulus [GO:0035924]; G protein-coupled receptor signaling pathway, coupled to cyclic nucleotide second messenger [GO:0007187]; granulocyte chemotaxis [GO:0071621]; inflammatory response [GO:0006954]; innate immune response [GO:0045087]; keratinocyte differentiation [GO:0030216]; monocyte chemotaxis [GO:0002548]; myoblast migration involved in skeletal muscle regeneration [GO:0014839]; negative regulation of apoptotic process [GO:0043066]; negative regulation of exocytosis [GO:0045920]; negative regulation of interleukin-8 production [GO:0032717]; negative regulation of T- |
| 742.79 | 573.47 | 1639.54 | 384.37 | 916.96 | 1635.6 | 6.87 | 1263.77 | 711.93 | 567.31 | 702.5 | 1679.73 | 783.07 | 1806.7 | 433.81 | 119.27 | 3207.43 | 499.19 | 17674.31 | Centrosomal protein of 89 kDa (Cep89) (Coiled-coil domain-containing protein 123) | cep89 ccdc123 zgc:63648 | chemical synaptic transmission [GO:0007268]; cilium assembly [GO:0060271]; mitochondrion organization [GO:0007005] |

| | | | | | | | | | | | | | | | | | | | | | |
|---|---|---|---|---|---|---|---|---|---|---|---|---|---|---|---|---|---|---|---|---|---|
| 22.61 | 2102.34 | 80.27 | 1.78 | 801.54 | 76.75 | 0 | 4076.29 | 0 | 32.59 | 864.17 | 470.12 | 57.07 | 1331.06 | 16.68 | 156.32 | 7452.01 | 104.78 | 17646.38 | Doublesex- and mab-3-related transcription factor A2 (Doublesex- and mab-3-related transcription factor 5) | dmrta2 dmrt5 | germ cell development [GO:0007281]; sex differentiation [GO:0007548] |
| 904.26 | 864.93 | 629.32 | 357.36 | 614.5 | 925.55 | 3825.99 | 184.06 | 386.13 | 296.94 | 243.3 | 390.15 | 1208.13 | 1009.09 | 542.52 | 4112.91 | 599.96 | 546.56 | 17641.66 | Protocadherin gamma-B5 (PCDH-gamma-B5) | PCDHGB5 | cell adhesion [GO:0007155]; homophilic cell adhesion via plasma membrane adhesion molecules [GO:0007156]; nervous system development [GO:0007399] |
| 711.41 | 978.32 | 1132.94 | 337.15 | 1104.12 | 918.26 | 3101.84 | 401.81 | 784.34 | 275.16 | 383.91 | 651.24 | 793.44 | 1551.54 | 1054.49 | 1573.61 | 920.02 | 956.46 | 17630.06 | Tomoregulin-2 (TR-2) (Transmembrane protein with EGF-like and two follistatin-like domains) | TMEFF2 | cell differentiation [GO:0030154]; negative regulation of cell migration [GO:0030336]; negative regulation of integrin biosynthetic process [GO:0045720]; negative regulation of stress fiber assembly [GO:0051497]; wound healing, spreading of cells [GO:0044319] |

| | | | | | | | | | | | | | | | | | | | | | |
|---|---|---|---|---|---|---|---|---|---|---|---|---|---|---|---|---|---|---|---|---|---|
| 1108.37 | 769.75 | 721.8 | 456.8 | 1608.17 | 974.39 | 12.08 | 1247.38 | 156.77 | 656.6 | 902.45 | 947.09 | 2600.23 | 1959.07 | 1430.31 | 88.92 | 1044 | 932.27 | 17616.45 | DCN1-like protein 2 (DCNL2) (DCUN1 domain-containing protein 2) (Defective in cullin neddylation protein 1-like protein 2) | DCUN1D2 C13orf17 DCUN1L2 | positive regulation of protein neddylation [GO:2000436]; protein neddylation [GO:0045116]; regulation of protein neddylation [GO:2000434] |
| 774.85 | 420.91 | 1262.8 | 411.81 | 599.21 | 1573.09 | 490.98 | 204.53 | 1299.98 | 374.65 | 251.41 | 1258.81 | 1341.6 | 810.93 | 1385.49 | 1064.53 | 1118.57 | 2949.77 | 17593.92 | Cholesterol 25-hydroxylase-like protein (EC 1.14.99.38) | ch25h zgc:101688 | B cell chemotaxis [GO:0035754]; cholesterol metabolic process [GO:0008203]; defense response to virus [GO:0051607]; sterol biosynthetic process [GO:0016126] |
| 63.94 | 5836.07 | 181.05 | 470.77 | 2423.03 | 1741.25 | 5.43 | 1700.82 | 6.05 | 117.62 | 227.77 | 665.49 | 38.32 | 2540.06 | 10.37 | 66.58 | 1381.8 | 60.53 | 17536.95 | mRNA-decapping enzyme 1B (EC 3.6.1.62) | DCP1B | deadenylation-dependent decapping of nuclear-transcribed mRNA [GO:0000290]; deadenylation-independent decapping of nuclear-transcribed mRNA [GO:0031087]; nuclear-transcribed mRNA catabolic process, nonsense-mediated decay [GO:0000184] |

| | | | | | | | | | | | | | | | | | | | | | |
|---|---|---|---|---|---|---|---|---|---|---|---|---|---|---|---|---|---|---|---|---|---|
| 0 | 0 | 578.93 | 15.59 | 107.77 | 5742.61 | 13.25 | 28.61 | 48.3 | 20.98 | 7.37 | 328.31 | 0 | 82.39 | 10339.37 | 61.96 | 76.36 | 0 | 17451.8 | Thrombospondin-3a (Thbs3a) | thbs3a thbs3 tsp3 zgc:103461 | cell adhesion [GO:0007155]; mesenchymal to epithelial transition [GO:0060231]; somite development [GO:0061053] |
| 0 | 123.27 | 0 | 9.03 | 4036.8 | 14.34 | 2.45 | 2436.46 | 0 | 10.06 | 7539.26 | 0 | 0 | 2689.3 | 0 | 10.63 | 484.86 | 3.45 | 17359.91 | Apelin receptor A (Angiotensin II receptor-like 1a) (Angiotensin receptor-like 1a) (G-protein coupled receptor APJ A) | aplnra agtrl1 agtrl1a | angioblast cell migration from lateral mesoderm to midline [GO:0035479]; blood vessel development [GO:0001568]; cardiac muscle tissue development [GO:0048738]; cell migration [GO:0016477]; cell migration involved in gastrulation [GO:0042074]; coronary vasculature development [GO:0060976]; determination of heart left/right asymmetry [GO:0061371]; determination of left/right symmetry [GO:0007368]; determination of liver left/right asymmetry [GO:0071910]; endodermal cell differentiation [GO:0035987]; epiboly involved in gastrulation with mouth forming second [GO:0055113]; G protein-coupled receptor signaling pathway [GO:0007186]; gastrulation with mouth forming second [GO:0001702]; heart development [GO:0007507]; Kupffer's vesicle development [GO:0070121]; lymph vessel development [GO:0001945]; negative regulation of cAMP-mediated signaling [GO:0043951]; positive regulation of [illegible] |
| 1721.51 | 1281.1 | 514.05 | 947.94 | 3244.38 | 760.55 | 343.32 | 420.06 | 62.06 | 557.37 | 990.66 | 210.47 | 1618.59 | 3091.89 | 161.18 | 260.68 | 1120.96 | 41.45 | 17348.22 | Signal transducer and activator of transcription 4 | STAT4 | cell surface receptor signaling pathway via JAK-STAT [GO:0007259]; cytokine-mediated signaling pathway [GO:0019221]; defense response [GO:0006952]; interleukin-12-mediated signaling pathway [GO:0035722]; positive regulation of transcription by RNA polymerase II [GO:0045944]; regulation of cell population proliferation [GO:0042127]; regulation of transcription by RNA polymerase II [GO:0006357]; response to interleukin-6 [GO:0070741]; response to peptide hormone [GO:0043434]; T-helper 1 cell differentiation [GO:0045063] |

| | | | | | | | | | | | | | | | | | | | | | |
|---|---|---|---|---|---|---|---|---|---|---|---|---|---|---|---|---|---|---|---|---|---|
| 575.25 | 310.17 | 1018.71 | 214.46 | 254.28 | 747.45 | 1629.05 | 168.12 | 2772.26 | 199.02 | 157.67 | 1323.54 | 800.06 | 705.59 | 2241.69 | 1038.95 | 327.24 | 2843.31 | 17326.82 | Inositol hexakisphosphate and diphosphoinositol-pentakisphosphate kinase 2 (EC 2.7.4.24) (Diphosphoinositol pentakisphosphate kinase 2) (Histidine acid phosphatase domain-containing protein 1) (InsP6 and PP-IP5 kinase 2) (VIP1 homolog 2) (mmVIP2) | Ppip5k2 Hisppd1 Kiaa0433 Vip2 | inositol metabolic process [GO:0006020]; sensory perception of sound [GO:0007605] |
| 1126.22 | 490.59 | 1531.03 | 343.83 | 434.54 | 816.59 | 1525.64 | 170.44 | 1827.16 | 381.3 | 252.52 | 1068.65 | 1183.36 | 625.69 | 1751.95 | 1102.11 | 423.25 | 2241.77 | 17296.64 | Exosome complex component RRP42 (Exosome component 7) (Ribosomal RNA-processing protein 42) (p8) | EXOSC7 KIAA0116 RRP42 | exonucleolytic trimming to generate mature 3'-end of 5.8S rRNA from tricistronic rRNA transcript (SSU-rRNA, 5.8S rRNA, LSU-rRNA) [GO:0000467]; nuclear mRNA surveillance [GO:0071028]; nuclear polyadenylation-dependent rRNA catabolic process [GO:0071035]; RNA catabolic process [GO:0006401]; RNA processing [GO:0006396]; rRNA catabolic process [GO:0016075]; rRNA processing [GO:0006364]; TRAMP-dependent tRNA surveillance pathway [GO:0071038]; U1 snRNA 3'-end processing [GO:0034473]; U4 snRNA 3'-end processing [GO:0034475]; U5 snRNA 3'-end processing [GO:0034476] |
| 1121.03 | 2388.9 | 951.53 | 283.06 | 2299.46 | 872.44 | 377.93 | 1165.56 | 236.72 | 281.74 | 1060.63 | 386.84 | 418.35 | 3590.56 | 162.07 | 297.85 | 1188.86 | 205.66 | 17289.19 | Ankyrin repeat and SOCS box protein 16 (ASB-16) | ASB16 | intracellular signal transduction [GO:0035556]; protein ubiquitination [GO:0016567] |

| | | | | | | | | | | | | | | | | | | | | | |
|---|---|---|---|---|---|---|---|---|---|---|---|---|---|---|---|---|---|---|---|---|---|
| 952.94 | 1066.63 | 871.71 | 545.71 | 1297.76 | 1186.35 | 1709.94 | 579.99 | 786.84 | 458.18 | 530.34 | 1075.86 | 1002.84 | 1633.75 | 710.83 | 1152.03 | 910.7 | 798.96 | 17271.36 | Histone H4 transcription factor (Histone nuclear factor P) (HiNF-P) (MBD2-interacting zinc finger protein) (Methyl-CpG-binding protein 2-interacting zinc finger protein) | Hinfp Mizf | cell cycle G1/S phase transition [GO:0044843]; DNA damage checkpoint signaling [GO:0000077]; DNA repair [GO:0006281]; DNA-templated transcription [GO:0006351]; establishment of protein localization [GO:0045184]; G1/S transition of mitotic cell cycle [GO:0000082]; in utero embryonic development [GO:0001701]; myoblast differentiation [GO:0045445]; negative regulation of DNA-templated transcription [GO:0045892]; negative regulation of gene expression [GO:0010629]; negative regulation of transcription by RNA polymerase II [GO:0000122]; positive regulation of DNA-templated transcription [GO:0045893]; positive regulation of gene expression [GO:0010628]; regulation of DNA-templated transcription [GO:0006355] |
| 1247.4 | 1320.68 | 6.16 | 2626.28 | 1125.17 | 16.14 | 9.07 | 931.6 | 21.42 | 3034.44 | 3266.41 | 7.97 | 596.02 | 708.21 | 9.78 | 720.64 | 1585.13 | 23.64 | 17256.16 | Myosin light chain 1, skeletal muscle isoform (LC-1) (LC1) (Myosin light chain alkali 1) (Myosin light chain A1) | | |
| 0.78 | 0.45 | 1157.52 | 2.3 | 0 | 5.46 | 1.4 | 0 | 4808.59 | 0 | 1.5 | 1813 | 2.29 | 1.62 | 3853.97 | 2.41 | 5.94 | 5598.56 | 17255.79 | Zinc finger CCHC domain-containing protein 8 (TRAMP-like complex RNA-binding factor ZCCHC8) | ZCCHC8 | mRNA 3'-end processing [GO:0031124]; RNA splicing [GO:0008380] |

| | | | | | | | | | | | | | | | | | | | | | |
|---|---|---|---|---|---|---|---|---|---|---|---|---|---|---|---|---|---|---|---|---|---|
| 76.12 | 66.41 | 267.42 | 407.08 | 45.7 | 412.25 | 8189.23 | 17.25 | 32.3 | 121.5 | 252.85 | 89.99 | 70.59 | 274.9 | 129.08 | 6473.02 | 285.97 | 25.19 | 17236.85 | Transmembrane protein 182 | Tmem182 | muscle organ development [GO:0007517]; myotube cell development involved in skeletal muscle regeneration [GO:0014906]; myotube differentiation involved in skeletal muscle regeneration [GO:0014908]; negative regulation of myoblast differentiation [GO:0045662]; negative regulation of myoblast fusion [GO:1901740] |
| 458.64 | 1378.54 | 713.44 | 229.49 | 2539.36 | 1231.32 | 381.14 | 1360.55 | 141.42 | 268.28 | 799.23 | 385.58 | 1009.39 | 3116.87 | 589.05 | 515 | 1596.04 | 523.23 | 17236.57 | Teashirt homolog 1 (Teashirt-like zinc finger protein) | tshz1 sdccag33 tsrt | regulation of transcription by RNA polymerase II [GO:0006357] |
| 5118.57 | 536.92 | 705 | 1059.61 | 595.18 | 982.65 | 847.61 | 244.93 | 155.73 | 883.27 | 317.66 | 291.52 | 2166.07 | 1293.74 | 282.99 | 848.5 | 732.9 | 164.92 | 17227.77 | Inositol hexakisphosphate kinase 1 (InsP6 kinase 1) (EC 2.7.4.21) (Inositol hexaphosphate kinase 1) | Ip6k1 Ihpk1 | inositol metabolic process [GO:0006020]; inositol phosphate biosynthetic process [GO:0032958]; negative regulation of cold-induced thermogenesis [GO:0120163]; phosphatidylinositol phosphate biosynthetic process [GO:0046854] |

| | | | | | | | | | | | | | | | | | | | | | |
|---|---|---|---|---|---|---|---|---|---|---|---|---|---|---|---|---|---|---|---|---|---|
| 1837.35 | 1005.42 | 1353.97 | 442.83 | 1141.39 | 1195.18 | 401.88 | 480.25 | 999.95 | 547.14 | 428.96 | 843.42 | 1346.94 | 1319.56 | 1159.26 | 461.78 | 934.84 | 1289.99 | 17190.11 | Protein-lysine N-methyltransferase SMYD4 (EC 2.1.1.-) (SET and MYND domain-containing protein 4) | smyd4 | determination of heart left/right asymmetry [GO:0061371]; heart development [GO:0007507]; heart looping [GO:0001947]; heart morphogenesis [GO:0003007]; methylation [GO:0032259] |
| 2339.65 | 655.56 | 571.27 | 817.78 | 2049.1 | 1827.14 | 13.62 | 843.26 | 40.24 | 499.86 | 331.59 | 273.51 | 2909.47 | 1869.22 | 765.45 | 237.6 | 915.46 | 94.47 | 17054.25 | Bone morphogenetic protein receptor type-1A (BMP type-1A receptor) (BMPR-1A) (EC 2.7.11.30) (Activin receptor-like kinase 3) (ALK-3) (Bone morphogenetic protein 4 receptor) (CD antigen CD292) | Bmpr1a | angiogenesis [GO:0001525]; anterior/posterior pattern specification [GO:0009952]; atrioventricular node cell development [GO:0060928]; atrioventricular valve development [GO:0003171]; BMP signaling pathway [GO:0030509]; cardiac conduction system development [GO:0003161]; cardiac right ventricle morphogenesis [GO:0003215]; cartilage development [GO:0051216]; cell differentiation [GO:0030154]; cellular response to BMP stimulus [GO:0071773]; cellular response to growth factor stimulus [GO:0071363]; central nervous system neuron differentiation [GO:0021953]; chondrocyte differentiation [GO:0002062]; developmental growth [GO:0048589]; dorsal aorta morphogenesis [GO:0035912]; dorsal/ventral axis specification [GO:0009950]; dorsal/ventral pattern formation [GO:0009953]; ectoderm development [GO:0007398]; embryonic digit morphogenesis [GO:0042733]; embryonic morphogenesis |

| | | | | | | | | | | | | | | | | | | | | | |
|---|---|---|---|---|---|---|---|---|---|---|---|---|---|---|---|---|---|---|---|---|---|
| 560.91 | 527.06 | 1926.11 | 202.44 | 539.34 | 1340.27 | 1723.3 | 197.23 | 1394.13 | 211.93 | 211.02 | 1204.79 | 1044.1 | 512.38 | 1566.43 | 1628.59 | 537.65 | 1703.27 | 17030.95 | m7GpppN-mRNA hydrolase (EC 3.6.1.62) (Nucleoside diphosphate-linked moiety X motif 20) (Nudix motif 20) (mRNA-decapping enzyme 2) (hDpc) | DCP2 NUDT20 | deadenylation-dependent decapping of nuclear-transcribed mRNA [GO:0000290]; histone mRNA catabolic process [GO:0071044]; mRNA catabolic process [GO:0006402]; negative regulation of telomere maintenance via telomerase [GO:0032211]; nuclear-transcribed mRNA catabolic process, deadenylation-dependent decay [GO:0000288]; nuclear-transcribed mRNA catabolic process, nonsense-mediated decay [GO:0000184]; regulation of mRNA stability [GO:0043488]; regulation of telomerase RNA localization to Cajal body [GO:1904872] |
| 1791.59 | 355.24 | 1531.69 | 537.4 | 596.8 | 792.92 | 780.71 | 118.17 | 2116.85 | 609.89 | 194.36 | 973.46 | 1256.62 | 305.24 | 1646.89 | 834.7 | 596.05 | 1975.85 | 17014.43 | Polysialoglycoprotein (PSGP) (Apopolysialoglycoprotein) (apoPSGP) | | |
| 401.63 | 530.87 | 1923.38 | 156.07 | 497.11 | 1162.87 | 70.89 | 237.31 | 1974.09 | 227.45 | 319.96 | 1777.71 | 360.09 | 497.89 | 2655.35 | 132.65 | 270.44 | 3808.02 | 17003.78 | Zinc finger and BTB domain-containing protein 39 | ZBTB39 KIAA0352 | negative regulation of transcription by RNA polymerase II [GO:0000122]; regulation of cytokine production [GO:0001817]; regulation of immune system process [GO:0002682] |

| | | | | | | | | | | | | | | | | | | | | | |
|---|---|---|---|---|---|---|---|---|---|---|---|---|---|---|---|---|---|---|---|---|---|
| 897.1 | 843.31 | 1268.1 | 540.51 | 981.32 | 1397.39 | 693.57 | 276.38 | 911.51 | 429.41 | 602.78 | 1272.13 | 1207.95 | 2413.49 | 817.01 | 809.48 | 886.41 | 754.26 | 17002.11 | CMP-N-acetylneuraminate-beta-galactosamide-alpha-2,3-sialyltransferase 1 (Alpha 2,3-ST 1) (Beta-galactoside alpha-2,3-sialyltransferase 1) (EC 2.4.3.4) (Gal-NAc6S) (Gal-beta-1,3-GalNAc-alpha-2,3-sialyltransferase) (Monosialoganglioside sialyltransferase) (EC 2.4.3.2) (ST3Gal I) (ST3GalI) (ST3GalA.1) (ST3O) (Sialyltransferase 4A) (SIAT4-A) | ST3GAL1 SIAT4A | ganglioside biosynthetic process via lactosylceramide [GO:0010706]; N-acetylneuraminate metabolic process [GO:0006054]; protein glycosylation [GO:0006486]; protein N-linked glycosylation [GO:0006487]; sialylation [GO:0097503] |
| 1117.66 | 449.13 | 1383.65 | 327.86 | 642.94 | 1191.55 | 303.45 | 333.75 | 2121.06 | 398.4 | 287.14 | 1262.39 | 977.82 | 911.82 | 1739.12 | 409.46 | 630.99 | 2512.05 | 17000.24 | Neuronal acetylcholine receptor subunit beta-2 (GF-beta-2) | chrnb2 | acetylcholine receptor signaling pathway [GO:0095500]; behavioral response to nicotine [GO:0035095]; cognition [GO:0050890]; memory [GO:0007613]; nervous system process [GO:0050877]; response to hypoxia [GO:0001666]; signal transduction [GO:0007165]; synaptic transmission, cholinergic [GO:0007271]; visual learning [GO:0008542] |
| 626.54 | 850.79 | 1102.21 | 228.85 | 828.5 | 761.76 | 3610.77 | 388.22 | 1253 | 179.35 | 390.72 | 1200.57 | 569.45 | 1209.61 | 816.09 | 1827.79 | 457.36 | 663.81 | 16965.39 | Small integral membrane protein 7 | smim7 | |
| 1014.04 | 294.64 | 813.62 | 444.45 | 380.45 | 1281.75 | 3645.59 | 210.83 | 519.59 | 656.29 | 284.37 | 799.65 | 907.65 | 479.01 | 923.06 | 3015.78 | 603.18 | 682.45 | 16956.4 | Butyrophilin subfamily 1 member A1 (BT) | BTN1A1 BTN | regulation of cytokine production [GO:0001817]; T cell receptor signaling pathway [GO:0050852] |

| | | | | | | | | | | | | | | | | | | | | | |
|---|---|---|---|---|---|---|---|---|---|---|---|---|---|---|---|---|---|---|---|---|---|
| 8.99 | 4817.37 | 0 | 20.8 | 3042.57 | 19.83 | 0 | 2064.82 | 1.18 | 32.32 | 1213.41 | 0 | 3.49 | 4203.89 | 0 | 11.62 | 1494.88 | 2.87 | 16938.04 | Guanine nucleotide exchange factor DBS (DBL's big sister) (MCF2-transforming sequence-like protein) (OST oncogene) | Mcf2l Ost | intracellular signal transduction [GO:0035556]; positive regulation of transcription by RNA polymerase II [GO:0045944]; Rho protein signal transduction [GO:0007266] |
| 5708.67 | 469.89 | 1833.26 | 1041.27 | 483.74 | 348.84 | 21.75 | 266.13 | 753.06 | 1284.11 | 245.4 | 658.33 | 751.96 | 238.7 | 335.76 | 96.63 | 512.21 | 1887.72 | 16937.43 | Nuclear protein MDM1 | mdm1 si:ch211-266a5.6 | negative regulation of centriole replication [GO:0046600] |
| 0 | 0 | 542.39 | 2.21 | 0 | 70.28 | 2.24 | 0 | 1945.17 | 0 | 0 | 673.3 | 0 | 0 | 5675.89 | 0.96 | 2.49 | 8010.94 | 16925.87 | Sialin (H(+)/nitrate cotransporter) (H(+)/sialic acid cotransporter) (AST) (Solute carrier family 17 (Anion/sugar transporter), member 5) (Vesicular excitatory amino acid transporter) (VEAT) | Slc17a5 | neurotransmitter loading into synaptic vesicle [GO:0098700]; response to bacterium [GO:0009617]; sialic acid transport [GO:0015739] |
| 2073.2 | 1060.97 | 2098.92 | 606.64 | 1028.2 | 1174.9 | 773.56 | 327.76 | 833.19 | 460.01 | 289.14 | 828 | 990.45 | 1039.44 | 972.83 | 709.62 | 528.13 | 1113.67 | 16908.63 | Transcription elongation factor A protein 3 (Transcription elongation factor S-II protein 3) (Transcription elongation factor TFIIS.h) | TCEA3 TFIISH | regulation of transcription by RNA polymerase II [GO:0006357]; transcription elongation by RNA polymerase II [GO:0006368] |

| | | | | | | | | | | | | | | | | | | | | | |
|---|---|---|---|---|---|---|---|---|---|---|---|---|---|---|---|---|---|---|---|---|---|
| 363.31 | 254.62 | 951.5 | 289.19 | 241.73 | 500.18 | 5425.78 | 134.34 | 398.53 | 196.25 | 284.3 | 477.45 | 401.2 | 359.61 | 555.62 | 5167.21 | 327.67 | 574.77 | 16903.26 | Bridging integrator 3 | Bin3 | actin cortical patch localization [GO:0051666]; cell division [GO:0051301]; endocytosis [GO:0006897]; myoblast migration involved in skeletal muscle regeneration [GO:0014839]; plasma membrane tubulation [GO:0097320]; protein localization [GO:0008104]; regulation of lamellipodium assembly [GO:0010591]; skeletal muscle fiber development [GO:0048741]; skeletal muscle tissue regeneration [GO:0043403]; unidimensional cell growth [GO:0009826] |
| 165.43 | 1075.04 | 480.09 | 149.52 | 3458.44 | 164.34 | 16.66 | 505.86 | 237.54 | 95.27 | 415.8 | 240.75 | 260.61 | 5194.45 | 152.48 | 296.33 | 3735.88 | 255.6 | 16900.09 | Rhodopsin kinase GRK1 (RK) (EC 2.7.11.14) (G protein-coupled receptor kinase 1) | GRK1 RHOK | G protein-coupled opsin signaling pathway [GO:0016056]; protein autophosphorylation [GO:0046777]; regulation of G protein-coupled receptor signaling pathway [GO:0008277]; regulation of opsin-mediated signaling pathway [GO:0022400]; regulation of signal transduction [GO:0009966]; visual perception [GO:0007601] |
| 825.41 | 938.82 | 1526.42 | 402.89 | 855.06 | 898.22 | 37.92 | 465.06 | 1601.64 | 636.25 | 426.28 | 883 | 981.74 | 1727.6 | 1759.76 | 269.4 | 750.9 | 1886.88 | 16873.25 | Cubilin (Intrinsic factor-cobalamin receptor) | CUBN IFCR | cholesterol metabolic process [GO:0008203]; protein transport [GO:0015031] |

| | | | | | | | | | | | | | | | | | | | | | |
|---|---|---|---|---|---|---|---|---|---|---|---|---|---|---|---|---|---|---|---|---|---|
| 498.12 | 7.12 | 0.65 | 2155.65 | 0 | 38.6 | 4550.98 | 0 | 0 | 142.69 | 153.9 | 1.96 | 0 | 0 | 0 | 9025.17 | 223.92 | 14.51 | 16813.27 | Contactin-associated protein-like 4 (Cell recognition molecule Caspr4) | CNTNAP4 CASPR4 KIAA1763 | cell adhesion [GO:0007155]; regulation of grooming behavior [GO:2000821]; regulation of synaptic transmission, dopaminergic [GO:0032225]; regulation of synaptic transmission, GABAergic [GO:0032228] |
| 18.97 | 32.34 | 624.09 | 22.6 | 83.71 | 193.46 | 33.56 | 18.98 | 4953.97 | 15.72 | 41.28 | 1407.98 | 9.13 | 164.1 | 4247 | 78.31 | 117.7 | 4698.94 | 16761.84 | Reelin (EC 3.4.21.-) (Reeler protein) | Reln Rl | associative learning [GO:0008306]; axon guidance [GO:0007411]; brain development [GO:0007420]; cell adhesion [GO:0007155]; cell migration [GO:0016477]; cell morphogenesis [GO:0000902]; central nervous system development [GO:0007417]; cerebral cortex development [GO:0021987]; cerebral cortex tangential migration [GO:0021800]; dendrite development [GO:0016358]; forebrain development [GO:0030900]; glial cell differentiation [GO:0010001]; hippocampus development [GO:0021766]; interneuron migration [GO:1904936]; lateral motor column neuron migration [GO:0097477]; layer formation in cerebral cortex [GO:0021819]; learning [GO:0007612]; locomotory behavior [GO:0007626]; long-term memory [GO:0007616]; long-term synaptic potentiation [GO:0060291]; modulation of chemical synaptic transmission [GO:0050804]; motor neuron migration [GO:0097475]; neuron migration [GO:0001764]; NMDA glutamate [illegible] |

| | | | | | | | | | | | | | | | | | | | | | |
|---|---|---|---|---|---|---|---|---|---|---|---|---|---|---|---|---|---|---|---|---|---|
| 585.37 | 815.33 | 205.14 | 264.35 | 877.7 | 139.49 | 6321.86 | 340.69 | 151.98 | 182.49 | 407.91 | 139.06 | 551.88 | 1088.62 | 187.46 | 3707.97 | 578.41 | 188.58 | 16734.29 | LETM1 domain-containing protein LETM2, mitochondrial (LETM1 and EF-hand domain-containing protein 2) (Leucine zipper-EF-hand-containing transmembrane protein 1-like) | letm2 TEgg018g03.1 | |
| 0 | 0 | 631.24 | 0 | 0 | 8750.21 | 8.24 | 0 | 0 | 12.8 | 6.23 | 0 | 0 | 0 | 7290.7 | 8.66 | 9.97 | 0 | 16718.05 | Cyclic AMP-responsive element-binding protein 3-like protein 3-B (cAMP-responsive element-binding protein 3-like protein 3-B) [Cleaved into: Processed cyclic AMP-responsive element-binding protein 3-like protein 3-B] | creb3l3b creb3l3 si:dkey-110c1.2 | regulation of transcription by RNA polymerase II [GO:0006357] |

| | | | | | | | | | | | | | | | | | | | | | |
|---|---|---|---|---|---|---|---|---|---|---|---|---|---|---|---|---|---|---|---|---|---|
| 48.16 | 24.62 | 1.33 | 37.97 | 31.72 | 0.97 | 21.19 | 26.25 | 1.24 | 3.05 | 321.37 | 11.5 | 0 | 7772.41 | 0 | 1437.58 | 6897.77 | 72.72 | 16709.85 | Nesprin-2 (KASH domain-containing protein 2) (KASH2) (Nuclear envelope spectrin repeat protein 2) (Nucleus and actin connecting element protein) (Protein NUANCE) (Synaptic nuclear envelope protein 2) (Syne-2) | SYNE2 KIAA1011 NUA | centrosome localization [GO:0051642]; nuclear migration [GO:0007097]; nuclear migration along microfilament [GO:0031022]; nucleokinesis involved in cell motility in cerebral cortex radial glia guided migration [GO:0021817]; positive regulation of cell migration [GO:0030335]; regulation of cilium assembly [GO:1902017] |
| 612.42 | 887.29 | 1079.5 | 2652.33 | 1716.94 | 1212.78 | 88.56 | 1197.71 | 511.43 | 508.4 | 783.79 | 215.22 | 909.44 | 698.74 | 1192.78 | 541.69 | 1303.91 | 545.91 | 16658.84 | Inactive serine protease PAMR1 (Peptidase domain-containing protein associated with muscle regeneration 1) (Regeneration-associated muscle protease homolog) | PAMR1 RAMP | |
| 1526.27 | 126.11 | 1856.24 | 178.43 | 786.16 | 3095.8 | 13.28 | 78.59 | 164.99 | 649.88 | 385.18 | 2513.57 | 1173.38 | 2333.08 | 343.4 | 199.12 | 967.77 | 235.17 | 16626.42 | Kinesin heavy chain isoform 5A (EC 5.6.1.3) (Kinesin heavy chain neuron-specific 1) (Neuronal kinesin heavy chain) (NKHC) | Kif5a Kiaa4086 Kif5 Nkhc1 | anterograde axonal protein transport [GO:0099641]; anterograde dendritic transport of neurotransmitter receptor complex [GO:0098971]; vesicle-mediated transport [GO:0016192] |

| | | | | | | | | | | | | | | | | | | | | | |
|---|---|---|---|---|---|---|---|---|---|---|---|---|---|---|---|---|---|---|---|---|---|
| 786.55 | 553.34 | 3248.86 | 262.52 | 347.01 | 4427.44 | 326.47 | 179.52 | 186.2 | 330.9 | 239.59 | 1492.22 | 1491.49 | 377.25 | 1080.31 | 403.02 | 425.2 | 467.6 | 16625.49 | Interferon regulatory factor 4 (IRF-4) (Lymphocyte-specific interferon regulatory factor) (LSIRF) (NF-EM5) (PU.1 interaction partner) (Transcriptional activator PIP) | Irf4 Spip | chromatin remodeling [GO:0006338]; defense response to protozoan [GO:0042832]; myeloid dendritic cell differentiation [GO:0043011]; negative regulation of toll-like receptor signaling pathway [GO:0034122]; positive regulation of cold-induced thermogenesis [GO:0120162]; positive regulation of DNA-templated transcription [GO:0045893]; positive regulation of interleukin-10 production [GO:0032733]; positive regulation of interleukin-13 production [GO:0032736]; positive regulation of interleukin-2 production [GO:0032743]; positive regulation of interleukin-4 production [GO:0032753]; positive regulation of transcription by RNA polymerase II [GO:0045944]; T-helper 17 cell lineage commitment [GO:0072540] |
| 26.29 | 0.69 | 263.88 | 113.23 | 1.34 | 498.38 | 9126.62 | 1.49 | 24.37 | 49.92 | 188.3 | 305.36 | 8.51 | 0.84 | 126.96 | 5675.3 | 179.12 | 19.44 | 16610.04 | Malate dehydrogenase, cytoplasmic (EC 1.1.1.37) (Cytosolic malate dehydrogenase) | MDH1 RCJMB04_2g5 | gluconeogenesis [GO:0006094]; malate metabolic process [GO:0006108]; tricarboxylic acid cycle [GO:0006099] |
| 684.06 | 737.89 | 1676.84 | 336.06 | 859.43 | 1621.28 | 1263.82 | 318.51 | 854.97 | 322.96 | 408.77 | 1789.88 | 739.7 | 1498.47 | 816.66 | 1280.2 | 788.24 | 589.84 | 16587.58 | LanC-like protein 2 (Testis-specific adriamycin sensitivity protein) | LANCL2 GPR69B TASP | carbohydrate metabolic process [GO:0005975]; negative regulation of DNA-templated transcription [GO:0045892]; peptide modification [GO:0031179]; positive regulation of abscisic acid-activated signaling pathway [GO:0009789] |

| | | | | | | | | | | | | | | | | | | | | | |
|---|---|---|---|---|---|---|---|---|---|---|---|---|---|---|---|---|---|---|---|---|---|
| 891.95 | 851.5 | 1689.18 | 139.4 | 1042.53 | 172.49 | 153.97 | 746.29 | 2176.81 | 71.66 | 449.28 | 625.43 | 313.78 | 3389.03 | 1522.16 | 94.88 | 779.98 | 1458.07 | 16568.39 | Prosaposin receptor GPR37 (Endothelin B receptor-like protein 1) (ETBR-LP-1) (G-protein coupled receptor 37) (Parkin-associated endothelin receptor-like receptor) (PAELR) | GPR37 | adenylate cyclase-inhibiting G protein-coupled receptor signaling pathway [GO:0007193]; cellular response to reactive oxygen species [GO:0034614]; dendrite development [GO:0016358]; dopamine biosynthetic process [GO:0042416]; G protein-coupled receptor signaling pathway [GO:0007186]; locomotion involved in locomotory behavior [GO:0031987]; neuropeptide signaling pathway [GO:0007218]; positive regulation of dopamine metabolic process [GO:0045964]; positive regulation of MAPK cascade [GO:0043410] |
| 0 | 1.63 | 651.44 | 20.23 | 0 | 44.74 | 24.26 | 0.67 | 4934.81 | 0 | 18.04 | 3316.27 | 0 | 0 | 2264.58 | 0 | 16.45 | 5260.8 | 16553.92 | Pseudouridylate synthase TRUB1 (EC 5.4.99.-) (TruB pseudouridine synthase homolog 1) (tRNA pseudouridine 55 synthase TRUB1) (Psi55 synthase TRUB1) (EC 5.4.99.25) | TRUB1 PUS4 | mRNA processing [GO:0006397]; mRNA pseudouridine synthesis [GO:1990481]; positive regulation of pre-miRNA processing [GO:2000633]; tRNA modification [GO:0006400] |

| | | | | | | | | | | | | | | | | | | | | | |
|---|---|---|---|---|---|---|---|---|---|---|---|---|---|---|---|---|---|---|---|---|---|
| 172.22 | 276.99 | 2463.49 | 300.92 | 142.38 | 3297.87 | 21.08 | 82.32 | 22.86 | 316.14 | 290.96 | 8107.02 | 135.72 | 180.15 | 46.38 | 193.12 | 291.56 | 207.91 | 16549.09 | Sterol regulatory element-binding protein 2 (SREBP-2) (Sterol regulatory element-binding transcription factor 2) [Cleaved into: Processed sterol regulatory element-binding protein 2 (Transcription factor SREBF2)] | srebf2 SREBP2 zgc:158371 | cholesterol metabolic process [GO:0008203]; liver development [GO:0001889]; positive regulation of cholesterol storage [GO:0010886]; positive regulation of transcription by RNA polymerase II [GO:0045944]; regulation of Notch signaling pathway [GO:0008593] |
| 65.37 | 4730.53 | 62.41 | 17.72 | 5433.15 | 40.77 | 0 | 3749.26 | 51.23 | 6.33 | 206.12 | 32.15 | 52.59 | 900.75 | 71.08 | 5.05 | 1008.21 | 92.78 | 16525.5 | Ryanodine receptor 2 (RYR-2) (RyR2) (Cardiac muscle ryanodine receptor) (Cardiac muscle ryanodine receptor-calcium release channel) (Cardiac muscle-type ryanodine receptor) (Type 2 ryanodine receptor) | RYR2 | calcium ion transport [GO:0006816]; calcium-mediated signaling [GO:0019722]; cellular response to caffeine [GO:0071313]; embryonic heart tube morphogenesis [GO:0003143]; intracellular calcium ion homeostasis [GO:0006874]; release of sequestered calcium ion into cytosol by sarcoplasmic reticulum [GO:0014808]; striated muscle contraction [GO:0006941] |
| 1237.36 | 1129.37 | 342.03 | 862.24 | 2562.25 | 1493.44 | 56.39 | 906.51 | 157.06 | 605.76 | 444.16 | 492.94 | 936.92 | 2331.28 | 394.53 | 157.75 | 2083.49 | 321.88 | 16515.36 | Anosmin-1 (Adhesion molecule-like X-linked) (Kallmann syndrome protein) | ANOS1 ADMLX KAL KAL1 KALIG1 | axon guidance [GO:0007411]; cell adhesion [GO:0007155]; chemotaxis [GO:0006935]; neuron differentiation [GO:0030182] |

| | | | | | | | | | | | | | | | | | | | | | |
|---|---|---|---|---|---|---|---|---|---|---|---|---|---|---|---|---|---|---|---|---|---|
| 240.41 | 274.32 | 2085.48 | 98.94 | 283.64 | 968.77 | 590.71 | 157 | 2609.5 | 102.96 | 178.26 | 1436.05 | 228.84 | 336.11 | 2245.06 | 546.63 | 307.7 | 3814.35 | 16504.73 | Ubiquitin carboxyl-terminal hydrolase MINDY-1 (EC 3.4.19.12) (Deubiquitinating enzyme MINDY-1) (Protein FAM63A) | mindy1 fam63a si:ch211-210h11.5 | proteolysis [GO:0006508] |
| 885.39 | 76.54 | 1702.22 | 271.26 | 128.54 | 5285.04 | 10.93 | 76.88 | 1031.58 | 228.23 | 59.36 | 593.08 | 518.63 | 382.9 | 3937.29 | 34.53 | 144.83 | 1137.47 | 16504.7 | Microtubule-actin cross-linking factor 1, isoforms 6/7 (Uncharacterized protein KIAA0754) | MACF1 KIAA0754 | |
| 1074.29 | 1029.34 | 704.85 | 355.83 | 623.3 | 908.58 | 3218.44 | 437.6 | 743.09 | 327.3 | 302.82 | 474.27 | 1317.68 | 1020.02 | 654.32 | 1752.59 | 765.43 | 792.81 | 16502.56 | Cytochrome c oxidase assembly protein COX18, mitochondrial | Cox18 Oxa1l2 | mitochondrial cytochrome c oxidase assembly [GO:0033617]; protein insertion into mitochondrial inner membrane from matrix [GO:0032979]; protein insertion into mitochondrial membrane [GO:0051204]; respiratory chain complex IV assembly [GO:0008535] |

| | | | | | | | | | | | | | | | | | | | | | |
|---|---|---|---|---|---|---|---|---|---|---|---|---|---|---|---|---|---|---|---|---|---|
| 157.34 | 59.12 | 267.08 | 141.13 | 34.28 | 477.82 | 7261.8 | 20.19 | 25.32 | 180.59 | 312.42 | 322.04 | 182.04 | 31.78 | 120.05 | 6615.83 | 190.78 | 96.51 | 16496.12 | Glycerophosphodiester phosphodiesterase 1 (Glycerophosphoinositol glycerophosphodiesterase GDE1) (EC 3.1.4.44) (Lysophospholipase D GDE1) (EC 3.1.4.-) (Membrane-interacting protein of RGS16) | Gde1 Mir16 | ethanolamine metabolic process [GO:0006580]; G protein-coupled receptor signaling pathway [GO:0007186]; N-acylethanolamine metabolic process [GO:0070291]; phospholipid metabolic process [GO:0006644] |
| 775.94 | 1522.77 | 56.44 | 2764.22 | 2045.15 | 43.9 | 142.51 | 1222.27 | 57.4 | 1116.95 | 1661.92 | 90.24 | 879.28 | 1392.24 | 57.09 | 1032.92 | 1517.62 | 110.05 | 16488.91 | ADP-ribosylation factor-like protein 5A | ARL5A | intracellular protein transport [GO:0006886]; protein localization to Golgi membrane [GO:1903292]; vesicle-mediated transport [GO:0016192] |
| 146.38 | 521.18 | 1126.6 | 134.97 | 540.6 | 537.33 | 45.73 | 189.67 | 2669.44 | 90.85 | 184.67 | 1506.91 | 131.54 | 642.31 | 3625.05 | 92.85 | 531.97 | 3744.46 | 16462.51 | X-linked retinitis pigmentosa GTPase regulator (mRpgr) | Rpgr | cellular response to light stimulus [GO:0071482]; eye photoreceptor cell development [GO:0042462]; intraciliary transport [GO:0042073]; positive regulation of autophagy [GO:0010508]; protein localization to non-motile cilium [GO:0097499]; retina morphogenesis in camera-type eye [GO:0060042]; visual perception [GO:0007601] |

| | | | | | | | | | | | | | | | | | | | | | |
|---|---|---|---|---|---|---|---|---|---|---|---|---|---|---|---|---|---|---|---|---|---|
| 84.9 | 5757.17 | 1.01 | 113.54 | 2928.36 | 9.43 | 0 | 1038.68 | 0 | 17.77 | 477.64 | 0 | 35.7 | 2900 | 0 | 26.81 | 3050.06 | 7.15 | 16448.22 | WD repeat-containing protein 47 (Neuronal enriched MAP-interacting protein) (Nemitin) | WDR47 KIAA0893 | adult locomotory behavior [GO:0008344]; anterior commissure morphogenesis [GO:0021960]; autophagy [GO:0006914]; cerebral cortex radial glia-guided migration [GO:0021801]; corpus callosum development [GO:0022038]; detection of hot stimulus involved in thermoception [GO:0120168]; microtubule cytoskeleton organization [GO:0000226]; motor behavior [GO:0061744]; negative regulation of microtubule depolymerization [GO:0007026]; neural precursor cell proliferation [GO:0061351]; neuronal stem cell population maintenance [GO:0097150] |
| 11.94 | 9.8 | 227.43 | 3.16 | 7.59 | 20.1 | 1.26 | 3.36 | 5542.81 | 4.51 | 6.39 | 1279.13 | 15.02 | 19.28 | 3550.31 | 5.89 | 19.38 | 5708.43 | 16435.79 | A-type potassium channel modulatory protein DPP6 (DPPX) (Dipeptidyl aminopeptidase-like protein 6) (Dipeptidyl aminopeptidase-related protein) (Dipeptidyl peptidase 6) (Dipeptidyl peptidase IV-like protein) (Dipeptidyl peptidase VI) (DPP VI) | Dpp6 Dpp-6 | establishment of localization in cell [GO:0051649]; neuronal action potential [GO:0019228]; positive regulation of potassium ion transmembrane transport [GO:1901381]; potassium ion transmembrane transport [GO:0071805]; protein localization to plasma membrane [GO:0072659]; proteolysis [GO:0006508]; regulation of membrane potential [GO:0042391]; regulation of postsynaptic membrane potential [GO:0060078]; regulation of potassium ion transmembrane transport [GO:1901379]; regulation of potassium ion transport [GO:0043266] |

| | | | | | | | | | | | | | | | | | | | | | |
|---|---|---|---|---|---|---|---|---|---|---|---|---|---|---|---|---|---|---|---|---|---|
| 941.75 | 480.36 | 987.47 | 358.01 | 449.44 | 1080.12 | 3055.99 | 257 | 809.1 | 399.06 | 246.93 | 662.22 | 1538.63 | 662.93 | 1048.16 | 1830.54 | 561.91 | 1046.41 | 16416.03 | Microtubule-associated protein RP/EB family member 1 (APC-binding protein EB1) (End-binding protein 1) (EB1) | Mapre1 | attachment of mitotic spindle microtubules to kinetochore [GO:0051315]; cell division [GO:0051301]; cell migration [GO:0016477]; establishment of mitotic spindle orientation [GO:0000132]; microtubule bundle formation [GO:0001578]; microtubule polymerization [GO:0046785]; negative regulation of microtubule polymerization [GO:0031115]; non-motile cilium assembly [GO:1905515]; positive regulation of microtubule polymerization [GO:0031116]; protein localization to astral microtubule [GO:1902888]; protein localization to centrosome [GO:0071539]; protein localization to microtubule [GO:0035372]; protein localization to mitotic spindle [GO:1902480]; regulation of microtubule polymerization or depolymerization [GO:0031110]; spindle assembly [GO:0051225] |
| 4241.36 | 665.69 | 555.62 | 1079.12 | 1642.93 | 652.45 | 26.46 | 559.27 | 66.03 | 1563.96 | 507.19 | 612.26 | 1464.45 | 709.91 | 258.49 | 434.04 | 1212.13 | 160.79 | 16412.15 | ADP/ATP translocase 2 (ADP,ATP carrier protein 2) (Adenine nucleotide translocator 2) (ANT 2) (Solute carrier family 25 member 5) [Cleaved into: ADP/ATP translocase 2, N-terminally processed] | Slc25a5 Ant2 | adaptive thermogenesis [GO:1990845]; adenine nucleotide transport [GO:0051503]; B cell differentiation [GO:0030183]; cellular response to leukemia inhibitory factor [GO:1990830]; chromosome segregation [GO:0007059]; erythrocyte differentiation [GO:0030218]; mitochondrial ADP transmembrane transport [GO:0140021]; mitochondrial ATP transmembrane transport [GO:1990544]; negative regulation of mitochondrial outer membrane permeabilization involved in apoptotic signaling pathway [GO:1901029]; positive regulation of cell population proliferation [GO:0008284]; positive regulation of mitophagy [GO:1901526]; regulation of mitochondrial membrane permeability [GO:0046902] |
| 0.46 | 0.48 | 566.67 | 2.31 | 0.46 | 3.49 | 32.02 | 0.52 | 5277.4 | 1.01 | 2.23 | 1659.87 | 0 | 2.64 | 3836.35 | 27.04 | 3.81 | 4979.36 | 16396.12 | Large ribosomal subunit protein uL2m (39S ribosomal protein L2, mitochondrial) (L2mt) (MRP-L2) | MRPL2 CGI-22 | mitochondrial translation [GO:0032543] |

| | | | | | | | | | | | | | | | | | | | | | |
|---|---|---|---|---|---|---|---|---|---|---|---|---|---|---|---|---|---|---|---|---|---|
| 1328.07 | 1038.18 | 1616.39 | 126.26 | 288.44 | 104.61 | 1145.3 | 537.29 | 553.03 | 818.4 | 517.19 | 2168.85 | 709.76 | 2713.67 | 373.48 | 955.54 | 895.74 | 433.93 | 16324.13 | Matrix metalloproteinase-24 (MMP-24) (EC 3.4.24.-) (Membrane-type matrix metalloproteinase 5) (MT-MMP 5) (MTMMP5) (Membrane-type-5 matrix metalloproteinase) (MT5-MMP) (MT5MMP) [Cleaved into: Processed matrix metalloproteinase-24] | Mmp24 Mt5mmp | cell-cell adhesion mediated by cadherin [GO:0044331]; cell-cell adhesion via plasma-membrane adhesion molecules [GO:0098742]; collagen catabolic process [GO:0030574]; detection of temperature stimulus involved in sensory perception of pain [GO:0050965]; extracellular matrix organization [GO:0030198]; glial cell differentiation [GO:0010001]; neuronal stem cell population maintenance [GO:0097150]; proteolysis [GO:0006508] |
| 885.56 | 839.51 | 682.64 | 533.51 | 895.72 | 947.6 | 2828.42 | 378.95 | 549.23 | 410.88 | 503.52 | 873.92 | 993.42 | 1037.16 | 553.13 | 2151.53 | 762.5 | 474.7 | 16301.9 | Histone-lysine N-methyltransferase KMT5C (Lysine N-methyltransferase 5C) (Lysine-specific methyltransferase 5C) (Suppressor of variegation 4-20 homolog 2) (Su(var)4-20 homolog 2) (Suv4-20h2) ([histone H4]-N-methyl-L-lysine20 N-methyltransferase KMT5B) (EC 2.1.1.362) ([histone H4]-lysine20 N-methyltransferase KMT5B) (EC 2.1.1.361) | KMT5C SUV420H2 PP7130 | DNA repair [GO:0006281]; methylation [GO:0032259]; positive regulation of double-strand break repair via nonhomologous end joining [GO:2001034]; positive regulation of isotype switching [GO:0045830] |

| | | | | | | | | | | | | | | | | | | | | | |
|---|---|---|---|---|---|---|---|---|---|---|---|---|---|---|---|---|---|---|---|---|---|
| 4697.76 | 3280.32 | 2016.59 | 129.77 | 345.2 | 563.43 | 11.44 | 139.73 | 37.22 | 1155.76 | 993.18 | 1318.41 | 507.59 | 560.09 | 396.63 | 56.64 | 72.23 | 9.06 | 16291.05 | Endoplasmic reticulum-Golgi intermediate compartment protein 1 (ER-Golgi intermediate compartment 32 kDa protein) (ERGIC-32) | ergic1 ergic32 zgc:114085 | endoplasmic reticulum to Golgi vesicle-mediated transport [GO:0006888]; retrograde vesicle-mediated transport, Golgi to endoplasmic reticulum [GO:0006890] |
| 546.38 | 606.87 | 821.56 | 332.2 | 650.73 | 792.85 | 2386.68 | 281.9 | 1160.17 | 203.72 | 252.87 | 1665.46 | 859.51 | 965.77 | 1041.48 | 1691.01 | 670.7 | 1353.56 | 16283.42 | E3 ubiquitin-protein ligase ZNRF1 (EC 2.3.2.27) (RING-type E3 ubiquitin transferase ZNRF1) (Zinc/RING finger protein 1) | znrf1 | proteasome-mediated ubiquitin-dependent protein catabolic process [GO:0043161]; protein K48-linked ubiquitination [GO:0070936] |
| 0 | 0 | 0 | 7.02 | 2.3 | 0 | 1.97 | 3.6 | 14.17 | 0 | 14.57 | 4.65 | 0 | 14030.55 | 53.41 | 2.02 | 2113.59 | 23.61 | 16271.46 | Mitochondrial glutamate carrier 1 (GC-1) (Glutamate/H(+) symporter 1) (Solute carrier family 25 member 22) | SLC25A22 GC1 | aspartate transmembrane transport [GO:0015810]; L-glutamate transmembrane transport [GO:0015813]; malate-aspartate shuttle [GO:0043490]; regulation of insulin secretion [GO:0050796] |

| | | | | | | | | | | | | | | | | | | | | | |
|---|---|---|---|---|---|---|---|---|---|---|---|---|---|---|---|---|---|---|---|---|---|
| 472.45 | 1183.51 | 1106.94 | 275.9 | 2285.01 | 1192.93 | 203.37 | 903.22 | 517.15 | 241.37 | 524.78 | 713.49 | 1063.92 | 2356.73 | 937.68 | 302.31 | 991.27 | 995.35 | 16267.38 | Juxtaposed with another zinc finger protein 1 (TAK1-interacting protein 27) (Zinc finger protein 802) | JAZF1 TIP27 ZNF802 | lipid metabolic process [GO:0006629]; negative regulation of transcription by RNA polymerase II [GO:0000122] |
| 27.72 | 0 | 6030.03 | 27.21 | 0 | 1408.88 | 0 | 2.19 | 27.31 | 14.69 | 3.51 | 8627.06 | 0 | 0 | 68.13 | 12.14 | 2.62 | 5.16 | 16256.65 | 5-hydroxytryptamine receptor 2A (5-HT-2) (5-HT-2A) (Serotonin receptor 2A) | HTR2A HTR2 | artery smooth muscle contraction [GO:0014824]; behavioral response to cocaine [GO:0048148]; chemical synaptic transmission [GO:0007268]; detection of mechanical stimulus involved in sensory perception of pain [GO:0050966]; detection of temperature stimulus involved in sensory perception of pain [GO:0050965]; G protein-coupled receptor signaling pathway, coupled to cyclic nucleotide second messenger [GO:0007187]; G protein-coupled serotonin receptor signaling pathway [GO:0098664]; glycolytic process [GO:0006096]; intracellular calcium ion homeostasis [GO:0006874]; memory [GO:0007613]; negative regulation of potassium ion transport [GO:0043267]; negative regulation of synaptic transmission, glutamatergic [GO:0051967]; phospholipase C-activating serotonin receptor signaling pathway [GO:0007208]; positive regulation of cell population proliferation [GO:0008284]; positive |
| 1733.59 | 1099.27 | 935.95 | 1143.97 | 1058.14 | 1807.51 | 407.43 | 997.35 | 1218.31 | 1224.12 | 654.87 | 541.85 | 606.3 | 689.69 | 399.35 | 514.58 | 711.17 | 508.39 | 16251.84 | Bifunctional glutamate/proline--tRNA ligase (Bifunctional aminoacyl-tRNA synthetase) (Glutamyl-prolyl-tRNA synthetase) [Includes: Glutamate--tRNA ligase (EC 6.1.1.17) (Glutamyl-tRNA synthetase) (GluRS);Proline--tRNA ligase (EC 6.1.1.15) (Prolyl-tRNA synthetase) (ProRS)] | EPRS1 EPRS QPRS | cellular response to insulin stimulus [GO:0032869]; cellular response to type II interferon [GO:0071346]; glutamyl-tRNA aminoacylation [GO:0006424]; negative regulation of translation [GO:0017148]; prolyl-tRNA aminoacylation [GO:0006433]; regulation of long-chain fatty acid import into cell [GO:0140212] |

| | | | | | | | | | | | | | | | | | | | | | |
|---|---|---|---|---|---|---|---|---|---|---|---|---|---|---|---|---|---|---|---|---|---|
| 2.09 | 0.88 | 2305.76 | 0.78 | 0.43 | 492.49 | 67.55 | 0 | 48.65 | 4.48 | 0.85 | 13121.68 | 0.97 | 0.58 | 98.26 | 53.04 | 2.49 | 49.83 | 16250.81 | Carnitine O-palmitoyltransferase 1, liver isoform (CPT1-L) (EC 2.3.1.21) (Carnitine O-palmitoyltransferase I, liver isoform) (CPT I) (CPTI-L) (Carnitine palmitoyltransferase 1A) (Succinyltransferase CPT1A) (EC 2.3.1.-) | CPT1A CPT1 | aflatoxin metabolic process [GO:0046222]; carnitine metabolic process [GO:0009437]; carnitine shuttle [GO:0006853]; cellular response to fatty acid [GO:0071398]; eating behavior [GO:0042755]; epithelial cell differentiation [GO:0030855]; fatty acid beta-oxidation [GO:0006635]; fatty acid metabolic process [GO:0006631]; glucose metabolic process [GO:0006006]; liver regeneration [GO:0097421]; long-chain fatty acid metabolic process [GO:0001676]; positive regulation of fatty acid beta-oxidation [GO:0032000]; positive regulation of innate immune response [GO:0045089]; regulation of insulin secretion [GO:0050796]; regulation of lipid storage [GO:0010883]; response to alkaloid [GO:0043279]; response to ethanol [GO:0045471]; response to hypoxia [GO:0001666]; response to nutrient [GO:0007584]; response to tetrachloromethane [GO:1904772]; response to xenobiotic stimulus [illegible] |
| 1842.85 | 522.68 | 1590.23 | 483.96 | 1185.2 | 2550.83 | 7.12 | 367.77 | 236.3 | 516.49 | 928.16 | 1142.47 | 589.02 | 1393.53 | 433.18 | 159.09 | 1670.48 | 628.07 | 16247.43 | Sodium/hydrogen exchanger 7 (Na(+)/H(+) exchanger 7) (NHE-7) (Solute carrier family 9 member 7) | SLC9A7 NHE7 | monoatomic ion transport [GO:0006811]; potassium ion transmembrane transport [GO:0071805]; regulation of Golgi lumen acidification [GO:1905526]; regulation of intracellular pH [GO:0051453]; regulation of pH [GO:0006885]; sodium ion import across plasma membrane [GO:0098719] |
| 423.72 | 29.18 | 553.16 | 123.48 | 64.61 | 539.9 | 653.51 | 86.58 | 164.13 | 329.55 | 70.3 | 230.22 | 1484.8 | 1667.34 | 3614.85 | 1431.96 | 1601.8 | 3097.92 | 16167.01 | Tetratricopeptide repeat protein 24 (TPR repeat protein 24) | TTC24 | |

| | | | | | | | | | | | | | | | | | | | | | |
|---|---|---|---|---|---|---|---|---|---|---|---|---|---|---|---|---|---|---|---|---|---|
| 615.64 | 343.14 | 1420.5 | 241.36 | 557.27 | 853.45 | 1062.01 | 211.81 | 2259.38 | 323.31 | 219.07 | 1172.17 | 608.88 | 686.28 | 2332.43 | 578.02 | 306.7 | 2350.09 | 16141.51 | Myocardin | Myocd Mycd | cardiac muscle cell apoptotic process [GO:0010659]; cardiac muscle cell differentiation [GO:0055007]; cardiac muscle cell myoblast differentiation [GO:0060379]; cardiac vascular smooth muscle cell differentiation [GO:0060947]; cardiac ventricle development [GO:0003231]; cell growth involved in cardiac muscle cell development [GO:0061049]; cellular component maintenance [GO:0043954]; cellular response to angiotensin [GO:1904385]; cellular response to growth factor stimulus [GO:0071363]; cellular response to hypoxia [GO:0071456]; cellular response to transforming growth factor beta stimulus [GO:0071560]; chromatin organization [GO:0006325]; digestive tract development [GO:0048565]; ductus arteriosus closure [GO:0097070]; heart development [GO:0007507]; hepatic stellate cell activation [GO:0035733]; lung alveolus development [GO:0048286]; negative regulation of amyloid-beta clearance |
| 952.24 | 319.62 | 826.04 | 142.69 | 200.16 | 501.8 | 265.45 | 70.66 | 2835.43 | 171.73 | 67.63 | 1297.27 | 1502.52 | 316.62 | 2765.37 | 282.17 | 149.72 | 3469.27 | 16136.39 | Microtubule-associated protein 4 (MAP-4) | MAP4 | cell division [GO:0051301]; cilium disassembly [GO:0061523]; establishment of spindle orientation [GO:0051294]; microtubule cytoskeleton organization [GO:0000226]; microtubule polymerization [GO:0046785]; microtubule sliding [GO:0051012]; mitotic spindle organization [GO:0007052]; negative regulation of non-motile cilium assembly [GO:1902856]; neuron projection development [GO:0031175] |
| 587.35 | 1219.97 | 1384.96 | 336.77 | 1070.85 | 971.77 | 1335.03 | 390.15 | 740.96 | 171.12 | 349.19 | 1666.7 | 711.82 | 1697.28 | 880.86 | 887.55 | 838.31 | 893.48 | 16134.12 | RNA-binding motif, single-stranded-interacting protein 2 (Suppressor of CDC2 with RNA-binding motif 3) | RBMS2 SCR3 | RNA processing [GO:0006396] |

| | | | | | | | | | | | | | | | | | | | | | |
|---|---|---|---|---|---|---|---|---|---|---|---|---|---|---|---|---|---|---|---|---|---|
| 359.61 | 498.34 | 1204.71 | 346.48 | 350.49 | 521.07 | 1911.97 | 60.14 | 2213.08 | 152.97 | 179.31 | 1985.05 | 693.15 | 873.22 | 1049.25 | 1620.07 | 481.57 | 1630.27 | 16130.75 | rRNA N(6)-adenosine-methyltransferase METTL5 (EC 2.1.1.-) (Methyltransferase-like protein 5) | mettl5 | positive regulation of translation [GO:0045727]; rRNA methylation [GO:0031167]; stem cell differentiation [GO:0048863] |
| 248.41 | 644.01 | 3335.37 | 203.24 | 302.27 | 3664.19 | 10.66 | 143.23 | 175.86 | 473.58 | 518.5 | 4105.03 | 137.67 | 368.17 | 556.97 | 498.75 | 362.65 | 381.81 | 16130.37 | Sulfotransferase 1C1 (ST1C1) (EC 2.8.2.-) (Phenol sulfotransferase) | Sult1c1 Sult1a2 | 3'-phosphoadenosine 5'-phosphosulfate metabolic process [GO:0050427]; sulfur compound metabolic process [GO:0006790]; xenobiotic metabolic process [GO:0006805] |
| 111.56 | 87.84 | 890.59 | 31.8 | 33.95 | 105.57 | 16.65 | 18.91 | 4337.82 | 29.93 | 26.18 | 1361.8 | 139.13 | 95.97 | 4121.42 | 32.62 | 51.24 | 4631.58 | 16124.56 | Zinc finger FYVE domain-containing protein 16 (Endofin) (Endosomal-associated FYVE domain protein) | Zfyve16 Kiaa0305 | |

| | | | | | | | | | | | | | | | | | | | | |
|---|---|---|---|---|---|---|---|---|---|---|---|---|---|---|---|---|---|---|---|---|
| 445.1 | 474.37 | 641.82 | 238.73 | 539.47 | 734.79 | 3777.64 | 249.26 | 1475.42 | 284.16 | 350.89 | 754.81 | 477.69 | 638.32 | 924.96 | 2263.18 | 440.58 | 1400.95 | 16112.14 | 26S proteasome non-ATPase regulatory subunit 3 (26S proteasome regulatory subunit RPN3) (26S proteasome regulatory subunit S3) (Proteasome subunit p58) (Transplantation antigen P91A) (Tum-P91A antigen) | Psmd3 P91a Tstap91a | regulation of protein catabolic process [GO:0042176] |
| 178.26 | 162.52 | 550.13 | 64.47 | 196.73 | 252.51 | 1418.79 | 121.75 | 3871.69 | 74.98 | 113.38 | 1367.33 | 131.09 | 286.05 | 3339.54 | 520.85 | 135.41 | 3322.12 | 16107.6 | Sodium/potassium-transporting ATPase subunit alpha-2 (Na(+)/K(+) ATPase alpha-2 subunit) (EC 7.2.2.13) (Sodium pump subunit alpha-2) | ATP1A2 KIAA0778 | adult locomotory behavior [GO:0008344]; amygdala development [GO:0021764]; ATP metabolic process [GO:0046034]; behavioral fear response [GO:0001662]; cardiac muscle contraction [GO:0060048]; cell communication by electrical coupling involved in cardiac conduction [GO:0086064]; cellular response to mechanical stimulus [GO:0071260]; cellular response to steroid hormone stimulus [GO:0071383]; intracellular potassium ion homeostasis [GO:0030007]; intracellular sodium ion homeostasis [GO:0006883]; L-ascorbic acid metabolic process [GO:0019852]; locomotion [GO:0040011]; locomotory exploration behavior [GO:0035641]; membrane depolarization during cardiac muscle cell action potential [GO:0086012]; membrane repolarization [GO:0086009]; monoatomic cation transmembrane transport [GO:0098655]; negative regulation of calcium ion transmembrane transport |

| | | | | | | | | | | | | | | | | | | | | | |
|---|---|---|---|---|---|---|---|---|---|---|---|---|---|---|---|---|---|---|---|---|---|
| 55.45 | 0 | 10.41 | 184.71 | 0 | 6.21 | 8322.97 | 0 | 2.82 | 92.22 | 287.22 | 3.85 | 12 | 0 | 4.11 | 6888.33 | 227.26 | 0 | 16097.56 | Very-long-chain (3R)-3-hydroxyacyl-CoA dehydratase (EC 4.2.1.134) (3-hydroxyacyl-CoA dehydratase) (HACD) (Protein-tyrosine phosphatase-like A domain-containing protein 1) | hacd3 ptplad1 si:ch211-117i10.7 zgc:63632 | fatty acid elongation [GO:0030497]; sphingolipid biosynthetic process [GO:0030148]; very long-chain fatty acid biosynthetic process [GO:0042761] |
| 1094.14 | 507.55 | 1189.95 | 282.55 | 522.99 | 969.46 | 82.71 | 229.71 | 1814.97 | 326.73 | 161.07 | 1125.7 | 774.56 | 1136.23 | 2126.83 | 315.84 | 659.19 | 2759.52 | 16079.7 | 26S proteasome non-ATPase regulatory subunit 4 (26S proteasome regulatory subunit RPN10) (26S proteasome regulatory subunit S5A) (Multiubiquitin chain-binding protein) | Psmd4 Mcb1 | |
| 893.9 | 1073.65 | 1404.49 | 320.64 | 899.54 | 1904.15 | 135.24 | 392.84 | 110.5 | 436.87 | 411.18 | 880 | 2679.33 | 1655.41 | 1134.15 | 353.13 | 760.35 | 600.71 | 16046.08 | Neutrophil cytosolic factor 1 (NCF-1) | Ncf1 | cellular defense response [GO:0006968]; cellular response to glucose stimulus [GO:0071333]; cellular response to testosterone stimulus [GO:0071394]; defense response to bacterium [GO:0042742]; defense response to fungus [GO:0050832]; defense response to Gram-positive bacterium [GO:0050830]; epithelial cell proliferation [GO:0050673]; hydrogen peroxide biosynthetic process [GO:0050665]; inflammatory response [GO:0006954]; leukocyte mediated cytotoxicity [GO:0001909]; leukotriene metabolic process [GO:0006691]; neutrophil-mediated killing of fungus [GO:0070947]; neutrophil-mediated killing of gram-positive bacterium [GO:0070946]; protein targeting to membrane [GO:0006612]; reactive oxygen species biosynthetic process [GO:1903409]; regulation of respiratory burst involved in inflammatory response [GO:0060264]; respiratory burst [GO:0045730]; respiratory burst involved in defense response [illegible] |

| | | | | | | | | | | | | | | | | | | | | | |
|---|---|---|---|---|---|---|---|---|---|---|---|---|---|---|---|---|---|---|---|---|---|
| 1716.93 | 608.58 | 1295.44 | 497.06 | 1224.26 | 2308.41 | 10.1 | 609.35 | 301.02 | 446.43 | 280.4 | 583.88 | 2021.27 | 1726.58 | 982.79 | 294.23 | 667.87 | 445.33 | 16019.93 | Kinesin-like protein KIF1C | Kif1c | microtubule-based movement [GO:0007018]; retrograde vesicle-mediated transport, Golgi to endoplasmic reticulum [GO:0006890] |
| 894.21 | 630.49 | 1314.85 | 233.47 | 647.49 | 748.98 | 175.49 | 339.32 | 2497.41 | 300.58 | 306.9 | 1214.74 | 665.04 | 876.34 | 2110.19 | 318.95 | 385.91 | 2347.72 | 16008.08 | ATP-dependent RNA helicase DHX58 (EC 3.6.4.13) (ATP-dependent helicase LGP2) (Protein D11Lgp2 homolog) (RIG-I-like receptor 3) (RLR-3) (RIG-I-like receptor LGP2) (RLR) | DHX58 D11LGP2E LGP2 | antiviral innate immune response [GO:0140374]; cytoplasmic pattern recognition receptor signaling pathway [GO:0002753]; negative regulation of innate immune response [GO:0045824]; negative regulation of MDA-5 signaling pathway [GO:0039534]; negative regulation of RIG-I signaling pathway [GO:0039536]; negative regulation of type I interferon production [GO:0032480]; positive regulation of MDA-5 signaling pathway [GO:1900245]; positive regulation of RIG-I signaling pathway [GO:1900246]; positive regulation of type I interferon production [GO:0032481]; regulation of innate immune response [GO:0045088]; response to bacterium [GO:0009617]; response to virus [GO:0009615] |
| 1175.11 | 796.22 | 826.72 | 456.91 | 832.89 | 1407.01 | 2860.51 | 407.43 | 358.74 | 467.99 | 371.32 | 509.42 | 1237.43 | 880.75 | 549.47 | 1757.12 | 600.98 | 504.28 | 16000.3 | Diablo IAP-binding mitochondrial protein (Diablo homolog, mitochondrial) (Direct IAP-binding protein with low pI) (Second mitochondria-derived activator of caspase) (Smac) [Cleaved into: Diablo IAP-binding mitochondrial protein, cleaved form] | Diablo Smac | intrinsic apoptotic signaling pathway [GO:0097193]; intrinsic apoptotic signaling pathway in response to oxidative stress [GO:0008631]; neuron apoptotic process [GO:0051402]; positive regulation of apoptotic process [GO:0043065] |

| | | | | | | | | | | | | | | | | | | | | | |
|---|---|---|---|---|---|---|---|---|---|---|---|---|---|---|---|---|---|---|---|---|---|
| 1005.03 | 1430.14 | 2818.84 | 203.33 | 707.88 | 1655.32 | 444.18 | 177.34 | 421.32 | 435.64 | 334.95 | 908.34 | 649.05 | 1544.48 | 827.81 | 265.45 | 466.81 | 1652 | 15947.91 | E3 SUMO-protein ligase PIAS2 (EC 2.3.2.-) (Androgen receptor-interacting protein 3) (ARIP3) (DAB2-interacting protein) (DIP) (E3 SUMO-protein transferase PIAS2) (Msx-interacting zinc finger protein) (Miz1) (PIAS-NY protein) (Protein inhibitor of activated STAT x) (Protein inhibitor of activated STAT2) | PIAS2 PIASX | DNA-templated transcription [GO:0006351]; negative regulation of androgen receptor signaling pathway [GO:0060766]; negative regulation of DNA-binding transcription factor activity [GO:0043433]; protein sumoylation [GO:0016925]; regulation of transcription by RNA polymerase II [GO:0006357] |
| 1435.76 | 96.28 | 1674.58 | 275.94 | 436.91 | 1799.74 | 13.45 | 69.74 | 1193.77 | 670.27 | 260.18 | 2301.87 | 1370.83 | 342.25 | 1495.18 | 283.86 | 448.28 | 1760.61 | 15929.5 | Protein ABHD18 (Alpha/beta hydrolase domain-containing protein 18) (Abhydrolase domain-containing protein 18) | Abhd18 | |
| 18.82 | 0 | 4.86 | 227.78 | 0 | 0 | 10580.49 | 0 | 0 | 79.05 | 181.35 | 0 | 0 | 0 | 0 | 4742.11 | 84.03 | 0 | 15918.49 | FRAS1-related extracellular matrix protein 2 (ECM3 homolog) | FREM2 | anatomical structure morphogenesis [GO:0009653]; cell adhesion [GO:0007155]; cell communication [GO:0007154]; embryonic digit morphogenesis [GO:0042733]; eye development [GO:0001654]; heart development [GO:0007507]; inner ear development [GO:0048839]; kidney development [GO:0001822]; morphogenesis of an epithelium [GO:0002009] |

| | | | | | | | | | | | | | | | | | | | | | |
|---|---|---|---|---|---|---|---|---|---|---|---|---|---|---|---|---|---|---|---|---|---|
| 2903.86 | 31.79 | 18.6 | 764.08 | 216.16 | 525.88 | 11.84 | 18.16 | 0.33 | 2876.04 | 149.74 | 124.87 | 7919.32 | 24.04 | 25.55 | 213.4 | 46.38 | 0.75 | 15870.79 | Ankyrin repeat and SAM domain-containing protein 1A (Odin) | ANKS1A ANKS1 KIAA0229 ODIN | ephrin receptor signaling pathway [GO:0048013]; neuron remodeling [GO:0016322]; substrate-dependent cell migration [GO:0006929] |
| 0 | 0 | 4330.24 | 0 | 0 | 0 | 142.25 | 0 | 3858.35 | 0 | 6.1 | 1053.14 | 0 | 0 | 1971.12 | 83.26 | 13.99 | 4386.85 | 15845.3 | Gap junction beta-1 protein (Connexin-32) (Cx32) | GJB1 CXN-32 | cell-cell signaling [GO:0007267]; purine ribonucleotide transport [GO:0015868] |
| 3604.03 | 44.26 | 28.02 | 833.73 | 214.06 | 375.83 | 28.26 | 16.47 | 0 | 2910.67 | 132.2 | 131.37 | 7169.17 | 20.95 | 17.15 | 200.51 | 41.78 | 4.63 | 15773.09 | Cadherin EGF LAG seven-pass G-type receptor 2 (Cadherin family member 10) (Epidermal growth factor-like protein 2) (EGF-like protein 2) (Flamingo homolog 3) (Multiple epidermal growth factor-like domains protein 3) (Multiple EGF-like domains protein 3) | CELSR2 CDHF10 EGFL2 KIAA0279 MEGF3 | cell-cell adhesion [GO:0098609]; cerebrospinal fluid secretion [GO:0033326]; cilium assembly [GO:0060271]; cilium movement [GO:0003341]; dendrite morphogenesis [GO:0048813]; G protein-coupled receptor signaling pathway [GO:0007186]; homophilic cell adhesion via plasma membrane adhesion molecules [GO:0007156]; motor neuron migration [GO:0097475]; neural plate anterior/posterior regionalization [GO:0021999]; regulation of cell-cell adhesion [GO:0022407]; regulation of DNA-templated transcription [GO:0006355]; regulation of protein localization [GO:0032880]; ventricular system development [GO:0021591]; Wnt signaling pathway [GO:0016055]; Wnt signaling pathway, planar cell polarity pathway [GO:0060071] |

| | | | | | | | | | | | | | | | | | | | | | |
|---|---|---|---|---|---|---|---|---|---|---|---|---|---|---|---|---|---|---|---|---|---|
| 1387 | 670.4 | 881.96 | 446.87 | 698.38 | 2720.88 | 1190.86 | 251.41 | 215.74 | 814 | 374.67 | 884.45 | 2039.62 | 740.96 | 734.47 | 760.96 | 633.07 | 298.38 | 15744.08 | Rapamycin-insensitive companion of mTOR (AVO3 homolog) (hAVO3) | RICTOR KIAA1999 | actin cytoskeleton organization [GO:0030036]; cellular response to nutrient levels [GO:0031669]; cytoskeleton organization [GO:0007010]; embryo development ending in birth or egg hatching [GO:0009792]; lipid biosynthetic process [GO:0008610]; negative regulation of apoptotic process [GO:0043066]; positive regulation of actin filament polymerization [GO:0030838]; positive regulation of cell growth [GO:0030307]; positive regulation of cell migration [GO:0030335]; positive regulation of endothelial cell proliferation [GO:0001938]; positive regulation of phosphatidylinositol 3-kinase/protein kinase B signal transduction [GO:0051897]; positive regulation of TOR signaling [GO:0032008]; regulation of actin cytoskeleton organization [GO:0032956]; regulation of establishment of cell polarity [GO:2000114]; regulation of gene expression [GO:0010468]; regulation of |
| 702.48 | 1110.15 | 1240.99 | 291.85 | 1406.51 | 3072.01 | 430.54 | 397.17 | 229.58 | 350.13 | 391.17 | 748.03 | 680.48 | 1485.41 | 1241.37 | 537.68 | 1140.94 | 270.88 | 15727.37 | Palmitoyltransferase ZDHHC17 (EC 2.3.1.225) (Acyltransferase ZDHHC17) (EC 2.3.1.-) (DHHC domain-containing cysteine-rich protein 17) (Zinc finger DHHC domain-containing protein 17) | zdhhc17 dhhc17 | axonogenesis [GO:0007409]; habituation [GO:0046959]; regulation of ERK1 and ERK2 cascade [GO:0070372]; regulation of neurotrophin TRK receptor signaling pathway [GO:0051386] |
| 862.84 | 1386.7 | 1090.84 | 672.01 | 1193.78 | 1165.53 | 782.97 | 810.89 | 494.63 | 388.79 | 513.69 | 671.47 | 1213.18 | 1087.59 | 813.44 | 657.19 | 1232.98 | 686.59 | 15725.11 | Myophilin | | actin filament organization [GO:0007015] |
| 32.62 | 20.2 | 3876.7 | 7.9 | 13.86 | 17.61 | 0.78 | 2.41 | 3965.97 | 7.52 | 10.39 | 1151.69 | 20.22 | 9.61 | 2981.54 | 2.75 | 11.65 | 3566.29 | 15699.71 | GTP-binding protein RAD (RAD1) (Ras associated with diabetes) | RRAD RAD | small GTPase-mediated signal transduction [GO:0007264] |

| | | | | | | | | | | | | | | | | | | | | | |
|---|---|---|---|---|---|---|---|---|---|---|---|---|---|---|---|---|---|---|---|---|---|
| 529.88 | 274.08 | 1970.43 | 161.91 | 352.58 | 1609.33 | 51.07 | 127.43 | 2423.51 | 138.84 | 134.36 | 1787.97 | 493.47 | 464.21 | 2231.27 | 97.54 | 156.83 | 2683.4 | 15688.11 | Voltage-gated inwardly rectifying potassium channel KCNH6 (Ether-a-go-go-related gene potassium channel 2) (ERG-2) (Eag-related protein 2) (Ether-a-go-go-related protein 2) (hERG-2) (hERG2) (Potassium voltage-gated channel subfamily H member 6) (Voltage-gated potassium channel subunit Kv11.2) | KCNH6 ERG2 | membrane repolarization during cardiac muscle cell action potential [GO:0086013]; potassium ion transmembrane transport [GO:0071805]; regulation of heart rate by cardiac conduction [GO:0086091]; regulation of ventricular cardiac muscle cell membrane repolarization [GO:0060307] |
| 1.9 | 1.08 | 825.4 | 0 | 1.64 | 14.8 | 0 | 1.3 | 3816.26 | 2.92 | 2.33 | 1357.14 | 181.34 | 14.9 | 4390.13 | 2.05 | 8.57 | 5048.5 | 15670.26 | Voltage-dependent T-type calcium channel subunit alpha-1H (Voltage-gated calcium channel subunit alpha Cav3.2) | Cacna1h | aldosterone biosynthetic process [GO:0032342]; calcium ion import [GO:0070509]; calcium ion transport [GO:0006816]; cellular response to hormone stimulus [GO:0032870]; cellular response to potassium ion [GO:0035865]; cortisol biosynthetic process [GO:0034651]; inorganic cation transmembrane transport [GO:0098662]; membrane depolarization during action potential [GO:0086010]; positive regulation of acrosome reaction [GO:2000344]; positive regulation of calcium ion-dependent exocytosis [GO:0045956]; positive regulation of cardiac muscle cell contraction [GO:0106134]; regulation of membrane potential [GO:0042391] |
| 5.1 | 0.4 | 762.63 | 5.3 | 0.36 | 7.64 | 16.43 | 0.22 | 4859.12 | 1.11 | 1.86 | 1183.94 | 0 | 0 | 3981.39 | 30.5 | 9.96 | 4803.07 | 15669.03 | Enhancer of mRNA-decapping protein 4 | edc4 si:dkey-16n13.6 | deadenylation-independent decapping of nuclear-transcribed mRNA [GO:0031087] |

| | | | | | | | | | | | | | | | | | | | | | |
|---|---|---|---|---|---|---|---|---|---|---|---|---|---|---|---|---|---|---|---|---|---|
| 378.99 | 554.4 | 2296.41 | 198.4 | 640.32 | 1116.87 | 3.22 | 365.94 | 1200.44 | 158.74 | 200.26 | 1120.34 | 656.51 | 1116.44 | 2568.4 | 171.92 | 498.04 | 2370.13 | 15615.77 | Cholesterol 7-desaturase nvd (EC 1.14.19.21) (Protein neverland) (Nvd protein) (nvd-Dr) | nvd zgc:92275 | cholesterol metabolic process [GO:0008203] |
| 755.74 | 885.17 | 1598.58 | 398.07 | 1194.18 | 2484.43 | 82.47 | 192.35 | 44.82 | 400.96 | 545.83 | 1745.17 | 1835.27 | 1528.13 | 510.25 | 382.18 | 913.71 | 104.3 | 15601.61 | Myosin light chain kinase family member 4 (EC 2.7.11.1) (Sugen kinase 85) (SgK085) | MYLK4 SGK085 | signal transduction [GO:0007165] |
| 798.06 | 846.57 | 8288.92 | 147.71 | 365.09 | 413.51 | 40.23 | 102.99 | 2238.83 | 166.9 | 217.76 | 1663.39 | 33.47 | 6.27 | 61.67 | 0 | 0 | 207.92 | 15599.29 | Large ribosomal subunit protein uL10 (60S acidic ribosomal protein P0) (60S ribosomal protein L10E) | RPLP0 | cytoplasmic translation [GO:0002181]; ribosomal large subunit assembly [GO:0000027] |
| 5.19 | 6953.38 | 21.84 | 2.61 | 978.16 | 4.31 | 0 | 2820.16 | 17.73 | 9.66 | 1421.13 | 0 | 0.77 | 2470.38 | 24.14 | 6.55 | 821.61 | 29.31 | 15586.93 | Plasma kallikrein (EC 3.4.21.34) (Fletcher factor) (Kininogenin) (Plasma prekallikrein) [Cleaved into: Plasma kallikrein heavy chain;Plasma kallikrein light chain] | Klkb1 Klk3 Pk | blood coagulation [GO:0007596]; fibrinolysis [GO:0042730]; inflammatory response [GO:0006954]; liver regeneration [GO:0097421]; plasminogen activation [GO:0031639]; positive regulation of fibrinolysis [GO:0051919] |

| | | | | | | | | | | | | | | | | | | | | | |
|---|---|---|---|---|---|---|---|---|---|---|---|---|---|---|---|---|---|---|---|---|---|
| 236.08 | 112.53 | 74.39 | 90.43 | 100.76 | 344.34 | 9796.29 | 41.47 | 187.35 | 115.28 | 108.2 | 122.79 | 326.42 | 219.13 | 240.53 | 3205 | 117.63 | 134.99 | 15573.61 | E3 ubiquitin-protein ligase CHIP (EC 2.3.2.27) (Carboxy terminus of Hsp70-interacting protein) (RING-type E3 ubiquitin transferase CHIP) (STIP1 homology and U box-containing protein 1) | Stub1 Chip | cellular response to heat [GO:0034605]; cellular response to hypoxia [GO:0071456]; cellular response to misfolded protein [GO:0071218]; chaperone-mediated autophagy [GO:0061684]; DNA repair [GO:0006281]; endoplasmic reticulum unfolded protein response [GO:0030968]; ERAD pathway [GO:0036503]; MAPK cascade [GO:0000165]; negative regulation of apoptotic process [GO:0043066]; negative regulation of cardiac muscle hypertrophy [GO:0010614]; negative regulation of peroxisome proliferator activated receptor signaling pathway [GO:0035359]; negative regulation of smooth muscle cell apoptotic process [GO:0034392]; negative regulation of transforming growth factor beta receptor signaling pathway [GO:0030512]; negative regulation of vascular associated smooth muscle contraction [GO:1904694]; positive regulation of chaperone-mediated protein complex assembly |
| 2391.37 | 245.29 | 3213.48 | 361.32 | 141.72 | 1059.15 | 33.82 | 101.68 | 789.76 | 636.37 | 85.86 | 844.12 | 1373.86 | 287.6 | 1837.63 | 253.18 | 265.42 | 1633.11 | 15554.74 | 1-phosphatidylinositol 4,5-bisphosphate phosphodiesterase beta-1 (EC 3.1.4.11) (PLC-154) (Phosphoinositide phospholipase C-beta-1) (Phospholipase C-beta-1) (PLC-beta-1) | PLCB1 | G protein-coupled receptor signaling pathway [GO:0007186]; interleukin-1-mediated signaling pathway [GO:0070498]; interleukin-12-mediated signaling pathway [GO:0035722]; interleukin-15-mediated signaling pathway [GO:0035723]; lipid catabolic process [GO:0016042]; memory [GO:0007613]; phosphatidylinositol metabolic process [GO:0046488]; phosphatidylinositol-mediated signaling [GO:0048015]; positive regulation of JNK cascade [GO:0046330]; release of sequestered calcium ion into cytosol [GO:0051209] |

| | | | | | | | | | | | | | | | | | | | | | |
|---|---|---|---|---|---|---|---|---|---|---|---|---|---|---|---|---|---|---|---|---|---|
| 9.49 | 3832.7 | 43.4 | 27.11 | 2761.36 | 37.72 | 0 | 1762.19 | 21.05 | 6.1 | 615.23 | 16.88 | 15.29 | 4318.16 | 26.02 | 2.88 | 2009.08 | 39.66 | 15544.32 | DNA topoisomerase 2-binding protein 1-A (Cut5 protein) (DNA topoisomerase II-binding protein 1-A) (TopBP1-A) (XtopBP) | topbp1-A cut5 | chromosome organization [GO:0051276]; DNA amplification [GO:0006277]; DNA damage response [GO:0006974]; DNA replication checkpoint signaling [GO:0000076]; DNA replication initiation [GO:0006270]; DNA replication preinitiation complex assembly [GO:0071163]; double-strand break repair via alternative nonhomologous end joining [GO:0097681]; double-strand break repair via homologous recombination [GO:0000724]; GINS complex assembly [GO:0071165]; mitotic DNA replication checkpoint signaling [GO:0033314]; mitotic G2 DNA damage checkpoint signaling [GO:0007095]; positive regulation of chromatin binding [GO:0035563]; positive regulation of peptidyl-serine phosphorylation [GO:0033138]; positive regulation of protein kinase activity [GO:0045860]; protein localization to chromatin [GO:0071168]; regulation of DNA-templated DNA replication initiation [GO:0030174] |
| 180.38 | 749.72 | 1200.26 | 194.3 | 1322.22 | 900.22 | 110.97 | 320.39 | 1188.98 | 8.18 | 218.02 | 744.33 | 56.1 | 3144.78 | 1624.25 | 846.06 | 1626.09 | 1105.8 | 15541.05 | DCN1-like protein 5 (DCNL5) (DCUN1 domain-containing protein 5) (Defective in cullin neddylation protein 1-like protein 5) (Squamous cell carcinoma-related oncogene 5) | DCUN1D5 SCCRO5 | DNA damage response [GO:0006974]; positive regulation of protein neddylation [GO:2000436]; protein neddylation [GO:0045116]; regulation of cell growth [GO:0001558]; regulation of protein neddylation [GO:2000434] |
| 1550.75 | 721.08 | 904.8 | 297.22 | 903.1 | 939.91 | 77.02 | 445.01 | 1599.42 | 438.46 | 328.25 | 917.11 | 949.41 | 1157.2 | 1312.25 | 279.1 | 690.34 | 2028.98 | 15539.41 | NHS-like protein 2 | NHSL2 | cell differentiation [GO:0030154] |

| | | | | | | | | | | | | | | | | | | | | | |
|---|---|---|---|---|---|---|---|---|---|---|---|---|---|---|---|---|---|---|---|---|---|
| 1147.35 | 835.48 | 644.48 | 225.05 | 1588.82 | 1046.38 | 711 | 1193.7 | 295.56 | 370.32 | 687.34 | 522.78 | 2073.13 | 1641.58 | 487.22 | 494.95 | 1076.5 | 465.02 | 15506.66 | Transcription factor SOX-5 | SOX5 | asymmetric neuroblast division [GO:0055059]; cartilage condensation [GO:0001502]; cartilage development [GO:0051216]; cell fate commitment [GO:0045165]; cellular response to transforming growth factor beta stimulus [GO:0071560]; chondrocyte differentiation [GO:0002062]; positive regulation of cartilage development [GO:0061036]; positive regulation of chondrocyte differentiation [GO:0032332]; positive regulation of mesenchymal stem cell differentiation [GO:2000741]; regulation of transcription by RNA polymerase II [GO:0006357]; transcription by RNA polymerase II [GO:0006366] |
| 308.64 | 501.6 | 1021.07 | 66.64 | 599.41 | 1351.76 | 822.38 | 256.38 | 1568.16 | 112.88 | 341.99 | 1529.56 | 333.35 | 961.57 | 2647.56 | 509.45 | 341.34 | 2228.08 | 15501.82 | Methylcytosine dioxygenase TET2 (EC 1.14.11.80) | TET2 KIAA1546 Nbla00191 | chromosomal 5-methylcytosine DNA demethylation, oxidation pathway [GO:0141167]; leukocyte differentiation [GO:0002521]; myeloid cell differentiation [GO:0030099]; positive regulation of gene expression via chromosomal CpG island demethylation [GO:0044029]; positive regulation of transcription by RNA polymerase II [GO:0045944]; protein O-linked glycosylation [GO:0006493] |
| 26.8 | 3.98 | 897.21 | 15.89 | 11.68 | 1.87 | 32.54 | 1.72 | 4813.25 | 12.84 | 7.46 | 1546.11 | 85.71 | 11.58 | 2481.51 | 35.49 | 26.91 | 5487.6 | 15500.15 | FGFR1 oncogene partner 2 homolog | fgfr1op2 si:ch211-203h15.2 zgc:64126 | response to wounding [GO:0009611] |
| 832.99 | 1500.97 | 1262.19 | 421.12 | 1322.18 | 1654.58 | 319.4 | 606.06 | 506.06 | 361.56 | 448.2 | 882.62 | 1108.99 | 1461.1 | 831.1 | 446.89 | 800.65 | 732.62 | 15499.28 | Echinoderm microtubule-associated protein-like 5 (EMAP-5) | Eml5 | |

| | | | | | | | | | | | | | | | | | | | | | |
|---|---|---|---|---|---|---|---|---|---|---|---|---|---|---|---|---|---|---|---|---|---|
| 2.6 | 3965.33 | 54.55 | 28.05 | 3073.24 | 39.06 | 473.21 | 1069.5 | 14.71 | 7.42 | 487 | 1.27 | 0.85 | 4032.19 | 1.59 | 385.28 | 1828.92 | 18.73 | 15483.5 | ATP-sensitive inward rectifier potassium channel 8 (Inward rectifier K(+) channel Kir6.1) (Potassium channel, inwardly rectifying subfamily J member 8) (uKATP-1) | KCNJ8 | adaptive immune response [GO:0002250]; apoptotic process [GO:0006915]; atrioventricular node cell differentiation [GO:0060922]; calcium ion transmembrane transport [GO:0070588]; CAMKK-AMPK signaling cascade [GO:0061762]; coronary vasculature development [GO:0060976]; defense response to virus [GO:0051607]; determination of adult lifespan [GO:0008340]; establishment of cell polarity [GO:0030010]; fat cell differentiation [GO:0045444]; fatty acid transport [GO:0015908]; fibroblast proliferation [GO:0048144]; gene expression [GO:0010467]; glutamate secretion, neurotransmission [GO:0061535]; heart morphogenesis [GO:0003007]; inorganic cation transmembrane transport [GO:0098662]; kidney development [GO:0001822]; membrane repolarization during ventricular cardiac muscle cell action potential [GO:0098915]; microglial cell activation [GO:0001774]; neuromuscular process |
| 287.66 | 233.19 | 931.2 | 295.74 | 220.17 | 533.38 | 3941.58 | 112.55 | 1256.6 | 164.09 | 163.84 | 733.13 | 611.88 | 335.22 | 1491.18 | 2001.25 | 308.73 | 1789.75 | 15411.14 | Phospholipase B-like 1 (EC 3.1.1.-) (LAMA-like protein 1) (Lamina ancestor homolog 1) (Phospholipase B domain-containing protein 1) [Cleaved into: Phospholipase B-like 1 chain A;Phospholipase B-like 1 chain B;Phospholipase B-like 1 chain C] | Plbd1 | phospholipid catabolic process [GO:0009395] |

| | | | | | | | | | | | | | | | | | | | | | |
|---|---|---|---|---|---|---|---|---|---|---|---|---|---|---|---|---|---|---|---|---|---|
| 0 | 30.62 | 0 | 9.66 | 16.86 | 0 | 1.56 | 4.59 | 1.46 | 0 | 28.66 | 0 | 9 | 11573.21 | 0 | 2.35 | 3705.83 | 20.74 | 15404.54 | Cystathionine gamma-lyase (EC 4.4.1.1) (Cysteine-protein sulfhydrase) (Gamma-cystathionase) | CTH | cysteine biosynthetic process [GO:0019344]; cysteine biosynthetic process via cystathionine [GO:0019343]; hydrogen sulfide biosynthetic process [GO:0070814]; lipid metabolic process [GO:0006629]; negative regulation of apoptotic process [GO:0043066]; positive regulation of canonical NF-kappaB signal transduction [GO:0043123]; positive regulation of NF-kappaB transcription factor activity [GO:0051092]; protein sulfhydration [GO:0044524]; protein-pyridoxal-5-phosphate linkage via peptidyl-N6-pyridoxal phosphate-L-lysine [GO:0018272]; transsulfuration [GO:0019346] |
| 1608.07 | 343.69 | 2280.94 | 366.82 | 501.86 | 1933.73 | 239.46 | 175.35 | 686.05 | 474.33 | 265.56 | 1664.32 | 1821.92 | 288.63 | 848.25 | 363.48 | 476.74 | 1023.87 | 15363.07 | DENN domain-containing protein 2C | DENND2C | |
| 10.33 | 1.44 | 2185.46 | 0.91 | 0.62 | 602.44 | 2.07 | 2.64 | 2227.82 | 2.7 | 2.25 | 6548.05 | 29.91 | 2.12 | 1397.67 | 3.84 | 1.72 | 2310.5 | 15332.49 | Alanine--tRNA ligase, mitochondrial (EC 6.1.1.7) (Alanyl-tRNA synthetase) (AlaRS) (Protein lactyltransferase AARS2) (EC 6.-.-.-) | Aars2 Aarsl Gm89 Kiaa1270 | mitochondrial alanyl-tRNA aminoacylation [GO:0070143]; negative regulation of cGAS/STING signaling pathway [GO:0160049] |
| 389.29 | 530.04 | 688.13 | 563.25 | 1418.36 | 2131.86 | 1099.13 | 666.24 | 930.49 | 288.9 | 270.55 | 445.79 | 1330.98 | 1533.24 | 987.31 | 682.63 | 638.59 | 697.01 | 15291.79 | Serine/threonine-protein kinase RIO1 (EC 2.7.11.1) (EC 3.6.1.-) (RIO kinase 1) | RIOK1 RIO1 | maturation of SSU-rRNA [GO:0030490]; positive regulation of rRNA processing [GO:2000234]; ribosomal small subunit biogenesis [GO:0042274] |
| 761.1 | 605.87 | 1651.27 | 354.03 | 707.7 | 1426.47 | 542.24 | 217.3 | 1621.6 | 291.14 | 216.96 | 1223.77 | 712.38 | 608.91 | 1517.87 | 512.39 | 393.16 | 1845.45 | 15209.61 | Actin-2 (EC 3.6.4.-) | | |

| | | | | | | | | | | | | | | | | | | | | | |
|---|---|---|---|---|---|---|---|---|---|---|---|---|---|---|---|---|---|---|---|---|---|
| 904.36 | 450.04 | 344.13 | 243.7 | 548.37 | 414.99 | 3279.13 | 267.14 | 95.45 | 205.97 | 295.97 | 194.36 | 1575.02 | 1272.65 | 364.09 | 3601.1 | 856.54 | 287.37 | 15200.38 | Bridge-like lipid transfer protein family member 3B (Syntaxin-6 Habc-interacting protein of 164 kDa) (UHRF1-binding protein 1-like) | Bltp3b Kiaa0701 Ship164 Uhrf1bp1l | early endosome to Golgi transport [GO:0034498]; intermembrane lipid transfer [GO:0120009] |
| 4477.76 | 199.04 | 1010.57 | 652.02 | 178.29 | 1286.38 | 6.94 | 76.96 | 55.6 | 489.72 | 108.07 | 531.54 | 3297.28 | 614.42 | 1086.77 | 295.93 | 604.28 | 227.38 | 15198.95 | Mitotic interactor and substrate of PLK1 (Mitotic spindle positioning protein) | MISP C19orf21 | cell division [GO:0051301]; cell migration [GO:0016477]; establishment of centrosome localization [GO:0051660]; establishment of mitotic spindle orientation [GO:0000132]; mitotic spindle assembly [GO:0090307]; organelle localization [GO:0051640]; regulation of protein localization to cell cortex [GO:1904776] |
| 0 | 0 | 8.68 | 256.69 | 0 | 9.04 | 8209.88 | 3.36 | 0 | 83.56 | 207.97 | 7.55 | 0 | 0 | 0 | 6237.82 | 120.23 | 6.85 | 15151.63 | Fibroblast growth factor receptor substrate 2 (FGFR substrate 2) (FGFR-signaling adaptor SNT) (Suc1-associated neurotrophic factor target 1) (SNT-1) | FRS2 | anterior/posterior axis specification, embryo [GO:0008595]; cell surface receptor protein tyrosine phosphatase signaling pathway [GO:0007185]; fibroblast growth factor receptor signaling pathway [GO:0008543]; forebrain development [GO:0030900]; G protein-coupled receptor signaling pathway [GO:0007186]; gastrulation with mouth forming second [GO:0001702]; lens fiber cell development [GO:0070307]; lens placode formation involved in camera-type eye formation [GO:0046619]; negative regulation of cardiac muscle cell differentiation [GO:2000726]; neuroblast proliferation [GO:0007405]; organ induction [GO:0001759]; positive regulation of MAPK cascade [GO:0043410]; prostate epithelial cord arborization involved in prostate glandular acinus morphogenesis [GO:0060527]; regulation of apoptotic process [GO:0042981]; regulation of epithelial cell proliferation [GO:0050678]; regulation of ERK1 and [illegible] |

| | | | | | | | | | | | | | | | | | | | | | |
|---|---|---|---|---|---|---|---|---|---|---|---|---|---|---|---|---|---|---|---|---|---|
| 5795.12 | 327.24 | 514.71 | 469.78 | 160.45 | 199.2 | 7.49 | 106.27 | 109.13 | 630.57 | 101.02 | 199.36 | 4024.84 | 754.26 | 396.03 | 376.04 | 810.09 | 164.18 | 15145.78 | Slit homolog 2 protein (Slit-2) [Cleaved into: Slit homolog 2 protein N-product;Slit homolog 2 protein C-product] | Slit2 | aortic valve morphogenesis [GO:0003180]; axon guidance [GO:0007411]; axonogenesis [GO:0007409]; cell-cell adhesion [GO:0098609]; dorsal/ventral axon guidance [GO:0033563]; in utero embryonic development [GO:0001701]; kidney development [GO:0001822]; mammary duct terminal end bud growth [GO:0060763]; mammary gland duct morphogenesis [GO:0060603]; metanephros development [GO:0001656]; negative chemotaxis [GO:0050919]; negative regulation of axon extension [GO:0030517]; negative regulation of cell population proliferation [GO:0008285]; negative regulation of gene expression [GO:0010629]; neuron projection morphogenesis [GO:0048812]; olfactory bulb development [GO:0021772]; pulmonary valve morphogenesis [GO:0003184]; retinal ganglion cell axon guidance [GO:0031290]; telencephalon cell migration [GO:0022029]; ventricular septum morphogenesis |
| 41.89 | 35.63 | 712.5 | 4.11 | 29.83 | 174.44 | 2.54 | 8.57 | 5352.69 | 18.22 | 4.91 | 1858.89 | 41.67 | 14.86 | 2794.29 | 13.72 | 22.65 | 3940.57 | 15071.98 | Protein O-linked-mannose beta-1,2-N-acetylglucosaminyltransferase 1 (POMGnT1) (EC 2.4.1.-) | POMGNT1 | O-glycan processing [GO:0016266]; protein O-linked glycosylation [GO:0006493] |
| 637.86 | 1624.01 | 1162.67 | 236.07 | 1104.71 | 944.28 | 521.45 | 411.99 | 987.81 | 256.06 | 409.19 | 718.16 | 440.04 | 1879.28 | 1033.85 | 608.06 | 924.23 | 1155.68 | 15055.4 | PDZ domain-containing protein 8 | Pdzd8 | cytoskeleton organization [GO:0007010]; lipid transport [GO:0006869]; mitochondrial calcium ion homeostasis [GO:0051560]; mitochondrion-endoplasmic reticulum membrane tethering [GO:1990456]; regulation of cell morphogenesis [GO:0022604] |

| | | | | | | | | | | | | | | | | | | | | | |
|---|---|---|---|---|---|---|---|---|---|---|---|---|---|---|---|---|---|---|---|---|---|
| 0 | 0 | 542.84 | 0 | 0 | 0 | 0 | 0 | 5703.39 | 0 | 0 | 992.03 | 0 | 0 | 3021.25 | 0 | 0 | 4776.86 | 15036.37 | Potassium channel subfamily K member 3 (Acid-sensitive potassium channel protein TASK-1) (TWIK-related acid-sensitive K(+) channel 1) (Two pore potassium channel KT3.1) (Two pore K(+) channel KT3.1) | Kcnk3 Task Task1 | cellular response to acidic pH [GO:0071468]; cellular response to hypoxia [GO:0071456]; cellular response to zinc ion [GO:0071294]; cochlea development [GO:0090102]; detection of hypoxic conditions in blood by carotid body chemoreceptor signaling [GO:0003029]; monoatomic ion transmembrane transport [GO:0034220]; negative regulation of cytosolic calcium ion concentration [GO:0051481]; potassium ion transmembrane transport [GO:0071805]; potassium ion transport [GO:0006813]; regulation of action potential firing rate [GO:0099605]; regulation of resting membrane potential [GO:0060075]; response to xenobiotic stimulus [GO:0009410] |
| 756.88 | 666.27 | 381.8 | 289.87 | 601.47 | 905.59 | 3531.43 | 415.92 | 541.53 | 295.99 | 268.62 | 352.21 | 1155.1 | 1057.51 | 597.02 | 1892.88 | 676.79 | 561.43 | 14948.31 | TNF receptor-associated factor 3 (EC 2.3.2.27) (CD40 receptor-associated factor 1) (CRAF1) (CD40-binding protein) (CD40BP) (RING-type E3 ubiquitin transferase TRAF3) | Traf3 Cap-1 Craf1 Trafamn | apoptotic process [GO:0006915]; innate immune response [GO:0045087]; negative regulation of NF-kappaB transcription factor activity [GO:0032088]; positive regulation of type I interferon production [GO:0032481]; regulation of apoptotic process [GO:0042981]; regulation of cytokine production [GO:0001817]; regulation of defense response to virus [GO:0050688]; regulation of interferon-beta production [GO:0032648]; regulation of proteolysis [GO:0030162]; Toll signaling pathway [GO:0008063]; toll-like receptor 4 signaling pathway [GO:0034142]; toll-like receptor signaling pathway [GO:0002224]; tumor necrosis factor-mediated signaling pathway [GO:0033209] |
| 1726.88 | 983.04 | 1182.26 | 752.9 | 775.37 | 1595.04 | 192.75 | 263.19 | 2079.29 | 1173.19 | 679.13 | 353.89 | 714.04 | 641.34 | 609.52 | 242.82 | 396.42 | 582.07 | 14943.14 | Lecithin retinol acyltransferase (EC 2.3.1.135) (Phosphatidylcholine--retinol O-acyltransferase) | LRAT | retinol metabolic process [GO:0042572]; visual perception [GO:0007601]; vitamin A metabolic process [GO:0006776] |

| | | | | | | | | | | | | | | | | | | | | | | |
|---|---|---|---|---|---|---|---|---|---|---|---|---|---|---|---|---|---|---|---|---|---|---|
| 572.53 | | 1558.85 | 1026.8 | 233.44 | 803.85 | 1044.61 | 1448.54 | 599.7 | 734.98 | 173.81 | 325.09 | 1212.49 | 785.45 | 904.24 | 877.74 | 1097.93 | 597.94 | 938.61 | 14936.6 | Rho GTPase-activating protein 6 (Rho-type GTPase-activating protein 6) (Rho-type GTPase-activating protein RhoGAPX-1) | ARHGAP6 RHOGAP6 | actin filament organization [GO:0007015]; actin filament polymerization [GO:0030041]; focal adhesion assembly [GO:0048041]; negative regulation of focal adhesion assembly [GO:0051895]; negative regulation of stress fiber assembly [GO:0051497]; positive regulation of intracellular signal transduction [GO:1902533]; positive regulation of phospholipase C/protein kinase C signal transduction [GO:0141214]; regulation of small GTPase mediated signal transduction [GO:0051056]; Rho protein signal transduction [GO:0007266] |
| 1641.58 | | 689.9 | 431.83 | 693.43 | 1155.49 | 1503.63 | 85.07 | 309.03 | 510.48 | 834.07 | 354.8 | 490.63 | 1165.84 | 2373.58 | 550.94 | 514.04 | 814.32 | 816.11 | 14934.77 | Tyrosine-protein phosphatase non-receptor type 23 (EC 3.1.3.48) | Ptpn23 Kiaa1471 | cilium assembly [GO:0060271]; early endosome to late endosome transport [GO:0045022]; endocytic recycling [GO:0032456]; negative regulation of epithelial cell migration [GO:0010633]; positive regulation of adherens junction organization [GO:1903393]; positive regulation of early endosome to late endosome transport [GO:2000643]; positive regulation of homophilic cell adhesion [GO:1903387]; positive regulation of Wnt protein secretion [GO:0061357]; protein transport [GO:0015031]; ubiquitin-dependent protein catabolic process via the multivesicular body sorting pathway [GO:0043162] |

| | | | | | | | | | | | | | | | | | | | | | |
|---|---|---|---|---|---|---|---|---|---|---|---|---|---|---|---|---|---|---|---|---|---|
| 340.86 | 1279.02 | 1076 | 345.9 | 1491.66 | 960.34 | 1594.51 | 652.81 | 430.47 | 181.66 | 481.56 | 1260.03 | 609.45 | 1475.02 | 612.24 | 911.01 | 677.73 | 548.76 | 14929.03 | Replication protein A 32 kDa subunit-B (RP-A p32) (Replication factor A protein 2) (RF-A protein 2) (Replication protein A 34 kDa subunit) (RP-A p34) | rpa2-b repa2 rpa32 rpa34 | base-excision repair [GO:0006284]; DNA replication [GO:0006260]; DNA replication checkpoint signaling [GO:0000076]; double-strand break repair via homologous recombination [GO:0000724]; mismatch repair [GO:0006298]; nucleotide-excision repair [GO:0006289]; protein localization to chromosome [GO:0034502]; regulation of DNA damage checkpoint [GO:2000001]; telomere maintenance [GO:0000723] |
| 1818.58 | 859.05 | 1322.84 | 373.7 | 1094.03 | 1670.55 | 260.56 | 391.39 | 91.23 | 321.37 | 396.55 | 642.22 | 1148.55 | 1914.44 | 878.19 | 466.5 | 941.93 | 310.08 | 14901.76 | Carboxy-terminal domain RNA polymerase II polypeptide A small phosphatase 2 (EC 3.1.3.16) (Nuclear LIM interactor-interacting factor 2) (NLI-interacting factor 2) (Protein OS-4) (Small C-terminal domain phosphatase 2) (Small CTD phosphatase 2) (SCP2) | CTDSP2 NIF2 OS4 SCP2 | negative regulation of G1/S transition of mitotic cell cycle [GO:2000134]; protein dephosphorylation [GO:0006470] |
| 21.23 | 1.04 | 0 | 273.87 | 0 | 6.1 | 7641.98 | 0.75 | 0 | 64.18 | 189.91 | 1.43 | 12.88 | 1.26 | 0 | 6511.14 | 155.27 | 5.28 | 14886.32 | Pre-mRNA-splicing factor 18 (PRP18 homolog) | prpf18 si:ch211-220f12.3 | generation of catalytic spliceosome for second transesterification step [GO:0000350] |
| 3676.79 | 591.66 | 1172.03 | 305.6 | 288.85 | 3343.24 | 23.2 | 156.18 | 239.5 | 383.57 | 224.04 | 750.73 | 2029.27 | 273.18 | 833.94 | 110.93 | 127.45 | 347.32 | 14877.48 | Complement C1q tumor necrosis factor-related protein 6 | C1QTNF6 CTRP6 UNQ581/PRO1151 | |

| | | | | | | | | | | | | | | | | | | | | | | |
|---|---|---|---|---|---|---|---|---|---|---|---|---|---|---|---|---|---|---|---|---|---|---|
| 1152.57 | | 632.81 | 1098.38 | 609.66 | 567.06 | 1590.27 | 1054.29 | 260.3 | 394.85 | 357.81 | 233.27 | 765.96 | 2068.93 | 809.52 | 737.82 | 1196.07 | 661.41 | 674.85 | 14865.83 | T-cell surface glycoprotein CD4 (T-cell surface antigen T4/Leu-3) (CD antigen CD4) | CD4 | adaptive immune response [GO:0002250]; calcium-mediated signaling [GO:0019722]; cell adhesion [GO:0007155]; cell surface receptor protein tyrosine kinase signaling pathway [GO:0007169]; cell surface receptor signaling pathway [GO:0007166]; cellular response to granulocyte macrophage colony-stimulating factor stimulus [GO:0097011]; cellular response to ionomycin [GO:1904637]; defense response to Gram-negative bacterium [GO:0050829]; enzyme-linked receptor protein signaling pathway [GO:0007167]; helper T cell enhancement of adaptive immune response [GO:0035397]; immune response [GO:0006955]; interleukin-15-mediated signaling pathway [GO:0035723]; macrophage differentiation [GO:0030225]; maintenance of protein location in cell [GO:0032507]; positive regulation of calcium ion transport into cytosol [GO:0010524]; positive regulation of |
| 956.27 | | 1107.59 | 1011.92 | 200.08 | 1252.97 | 1680.08 | 128.02 | 728.75 | 290.51 | 304.79 | 602.5 | 1057.3 | 870.15 | 2326.31 | 590.14 | 188.26 | 1000.61 | 541.66 | 14837.91 | Erythropoietin receptor (EPO-R) | EPOR | brain development [GO:0007420]; cytokine-mediated signaling pathway [GO:0019221]; decidualization [GO:0046697]; heart development [GO:0007507]; positive regulation of cell population proliferation [GO:0008284] |

| | | | | | | | | | | | | | | | | | | | | | |
|---|---|---|---|---|---|---|---|---|---|---|---|---|---|---|---|---|---|---|---|---|---|
| 1584.41 | 861.04 | 799.65 | 504.04 | 970.66 | 1001.85 | 499.35 | 501.32 | 278.5 | 652.15 | 439.68 | 677.51 | 2059.17 | 992.29 | 388.69 | 606.65 | 1406.49 | 600.13 | 14823.58 | Fibroblast growth factor 11 (FGF-11) (Fibroblast growth factor homologous factor 3) (FHF-3) | FGF11 FHF3 | cell-cell signaling [GO:0007267]; nervous system development [GO:0007399]; neurogenesis [GO:0022008]; signal transduction [GO:0007165] |
| 446.7 | 1365.77 | 369.62 | 98.92 | 1721.15 | 331.75 | 14.19 | 673.19 | 1092.98 | 50.81 | 435.19 | 489.61 | 426.91 | 3505.82 | 1250.93 | 320.5 | 819.29 | 1408.81 | 14822.14 | LIM domain-binding protein 1 (LDB-1) (Carboxyl-terminal LIM domain-binding protein 2) (CLIM-2) (LIM domain-binding factor CLIM2) (cLdb1) (Neural Src-interacting protein) (Nuclear LIM interactor) | LDB1 CLIM2 NSIP | anterior/posterior axis specification [GO:0009948]; negative regulation of erythrocyte differentiation [GO:0045647]; negative regulation of transcription by RNA polymerase II [GO:0000122]; nervous system development [GO:0007399]; neuron differentiation [GO:0030182]; positive regulation of transcription by RNA polymerase II [GO:0045944]; Wnt signaling pathway [GO:0016055] |
| 1562.57 | 768.6 | 630.08 | 362.45 | 629.93 | 1183.02 | 1772 | 295.24 | 224.52 | 382.39 | 200.72 | 461.25 | 2333.4 | 787.31 | 850.41 | 1331.33 | 603.41 | 436.49 | 14815.12 | Charged multivesicular body protein 2a (Chromatin-modifying protein 2a) (CHMP2a) | chmp2a TNeu078g14.1 | exit from mitosis [GO:0010458]; nuclear membrane reassembly [GO:0031468]; protein transport [GO:0015031]; vacuolar transport [GO:0007034] |

| | | | | | | | | | | | | | | | | | | | | | |
|---|---|---|---|---|---|---|---|---|---|---|---|---|---|---|---|---|---|---|---|---|---|
| 838.95 | 1484.52 | 944.43 | 232.73 | 1220.83 | 963.12 | 797.45 | 857.72 | 597.15 | 205.06 | 433.83 | 694.42 | 1068.9 | 1316.66 | 730.08 | 608.81 | 1151.23 | 631.07 | 14776.96 | Large ribosomal subunit protein uL15m (39S ribosomal protein L15, mitochondrial) (L15mt) (MRP-L15) | mrpl15 | translation [GO:0006412] |
| 31.69 | 0 | 2.55 | 232.97 | 0 | 10.1 | 7273.01 | 0 | 0 | 61.74 | 194.32 | 0.74 | 3.84 | 0 | 0 | 6748.63 | 173.55 | 10.6 | 14743.74 | Ankycorbin (Ankyrin repeat and coiled-coil structure-containing protein) (Retinoic acid-induced protein 14) | Rai14 | cell differentiation [GO:0030154]; spermatogenesis [GO:0007283] |
| 416 | 732.97 | 1761.57 | 323.91 | 540.05 | 477.1 | 1313.73 | 304.1 | 986.55 | 262.47 | 357.55 | 1993.65 | 587.75 | 1150.67 | 1058.92 | 793.66 | 609.35 | 1070.5 | 14740.5 | Peptidyl-prolyl cis-trans isomerase FKBP14 (PPIase FKBP14) (EC 5.2.1.8) (22 kDa FK506-binding protein) (22 kDa FKBP) (FKBP-22) (FK506-binding protein 14) (FKBP-14) (Rotamase) | FKBP14 FKBP22 UNQ322/PRO381 | |
| 533.94 | 385.01 | 345.69 | 76.81 | 265.77 | 589.26 | 176.36 | 135.95 | 38.38 | 221.46 | 122.59 | 164.39 | 524.47 | 459.79 | 188.94 | 3006.47 | 7277.63 | 219.2 | 14732.11 | Protein jagged-2 (Jagged2) | Jag2 | auditory receptor cell fate commitment [GO:0009912]; cell fate determination [GO:0001709]; epithelial cell apoptotic process involved in palatal shelf morphogenesis [GO:1990134]; gamma-delta T cell differentiation [GO:0042492]; in utero embryonic development [GO:0001701]; morphogenesis of embryonic epithelium [GO:0016331]; Notch signaling pathway [GO:0007219]; odontogenesis of dentin-containing tooth [GO:0042475]; positive regulation of Notch signaling pathway [GO:0045747]; regulation of cell adhesion [GO:0030155]; regulation of cell population proliferation [GO:0042127]; respiratory system process [GO:0003016]; sensory perception of sound [GO:0007605]; skeletal system development [GO:0001501]; spermatogenesis [GO:0007283]; thymic T cell selection [GO:0045061] |

| | | | | | | | | | | | | | | | | | | | | | |
|---|---|---|---|---|---|---|---|---|---|---|---|---|---|---|---|---|---|---|---|---|---|
| 1681.17 | 19.44 | 672.03 | 465.42 | 34.33 | 1612.5 | 13.65 | 14.65 | 3162.31 | 643.93 | 13.77 | 813.08 | 393.99 | 28.88 | 1947.05 | 26.22 | 33.6 | 3119.3 | 14695.32 | Little elongation complex subunit 2 (Interactor of little elongator complex ELL subunit 2) (NMDA receptor-regulated protein 2) | ICE2 BRCC1 NARG2 UNQ3101/PRO10100 | positive regulation of transcription by RNA polymerase III [GO:0045945]; snRNA transcription by RNA polymerase II [GO:0042795]; snRNA transcription by RNA polymerase III [GO:0042796] |
| 251.07 | 4033.86 | 334.79 | 429.06 | 3467.49 | 95.66 | 2.33 | 2570.92 | 7.59 | 148.54 | 657.47 | 546.71 | 54.75 | 814.71 | 12.59 | 31.7 | 1184.18 | 14.15 | 14657.57 | GON-4-like protein (GON-4 homolog) | GON4L GON4 KIAA1606 | regulation of DNA-templated transcription [GO:0006355] |
| 476.81 | 143.6 | 1460.28 | 241.86 | 206.79 | 706.45 | 5229.76 | 132.97 | 206.46 | 230.42 | 163.57 | 422.7 | 663.48 | 235.77 | 362.42 | 3040.98 | 393.62 | 339.04 | 14656.98 | Palmitoyltransferase ZDHHC16A (EC 2.3.1.225) (Zinc finger DHHC domain-containing protein 16A) (DHHC-16A) | zdhhc16a zdhhc16 | commitment of multipotent stem cells to neuronal lineage in forebrain [GO:0021898]; DNA damage response [GO:0006974]; eye development [GO:0001654]; fibroblast growth factor receptor signaling pathway involved in forebrain neuron fate commitment [GO:0021899]; heart development [GO:0007507]; protein palmitoylation [GO:0018345]; telencephalon development [GO:0021537] |

| | | | | | | | | | | | | | | | | | | | | | | |
|---|---|---|---|---|---|---|---|---|---|---|---|---|---|---|---|---|---|---|---|---|---|---|
| 484.41 | | 1141.63 | 1122.63 | 251.76 | 1005.86 | 828.32 | 1526.96 | 434.38 | 828.52 | 207.34 | 316.29 | 772.83 | 864.16 | 1147.2 | 892.59 | 1314.01 | 605.65 | 911.34 | 14655.88 | Actin-related protein 3 (Actin-like protein 3) | actr3 arp3 | Arp2/3 complex-mediated actin nucleation [GO:0034314]; positive regulation of transcription by RNA polymerase II [GO:0045944] |
| 406.43 | | 779.94 | 478.99 | 192.89 | 628.78 | 620.49 | 2966.66 | 276.64 | 861.29 | 122.58 | 265.78 | 849.8 | 600.88 | 1671.69 | 889.98 | 1655.09 | 631.69 | 755.34 | 14654.94 | Protein FAM210B, mitochondrial | FAM210B C20orf108 PSEC0265 | cellular response to estradiol stimulus [GO:0071392]; erythrocyte maturation [GO:0043249]; inflammatory response [GO:0006954]; positive regulation of erythrocyte differentiation [GO:0045648]; reactive oxygen species metabolic process [GO:0072593]; skin development [GO:0043588]; spleen development [GO:0048536] |
| 532.32 | | 322.74 | 1566.64 | 211.51 | 365.59 | 645.84 | 835.04 | 129.59 | 2436.59 | 180.47 | 146.26 | 1165.26 | 652.68 | 600.62 | 1535.14 | 828.28 | 361.47 | 2126.45 | 14642.49 | Clathrin light chain B (Lcb) | CLTB CLTLB | clathrin-dependent endocytosis [GO:0072583]; intracellular protein transport [GO:0006886] |

| | | | | | | | | | | | | | | | | | | | | | |
|---|---|---|---|---|---|---|---|---|---|---|---|---|---|---|---|---|---|---|---|---|---|
| 774.89 | 942.41 | 1122.15 | 184.57 | 702.07 | 900.88 | 335.61 | 279.85 | 1513.87 | 172.56 | 270.02 | 1050.71 | 644.89 | 1653.14 | 1382.6 | 358.17 | 591.18 | 1751.4 | 14630.97 | Protein lyl-1 (Class A basic helix-loop-helix protein 18) (bHLHa18) (Lymphoblastic leukemia-derived sequence 1) | LYL1 BHLHA18 | B cell differentiation [GO:0030183]; blood vessel maturation [GO:0001955]; definitive hemopoiesis [GO:0060216]; positive regulation of DNA-templated transcription [GO:0045893]; regulation of DNA-templated transcription [GO:0006355]; regulation of transcription by RNA polymerase II [GO:0006357] |
| 508.68 | 412.09 | 680.59 | 237.69 | 411.14 | 737.56 | 3377.51 | 165.55 | 1067.03 | 199.24 | 253.77 | 518.42 | 436.88 | 704.62 | 845.8 | 2760.16 | 375.21 | 936.31 | 14628.25 | RNA-binding protein 25 (Arg/Glu/Asp-rich protein of 120 kDa) (RED120) (Protein S164) (RNA-binding motif protein 25) (RNA-binding region-containing protein 7) | RBM25 RNPC7 | mRNA processing [GO:0006397]; regulation of alternative mRNA splicing, via spliceosome [GO:0000381]; regulation of apoptotic process [GO:0042981]; RNA splicing [GO:0008380] |
| 0 | 7.8 | 1864.4 | 0 | 0.96 | 2.61 | 0.35 | 0.74 | 4058.52 | 1.18 | 1.31 | 880.08 | 1.53 | 0.77 | 3479.17 | 14.35 | 2.93 | 4305.46 | 14622.16 | Endophilin-B2 (SH3 domain-containing GRB2-like protein B2) | SH3GLB2 KIAA1848 PP578 | membrane organization [GO:0061024] |
| 1224.02 | 856.06 | 1224.55 | 360.25 | 816.8 | 1301.47 | 56.01 | 393.08 | 1443.43 | 395.12 | 262.73 | 988.23 | 992.86 | 938.97 | 1291.26 | 154.26 | 507.92 | 1411.55 | 14618.57 | Serine/threonine-protein kinase tousled-like 2 (EC 2.7.11.1) (PKU-alpha) (Tousled-like kinase 2) | tlk2 zgc:136697 | chromosome segregation [GO:0007059]; intracellular signal transduction [GO:0035556] |

| | | | | | | | | | | | | | | | | | | | | | |
|---|---|---|---|---|---|---|---|---|---|---|---|---|---|---|---|---|---|---|---|---|---|
| 5491.07 | 127.53 | 1726.74 | 666.94 | 101.88 | 3870.48 | 5.83 | 52.32 | 344.18 | 892.94 | 50.01 | 333.01 | 211.53 | 99.57 | 49.79 | 40.26 | 242.88 | 260.95 | 14567.91 | Membrane-bound transcription factor site-2 protease (EC 3.4.24.85) (Endopeptidase S2P) (Sterol regulatory element-binding proteins intramembrane protease) (SREBPs intramembrane protease) | MBTPS2 S2P | ATF6-mediated unfolded protein response [GO:0036500]; bone maturation [GO:0070977]; cholesterol metabolic process [GO:0008203]; endoplasmic reticulum unfolded protein response [GO:0030968]; membrane protein intracellular domain proteolysis [GO:0031293]; mitotic G2 DNA damage checkpoint signaling [GO:0007095]; positive regulation of cholesterol biosynthetic process [GO:0045542]; positive regulation of transcription by RNA polymerase II [GO:0045944]; protein maturation [GO:0051604]; regulation of cholesterol biosynthetic process [GO:0045540]; regulation of response to endoplasmic reticulum stress [GO:1905897]; response to endoplasmic reticulum stress [GO:0034976] |
| 34.53 | 3.99 | 771.43 | 4.7 | 9.71 | 137.96 | 3.11 | 9.45 | 4415.88 | 3.85 | 2.86 | 1650.34 | 35.51 | 6.9 | 3251.09 | 2.91 | 4.09 | 4207.55 | 14555.86 | Semaphorin-4B (Semaphorin-C) (Sema C) | Sema4b Kiaa1745 Semac SemC | cell differentiation [GO:0030154]; nervous system development [GO:0007399] |
| 0.52 | 4.05 | 1858.19 | 0.15 | 5.9 | 2.21 | 98.99 | 1 | 4197.62 | 1.31 | 3.8 | 1236.62 | 1.46 | 7.53 | 3802.15 | 54.62 | 5.99 | 3266.47 | 14548.58 | Low-density lipoprotein receptor-related protein 1 (LRP-1) (Alpha-2-macroglobulin receptor) (A2MR) | LRP1 | endocytosis [GO:0006897] |

| | | | | | | | | | | | | | | | | | | | | | |
|---|---|---|---|---|---|---|---|---|---|---|---|---|---|---|---|---|---|---|---|---|---|
| 332.81 | 314.81 | 1744.34 | 178.12 | 305.31 | 1214.32 | 16.51 | 282.17 | 1856.2 | 100.17 | 145.33 | 1018.95 | 609.5 | 562.2 | 3312.33 | 63.53 | 448.39 | 2032.64 | 14537.63 | Chemokine-like protein TAFA-5 | TAFA5 FAM19A5 | G protein-coupled receptor signaling pathway [GO:0007186]; negative regulation of vascular associated smooth muscle cell proliferation [GO:1904706]; negative regulation of vascular wound healing [GO:0061044] |
| 2.7 | 2.38 | 5303.63 | 2.72 | 7.32 | 1041.27 | 0.37 | 4.05 | 23.59 | 4.5 | 5.25 | 7965.58 | 2.14 | 4.64 | 78.27 | 21.56 | 26.74 | 7.94 | 14504.65 | Pre-mRNA-processing-splicing factor 8 (220 kDa U5 snRNP-specific protein) (PRP8 homolog) (Splicing factor Prp8) (p220) | PRPF8 PRPC8 | cellular response to lipopolysaccharide [GO:0071222]; cellular response to tumor necrosis factor [GO:0071356]; mRNA processing [GO:0006397]; mRNA splicing, via spliceosome [GO:0000398]; RNA splicing [GO:0008380]; RNA splicing, via transesterification reactions [GO:0000375]; spliceosomal tri-snRNP complex assembly [GO:0000244] |
| 124.2 | 733.48 | 209.63 | 172.76 | 1494.82 | 1425.63 | 3.34 | 1415.57 | 105.34 | 215.04 | 926.72 | 840.48 | 1165.11 | 2275.63 | 792.21 | 149.82 | 1583.39 | 866.78 | 14499.95 | THAP domain-containing protein 11 | thap11 zgc:65871 | regulation of transcription by RNA polymerase II [GO:0006357] |

| | | | | | | | | | | | | | | | | | | | | | |
|---|---|---|---|---|---|---|---|---|---|---|---|---|---|---|---|---|---|---|---|---|---|
| 309.75 | 848.31 | 590.59 | 634.17 | 1536.71 | 738.07 | 976.73 | 818.56 | 574.19 | 366.67 | 709.02 | 964.83 | 89.23 | 586.41 | 52.63 | 1593.6 | 2226.65 | 882.84 | 14498.96 | Heat shock cognate 71 kDa protein (EC 3.6.4.10) (Heat shock 70 kDa protein 8) | HSPA8 HSC70 | chaperone cofactor-dependent protein refolding [GO:0051085]; clathrin coat disassembly [GO:0072318]; mRNA processing [GO:0006397]; negative regulation of DNA-templated transcription [GO:0045892]; protein refolding [GO:0042026]; protein targeting to lysosome involved in chaperone-mediated autophagy [GO:0061740]; RNA splicing [GO:0008380]; synaptic vesicle uncoating [GO:0016191] |
| 1203.97 | 326.5 | 2425.92 | 379.81 | 1514.1 | 1105.54 | 118.17 | 450.14 | 279.79 | 707.33 | 671.91 | 1286.29 | 711.96 | 310.47 | 666.33 | 318.23 | 1627.54 | 383.95 | 14487.95 | Guanine nucleotide exchange factor VAV2 (VAV-2) | Vav2 | angiogenesis [GO:0001525]; cell migration [GO:0016477]; cell projection assembly [GO:0030031]; lamellipodium assembly [GO:0030032]; positive regulation of phosphatidylinositol 3-kinase/protein kinase B signal transduction [GO:0051897]; regulation of cell size [GO:0008361]; small GTPase-mediated signal transduction [GO:0007264] |

| | | | | | | | | | | | | | | | | | | | | | |
|---|---|---|---|---|---|---|---|---|---|---|---|---|---|---|---|---|---|---|---|---|---|
| 155.06 | 532.82 | 887.78 | 134.21 | 372.56 | 776.62 | 21.41 | 224.7 | 2246.02 | 114.77 | 220.78 | 1044.84 | 76.18 | 969.16 | 2614.88 | 262.32 | 667.85 | 3161.64 | 14483.6 | Probable E3 ubiquitin-protein ligase HERC1 (EC 2.3.2.26) (HECT domain and RCC1-like domain-containing protein 1) (HECT-type E3 ubiquitin transferase HERC1) (p532) (p619) | HERC1 | autophagy [GO:0006914]; bone mineralization [GO:0030282]; bone remodeling [GO:0046849]; cerebellar Purkinje cell differentiation [GO:0021702]; corpus callosum development [GO:0022038]; gene expression [GO:0010467]; negative regulation of autophagy [GO:0010507]; neuromuscular process controlling balance [GO:0050885]; neuron projection development [GO:0031175]; protein ubiquitination [GO:0016567] |
| 1071.93 | 765.15 | 942.39 | 427.04 | 1070.64 | 626.14 | 7.57 | 344.59 | 2030.94 | 321.43 | 275.65 | 1028.48 | 349.07 | 562.08 | 1713.18 | 234.35 | 943.76 | 1732.4 | 14446.79 | Inhibitor of nuclear factor kappa-B kinase subunit alpha (I kappa-B kinase alpha) (IKK-A) (IKK-alpha) (IkBKA) (IkappaB kinase) (EC 2.7.11.10) (Conserved helix-loop-helix ubiquitous kinase) (I-kappa-B kinase 1) (IKK1) (Nuclear factor NF-kappa-B inhibitor kinase alpha) (NFKBIKA) | chuk ikk1 zgc:56539 | cellular response to tumor necrosis factor [GO:0071356]; central nervous system development [GO:0007417]; epiboly involved in gastrulation with mouth forming second [GO:0055113]; negative regulation of apoptotic process [GO:0043066]; negative regulation of canonical NF-kappaB signal transduction [GO:0043124]; positive regulation of canonical NF-kappaB signal transduction [GO:0043123]; positive regulation of transcription by RNA polymerase II [GO:0045944]; somite specification [GO:0001757]; tumor necrosis factor-mediated signaling pathway [GO:0033209] |
| 20.66 | 21.15 | 1011.79 | 36.13 | 7.46 | 22.46 | 4639.47 | 18.69 | 2064.52 | 23.81 | 140.23 | 531.71 | 11.72 | 38.68 | 1194.01 | 2942.57 | 105.14 | 1599.68 | 14429.88 | RNA-binding protein NOB1 (EC 3.1.-.-) | NOB1 | maturation of SSU-rRNA [GO:0030490] |

| | | | | | | | | | | | | | | | | | | | | | |
|---|---|---|---|---|---|---|---|---|---|---|---|---|---|---|---|---|---|---|---|---|---|
| 814.08 | 1373.7 | 821.81 | 265.84 | 908.47 | 1296.52 | 110.39 | 523.29 | 274.03 | 297.96 | 582.66 | 710.65 | 929.14 | 2055.58 | 780.62 | 515.02 | 1339.37 | 824.05 | 14423.18 | Collagen alpha-1(XXVII) chain B | col27a1b col27a1 | bone mineralization [GO:0030282]; notochord morphogenesis [GO:0048570]; skeletal system development [GO:0001501] |
| 959.85 | 763.37 | 795.87 | 351.2 | 638.38 | 1255.18 | 2344.39 | 418.02 | 480.71 | 364.62 | 322.01 | 598.94 | 913.54 | 783.45 | 630.69 | 1790.71 | 422.5 | 568.9 | 14402.33 | TBC1 domain family member 1 | TBC1D1 KIAA1108 | |
| 250.24 | 758.99 | 245.61 | 599.92 | 1774.51 | 1628.47 | 10.97 | 627.5 | 1097.26 | 1565.95 | 1078.97 | 531.41 | 508.71 | 1297.28 | 968.54 | 197.8 | 622.3 | 637.44 | 14401.87 | Anoctamin-7 (New gene expressed in prostate homolog) (Transmembrane protein 16G) | Ano7 Ngep Tmem16g | calcium activated galactosylceramide scrambling [GO:0061591]; calcium activated phosphatidylcholine scrambling [GO:0061590]; calcium activated phosphatidylserine scrambling [GO:0061589]; chloride transport [GO:0006821]; establishment of localization in cell [GO:0051649] |

| | | | | | | | | | | | | | | | | | | | | | |
|---|---|---|---|---|---|---|---|---|---|---|---|---|---|---|---|---|---|---|---|---|---|
| 586.62 | 749.4 | 719.17 | 257.81 | 685.87 | 981.74 | 3249.47 | 286.18 | 522.02 | 282.72 | 273.43 | 620.11 | 608.99 | 616.52 | 606.16 | 2182.97 | 440.25 | 719.01 | 14388.44 | Delta-1-pyrroline-5-carboxylate synthase (P5CS) (Aldehyde dehydrogenase family 18 member A1) [Includes: Glutamate 5-kinase (GK) (EC 2.7.2.11) (Gamma-glutamyl kinase);Gamma-glutamyl phosphate reductase (GPR) (EC 1.2.1.41) (Glutamate-5-semialdehyde dehydrogenase) (Glutamyl-gamma-semialdehyde dehydrogenase)] | ALDH18A1 | citrulline biosynthetic process [GO:0019240]; glutamate metabolic process [GO:0006536]; L-proline biosynthetic process [GO:0055129]; ornithine biosynthetic process [GO:0006592]; proline biosynthetic process [GO:0006561] |
| 986.04 | 1190.91 | 567.77 | 225.33 | 1114.33 | 666.8 | 208.84 | 251.59 | 186.84 | 223.72 | 738.12 | 429.4 | 2638.26 | 1757.34 | 587.27 | 1057.4 | 1180.02 | 367.37 | 14377.35 | Very-long-chain 3-oxoacyl-CoA reductase-A (EC 1.1.1.330) (17-beta-hydroxysteroid dehydrogenase 12-A) (17-beta-HSD 12-A) (zf3.1) (zfHSD17B12A) (3-ketoacyl-CoA reductase) (KAR) (Estradiol 17-beta-dehydrogenase 12-A) (EC 1.1.1.62) | hsd17b12a zgc:55589 | estrogen biosynthetic process [GO:0006703]; fatty acid biosynthetic process [GO:0006633] |
| 1299.37 | 1722.96 | 1468.92 | 256.75 | 940.87 | 1019.7 | 46.75 | 779.45 | 848.26 | 406.31 | 505.57 | 1497.79 | 559.75 | 921.86 | 278.17 | 155.24 | 1364.96 | 304.64 | 14377.32 | PI-PLC X domain-containing protein 1 | PLCXD1 | lipid metabolic process [GO:0006629] |

| | | | | | | | | | | | | | | | | | | | | | |
|---|---|---|---|---|---|---|---|---|---|---|---|---|---|---|---|---|---|---|---|---|---|
| 447.15 | 754.9 | 133.24 | 830.32 | 344.37 | 728.11 | 4979.7 | 675.28 | 209.43 | 921.23 | 533.8 | 388.06 | 366.5 | 259.99 | 155.33 | 2236.61 | 276.67 | 136.19 | 14376.88 | Mucosa-associated lymphoid tissue lymphoma translocation protein 1 (EC 3.4.22.-) (MALT lymphoma-associated translocation) (Paracaspase) | MALT1 MLT | B cell activation [GO:0042113]; B-1 B cell differentiation [GO:0001923]; defense response [GO:0006952]; innate immune response [GO:0045087]; lipopolysaccharide-mediated signaling pathway [GO:0031663]; negative regulation of apoptotic process [GO:0043066]; nuclear export [GO:0051168]; positive regulation of canonical NF-kappaB signal transduction [GO:0043123]; positive regulation of interleukin-1 beta production [GO:0032731]; positive regulation of interleukin-2 production [GO:0032743]; positive regulation of protein ubiquitination [GO:0031398]; positive regulation of T cell cytokine production [GO:0002726]; positive regulation of T-helper 17 cell differentiation [GO:2000321]; proteolysis [GO:0006508]; proteolysis involved in protein catabolic process [GO:0051603]; regulation of apoptotic process [GO:0042981]; regulation of T cell receptor signaling pathway [GO:0050856]; response to fungus |
| 3084.74 | 238.11 | 179.55 | 775.68 | 267.21 | 253.17 | 6.55 | 115.24 | 30.7 | 870.93 | 201.01 | 109.49 | 6516.93 | 617.62 | 334.04 | 245.8 | 406.76 | 116.25 | 14369.78 | Neural cell adhesion molecule L1-like protein (Close homolog of L1) [Cleaved into: Processed neural cell adhesion molecule L1-like protein] | CHL1 CALL | adult locomotory behavior [GO:0008344]; axon guidance [GO:0007411]; cell adhesion [GO:0007155]; cognition [GO:0050890]; exploration behavior [GO:0035640]; negative regulation of neuron apoptotic process [GO:0043524]; neuron migration [GO:0001764]; signal transduction [GO:0007165] |

| | | | | | | | | | | | | | | | | | | | | | |
|---|---|---|---|---|---|---|---|---|---|---|---|---|---|---|---|---|---|---|---|---|---|
| 990.64 | 794.69 | 1150.78 | 289.8 | 822.28 | 1014.21 | 162.91 | 538.66 | 1016.54 | 326.35 | 392.62 | 866.4 | 608.7 | 1336.05 | 1278.71 | 225.76 | 850.35 | 1696.83 | 14362.28 | Spliceosome associated factor 3, U4/U6 recycling protein (Squamous cell carcinoma antigen recognized by T-cells 3) (SART-3) | sart3 egy si:ch211-191d15.4 wu:fc51h03 | exocrine pancreas development [GO:0031017]; hematopoietic stem cell homeostasis [GO:0061484]; lymphocyte differentiation [GO:0030098]; mRNA splicing, via spliceosome [GO:0000398]; nucleosome assembly [GO:0006334]; regulation of hematopoietic stem cell proliferation [GO:1902033]; regulation of intrinsic apoptotic signaling pathway by p53 class mediator [GO:1902253]; spliceosomal complex assembly [GO:0000245]; spliceosomal snRNP assembly [GO:0000387]; thymus development [GO:0048538]; transcription elongation-coupled chromatin remodeling [GO:0140673] |
| 1289.95 | 637.21 | 2053.31 | 465.17 | 791.78 | 2029.69 | 166.78 | 248.88 | 2196.44 | 448.63 | 254.13 | 1241.1 | 469.99 | 137.22 | 715.73 | 269.68 | 90.33 | 834.87 | 14340.89 | Dihydrolipoyl dehydrogenase, mitochondrial (EC 1.8.1.4) (Dihydrolipoamide dehydrogenase) | DLD | 2-oxoglutarate metabolic process [GO:0006103]; branched-chain amino acid catabolic process [GO:0009083]; gastrulation [GO:0007369]; mitochondrial electron transport, NADH to ubiquinone [GO:0006120]; proteolysis [GO:0006508]; pyruvate metabolic process [GO:0006090]; regulation of membrane potential [GO:0042391]; sperm capacitation [GO:0048240] |

| | | | | | | | | | | | | | | | | | | | | | |
|---|---|---|---|---|---|---|---|---|---|---|---|---|---|---|---|---|---|---|---|---|---|
| 29.59 | 43.08 | 531.63 | 5.44 | 56.05 | 38.67 | 1.76 | 27.6 | 3647.75 | 3.67 | 12.58 | 1399.3 | 9.6 | 9.87 | 3984.53 | 2.09 | 28.53 | 4497.64 | 14329.38 | Delta-like protein 4 (Drosophila Delta homolog 4) (Delta4) | DLL4 UNQ1895/PRO 4341 | angiogenesis [GO:0001525]; aortic valve morphogenesis [GO:0003180]; blood vessel lumenization [GO:0072554]; blood vessel remodeling [GO:0001974]; branching involved in blood vessel morphogenesis [GO:0001569]; cardiac atrium morphogenesis [GO:0003209]; cardiac ventricle morphogenesis [GO:0003208]; cellular response to fibroblast growth factor stimulus [GO:0044344]; cellular response to vascular endothelial growth factor stimulus [GO:0035924]; dorsal aorta morphogenesis [GO:0035912]; negative regulation of blood vessel endothelial cell proliferation involved in sprouting angiogenesis [GO:1903588]; negative regulation of cell migration involved in sprouting angiogenesis [GO:0090051]; negative regulation of cell population proliferation [GO:0008285]; negative regulation of endothelial cell migration [GO:0010596]; negative regulation of gene expression [GO:0010629]; negative regulation of Notch signaling pathway [GO:0045746]; negative regulation of |
| 698.79 | 550.29 | 777.69 | 340.95 | 499.36 | 719.54 | 3868.65 | 317.37 | 365.46 | 194.55 | 230.79 | 452.43 | 672.47 | 723.57 | 496.42 | 2465.39 | 426.99 | 496.36 | 14297.07 | GDNF family receptor alpha-2 (GDNF receptor alpha-2) (GDNFR-alpha-2) (GFR-alpha-2) | GFRA2 | glial cell-derived neurotrophic factor receptor signaling pathway [GO:0035860]; nervous system development [GO:0007399] |
| 260.39 | 634.98 | 2882.95 | 494.4 | 806.84 | 543.82 | 467.52 | 433.94 | 1207.43 | 196.06 | 350.22 | 1751.13 | 235.71 | 752 | 867.12 | 671.34 | 554.05 | 1104.17 | 14214.07 | Snaclec 7 (C-type lectin-like 7) | | |

| | | | | | | | | | | | | | | | | | | | | | |
|---|---|---|---|---|---|---|---|---|---|---|---|---|---|---|---|---|---|---|---|---|---|
| 722.48 | 1675.32 | 857.86 | 201.92 | 1447.97 | 890.53 | 687.51 | 947.97 | 605.32 | 216.66 | 363.09 | 549.34 | 920.5 | 1125.22 | 785.37 | 530.74 | 944.81 | 734.99 | 14207.6 | Serine/threonine-protein kinase PAK 5 (EC 2.7.11.1) (p21-activated kinase 5) (PAK-5) (p21-activated kinase 7) (PAK-7) | Pak5 Pak7 | apoptotic process [GO:0006915]; cytoskeleton organization [GO:0007010]; learning [GO:0007612]; locomotory behavior [GO:0007626]; memory [GO:0007613]; negative regulation of extrinsic apoptotic signaling pathway [GO:2001237] |
| 504.98 | 415.09 | 449.96 | 316.29 | 420.93 | 485.24 | 4039.39 | 152.32 | 271.79 | 202.71 | 305.24 | 360.87 | 1114.1 | 366.18 | 535.58 | 3294.59 | 421.38 | 526.02 | 14182.66 | Sulfhydryl oxidase 2 (EC 1.8.3.2) (Neuroblastoma-derived sulfhydryl oxidase) (Quiescin Q6-like protein 1) | QSOX2 QSCN6L1 SOXN | protein folding [GO:0006457] |
| 497.31 | 772.83 | 1120.09 | 260.82 | 852.37 | 871.14 | 1843.4 | 363.64 | 1024.16 | 233.25 | 324.42 | 923.78 | 712.39 | 997.39 | 930.07 | 763.39 | 597.8 | 1089.28 | 14177.53 | Zinc finger protein 366 (Dendritic cell-specific transcript protein) (DC-SCRIPT) | ZNF366 | negative regulation of intracellular estrogen receptor signaling pathway [GO:0033147]; negative regulation of transcription by RNA polymerase II [GO:0000122]; response to estrogen [GO:0043627] |

| | | | | | | | | | | | | | | | | | | | | | |
|---|---|---|---|---|---|---|---|---|---|---|---|---|---|---|---|---|---|---|---|---|---|
| 722.29 | 1075.75 | 1198.23 | 321.16 | 634.4 | 1470.12 | 98.02 | 337.26 | 974.54 | 318.83 | 245.95 | 906.13 | 797.65 | 1538.75 | 1120.17 | 319.65 | 708.39 | 1381.59 | 14168.88 | Indoleamine 2,3-dioxygenase 2 (IDO-2) (EC 1.13.11.-) (Indoleamine 2,3-dioxygenase-like protein 1) (Indoleamine-pyrrole 2,3-dioxygenase-like protein 1) | IDO2 INDOL1 | 'de novo' NAD biosynthetic process from L-tryptophan [GO:0034354]; immune system process [GO:0002376]; L-tryptophan catabolic process to kynurenine [GO:0019441] |
| 386.93 | 493.45 | 1240.7 | 178.31 | 566.14 | 984.14 | 4.28 | 306.97 | 1675.16 | 212.25 | 205.6 | 949.24 | 619.53 | 1168.45 | 2480.43 | 262.37 | 449.61 | 1958.11 | 14141.67 | Propionyl-CoA carboxylase beta chain, mitochondrial (PCCase subunit beta) (EC 6.4.1.3) (Propanoyl-CoA:carbon dioxide ligase subunit beta) | Pccb | |
| 82.82 | 3430.48 | 0 | 176.49 | 3225.35 | 16.76 | 0 | 774.17 | 0 | 76.06 | 1045.15 | 5.9 | 43.63 | 3715.19 | 0 | 222.62 | 1306.81 | 6.06 | 14127.49 | Pleckstrin homology-like domain family B member 1 (Protein LL5-alpha) | Phldb1 Ll5 | positive regulation of basement membrane assembly involved in embryonic body morphogenesis [GO:1904261]; regulation of epithelial to mesenchymal transition [GO:0010717]; regulation of gastrulation [GO:0010470]; regulation of microtubule cytoskeleton organization [GO:0070507] |

| | | | | | | | | | | | | | | | | | | | | | |
|---|---|---|---|---|---|---|---|---|---|---|---|---|---|---|---|---|---|---|---|---|---|
| 874.97 | 608.69 | 2381.27 | 418.33 | 738.99 | 1472.05 | 645.42 | 281.92 | 804.66 | 371.78 | 280.11 | 1169.27 | 763.51 | 658.93 | 684.11 | 653.79 | 500.61 | 807.41 | 14115.82 | Y+L amino acid transporter 2 (Solute carrier family 7 member 6) (zfSlc7a6) (y(+)L-type amino acid transporter 2) (Y+LAT2) (y+LAT-2) | slc7a6 | amino acid transmembrane transport [GO:0003333]; glycine betaine transport [GO:0031460]; L-arginine transmembrane transport [GO:1903826]; neutral amino acid transport [GO:0015804]; nitric oxide biosynthetic process [GO:0006809] |
| 36.07 | 0 | 14.2 | 113.2 | 9.73 | 6.82 | 7989.25 | 6.3 | 9.66 | 81.96 | 206.97 | 0 | 37.37 | 1.18 | 10.03 | 5378.08 | 171.21 | 20.62 | 14092.65 | Stonustoxin subunit beta (SNTX subunit beta) (DELTA-synanceitoxin-Sh1b) (DELTA-SYTX-Sh1b) (Trachynilysin subunit beta) (TLY subunit beta) | | killing of cells of another organism [GO:0031640] |
| 1019.24 | 874.67 | 1162.62 | 391.24 | 901.45 | 1399.99 | 530.24 | 487.49 | 761.68 | 354.49 | 356.96 | 918.68 | 506.96 | 1294.43 | 861.01 | 646.39 | 776.84 | 847.05 | 14091.43 | Zinc finger protein AEBP2 (Adipocyte enhancer-binding protein 2 homolog) (AE-binding protein 2 homolog) | aebp2 si:dkey-158p11.2 zgc:63755 | chromatin organization [GO:0006325]; regulation of transcription by RNA polymerase II [GO:0006357] |

| | | | | | | | | | | | | | | | | | | | | | |
|---|---|---|---|---|---|---|---|---|---|---|---|---|---|---|---|---|---|---|---|---|---|
| 9.14 | 1.14 | 437.34 | 5.1 | 4.55 | 19.46 | 0 | 5.27 | 2968.55 | 4.55 | 2.38 | 874.52 | 4.25 | 17.35 | 5096.22 | 1.61 | 0.7 | 4624.6 | 14076.73 | Unconventional myosin-Ic (Myosin I beta) (MMI-beta) (MMIb) | MYO1C RCJMB04_6o17 | actin filament organization [GO:0007015]; actin filament-based movement [GO:0030048]; endocytosis [GO:0006897] |
| 440.04 | 186.07 | 5628.69 | 81.34 | 76.56 | 472.66 | 3.76 | 60.62 | 984.65 | 111.5 | 34.31 | 3222.34 | 83.74 | 292.68 | 827.16 | 15.53 | 49.9 | 1418.78 | 13990.33 | A-kinase anchor protein 13 (AKAP-13) (AKAP-Lbc) (Breast cancer nuclear receptor-binding auxiliary protein) (Guanine nucleotide exchange factor Lbc) (Human thyroid-anchoring protein 31) (Lymphoid blast crisis oncogene) (LBC oncogene) (Non-oncogenic Rho GTPase-specific GTP exchange factor) (Protein kinase A-anchoring protein 13) (PRKA13) (p47) | AKAP13 BRX HT31 LBC | adrenergic receptor signaling pathway [GO:0071875]; bone development [GO:0060348]; cardiac muscle cell differentiation [GO:0055007]; G protein-coupled receptor signaling pathway [GO:0007186]; heart development [GO:0007507]; positive regulation of canonical NF-kappaB signal transduction [GO:0043123]; positive regulation of Rho protein signal transduction [GO:0035025]; regulation of Rho protein signal transduction [GO:0035023]; regulation of sarcomere organization [GO:0060297]; regulation of small GTPase mediated signal transduction [GO:0051056] |
| 675.58 | 951.65 | 476.41 | 147.08 | 774.64 | 457.35 | 865.02 | 359.38 | 602.22 | 94.71 | 301.78 | 329.66 | 1529.27 | 2206.12 | 921.99 | 1340.14 | 1135.78 | 777.15 | 13945.93 | Calmodulin-lysine N-methyltransferase (CLNMT) (CaM KMT) (EC 2.1.1.60) | Camkmt Clnmt | methylation [GO:0032259]; mitochondrion organization [GO:0007005] |
| 2048.89 | 867.85 | 238.06 | 293.73 | 1568.2 | 244.25 | 110.77 | 416.57 | 46.45 | 355.56 | 293.89 | 122.62 | 2623.26 | 2196.98 | 287.08 | 719.74 | 1311.5 | 199.87 | 13945.27 | Casein kinase I isoform gamma-2 (CKI-gamma 2) (EC 2.7.11.1) | Csnk1g2 Ck1g2 | Wnt signaling pathway [GO:0016055] |
| 416.61 | 616.97 | 464.09 | 219.46 | 550.35 | 833.49 | 3618.05 | 495.5 | 533.99 | 172.78 | 151.03 | 427.85 | 840.83 | 813.32 | 410.05 | 1838.49 | 669.98 | 853.37 | 13926.21 | Uncharacterized oxidoreductase YtbE (EC 1.-.-.-) | ytbE BSU29050 | |

| | | | | | | | | | | | | | | | | | | | | | |
|---|---|---|---|---|---|---|---|---|---|---|---|---|---|---|---|---|---|---|---|---|---|
| 1500.77 | 113.75 | 2537.09 | 146.62 | 110.38 | 2186.96 | 4.03 | 21.14 | 58.18 | 303.57 | 23.34 | 2673.44 | 3314.78 | 52.36 | 553.71 | 79.28 | 17.58 | 200.87 | 13897.85 | Alpha-soluble NSF attachment protein (SNAP-alpha) (N-ethylmaleimide-sensitive factor attachment protein alpha) | NAPA SNAPA | apical protein localization [GO:0045176]; brain development [GO:0007420]; intra-Golgi vesicle-mediated transport [GO:0006891]; intracellular protein transport [GO:0006886]; membrane fusion [GO:0061025]; neuron differentiation [GO:0030182]; regulation of synaptic vesicle priming [GO:0010807]; SNARE complex disassembly [GO:0035494]; synaptic transmission, glutamatergic [GO:0035249]; synaptic vesicle priming [GO:0016082] |
| 13.62 | 18.08 | 3139.91 | 36.02 | 33.68 | 569.69 | 1011.8 | 31.35 | 245.56 | 37.47 | 90.75 | 5902.85 | 29 | 22.15 | 188.27 | 2332.07 | 62.45 | 99.88 | 13864.6 | Ubiquitin thioesterase otulin (EC 3.4.19.12) (Deubiquitinating enzyme otulin) (OTU domain-containing deubiquitinase with linear linkage specificity) (Ubiquitin thioesterase Gumby) | Otulin Fam105b Gum | innate immune response [GO:0045087]; negative regulation of inflammatory response [GO:0050728]; negative regulation of NF-kappaB transcription factor activity [GO:0032088]; nucleotide-binding oligomerization domain containing 2 signaling pathway [GO:0070431]; protein linear deubiquitination [GO:1990108]; proteolysis [GO:0006508]; regulation of canonical Wnt signaling pathway [GO:0060828]; regulation of tumor necrosis factor-mediated signaling pathway [GO:0010803]; sprouting angiogenesis [GO:0002040]; Wnt signaling pathway [GO:0016055] |

| | | | | | | | | | | | | | | | | | | | | | |
|---|---|---|---|---|---|---|---|---|---|---|---|---|---|---|---|---|---|---|---|---|---|
| 1025.04 | 952.55 | 895.93 | 336.2 | 1032.65 | 984.01 | 205.85 | 530.93 | 816.37 | 386.66 | 437.9 | 760.08 | 826.86 | 1364.06 | 1112.93 | 324.31 | 794.81 | 1051.64 | 13838.78 | ATP-binding cassette sub-family G member 4 (EC 7.6.2.-) | ABCG4 WHITE2 | cellular response to high density lipoprotein particle stimulus [GO:0071403]; cellular response to leukemia inhibitory factor [GO:1990830]; cholesterol efflux [GO:0033344]; cholesterol homeostasis [GO:0042632]; positive regulation of cholesterol biosynthetic process [GO:0045542]; positive regulation of cholesterol efflux [GO:0010875]; regulation of DNA-templated transcription [GO:0006355]; transmembrane transport [GO:0055085] |
| 881.85 | 632.73 | 3950.44 | 399.25 | 999.38 | 531.59 | 6.51 | 196.09 | 1011.53 | 371.41 | 262.54 | 896.14 | 2266.21 | 313.61 | 303.81 | 138.41 | 314.06 | 361.38 | 13836.94 | Mitochondrial glutathione transporter SLC25A40 (Solute carrier family 25 member 40) | slc25a40 zgc:92520 | import into the mitochondrion [GO:0170036] |
| 30.03 | 4092.85 | 190.74 | 29.09 | 3064.11 | 288.34 | 4.08 | 1589.7 | 13.7 | 20.1 | 495.66 | 104.47 | 23.09 | 2295.45 | 92.55 | 60.82 | 1359.26 | 80.16 | 13834.2 | DCC-interacting protein 13-alpha (Dip13-alpha) (Adapter protein containing PH domain, PTB domain and leucine zipper motif 1) | Appl1 Dip13a Kiaa1428 | adiponectin-activated signaling pathway [GO:0033211]; cellular response to hepatocyte growth factor stimulus [GO:0035729]; insulin receptor signaling pathway [GO:0008286]; maintenance of synapse structure [GO:0099558]; negative regulation of Fc-gamma receptor signaling pathway involved in phagocytosis [GO:1905450]; positive regulation of cytokine production involved in inflammatory response [GO:1900017]; positive regulation of D-glucose import [GO:0046326]; positive regulation of macropinocytosis [GO:1905303]; positive regulation of melanin biosynthetic process [GO:0048023]; protein import into nucleus [GO:0006606]; regulation of fibroblast migration [GO:0010762]; regulation of G1/S transition of mitotic cell cycle [GO:2000045]; regulation of innate immune response [GO:0045088]; regulation of protein localization to plasma membrane [GO:1903076]; regulation of toll-like receptor 4 |

| | | | | | | | | | | | | | | | | | | | | | |
|---|---|---|---|---|---|---|---|---|---|---|---|---|---|---|---|---|---|---|---|---|---|
| 497.11 | 974.72 | 979.99 | 215.67 | 1023.89 | 933.91 | 234.19 | 374.39 | 1120.43 | 155.48 | 326.67 | 1040.28 | 417.64 | 1865.12 | 1428.29 | 342.61 | 621.45 | 1270.66 | 13822.5 | Phosphomevalonate kinase (PMKase) (EC 2.7.4.2) | PMVK | cholesterol biosynthetic process [GO:0006695]; isopentenyl diphosphate biosynthetic process, mevalonate pathway [GO:0019287]; response to cholesterol [GO:0070723] |
| 511.07 | 428.16 | 1618.87 | 303.3 | 812.78 | 998.61 | 43.96 | 206.73 | 1021.4 | 187.41 | 484.27 | 2872.91 | 530.47 | 602.02 | 888.69 | 260.7 | 955.5 | 1077.78 | 13804.63 | Low-density lipoprotein receptor (LDL receptor) | Ldlr | amyloid-beta clearance [GO:0097242]; amyloid-beta clearance by cellular catabolic process [GO:0150094]; artery morphogenesis [GO:0048844]; cellular response to fatty acid [GO:0071398]; cellular response to low-density lipoprotein particle stimulus [GO:0071404]; cholesterol homeostasis [GO:0042632]; cholesterol import [GO:0070508]; cholesterol metabolic process [GO:0008203]; cholesterol transport [GO:0030301]; endocytosis [GO:0006897]; establishment of localization in cell [GO:0051649]; high-density lipoprotein particle clearance [GO:0034384]; intestinal cholesterol absorption [GO:0030299]; lipid metabolic process [GO:0006629]; lipoprotein catabolic process [GO:0042159]; lipoprotein metabolic process [GO:0042157]; long-term memory [GO:0007616]; low-density lipoprotein particle clearance [GO:0034383]; negative regulation of amyloid fibril formation [GO:1905907]; negative regulation of astrocyte [illegible] |

| | | | | | | | | | | | | | | | | | | | | | |
|---|---|---|---|---|---|---|---|---|---|---|---|---|---|---|---|---|---|---|---|---|---|
| 679.1 | 589.93 | 791.14 | 271.3 | 738.52 | 971.65 | 1662.3 | 245.2 | 939.17 | 260.13 | 220.31 | 797.47 | 892.08 | 604.19 | 1073.55 | 1263.39 | 429.46 | 1354.04 | 13782.93 | Transcription factor AP-2 gamma (AP2-gamma) (Activating enhancer-binding protein 2 gamma) (Transcription factor ERF-1) | TFAP2C | cell-cell signaling [GO:0007267]; cerebral cortex development [GO:0021987]; dichotomous subdivision of terminal units involved in mammary gland duct morphogenesis [GO:0060598]; epithelial cell proliferation involved in mammary gland duct elongation [GO:0060750]; forebrain neuron fate commitment [GO:0021877]; germ-line stem cell population maintenance [GO:0030718]; hair follicle development [GO:0001942]; inner cell mass cell differentiation [GO:0001826]; keratinocyte development [GO:0003334]; male gonad development [GO:0008584]; morula formation [GO:0140001]; negative regulation of gene expression, epigenetic [GO:0045814]; positive regulation of transcription by RNA polymerase II [GO:0045944]; regulation of cell population proliferation [GO:0042127]; regulation of epidermis development [GO:0045682]; regulation of transcription by RNA polymerase II [GO:0006357]; sebaceous gland |
| 876.25 | 602.77 | 806.99 | 320.49 | 568.33 | 1259.67 | 1805.66 | 347.76 | 288.56 | 331.29 | 316.33 | 495.69 | 1412.97 | 791.79 | 673.64 | 1878.14 | 535.26 | 465.14 | 13776.73 | Glutamate receptor 3 (GluR-3) (AMPA-selective glutamate receptor 3) (GluR-C) (GluR-K3) (Glutamate receptor ionotropic, AMPA 3) | Gria3 GluA3 Glur3 | long-term synaptic potentiation [GO:0060291]; modulation of chemical synaptic transmission [GO:0050804]; protein heterotetramerization [GO:0051290]; protein homotetramerization [GO:0051289]; regulation of receptor recycling [GO:0001919]; response to fungicide [GO:0060992]; response to lithium ion [GO:0010226]; synaptic transmission, glutamatergic [GO:0035249] |

| | | | | | | | | | | | | | | | | | | | | | |
|---|---|---|---|---|---|---|---|---|---|---|---|---|---|---|---|---|---|---|---|---|---|
| 737.93 | 550.26 | 675.28 | 254.58 | 624.19 | 987.86 | 2187.33 | 248.59 | 809.4 | 198.04 | 211.13 | 563.78 | 1063.71 | 847.57 | 988.37 | 1378.08 | 446.51 | 1000.63 | 13773.24 | Tyrosine-protein kinase Blk (EC 2.7.10.2) (B lymphocyte kinase) (p55-Blk) | BLK | B cell receptor signaling pathway [GO:0050853]; cell differentiation [GO:0030154]; cell surface receptor protein tyrosine kinase signaling pathway [GO:0007169]; intracellular signal transduction [GO:0035556]; peptidyl-tyrosine phosphorylation [GO:0018108]; positive regulation of insulin secretion [GO:0032024] |
| 192.77 | 2286.72 | 100.94 | 98.59 | 1733.51 | 14.51 | 0 | 791.62 | 12.11 | 62.27 | 868.87 | 47.43 | 498.27 | 4193.86 | 148.33 | 392.71 | 2056.41 | 267.28 | 13766.2 | NACHT, LRR and PYD domains-containing protein 12 (Monarch-1) (PYRIN-containing APAF1-like protein 7) (Regulated by nitric oxide) | NLRP12 NALP12 PYPAF7 RNO | cellular response to cytokine stimulus [GO:0071345]; dendritic cell migration [GO:0036336]; ERK1 and ERK2 cascade [GO:0070371]; negative regulation of canonical NF-kappaB signal transduction [GO:0043124]; negative regulation of cytokine production [GO:0001818]; negative regulation of ERK1 and ERK2 cascade [GO:0070373]; negative regulation of inflammatory response [GO:0050728]; negative regulation of interleukin-1 production [GO:0032692]; negative regulation of interleukin-6 production [GO:0032715]; negative regulation of non-canonical NF-kappaB signal transduction [GO:1901223]; negative regulation of signal transduction [GO:0009968]; negative regulation of Toll signaling pathway [GO:0045751]; positive regulation of inflammatory response [GO:0050729]; positive regulation of interleukin-1 beta production [GO:0032731]; positive regulation of MHC class I biosynthetic process [GO:0045345]; positive regulation of non- |
| 2233.51 | 370.67 | 661.11 | 872.28 | 831.12 | 830.74 | 31.55 | 235.07 | 434.25 | 1006.49 | 209.9 | 2641.46 | 1040.53 | 206.98 | 321.98 | 211.17 | 590.1 | 1036 | 13764.91 | Guanine nucleotide-binding protein G(s) subunit alpha (Adenylate cyclase-stimulating G alpha protein) | gnas | adenylate cyclase-activating dopamine receptor signaling pathway [GO:0007191]; sensory perception of chemical stimulus [GO:0007606] |

| | | | | | | | | | | | | | | | | | | | | | |
|---|---|---|---|---|---|---|---|---|---|---|---|---|---|---|---|---|---|---|---|---|---|
| 3835.83 | 26.49 | 15.63 | 749 | 66.24 | 282.82 | 39.14 | 4.42 | 0 | 2965.2 | 43.1 | 119.17 | 5198.09 | 15.46 | 46.49 | 325.86 | 25.32 | 0.73 | 13758.99 | COMM domain-containing protein 5 (Hypertension-related calcium-regulated gene protein) (HCaRG) | COMMD5 HT002 | |
| 782.84 | 984.42 | 1362.22 | 290.44 | 943.9 | 1225.78 | 501.31 | 321.96 | 707.39 | 234.87 | 289.97 | 1051.91 | 977.52 | 1243.08 | 800.24 | 464.24 | 812.78 | 763.91 | 13758.78 | Fibrinogen alpha chain [Cleaved into: Fibrinopeptide A;Fibrinogen alpha chain] | FGA | blood coagulation, common pathway [GO:0072377]; fibrinolysis [GO:0042730]; platelet aggregation [GO:0070527]; positive regulation of heterotypic cell-cell adhesion [GO:0034116]; protein polymerization [GO:0051258] |
| 517.72 | 1099.29 | 593.8 | 278.76 | 701.57 | 225.45 | 1.16 | 528.29 | 177.53 | 308.93 | 287.8 | 296.16 | 198.21 | 6980.26 | 240.38 | 48.61 | 954.76 | 319.29 | 13757.97 | Zinc finger protein Pegasus (Ikaros family zinc finger protein 5) | ikzf5 zgc:92405 | determination of intestine left/right asymmetry [GO:0071908]; determination of left/right asymmetry in lateral mesoderm [GO:0003140]; determination of left/right symmetry [GO:0007368]; determination of pancreatic left/right asymmetry [GO:0035469]; regulation of transcription by RNA polymerase II [GO:0006357] |
| 127.69 | 1372.53 | 201.75 | 395.68 | 1202.88 | 125.56 | 496.69 | 603.07 | 164.11 | 94.74 | 381.36 | 241.76 | 195.79 | 4536.9 | 242 | 758.03 | 2248.99 | 358.89 | 13748.42 | Myeloid-associated differentiation marker homolog | myadm | |

| | | | | | | | | | | | | | | | | | | | | | |
|---|---|---|---|---|---|---|---|---|---|---|---|---|---|---|---|---|---|---|---|---|---|
| 136.02 | 31.29 | 83.76 | 284.26 | 28.67 | 132.05 | 8444.35 | 5.59 | 21.04 | 96.12 | 137.9 | 57.54 | 608.31 | 36.1 | 48.54 | 3366.89 | 180.09 | 37.21 | 13735.73 | 5'-AMP-activated protein kinase subunit gamma-2 (AMPK gamma2) (AMPK subunit gamma-2) (H91620p) | PRKAG2 | ATP biosynthetic process [GO:0006754]; cellular response to glucose starvation [GO:0042149]; cellular response to nutrient levels [GO:0031669]; fatty acid biosynthetic process [GO:0006633]; glycogen metabolic process [GO:0005977]; intracellular signal transduction [GO:0035556]; positive regulation of gluconeogenesis [GO:0045722]; regulation of carbon utilization [GO:0043609]; regulation of cell cycle [GO:0051726]; regulation of D-glucose import [GO:0046324]; regulation of fatty acid metabolic process [GO:0019217]; regulation of fatty acid oxidation [GO:0046320]; regulation of glycolytic process [GO:0006110]; sterol biosynthetic process [GO:0016126] |
| 232.49 | 88.5 | 2870.56 | 73.12 | 41.02 | 1364.55 | 12.1 | 9.34 | 2889.79 | 72.13 | 75.84 | 1319.96 | 312.38 | 341.76 | 1404.8 | 132.74 | 517.49 | 1921.86 | 13680.43 | Occludin | Ocln | bicellular tight junction assembly [GO:0070830]; cell-cell junction organization [GO:0045216]; cellular response to tumor necrosis factor [GO:0071356]; negative regulation of gene expression [GO:0010629]; positive regulation of blood-brain barrier permeability [GO:1905605]; positive regulation of D-glucose import [GO:0046326]; positive regulation of gene expression [GO:0010628]; positive regulation of lamellipodium assembly [GO:0010592]; positive regulation of microtubule polymerization [GO:0031116]; positive regulation of wound healing [GO:0090303]; regulation of D-glucose transmembrane transport [GO:0010827]; response to ethanol [GO:0045471]; response to Gram-positive bacterium [GO:0140459]; response to interleukin-18 [GO:0070673]; tight junction organization [GO:0120193] |
| 998.48 | 921.05 | 932.87 | 267.79 | 827.77 | 950.3 | 670.23 | 555.47 | 891.38 | 320.88 | 331.75 | 713.01 | 800.86 | 1245.55 | 910.64 | 518.49 | 728.49 | 1068.24 | 13653.25 | Protein PRRC2A (HLA-B-associated transcript 2) (Large proline-rich protein BAT2) (Proline-rich and coiled-coil-containing protein 2A) (Protein G2) | PRRC2A BAT2 G2 | cell differentiation [GO:0030154] |

| | | | | | | | | | | | | | | | | | | | | | |
|---|---|---|---|---|---|---|---|---|---|---|---|---|---|---|---|---|---|---|---|---|---|
| 1132 | 619.92 | 1507.86 | 472.83 | 1100.16 | 1719.02 | 4.97 | 425.93 | 166.35 | 513.07 | 441.08 | 1212.74 | 1340.77 | 1287.14 | 613.43 | 210.23 | 532.01 | 310.4 | 13609.91 | G protein-coupled receptor 137Ba | gpr137ba | autophagy [GO:0006914]; bone remodeling [GO:0046849]; negative regulation of bone resorption [GO:0045779]; negative regulation of osteoclast differentiation [GO:0045671]; positive regulation of protein localization to lysosome [GO:0150032]; positive regulation of TORC1 signaling [GO:1904263]; regulation of autophagy [GO:0010506]; regulation of GTPase activity [GO:0043087] |
| 1059.86 | 599.22 | 551.52 | 470.85 | 750.83 | 909.77 | 1361.94 | 615.54 | 385.72 | 446.76 | 452.51 | 469.01 | 985.68 | 1189.2 | 560.76 | 1186.5 | 1109.64 | 500.4 | 13605.71 | Host cell factor 2 (HCF-2) (C2 factor) | Hcfc2 | antiviral innate immune response [GO:0140374]; immune response involved in response to exogenous dsRNA [GO:1902615]; negative regulation of transcription by RNA polymerase II [GO:0000122]; regulation of toll-like receptor 3 signaling pathway [GO:0034139]; regulation of transcription by RNA polymerase II [GO:0006357] |

| | | | | | | | | | | | | | | | | | | | | | |
|---|---|---|---|---|---|---|---|---|---|---|---|---|---|---|---|---|---|---|---|---|---|
| 884.89 | | 619.48 | 808.63 | 206.65 | 1438.65 | 1768.29 | 6.58 | 344.47 | 355.2 | 371.62 | 638.64 | 884.42 | 638.56 | 1019.88 | 202.94 | 461.17 | 2041.98 | 912 | 13604.05 | Alpha-internexin (Alpha-Inx) (66 kDa neurofilament protein) (NF-66) (Neurofilament-66) (Neurofilament 5) | INA NEF5 | cell differentiation [GO:0030154]; cellular response to leukemia inhibitory factor [GO:1990830]; intermediate filament organization [GO:0045109]; neurofilament cytoskeleton organization [GO:0060052]; postsynaptic modulation of chemical synaptic transmission [GO:0099170]; substantia nigra development [GO:0021762] |
| 985.87 | | 974.38 | 865.16 | 270.48 | 722.55 | 982.92 | 824.76 | 376.08 | 742.04 | 244.1 | 272.62 | 830.18 | 896.88 | 1319.15 | 858.44 | 884.13 | 662.01 | 884.57 | 13596.32 | Insulin-like growth factor 1 (Insulin-like growth factor I) (IGF-I) (Somatomedin) | igf1 igf-1 | cell population proliferation [GO:0008283]; insulin-like growth factor receptor signaling pathway [GO:0048009]; negative regulation of apoptotic process [GO:0043066]; negative regulation of release of cytochrome c from mitochondria [GO:0090201]; positive regulation of cell population proliferation [GO:0008284]; positive regulation of phosphatidylinositol 3-kinase/protein kinase B signal transduction [GO:0051897] |

| | | | | | | | | | | | | | | | | | | | | | |
|---|---|---|---|---|---|---|---|---|---|---|---|---|---|---|---|---|---|---|---|---|---|
| 234.22 | 570.68 | 2655.79 | 105.79 | 967.41 | 1246.64 | 925.94 | 202.94 | 666.99 | 70.28 | 494.96 | 928.8 | 222.64 | 1203.47 | 1110.89 | 727.51 | 747.39 | 513.9 | 13596.24 | Gamma-aminobutyric acid receptor subunit rho-3 (GABA(A) receptor subunit rho-3) (GABAAR subunit rho-3) (GABA(C) receptor) | GABRR3 | chemical synaptic transmission [GO:0007268]; chloride transmembrane transport [GO:1902476]; gamma-aminobutyric acid signaling pathway [GO:0007214] |
| 746.96 | 763.57 | 1108.96 | 192.46 | 742.91 | 1127.31 | 907.51 | 356.71 | 669.07 | 349.7 | 333.23 | 939.59 | 824.01 | 1022.58 | 1178.87 | 596.08 | 577.58 | 1149.07 | 13586.17 | Protein sidekick-2 | SDK2 KIAA1514 | camera-type eye photoreceptor cell differentiation [GO:0060219]; homophilic cell adhesion via plasma membrane adhesion molecules [GO:0007156]; retina layer formation [GO:0010842]; synapse assembly [GO:0007416] |

| | | | | | | | | | | | | | | | | | | | | | |
|---|---|---|---|---|---|---|---|---|---|---|---|---|---|---|---|---|---|---|---|---|---|
| 991.45 | 746.72 | 850.09 | 548.08 | 616.69 | 2916.7 | 543.8 | 264.82 | 193.64 | 573.97 | 242.66 | 1772.54 | 1255.35 | 420.23 | 565.22 | 434.4 | 394.03 | 230.66 | 13561.05 | SAGA-associated factor 29 (rSGF29) (Coiled-coil domain-containing protein 101) (SAGA complex-associated factor 29) | Sgf29 Ccdc101 | negative regulation of transcription by RNA polymerase II [GO:0000122]; regulation of cell cycle [GO:0051726]; regulation of cell division [GO:0051302]; regulation of DNA repair [GO:0006282]; regulation of DNA-templated transcription [GO:0006355]; regulation of embryonic development [GO:0045995]; regulation of transcription by RNA polymerase II [GO:0006357]; response to endoplasmic reticulum stress [GO:0034976]; transcription initiation-coupled chromatin remodeling [GO:0045815] |
| 449.87 | 612.57 | 816.55 | 206.85 | 632.77 | 1436.32 | 215.82 | 682.33 | 97.95 | 417 | 477.17 | 1171.57 | 1398.97 | 1730.1 | 1018.87 | 622.83 | 899.8 | 645.14 | 13532.48 | Zinc finger protein 518A | ZNF518A KIAA0335 ZNF518 | chromatin organization [GO:0006325]; regulation of transcription by RNA polymerase II [GO:0006357] |
| 42.31 | 4920.04 | 146.06 | 163.22 | 2621.49 | 48.31 | 12.48 | 1428.32 | 5.13 | 47.32 | 533.6 | 64.28 | 18.74 | 2305.3 | 33.02 | 29.14 | 1091.37 | 19.51 | 13529.64 | Proto-oncogene tyrosine-protein kinase Src (EC 2.7.10.2) (Proto-oncogene c-Src) (pp60c-src) (p60-Src) | src | cell adhesion [GO:0007155]; cell differentiation [GO:0030154]; epidermal growth factor receptor signaling pathway [GO:0007173]; negative regulation of extrinsic apoptotic signaling pathway [GO:2001237]; negative regulation of intrinsic apoptotic signaling pathway [GO:2001243]; progesterone receptor signaling pathway [GO:0050847]; regulation of vascular permeability [GO:0043114]; response to yeast [GO:0001878] |

| | | | | | | | | | | | | | | | | | | | | | |
|---|---|---|---|---|---|---|---|---|---|---|---|---|---|---|---|---|---|---|---|---|---|
| 5.59 | 13.07 | 7118.66 | 16.91 | 12.62 | 60.72 | 47.74 | 7.68 | 1777.37 | 10.26 | 11.54 | 1011.76 | 38.46 | 22.4 | 1384.62 | 90.04 | 24.81 | 1862.87 | 13517.12 | DnaJ homolog subfamily B member 6 | dnajb6 | chaperone-mediated protein folding [GO:0061077] |
| 21.98 | 297.5 | 523.12 | 31.28 | 289.27 | 457.18 | 26.28 | 94.38 | 2326.08 | 21.78 | 126.23 | 1042.25 | 7.13 | 769.5 | 3366.21 | 185.32 | 364.29 | 3521.48 | 13471.26 | Protein virilizer homolog | Virma Kiaa1429 | mRNA processing [GO:0006397]; RNA splicing [GO:0008380] |
| 604.78 | 615.89 | 969.8 | 254.18 | 501.31 | 552.36 | 2027.62 | 250.22 | 836.64 | 195.28 | 262.3 | 1179.41 | 761.71 | 855.05 | 737.05 | 1445.02 | 522.38 | 850.82 | 13421.82 | Ankyrin repeat domain-containing protein 40 | ANKRD40 | |
| 5.85 | 47.4 | 259.97 | 6.41 | 12.89 | 16.44 | 285.28 | 2.38 | 3840.05 | 5.63 | 9.53 | 1339.23 | 3.42 | 55.22 | 3815.05 | 53.52 | 33.16 | 3620.04 | 13411.47 | Nucleus accumbens-associated protein 1 (NAC-1) (BTB/POZ domain-containing protein 14B) | Nacc1 Btbd14b Nac1 | negative regulation of DNA-templated transcription [GO:0045892]; positive regulation of cell population proliferation [GO:0008284] |
| 470.58 | 1103.63 | 1287.21 | 395.04 | 948.86 | 1012.83 | 41.72 | 738.66 | 479.77 | 386.38 | 637.3 | 2151.13 | 526.25 | 1066.95 | 871.49 | 102 | 557.75 | 626.07 | 13403.62 | Protection of telomeres protein 1 (cPot1) (POT1-like telomere end-binding protein) | POT1 | regulation of telomere maintenance via telomerase [GO:0032210]; telomere capping [GO:0016233]; telomere maintenance via telomerase [GO:0007004] |

| | | | | | | | | | | | | | | | | | | | | | |
|---|---|---|---|---|---|---|---|---|---|---|---|---|---|---|---|---|---|---|---|---|---|
| 823.76 | 887.25 | 510.12 | 374.7 | 756.95 | 989.97 | 1760.06 | 377.16 | 552.43 | 306.05 | 375.17 | 539.98 | 1019.98 | 915.37 | 554.37 | 1311.63 | 706.88 | 641.36 | 13403.19 | Hairy/enhancer-of-split related with YRPW motif protein 2 (Protein gridlock) | hey2 grl zgc:136746 | artery development [GO:0060840]; artery morphogenesis [GO:0048844]; blood vessel development [GO:0001568]; blood vessel morphogenesis [GO:0048514]; cardiac muscle cell proliferation [GO:0060038]; circulatory system development [GO:0072359]; dorsal aorta morphogenesis [GO:0035912]; heart development [GO:0007507]; negative regulation of DNA-templated transcription [GO:0045892]; Notch signaling pathway [GO:0007219]; Notch signaling pathway involved in arterial endothelial cell fate commitment [GO:0060853]; regulation of neurogenesis [GO:0050767]; vasculogenesis [GO:0001570] |
| 1398.97 | 759.47 | 765.18 | 298.95 | 683.35 | 1135.1 | 371.31 | 279.46 | 722.22 | 414.59 | 490.73 | 810.23 | 847.11 | 1479.81 | 1003.6 | 380.46 | 636.26 | 895.04 | 13371.84 | Leucyl-cystinyl aminopeptidase (Cystinyl aminopeptidase) (EC 3.4.11.3) (Oxytocinase) (OTase) | Lnpep | negative regulation of cold-induced thermogenesis [GO:0120163]; protein catabolic process [GO:0030163]; proteolysis [GO:0006508] |
| 1099.78 | 737.94 | 1237.39 | 390.36 | 673.77 | 1171.34 | 366.19 | 237.63 | 921.76 | 330.96 | 293.85 | 743.7 | 1117.35 | 892.47 | 1149.3 | 311.54 | 555.25 | 1134.71 | 13365.29 | Transmembrane protein 266 (hTMEM266) (HV1-related protein 1) (HsHVRP1) | TMEM266 C15orf27 HVRP1 | |

| | | | | | | | | | | | | | | | | | | | | | |
|---|---|---|---|---|---|---|---|---|---|---|---|---|---|---|---|---|---|---|---|---|---|
| 699.37 | 542.95 | 878.44 | 246.07 | 707.72 | 1319.46 | 901.49 | 306.6 | 1041.62 | 303.07 | 290.85 | 818.73 | 746.05 | 817.63 | 1179.92 | 674.23 | 535.62 | 1322.38 | 13332.2 | Anaphase-promoting complex subunit 1 (APC1) (Cyclosome subunit 1) (Mitotic checkpoint regulator) (Testis-specific gene 24 protein) | ANAPC1 TSG24 | anaphase-promoting complex-dependent catabolic process [GO:0031145]; cell division [GO:0051301]; metaphase/anaphase transition of mitotic cell cycle [GO:0007091]; protein branched polyubiquitination [GO:0141198]; protein K11-linked ubiquitination [GO:0070979]; protein K48-linked ubiquitination [GO:0070936]; regulation of meiotic cell cycle [GO:0051445]; regulation of mitotic cell cycle [GO:0007346] |
| 737.71 | 562.1 | 1566.08 | 301.94 | 470.86 | 821.13 | 1626.95 | 290.14 | 577.14 | 245.83 | 201.12 | 546.63 | 425.21 | 1039.82 | 674.25 | 1915.16 | 574.99 | 739.17 | 13316.23 | Vacuolar protein sorting-associated protein 33B | vps33b | intracellular protein transport [GO:0006886]; intrahepatic bile duct development [GO:0035622]; vesicle-mediated transport [GO:0016192] |
| 1069.59 | 736.67 | 836.51 | 423.8 | 934.17 | 787.03 | 568.05 | 227.22 | 125.55 | 260.01 | 240.33 | 616.67 | 1836.4 | 1908.29 | 511 | 1010.77 | 895.72 | 320.95 | 13308.73 | Elongation factor 1-delta (EF-1-delta) (Antigen NY-CO-4) | EEF1D EF1D | cellular response to heat [GO:0034605]; cellular response to ionizing radiation [GO:0071479]; cytoplasmic translational elongation [GO:0002182]; positive regulation of transcription by RNA polymerase II [GO:0045944]; translational elongation [GO:0006414] |

| | | | | | | | | | | | | | | | | | | | | | |
|---|---|---|---|---|---|---|---|---|---|---|---|---|---|---|---|---|---|---|---|---|---|
| 5.89 | 15.44 | 323.12 | 1.21 | 5.7 | 4.84 | 3.03 | 10.45 | 4399.66 | 0 | 4.18 | 1241.47 | 3.27 | 10.94 | 3274.48 | 3.88 | 22.53 | 3978 | 13308.09 | Transcriptional repressor NF-X1 (EC 2.3.2.-) (Nuclear transcription factor, X box-binding protein 1) | NFX1 NFX2 | inflammatory response [GO:0006954]; negative regulation of MHC class II biosynthetic process [GO:0045347]; negative regulation of transcription by RNA polymerase II [GO:0000122]; protein autoubiquitination [GO:0051865]; protein ubiquitination [GO:0016567]; transcription by RNA polymerase II [GO:0006366] |
| 64.63 | 1.15 | 11.22 | 241.14 | 7.12 | 17.32 | 7472.62 | 1.25 | 38.89 | 79.67 | 183.36 | 5.5 | 164.49 | 6.84 | 41.79 | 4791.46 | 127.12 | 45.42 | 13300.99 | Cerebral cavernous malformations protein 2 homolog (Malcavernin) (Valentine) | ccm2 vtn | anterior/posterior axis specification [GO:0009948]; cardiac jelly development [GO:1905072]; cell-cell junction assembly [GO:0007043]; heart contraction [GO:0060047]; heart development [GO:0007507]; vasculogenesis [GO:0001570] |

| | | | | | | | | | | | | | | | | | | | | | | |
|---|---|---|---|---|---|---|---|---|---|---|---|---|---|---|---|---|---|---|---|---|---|---|
| 313.14 | | 2029.74 | 535.32 | 150.94 | 1459.82 | 663.74 | 106.45 | 769.08 | 760.26 | 121.81 | 505.66 | 483.87 | 478.25 | 1840.74 | 952.39 | 164.86 | 867.22 | 1094.31 | 13297.6 | SWI/SNF-related matrix-associated actin-dependent regulator of chromatin subfamily A member 2 (SAMRCA2) (EC 3.6.4.-) (BRG1-associated factor 190B) (BAF190B) (Probable global transcription activator SNF2L2) (Protein brahma homolog) (SNF2-alpha) | Smarca2 Baf190b Brm Snf2a Snf2l2 | aortic smooth muscle cell differentiation [GO:0035887]; cell population proliferation [GO:0008283]; chromatin organization [GO:0006325]; chromatin remodeling [GO:0006338]; fibroblast proliferation [GO:0048144]; hematopoietic stem cell homeostasis [GO:0061484]; negative regulation of cell differentiation [GO:0045596]; negative regulation of cell population proliferation [GO:0008285]; negative regulation of fibroblast proliferation [GO:0048147]; nervous system development [GO:0007399]; nucleosome assembly [GO:0006334]; positive regulation of cell differentiation [GO:0045597]; positive regulation of cell population proliferation [GO:0008284]; positive regulation of double-strand break repair [GO:2000781]; positive regulation of myoblast differentiation [GO:0045663]; positive regulation of stem cell population maintenance [GO:1902459]; positive regulation of T cell differentiation [GO:0045582]; regulation of G0 to G1 transition |
| 635.08 | | 497.91 | 753.34 | 348.46 | 360.55 | 981.34 | 2493.7 | 249.35 | 509.17 | 305.58 | 177.86 | 782.74 | 700.33 | 470.52 | 731.63 | 1900.31 | 362.27 | 1024.32 | 13284.46 | Methionine-R-sulfoxide reductase B3, mitochondrial (MsrB3) (EC 1.8.4.12) (EC 1.8.4.14) | Msrb3 | protein repair [GO:0030091]; response to oxidative stress [GO:0006979] |
| 582.32 | | 484.86 | 884.36 | 153.69 | 442.6 | 763.73 | 696.04 | 235.91 | 1800.14 | 218.6 | 209.21 | 803.87 | 738.3 | 679.95 | 1616.54 | 463.86 | 394.55 | 2109.11 | 13277.64 | Endoribonuclease Dicer (EC 3.1.26.3) | dicer1 | apoptotic DNA fragmentation [GO:0006309]; cranial skeletal system development [GO:1904888]; global gene silencing by mRNA cleavage [GO:0098795]; heterochromatin formation [GO:0031507]; melanocyte differentiation [GO:0030318]; miRNA processing [GO:0035196]; miRNA-mediated gene silencing by mRNA destabilization [GO:0035279]; miRNA-mediated post-transcriptional gene silencing [GO:0035195]; neural crest cell development [GO:0014032]; pre-miRNA processing [GO:0031054]; regulation of neurogenesis [GO:0050767]; siRNA processing [GO:0030422]; ventricular system development [GO:0021591] |
| 492.23 | | 1050.08 | 466.84 | 208.71 | 1474.54 | 990.93 | 259.18 | 1520.67 | 105.91 | 514.24 | 757.67 | 324.81 | 915.19 | 2105.12 | 220.78 | 344.69 | 1392.9 | 115.77 | 13260.26 | PHD finger protein 11 | PHF11 | |

| | | | | | | | | | | | | | | | | | | | | | |
|---|---|---|---|---|---|---|---|---|---|---|---|---|---|---|---|---|---|---|---|---|---|
| 1113.49 | 692.31 | 953.22 | 251.57 | 569.91 | 824.89 | 184.21 | 307.54 | 1617.61 | 325.27 | 245.62 | 733.8 | 659.43 | 913.59 | 1407.45 | 324.64 | 511.74 | 1614.92 | 13251.21 | Protein mono-ADP-ribosyltransferase PARP9 (EC 2.4.2.-) (ADP-ribosyltransferase diphtheria toxin-like 9) (ARTD9) (B aggressive lymphoma protein homolog) (Poly [ADP-ribose] polymerase 9) (PARP-9) | Parp9 Bal | defense response to virus [GO:0051607]; DNA damage checkpoint signaling [GO:0000077]; double-strand break repair [GO:0006302]; innate immune response [GO:0045087]; negative regulation of gene expression [GO:0010629]; negative regulation of transcription by RNA polymerase II [GO:0000122]; positive regulation of defense response to virus by host [GO:0002230]; positive regulation of DNA-templated transcription [GO:0045893]; positive regulation of protein localization to nucleus [GO:1900182]; positive regulation of type II interferon-mediated signaling pathway [GO:0060335]; post-transcriptional regulation of gene expression [GO:0010608]; regulation of response to type II interferon [GO:0060330] |
| 1104.91 | 781.55 | 1136.83 | 299.97 | 647.52 | 938.64 | 886.11 | 410.39 | 799.03 | 333.13 | 319.41 | 609.08 | 911.97 | 911.01 | 841.27 | 762.37 | 622.25 | 919.33 | 13234.77 | La-related protein 4 (La ribonucleoprotein domain family member 4) | LARP4 PP13296 | cytoskeleton organization [GO:0007010]; positive regulation of translation [GO:0045727]; post-transcriptional regulation of gene expression [GO:0010608]; regulation of cell morphogenesis [GO:0022604]; translation [GO:0006412] |
| 462.28 | 905.43 | 1565.24 | 151.34 | 1230.59 | 1511.01 | 69.17 | 923.57 | 211.77 | 225.44 | 506.77 | 769.07 | 748.41 | 1999.09 | 562.56 | 142.02 | 476.81 | 757.91 | 13218.48 | Bromodomain adjacent to zinc finger domain protein 2B (hWALp4) | BAZ2B KIAA1476 | chromatin remodeling [GO:0006338]; regulation of transcription by RNA polymerase II [GO:0006357] |

| | | | | | | | | | | | | | | | | | | | | | | |
|---|---|---|---|---|---|---|---|---|---|---|---|---|---|---|---|---|---|---|---|---|---|---|
| 508.15 | | 1080.31 | 958.84 | 511.7 | 1399.77 | 922.88 | 853.62 | 529.77 | 376.45 | 301.95 | 635.63 | 720.61 | 548.56 | 1343.07 | 358.16 | 746.67 | 895.14 | 520.62 | 13211.9 | Neuroepithelial cell-transforming gene 1 protein (Rho guanine nucleotide exchange factor 8) | Net1 Arhgef8 | cellular response to hydrogen peroxide [GO:0070301]; cellular response to ionizing radiation [GO:0071479]; intracellular signal transduction [GO:0035556]; myoblast migration [GO:0051451]; positive regulation of apoptotic process [GO:0043065]; positive regulation of GTPase activity [GO:0043547]; positive regulation of substrate adhesion-dependent cell spreading [GO:1900026] |
| 400.54 | | 503.15 | 857.12 | 299.32 | 642.42 | 1090.96 | 2004.15 | 247.05 | 563.85 | 212.98 | 277.9 | 863.91 | 821.14 | 847.85 | 707.5 | 1528.8 | 467.81 | 858.6 | 13195.05 | Thyroid hormone receptor beta (TR-beta) (TRb) (Nuclear receptor subfamily 1 group A member 2) (Thyroid hormone receptor beta-1) (TRbeta1) | thrb nr1a2 trb si:ch211-264a6.2 | camera-type eye development [GO:0043010]; cell differentiation [GO:0030154]; inner ear development [GO:0048839]; negative regulation of transcription by RNA polymerase II [GO:0000122]; positive regulation of transcription by RNA polymerase II [GO:0045944]; regulation of DNA-templated transcription [GO:0006355]; retinal cone cell development [GO:0046549]; retinal cone cell differentiation [GO:0042670]; retinal cone cell fate determination [GO:0042671]; retinoic acid receptor signaling pathway [GO:0048384]; thyroid hormone receptor signaling pathway [GO:0002154] |

| | | | | | | | | | | | | | | | | | | | | | |
|---|---|---|---|---|---|---|---|---|---|---|---|---|---|---|---|---|---|---|---|---|---|
| 0 | 0 | 3396.42 | 0 | 0.49 | 0.61 | 0.42 | 0 | 3127.32 | 0.33 | 0 | 1152.71 | 1.87 | 0 | 2244.51 | 0.44 | 0.46 | 3252.91 | 13178.49 | Sphingosine-1-phosphate phosphatase 2 (SPPase2) (Spp2) (hSPP2) (EC 3.1.3.-) (Sphingosine-1-phosphatase 2) | SGPP2 | phospholipid dephosphorylation [GO:0046839]; regulation of type B pancreatic cell proliferation [GO:0061469]; sphingolipid catabolic process [GO:0030149]; sphingosine metabolic process [GO:0006670] |
| 980.54 | 946.75 | 1349.29 | 118.44 | 1430.81 | 2340.92 | 40.26 | 443.53 | 10.72 | 112.94 | 385.12 | 1461.92 | 550.8 | 1903.37 | 652.38 | 125.66 | 231.71 | 90.21 | 13175.37 | Myocardial zonula adherens protein (GRINL1A upstream protein) (Gup) | MYZAP MYOZAP | intracellular signal transduction [GO:0035556] |
| 153.18 | 205.15 | 755.83 | 40.55 | 130.21 | 340.37 | 173.71 | 50.44 | 3149.63 | 57.08 | 55.59 | 1066.4 | 229.8 | 348.05 | 2562.47 | 189.58 | 161.25 | 3499.93 | 13169.22 | Protein PRRC2B (HLA-B-associated transcript 2-like 1) (Proline-rich coiled-coil protein 2B) | PRRC2B BAT2L BAT2L1 KIAA0515 | cell differentiation [GO:0030154] |
| 1185.68 | 577.82 | 735.98 | 351.1 | 866.23 | 1488.66 | 109.38 | 538.19 | 222.83 | 453.98 | 388.04 | 610.21 | 2491.24 | 1014.39 | 748.15 | 253.35 | 694.39 | 420 | 13149.62 | Troponin I, fast skeletal muscle (Troponin I, fast-twitch isoform) | TNNI2 | cardiac muscle contraction [GO:0060048]; skeletal muscle contraction [GO:0003009] |

| | | | | | | | | | | | | | | | | | | | | | |
|---|---|---|---|---|---|---|---|---|---|---|---|---|---|---|---|---|---|---|---|---|---|
| 478.47 | 923.17 | 679.03 | 381.47 | 966.89 | 740.08 | 15.14 | 351.09 | 476.23 | 165.19 | 185.5 | 1046.91 | 875.51 | 2549.2 | 1132.9 | 347.56 | 711.05 | 1101.1 | 13126.49 | Serine/threonine-protein kinase Nek1 (EC 2.7.11.1) (Never in mitosis A-related kinase 1) (NimA-related protein kinase 1) (Renal carcinoma antigen NY-REN-55) | NEK1 KIAA1901 | cell division [GO:0051301]; cilium assembly [GO:0060271]; protein phosphorylation [GO:0006468] |
| 843.98 | 171.28 | 1539.41 | 197.41 | 251.65 | 1490.55 | 47.44 | 47.4 | 268.92 | 236.55 | 97.94 | 2758.29 | 2038.7 | 741.25 | 845.54 | 605.44 | 465.23 | 460.25 | 13107.23 | Alpha-1,3-mannosyl-glycoprotein 4-beta-N-acetylglucosaminyltransferase C (EC 2.4.1.145) (N-glycosyl-oligosaccharide-glycoprotein N-acetylglucosaminyltransferase IVc) (GnT-IVc) (N-acetylglucosaminyltransferase IVc) (UDP-N-acetylglucosamine: alpha-1,3-D-mannoside beta-1,4-N-acetylglucosaminyltransferase IVc) | mgat4c zgc:101663 | protein N-linked glycosylation [GO:0006487] |

| | | | | | | | | | | | | | | | | | | | | | | |
|---|---|---|---|---|---|---|---|---|---|---|---|---|---|---|---|---|---|---|---|---|---|---|
| 1447.74 | 480.59 | 649.85 | 532.8 | 468.86 | 1012.05 | 32.5 | 298.76 | 44.05 | 456.14 | 267.42 | 440.7 | 2900.98 | 1511.86 | 1079.99 | 489.79 | 829.36 | 156.75 | 13100.19 | Cathepsin S (EC 3.4.22.27) | CTSS | | adaptive immune response [GO:0002250]; antigen processing and presentation [GO:0019882]; antigen processing and presentation of exogenous peptide antigen via MHC class II [GO:0019886]; antigen processing and presentation of peptide antigen [GO:0048002]; basement membrane disassembly [GO:0034769]; cellular response to thyroid hormone stimulus [GO:0097067]; collagen catabolic process [GO:0030574]; extracellular matrix disassembly [GO:0022617]; immune response [GO:0006955]; positive regulation of cation channel activity [GO:2001259]; protein processing [GO:0016485]; proteolysis [GO:0006508]; proteolysis involved in protein catabolic process [GO:0051603]; regulation of antigen processing and presentation [GO:0002577]; response to acidic pH [GO:0010447]; toll-like receptor signaling pathway [GO:0002224] |
| 145.8 | 276.41 | 425.75 | 41.57 | 333.5 | 435.76 | 16.38 | 171.02 | 3804.37 | 39.73 | 157.09 | 1171.96 | 167.09 | 673.84 | 1235.06 | 187.48 | 305.53 | 3509.43 | 13097.77 | Clathrin coat assembly protein AP180 (91 kDa synaptosomal-associated protein) (Clathrin coat-associated protein AP180) (Phosphoprotein F1-20) | SNAP91 KIAA0656 | | clathrin coat assembly [GO:0048268]; clathrin-dependent endocytosis [GO:0072583]; protein transport [GO:0015031]; regulation of clathrin-dependent endocytosis [GO:2000369]; vesicle budding from membrane [GO:0006900] |

| | | | | | | | | | | | | | | | | | | | | | |
|---|---|---|---|---|---|---|---|---|---|---|---|---|---|---|---|---|---|---|---|---|---|
| 845.08 | 708.79 | 738 | 339.58 | 638.14 | 1155.55 | 1839.87 | 396.29 | 615.65 | 308.69 | 260.34 | 582.65 | 749.89 | 887.56 | 551.18 | 1110.77 | 583.55 | 776.33 | 13087.91 | RalA-binding protein 1 (RalBP1) (Cytocentrin) (Dinitrophenyl S-glutathione ATPase) (DNP-SG ATPase) (EC 7.6.2.2, EC 7.6.2.3) (Ral-interacting protein 1) | Ralbp1 | cell division [GO:0051301]; doxorubicin transport [GO:1900753]; endocytosis [GO:0006897]; negative regulation of smooth muscle cell proliferation [GO:0048662]; positive regulation of mitochondrial fission [GO:0090141]; positive regulation of oxidative stress-induced neuron intrinsic apoptotic signaling pathway [GO:1903378]; positive regulation of protein phosphorylation [GO:0001934]; receptor-mediated endocytosis [GO:0006898]; regulation of Cdc42 protein signal transduction [GO:0032489]; regulation of postsynaptic membrane neurotransmitter receptor levels [GO:0099072]; small GTPase-mediated signal transduction [GO:0007264]; transmembrane transport [GO:0055085]; xenobiotic detoxification by transmembrane export across the plasma membrane [GO:1990961] |
| 1026.79 | 670.71 | 959.46 | 458.7 | 779.87 | 965.21 | 1301.26 | 350.28 | 629.42 | 265.3 | 418.85 | 611.11 | 872.3 | 1108.62 | 621.4 | 776.23 | 665.19 | 569.79 | 13050.49 | Integral membrane protein 2C [Cleaved into: CT-BRI3] | Itm2c | negative regulation of amyloid precursor protein biosynthetic process [GO:0042985]; negative regulation of neuron projection development [GO:0010977]; neuron differentiation [GO:0030182]; positive regulation of extrinsic apoptotic signaling pathway [GO:2001238] |
| 1.96 | 2548.83 | 0 | 0 | 3872.9 | 28.08 | 68.61 | 1483.55 | 6.52 | 1.5 | 826.19 | 2.03 | 6.79 | 3274.64 | 15.63 | 19.16 | 868.69 | 9.38 | 13034.46 | Transmembrane protein 87A (Elkin1) | tmem87a TGas047b06.1 | cellular response to mechanical stimulus [GO:0071260]; detection of mechanical stimulus involved in sensory perception of touch [GO:0050976] |

| | | | | | | | | | | | | | | | | | | | | | |
|---|---|---|---|---|---|---|---|---|---|---|---|---|---|---|---|---|---|---|---|---|---|
| 0 | 1926.9 | 0 | 20.81 | 431.29 | 0 | 0 | 996.52 | 0 | 0 | 2368 | 8.23 | 0 | 2966.54 | 0 | 0 | 4315.19 | 0 | 13033.48 | TLC domain-containing protein 3A (Protein CT120) (Protein FAM57A) | TLCD3A FAM57A | lipid homeostasis [GO:0055088] |
| 166.39 | 62.96 | 3410.95 | 23.89 | 96.38 | 143.65 | 32.76 | 18.47 | 3252.56 | 42.32 | 31.45 | 772.91 | 187.27 | 66.72 | 2096.42 | 47.49 | 67.3 | 2509.3 | 13029.19 | Pigment epithelium-derived factor (PEDF) (Serpin F1) | SERPINF1 PEDF | negative regulation of angiogenesis [GO:0016525]; positive regulation of neurogenesis [GO:0050769] |
| 609.57 | 975.48 | 629.93 | 362.51 | 635.97 | 766.5 | 223.9 | 312.08 | 1131.72 | 610.39 | 437.24 | 1240.68 | 1180.25 | 988.2 | 928.31 | 336.79 | 422.18 | 1228.02 | 13019.72 | Spermatogenesis-associated protein 7 homolog | Spata7 | microtubule cytoskeleton organization [GO:0000226]; photoreceptor cell maintenance [GO:0045494]; protein localization to photoreceptor connecting cilium [GO:1903621]; protein localization to photoreceptor outer segment [GO:1903546]; visual perception [GO:0007601] |

| | | | | | | | | | | | | | | | | | | | | | |
|---|---|---|---|---|---|---|---|---|---|---|---|---|---|---|---|---|---|---|---|---|---|
| 1544.74 | 534.84 | 600.17 | 431.28 | 783.53 | 378.36 | 118.73 | 632.34 | 754.84 | 585.5 | 522.07 | 408.08 | 727.7 | 1254.53 | 934.36 | 294.69 | 1187.44 | 1316.81 | 13010.01 | Signal transducer and activator of transcription 1-alpha/beta (Transcription factor ISGF-3 components p91/p84) | STAT1 | blood circulation [GO:0008015]; cell surface receptor signaling pathway via JAK-STAT [GO:0007259]; cell surface receptor signaling pathway via STAT [GO:0097696]; cellular response to insulin stimulus [GO:0032869]; cellular response to interferon-beta [GO:0035458]; cellular response to type II interferon [GO:0071346]; defense response [GO:0006952]; defense response to virus [GO:0051607]; interleukin-27-mediated signaling pathway [GO:0070106]; interleukin-7-mediated signaling pathway [GO:0038111]; interleukin-9-mediated signaling pathway [GO:0038113]; metanephric mesenchymal cell differentiation [GO:0072162]; metanephric mesenchymal cell proliferation involved in metanephros development [GO:0072136]; negative regulation by virus of viral protein levels in host cell [GO:0046725]; negative regulation of angiogenesis [GO:0016525]; negative regulation of canonical NF-kappaB signal |
| 975.88 | 674.91 | 297.04 | 281.87 | 769.44 | 922.69 | 1899.67 | 309.06 | 551.2 | 314.23 | 303.74 | 719.49 | 1116.75 | 703.02 | 599.5 | 1173.31 | 750.71 | 617.53 | 12980.04 | Polycomb group RING finger protein 3 (RING finger protein 3A) | PCGF3 RNF3 RNF3A | random inactivation of X chromosome [GO:0060816]; regulation of transcription by RNA polymerase II [GO:0006357] |
| 570.82 | 849.96 | 247.56 | 974.97 | 890.63 | 2564.7 | 0 | 609.41 | 198.13 | 310.47 | 337.33 | 2971.34 | 1039.55 | 423.85 | 260.2 | 65.73 | 263.35 | 399.97 | 12977.97 | Terminal uridylyltransferase 7 (TUTase 7) (EC 2.7.7.52) (Zinc finger CCHC domain-containing protein 6) | TUT7 HS2 KIAA1711 ZCCHC6 | miRNA metabolic process [GO:0010586]; oocyte maturation [GO:0001556]; polyuridylation-dependent mRNA catabolic process [GO:1990074]; pre-miRNA processing [GO:0031054]; RNA 3'-end processing [GO:0031123]; transposable element silencing by mRNA destabilization [GO:0141008] |

| | | | | | | | | | | | | | | | | | | | | | | |
|---|---|---|---|---|---|---|---|---|---|---|---|---|---|---|---|---|---|---|---|---|---|---|
| 372.89 | | 1528.23 | 527.95 | 285.29 | 2514.89 | 400.91 | 21.61 | 739.03 | 304.76 | 313.38 | 509.94 | 1300.45 | 189.79 | 1115.79 | 327.22 | 346.53 | 1717.08 | 460.61 | 12976.35 | Protein Red (Cytokine IK) (IK factor) | Ik | mitotic cell cycle [GO:0000278]; mitotic spindle assembly checkpoint signaling [GO:0007094]; mRNA splicing, via spliceosome [GO:0000398]; protein localization to kinetochore [GO:0034501] |
| 635.71 | | 572.65 | 406.65 | 309.9 | 550.67 | 821.95 | 2743.2 | 267.35 | 433.47 | 226.41 | 210.14 | 453.92 | 909.5 | 988.68 | 521.79 | 1765.6 | 539.42 | 618.57 | 12975.58 | Cyclic AMP receptor-like protein A | crlA DDB_G0280983 | adenylate cyclase-activating G protein-coupled receptor signaling pathway [GO:0007189]; negative regulation of cell growth [GO:0030308]; signal transduction [GO:0007165]; sporulation resulting in formation of a cellular spore [GO:0030435] |

| | | | | | | | | | | | | | | | | | | | | | |
|---|---|---|---|---|---|---|---|---|---|---|---|---|---|---|---|---|---|---|---|---|---|
| 664.99 | 259.34 | 1060.04 | 194.08 | 243.46 | 775.34 | 203.72 | 98.87 | 2024.3 | 199.42 | 127.12 | 1024.3 | 750.24 | 680.54 | 1733.3 | 290.34 | 387.19 | 2257.63 | 12974.22 | Solute carrier family 4 member 11 (Sodium borate cotransporter 1) (NaBC1) | SLC4A11 BTR1 | bicarbonate transport [GO:0015701]; borate transport [GO:0046713]; cellular hypotonic response [GO:0071476]; cellular response to oxidative stress [GO:0034599]; fluid transport [GO:0042044]; intracellular monoatomic cation homeostasis [GO:0030003]; monoatomic anion transport [GO:0006820]; monoatomic ion homeostasis [GO:0050801]; proton transmembrane transport [GO:1902600]; regulation of mesenchymal stem cell differentiation [GO:2000739]; regulation of mitochondrial membrane potential [GO:0051881]; sodium ion transport [GO:0006814]; transmembrane transport [GO:0055085] |
| 499.03 | 681.45 | 1187.2 | 104.94 | 670.86 | 932.83 | 58.92 | 323 | 1496.4 | 205.34 | 247.42 | 1048.78 | 609.91 | 953.42 | 1653.64 | 143.58 | 401.8 | 1731.59 | 12950.11 | Histone-lysine N-methyltransferase 2C (Lysine N-methyltransferase 2C) (EC 2.1.1.364) (Homologous to ALR protein) (Myeloid/lymphoid or mixed-lineage leukemia protein 3) | KMT2C HALR KIAA1506 MLL3 | methylation [GO:0032259]; positive regulation of transcription by RNA polymerase II [GO:0045944] |

| | | | | | | | | | | | | | | | | | | | | | | |
|---|---|---|---|---|---|---|---|---|---|---|---|---|---|---|---|---|---|---|---|---|---|---|
| 312.69 | | 221.49 | 351.74 | 217.4 | 198.21 | 324.66 | 4792.1 | 103.42 | 540.22 | 163.48 | 156.31 | 414.01 | 448.65 | 276.08 | 544.51 | 2921.96 | 240.07 | 696.03 | 12923.03 | Chromodomain-helicase-DNA-binding protein 2 (CHD-2) (EC 3.6.4.-) (ATP-dependent helicase CHD2) | CHD2 | DNA damage response [GO:0006974]; gene expression [GO:0010467]; hematopoietic stem cell differentiation [GO:0060218]; muscle organ development [GO:0007517]; nucleosome organization [GO:0034728] |
| 643.26 | | 590.49 | 1952.84 | 326.61 | 586.1 | 1027.51 | 233.29 | 421.97 | 1006.33 | 368.9 | 445.8 | 483.81 | 580.47 | 1006.11 | 1253.73 | 299.05 | 362.59 | 1330.04 | 12918.9 | 3'-5' exoribonuclease HELZ2 (EC 3.1.13.1) (ATP-dependent RNA helicase PRIC285) (EC 3.6.4.13) (Helicase with zinc finger 2, transcriptional coactivator) (Helicase with zinc finger domain 2) (PPAR-alpha-interacting complex protein 285) (PPAR-gamma DNA-binding domain-interacting protein 1) (PDIP1) (PPAR-gamma DBD-interacting protein 1) (Peroxisomal [illegible] | HELZ2 KIAA1769 PRIC285 | positive regulation of transcription by RNA polymerase II [GO:0045944]; regulatory ncRNA-mediated post-transcriptional gene silencing [GO:0035194] |

| | | | | | | | | | | | | | | | | | | | | | |
|---|---|---|---|---|---|---|---|---|---|---|---|---|---|---|---|---|---|---|---|---|---|
| 0 | 0 | 12906.72 | 0 | 0 | 0 | 2.51 | 0 | 0 | 0 | 0 | 0 | 0 | 0 | 0 | 0 | 0 | 3.59 | 12912.82 | Cadherin-18 (Cadherin-14) | CDH18 CDH14 | adherens junction organization [GO:0034332]; calcium-dependent cell-cell adhesion via plasma membrane cell adhesion molecules [GO:0016339]; cell migration [GO:0016477]; cell morphogenesis [GO:0000902]; cell-cell adhesion mediated by cadherin [GO:0044331]; cell-cell junction assembly [GO:0007043]; homophilic cell adhesion via plasma membrane adhesion molecules [GO:0007156] |
| 966.57 | 641.19 | 2234.35 | 183.52 | 238.13 | 1455.21 | 3.84 | 157.21 | 795.88 | 194.75 | 193.33 | 491.5 | 2558.65 | 716.3 | 481.63 | 189.18 | 283.91 | 1125.92 | 12911.07 | Carboxypeptidase B2 (EC 3.4.17.20) (Carboxypeptidase R) (CPR) (Carboxypeptidase U) (CPU) (Thrombin-activable fibrinolysis inhibitor) (TAFI) | Cpb2 | blood coagulation [GO:0007596]; cellular response to glucose stimulus [GO:0071333]; fibrinolysis [GO:0042730]; liver regeneration [GO:0097421]; negative regulation of fibrinolysis [GO:0051918]; negative regulation of hepatocyte proliferation [GO:2000346]; negative regulation of plasminogen activation [GO:0010757]; positive regulation of extracellular matrix constituent secretion [GO:0003331]; protein catabolic process [GO:0030163]; proteolysis [GO:0006508]; response to xenobiotic stimulus [GO:0009410] |
| 563.95 | 661.19 | 554.15 | 312.4 | 682.71 | 1096.28 | 2457.7 | 202.99 | 340.57 | 192.95 | 232.21 | 630.75 | 946.61 | 1035.91 | 456.17 | 1469.33 | 636.45 | 427.22 | 12899.54 | Cytosolic non-specific dipeptidase (EC 3.4.13.18) (CNDP dipeptidase 2) (Threonyl dipeptidase) | CNDP2 | proteolysis [GO:0006508] |

| | | | | | | | | | | | | | | | | | | | | | |
|---|---|---|---|---|---|---|---|---|---|---|---|---|---|---|---|---|---|---|---|---|---|
| 27.52 | 29.53 | 2600.65 | 14.06 | 9.03 | 359.73 | 1.1 | 8.08 | 299.11 | 13.41 | 8.26 | 8307.49 | 35.8 | 9.86 | 483.44 | 17.67 | 28.57 | 616.73 | 12870.04 | Phospholipase DDHD1 (EC 3.1.1.111) (EC 3.1.1.32) (DDHD domain-containing protein 1) (Phosphatidic acid-preferring phospholipase A1 homolog) (PA-PLA1) (EC 3.1.1.118) (Phospholipid sn-1 acylhydrolase) | DDHD1 KIAA1705 | phosphatidylinositol metabolic process [GO:0046488]; positive regulation of mitochondrial fission [GO:0090141] |
| 816.21 | 1465.53 | 455.66 | 408.53 | 1028.81 | 672.9 | 420.58 | 626.99 | 380.3 | 261.26 | 319.44 | 575.91 | 1568.22 | 908.71 | 535.63 | 564.25 | 1313.05 | 547.88 | 12869.86 | Excitatory amino acid transporter 5 (Retinal glutamate transporter) (Solute carrier family 1 member 7) | SLC1A7 EAAT5 | chloride transmembrane transport [GO:1902476]; dicarboxylic acid transport [GO:0006835]; L-glutamate transmembrane transport [GO:0015813]; monoatomic ion transport [GO:0006811]; neurotransmitter transport [GO:0006836]; neurotransmitter uptake [GO:0001504] |
| 1910.24 | 464.04 | 406.95 | 817.53 | 304.01 | 1417.47 | 245.39 | 108.1 | 404.2 | 486.39 | 158.5 | 308.65 | 2536.98 | 1158.28 | 811.35 | 426.66 | 450.69 | 426.12 | 12841.55 | ATP synthase peripheral stalk subunit OSCP, mitochondrial (ATP synthase subunit O) (Oligomycin sensitivity conferral protein) (OSCP) | ATP5PO ATP5O | proton motive force-driven mitochondrial ATP synthesis [GO:0042776] |

| | | | | | | | | | | | | | | | | | | | | | |
|---|---|---|---|---|---|---|---|---|---|---|---|---|---|---|---|---|---|---|---|---|---|
| 503.27 | 901.89 | 578.78 | 263.02 | 969.9 | 645.33 | 595.04 | 385.73 | 713.21 | 232.08 | 329.42 | 487.45 | 471.08 | 2001.59 | 806.2 | 1137.16 | 691.97 | 1126.62 | 12839.74 | Transcription factor SOX-6 | SOX6 | brain development [GO:0007420]; cartilage condensation [GO:0001502]; cartilage development [GO:0051216]; cell fate commitment [GO:0045165]; chondrocyte differentiation [GO:0002062]; positive regulation of chondrocyte differentiation [GO:0032332]; spinal cord oligodendrocyte cell differentiation [GO:0021529] |
| 1606.55 | 482.17 | 1266.95 | 449.64 | 417.26 | 2108.22 | 284.52 | 224.8 | 291.26 | 587.05 | 263.96 | 509.84 | 1554.78 | 809.23 | 560.94 | 389.18 | 637.85 | 362.59 | 12806.79 | Inactive tyrosine-protein kinase PRAG1 (PEAK1-related kinase-activating pseudokinase 1) (Pragmin) (Sugen kinase 223) (SgK223) | PRAG1 SGK223 | cell migration [GO:0016477]; negative regulation of neuron projection development [GO:0010977]; positive regulation of Rho protein signal transduction [GO:0035025]; regulation of cell motility [GO:2000145]; regulation of cell shape [GO:0008360]; regulation of Notch signaling pathway [GO:0008593] |

| | | | | | | | | | | | | | | | | | | | | | |
|---|---|---|---|---|---|---|---|---|---|---|---|---|---|---|---|---|---|---|---|---|---|
| 916.94 | 944.76 | 890.07 | 374.42 | 1101.26 | 1499.58 | 359.38 | 489.98 | 186.85 | 351.75 | 522.41 | 604.73 | 1178.36 | 1071.5 | 572.94 | 460.16 | 962.84 | 311.95 | 12799.88 | Cytochrome P450 7B1 (24-hydroxycholesterol 7-alpha-hydroxylase) (EC 1.14.14.26) (25/26-hydroxycholesterol 7-alpha-hydroxylase) (EC 1.14.14.29) (3-hydroxysteroid 7-alpha hydroxylase) (Oxysterol 7-alpha-hydroxylase) | CYP7B1 | B cell chemotaxis [GO:0035754]; bile acid biosynthetic process [GO:0006699]; cholesterol homeostasis [GO:0042632]; cholesterol metabolic process [GO:0008203]; epithelial cell proliferation [GO:0050673]; estrogen receptor signaling pathway [GO:0030520]; negative regulation of intracellular estrogen receptor signaling pathway [GO:0033147]; positive regulation of epithelial cell proliferation [GO:0050679]; prostate gland epithelium morphogenesis [GO:0060740]; steroid biosynthetic process [GO:0006694]; sterol metabolic process [GO:0016125] |
| 416.25 | 148.51 | 203.9 | 342.62 | 157.39 | 362.94 | 5511.12 | 46.41 | 270.45 | 209.19 | 115.94 | 188.66 | 468.69 | 223.16 | 275.65 | 3269.87 | 234.68 | 338.47 | 12783.9 | Large ribosomal subunit protein mL65 (28S ribosomal protein S30, mitochondrial) (MRP-S30) (S30mt) | MRPS30 | translation [GO:0006412] |
| 1110.36 | 458.46 | 132.31 | 370.14 | 1215.64 | 1303.9 | 903.88 | 265.07 | 46.29 | 680.72 | 385.71 | 578.61 | 1679.86 | 1217.97 | 670.62 | 702.52 | 796.24 | 258.53 | 12776.83 | Spermatogenesis-associated protein 2 | spata2 | programmed cell death [GO:0012501]; protein K63-linked deubiquitination [GO:0070536]; protein linear deubiquitination [GO:1990108]; regulation of inflammatory response [GO:0050727]; regulation of necroptotic process [GO:0060544]; regulation of tumor necrosis factor-mediated signaling pathway [GO:0010803] |

| | | | | | | | | | | | | | | | | | | | | | |
|---|---|---|---|---|---|---|---|---|---|---|---|---|---|---|---|---|---|---|---|---|---|
| 933.92 | 368.87 | 2404.46 | 191.26 | 622.7 | 760.86 | 991.1 | 246.74 | 673.38 | 299.17 | 322.2 | 1076.66 | 470.12 | 264.02 | 923.87 | 709.86 | 284.16 | 1220.32 | 12763.67 | Formin-binding protein 1 (Formin-binding protein 17) (hFBP17) | FNBP1 FBP17 KIAA0554 | endocytosis [GO:0006897]; signal transduction [GO:0007165] |
| 2963.77 | 424.32 | 1196.38 | 599.72 | 262.72 | 1105.86 | 6.85 | 216.74 | 112.73 | 569.88 | 244.89 | 753.17 | 2113.03 | 809.37 | 260.28 | 131.2 | 803.4 | 160.68 | 12734.99 | Septin-9 (MLL septin-like fusion protein MSF-A) (MLL septin-like fusion protein) (Ovarian/Breast septin) (Ov/Br septin) (Septin D1) | SEPTIN9 KIAA0991 MSF SEPT9 | actin cytoskeleton organization [GO:0030036]; cytoskeleton-dependent cytokinesis [GO:0061640]; positive regulation of non-motile cilium assembly [GO:1902857]; protein localization [GO:0008104]; septin cytoskeleton organization [GO:0032185] |
| 2623.58 | 1026.5 | 527.82 | 429.68 | 639.21 | 668.28 | 225.67 | 1128.44 | 76.19 | 516.44 | 723.14 | 638.95 | 970.04 | 763.46 | 421.93 | 135.25 | 1042.79 | 117.1 | 12674.47 | 5'-3' exoribonuclease 2 (EC 3.1.13.-) (Protein Dhm1) | Xrn2 Dhm1 | DNA recombination [GO:0006310]; DNA repair [GO:0006281]; hippocampus development [GO:0021766]; microtubule-based process [GO:0007017]; mRNA processing [GO:0006397]; neuron differentiation [GO:0030182]; retina development in camera-type eye [GO:0060041]; RNA metabolic process [GO:0016070]; spermatogenesis [GO:0007283]; termination of RNA polymerase II transcription [GO:0006369] |

| | | | | | | | | | | | | | | | | | | | | | |
|---|---|---|---|---|---|---|---|---|---|---|---|---|---|---|---|---|---|---|---|---|---|
| 1410.76 | 437.95 | 550.04 | 371.4 | 333.4 | 883.74 | 138.3 | 176.34 | 673.76 | 282.35 | 124.63 | 544.93 | 3632.61 | 696.11 | 847.82 | 336.74 | 371.26 | 856.56 | 12668.7 | Protein wntless homolog (Integral membrane protein GPR177) | WLS GPR177 RCJMB04_5c7 | intracellular protein transport [GO:0006886]; positive regulation of Wnt protein secretion [GO:0061357]; positive regulation of Wnt signaling pathway [GO:0030177]; Wnt protein secretion [GO:0061355]; Wnt signaling pathway [GO:0016055] |
| 854.67 | 212 | 766.79 | 487.91 | 190.01 | 921.54 | 1562.68 | 69.74 | 569.56 | 508.27 | 127.2 | 749.56 | 3125.17 | 186.55 | 496.71 | 1222.27 | 125.12 | 489.09 | 12664.84 | Interferon gamma receptor 1 (Interferon gamma receptor 1-1) | ifngr1 ifngr1-1 | |
| 1240.3 | 382.8 | 650.42 | 676.6 | 467 | 1072.31 | 229.53 | 570.35 | 28.72 | 557.8 | 529.23 | 1464.67 | 2055.67 | 883.8 | 372.05 | 421.34 | 892.13 | 168.42 | 12663.14 | Mitogen-activated protein kinase 1 (MAP kinase 1) (MAPK 1) (EC 2.7.11.24) (ERT1) (Extracellular signal-regulated kinase 2) (ERK-2) (MAP kinase isoform p42) (p42-MAPK) (Mitogen-activated protein kinase 2) (MAP kinase 2) (MAPK 2) | Mapk1 Erk2 Mapk Prkm1 | androgen receptor signaling pathway [GO:0030521]; animal organ morphogenesis [GO:0009887]; apoptotic process [GO:0006915]; B cell receptor signaling pathway [GO:0050853]; Bergmann glial cell differentiation [GO:0060020]; cardiac neural crest cell development involved in heart development [GO:0061308]; caveolin-mediated endocytosis [GO:0072584]; cellular response to amino acid starvation [GO:0034198]; cellular response to tumor necrosis factor [GO:0071356]; cytosine metabolic process [GO:0019858]; DNA damage response [GO:0006974]; DNA-templated transcription [GO:0006351]; epidermal growth factor receptor signaling pathway [GO:0007173]; ERBB2-ERBB3 signaling pathway [GO:0038133]; ERK1 and ERK2 cascade [GO:0070371]; face development [GO:0060324]; heart development [GO:0007507]; insulin receptor signaling pathway [GO:0008286]; insulin-like growth factor receptor signaling pathway [illegible] |

| | | | | | | | | | | | | | | | | | | | | | |
|---|---|---|---|---|---|---|---|---|---|---|---|---|---|---|---|---|---|---|---|---|---|
| 761.96 | 706.15 | 701 | 322.85 | 710.38 | 1211.09 | 1452.59 | 319.14 | 332.23 | 212.56 | 269.06 | 1026.25 | 668.34 | 782.18 | 524.49 | 1583.44 | 494.8 | 554.96 | 12633.47 | Collagen alpha-1(VIII) chain (Endothelial collagen) [Cleaved into: Vastatin] | COL8A1 C3orf7 | angiogenesis [GO:0001525]; camera-type eye morphogenesis [GO:0048593]; cell adhesion [GO:0007155]; endodermal cell differentiation [GO:0035987]; endothelial cell proliferation [GO:0001935]; positive regulation of cell-substrate adhesion [GO:0010811] |
| 808.88 | 455.51 | 1209.57 | 239.11 | 410.95 | 726.3 | 941.56 | 178.69 | 1157.74 | 169.02 | 153.51 | 830.03 | 819.86 | 616.87 | 790.78 | 1014.68 | 412.54 | 1689.85 | 12625.45 | Probable N-acetyltransferase 14 (EC 2.3.1.-) | nat14 zgc:153234 | |
| 719.17 | 373.11 | 2246.36 | 278.73 | 557.4 | 766.32 | 3.11 | 688.17 | 819.62 | 457.53 | 598.07 | 2382.58 | 534.92 | 145.45 | 846.25 | 38.6 | 172.5 | 987.57 | 12615.46 | Polymeric immunoglobulin receptor (PIgR) (Poly-Ig receptor) [Cleaved into: Secretory component] | PIGR | immunoglobulin transcytosis in epithelial cells mediated by polymeric immunoglobulin receptor [GO:0002415] |

| | | | | | | | | | | | | | | | | | | | | | |
|---|---|---|---|---|---|---|---|---|---|---|---|---|---|---|---|---|---|---|---|---|---|
| 819.98 | 607.55 | 970.99 | 187.17 | 532.91 | 1072.77 | 1072.9 | 270.5 | 775.6 | 232.96 | 217 | 593.84 | 626.69 | 723.83 | 1455.79 | 755.14 | 526.91 | 1161.35 | 12603.88 | Myelin-associated glycoprotein (Siglec-4a) | MAG GMA | axon regeneration [GO:0031103]; cell adhesion [GO:0007155]; cell-cell adhesion via plasma-membrane adhesion molecules [GO:0098742]; cellular response to mechanical stimulus [GO:0071260]; central nervous system myelin formation [GO:0032289]; negative regulation of axon extension [GO:0030517]; negative regulation of neuron apoptotic process [GO:0043524]; negative regulation of neuron differentiation [GO:0045665]; negative regulation of neuron projection development [GO:0010977]; positive regulation of astrocyte differentiation [GO:0048711]; positive regulation of myelination [GO:0031643]; substantia nigra development [GO:0021762]; transmission of nerve impulse [GO:0019226] |
| 1008.8 | 557.92 | 856.82 | 304.67 | 447.07 | 934.86 | 1475.18 | 228.44 | 740.65 | 288.57 | 234.65 | 637.23 | 834.24 | 599.45 | 842.17 | 1333.12 | 444.69 | 832.89 | 12601.42 | Prominin-1-A (Prominin-like protein 1) | prom1a proml1 | |
| 4191.03 | 0 | 7.13 | 610.34 | 78.27 | 114.82 | 7.24 | 0 | 0 | 2274.54 | 51.35 | 91.28 | 4954.13 | 6.38 | 0 | 188.63 | 7.1 | 0 | 12582.24 | Galectin-3-binding protein B (Lectin galactoside-binding soluble 3-binding protein B) | lgals3bpb zgc:77059 | cell adhesion [GO:0007155] |
| 658.87 | 755.87 | 1006.49 | 246.66 | 571.09 | 1162.24 | 788.89 | 263.65 | 1023.55 | 208.36 | 236.23 | 1169.08 | 728.55 | 819.84 | 1003.66 | 443.12 | 483.31 | 1009.36 | 12578.82 | Metal transporter CNNM1 (Ancient conserved domain-containing protein 1) (Cyclin-M1) | CNNM1 ACDP1 | magnesium ion homeostasis [GO:0010960]; monoatomic ion transport [GO:0006811] |
| 449.75 | 534.07 | 536.43 | 193.07 | 525.57 | 454.13 | 1685.07 | 287.99 | 1136.64 | 164.87 | 189.45 | 587.92 | 488.92 | 651.59 | 1549.38 | 945.67 | 346.48 | 1817.51 | 12544.51 | Actinia tenebrosa protease inhibitors (Carboxypeptidase inhibitor SmCI-like) [Cleaved into: ATPI-I;ATPI-II] | | |

| | | | | | | | | | | | | | | | | | | | | |
|---|---|---|---|---|---|---|---|---|---|---|---|---|---|---|---|---|---|---|---|---|
| 111.2 | 23.74 | 18.15 | 84.14 | 32.45 | 336.65 | 18.78 | 1.21 | 4.19 | 47.97 | 59.24 | 115.07 | 7.72 | 6480.89 | 216.46 | 1584.34 | 3349.2 | 28.37 | 12519.77 | Serine/threonine-protein kinase 38-like (EC 2.7.11.1) (NDR2 protein kinase) (Nuclear Dbf2-related kinase 2) | STK38L KIAA0965 NDR2 | intracellular signal transduction [GO:0035556]; negative regulation of autophagy [GO:0010507]; protein phosphorylation [GO:0006468]; regulation of cellular component organization [GO:0051128] |
| 1609 | 535.03 | 1277.5 | 276.46 | 678.39 | 687.63 | 311.72 | 255.5 | 1171.34 | 492.94 | 235.61 | 725.27 | 472.19 | 659.27 | 1140.01 | 342.73 | 597.62 | 1035.34 | 12503.55 | Large neutral amino acids transporter small subunit 3 (L-type amino acid transporter 3) (Prostate cancer overexpressed gene 1 protein) (Solute carrier family 43 member 1) | SLC43A1 LAT3 PB39 POV1 | amino acid transport [GO:0006865]; isoleucine transport [GO:0015818]; L-leucine transport [GO:0015820]; L-valine transmembrane transport [GO:1903785]; negative regulation of amino acid transport [GO:0051956]; negative regulation of L-leucine import across plasma membrane [GO:1905533]; neutral amino acid transport [GO:0015804] |
| 1230.63 | 277.12 | 2466.18 | 354.16 | 216.73 | 1242.63 | 114.33 | 76.23 | 1794.85 | 297.32 | 54.17 | 617.02 | 1170.93 | 93.37 | 961.75 | 240.09 | 338.85 | 936.17 | 12482.53 | Probable E3 ubiquitin-protein ligase DTX3 (EC 2.3.2.27) (Protein deltex-3) (Deltex3) (mDTX3) (RING-type E3 ubiquitin transferase DTX3) | Dtx3 | Notch signaling pathway [GO:0007219]; protein ubiquitination [GO:0016567] |

| | | | | | | | | | | | | | | | | | | | | | |
|---|---|---|---|---|---|---|---|---|---|---|---|---|---|---|---|---|---|---|---|---|---|
| 4.18 | 5.86 | 413.85 | 1.23 | 2.54 | 3.06 | 9.51 | 3.38 | 4147.48 | 2.17 | 0.44 | 1066.34 | 108.2 | 13.52 | 2655.79 | 18.92 | 8.76 | 4016.97 | 12482.2 | Zinc finger SWIM domain-containing protein 8 | ZSWIM8 | positive regulation of miRNA catabolic process [GO:2000627]; proteasome-mediated ubiquitin-dependent protein catabolic process [GO:0043161]; protein ubiquitination [GO:0016567] |
| 1678.88 | 880.6 | 880.28 | 323.14 | 989.4 | 1372.73 | 155.5 | 670.75 | 105.85 | 414.99 | 447.15 | 883.4 | 896.77 | 1050.44 | 436.96 | 184.98 | 853.36 | 253.02 | 12478.2 | Pericentriolar material 1 protein (PCM-1) | PCM1 | cilium assembly [GO:0060271]; intraciliary transport involved in cilium assembly [GO:0035735]; microtubule anchoring at centrosome [GO:0034454]; non-motile cilium assembly [GO:1905515]; positive regulation of intracellular protein transport [GO:0090316]; protein localization to centrosome [GO:0071539] |

| | | | | | | | | | | | | | | | | | | | | | |
|---|---|---|---|---|---|---|---|---|---|---|---|---|---|---|---|---|---|---|---|---|---|
| 174.54 | 0 | 2092.92 | 18.23 | 0.19 | 67.51 | 4.38 | 0.92 | 3229.18 | 54.88 | 2.78 | 757.12 | 49.04 | 0 | 2203.51 | 3.19 | 3.6 | 3801.85 | 12463.84 | Suppressor of hairless protein homolog (X-Su(H)) | rbpj rbpsuh suh | Notch signaling pathway [GO:0007219]; positive regulation of transcription by RNA polymerase II [GO:0045944]; positive regulation of transcription of Notch receptor target [GO:0007221] |
| 52.51 | 2724.1 | 307.77 | 87.24 | 2131.45 | 323.1 | 121.36 | 1097.72 | 143.24 | 35.78 | 717.29 | 283.1 | 52.09 | 2573.96 | 145.83 | 177.2 | 1328.01 | 150.33 | 12452.08 | X-linked retinitis pigmentosa GTPase regulator-interacting protein 1 (RPGR-interacting protein 1) | RPGRIP1 | non-motile cilium assembly [GO:1905515]; visual perception [GO:0007601] |
| 578.81 | 613.78 | 667.34 | 457.65 | 644.71 | 817.14 | 1534.47 | 235.54 | 535.48 | 224.01 | 250.35 | 726.13 | 901.09 | 982.49 | 732 | 1198.76 | 540.23 | 810.06 | 12450.04 | Unconventional myosin-VI (Unconventional myosin-6) | MYO6 | actin filament organization [GO:0007015]; actin filament-based movement [GO:0030048]; DNA damage response, signal transduction by p53 class mediator [GO:0030330]; endocytosis [GO:0006897]; inner ear auditory receptor cell differentiation [GO:0042491]; inner ear morphogenesis [GO:0042472]; intracellular protein transport [GO:0006886]; protein localization [GO:0008104]; regulation of secretion [GO:0051046]; sensory perception of sound [GO:0007605] |
| 420.43 | 551.61 | 1303.94 | 345.19 | 386.74 | 2483.99 | 164.83 | 166.27 | 569.81 | 224.98 | 136.44 | 1071.66 | 1392.41 | 568.73 | 1064.39 | 328.24 | 497.23 | 738.23 | 12415.12 | Tropomodulin-4 (Skeletal muscle tropomodulin) (Sk-Tmod) | Tmod4 | pointed-end actin filament capping [GO:0051694] |

| | | | | | | | | | | | | | | | | | | | | | |
|---|---|---|---|---|---|---|---|---|---|---|---|---|---|---|---|---|---|---|---|---|---|
| 644.14 | 793.04 | 434.14 | 308.9 | 646.14 | 1344.13 | 1905.1 | 235.62 | 252.38 | 262.98 | 286.05 | 956.09 | 1031.04 | 522.47 | 412.5 | 1621.63 | 487.74 | 265.25 | 12409.34 | Leucine-rich repeat and fibronectin type III domain-containing protein 1-like protein | lrfn1l zgc:172282 | |
| 417.35 | 983.81 | 547.72 | 323 | 789.41 | 1341 | 1365.34 | 218.41 | 227.66 | 156.78 | 232 | 912.96 | 1005.76 | 1044.66 | 557.7 | 1223.85 | 656.78 | 403.2 | 12407.39 | Inositol hexakisphosphate kinase 2 (InsP6 kinase 2) (EC 2.7.4.-) (P(i)-uptake stimulator) (PiUS) | IP6K2 IHPK2 | cellular response to flavonoid [GO:1905396]; inositol phosphate biosynthetic process [GO:0032958]; inositol phosphate metabolic process [GO:0043647]; negative regulation of cell growth [GO:0030308]; phosphatidylinositol phosphate biosynthetic process [GO:0046854]; positive regulation of apoptotic process [GO:0043065]; protein stabilization [GO:0050821] |
| 50.45 | 2447.61 | 225.93 | 37.3 | 1365.7 | 138.71 | 22.37 | 1070.57 | 175.92 | 36.38 | 611.09 | 210.24 | 28.51 | 3862.16 | 200.62 | 40.74 | 1610.21 | 268.68 | 12403.19 | Dol-P-Man:Man(5)GlcNAc(2)-PP-Dol alpha-1,3-mannosyltransferase (EC 2.4.1.258) (Asparagine-linked glycosylation protein 3 homolog) (Dol-P-Man-dependent alpha(1-3)-mannosyltransferase) (Dolichyl-P-Man:Man(5)GlcNAc(2)-PP-dolichyl mannosyltransferase) (Dolichyl-phosphate-mannose--glycolipid alpha-mannosyltransferase) | Alg3 | dolichol-linked oligosaccharide biosynthetic process [GO:0006488]; protein N-linked glycosylation [GO:0006487] |

| | | | | | | | | | | | | | | | | | | | | | |
|---|---|---|---|---|---|---|---|---|---|---|---|---|---|---|---|---|---|---|---|---|---|
| 199.12 | 260.22 | 490.58 | 110.35 | 263.49 | 457.57 | 4867.86 | 102.29 | 466.49 | 183.74 | 186.39 | 411.14 | 310.5 | 241.3 | 503.6 | 2487.17 | 280.65 | 573.48 | 12395.94 | Protein mono-ADP-ribosyltransferase PARP11 (EC 2.4.2.-) (ADP-ribosyltransferase diphtheria toxin-like 11) (ARTD11) (Poly [ADP-ribose] polymerase 11) (PARP-11) | Parp11 | cell differentiation [GO:0030154]; mRNA transport [GO:0051028]; nuclear envelope organization [GO:0006998]; protein auto-ADP-ribosylation [GO:0070213]; protein transport [GO:0015031]; spermatogenesis [GO:0007283] |
| 978.78 | 805.61 | 1232.94 | 215.02 | 726.88 | 1010.03 | 20.13 | 290.33 | 915.93 | 274.22 | 274.97 | 806.16 | 694.09 | 1064.39 | 1174.45 | 141.57 | 492.02 | 1265.63 | 12383.15 | Mesoderm induction early response protein 1 (Early response 1) (Er1) (Mi-er1) | Mier1 Kiaa1610 | chromatin remodeling [GO:0006338]; regulation of DNA-templated transcription [GO:0006355] |
| 949 | 301.66 | 732.21 | 124.99 | 108.32 | 301.67 | 2114.96 | 111.39 | 410.97 | 88.08 | 115.28 | 342.78 | 709.46 | 2589.48 | 517.21 | 1401.22 | 1193.47 | 255.66 | 12367.81 | DBIRD complex subunit ZNF326 (Zinc finger protein 326) (Zinc finger protein interacting with mRNPs) | ZNF326 ZIRD | mRNA processing [GO:0006397]; regulation of DNA-templated transcription elongation [GO:0032784]; regulation of RNA splicing [GO:0043484]; RNA splicing [GO:0008380] |

| | | | | | | | | | | | | | | | | | | | | | |
|---|---|---|---|---|---|---|---|---|---|---|---|---|---|---|---|---|---|---|---|---|---|
| 759.43 | 980.76 | 1282.41 | 827.96 | 938.55 | 1613.28 | 0 | 18.23 | 26.04 | 496.42 | 401.26 | 1224.22 | 1228.82 | 1775.3 | 737.61 | 4.11 | 21.58 | 27.97 | 12363.95 | Serine/threonine-protein phosphatase PP1-beta (EC 3.1.3.16) (Glc seven-like phosphatase 2) | gsp-2 CBG00598 | cell division [GO:0051301]; chromatin organization [GO:0006325]; meiotic cell cycle [GO:0051321] |
| 29.64 | 0 | 0.93 | 170.96 | 1.78 | 3.04 | 6201.13 | 0 | 0 | 41.5 | 175.56 | 0 | 18.88 | 2.44 | 0 | 5607.4 | 105.29 | 1.1 | 12359.65 | GRB2-associated-binding protein 1 (GRB2-associated binder 1) (Growth factor receptor bound protein 2-associated protein 1) | GAB1 | signal transduction [GO:0007165] |
| 0 | 0.61 | 4714.48 | 0 | 1.8 | 12.24 | 8.06 | 0 | 2349.14 | 0 | 0 | 746.59 | 0 | 6.09 | 1472.11 | 6.41 | 2.59 | 3036.54 | 12356.66 | Glutamate receptor ionotropic, NMDA 3B (GluN3B) (N-methyl-D-aspartate receptor subtype 3B) (NMDAR3B) (NR3B) | GRIN3B | ionotropic glutamate receptor signaling pathway [GO:0035235]; modulation of chemical synaptic transmission [GO:0050804]; monoatomic cation transmembrane transport [GO:0098655]; protein insertion into membrane [GO:0051205]; regulation of calcium ion transport [GO:0051924]; regulation of synaptic plasticity [GO:0048167]; synaptic transmission, glutamatergic [GO:0035249] |

| | | | | | | | | | | | | | | | | | | | | | |
|---|---|---|---|---|---|---|---|---|---|---|---|---|---|---|---|---|---|---|---|---|---|
| 480.52 | 410.92 | 831.21 | 130.95 | 409.7 | 622.46 | 2668.56 | 181.27 | 689.55 | 215.49 | 205.8 | 635.51 | 547.46 | 513.32 | 961.79 | 1545.7 | 330.08 | 966.47 | 12346.76 | Macrophage mannose receptor 1 (MMR) (C-type lectin domain family 13 member D) (C-type lectin domain family 13 member D-like) (Human mannose receptor) (hMR) (Macrophage mannose receptor 1-like protein 1) (CD antigen CD206) | MRC1 CLEC13D CLEC13DL MRC1L1 | cellular response to interleukin-4 [GO:0071353]; cellular response to lipopolysaccharide [GO:0071222]; cellular response to type II interferon [GO:0071346]; receptor-mediated endocytosis [GO:0006898] |
| 1022.83 | 741.78 | 1019.27 | 298.08 | 677.49 | 1278.53 | 94.66 | 528.08 | 525.62 | 377.95 | 203.7 | 1110.91 | 1156.57 | 1139.52 | 700.47 | 171.05 | 578.23 | 706.44 | 12331.18 | SEC14-like protein 2 (Alpha-tocopherol-associated protein) (TAP) (bTAP) | SEC14L2 | |
| 412.13 | 829.39 | 2113.89 | 153.53 | 793.47 | 2418.82 | 29.88 | 320.13 | 223.94 | 184.15 | 340.53 | 997.14 | 221.67 | 905.82 | 718.49 | 179.01 | 867.64 | 616.34 | 12325.97 | Membrane-associated guanylate kinase, WW and PDZ domain-containing protein 1 (Atrophin-1-interacting protein 3) (AIP-3) (BAI1-associated protein 1) (BAP-1) (Membrane-associated guanylate kinase inverted 1) (MAGI-1) (Trinucleotide repeat-containing gene 19 protein) (WW domain-containing protein 3) (WWP3) | MAGI1 AIP3 BAIAP1 BAP1 TNRC19 | cell adhesion [GO:0007155]; cell surface receptor signaling pathway [GO:0007166]; endothelial cell morphogenesis [GO:0001886]; positive regulation of cell-cell adhesion [GO:0022409]; protein-containing complex assembly [GO:0065003]; signal transduction [GO:0007165] |
| 627.6 | 555.65 | 2831.4 | 219.14 | 555.28 | 1289.45 | 186.14 | 188.18 | 418.53 | 257.12 | 194.42 | 1783.11 | 438.63 | 911.24 | 618.87 | 404.72 | 398.42 | 443.87 | 12321.77 | Zinc finger protein 276 (Zfp-276) (Zinc finger protein 477) | ZNF276 CENP-Z ZFP276 ZNF477 | regulation of transcription by RNA polymerase II [GO:0006357] |

| | | | | | | | | | | | | | | | | | | | | | | |
|---|---|---|---|---|---|---|---|---|---|---|---|---|---|---|---|---|---|---|---|---|---|---|
| 468.02 | | 775.27 | 789.82 | 142.3 | 751.14 | 703.62 | 595.26 | 304.04 | 1301.39 | 158.64 | 278.54 | 752.39 | 368.35 | 1066.67 | 1340.54 | 553.84 | 435.04 | 1535.54 | 12320.41 | Progranulin (PGRN) (Acrogranin) (Epithelin precursor) (Glycoprotein of 88 Kda) (GP88) (Glycoprotein 88) (Granulin precursor) (PC cell-derived growth factor) (PCDGF) (Proepithelin) (PEPI) [Cleaved into: Paragranulin;Granulin-1 (Granulin G);Granulin-2 (Granulin F);Granulin-3 (Epithelin-2) (Granulin B);Granulin-4 (Epithelin-1) (Granulin | GRN | astrocyte activation involved in immune response [GO:0002265]; blastocyst hatching [GO:0001835]; embryo implantation [GO:0007566]; epithelial cell proliferation [GO:0050673]; locomotory exploration behavior [GO:0035641]; lysosomal lumen acidification [GO:0007042]; lysosomal protein catabolic process [GO:1905146]; lysosomal transport [GO:0007041]; lysosome organization [GO:0007040]; maintenance of synapse structure [GO:0099558]; microglial cell activation involved in immune response [GO:0002282]; negative regulation of microglial cell activation [GO:1903979]; negative regulation of neuron apoptotic process [GO:0043524]; negative regulation of neutrophil activation [GO:1902564]; negative regulation of respiratory burst involved in inflammatory response [GO:0060266]; positive regulation of angiogenesis [GO:0045766]; positive regulation of aspartic-type peptidase activity [GO:1905247]; positive regulation of |
| 450.49 | | 608.48 | 1561.63 | 243.11 | 766.55 | 916.92 | 172.83 | 526.65 | 778.16 | 195.2 | 284.47 | 1363.09 | 790.01 | 807.95 | 801.02 | 199.56 | 548.54 | 1301.01 | 12315.67 | Cysteine-rich and transmembrane domain-containing protein 1 | CYSTM1 | |
| 711.7 | | 878.06 | 496.83 | 255.56 | 447.38 | 866.28 | 1730.09 | 323.26 | 556.45 | 220.61 | 271.18 | 482.7 | 1025.53 | 930.11 | 564.65 | 1192.48 | 688.91 | 652.1 | 12293.88 | Capping protein, Arp2/3 and myosin-I linker protein 2 (Capping protein regulator and myosin 1 linker 2) (F-actin-uncapping protein RLTPR) (Leucine-rich repeat-containing protein 16C) (RGD, leucine-rich repeat, tropomodulin and proline-rich-containing protein) | CARMIL2 LRRC16C RLTPR | actin filament network formation [GO:0051639]; cell migration [GO:0016477]; establishment or maintenance of cell polarity [GO:0007163]; establishment or maintenance of monopolar cell polarity [GO:0061339]; negative regulation of barbed-end actin filament capping [GO:2000813]; positive regulation of cell migration [GO:0030335]; positive regulation of extracellular matrix disassembly [GO:0090091]; positive regulation of lamellipodium assembly [GO:0010592]; positive regulation of lamellipodium organization [GO:1902745]; positive regulation of ruffle assembly [GO:1900029]; regulation of Arp2/3 complex-mediated actin nucleation [GO:0034315]; wound healing, spreading of cells [GO:0044319] |

| | | | | | | | | | | | | | | | | | | | | | |
|---|---|---|---|---|---|---|---|---|---|---|---|---|---|---|---|---|---|---|---|---|---|
| 450.45 | 558.66 | 1055.8 | 220.21 | 394.33 | 1333.64 | 1427.19 | 181.71 | 688.17 | 238.1 | 189.09 | 986.3 | 613.48 | 591 | 681.77 | 1228.47 | 453.13 | 1000.17 | 12291.67 | Exocyst complex component 4 (Exocyst complex component Sec8) | EXOC4 KIAA1699 SEC8 SEC8L1 | chemical synaptic transmission [GO:0007268]; exocytosis [GO:0006887]; Golgi to plasma membrane transport [GO:0006893]; membrane fission [GO:0090148]; mitotic cytokinesis [GO:0000281]; paraxial mesoderm formation [GO:0048341]; protein transmembrane transport [GO:0071806]; regulation of macroautophagy [GO:0016241]; vesicle docking involved in exocytosis [GO:0006904]; vesicle tethering involved in exocytosis [GO:0090522] |
| 0 | 0 | 12285.47 | 0 | 0 | 0 | 0 | 0 | 0 | 0 | 0 | 3.4 | 0 | 0 | 0 | 0 | 0 | 0 | 12288.87 | DELTA-sagatoxin-Srs1a (DELTA-SATX-Srs1a) (Cytolysin Src-1) (Src I) | | cytolysis in another organism [GO:0051715]; monoatomic cation transport [GO:0006812]; pore complex assembly [GO:0046931] |
| 3.08 | 1.16 | 423.13 | 0 | 1.67 | 15.72 | 0 | 0 | 4284.95 | 3.98 | 5.8 | 1060.26 | 0 | 0.71 | 2879.11 | 3.78 | 4.46 | 3591.77 | 12279.58 | Uncharacterized protein 058R | FV3-058R | |

| | | | | | | | | | | | | | | | | | | | | | | |
|---|---|---|---|---|---|---|---|---|---|---|---|---|---|---|---|---|---|---|---|---|---|---|
| 2708.84 | 59.3 | 68.32 | 135.97 | 108.89 | 21.48 | 0 | 13.3 | 0.99 | 70.9 | 9.08 | 61.18 | 6412.11 | 426.38 | 175.46 | 1855.66 | 135.83 | 12.11 | 12275.8 | Survival motor neuron protein 1 | smn1 smn | | axon arborization [GO:0140060]; axon development [GO:0061564]; axon extension [GO:0048675]; axonogenesis [GO:0007409]; dendrite development [GO:0016358]; motor neuron axon guidance [GO:0008045]; mRNA splicing, via spliceosome [GO:0000398]; neuromast regeneration [GO:0070657]; neuromuscular junction development [GO:0007528]; neuron cellular homeostasis [GO:0070050]; peripheral nervous system neuron axonogenesis [GO:0048936]; positive regulation of oxidative phosphorylation [GO:1903862]; protein complex oligomerization [GO:0051259]; regulation of axon extension involved in axon guidance [GO:0048841]; regulation of neuromuscular junction development [GO:1904396]; spliceosomal snRNP assembly [GO:0000387]; synaptic assembly at neuromuscular junction [GO:0051124]; tissue regeneration [GO:0042246] |
| 18.02 | 9.92 | 604.32 | 3.05 | 3.07 | 149.71 | 0 | 11.81 | 3674.43 | 7.92 | 11.36 | 993.03 | 6.78 | 31.18 | 3739.59 | 20.98 | 2.84 | 2978.87 | 12266.88 | Speriolin-like protein (Spermatogenesis and centriole-associated protein 1-like protein) | SPATC1L C21orf56 | | actin polymerization or depolymerization [GO:0008154]; positive regulation of cAMP/PKA signal transduction [GO:0141163]; spermatogenesis [GO:0007283] |
| 254.28 | 285.37 | 535.14 | 374.94 | 276.62 | 176.8 | 1093.42 | 115.5 | 1361.14 | 194.17 | 285.27 | 635.13 | 250.17 | 1226.94 | 1447.63 | 1059.82 | 870.91 | 1814.91 | 12258.16 | Ubiquitin carboxyl-terminal hydrolase 21 (EC 3.4.19.12) (Deubiquitinating enzyme 21) (Ubiquitin thioesterase 21) (Ubiquitin-specific-processing protease 21) | Usp21 | | neuron projection development [GO:0031175]; protein deubiquitination [GO:0016579]; proteolysis [GO:0006508]; transcription initiation-coupled chromatin remodeling [GO:0045815] |

| | | | | | | | | | | | | | | | | | | | | | |
|---|---|---|---|---|---|---|---|---|---|---|---|---|---|---|---|---|---|---|---|---|---|
| 327.8 | 270.4 | 1030.31 | 101.09 | 278.99 | 643.64 | 12.86 | 145.02 | 1959.07 | 165.7 | 166.66 | 1113.11 | 306.92 | 512.58 | 2430.45 | 46 | 217.8 | 2527.14 | 12255.54 | Thrombospondin-1 (Glycoprotein G) | THBS1 TSP-1 TSP1 | behavioral response to pain [GO:0048266]; cell adhesion [GO:0007155]; negative regulation of angiogenesis [GO:0016525]; response to endoplasmic reticulum stress [GO:0034976]; response to unfolded protein [GO:0006986] |
| 759.21 | 751.83 | 759.89 | 311.48 | 782.19 | 1006.8 | 665.49 | 461.92 | 824.17 | 281.32 | 375.36 | 796.75 | 913.51 | 820.35 | 796.64 | 718.56 | 551.74 | 673.39 | 12250.6 | BTB/POZ domain-containing protein 6-A | btbd6a zgc:172197 | neurogenesis [GO:0022008]; regulation of neurogenesis [GO:0050767] |
| 1361.53 | 1610.88 | 359.14 | 361.45 | 2225.08 | 575.25 | 132.97 | 687.49 | 67.21 | 276.02 | 416.87 | 187.42 | 726.3 | 2106.08 | 225.98 | 182.54 | 613.89 | 127.11 | 12243.21 | Integrator complex subunit 9 (Int9) | Ints9 D14Ertd231e | negative regulation of transforming growth factor beta receptor signaling pathway [GO:0030512]; RNA polymerase II transcription initiation surveillance [GO:0160240]; snRNA 3'-end processing [GO:0034472] |

| | | | | | | | | | | | | | | | | | | | | | |
|---|---|---|---|---|---|---|---|---|---|---|---|---|---|---|---|---|---|---|---|---|---|
| 647.98 | 1428.01 | 366.4 | 247.87 | 1227.71 | 936.71 | 2.02 | 1405.27 | 24.79 | 324.02 | 577.71 | 258.4 | 1204.74 | 1484.67 | 308.92 | 68.81 | 1511.21 | 212.82 | 12238.06 | FAD synthase (EC 2.7.7.2) (FAD pyrophosphorylase) (FMN adenylyltransferase) (Flavin adenine dinucleotide synthase) [Includes: Molybdenum cofactor biosynthesis protein-like region;FAD synthase region] | flad1 zgc:91843 | FAD biosynthetic process [GO:0006747] |
| 681.66 | 434.47 | 638.14 | 254.27 | 275.07 | 562.19 | 1722.49 | 86.24 | 1179.16 | 159.43 | 139.19 | 592.93 | 1017.3 | 393.88 | 1309.21 | 1242.56 | 262.78 | 1284.25 | 12235.22 | Selenocysteine insertion sequence-binding protein 2 (SECIS-binding protein 2) | SECISBP2 SBP2 | forebrain neuron development [GO:0021884]; mRNA stabilization [GO:0048255]; negative regulation of nuclear-transcribed mRNA catabolic process, nonsense-mediated decay [GO:2000623]; RNA catabolic process [GO:0006401]; selenocysteine incorporation [GO:0001514]; striatum development [GO:0021756] |

| | | | | | | | | | | | | | | | | | | | | | |
|---|---|---|---|---|---|---|---|---|---|---|---|---|---|---|---|---|---|---|---|---|---|
| 9.94 | 23.86 | 615.36 | 9.55 | 7.34 | 110.67 | 25.03 | 6.36 | 2336.46 | 4.32 | 30.59 | 576.51 | 0 | 28.35 | 3875.73 | 37.92 | 14.31 | 4515.39 | 12227.69 | Immunoglobulin-binding protein 1 (B-cell signal transduction molecule alpha 4) (Protein alpha-4) (CD79a-binding protein 1) (Protein phosphatase 2/4/6 regulatory subunit) (Renal carcinoma antigen NY-REN-16) | IGBP1 IBP1 | B cell activation [GO:0042113]; intracellular signal transduction [GO:0035556]; negative regulation of stress-activated MAPK cascade [GO:0032873]; negative regulation of transcription by RNA polymerase II [GO:0000122]; regulation of dephosphorylation [GO:0035303]; regulation of microtubule-based movement [GO:0060632]; response to interleukin-1 [GO:0070555]; response to tumor necrosis factor [GO:0034612]; signal transduction [GO:0007165] |
| 1447.06 | 964.78 | 1117.45 | 341.24 | 1016.61 | 828.04 | 13.04 | 350.32 | 727.76 | 297.86 | 318.46 | 659.5 | 698.69 | 1153.67 | 732.15 | 82.38 | 533.64 | 942.45 | 12225.1 | Interleukin-1 receptor accessory protein (IL-1 receptor accessory protein) (IL-1RAcP) (EC 3.2.2.6) | Il1rap | cytokine-mediated signaling pathway [GO:0019221]; inflammatory response [GO:0006954]; interleukin-1-mediated signaling pathway [GO:0070498]; interleukin-33-mediated signaling pathway [GO:0038172]; positive regulation of interleukin-13 production [GO:0032736]; positive regulation of interleukin-4 production [GO:0032753]; positive regulation of interleukin-5 production [GO:0032754]; positive regulation of interleukin-6 production [GO:0032755]; positive regulation of synapse assembly [GO:0051965]; regulation of postsynaptic density assembly [GO:0099151]; regulation of presynapse assembly [GO:1905606]; synaptic membrane adhesion [GO:0099560]; trans-synaptic signaling by trans-synaptic complex [GO:0099545] |

| | | | | | | | | | | | | | | | | | | | | | |
|---|---|---|---|---|---|---|---|---|---|---|---|---|---|---|---|---|---|---|---|---|---|
| 2048.85 | 240.46 | 376.7 | 839.74 | 245.24 | 472.17 | 2.38 | 98.35 | 45.72 | 590.7 | 163.87 | 416.33 | 5010.74 | 502.74 | 337.72 | 242.27 | 458.17 | 132.76 | 12224.91 | Zinc transporter ZIP1 (Solute carrier family 39 member 1) (Zinc-iron-regulated transporter-like) (Zrt- and Irt-like protein 1) (ZIP-1) (hZIP1) | SLC39A1 IRT1 ZIP1 ZIRTL CGI-08 CGI-71 | embryonic cranial skeleton morphogenesis [GO:0048701]; in utero embryonic development [GO:0001701]; limb development [GO:0060173]; monoatomic cation transport [GO:0006812]; zinc ion transmembrane transport [GO:0071577] |
| 1784.39 | 686.19 | 666.3 | 262.84 | 759.54 | 865.88 | 1777.64 | 292.82 | 399.68 | 174.72 | 259.92 | 429.94 | 497.67 | 918.07 | 346.8 | 890.61 | 656.33 | 551.15 | 12220.49 | Cytokine receptor-like factor 3 (Clf-3 protein) | crlf3 clf-3 zgc:110212 | |
| 714.08 | 460.03 | 1391.8 | 278.83 | 505.7 | 1845.88 | 347.83 | 236.07 | 686.53 | 301.13 | 185.81 | 832.8 | 1147.02 | 483.65 | 1079.7 | 451.52 | 246.55 | 1017.28 | 12212.21 | Kinesin-like protein KIF16B (Sorting nexin-23) | KIF16B C20orf23 KIAA1590 SNX23 | early endosome to late endosome transport [GO:0045022]; endoderm development [GO:0007492]; epidermal growth factor receptor signaling pathway [GO:0007173]; fibroblast growth factor receptor signaling pathway [GO:0008543]; formation of primary germ layer [GO:0001704]; Golgi to endosome transport [GO:0006895]; receptor catabolic process [GO:0032801]; regulation of receptor recycling [GO:0001919]; vesicle transport along microtubule [GO:0047496] |
| 857.78 | 759.7 | 513.41 | 296.97 | 574.03 | 705.29 | 1794.73 | 317.25 | 517.47 | 233.71 | 271.16 | 443.34 | 1026.8 | 977.22 | 555.13 | 1073.58 | 696.05 | 590.82 | 12204.44 | Glycogen phosphorylase, muscle form (EC 2.4.1.1) (Myophosphorylase) | Pygm | glycogen catabolic process [GO:0005980] |

| | | | | | | | | | | | | | | | | | | | | | |
|---|---|---|---|---|---|---|---|---|---|---|---|---|---|---|---|---|---|---|---|---|---|
| 2.58 | 0.84 | 968.51 | 4.66 | 1.91 | 165.27 | 2.29 | 1.21 | 3273.39 | 2.58 | 3.64 | 1405.61 | 7.44 | 2.89 | 2476.99 | 7.77 | 8.8 | 3854.77 | 12191.15 | Sorting nexin-7 | Snx7 | positive regulation of autophagosome assembly [GO:2000786]; protein transport [GO:0015031] |
| 188.86 | 425.07 | 1778.3 | 582.46 | 642.54 | 775.95 | 1417.08 | 420.96 | 372.6 | 213.37 | 378.38 | 1120.2 | 382.77 | 474.59 | 668.91 | 1111.07 | 518.21 | 708.05 | 12179.37 | Vitronectin (VN) (Glycoprotein 66) (S-protein) (Serum-spreading factor) | VTN | cell adhesion mediated by integrin [GO:0033627]; cell-matrix adhesion [GO:0007160]; immune response [GO:0006955] |
| 460.61 | 439.44 | 1047.02 | 133.93 | 331.18 | 617.78 | 753.7 | 177.22 | 1597.35 | 132.06 | 153.19 | 896.27 | 654.25 | 578.65 | 1369.62 | 658.71 | 344.51 | 1828.4 | 12173.89 | Coiled-coil domain-containing protein 50 | CCDC50 RCJMB04_2n23 | |
| 0 | 0 | 2032.02 | 0 | 0 | 350.89 | 0 | 0 | 38.39 | 0 | 0 | 9596.84 | 0 | 0 | 126.05 | 0 | 0 | 19.99 | 12164.18 | Saxitoxin and tetrodotoxin-binding protein 1 | psbp1 pstbp1 | response to toxic substance [GO:0009636] |
| 1861.43 | 79.94 | 412.56 | 596.79 | 193.8 | 565.75 | 411.78 | 50.46 | 182.59 | 2029.71 | 164.47 | 247.4 | 2586.84 | 236.73 | 255.09 | 438.45 | 422.25 | 1419.18 | 12155.22 | 28S rRNA (cytosine-C(5))-methyltransferase (EC 2.1.1.-) (Nucleolar protein 1) (Nucleolar protein 2 homolog) (Proliferating-cell nucleolar antigen p120) (Proliferation-associated nucleolar protein p120) | Nop2 Nol1 | blastocyst formation [GO:0001825]; regulation of signal transduction by p53 class mediator [GO:1901796]; ribosomal large subunit assembly [GO:0000027]; ribosomal large subunit biogenesis [GO:0042273]; rRNA processing [GO:0006364] |

| | | | | | | | | | | | | | | | | | | | | | |
|---|---|---|---|---|---|---|---|---|---|---|---|---|---|---|---|---|---|---|---|---|---|
| 1738.25 | 640.92 | 449.32 | 314.02 | 443.91 | 431.17 | 2570.9 | 233.82 | 136.13 | 253.84 | 423.77 | 401.38 | 198.84 | 1384.23 | 305.93 | 1730.96 | 332.99 | 143.01 | 12133.39 | Ribosomal protein eL22-like 1 (60S ribosomal protein L22-like 1) (Large ribosomal subunit protein eL22-like 1) | rpl22l1 | translation [GO:0006412] |
| 532.73 | 717.29 | 407.77 | 326.18 | 785.72 | 1933.09 | 47.62 | 649.45 | 715.34 | 548.8 | 753.18 | 389.53 | 533.78 | 1544.71 | 972.87 | 234.99 | 381.54 | 654.74 | 12129.33 | Ataxin-7-like protein 1 (Ataxin-7-like protein 4) | ATXN7L1 ATXN7L4 KIAA1218 | |
| 747.72 | 979.49 | 1191.3 | 215.28 | 866.93 | 933.02 | 8.45 | 567.01 | 904.67 | 262.12 | 327.29 | 650.49 | 719.07 | 1128.31 | 813.34 | 110.31 | 620.58 | 1080.43 | 12125.81 | Beta-1,4-mannosyl-glycoprotein 4-beta-N-acetylglucosaminyltransferase (EC 2.4.1.144) (N-glycosyl-oligosaccharide-glycoprotein N-acetylglucosaminyltransferase III) (GNT-III) (GlcNAc-T III) (N-acetylglucosaminyltransferase III) | Mgat3 Gnt3 | amyloid-beta metabolic process [GO:0050435]; cellular response to oxidative stress [GO:0034599]; cognition [GO:0050890]; lysosomal protein catabolic process [GO:1905146]; N-acetylglucosamine metabolic process [GO:0006044]; negative regulation of lysosomal protein catabolic process [GO:1905166]; positive regulation of protein localization to early endosome [GO:1902966]; protein localization [GO:0008104]; protein localization to early endosome [GO:1902946]; protein N-linked glycosylation [GO:0006487]; regulation of cell migration [GO:0030334] |
| 324.09 | 554.1 | 510.93 | 651.6 | 530.39 | 1712.41 | 6.21 | 547.65 | 851.54 | 383.12 | 335.23 | 2449.16 | 712.45 | 446.86 | 777.53 | 86.23 | 237.24 | 988.96 | 12105.7 | Transcription factor TFIIIB component B'' homolog (Transcription factor IIIB 150) (TFIIIB150) (Transcription factor-like nuclear regulator) | BDP1 KIAA1241 KIAA1689 TFNR | RNA polymerase III preinitiation complex assembly [GO:0070898] |

| | | | | | | | | | | | | | | | | | | | | | |
|---|---|---|---|---|---|---|---|---|---|---|---|---|---|---|---|---|---|---|---|---|---|
| 537.88 | 732.74 | 2081.43 | 315.9 | 673.51 | 1221.51 | 809.78 | 336.47 | 493.39 | 160.83 | 232.51 | 624.98 | 670.74 | 580.13 | 900.3 | 628.14 | 507.95 | 593.26 | 12101.45 | Probable ATP-dependent RNA helicase DDX5 (EC 3.6.4.13) (DEAD box protein 5) | DDX5 QtsA-17658 | alternative mRNA splicing, via spliceosome [GO:0000380]; androgen receptor signaling pathway [GO:0030521]; epithelial to mesenchymal transition [GO:0001837]; estrogen receptor signaling pathway [GO:0030520]; intrinsic apoptotic signaling pathway by p53 class mediator [GO:0072332]; miRNA transcription [GO:0061614]; myoblast differentiation [GO:0045445]; negative regulation of transcription by RNA polymerase II [GO:0000122]; nuclear-transcribed mRNA catabolic process [GO:0000956]; positive regulation of DNA damage response, signal transduction by p53 class mediator [GO:0043517]; regulation of alternative mRNA splicing, via spliceosome [GO:0000381]; regulation of androgen receptor signaling pathway [GO:0060765]; regulation of osteoblast differentiation [GO:0045667]; regulation of skeletal muscle cell differentiation [GO:2001014]; regulation of transcription by RNA polymerase II |
| 638.9 | 893.25 | 695.73 | 573.85 | 592.09 | 1437.33 | 6.83 | 1860.95 | 876.29 | 635.59 | 618.62 | 281.04 | 328.06 | 1394.88 | 332.55 | 89.44 | 450.36 | 394.54 | 12100.3 | Protein yippee-like 5 | YPEL5 CGI-127 | cell population proliferation [GO:0008283] |
| 360.55 | 3.61 | 2.6 | 1204.82 | 0 | 23.27 | 3509.03 | 0 | 0 | 0 | 0 | 0 | 0 | 0 | 0 | 6761.83 | 211.14 | 16.81 | 12093.66 | BAR/IMD domain-containing adapter protein 2-like 1 (Brain-specific angiogenesis inhibitor 1-associated protein 2-like protein 1) (BAI1-associated protein 2-like protein 1) | Baiap2l1 | actin crosslink formation [GO:0051764]; actin filament bundle assembly [GO:0051017]; plasma membrane organization [GO:0007009]; positive regulation of actin filament polymerization [GO:0030838]; regulation of actin cytoskeleton organization [GO:0032956] |

| | | | | | | | | | | | | | | | | | | | | | |
|---|---|---|---|---|---|---|---|---|---|---|---|---|---|---|---|---|---|---|---|---|---|
| 647.09 | 714.45 | 354.94 | 232.46 | 1178.84 | 573.06 | 454.86 | 277.06 | 286.65 | 266.45 | 564.31 | 638.47 | 880.85 | 2811.35 | 404.01 | 447.41 | 971.04 | 386.76 | 12090.06 | Microspherule protein 1 (58 kDa microspherule protein) (Cell cycle-regulated factor p78) (INO80 complex subunit J) (MCRS2) | MCRS1 INO80Q MSP58 | chromatin remodeling [GO:0006338]; DNA recombination [GO:0006310]; DNA repair [GO:0006281]; negative regulation of telomere maintenance via telomere lengthening [GO:1904357]; positive regulation of DNA repair [GO:0045739]; positive regulation of DNA-templated transcription [GO:0045893]; positive regulation of protein localization to nucleolus [GO:1904751]; positive regulation of telomere maintenance in response to DNA damage [GO:1904507]; positive regulation of transcription by RNA polymerase II [GO:0045944]; protein modification process [GO:0036211]; regulation of cell cycle [GO:0051726]; regulation of chromosome organization [GO:0033044]; regulation of DNA repair [GO:0006282]; regulation of DNA replication [GO:0006275]; regulation of DNA strand elongation [GO:0060382]; regulation of embryonic development [GO:0045995]; telomere maintenance [GO:0000723] |
| 488.31 | 529.71 | 393.03 | 196.18 | 350.93 | 434.78 | 3607.42 | 139.21 | 604.56 | 152.47 | 140.66 | 397.56 | 621.8 | 719.03 | 482.5 | 2002.95 | 226.82 | 599.34 | 12087.26 | Cyclin-dependent kinase-like 1 (EC 2.7.11.22) | cdkl1 zgc:101002 | |
| 0 | 3702.89 | 815.54 | 5.36 | 1721.85 | 319.94 | 11.75 | 1530.6 | 191.29 | 3.84 | 200.18 | 566.94 | 0 | 2276.61 | 371.87 | 9.88 | 180.49 | 173.42 | 12082.45 | CXXC motif containing zinc binding protein (UPF0587 protein C1orf123 homolog) | czib | |
| 1159.68 | 523.77 | 977.73 | 360.31 | 615.01 | 1477.42 | 1152.17 | 255.88 | 477.11 | 371.63 | 227.72 | 513.45 | 871.42 | 715.02 | 516.43 | 856.61 | 456.36 | 554.02 | 12081.74 | Tyrosine-protein phosphatase non-receptor type 13 (EC 3.1.3.48) (Fas-associated protein-tyrosine phosphatase 1) (FAP-1) (PTP-BAS) (Protein-tyrosine phosphatase 1E) (PTP-E1) (hPTPE1) (Protein-tyrosine phosphatase PTPL1) | PTPN13 PNP1 PTP1E PTPL1 | cellular response to toxic substance [GO:0097237]; negative regulation of excitatory synapse assembly [GO:1904890]; negative regulation of protein phosphorylation [GO:0001933]; peptidyl-tyrosine dephosphorylation [GO:0035335]; protein dephosphorylation [GO:0006470]; regulation of phosphatidylinositol 3-kinase/protein kinase B signal transduction [GO:0051896] |

| | | | | | | | | | | | | | | | | | | | | | |
|---|---|---|---|---|---|---|---|---|---|---|---|---|---|---|---|---|---|---|---|---|---|
| 464.44 | 431.91 | 885.03 | 172.64 | 425.1 | 231.31 | 398.13 | 183.1 | 1965.88 | 123.69 | 135.64 | 816.55 | 536.59 | 959.25 | 1516.54 | 488.2 | 354.82 | 1974.08 | 12062.9 | Liprin-alpha-4 (Protein tyrosine phosphatase receptor type f polypeptide-interacting protein alpha-4) (PTPRF-interacting protein alpha-4) | PPFIA4 KIAA0897 | synapse organization [GO:0050808] |
| 605.37 | 609.01 | 852.86 | 200.37 | 647.77 | 772.63 | 294.59 | 379.39 | 1414.48 | 242.41 | 262.64 | 814.27 | 576.16 | 947.28 | 1263.56 | 279.16 | 422.9 | 1475.14 | 12059.99 | NEDD4-binding protein 1 (N4BP1) (EC 3.1.-.-) | n4bp1 si:ch211-215n5.3 zgc:66437 | innate immune response [GO:0045087]; negative regulation of cytokine production [GO:0001818]; negative regulation of proteasomal ubiquitin-dependent protein catabolic process [GO:0032435]; negative regulation of protein ubiquitination [GO:0031397]; negative regulation of viral genome replication [GO:0045071]; regulation of innate immune response [GO:0045088] |
| 554.59 | 1468.57 | 1131 | 385.38 | 1591.04 | 992.12 | 102.45 | 313.88 | 345.69 | 274.64 | 430.12 | 601.19 | 617.71 | 942.26 | 530.12 | 279.57 | 979.23 | 518.74 | 12058.3 | Friend leukemia integration 1 transcription factor (Proto-oncogene Fli-1) | FLI1 | cell differentiation [GO:0030154]; regulation of transcription by RNA polymerase II [GO:0006357] |

| | | | | | | | | | | | | | | | | | | | | | |
|---|---|---|---|---|---|---|---|---|---|---|---|---|---|---|---|---|---|---|---|---|---|
| 880.43 | 519.05 | 1199.13 | 148.53 | 538.28 | 752.94 | 629.26 | 334.79 | 1238.57 | 222.47 | 186.22 | 740.26 | 643.61 | 664.36 | 1041.7 | 654.74 | 391.29 | 1262.1 | 12047.73 | Glia-derived nexin (GDN) (Peptidase inhibitor 7) (PI-7) (Protease nexin 1) (PN-1) (Protease nexin I) (Serpin E2) | Serpine2 Pi7 Pn1 | blood coagulation [GO:0007596]; cell differentiation [GO:0030154]; cerebellar granular layer morphogenesis [GO:0021683]; dense core granule biogenesis [GO:0061110]; detection of mechanical stimulus involved in sensory perception [GO:0050974]; embryo implantation [GO:0007566]; innervation [GO:0060384]; long-term synaptic potentiation [GO:0060291]; mating plug formation [GO:0042628]; negative regulation of blood coagulation [GO:0030195]; negative regulation of cell growth [GO:0030308]; negative regulation of cell population proliferation [GO:0008285]; negative regulation of phosphatidylinositol 3-kinase/protein kinase B signal transduction [GO:0051898]; negative regulation of plasminogen activation [GO:0010757]; negative regulation of platelet activation [GO:0010544]; negative regulation of platelet aggregation [GO:0090331]; negative regulation of protein catabolic process |
| 74.65 | 1610.26 | 190.16 | 87.17 | 2612.81 | 891.91 | 6.6 | 782.3 | 9.76 | 13.63 | 419.22 | 131.15 | 43.08 | 2748.82 | 462.83 | 276.17 | 1644.74 | 35.39 | 12040.65 | Voltage-dependent calcium channel subunit alpha-2/delta-4 (Voltage-gated calcium channel subunit alpha-2/delta-4) [Cleaved into: Voltage-dependent calcium channel subunit alpha-2-4;Voltage-dependent calcium channel subunit delta-4] | Cacna2d4 | |

| | | | | | | | | | | | | | | | | | | | | | |
|---|---|---|---|---|---|---|---|---|---|---|---|---|---|---|---|---|---|---|---|---|---|
| 3538.86 | 311.59 | 927.1 | 780.49 | 812.86 | 933.72 | 263.21 | 282.97 | 513.04 | 1433.77 | 350.15 | 314.21 | 424.86 | 75.34 | 103.44 | 163.32 | 405.26 | 403.29 | 12037.48 | Integrin alpha-11 | Itga11 | cell adhesion [GO:0007155]; fibroblast migration [GO:0010761]; integrin-mediated signaling pathway [GO:0007229]; substrate-dependent cell migration [GO:0006929] |
| 2095.2 | 210.93 | 622.75 | 572.84 | 241.24 | 1671.39 | 112.79 | 125.05 | 327.4 | 703.35 | 127.85 | 615.79 | 1630.7 | 918.49 | 608.5 | 418.01 | 574.31 | 452.6 | 12029.19 | Zinc finger SWIM domain-containing protein 6 | Zswim6 Kiaa1577 | neuron projection morphogenesis [GO:0048812]; regulation of neuron migration [GO:2001222]; striatal medium spiny neuron differentiation [GO:0021773] |
| 1905.41 | 244.1 | 1321.57 | 793.01 | 496.02 | 800.87 | 33.25 | 116.74 | 288.72 | 720.38 | 203.95 | 2123.31 | 1067.51 | 275.43 | 410.42 | 366.95 | 402.54 | 457.95 | 12028.13 | Probable RNA polymerase II nuclear localization protein SLC7A6OS (Solute carrier family 7 member 6 opposite strand transcript homolog) | slc7a6os zgc:103493 | central nervous system development [GO:0007417]; developmental process [GO:0032502]; midbrain-hindbrain boundary structural organization [GO:0021552]; protein transport [GO:0015031]; somite development [GO:0061053] |

| | | | | | | | | | | | | | | | | | | | | | |
|---|---|---|---|---|---|---|---|---|---|---|---|---|---|---|---|---|---|---|---|---|---|
| 101.31 | 2076 | 382.08 | 219.44 | 2006.71 | 563.03 | 134.54 | 1345.2 | 143.47 | 139.78 | 593.95 | 271.81 | 87.12 | 2311.54 | 246.8 | 105.42 | 973.11 | 326.42 | 12027.73 | Complement component C8 beta chain (Complement component 8 subunit beta) | c8b | complement activation, alternative pathway [GO:0006957]; complement activation, classical pathway [GO:0006958]; killing of cells of another organism [GO:0031640] |
| 555.23 | 320.61 | 1788.29 | 145.8 | 377.77 | 571.55 | 498.63 | 143.2 | 2024.56 | 140.81 | 96.84 | 710.69 | 317.17 | 362.99 | 1560.6 | 380.59 | 201.48 | 1828.58 | 12025.39 | ADP-ribosylhydrolase ARH1 (EC 3.2.2.19) (ADP-ribose-L-arginine cleaving enzyme) ([Protein ADP-ribosylarginine] hydrolase) (ADP-ribosylarginine hydrolase) (hARH1) | ADPRH ARH1 | protein de-ADP-ribosylation [GO:0051725]; protein modification process [GO:0036211] |
| 229.15 | 153.2 | 456.19 | 194.15 | 224.11 | 1794.22 | 3524.32 | 91.96 | 153.55 | 108.75 | 141.26 | 1035.76 | 430.82 | 289.01 | 515.95 | 2020.98 | 390.06 | 255.66 | 12009.1 | Golgi apparatus membrane protein TVP23 homolog B | Tvp23b Fam18b Fam18b1 | |

| | | | | | | | | | | | | | | | | | | | | | |
|---|---|---|---|---|---|---|---|---|---|---|---|---|---|---|---|---|---|---|---|---|---|
| 794.04 | 368.13 | 998.98 | 126.02 | 251.06 | 563.58 | 168.77 | 86.09 | 1718.02 | 177.84 | 98.67 | 1005.44 | 635.72 | 445.95 | 2008.04 | 193.86 | 178.97 | 2186.57 | 12005.75 | E3 ubiquitin-protein ligase COP1 (EC 2.3.2.27) (Constitutive photomorphogenesis protein 1 homolog) (mCOP1) (RING finger and WD repeat domain protein 2) (RING-type E3 ubiquitin transferase RFWD2) | Cop1 Rfwd2 RNF200 | positive regulation of proteasomal ubiquitin-dependent protein catabolic process [GO:0032436]; proteasome-mediated ubiquitin-dependent protein catabolic process [GO:0043161]; protein ubiquitination [GO:0016567]; response to ionizing radiation [GO:0010212] |
| 563.68 | 1619.2 | 458.06 | 206.48 | 1965.85 | 744.37 | 205.66 | 888.61 | 276.53 | 226.62 | 475.44 | 385.54 | 382.32 | 1735.64 | 387.58 | 189.71 | 975.78 | 318.09 | 12005.16 | Coiled-coil domain-containing protein 142 | CCDC142 PSEC0243 | |
| 2509.81 | 340.46 | 235.26 | 556.54 | 394.49 | 586.68 | 412 | 396.55 | 42.73 | 2157.36 | 525.37 | 450.57 | 2027.58 | 472.94 | 121.05 | 286.07 | 458.85 | 22.93 | 11997.24 | L-fucose dehydrogenase (EC 1.1.1.122) (17-beta-hydroxysteroid dehydrogenase DHRS10) (Dehydrogenase/reductase SDR family member 10) (Retinal short-chain dehydrogenase/reductase retSDR3) | HSD17B14 DHRS10 SDR3 | L-fucose catabolic process [GO:0042355]; steroid catabolic process [GO:0006706] |

| | | | | | | | | | | | | | | | | | | | | | |
|---|---|---|---|---|---|---|---|---|---|---|---|---|---|---|---|---|---|---|---|---|---|
| 46.26 | 0.36 | 6.23 | 58.8 | 0 | 3.66 | 7526.55 | 0.26 | 1.1 | 76.74 | 188.58 | 1 | 14.65 | 0 | 0.63 | 3919.51 | 123.06 | 6.45 | 11973.84 | Epidermal growth factor receptor (EC 2.7.10.1) | EGFR | cellular response to epidermal growth factor stimulus [GO:0071364]; cellular response to estradiol stimulus [GO:0071392]; epidermal growth factor receptor signaling pathway [GO:0007173]; learning or memory [GO:0007611]; negative regulation of apoptotic process [GO:0043066]; neuron differentiation [GO:0030182]; positive regulation of cell growth [GO:0030307]; positive regulation of epithelial cell proliferation [GO:0050679]; positive regulation of ERK1 and ERK2 cascade [GO:0070374]; positive regulation of MAPK cascade [GO:0043410]; positive regulation of protein localization to early endosome [GO:1902966]; positive regulation of protein phosphorylation [GO:0001934]; positive regulation of transcription by RNA polymerase II [GO:0045944] |
| 986.03 | 616.64 | 1146.14 | 20.94 | 581.04 | 122.56 | 1764.39 | 308.07 | 731.21 | 35.45 | 251.23 | 367.64 | 540.33 | 1074.33 | 915.23 | 1394.47 | 490.52 | 624 | 11970.22 | Trinucleotide repeat-containing gene 6C protein | TNRC6C KIAA1582 | miRNA-mediated gene silencing by inhibition of translation [GO:0035278]; miRNA-mediated post-transcriptional gene silencing [GO:0035195]; positive regulation of nuclear-transcribed mRNA catabolic process, deadenylation-dependent decay [GO:1900153]; positive regulation of nuclear-transcribed mRNA poly(A) tail shortening [GO:0060213] |

| | | | | | | | | | | | | | | | | | | | | | |
|---|---|---|---|---|---|---|---|---|---|---|---|---|---|---|---|---|---|---|---|---|---|
| 919.91 | 997.21 | 772.93 | 193.81 | 605.45 | 935.76 | 15.95 | 364.01 | 87.78 | 300.74 | 355.7 | 407.09 | 1906.13 | 1634.19 | 834.36 | 182.81 | 848.03 | 608.32 | 11970.18 | BBSome complex member BBS1 (Bardet-Biedl syndrome 1 protein homolog) | Bbs1 | adult behavior [GO:0030534]; brain morphogenesis [GO:0048854]; cartilage development [GO:0051216]; cerebral cortex development [GO:0021987]; cilium assembly [GO:0060271]; dendrite development [GO:0016358]; fat cell differentiation [GO:0045444]; fertilization [GO:0009566]; Golgi to plasma membrane protein transport [GO:0043001]; hippocampus development [GO:0021766]; hormone metabolic process [GO:0042445]; intraciliary retrograde transport [GO:0035721]; lipid metabolic process [GO:0006629]; microtubule cytoskeleton organization [GO:0000226]; neural precursor cell proliferation [GO:0061351]; neuron migration [GO:0001764]; non-motile cilium assembly [GO:1905515]; olfactory behavior [GO:0042048]; photoreceptor cell maintenance [GO:0045494]; photoreceptor cell morphogenesis [GO:0008594]; protein localization [GO:0008104]; protein localization to cilium [GO:0061512]; |
| 1 | 14.15 | 360.38 | 0.91 | 68.84 | 15.13 | 1.45 | 34.25 | 4142.2 | 1.57 | 22.98 | 1089.35 | 2.05 | 107.47 | 3165.23 | 6.06 | 6.34 | 2923.88 | 11963.24 | DmX-like protein 1 (X-like 1 protein) | DMXL1 XL1 | vacuolar acidification [GO:0007035] |
| 22.09 | 29.94 | 1427.46 | 64.03 | 56.91 | 482.6 | 3.23 | 22.13 | 3871.66 | 14.86 | 33.12 | 1275.88 | 21.58 | 56.15 | 2044.34 | 27.47 | 35.13 | 2468.17 | 11956.75 | CD209 antigen-like protein D (DC-SIGN-related protein 3) (DC-SIGNR3) (CD antigen CD209) | Cd209d | defense response to bacterium [GO:0042742]; endocytosis [GO:0006897]; positive regulation of cytokine production [GO:0001819] |

| | | | | | | | | | | | | | | | | | | | | | | |
|---|---|---|---|---|---|---|---|---|---|---|---|---|---|---|---|---|---|---|---|---|---|---|
| 505.39 | | 534.93 | 379.8 | 216.96 | 938.08 | 1364.89 | 155.88 | 736.76 | 118.61 | 315.58 | 447.21 | 626.82 | 1683.94 | 1614.3 | 735.74 | 254.73 | 842.91 | 468.73 | 11941.26 | WASH complex subunit 4 (Strumpellin and WASH-interacting protein) (SWIP) (WASH complex subunit SWIP) | WASHC4 KIAA1033 | cognition [GO:0050890]; endolysosomal toll-like receptor signaling pathway [GO:0140894]; endosomal transport [GO:0016197]; endosome organization [GO:0007032]; neuromuscular process [GO:0050905]; nuclear envelope budding [GO:0140591]; protein transport [GO:0015031]; regulation of Arp2/3 complex-mediated actin nucleation [GO:0034315]; regulation of protein complex stability [GO:0061635] |
| 792.24 | | 1106.58 | 555.68 | 326.91 | 987.5 | 523.1 | 11.1 | 683.12 | 51.75 | 439.19 | 358.83 | 303.53 | 2657.23 | 1568.8 | 427.77 | 141.22 | 832.92 | 171.99 | 11939.46 | Thioredoxin domain-containing protein 5 (EC 1.8.4.-) (EC 5.3.4.1) (Endoplasmic reticulum resident protein 46) (ER protein 46) (ERp46) (Plasma cell-specific thioredoxin-related protein) (PC-TRP) (Thioredoxin-like protein p46) | Txndc5 Tlp46 | |

| | | | | | | | | | | | | | | | | | | | | | |
|---|---|---|---|---|---|---|---|---|---|---|---|---|---|---|---|---|---|---|---|---|---|
| 656.51 | 1647.1 | 69.52 | 750.04 | 596.42 | 1516.08 | 0 | 1333.83 | 0 | 1471.54 | 1418.24 | 0 | 573.6 | 1559.9 | 133.6 | 29.08 | 159.14 | 8.48 | 11923.08 | Tetratricopeptide repeat protein 5 (TPR repeat protein 5) (Stress-responsive activator of p300) (Protein Strap) | TTC5 | cellular response to starvation [GO:0009267]; DNA damage response [GO:0006974]; DNA repair [GO:0006281]; positive regulation of mRNA catabolic process [GO:0061014]; positive regulation of transcription by RNA polymerase II [GO:0045944] |
| 333.28 | 743.23 | 1240.99 | 133.49 | 702.42 | 1033.59 | 145.71 | 196.35 | 1391.01 | 122.99 | 184.69 | 1116.66 | 383.48 | 494.66 | 1646.63 | 166.9 | 317.38 | 1568.23 | 11921.69 | Cell death regulator Aven | Aven | apoptotic process [GO:0006915]; negative regulation of apoptotic process [GO:0043066] |
| 58.74 | 0.79 | 3.4 | 421.1 | 0 | 5.56 | 6481.65 | 0 | 0 | 68.9 | 118.51 | 0 | 7.34 | 0 | 4.08 | 4653.53 | 91.35 | 4.01 | 11918.96 | Kelch domain-containing protein 1 | Klhdc1 | protein ubiquitination [GO:0016567]; ubiquitin-dependent protein catabolic process via the C-end degron rule pathway [GO:0140627] |
| 1346.55 | 397.97 | 649.51 | 380.62 | 410.83 | 452.62 | 1798.01 | 218.93 | 243.81 | 440.5 | 312.28 | 542.82 | 775.94 | 572.79 | 316.69 | 2009.15 | 683.9 | 363.5 | 11916.42 | Metallophosphoesterase 1 (EC 3.1.-.-) (Post-GPI attachment to proteins factor 5) | mppe1 pgap5 zgc:112219 | endoplasmic reticulum to Golgi vesicle-mediated transport [GO:0006888]; GPI anchor biosynthetic process [GO:0006506] |

| | | | | | | | | | | | | | | | | | | | | | |
|---|---|---|---|---|---|---|---|---|---|---|---|---|---|---|---|---|---|---|---|---|---|
| 762.08 | 596.55 | 800.41 | 286.14 | 502.56 | 814.05 | 892.93 | 229.7 | 983.79 | 268.38 | 201.14 | 631.35 | 796.53 | 852.68 | 947.38 | 687.35 | 475.51 | 1177.26 | 11905.79 | Bromodomain-containing protein 1 (BR140-like protein) (Bromodomain and PHD finger-containing protein 2) | BRD1 BRL BRPF2 | chromatin remodeling [GO:0006338]; erythrocyte maturation [GO:0043249]; positive regulation of erythrocyte differentiation [GO:0045648]; regulation of developmental process [GO:0050793]; regulation of DNA-templated transcription [GO:0006355]; regulation of hemopoiesis [GO:1903706]; regulation of transcription by RNA polymerase II [GO:0006357]; response to electrical stimulus [GO:0051602]; response to immobilization stress [GO:0035902] |
| 648.6 | 372.01 | 1526.19 | 133.09 | 342.25 | 2040.49 | 271.85 | 180.72 | 788.85 | 188.43 | 162.88 | 921.06 | 777.6 | 305.32 | 1098.46 | 288.25 | 301.12 | 1557.66 | 11904.83 | 3-hydroxyisobutyryl-CoA hydrolase, mitochondrial (EC 3.1.2.4) (3-hydroxyisobutyryl-coenzyme A hydrolase) (HIB-CoA hydrolase) (HIBYL-CoA-H) | hibch zgc:110824 | valine catabolic process [GO:0006574] |
| 5.6 | 2804.36 | 110.7 | 4.73 | 2192.37 | 130.44 | 35.99 | 488.2 | 0 | 5.29 | 788.52 | 114.91 | 0 | 2813.45 | 22.96 | 16.46 | 2358.31 | 11.01 | 11903.3 | Lysyl oxidase homolog 2A (EC 1.4.3.13) (Lysyl oxidase-like protein 2A) | loxl2a | collagen fibril organization [GO:0030199]; endothelial cell migration [GO:0043542]; endothelial cell proliferation [GO:0001935]; epithelial to mesenchymal transition [GO:0001837]; heterochromatin organization [GO:0070828]; negative regulation of DNA-templated transcription [GO:0045892]; negative regulation of stem cell population maintenance [GO:1902455]; negative regulation of transcription by RNA polymerase II [GO:0000122]; peptidyl-lysine oxidation [GO:0018057]; positive regulation of chondrocyte differentiation [GO:0032332]; positive regulation of epithelial to mesenchymal transition [GO:0010718]; response to hypoxia [GO:0001666]; sprouting angiogenesis [GO:0002040] |

| | | | | | | | | | | | | | | | | | | | | | |
|---|---|---|---|---|---|---|---|---|---|---|---|---|---|---|---|---|---|---|---|---|---|
| 400.77 | 924.02 | 1356.51 | 125.51 | 899.69 | 1001.35 | 155.1 | 347.64 | 569.69 | 133.14 | 225.39 | 797.07 | 607.42 | 1171.92 | 1309.38 | 189.4 | 560.79 | 1117.68 | 11892.47 | Dipeptidyl peptidase 9 (EC 3.4.14.5) | dpp9 zgc:152900 | proteolysis [GO:0006508]; regulation of inflammatory response [GO:0050727] |
| 578.24 | 897.43 | 406.61 | 749 | 753.53 | 1665.85 | 257.26 | 938.5 | 443.04 | 877.82 | 953.84 | 684.51 | 386.27 | 975.43 | 342.8 | 328.29 | 327.74 | 323.66 | 11889.82 | Methionine aminopeptidase 1D, mitochondrial (MAP 1D) (MetAP 1D) (EC 3.4.11.18) (Methionyl aminopeptidase type 1D, mitochondrial) (Peptidase M 1D) | metap1d map1d zgc:110461 | proteolysis [GO:0006508] |
| 158.19 | 126.14 | 1177.22 | 98.14 | 89.16 | 184.17 | 721.35 | 32.61 | 2256.24 | 61.37 | 51.88 | 1000.05 | 74.03 | 155.74 | 2101.71 | 585.28 | 105.6 | 2901.86 | 11880.74 | Homeobox protein Nkx-6.1 (Homeobox protein NK-6 homolog A) | Nkx6-1 Nkx6.1 Nkx6a | cell differentiation [GO:0030154]; cellular response to cytokine stimulus [GO:0071345]; cellular response to peptide hormone stimulus [GO:0071375]; central nervous system neuron differentiation [GO:0021953]; endocrine pancreas development [GO:0031018]; negative regulation of oligodendrocyte differentiation [GO:0048715]; negative regulation of transcription by RNA polymerase II [GO:0000122]; neurogenesis [GO:0022008]; neuron differentiation [GO:0030182]; oligodendrocyte differentiation [GO:0048709]; pancreatic A cell differentiation [GO:0003310]; positive regulation of insulin secretion [GO:0032024]; positive regulation of neuron differentiation [GO:0045666]; positive regulation of oligodendrocyte differentiation [GO:0048714]; regulation of axon extension [GO:0030516]; regulation of neuron migration [GO:2001222]; regulation of transcription by RNA polymerase II [GO:0006357]; response |

| | | | | | | | | | | | | | | | | | | | | | |
|---|---|---|---|---|---|---|---|---|---|---|---|---|---|---|---|---|---|---|---|---|---|
| 0.25 | 1.54 | 102.64 | 4.64 | 1.32 | 16.59 | 1.63 | 3.43 | 4184.27 | 3.6 | 3.13 | 1277.17 | 4.92 | 7.09 | 2989.03 | 9.37 | 20.54 | 3236.38 | 11867.54 | Cytosolic carboxypeptidase 1 (EC 3.4.17.-) (EC 3.4.17.24) (ATP/GTP-binding protein 1) (Protein deglutamylase CCP1) | agtpbp1 ccp1 | C-terminal protein deglutamylation [GO:0035609]; cerebellar Purkinje cell differentiation [GO:0021702]; chordate embryonic development [GO:0043009]; eye photoreceptor cell differentiation [GO:0001754]; mitochondrion organization [GO:0007005]; neuromuscular process [GO:0050905]; olfactory bulb development [GO:0021772]; protein side chain deglutamylation [GO:0035610]; proteolysis [GO:0006508]; T cell differentiation in thymus [GO:0033077] |
| 400.02 | 184.49 | 499.04 | 221.41 | 221.43 | 370.29 | 4193.09 | 107.15 | 286.97 | 132.74 | 155.92 | 167.53 | 364.4 | 188.55 | 319.69 | 3537.37 | 218.88 | 296.06 | 11865.03 | Nck-associated protein 1 (NAP 1) | nckap1 si:dkey-234n3.1 wu:fd05c01 | cell migration [GO:0016477]; cell morphogenesis [GO:0000902]; cell projection assembly [GO:0030031]; cortical actin cytoskeleton organization [GO:0030866]; neuron projection morphogenesis [GO:0048812] |

**Table S3. Read depth and contigs among sample types.**

| Sample ID | Species | Tissue | Sample | Total reads | Contigs | N50 |
|---|---|---|---|---|---|---|
| W1.1 | Macropharyngodon choati | Spleen | W1-1 | 57,984,236 | 135331 | 1515 |
| W1.2 | Macropharyngodon choati | Liver | W1-2 | 55,237,154 | 146945 | 1443 |
| W1.3 | Macropharyngodon choati | Brain | W1-3 | 56,657,276 | 252810 | 1289 |
| W2.1 | Macropharyngodon choati | Spleen | W2-1 | 56,922,386 | 111341 | 1086 |
| W2.2 | Macropharyngodon choati | Liver | W2-2 | 52,630,940 | 136072 | 1393 |
| W2.3 | Macropharyngodon choati | Brain | W2-3 | 58,123,578 | 185322 | 1438 |
| W3.1 | Macropharyngodon choati | Spleen | W3-1 | 55,036,118 | 35305 | 2245 |
| W3.2 | Macropharyngodon choati | Liver | W3-2 | 51,281,314 | 125489 | 1236 |
| W3.3 | Macropharyngodon choati | Brain | W3-3 | 53,389,004 | 241874 | 1814 |
| W4.1 | Macropharyngodon choati | Spleen | W4-1 | 56,002,672 | 104651 | 1025 |
| W4.2 | Macropharyngodon choati | Liver | W4-2 | 52,781,266 | 117339 | 1119 |
| W4.3 | Macropharyngodon choati | Brain | W4-3 | 56,828,926 | 241328 | 1147 |
| W5.1 | Macropharyngodon choati | Spleen | W5-1 | 56,506,838 | 164824 | 1581 |
| W5.2 | Macropharyngodon choati | Liver | W5-2 | 52,832,582 | 164139 | 1407 |
| W5.3 | Macropharyngodon choati | Brain | W5-3 | 51,554,294 | 266697 | 1756 |
| W6.1 | Macropharyngodon choati | Spleen | W6-1 | 49,127,882 | 115055 | 1264 |
| W6.2 | Macropharyngodon choati | Liver | W6-2 | 49,481,968 | 151548 | 1429 |
| W6.3 | Macropharyngodon choati | Brain | W6-3 | 57,554,352 | 250964 | 1963 |
